# Rab2 and Arl8/BORC control retrograde axonal transport of dense core vesicles via Syd/dJIP3/4 and RUFY dynein adaptors

**DOI:** 10.1101/2025.05.28.656585

**Authors:** Viktor K. Lund, Antony Chirco, Michela Caliari, Andreas Haahr Larsen, Ulrik Gether, Kenneth Lindegaard Madsen, Michael Wierer, Ole Kjaerulff

## Abstract

Neuropeptide-containing dense core vesicles (DCVs) are generated in the neuronal cell body and circulate throughout the axonal arbor to supply distal release sites. This circulation depends on the anterograde kinesin-1 and kinesin-3 motors and the retrograde dynein-dynactin motor. While kinesin-3 is recruited to DCVs with the aid of the small GTPase Arl8, it is unclear how dynein and kinesin-1 are recruited and regulated. Here we show that DCV motility in Drosophila (fruit flies) depends on the dynein and kinesin-1 adaptor Sunday Driver/dJIP3/4 (Syd) and the novel dynein adaptor RUFY. Syd and RUFY bind each other; moreover, Syd binds the DCV-located GTPase Rab2 that controls retrograde DCV transport, and RUFY binds Arl8. Disruption of Rab2, Syd, RUFY, dynein, and the Arl8 activator BORC all produce a similar DCV axonal transport phenotype characterized by axonal accumulation of immobile DCVs and a selective reduction in retrograde DCV flux. Our data suggest a model where dynein is recruited and activated by a complex of Syd and RUFY, which is anchored to DCVs by a Rab2- and Arl8-dependent mechanism. Lastly, we show that loss of Rab2 results in missorting of the DCV membrane proteins VMAT and Synaptotagmin-α, similar to the reported effect of Rab2 deletion on the sorting of synaptic vesicle and active zone proteins. However, disruption of Syd, RUFY or dynein does not phenocopy the Rab2-specific VMAT sorting defect, suggesting that Rab2 employs separate effectors in DCV biogenesis and motility.

## Introduction

In neurons, long distance axonal transport along microtubules (MTs) mediated by molecular motors is critical for enabling axonal outgrowth, maintaining synaptic function and ensuring structural and functional synaptic plasticity. Neuropeptide/neurohormone-containing dense core vesicles (DCVs) constitute a particular logistical challenge in this context since, unlike small synaptic vesicles (SVs) that are generated locally at the presynapse through endocytosis, they are produced at the *trans*-Golgi network (TGN) in the soma and completely rely on axonal transport for delivery to distal synaptic release sites. Previous work in Drosophila ^1^ and hippocampal neurons ^2^ indicates that DCVs circulate throughout the axonal arbor, only reversing direction at distal axonal termini and the proximal axon, with only a relatively small probability of being deposited in any given synaptic bouton as they pass through it. This arrangement has been suggested to ensure an even distribution of DCVs between synaptic boutons within the arbor ^1, 3^ and the circulating DCVs also represent a large reserve pool that can be quickly drawn upon when synaptic release sites are depleted. Indeed, synaptic capture of circulating DCVs increases in response to neuronal activity ^4, 5, 6^. This means that anterograde transport (towards synaptic termini) and retrograde transport (towards the soma) are equally important for supplying synaptic boutons.

Axonal MTs are universally oriented with plus ends out towards distal axonal termini and minus ends in towards the soma. Anterograde transport is mediated by plus-end directed kinesin family motors, while retrograde transport is mediated by the minus-end directed cytoplasmic dynein motor ^7^. These motors are usually autoinhibited in their native, non-cargo coupled state and motor activation and cargo attachment is regulated by a large set of cargo and motor specific accessory and adaptor proteins, and often involves small GTPases of the Rab and Arf/Arl families ^8, 9^. Rab and Arf/Arl proteins behave like molecular switches cycling between an inactive GDP-loaded soluble state and an active GTP-loaded state where they are inserted into specific organellar membranes and recruit effector proteins such as tethers, vesicle coats and motor adaptors ^10^. This allows Rab and Arf/Arl proteins to link organellar identity to the characteristics and behavior of their cognate compartments, including motor-driven movement pattern and location within the cell.

In both invertebrates and mammals, axonal DCV transport depends on the fast kinesin-3 and the slower kinesin-1 anterograde motors, as well as dynein ^11, 12, 13, 14, 15^. Kinesin-3 is required to traverse a pre-axonal filtering region for cell body exit ^2, 12, 13^ and is regulated by the DCV-resident ^16^ small GTPase Arl8, which binds and activates it directly ^17, 18^. However, it is less clear how kinesin-1 and dynein are recruited to DCVs to maintain DCV circulation ^9^.

A potential clue to the mechanism governing retrograde DCV transport comes from our finding that the small GTPase Rab2 is required for axonal transport of DCVs and lysosomes in Drosophila ^16^. Notably, although we formerly speculated that Rab2 may control kinesin-3, the Rab2 loss-of-function phenotype was characterized by a strikingly selective reduction in retrograde DCV transport, implying that Rab2 may play a role in dynein regulation ^16^. Moreover, Rab2 overexpression caused the redistribution of DCVs from the axono-synaptic compartment to the soma in pupal neurons releasing the Bursicon neuropeptide hormone ^16^. Rab2 is a highly conserved member of the Rab protein family, which apart from its involvement in organelle motility is indispensable for lysosomal function ^19, 20^, autophagy ^21, 22^, synaptic protein sorting ^23^, and DCV biogenesis ^24, 25, 26, 27^. Work mostly done in *C. elegans* indicates that Rab2 and certain Rab2 effectors prevent the loss of a subset of DCV cargos to late endosomes/lysosomes during DCV maturation ^24, 25, 27^. Despite the identification of many molecular components, the exact details of the Rab2-dependent DCV cargo sorting pathway have remained mysterious, although an endosomal recycling mechanism seems to be involved ^28, 29^.

Here we use proximity proteomics to find that active Rab2 associates with the dynein/kinesin-1 adaptor Sunday Driver (Syd/dJIP3/4) and the novel dynein adaptor RUFY1 (RUFY) in living fly neurons. Biochemical experiments using Drosophila proteins expressed in HEK cells indicate that Rab2 physically interacts with Syd via the Syd RH2 cargo-binding domain, while RUFY binds Syd through the Syd C-terminal WD40 domain. Furthermore, defects in Rab2, Syd, RUFY, dynein, and partially kinesin-1, produce qualitatively similar effects on axonal transport of DCVs in fly motor neurons, characterized by a selective loss of retrograde transport and an increase in the proportion of static DCVs in the axons. In contrast, mutation of kinesin-3 primarily results in a strong symmetric reduction of anterograde and retrograde DCV fluxes due to a block in cell body exit, coupled with a severe slowing of anterograde transport velocity. We also show that, like the mammalian RUFY1-4 proteins, fly RUFY also binds Arl8, suggesting that the dynein-dynactin activating retrograde transport complex composed of Syd and RUFY is stabilized on DCVs in a Rab2- and Arl8-dependent manner. Consistent with this, knockout of the Arl8-activating BORC complex produces a phenotype closely resembling dynein/kinesin-1 loss-of-function phenotypes. Lastly, we find that in *Rab2* null neurons, DCV membrane cargos are missorted and accumulate in ectopic aggregates that likely represent stalled transport vesicles unable to fuse with the TGN or maturing DCVs, which mirrors recent findings for SV-components. Syd, RUFY and dynein are not responsible for the Rab2-dependent DCV membrane cargo sorting, but may together with Rab2 control DCV abundance.

## Results

### Proximity-dependent biotinylation identifies Sunday Driver (Syd)/dJIP3/4 as a DCV-resident Rab2-interacting protein

Rab2 is present on neuronal DCVs, and retrograde axonal transport of DCVs is severely and selectively disrupted in motor neurons of *Rab2* null third instar (L3) Drosophila larvae (**Fig 1A**) ^16^. We previously hypothesized that this reflects a function of Rab2 in recruitment of molecular motors to the DCV surface through adaptor proteins. However, none of the known Rab2 effectors that could reasonably be expected to fill this role, such as the BicD dynein adaptor ^30^, were required for normal axonal DCV transport in flies ^16^. To identify potential novel effector proteins that could link activated Rab2 to motors responsible for DCV motility, we employed in vivo proximity-dependent biotinylation (PDB) combined with quantitative mass spectrometry (MS), using a pan-neuronally expressed constitutively active GTP-locked TurboID-Rab2^Q65L^ chimera as bait (**Fig 1B**). TurboID is a promiscuous biotin ligase derived from BirA* ^31^, which when fused to a protein of interest and expressed in a desired tissue biotinylates proteins in its immediate vicinity (within ∼10 nm ^32^) that can then be isolated using streptavidin. To filter out proteins not specifically interacting with the active form of Rab2, the MS signal of purified biotinylated neuronal proteins from TurboID-Rab2^Q65L^-expressing adult flies was compared to that of control flies expressing the inactive GDP-locked TurboID-Rab2^S20N^ variant. This approach identified ∼300 proteins significantly enriched more than two-fold in the nano-environment of active neuronal Rab2 (**Fig. 1B, Supplementary Data 1, 2)**, including many known Rab2 effectors **(Fig. S1A)**. The remaining proteins in this group likely represent a mix of unknown effectors, constituents of effector complexes, and resident proteins of Rab2-associated vesicular compartments. The highest levels of enrichment were seen for proteins involved in lysosomal function and autophagy (with the most enriched protein being the transmembrane autophagy factor Atg9) (**Fig. 1B, C, Supplementary Data 2)**. This is in agreement with a critical role of Rab2 in lysosomal biogenesis and macroautophagy ^19, 20, 21^. Golgi apparatus-associated tethering proteins, many of them Rab2 effectors, were also well-represented, consistent with the involvement of Rab2 in Golgi function ^23, 33, 34^. In addition, there was a strong representation of early and recycling endosomal proteins (**Fig. 1C, Supplementary Data 2)**. Although this latter finding may in part reflect the difficulty of clearly differentiating between components belonging to the early and late stages of the endocytic pathway, it also fits with observations of Rab2 presence at a lower level throughout the endosomal system ^20, 21^.

**Figure 1.**
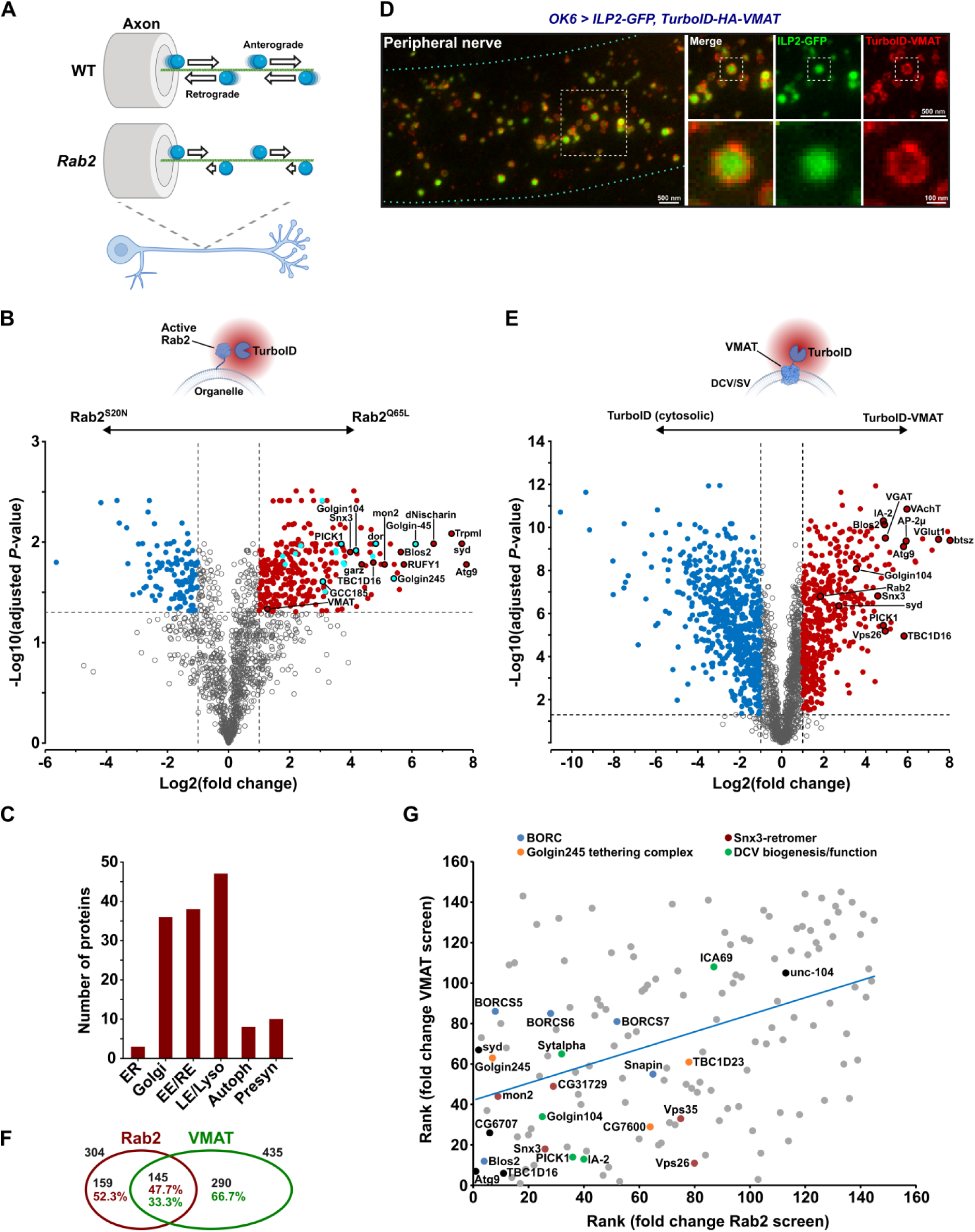
In vivo neuron-specific PDB/MS in Drosophila for detection of proteins interacting with active Rab2 and the DCV membrane protein VMAT. **A**, Schematic illustrating the effect of Rab2 loss on axonal transport of DCVs in flies ^16^. In wild type motor axons, DCVs (*blue spheres*) are transported bidirectionally with similar anterograde and retrograde flux. In *Rab2* null mutants, retrograde flux is strongly reduced, while anterograde flux is partially reduced. **B**, Volcano plot showing the fold change and Student’s *t*-test statistics of biotinylated protein label free quantification (LFQ) intensities from flies with pan-neuronal expression of TurboID-Rab2^Q65L^ (*elav > ILP2-GFP, 2xHA-TurboID-Rab2^Q65L^*) relative to TurboID-Rab2^S20N^ (*elav > ILP2-GFP, 2xHA-TurboID-Rab2^S20N^*). Proteins significantly upregulated in the active Rab2 condition (fold change > 2, FDR-adjusted *P*-value < 0.05) are highlighted in *red,* known Rab2 effectors in *light blue* (see also Fig. S1). **C**, Distribution in subcellular neuronal compartments of proteins specifically enriched in flies expressing TurboID-Rab2^Q65L^ compared to TurboID-Rab2^S20N^. ER, endoplasmic reticulum; EE, early endosomes; RE, recycling endosomes; LE, late endosomes; Lyso, lysosomes; Autoph, autophagosomes; Presyn, presynapse. **D**, Representative STED image showing the distribution of ILP2-GFP and hemagglutinin (HA)-tagged TurboID-VMAT fusion protein in motor axons in peripheral nerve A7 of third instar larva. Scale bars: *left*, 500 nm; *right*, 500 nm, 100 nm (inset). **E**, Student’s *t*-test of biotinylated protein LFQ intensities from flies with pan-neuronal expression of TurboID-VMAT (*elav > ILP2-GFP, TurboID-HA-VMAT*) relative to free cytosolic TurboID (*elav > ILP2-GFP, TurboID*). Proteins significantly enriched in the TurboID-HA-VMAT condition (fold change > 2, FDR-adjusted *P*-value < 2) are highlighted in *red*. **F**, Venn diagram of the overlap between proteins enriched both in flies expressing TurboID-Rab2^Q65L^ and TurboID-VMAT. **G**, Relationship between proteins significantly enriched in TurboID-Rab2^Q65L^ and TurboID-VMAT flies, when ranked by enrichment level. Spearman’s rank correlation 0.422, *P* < 10^-6^.

Strikingly, the second-most enriched protein for active Rab2 (∼200 fold enrichment over inactive Rab2, **Fig. 1B**) was Sunday Driver (Syd), the fly ortholog of mammalian JIP3 and JIP4, which function as activating adaptors for dynein ^35, 36^ and also bind kinesin-1 ^37, 38^. Other notable highly enriched hits were RUFY1/CG31064 (∼50 fold enrichment; from hereon called RUFY), the fly ortholog of the mammalian RUFY1-4 family of coiled-coil proteins, recently shown to link Arl8 and Rab14 to dynein, and possibly to function as dynein activating adaptors ^39, 40, 41^; and the ortholog of the mammalian Rab14 effector Nischarin (CG11807, ∼100 fold enrichment; from hereon called dNischarin) that shows distant homology to the SKIP motor adaptor ^42^. Interestingly, RUFY1 and RUFY2 were previously detected as unconfirmed potential effectors of human Rab2A, using the MitoID PDB protocol ^43^.

As a separate strategy to identify motor adaptors and their recruitment factors responsible for DCV motility, we also sought to determine the *in vivo* surface proteome of neuronal DCVs using the DCV-resident membrane protein Vesicular Monoamine Transporter (VMAT) as PDB bait. To this end, we generated a transgene encoding Drosophila VMAT fused through its cytosolic N-terminal tail to TurboID. Nanoscopic examination using stimulated emission depletion (STED) microscopy showed that when expressed in larval motor neurons together with the lumenal DCV cargo marker ILP2-GFP ^1^, most TurboID-VMAT decorates the limiting membrane of ILP2-positive DCVs, which were ∼130 nm in diameter (135±2.0 nm, mean±s.e.m., *n* = 1026 vesicles). The association of VMAT with DCVs was most clearly seen in axons (**Fig. 1D**), where the density of organelles is relatively low, but was also observed in somata and in synaptic boutons (**Fig. S1C**). In boutons, TurboID-VMAT was also present in ∼50 nm wide (52±1. nm, *n* = 25) punctate structures, possibly corresponding to small synaptic vesicles (SVs) (**Fig. S1C**). Comparative quantitative MS of biotinylated proteins from flies pan-neuronally expressing either TurboID-VMAT, or a free cytosolic TurboID control transgene, yielded ∼450 proteins significantly enriched in the TurboID-VMAT line (**Fig. 1E, Supplementary Data 3)**. Among the PDB hits enriched for TurboID-VMAT were well-known DCV membrane and peripheral membrane proteins such as IA-2 ^44^ and bitesize/Granuphilin/SYTL4 ^45, 46, 47^ (with the latter showing the highest enrichment of all proteins), as well as proteins involved in DCV biogenesis (**Fig. 1E, S1B)**. We also observed a strong enrichment for SV and endocytic proteins, consistent with VMAT also being targeted to SVs ^48^. Both Syd and Rab2 were also significantly enriched in the TurboID-VMAT dataset (**Fig. 1E, Supplementary Data 3)**.

Overall, the Rab2^Q65L^- and VMAT-enriched protein sets overlapped quite substantially (**Fig. 1F**), consistent with Rab2 and VMAT functioning within the same compartment(s). Moreover, we observed a significant correlation between enrichment levels across the two PDB datasets (**Fig. 1G**). Interestingly, among the highly enriched proteins in both the Rab2^Q65L^ and VMAT screens were many components of the Snx3-Retromer endosomal recycling complex ^49, 50^, a TGN vesicle tethering/fusion complex composed of TBC1D23, FAM91A1 (CG7600) and the Rab2-effector Golgin245 ^51^ (**Fig. 1G**). This suggests that Rab2 may be involved in recycling of VMAT from endosomes to TGN. Other hits ranking high in both data sets included proteins related to DCV biogenesis and subunits of the BORC Arl8 activator complex (**Fig. 1G**), consistent with the critical role of Arl8 in DCV motility ^16^.

Together, these data indicate that the dynein adaptors Syd and RUFY are spatially closely associated with active Rab2 in fly neurons in vivo, and that Syd and Rab2 may be present together at the surface of DCVs.

### Syd interacts with Rab2 via its RH2 domain and behaves as a Rab2 effector

To test if the high levels of PDB enrichment reflected physical interactions between Rab2 and Syd, RUFY and dNischarin, we performed co-immunoprecipitation (Co-IP) experiments using epitope-tagged versions of these proteins expressed in HEK293 (HEK) cells. Only myc-tagged full-length or truncated Syd (see below) was able to co-precipitate Rab2^Q65L^ in appreciable amounts, with RUFY and dNischarin producing yields barely above background (**Fig. 2A**). The Rab2:Syd interaction was relatively fragile, requiring a saponin-based lysis/binding buffer to achieve noticeable co-IP yields **(Fig. S2A)**. This is likely why this interaction was not found in earlier Rab:effector affinity-proteomic screening ^30^ and may indicate that it requires additional protein or lipid components. However, in Co-IP experiments Syd bound much stronger to active compared to inactive forms of Rab2, thus behaving as a classical Rab GTPase effector (**Fig. 2B, S2B)**. We therefore continued the investigation of the Rab2:Syd interaction.

**Figure 2.**
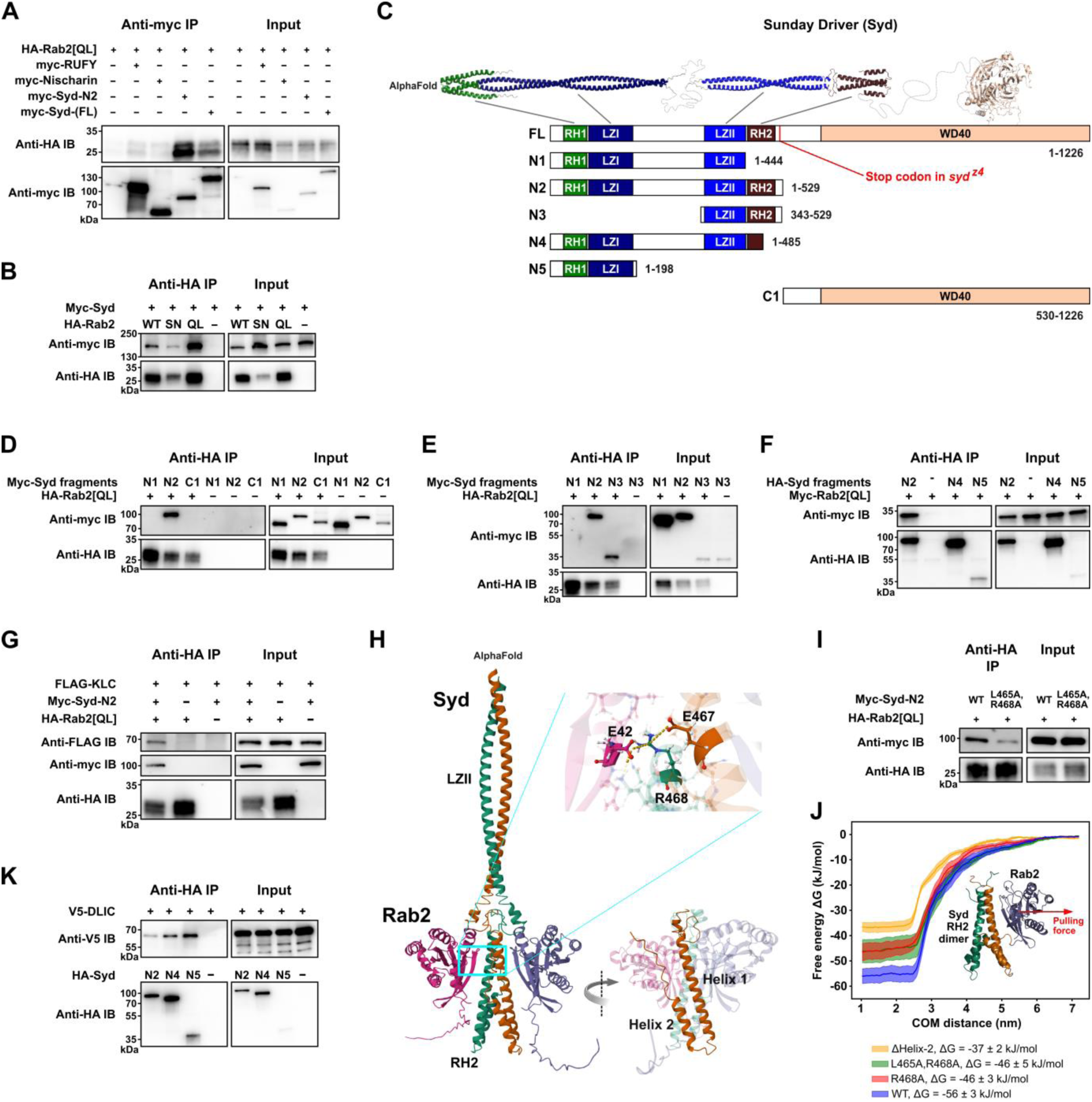
Syd binds active Rab2 via the RH2 domain and also binds kinesin-1 and dynein motors. Co-IP experiments performed on lysates from HEK cells transfected with constructs encoding epitope-tagged Drosophila proteins, and MD simulation of the Syd:Rab2 interaction. **A**, Myc-tagged full length Syd and truncated N2-Syd (Syd^1-529^) co-immunoprecipitates HA-tagged GTP-locked, consitutively active Rab2^Q65L^. In comparison, co-precipitation of HA-Rab2^Q65L^ by myc-tagged dNischarin and RUFY is near background levels. **B**, Wild type HA-Rab2 and HA-Rab2^Q65L^ co-immunoprecipitate myc-Syd more efficiently than GDP-locked, inactive HA-Rab2^S20N^. **C**, Structure of Syd. *Top*, Expected structure of Syd homodimer assembled from three separate AlphaFold predictions. The WD40 domain of only one Syd monomer is shown. *Bottom*, Schematic representation of the domain architecture of full length Syd (isoform A, UniProt Q9GQF1) and of truncated Syd variants. **D**, **E**, Co-IP of myc-tagged Syd fragments C1 and N1-N3 (shown in C) by HA-Rab2^Q65L^. Only myc-N2-Syd and myc-N3-Syd, which contain the RH2 domain, co-precipitate with Rab2. **F**, Co-IP of myc-Rab2^Q65L^ by HA-tagged Syd fragments N2, N4 and N5. **G**, FLAG-tagged Klc co-immunoprecipitates in complex with myc-N2-Syd and HA-Rab2^Q65L^ but not with HA-Rab2^Q65L^ alone. **H**, AlphaFold model of Syd LZII-RH2 (Syd^359-526^) dimer in complex with two copies of Rab2. *Inset*, R468 in Syd RH2 helix 1 is predicted to engage in ionic interactions with E42 in the Rab2 switch I region, and E467 in the other RH2 monomer helix 1. **I**, Comparison of co-IP of wild type myc-N2-Syd and mutant myc-N2-Syd^L465A,R468A^ with HA-Rab2^Q65L^. **J**, MD simulation. The Syd- RH2 dimer and one Rab2 moiety from the AlphaFold prediction in H were isolated in silico and pulled apart (*red arrow* represent pulling force direction) to estimate the free energy of Rab2:Syd-RH2 binding. Free energy curves (mean and standard error) as a function of Rab2:RH2 center-of-mass distance were calculated for wild type LZII-RH2 (*blue*), RH2^L465A,R468A^ (*green*), RH2^R468A^ (*red*) and RH2^ΔHelix-2^ (*yellow*) which was truncated after V504 removing the entirety of helix 2. **K.** Co-IP of V5-tagged DLIC with HA-tagged Syd fragments N2, N4 and N5. Note the higher co-IP efficiency for N4-Syd, where the C-terminal half of the RH2 domain (see Fig. S2C) is absent, compared to N2-Syd, which has an intact RH2 domain.

Structurally, Syd-family proteins (Syd/JIP3/JIP4) are large homodimers composed of an N-terminal region dominated by stretches of coiled-coil, followed by a C-terminal WD40 domain of unknown function (**Fig. 2C**). The N-terminal half of Syd/JIP3/4 contains the RILP homology domains 1 and 2 (RH1 and RH2; also found in the distantly related RILP/RILPL family of adaptors), flanking two short leucine zipper domains (LZI and LZII) separated by a lengthy unstructured region (**Fig. 2C**). The N-terminal RH1-LZI region of JIP3 binds in the cleft formed between dynein and dynactin and is sufficient to activate dynein motility ^35^. The more downstream LZII-RH2 region binds the kinesin-1 light chain (Klc) and Arf6 via the LZII domain ^38, 52, 53, 54^, and Rab8, 10 and 36 via the RH2 domain ^55, 56^ and is thought to be responsible for cargo binding.

Consistent with this pattern, truncation mapping showed that an intact RH2 domain is required for Rab2 binding to Syd in co-IP experiments (**Fig. 2C-F**). Furthermore, while the isolated RH2 domain failed to express in HEK cells, a fragment consisting of only the LZII and RH2 domains (N3-Syd) was sufficient to be precipitated by active Rab2^Q65L^ (**Fig. 2E**). Notably, the Syd^1-529^ fragment (N2-Syd) containing all N-terminal coiled-coil regions, but lacking the C-terminal region containing the WD40 domain, precipitated Rab2^Q65L^ better than full-length (FL) Syd (**Fig. 2A**), suggesting that the WD40 domain exerts an inhibitory influence on this interaction. Also, while Rab2^Q65L^ alone did not precipitate Drosophila Klc, it did so when co-overexpressed with N2-Syd (**Fig. 2G**). This shows that Syd can bridge active Rab2 and molecular motors, and that Rab2 binds Syd in a way that does not interfere with Klc binding at the LZII domain.

AlphaFold multimer modelling of the Syd LZII-RH2 dimer (Syd^359–526^) together with two Rab2 chains templated on the crystal structure of active GppNHp-bound Rab2 (PDB: 4rke)^57^ yielded a predicted structure or the Rab2:Syd^LZII-RH2^ complex (**Fig. 2H**). It broadly resembles the crystal structures of Rab7 bound to the RILP RH2-domain (PDB: 1yhn)^58^ and phospho-Rab8a bound to the RILPL2 RH2-domain (PDB: 6rir)^56^. In the AlphaFold prediction, the two Syd-RH2 monomers, each composed of two roughly antiparallel alpha helixes (Helix 1 and Helix 2), together form a four-helix bundle, with the N-terminal part of each Helix 1 constituting the main interaction surface with Rab2 (**Fig. 2H**). Alanine substitution of the highly conserved residues L465 and R468 **(Fig. S2C)** in Helix α1 that were predicted to form contacts with the Rab2 switch regions (**Fig. 2H**) substantially reduced Syd-N2 precipitation by Rab2^Q65L^ (**Fig. 2I**). These results were recapitulated by molecular dynamics (MD) simulations of the Rab2:Syd^LZII-RH2^ AlphaFold structure, which predict that the R468 residue significantly contributes to Rab2:Syd-RH2 binding energy (**Fig. 2J**). In addition, mutation of a cluster of nine conserved residues in Helix 2 (N2-Syd^7A^) **(Fig. S2C, D)**, or deletion of the entire Helix 2 together with the seven most C-terminal residues of Helix α1 (N4-Syd) (**Fig. 2F**) weakened and entirely abolished N2-Syd precipitation by Rab2^Q65L^, respectively. Together with MD modelling predicting that removal of Helix 2 would result in a substantial decrease in binding energy **(Fig. S2J)**, these data indicate that Helix 2, like Helix 1, plays an important role in Rab2 binding, perhaps by stabilizing the Helix 1 dimer conformation.

A critical early step during the JIP3-assisted assembly and activation of the dynein-dynactin complex is binding of the dynein light intermediate chain (DLIC) to the JIP3 RH1 domain ^35^. We confirmed that the Syd-dynein interaction is conserved in flies by showing that N2-Syd can co-precipitate Drosophila DLIC (**Fig. 2K**). Moreover, further truncated Syd variants lacking the RH2 Helix 2 (Syd^1-485^, N4-Syd) or containing only the RH1-LZI region (Syd^1-1^^98^, N5-Syd) precipitated DLIC noticeably better than N2-Syd **(Fig. S2C)**. These data mirror recent findings showing that the JIP3 RH1 domain is autoinhibited by a conserved motif in the RH2 domain ^35^.

Collectively, these data suggest that active Rab2 interacts with Syd via the cargo-binding Syd RH2 domain and that this interaction is compatible with kinesin-1 and dynein recruitment by Syd.

### Loss of Syd and Rab2 produce similar effects on axonal transport of DCVs and lysosomes

Loss of Syd causes a strong defect in the axonal transport of synaptic vesicle proteins in Drosophila larvae ^38^. If Syd constitutes the link between active DCV-localized Rab2 and molecular motors, one would expect disruption of Syd and Rab2 to produce similar effects on DCV transport. We therefore examined mid-axon transport of ILP2-GFP-positive DCVs in L3 larval motor neurons, targeted by the OK6-Gal4 driver, using the same live confocal imaging method employed in our previous study ^16^. During time-lapse imaging of a 130 µm long stretch of the A7 peripheral nerve in fillet-dissected larvae, we photobleached two 60 μm long flanking segments around a 10 μm central region of the nerve and then recorded the movement of fluorescent DCVs initially located in the unbleached center as well as those entering laterally from outside the field of view (**Fig. 3A**). This approach allows the examination of both the transport in the anterograde and retrograde directions (distinguishable because motor neuron axons are uniformly oriented towards the periphery) and the abundance of static vesicles. After converting the time-lapse movies of DCV transport to kymographs, we plotted the frequency distribution of the angle between vesicle trajectories and the vertical axis in the kymographs, using a fast Fourier transform algorithm (see Materials and Methods). The resulting “directional distributions” (**Fig. 3B-D**) are amenable to high throughput analysis and provide a convenient overview of the relative amounts of anterograde and retrograde transport (left- and rightmost peaks, respectively) and the relative amount of static cargo (middle peak at 0°). The relative amplitudes of the anterograde/retrograde and static peaks in the directional distributions (**Fig. 3B-F**) aligned well with absolute vesicle flux and static vesicle counts across the genotypes examined (**Fig. 3G, H**).

**Figure 3.**
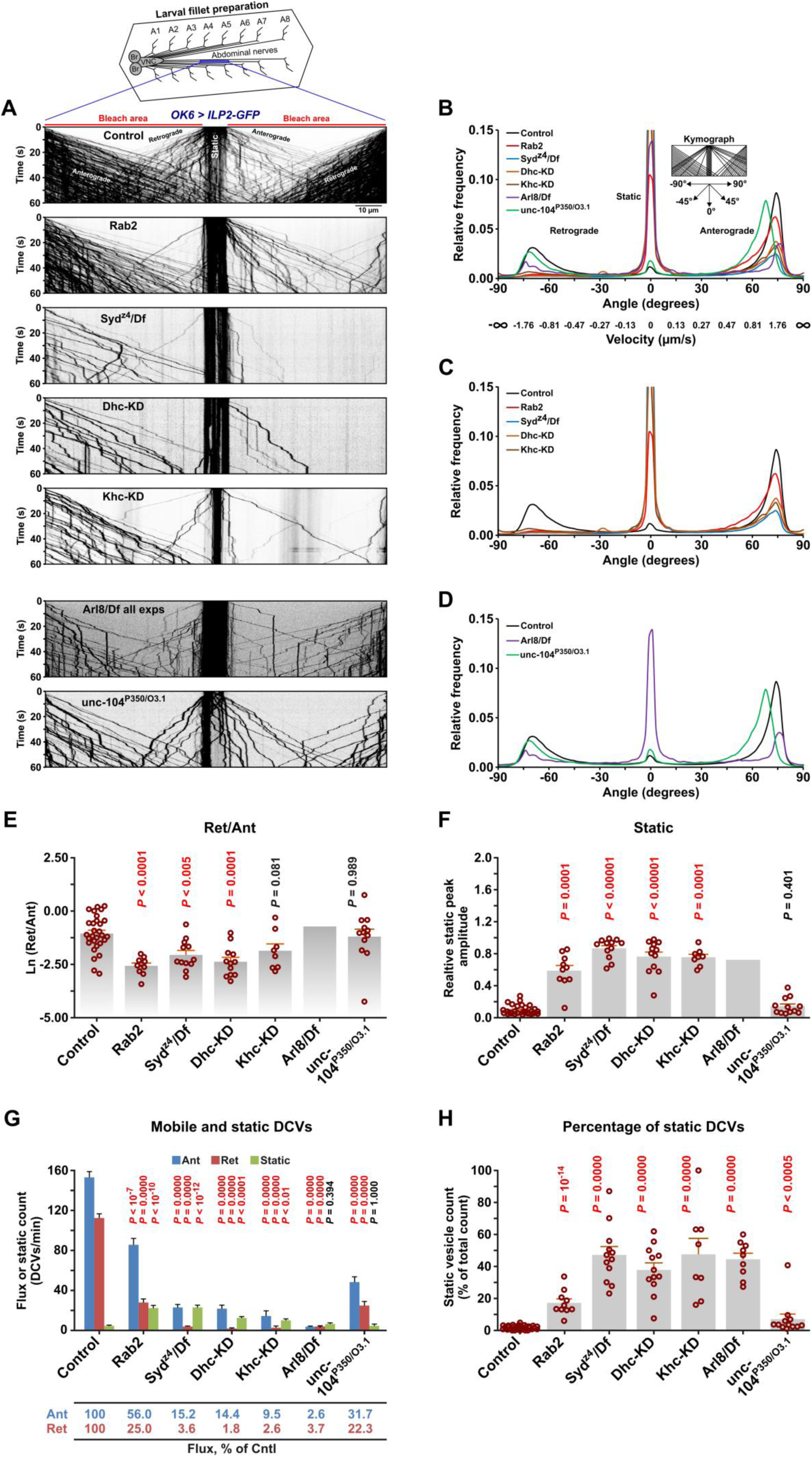
Disruption of Rab2, Syd and dynein result in similar DCV axonal transport phenotypes. **A**, Representative kymographs showing transport of ILP2-GFP-positive DCVs in motor axons in the A7 nerve of third instar larvae with the indicated genotypes. KD, motor neuron-specific knockdown (driven by *OK6*-Gal4). Time-lapse confocal imaging was performed immediately after bleaching the areas indicated with *red bars*. Each kymograph depicts a single recording, except for *Arl8/Df*, where 20 superimposed recordings are shown (see Methods). Scale bar: 10 µm. **B-D**, Directional distributions showing the relative frequency of DCV transport velocities, expressed as the angle between the DCV trajectories and the vertical axis in the kymographs in A (inset in B). Actual DCV velocities converted from angles have been added to the x-axis in B. For each genotype, the directional distribution is averaged from *n* larvae, where *n* is equal to the number of data points in E, F, and H. **E**, Logarithmic ratio of the retrograde to anterograde peak amplitude in the directional distributions in B-D (the retrograde and anterograde peak amplitude is the maximal relative frequency of angles less than -46° and more than 46°, respectively). **F**, The static peak amplitude relative to the sum of the static, retrograde and anterograde peak amplitudes (the static peak amplitude is the maximal relative frequency of angles within the central interval -13° to 13°). **G**, Counts of DCVs entering from the sides into the field of view in the anterograde or retrograde directions, and of static vesicles in the central unbleached area. Counts were done over 30 s and multiplied by two, converting the dynamic vesicle counts to DCV flux in vesicles per minute. **H**, Percentage of static vesicle counts relative to total vesicle counts for each genotype in G. Statistical tests of the data in E-H (represented as mean+SEM) are specified in the Source File. Results involving *Arl8/Df* represent re-analysis of data published earlier ^16^.

In wild type animals, apart from a smaller static component, we observed large anterograde and retrograde DCV fluxes with a moderate excess of anterograde transport (**Fig. 3A, B-E, G**), consistent with axonal DCV circulation. As reported previously ^16^, loss of Rab2 was associated with a pronounced DCV transport defect with a relative increase in the static vesicle signal, moderate reduction in anterograde transport and a disproportionately severe reduction in retrograde transport (**Fig 3A-C, E-H, Video S1**). The latter was evidenced by the almost complete disappearance of the sharp peak associated with retrograde transport in the directional distribution (**Fig. 3B, C**). Importantly, a qualitatively similar, but more severe phenotype was observed in larvae hemizygous for the *syd^z4^* strong loss-of-function allele ^38^ (*syd^z4^/Df*), or when dynein function was impaired in motor neurons by RNAi-mediated depletion of the dynein heavy chain (Dhc) (**Fig 3A-C, E-H, Video S1)**. The movement speed of the remaining retrograde vesicles in *Rab2* null and *syd^z4^/Df* animals was also considerably slower compared to wild type, similar to Dhc-depleted animals **(Fig. S3B)**. Suppression of the kinesin-1 motor by kinesin-1 heavy chain (Khc) depletion also resulted in a transport defect characterized by a selective deficit in retrograde DCV traffic (**Fig. 3A-C, E-H, S3A, B, Video S1)**, and featuring the appearance of prominent axonal DCV-filled focal accumulations (**Fig. 4A**). The selective effect of kinesin-1 dysfunction on the retrograde DCV flux in flies has been reported previously ^15^ and may be due to progressive stalling during anterograde transport, although there are also indications of direct and indirect co-dependence between kinesin-1 and dynein ^59, 60^. These findings suggest that Rab2 and Syd are involved in the function of dynein and/or kinesin-1 during DCV transport.

**Figure 4.**
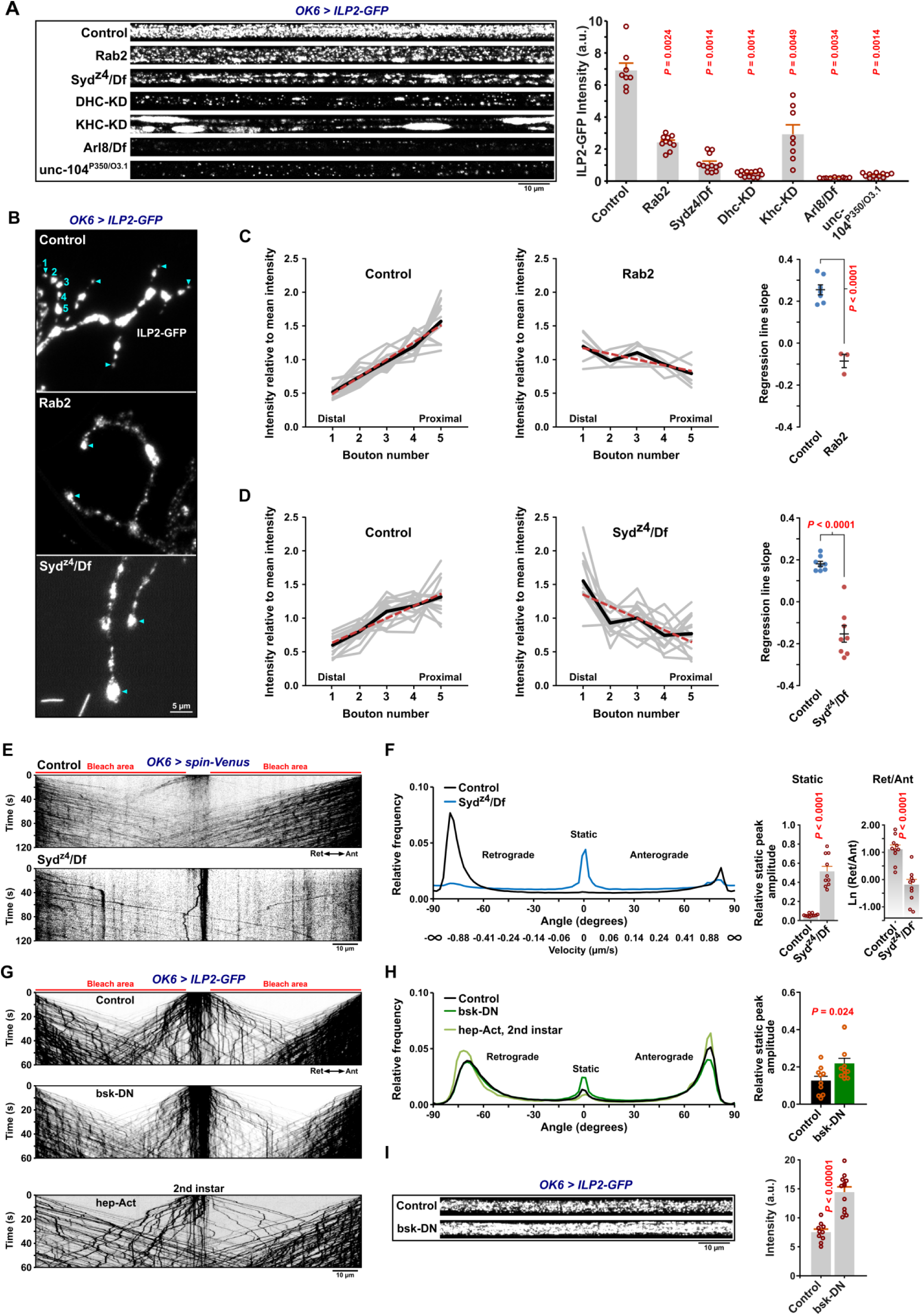
Axonal levels and distribution of DCV cargo, disrupted transport of lysosomal organelles in *syd* mutants, and effects of altered JNK activity on DCV transport. **A**, *Left*, Representative pre-bleach confocal micrographs of the A7 nerve in third instar larvae expressing ILP2-GFP in motor neurons. *Right*, Quantification of the axonal ILP2-GFP signal intensity. A.u., arbitrary units. Scale bar: 10 µm. **B**, Neuromuscular junction of muscle fiber 6 and 7 in control, *Rab2*, and *syd^z4^/Df* larval fillets. *Numbers in blue* indicate the distal five boutons in a single branch of a motor neuron ending. *Blue triangles* indicate the distalmost bouton in the same and other branches. Scale bar: 5 µm. **C**, **D**, *Left and middle*, ILP2-GFP signal intensity in the distal five boutons of individual branches. *Thick black lines* represent the mean intensity for each bouton number. *Dashed red lines* were produced by linear regression. *Right*, Regression line slopes. The mean±SEM is indicated. **E**, Representative kymographs showing transport of Spinster-positive organelles in A7 motor axons of control and *Syd^z4^/Df* third instar larvae. Scale bar: 10 µm. **F**, *Left*, Directional distributions of Spinster-positive organelle transport velocities, expressed as angles, cf. Fig. 3B-D. Actual velocities converted from angles have been added to the x-axis. For each genotype, the average directional distribution from *n* larvae is shown, where *n* is equal the number of data points in the bar graphs (*right*), which depict the relative static peak amplitude and the logarithmic ratio of retrograde to anterograde peak amplitude. **G**, Kymographs of DCV transport in motor axons of third instar control and *bsk-*DN larvae, and in second instar *hep-Act* larvae. Scale bar: 10 µm. **H**, Directional distributions derived from G. For control and *bsk-*DN larvae, the distributions are averaged from *n* larvae, where *n* is equal to the number of data points in the bar graph at the *right* showing relative static peak amplitude. For *hep-Act*, the directional distribution is the mean of three recordings from two larvae. **I**, Axonal pre-bleach intensity of control and *bsk-DN* larvae. Scale bar: 10 µm. Bar graphs in F, H, and I represent the mean+SEM. For the quantitative data in A, C, D, F, H, and I each data point represents one larva, and statistical tests are specified in the Source File.

In contrast, larvae carrying a heteroallelic combination of the *unc-104^P3^*^50^ null and *unc-104^O3.1^* hypomorphic mutations of the fast anterograde kinesin-3 family Unc-104 motor (ortholog of mammalian KIF1A/B/C) displayed a qualitatively different axonal transport phenotype characterized by a severe reduction in axonal DCV content (**Fig. 4A**) due to a failure of cell body exit ^13^, combined with more symmetrical bidirectional fluxes of remaining axonal DCVs **(Fig. 3A, B, D-H, Video S1)**. *unc-104^P3^*^50^*/unc-104^O3.1^*animals also displayed a strong reduction in the mean anterograde vesicle speed **(Fig. S3A)** reflected in a pronounced leftward shift (towards lower velocities) of the anterograde peak in the directional distribution (**Fig. 3B, D**). This is consistent with Unc-104 being responsible for fast anterograde DCV movement ^13, 15^. Applying the same analysis to previously published ^16^ DCV axonal transport recordings in animals lacking Arl8, thought to be responsible for Unc-104 activation ^17, 18, 61^, also showed a similar phenotype with very strong, but symmetrical reductions in the bidirectional DCV flux **(Fig. 3A, B, D-H)**. Interestingly, unlike *unc-104* mutants, but resembling Rab2, Syd, Dhc, and kinesin-1 deficient animals, *Arl8* nulls also displayed a large relative increase in the static DCV component suggesting that it may also be involved in regulation of kinesin-1 and dynein motors **(Fig. 3A, C-H)**.

Disruption of dynein function causes a pronounced accumulation of excess DCVs in the distal-most boutons of larval motor terminals ^1^. Consistent with this, in type Ib motor terminals on larval muscles 6 and 7, both *Syd* and *Rab2* mutants displayed a clear reversal of the usual trend of decreasing bouton content of ILP2-GFP in more distal boutons (**Fig. 4B-D**), although the Syd phenotype was again more severe. This further indicates that Rab2 and Syd are required for retrograde dynein-dependent transport.

Besides DCV transport, Rab2 is required for axonal transport of lysosomes and early/late endosomes in flies ^16^. JIP3/4 are also well known to mediate lysosomal motility in mammals ^36, 62, 63, 64^. We found that transport of lysosomes labelled with Spinster-Venus was severely disrupted in motor axons of *syd^z4^/Df* mutant larvae, with less bidirectional transport and a relative increase in static organelles (**Fig. 4E, F**). While direct comparisons with previously recorded data for *Rab2* nulls **(Fig. S3C, D)** ^16^ are difficult due to the use of different markers (Spinster-Venus vs Spinster-GFP, necessitated by the different chromosomal locations of *Rab2* and *syd*), the disruptions in lysosomal transport appeared to be similar in *Rab2* and *syd* mutants, albeit with a stronger defect in *syd^z4^/Df*.

In conclusion, loss of Rab2 and Syd produce qualitatively similar axonal transport defects for DCVs and lysosomes, although the Syd mutant phenotype is more severe. Moreover, the Rab2 and Syd-related DCV transport defect is consistent with a disruption of dynein-mediated retrograde motility.

### The Arl8 effector RUFY cooperates with Syd to drive retrograde axonal transport of DCVs

The less severe *Rab2* null DCV transport phenotype compared to the *syd* mutant phenotype strongly suggests the presence of additional vesicular Syd-recruitment factors. Syd/JIP3/4 proteins bind small GTPases Arf6 ^54^, Rab36, Rab8 and LRRK-phosphorylated Rab10 ^55, 56^, of which Rab8 and Rab10 were enriched in our VMAT-specific PDB dataset **(Fig. S4A, Supplementary Data 3)**. Moreover, Arf6 and Rab10 control JIP3/4-mediated axonal transport of mammalian autolysosomes ^36, 65^. We tested larvae with null mutations in Arf6, Rab8 and Rab10, or homozygous for a transposon insertion allele for the fly Rab36 ortholog, RabX5, but found no obvious disruption of axonal DCV transport **(Fig. S4B)**. Of multiple Rabs (besides Rab2) enriched in VMAT-proximity proteomics **(Fig. S4A),** only Rab1 or Rab11 produced any effect on axonal DCV transport when disrupted by mutation or motor neuron-specific depletion **(Fig. S4C-G)**. However, since no physical interactions between Rab1 or Rab11 and Syd family proteins have been reported, and the Rab1 and Rab11 depleted animals did not develop beyond late first or early second instar, we did not pursue this line of inquiry further. We also tested the ortholog of the mammalian TMEM55A/B transmembrane proteins (CG6707), which ranked high in both Rab2- and VMAT-PDB datasets (**Fig. 1G**). TMEM55B mediates the recruitment of JIP4 for dynein mediated lysosomal motility in mammals ^66^. However, depletion of CG6707 with two independent RNAi-transgenes did not affect axonal transport of DCVs **(Fig. S5A, B )**.

In our search for more components of the retrograde motor complex we next focused our attention on dNischarin and RUFY, which also ranked high in the Rab2-PDB dataset but did not interact strongly with Rab2 (**Fig. 1B, 2A)**. Interestingly, while motor neuron-specific dNischarin depletion produced no effect **(Fig. S5A, B)**, depletion of RUFY with either of two independent RNAi transgenes caused a pronounced DCV transport defect characterized by a selective reduction in retrograde movement and a relative increase in static DCV cargo (**Fig. 5C, D**). As almost all retrograde DCV transport is already lost in *syd^z4^/Df* animals (**Fig. 2A-C, G)**, we reasoned that Syd and RUFY function as part of the same mechanism to recruit/activate dynein. Although mammalian RUFY proteins have been proposed to function as dynein activating adaptors in their own right ^40, 41^, at least one, RUFY3, also binds JIP4 ^39^. Consistent with this observation, we found that fly RUFY immunoprecipitates Syd when the two proteins are expressed together in HEK cells. This interaction appears to depend on the Syd C-terminal WD40 domain, as the truncated Syd-N2 (Syd^1-529^) variant missing this region showed very little binding to RUFY compared to full-length Syd (**Fig. 5A**). Similar to its mammalian orthologs, which are ARL8A/B effectors ^39, 40, 41^, RUFY also bound fly Arl8 in co-IP experiments (**Fig. 5B**). The presence of Arl8 also increased the interaction between RUFY and Rab2^Q65L^, similar to the reported effect of Arl8b on the interaction between RUFY1 and the Rab2-related ^67^ Rab14 GTPase in mammals ^41^ (**Fig. 5B**). This suggests that Syd may be stabilized on the DCV surface by a combination of interactions including Rab2-binding via the RH2 domain, and WD40 domain-dependent binding to RUFY, which itself is recruited to the vesicles via Arl8 and possibly Rab2 (**Fig. 5J**).

**Figure 5.**
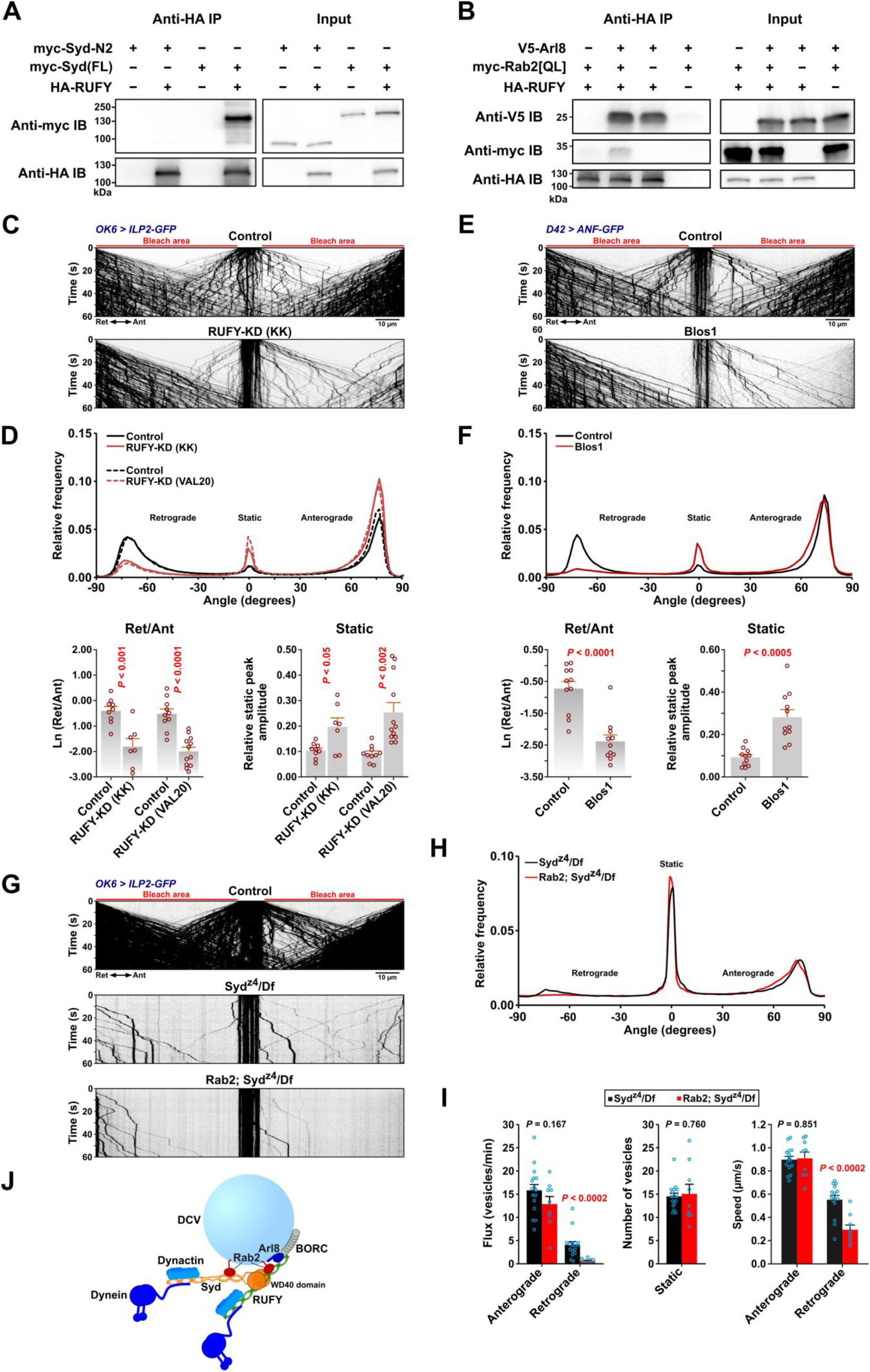
RUFY interacts with Syd, Arl8 and Rab2 and is required for axonal DCV transport. **A, B**, Western blots of the indicated co-IP eluates (∼40% eluate volume) and HEK cell lysates (∼1% reaction volume). **A**, HA-tagged RUFY co-immunoprecipitates myc-tagged full-length Syd but not truncated N2-Syd (Syd^1-529^) that lacks the C-terminal WD40 domain. **B,** HA-RUFY co-immunoprecipitates V5-tagged Arl8, and co-expression of V5-Arl8 enhances co-immunoprecipitation of myc-Rab2^Q65L^ with HA-RUFY. **C**, Representative kymographs showing transport of ILP2-GFP-positive DCVs in motor axons of control larvae and larvae subjected to motor neuron-specific knockdown of *RUFY*, driven by *OK6*-Gal4. Scale bar: 10 µm. **D**, *Top*, Directional distributions derived from C, averaged from *n* larvae, where *n* is equal to the number of data points in the bar graphs (*bottom*) that depict the logarithmic ratio of retrograde to anterograde peak amplitude, and the relative static peak amplitude. In C and D, “KK” and “VAL20” refer to *UAS-RNAi* lines from the KK collection and the VALIUM20 vector-based collection, respectively. For simplicity, only KK line data are illustrated in C. **E**, Representative kymographs showing transport of ANF-GFP-positive DCVs in motor axons of control and *Blos1* larvae. Scale bar: 10 µm. **F**, *Top*, Directional distributions derived from E, averaged from *n* larvae, where *n* is equal to the number of data points in the bar graphs (*bottom*) that depict the logarithmic ratio of retrograde to anterograde peak amplitude, and the relative static peak amplitude. **G**, Kymographs of transport of ILP2-GFP-positive DCVs in motor axons of control, *Syd^z4^/Df* single mutant, and *Rab2; Syd^z4^/Df* double mutant larvae. Scale bar: 10 µm. **H**, Directional distributions from G, averaged from *n* larvae, where *n* is equal to the number of data points in the bar graphs in I. **I**, Flux of dynamic vesicles, counts of static vesicles in the central unbleached region and speed of dynamic vesicles in *Syd^z4^/Df* single mutants, and *Rab2; Syd^z4^/Df* double mutants. **J,** Hypothetical model of the DCV dynein-dynactin recruitment complex, consisting of Syd and RUFY anchored to the vesicle by Rab2 and Arl8. Kinesin-1 bound by Syd and kinesin-3 regulated by Arl8-BORC are omitted. It is uncertain if RUFY can recruit and activate dynein-dynactin. Bar graphs in D, F, and I represent the mean+SEM, and each data point represents one larva. Statistical tests of the quantitative data in D, F, and I are specified in the Source File. For conversion of angles in D, F, and H to DCV velocities, see Fig. 3B. All data are from third instar larvae. Results in E and F represent re-analysis of data published earlier ^16^.

This model fits with our observations that *Arl8* nulls, besides showing a relatively symmetrical reduction in the extent of anterograde and retrograde transport due to a failure of cell body exit ^16^, also display a strong increase in the proportion of static DCVs **(Fig. 2A-D, F, H)** similar to the syd/dynein/kinesin-1 phenotypic group. An involvement of Arl8 in dynein regulation via RUFY can also explain our previously published data ^16^ showing that knockout of the critical BORC subunit, Blos1 ^68, 69^, produces a strikingly selective loss of retrograde axonal transport of ANF-positive DCVs **(reanalyzed in Fig. 5E, F**).

In addition to binding Arl8, RUFY proteins are also known Rab effectors; fly RUFY binds active Rab4 ^30^, and mammalian RUFY1 requires Rab14 for recruitment to endosomes ^41^. We observed that whereas Rab14 depletion did not affect DCV axonal transport, a small but significant increase in the static component resulted upon Rab4 depletion, suggesting that Rab4 may have a minor role in RUFY recruitment to DCVs **(Fig. S6)**.

A requirement for the Syd:RUFY interaction potentially explains the strong phenotype of the *syd^z4^* allele, which introduces a premature stop codon (at amino acid position 514 in isoform A) leading to the truncation of the entire Syd WD40 domain but leaving a protein roughly corresponding to our Syd-N2 construct ^38, 70^ (**Fig. 2C**), containing both the Rab2 binding RH2 domain and the dynein-dynactin activating region as defined by Singh et al. for JIP3 ^35^. Interestingly, although almost all retrograde (and most anterograde) transport is lost in *syd^z4^/Df* hemizygous larvae **(Fig. 3A-C, G)**, a small retrograde DCV flux remained (**Fig. 5G-I**). This residual retrograde flux was almost entirely eliminated in *Rab2; syd^z4^/Df* double mutants with the few remaining retrograde vesicles moving significantly slower, as would be expected if it was driven by lower affinity recruitment of the remaining Syd fragment by Rab2 alone (**Fig. 5G-I**). However, we cannot exclude an alternative model where RUFY functions directly as the activating adaptor for dynein and is recruited to DCVs by a combination of Syd (via the WD40 domain), Rab2 and Arl8 (**Fig. 5J**).

In conclusion, Syd and the Arl8 effector RUFY interact via the Syd C-terminal WD40 domain and together are required for retrograde axonal transport of DCVs. This implies that Arl8 and its activator, BORC, are also involved in retrograde dynein mediated DCV transport, in addition to fast anterograde transport mediated by kinesin-3.

### JNK promotes axonal DCV motility

Besides the vesicular motor adaptor role, Syd/JIP3/4 proteins function as important scaffolds and transport adaptors for the JNK MAP kinase ^71, 72, 73^. Active JNK is important for axonogenesis and is enriched in mature axons ^74^. Moreover, there are strong indications that JNK regulates Syd-linked motor activity or Syd-cargo binding ^70, 73^. We therefore tested if the activity level of the sole fly JNK ortholog, basket (bsk), affects axonal DCV-transport. Overexpression of dominant negative bsk (*bsk-DN*) in larval motor neurons increased the relative amount of static axonal DCV signal (**Fig. 4G-H**) accompanied by a doubling of total axonal DCV marker content, implying an increase in the total mid-axonal DCV population due to DCV stalling. Unfortunately, motor neuron overexpression of the constitutively active form of the upstream bsk activator hemipterous/MKK7 (hep-Act) blocked development around the L2 stage, making comparisons to controls or bsk-DN expressing larvae (which developed normally) difficult. However, hep-Act expressing larvae did not display any clear reduction in the motility of axonal DCVs (**Fig. 4G, H**). These results suggest that a baseline level of JNK activity is required to ensure normal axonal DCV circulation.

The LRRK2 kinase and its orthologs also bind to Syd family proteins ^75, 76^, and LRRK-phosphorylation of a subset of Rabs (including Rab8 and Rab10) on a conserved switch II residue promotes Rab binding to RH2 domains of RILP/RILPL and JIP3/4 ^56, 77^. However, when tested, animals lacking the sole fly LRRK ortholog did not show the DCV transport defects typically associated with Syd, RUFY, Rab2 or dynein dysfunction, although they did display a moderate decrease in the total axonal DCV cargo content **(Fig. S6A-F)**. The latter could be due to a perturbation of DCV biogenesis in *LRRK* nulls, as data from *C. elegans* suggests that LRRK is involved in protein and membrane sorting at the Golgi ^76^. The apparent lack of retrograde transport defects in *LRRK*-nulls is consistent with Rab10 being dispensable for DCV transport **(Fig. S4B, C)**, and the absence of any obvious effect on the Rab2^Q65L^:N2-Syd Co-IP yield during pharmacological inhibition of LRRK2 activity or overexpression of mutant gain-of-function LRRK2 in HEK cells **(Fig. S2E)**.

### DCV membrane proteins are missorted to ectopic vesicle aggregates in Rab2-null cell bodies

Apart from the lysosome- and transport-related functions of Rab2, it is involved in DCV biogenesis in *C. elegans* ^24, 25, 27^. Recent work in flies has further revealed that Rab2 plays a crucial role in generating specialized transport organelles that carry presynaptic components from somata to synapses ^23^. We therefore examined in more detail the distribution of lumenal and transmembrane DCV cargo in the form of ILP2-GFP and HA-VMAT, respectively, in Drosophila motor neurons in wild type and *Rab2* null backgrounds.

As described above (**Fig. 1D**), VMAT mostly co-localized with abundant ILP2-positive DCVs when observed using STED microscopy in wild type larval motor neuron cell bodies in the ventral nerve cord (VNC) (**Fig. 6A, B**). Strikingly, in *Rab2* null cell bodies the number of DCVs was severely reduced, and most VMAT signal was no longer associated with ILP2 but instead segregated into dense, roughly spherical aggregates averaging ∼0,5 µm in diameter (**Fig. 6A, B, S7E**). These VMAT aggregates had a granular internal structure, suggesting that they are composed of smaller vesicles (∼50 nm in diameter, see Methods) and are likely identical or similar to SV protein-containing tubulo-vesicular clusters that were previously described in *Rab2* null larval neurons ^23^. These clusters, located in the vicinity the TGN, were proposed to form as the result of failure to fuse small elongated (40 x 60 nm) Golgi-derived transport vesicles containing SV-proteins with morphologically similar vesicles containing presynaptic active zone scaffolds ^23^. The VMAT^Y600A^ mutant that is not efficiently targeted to SVs due to the disruption of a tyrosine-based AP-2 clathrin adaptor complex binding site, but remains associated with DCVs ^48^, still formed aggregates in Rab2-nulls **(Fig. S7A, B)**. In addition, DCV-specific Synaptotagmin-α tagged with mCherry ^78^ also redistributed away from somatal DCVs in *Rab2* mutants, although it was much less prone to form the dense spherical aggregates seen for VMAT but instead accumulated in more loosely organized vesicle clusters **(Fig. S7C, D)**. In some instances, VMAT and Syt-α vesicle aggregates abutted clusters of ILP2-positive vesicles, which may represent immature DCVs or TGN regions from where DCV-budding takes place **(arrow in Fig. S7C)**.

**Figure 6.**
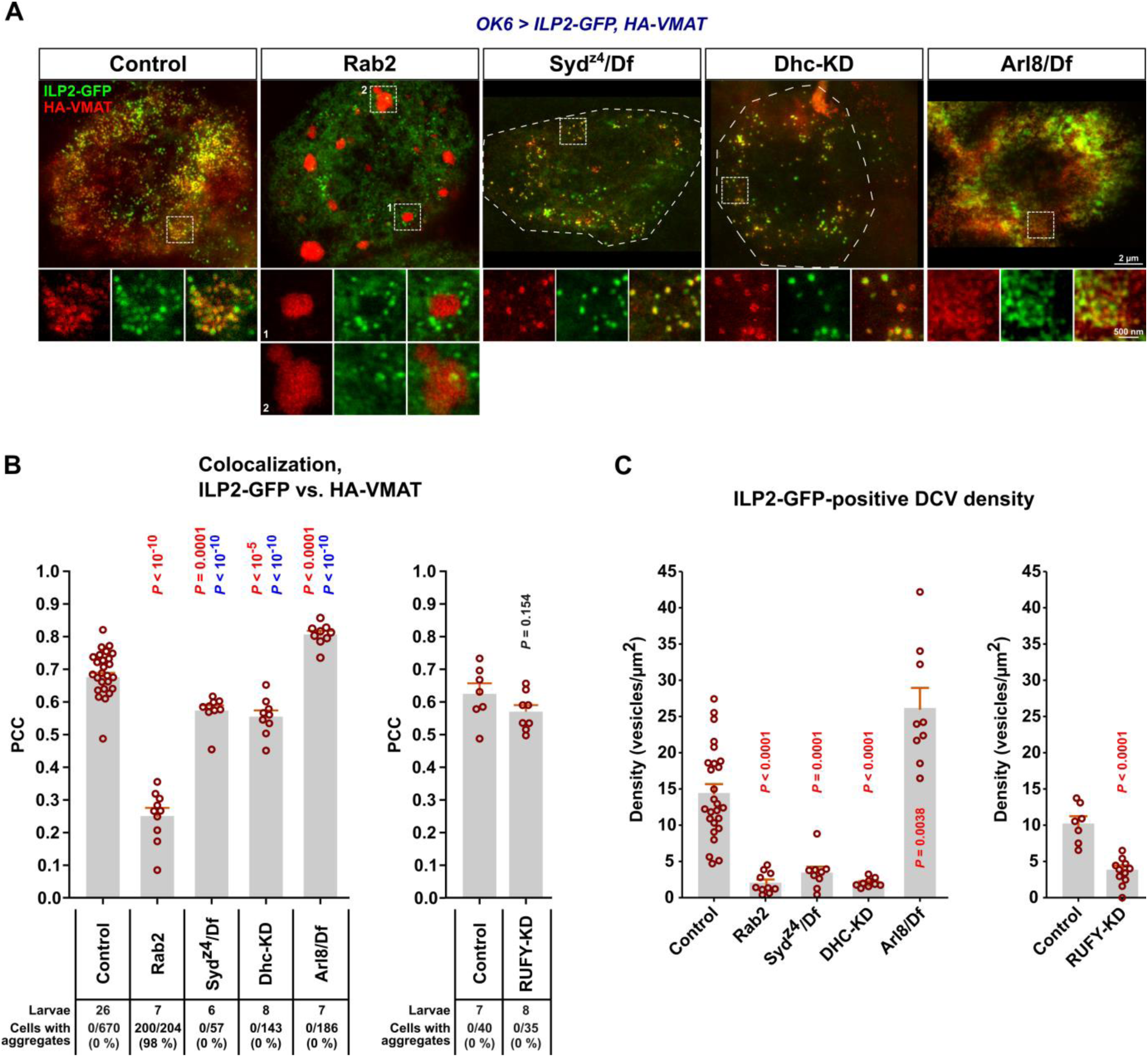
Missorting of HA-VMAT and reduction of DCV numbers in cell bodies of *Rab2* mutants. **A**, Representative STED images showing the distribution of ILP2-GFP and HA-tagged wild type VMAT in motor neuron cell bodies in VNCs of third instar larvae. KD, motor neuron-specific knockdown,. Scale bar: 2 µm, 500 nm (insets). **B**, *Left*, The co-localization of ILP2-GFP and HA-VMAT in A was quantified using Pearson’s correlation coefficient (PCC). *P*-values in *red* and *blue* result from comparison with the Control and *Rab2* group, respectively. *Right*, The correlation between ILP2-GFP and HA-VMAT in motor neuron cell bodies of *RUFY-KD* and control larvae. Counts of VMAT aggregates made in confocal images are shown (*numbers below the graphs*). **C**, *Left*, ILP2-GFP-positive vesicle densities in cell bodies for the larval genotypes in A, and in *RUFY-KD* larvae with controls (*right*). Bar graphs in B and C represent mean+SEM, and each data point represents one larva. Details of statistical tests of the data in B and C are contained in the Source File.

These findings show that, similar to SV- and active zone-proteins, DCV-specific membrane proteins aggregate in ectopic vesicle clusters in the somata of *Rab2* null fly neurons. We speculate that these vesicle clusters form as a result of their constituent transport vesicles being unable to fuse to maturing DCVs or the TGN in the absence of Rab2.

### Syd, RUFY and dynein are not responsible for Rab2-dependent sorting of DCV membrane proteins, but may play a role in biosynthetic transport of lumenal cargo

The findings described above raised the question of whether Syd and RUFY and their associated motors contribute to Rab2-dependent functions during DCV biogenesis in the cell body. Alternatively, it is possible that the *Rab2* null axonal transport phenotype is caused by a failure of Rab2-dependent sorting of critical DCV membrane proteins that serve as vesicular anchors for Syd and/or RUFY.

Neither *Syd^z4^/Df* larvae, nor larvae where RUFY or dynein were depleted by motor-neuron specific RNAi displayed the strong separation of the VMAT and ILP2 signals or the large VMAT aggregates characteristic of *Rab2* nulls (**Fig. 6A, B, S7F**). This shows that Syd, RUFY and dynein do not mediate the Rab2-dependent sorting of SV/DCV membrane proteins, and that this process must rely on one or more different Rab2 effectors.

Apart from the proximity and physical interactions between Rab2 and Syd (**Fig. 1B, 2)**, two lines of evidence suggest that the Rab2 axonal transport phenotype is not primarily due to the missorting of DCV membrane proteins. First, when we examined the axonal distribution of VMAT and ILP2 we found that *Rab2* null axons exhibited a dramatic accumulation of VMAT-containing vesicles and smaller irregularly shaped vesicle clusters that were not associated with ILP2-positive DCVs **(Fig. S7G, H)**, partially mirroring the cell body phenotype. However, the mean VMAT signal associated with individual axonal DCVs was not significantly reduced in *Rab2^Δ1^* compared to wild type **(Fig. S7H)**. This suggests that remaining DCVs present in axons of Rab2-deficient animals are not strongly depleted of DCV-specific membrane proteins. Second, suppression of AP-3, which is required for sorting of DCV membrane proteins (including VMAT ^79^), by depletion of the critical AP-3 β3-adaptin/ruby subunit did not noticeably impact axonal DCV transport **(Fig. S5C, D)**. In light of the high enrichment scores for Snx3-Retromer and the Golgin245-TBC1D23-FAM91A1 tethering/fusion complex in Rab2- and VMAT-specific proximity proteomics (**Fig. 1G**), we also considered the possibility that Rab2 functions in the retrieval of VMAT (and by extension other DCV membrane proteins) from endosomes to the TGN. However, depletion of the core retromer subunit Vps35 did not affect axonal transport of DCVs **(Fig. S5C, D)**, suggesting that defective endosomal retrieval is not the cause of the axonal transport defects in Rab2 mutants.

In contrast to the Rab2-specific effect on VMAT sorting, larvae deficient in Syd, RUFY or dynein all displayed dramatic reductions in the cell body density of ILP2-positive DCVs as was seen in Rab2 mutants (**Fig. 6A, C**). This effect can likely be partially explained by trapping of DCVs in the axono-synaptic compartment by the loss of retrograde axonal transport, consistent with the observed increases in both proportion and absolute numbers of static axonal DCV in these animals (**Fig. 3A, F-H, 4C, D**). In comparison, loss of Arl8, which together with unc-104 is required for DCV cell body exit ^13, 16^, resulted in a large increase the cell body DCV density (**Fig. 6A, C**). However, axonal DCV trapping cannot fully explain the decrease in somatal DCV numbers in Rab2-, Syd-, RUFY- and dynein-disrupted motor neurons, as these genotypes also display severe reductions in total axonal DCV cargo content (**Fig. 4A, S6E, F**). Furthermore, despite the relative DCV cargo enrichment in the most distal synaptic boutons, overall synaptic DCV cargo content was also reduced in Rab2 ^16^ and Syd mutants **(Fig. S6G, H)** compared to wild type controls. Because our previous work indicates that DCV lumenal cargo loading is close to normal in *Rab2* nulls ^16^, this suggests that the total DCV number is substantially reduced in Rab2- and Syd-deficient neurons, and likely also in RUFY- and dynein-depleted neurons. Since Rab2, dynein and Syd are important for the function of the Golgi apparatus and/or ER-Golgi transport ^23, 76, 80, 81, 82^, this may reflect a role for these proteins in trafficking of lumenal DCV cargo at early stages of the secretory pathway in a process that is not related to DCV membrane protein sorting.

Overall, we find that Rab2-dependent sorting of DCV membrane proteins is not mediated by Syd, RUFY or dynein and is also unlikely to be responsible for the DCV axonal transport defects observed upon loss of Rab2. However, loss of Rab2, Syd or dynein led to similar reductions in neuronal DCV numbers, possibly due to a dysfunction of lumenal DCV cargo trafficking at earlier stations of the secretory pathway.

## Discussion

We have investigated the machinery responsible for axonal circulation of DCVs in Drosophila, using its dependence on the small GTPase Rab2 as a starting point. We find that axonal DCV motility is mediated by the coiled-coil dynein and kinesin-1 adaptor Syd/dJIP3/4 and the novel coiled-coil dynein adaptor RUFY, which interact with the Rab2 and Arl8 GTPases. Given the severity of both the Syd loss-of-function phenotype and of the RUFY depletion phenotype, which is likely an underestimation of a full RUFY-null phenotype, it seems likely that these proteins function as part of the same mechanism to activate retrograde movement rather than in parallel. Furthermore, the two adaptors bind to each other in an interaction that requires the Syd C-terminal WD40 domain, and truncation of this region eliminates most or all Syd activity. These observations suggest a model where dynein-dynactin and kinesin-1 are activated and recruited by Syd anchored to the DCV membrane by a combination of Rab2 (potentially assisted by an unknown Rab2 effector) and RUFY, which itself is recruited by Arl8 and its activator BORC, and possibly Rab2 (**Fig. 5J**). However, RUFY also has the long coiled-coil structure characteristic of dynein activating adaptors and there is evidence that mammalian RUFY1/3 interact with dynein directly ^40, 41^. The RUN domain and central coiled-coil region 2 that are required for the interaction of RUFY1/3 with dynein ^40, 41^ are conserved in fly RUFY **(Fig. S2F)**. We can therefore also not exclude alternative scenarios where RUFY acts as the main activating adaptor for dynein, while Syd helps anchoring RUFY to the vesicle membrane (and at the same time interacts with kinesin-1), or where the active transport complex contains two or more dynein dimers bound to a Syd:RUFY heterotetramer. The latter scenario would mirror recent findings showing that two dynein dimers form a complex with two BICDR1 activating adaptor dimers during transport ^83^. Further research is required to understand how Syd-RUFY dependent motility functions and whether RUFY proteins can activate dynein directly.

The involvement of Arl8 in retrograde transport of DCVs via RUFY, in addition to its previously known role in anterograde transport via unc-104/kinesin-3, mirrors recent findings elucidating the role of mammalian ARL8 in regulation of lysosomal motility ^39, 40, 41, 84^. ARL8 also recruits kinesin-1 through the SKIP/PLEKHM2 adaptor ^85, 86^, but the fly SKIP ortholog, prd1, lacks the Arl8-binding RUN domain and prd1 knockout larvae did not exhibit any obvious DCV transport defects **(Fig. S3E, F)**. Our observed Arl8 null phenotype ^16^ has features in common with both unc-104 and Syd/dynein/kinesin-1 mutants, in that it both exhibited a cell body exit defect and a strong increase in the proportion of static vesicles among those remaining in the axons (**Fig. 3**). In comparison, the few remaining axonal DCVs in unc-104^P3^^50^/unc-104^O3.1^ hypomorphs were mostly motile, but much slower when moving in the anterograde direction (**Fig. 3**), as reported previously ^15^. Meanwhile, depletion of kinesin-1 or dynein, apart from selective reductions in retrograde transport flux and movement speed **(Fig. S3A, B)**, produced large increases in the proportion of static axonal DCVs (**Fig. 3**). As also noted by others ^14^, this suggests that the main role of kinesin-3 is to move DCVs out of the soma and to enhance the anterograde axonal DCV velocity, while kinesin-1 and dynein are critical for maintaining bidirectional motility among the axonal DCVs, and their absence leads to axonal vesicle stalling. This difference likely relates in large part to distinct interactions of kinesin-1 and -3 with differentially distributed MT-binding proteins ^12, 87^. Moreover, the kinesin-1- and dynein-mediated axonal DCV motility appears to be partially maintained by JNK signalling. Surprisingly, the knockout phenotype for the BORC subunit Blos1 (**Fig. 5E, F**) was closer to that exhibited by Syd/RUFY/kinesin-1/dynein than was the Arl8 knockout phenotype. This difference may reflect graded phenotypic effects of different levels of remaining Arl8 activity, or that BORC only activates Arl8 in the axonal compartment, while a different Arl8 activator facilitates DCV sorting from the soma to the axon. However, it is possible that BORC could directly interact with RUFY or Syd.

The identified components of the DCV transport complex, Syd/JIP3/4, RUFY, Arl8, BORC and Rab2 all have an extensive record as regulators of lysosomal positioning and/or axonal transport ^18, 36, 39, 40, 62, 63, 64, 85, 88^, revealing a striking similarity between the motility mechanisms of DCVs and lysosomes. These results are not due to simple organelle misidentification or inherently fuzzy organelle identities, as shown by our previous work, and also could explain the relatively similar pattern of axonal DCV and spinster-positive lysosome motility (**Fig. 3A, 4E, S3A**)^16^. Despite a bias towards anterograde (DCVs) or retrograde (lysosomes) transport, both organelles display extensive bidirectional movement at very similar speeds and characterized by long runs ^16^(**Fig. 3, 4E, S3A-B)** indicative of axonal circulation.

Interestingly, in our PDB experiments we observe strong enrichment for lysosomal/autophagic and Golgi-related proteins in the Rab2^Q65L^-specific dataset, but less so for endoplasmic reticulum (ER) proteins (**Fig. 1C**) as might have been expected from early studies that indicated that Rab2 is responsible for retrograde transport from Golgi-ER intermediates to the ER ^89, 90^. These findings are, however, consistent with newer work that places Rab2 as a critical factor in lysosome function and biogenesis ^19, 20^, autophagy initiation and clearance ^21, 22^ and tethering at the Golgi apparatus ^30, 33, 34^. Apart from most previously known Rab2 effectors, the Rab2^Q65L^-PDB data set also contained several proteins detected by Drosophila Rab2 affinity-purification MS or human Rab2A MitoID but not validated as effectors ^30, 43^, including RUFY and gartenzwerg/GBF1. GBF1 is a Golgi-localized Arf1 activator promoting COPI vesicle budding ^91^ and could thus provide a link between Rab2 and retrograde Golgi-to-ER trafficking. Curiously, both Rab2^Q65L^- and VMAT-PDB datasets showed high enrichment for the transmembrane autophagy factor Atg9 (**Fig. 1G**), which may reflect the newly uncovered close ties between the synaptic exo-endocytic cycle and synaptic autophagy ^92^ and/or that Atg9 follows the same Rab2-dependent sorting pathway in the soma as VMAT (see below).

In line with extensive *C. elegans* work demonstrating a role for Rab2 and its effectors in DCV biogenesis ^24, 25, 27, 28^ we also observed that the loss of Rab2 causes a severe DCV membrane protein sorting defect in flies. In Rab2 null fly neurons VMAT accumulated in the soma in densely packed ectopic aggregates of small vesicles closely resembling the TGN-proximate SV-component-containing vesicle clusters previously observed in Rab2 mutants ^23^. This suggests that DCV and SV membrane protein cargo is sorted into small transport vesicles during DCV/SV-transport organelle biogenesis, which fail to fuse to their target compartment in the absence of Rab2. Interestingly, the dense, droplet-like appearance of the ectopic VMAT vesicle aggregates (**Fig. 6A**) suggests that they undergo liquid-liquid phase separation (LLPS) from the surrounding cytoplasm; this could result from a high abundance of LLPS-prone ^93^ synaptic proteins in the vesicles. The vesicle target compartment may be immature DCVs or SV-carrier organelles or a TGN sub-compartment. In fact, substantial enrichment for both a retromer recycling complex variant and Rab2-binding TGN tethering factors (Golgin245 and Golgin-104) in the Rab2^Q65L^- and VMAT-PDB datasets suggests that these transport vesicles may originate in endosomes and carry their cargo to the TGN. This correlates with recent work showing that DCV membrane cargo (including VMAT) initially exits the Golgi apparatus separately from lumenal cargo ^94^ and that endosomal recycling plays a critical role in DCV-biogenesis ^29, 95^. While *C. elegans* reports indicate that some lumenal DCV cargo is degraded in late endosomes/lysosomes in Rab2 mutants ^24^, membrane cargo lost from DCVs was shown to accumulate in neuronal somata in an unidentified compartment ^25^, indicating that the Rab2 mechanism of action and loss-of-function phenotype may fundamentally be the same in flies and nematodes. While future experiments will need to address if Rab2 indeed mediates the retrieval of DCV/SV proteins from endosomes to the TGN, such a role would be in keeping with the high abundance of Rab2 at the Golgi and relatively close phylogenetic kinship of Rab2 and endosomal recycling associated Rab14 and Rab4 ^96^.

In summary, we find that coupling of kinesin-1 and dynein to DCVs for axonal transport happens via Syd/dJIP3/4 and RUFY that are controlled by Rab2 and Arl8/BORC, but that these adaptors are not required for Rab2-dependent in DCV membrane cargo sorting.

## Materials and Methods

### Fly husbandry and genetics

Fly stocks were maintained on Nutri-Fly Bloomington Formulation medium (Genesee Scientific, San Diego, California) at 25°C. For experiments involving live imaging of axonal transport or immunostaining of larval tissues all larvae were reared on apple juice (AJ) plates (27 g/L agar, 12 g/L sucrose, 1.875 g/L Nipagin [methyl 4-hydroxybenzoate]) supplemented with yeast paste. After an overnight lay, plates were incubated for ∼48h at 25°C to allow all eggs to hatch. L1-2 larvae of the desired genotype (determined by presence or absence of appropriate fluorescence) were then transferred to fresh yeasted AJ plates for another ∼48h (∼96h after end of egg laying) before being dissected. For RNAi experiments, flies were reared at 29°C and picked for dissection ∼72h after end of egg laying to account for the faster pace of development at higher temperature. To increase knockdown efficiency, UAS-Dicer-2 was co-overexpressed with the RNAi transgenes, except when short hairpin RNAs were used. In matching RNAi controls, UAS-Dicer-2 was overexpressed alone under control of the appropriate driver. Fly lines are listed in Supplementary Table 1.

### Drosophila Phi31C transformation

Fly embryo DNA injections and selection of Phi31C transformants was performed by BestGene Inc.. The UAS-HA-TurboID-VMAT and UAS-HA-TurboID transgenes were inserted into the M{3xP3-RFP.attP}ZH-86Fb attP site on the third chromosome using Phi31C transformation to ensure comparable levels of expression.

### Molecular biology

All constructs (listed in Supplementary Table 2) were generated by GenScript Biotech (Netherlands) BV, except pCMV5-FLAG-LRRK2[G2019S], which was obtained from MRC PPU Reagents and Services (University of Dundee, Scotland). All constructs for protein expression in mammalian cell culture generated for this study were made in the pCDNA3.1(+) vector and were based on the following protein isoforms: Syd isoform A (UniProt Q9GQF1), RUFY/CG31064 isoform G (UniProt A0A0B4LHR8), dNischarin/CG11807 (UniProt Q7K490), DLIC isoform A (UniProt Q9VZ20), and Klc isoform A (UniProt P46824), Rab2 (UniProt O18333), Arl8 (UniProt Q9VHV5). Constructs for Drosophila Phi31C transformation were generated in the pUASTattB vector.

### *In vivo* proximity biotinylation and purification of biotinylated proteins

Flies expressing TurboID-Rab2S20N, TurboID-Rab2Q65L, TurboID-VMAT together with ILP2-GFP, or free TurboID together with ILP2-GFP under control of the pan-neuronal elav-Gal4 driver were reared at 25°C on Nutri-Fly Bloomington Formulation medium supplemented with 100 μM Biotin. (ILP2-GFP was co-overexpressed with both TurboID-VMAT and free TurboID transgenes in order to stimulate DCV production.) Adult flies were collected 0-3 days after eclosion, flash-frozen in liquid nitrogen and stored at −80°C. For purification of biotinylated proteins, 0.5-1 mL of frozen flies were transferred to pre-cooled dounce homogenizers on ice, quickly dounced 5x, then dounced 15x in 3 mL RIPA lysis buffer (50 mM Tris pH 7.5, 150 mM NaCl, 0.5% Sodiumdeoxycholate, 1.0% NP-40, 0.1% SDS, 1 mM DTT, cOmplete Protease Inhibitor Cocktail [1 tablet/25 ml](Roche, Ref: 11836145001), 1 mM PMSF). Lysates were then incubated for 30 min on ice and again dounced 15x. Hereafter the lysates were centrifuged 3 times for 15 min at 50000 g and passed through 40 µm Cell Strainers (Fisherbrand, Cat. no. 22363547) to remove insoluble debris. Protein concentrations in the resulting lysate supernatants were measured using the BCA assay (Pierce™ BCA Protein Assay Kit, Thermo Fisher Scientific, Ref: 23225) and adjusted to 2.0 mg/ml. 3.4 ml of each lysate was pre-cleared for 1 h at 4°C under rotation with 300 µl Sepharose 4B beads (Sigma-aldrich 4B200-100ML, Lot# MKCJ6278), previously equilibrated in RIPA buffer. Sepharose 4B beads were then removed by gentle centrifugation and 1.5 ml of each supernatant was incubated over night at 4°C under rotation with 50 μl Dynabeads MyOne StreptavidinT1 (Thermo Fisher Scientific, Ref 65601, Lot: 00804134). Dynabeads were then magnetically concentrated and washed 1x with 1 mL high-SDS RIPA buffer (50 mM Tris ph7.5, 150 mM NaCl, 0.5% Sodiumdeoxycholate, 1.0% NP-40, 0.4% SDS) at room temperature (RT), then transferred to clean sample tubes in a new wash of high-SDS RIPA. They were then washed 2x with SDS wash buffer (50 mM Tris ph7.5, 2.0% SDS), 1x with high-SDS RIPA buffer, 1x with normal RIPA buffer, and finally 2x with PBS and again transferred to clean tubes in a third wash of PBS.

### LC-MS analysis of biotinylated proteins

Washed beads were eluted by for 30 minutes incubation at 37°C in elution buffer 1 (2 M Urea, 50 mM Tris-HCl pH 7.5, 2 mM DTT, 20 μg/ml trypsin) followed by a second elution step for 5 minutes in elution buffer 2 (2 M Urea, 50 mM Tris-HCl pH 7.5, 10 mM Chloroacetamide). Both eluates were combined at further incubated at room temperature overnight. Tryptic peptide mixtures were acidified to 1% TFA and loaded onto Evotips (Evosep). Peptides were separated on a Pepsep 15 cm, 150 μM ID column packed with C18 beads (1.5 μm) using an Evosep ONE HPLC system applying the default 30-SPD (30 samples per day) method. The column temperature was maintained at 50 °C. Peptides were injected via a CaptiveSpray source and 20 μm emitter into a timsTOF pro2 mass spectrometer (Bruker) operated in PASEF mode. MS data was collected over a range of 100-1700 m/z with a TIMS mobility range of 0.6-1.6 1/K0. TIMS ramp and accumulation times were set to 100 milliseconds, with 10 PASEF ramps recorded for a total cycle time of 1.17 seconds. The MS/MS target intensity and intensity threshold were set to 20,000 and 2,500, respectively. An exclusion list of 0.4 min was activated for precursors within 0.015 m/z and 0.015 V cm^−2^ width.

### MS data analysis

Raw mass spectrometry data were analyzed using MaxQuant (version 1.6.15.0). Peak lists were searched against the human Uniprot FASTA database, combined with 262 common contaminants, using the integrated Andromeda search engine. A false discovery rate of 1% was set for both peptides (minimum length of 7 amino acids) and proteins. Carbamidomethylation of cysteine was specified as a fixed modification, while oxidation of methionine, acetylation at the protein N-terminus, acetylation of lysine, and phosphorylation of serine, threonine, and tyrosine were considered variable modifications. Additionally, “Match between runs” (MBR) was enabled with a Match time window of 0.7 minutes and a Match ion mobility window of 0.05 minutes.

All statistical analysis was conducted using in-house developed Python code ^97^. LFQ intensity values were log2-transformed, and features with less than 70% of valid values in at least one group were eliminated. Remaining missing values were replaced by mixed imputation, where kNN and MinProb (width=0.3 and shift=1.8) methods are used for values missing at random (MAR) and values missing not at random (MNAR), respectively ^98^. MAR is defined when minimum 60% of the samples within a given group have an existing value. Differentially expressed features were identified by unpaired Student’s *t*-tests, followed by Benjamini-Hochberg correction for multiple hypothesis testing with a False Discovery Rate (FDR) threshold of 0.05 and a fold-change of 2.

### HEK cell transfection and co-immunoprecipitation

HEK293 (ATCC, #CRL-1573) cells were maintained in DMEM w. HEPES and NaHCO3 (University of Copenhagen, Substrat og SterilCentralen, Ref #: 12) supplemented with 10% Standard Fetal Bovine Serum (GIBCO, Ref: 10270-106), 200 U/ml penicillin and 50 μg/mg streptomycin.

For all Co-IP experiments, T75 cell culture flasks with ∼7*10^6^ cells per flask (corresponding to a cell density of ∼10^5^ cells/cm^2^) were transfected with a total of 1-7µg of DNA using 3µl Lipofectamine 2000 (Invitrogen, Ref: 11668-019) per 1µg of DNA. Cells were grown for ∼48 h after transfection to allow for recombinant protein expression. In the experiment to test for dependence of Rab2:Syd complex formation on LRRK phosphorylation, cells were also exposed to 2 µM MLi-2 in the medium for 2h immediately before being harvested.

At the end of the expression period cells were washed twice in ice-cold PBS and harvested in 1.5 mL ice-cold co-IP lysis buffer (20 mM HEPES pH 7.4, 130 mM NaCl, 2 mM MgCl2, 0.1% w/v Saponin, cOmplete Protease Inhibitor Cocktail [1.5 tablet/50 ml](Roche, Ref: 11836145001)) using a cell scraper. The co-IP lysis buffer used to handle samples with active GTPases (Rab2(wt) and Rab2[Q65L], but not Rab2[S20N]) was supplemented with 60 μM GppNHp, to lock the GTPases in their active conformation. In some experiments, the lysis buffer was also supplemented with 1:285 Phosophatase Inhibitor Cocktail 2 (Sigma-Aldrich, P5726), but this practice was discontinued after it was established that LRRK activity is not required for Rab2 binding to Syd. Cell suspensions were lysed by being forced through a 25 G needle 6 times. Resulting lysates were incubated on ice for 30 min and insoluble debris was removed by a pair of consecutive 12k g centrifugation steps of 10 and 5 min respectively. Supernatant protein concentrations were measured using the BCA assay (Pierce™ BCA Protein Assay Kit, Thermo Fisher Scientific, Ref: 23225) and adjusted to 1.0 mg/ml. 1.2 mL of each lysate supernatant was incubated with 22 μl Pierce™ Anti-HA Magnetic Beads (Thermo Fisher Scientific, Ref: 88836) or Pierce™ Anti-c-Myc Magnetic Beads (Thermo Fisher Scientific, Ref: 88842) for 2 h at 4°C under rotation. Magnetic beads were then washed 4x with standard co-IP lysis buffer with 0.1% saponin (anti-HA beads), or 2x with co-IP lysis buffer with 0.25% saponin followed by 2x co-IP lysis buffer with 0.1% saponin (anti-c-Myc beads). During each wash beads were resuspended by pipetting and then separated from the supernatant using a DynaMag™-2 magnetic rack (Thermo Fisher Scientific, Ref: 12321D). Beads were also transferred to new Eppendorf tubes in the second wash. Bound proteins were eluted form anti-HA beads by either by incubation in 40 µl 100 mM NaOH for 10 min at RT or with 40 µl 2.5 mg/ml HA peptide (Thermo Scientific Cat #: 26184) for 20 min at 37°C, and from anti-c-Myc beads by heating in SDS-PAGE loading buffer diluted 1:2 in wash buffer. Eluates and lysate aliquots were mixed with standard Laemmli SDS-PAGE loading buffer and heated to 99.9°C for 10 min in preparation for SDS-PAGE. All co-IP experiments except the experiment in Fig. S2A were replicated at least once.

### Western Blotting

For SDS-PAGE, protein samples were run on AnyKD Mini-PROTEAN® TGX™ Precast Protein Gels (BioRad, Ref: 4569033) clamped to 100V. Separated proteins were transferred to PVDF membranes using the Trans-Blot Turbo Transfer System (BioRad, Ref: 1704150) running a custom transfer program (1.3A, 25V, 20 min, current clamped). Membranes were blocked overnight at 4°C, then probed with primary antibody in blocking buffer (PBS ph7.4, 0.05% Tween-20 v/v, 5% v/w Skim Milk Powder [Sigma-Aldrich, Ref: 70166]) for 1 h at RT. After 3x 10 min washes in wash buffer (PBS ph7.4, 0.05% Tween-20 v/v), membranes were then incubated with HRP-conjugated secondary antibody in blocking buffer for 1 h at RT. Membranes were then again washed for 3x 10 min in wash buffer, and then 1x 5 min in PBS, before being deposited in deionized water. Chemiluminescent signals were developed by incubating the membranes for 10 min in SuperSignal ELISA Femto Maximum Sensitivity Substrate (Thermo Fisher Scientific, Ref: 37075). Membrane imaging was performed on an Amersham ImageQuant 800 luminescence imager using the Signal-to-Noise Optimization Watch (SNOW) capture mode. Antibodies used for western blotting are listed in Supplementary Table 3.

For co-IP experiments, between 40% and 15% of the total eluate volume and a lysate volume corresponding to ∼1% of the binding reaction was loaded on each gel. In experiments shown in Fig. 2B, D, E, F, G, I, K and Fig. S2B, D, E membranes with lysate and eluate samples were developed separately as the yield co-IP yields were relatively low. In Fig. 2A and 5A-B lysate and eluate samples were developed together on the same membrane.

### Immunohistochemistry

For staining of neuronal somata in the larval VNC and of larval peripheral nerves, third instar larvae were dissected in phosphate-buffered saline (PBS) containing 137 mM NaCl, 2.7 mM KCl, 1.5 mM KH2PO4, 6.5 mM NaH2PO4, pH 7.4. Larval CNS (still attached to a piece anterior cuticle for easier handling) were extracted and briefly stored in Schneider’s insect cell medium (Life Technologies, A820) supplemented with 5% heat-inactivated fetal bovine serum (FBS) at RT prior to fixation. For staining of larval NMJ synapses, third instar larvae were pinned down using 0.1 mm Minutien pins (Fine Science Tools) on ∼1.3 cm Ø Sylgard slabs and fillet-dissected in modified HL3 solution (70 mM NaCl, 5 mM KCl, 10 mM NaHCO3, 20 mM MgCl2, 5 mM trehalose, 115 mM sucrose, 0.5 mM EGTA, 5 mM HEPES, pH 7.2). After dissection, isolated CNSs and larval fillets were fixed in 3.7% formaldehyde in PBS at RT for 50 min. Specimens were then washed 6x10 min in PBX (PBS with 0.3% Triton X-100 [Sigma-Aldrich, Ref: T8787]) and blocked for 2 h at RT in blocking buffer (PBX with 10% goat serum [Sigma-Aldrich, Ref: G9023]). After this they were incubated for 4°C 72h in primary antibodies in antibody incubation buffer (PBX with 5% goat serum). This was followed by 6x10 min washes in PBX at RT, and an overnight incubation at 4°C with secondary antibodies in antibody incubation buffer. Finally, specimens were subjected to another set of 6x10 min PBX washes, followed by 2x5 min washes in PBS, and mounted in ProLong® Gold antifade reagent (Life Technologies, P36934). All incubations were done under gentle agitation.

For standard confocal microscopy, secondary or primary antibodies were labelled with Alexa 488, Alexa 647, or Rhodamine Red-X dyes. For STED microscopy, antibodies were labelled with Abberior STAR RED and Abberior STAR ORANGE. Antibodies used for immunohistochemistry are listed in Supplementary Table 3.

### Conjugation of fluorophore to nanobody

In prepartion for STED imaging, alpaca anti-GFP V_H_H single domain antibody/nanobody (Chromotech, gt-250, Lot: 71017001U) was conjugated to abberior STAR ORANGE NHS ester (abberior, Ref: STORANGE-0002-1MG, Lot: 10319RK-1), then isolated through Zeba Spin Desalting Columns, 7K MWCO (Thermo Fisher Scientific, Ref: 89883). This was completed by first washing the column three times with 300 μl of 100 mM NaHCO3 in PBS. Following the washes, 100 μg of nanobody in 200 μL of PBS was added and spun at 1500 g for 2 minutes. The flow-through was collected and combined with a 5-fold molar excess of NHS ester fluorophore. This solution was incubated in the dark at RT, shaking, for 2.5 hours. A new spin column was washed 3 times with 0.02% NaN3 in PBS. The antibody sample with the dye was then added to the column and centrifuged for 2 minutes at 1500 g. Protein and label concentrations were measured on an Eppendorf BioPhotometer Plus spectrophotometer. The resulting STAR ORANGE-conjugated nanobody had a labelling rate of >0.4 fluorophores/molecule.

### Confocal microscopy

Confocal microscopy was carried out at the Core Facility for Integrated Microscopy (Department of Biomedical Sciences, University of Copenhagen) using a LSM 700 confocal microscope (Carl Zeiss Microscopy GmbH, Jena, Germany) and the following objectives: Plan-APOCHROMAT 63x/1.4 Oil DIC (for IHC samples), W APOCHROMAT 40x/1.0 DIC VIS-IR #421462-9900 Water dipping (for live imaging). Live confocal microscopy was performed as previously described ^16^. In brief, fillet-dissected third instar larvae pinned down in Sylgard dishes were imaged directly in modified HL3 using the W APOCHROMAT 40x/1.0 DIC VIS-IR #421462-9900 water dipping objective. For assessment of axonal transport of DCVs and lysosomes A7 peripheral nerves were imaged in a 128 µm long segment 0.5-1.0 mm from the nerve egress from the VNC. After recording a prebleach image, the 60 µm flanking sections were photobleached using a 405 nm laser and subsequent time-lapse imaging performed for 499 frames (corresponding to ∼106 s for DCVs, and ∼212 s for lysosomes due to higher averaging). In a few cases, imaging was performed in younger larvae, as specified in the Figures.

### STED microscopy

STED microscopy was performed at the Core Facility for Integrated Microscopy (Department of Biomedical Sciences, University of Copenhagen) using an Abberior STEDYCON system mounted on a Zeiss AxioImager Z1 widefield microscope with an alpha Plan-Apochromat 100x/1.46 Oil DIC M27 objective. Pixel sizes of 20 or 15 nm were used during imaging.

### Image analysis

Confocal and STED images were analyzed using the Fiji/ImageJ package ^99^. Prebleach nerve fluorescence was quantified as the total integrated density of the A7 mid-nerve segment subsequently undergoing time-lapse imaging of vesicle transport.

### Axonal vesicle transport

A custom algorithm was used to generate kymographs from time-lapse recordings. For presentation in Figures, kymographs were digitally inverted. To produce directional distributions, the kymographs were rotated 90° counterclockwise and subjected to the Directionality plugin in Fiji, selecting Fourier spectrum analysis. For statistical analysis, the peak relative frequency of directional distribution angles was determined within the following intervals: -87.98° to -45.51° (retrograde peak, *Pret*), 87.98° to 45.51° (anterograde peak, *Pant*), -13.14° to 13.14° (static peak, *Pstat*). The static relative peak amplitude, *Pstat(rel)*, was calculated using the expression:

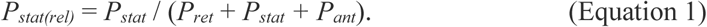

Directional distribition angles were converted to transport velocities using the expression:

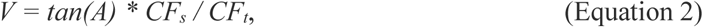

where *V* is transport velocity in µm/seconds*, A* is the angle, and *CFs* and *CFt* are conversion factors for the space and time axes of the kymograph, respectively. For DCV experiments, *CFs* was 0.1 µm/pixel and *CFt* was 0.2125 seconds/pixel. For all other genotypes than *Arl8/Df*, a directional distribution was generated from each kymograph. The paucity of DCVs in Arl8 mutant nerves made this approach less attractive. Instead, one single directional distribution was produced from a maximum intensity projection of 20 stacked kymographs (reanalysis of data from nine Arl8 larvae, published previously ^16^).

Anterograde and retrograde DCV flux in axons during the initial 30 seconds of the recording session was quantified by counting unbleached DCVs that entered the bleached areas from the left and right side, respectively, and travelled at least 1.8 µm further along the axon. To facilitate DCV tracking, images were Gaussian blurred and DCV centers marked with a black dot using the Find Maxima Plugin in Fiji to locate fluorescence peak intensities. DCV flux in *Syd* and in *Rab2, Syd* double mutants was quantified using kymographs. Static (i.e., not moving in 30 seconds) vesicles were counted by generating “kymostacks” of the central unbleached region, as described earlier ^16^.

To quantify DCV speed, a segmented line was fitted to the trajectory of individual vesicles in the kymographs, with each line segment representing an anterograde run, a retrograde run, or a pause if DCV speed was lower than 0.015 μm/s ^16^. Pauses were excluded before calculating the average run speed per vesicle.

As for DCVs, directional distributions were produced to quantify axonal transport of lysosomal organelles labelled with Spinster-Venus in motor axons of *Syd* larvae and matching controls. To relate these results to Rab2’s role in lysosomal transport, directional distributions were also generated of Spinster-GFP-positive lysosome transport in *Rab2* larvae and their controls (reanalysis of data published earlier ^16^). Equation 2 was used to convert the directional angles to lysosome transport velocities, with *CFs* equal to 0.1 µm/pixel and *CFt* equal to 0.4251 seconds/pixel.

### ILP2-GFP intensity in nerve terminals

The overall ILP2-GFP immunosignal was measured as the integrated density on background-subtracted sum projections of z-stacks traversing the entire nerve terminal. To quantify the intensity gradient of the ILP2-GFP signal in distal boutons of individual end branches, images were Gaussian blurred before drawing segmented lines through the boutons in each branch, starting with the end bouton. The amplitudes of the first five peaks in the corresponding plot profiles (where the x-axis represents distance along the line and the y-axis pixel intensity) were taken as the intensity of the distalmost five boutons. Amplitudes were standardized by dividing by their mean. Slopes of linear regression lines in plots of the five peak amplitudes against their x-positions were calculated using Excel software.

### HA-VMAT aggregate frequency

To determine the abundance of HA-VMAT aggregates, motor neuron somata found to contain rounded, sharply demarcated “drop-like” aggregates of HA-VMAT-positive vesicles were counted using confocal micrographs of the dorsal surface of the larval ventral nerve cord.

### STED micrograph analysis

To quantify the extravesicular percentage of the HA-VMAT immunosignal in axons, images were background-subtracted and the ILP2-GFP channel thresholded to include the 10% most intense pixels. This threshold was converted to a region of interest (ROI) set that was restored on the HA-VMAT channel. The HA-VMAT intensity (integrated density) of the ILP2-GFP-associated ROI set was divided by the total HA-VMAT intensity obtained after also thresholding the HA-VMAT channel to 10%. The resulting intravesicular HA-VMAT signal percentage was subtracted from 100 to obtain the extravesicular percentage. The mean vesicular HA-VMAT intensity was measured in individual ROIs from the ILP2-GFP-associated ROI set.

To estimate the density of ILP2-GFP-containing vesicles in cell bodies, the intensity of the total ILP2-GFP immunosignal was divided by the intensity of individual vesicles, calculated as an average of generally 5-10 isolated vesicles. The resulting vesicle count was finally divided by the cell area. Calculation of Pearson’s correlation coefficient to quantify ILP2-GFP vs. HA-VMAT, and ILP2-GFP vs. Sytα-mCherry colocalization was restricted to cell body ROIs.

The size of ILP2-GFP-positive DCVs and small HA-TurboID-VMAT-positive vesicles in presynaptic boutons was quantified using background-subtracted images of presynaptic type Ib and II boutons. To obtain DCV size, the ILP2-GFP channel was subjected to Gaussian blur (σ = 20 nm) and thresholded, before using Fiji’s particle analysis feature to obtain the area of individual particles, from which the diameter was calculated by assuming a circular shape. The analysis was restricted to particles with a circularity above 0.80. To obtain the size of small VMAT-positive vesicles, representative isolated vesicles were selected in the VMAT channel in areas devoid of ILP2-GFP signal. The vesicle diameter was measured as the full width at half maximum (FWHM) on a Gaussian fit of the intensity profile.

To determine the size of the vesicles constituting the HA-VMAT aggregates in cell bodies of *Rab2* larvae, 50 representative, isolated vesicles on STED images of 10 cell bodies in four larvae, located either in the rim of densely packed aggregates or in aggregates with moderate vesicle density, were selected for measuring the FWHM of Gaussian fitted intensity profiles.

### Candidate protein analysis

Documentation concerning the subcellular localization and functional annotation of biotinylated candidate proteins was obtained from open source bioinformatics databases (https://flybase.org/, https://www.uniprot.org/), updated through searches of the recent literature. The information summarized in Supplementary Table 2 is based on studies of both the Drosophila proteins and their closest human orthologs.

Cutoff criteria for significantly biotinylated candidate proteins in volcano plots were an adjusted *P*-value less than 0.05, and a fold change exceeding 2. When calculating the rank correlation between Rab2-related and VMAT-related candidate proteins the highest rank was used in cases where the same protein was detected more than once,

### AlphaFold modelling

Protein structure modelling of the Drosophila Rab2:Syd (2:2) complex was performed with ColabFold ^100^ version 1.5.5 (AlphaFold2 ^101^ using MMSeq2) (https://colab.research.google.com/github/sokrypton/ColabFold/blob/main/AlphaFold2.ipynb) using the AlphaFold2_Multimer setting and relaxation. The crystal structure of active Drosophila Rab2 bound to GppNHp (pdb: 4rke, ^57^) was used as a template.

Modelling of the RUFY dimer was performed with AlphaFold 3 ^102^ (https://alphafoldserver.com/).

### Molecular dynamics simulations

The AlphaFold structure was used for the Syd dimers. A Rab2 structure with bound Mg and GNP (PDB 4RKE) ^57^ (without N-terminal residues GAMG and bound water) was aligned with the Rab2 from the AlphaFold structure, which was then removed, except for C-terminus, which was not resolved in the crystal structure. GNP was replaced with GTP. The construct was truncated to reduce computational time. The Syd dimers were truncated to residues L450 – G526 and only one truncated Rab2 (M1-G188) was maintained in the complex. Mutant R468A, double mutant L465/R468A and truncated construct ΔHelix-2 (Syd truncated after V504) were generated in PyMOL (The PyMOL Molecular Graphics System, Version 3.0 Schrödinger, LLC). The complex was solvated in a 20x9x9 nm box with TIP3P water and 100 mM NaCl. The simulations were run with GROMACS 2021.4 ^103^ and the CHARMM36m force field ^104^. The system was minimized, then equilibrated in 10 ps with constant number of particles, volume and temperature (NVT) and 100 ps with constant number of particles, pressure and temperature (NPT). Both equilibrations were run with 2 fs step size, v-rescale temperature coupling (time constant 0.1 ps) to keep temperature at 300 K, and the NPT equilibration was run with Berendsen isotropic pressure coupling (time constant 2 ps) to keep pressure at 1 bar. The proteins were restrained with position restraints during equilibration. The Particle mesh Ewald (PME) algorithm ^105^ was used for long range and LINCS algorithm to constrain hydrogens ^106^. The restraints were relieved and a simulation was run for 2 ns with isotropic Parinello-Rahmen pressure coupling. The Rab2 (including Mg and GTP) was then pulled away from the Syd dimer using GROMACS built-in umbrella biasing potential with a rate of 0.01 nm/ps. Frames were taken from this pull simulation with Syd-Rab2 centre-of-mass distances up to 7.5 nm in steps of 0.15 nm. Each of these frames were used for 10 ns simulations, with COM distance fixed using umbrella biasing force of 1000 kJ/mol/nm^2^. Potential of mean force (free energy of binding) was calculated from the umbrella simulations using the weighted histogram average method (WHAM) ^107^. The whole process (including solvation and minimization) was repeated 10 times for each construct, and the mean values and SEM were calculated.

### Sequence handling and alignment

Protein sequence alignment was performed in BioEdit (Hall T.A. 1999, BioEdit: a user-friendly biological sequence alignment editor and analysis program for Windows 95/98/NT) using the ClustalW multiple alignment function.

### Helical propensity estimation

Alpha helical propensity of the different regions of the Syd RH2 domain were evaluated using the NetSurfP - 3.0 online tool from the Technical University of Denmark ^108^. https://services.healthtech.dtu.dk/services/NetSurfP-3.0/

### Statistical analysis

Data visualization in graphs was performed with Excel software (Microsoft), which was also used for *t*-tests. Analysis of variance (ANOVA) and Dunnett’s test were executed with JMP sofware (JMP Statistical Discovery). Before analysis, datasets were assessed for homogeneity of variances with a test battery including Bartlett’s test, and the residuals were checked for normal distribution with Shapiro-Wilk’s test. Data failing to conform to normality or homoscedasticity were logarithmically transformed, or the non-parametric Steel with control test was applied, as appropriate. The rank correlation between Rab2-related and VMAT-related candidate proteins was calculated using Excel. *P* values less than 0.05 were considered significant, indicated with red text color in the Figures. All perfomed Dunnett’s tests and *t*-tests were two-sided. The experimental unit was larva (i.e., one larva was represented with one value, usually the average of repeated measurements). In tests with a non-significant outcome when comparing group means, the sample size was at least four third instar larvae per group. Details of individual statistical tests, including sample size, are provided in the Source File.

## Acknowledgements

We thank the staff at the Core Facility for Integrated Microscopy and Anders Bohl Pedersen for technical assistance. We thank Dr. Alyssa Johnson, Dr. Graeme Davis, Dr. Peter Robin Hiesinger, Dr. William M. Saxton and Dr. David Krantz for sharing fly lines, and Matthew Domenic Lycas for his help with nanobody fluorophore conjugation. Andrew Carter and Sami Chaaban are thanked for useful suggestions and for sharing unpublished information on JIP3 autoinhibition. The MD simulations were performed using the Danish National Life Science Supercomputing Center, Computerome. We thank the staff at the Proteomics Research Infrastructure for expert technical assistance. Mass spectrometry-based proteomics analyses were performed by the Proteomics Research Infrastructure (PRI) at the University of Copenhagen (UCPH), supported by the Novo Nordisk Foundation (NNF) (grant agreement number NNF19SA0059305). This work was supported by Aage og Johanne Louis-Hansens Fond, The Lundbeck Foundation (grants RF347-2020-2339 and R359-2020-2301), NNF (grant agrrement number 70287), and the ERASMUS programme.

## Data availability

The mass spectrometry proteomics data have been deposited to the ProteomeXchange Consortium (http://proteomecentral.proteomexchange.org) through the PRIDE partner repository. The data set identifiers are PXD063196 (https://www.ebi.ac.uk/pride/archive/projects/PXD063196), and PXD063200 (https://www.ebi.ac.uk/pride/archive/projects/PXD063200).

## Supplementary Material

Supplementary Figures S1-S7 with legends

**Figure S1.**
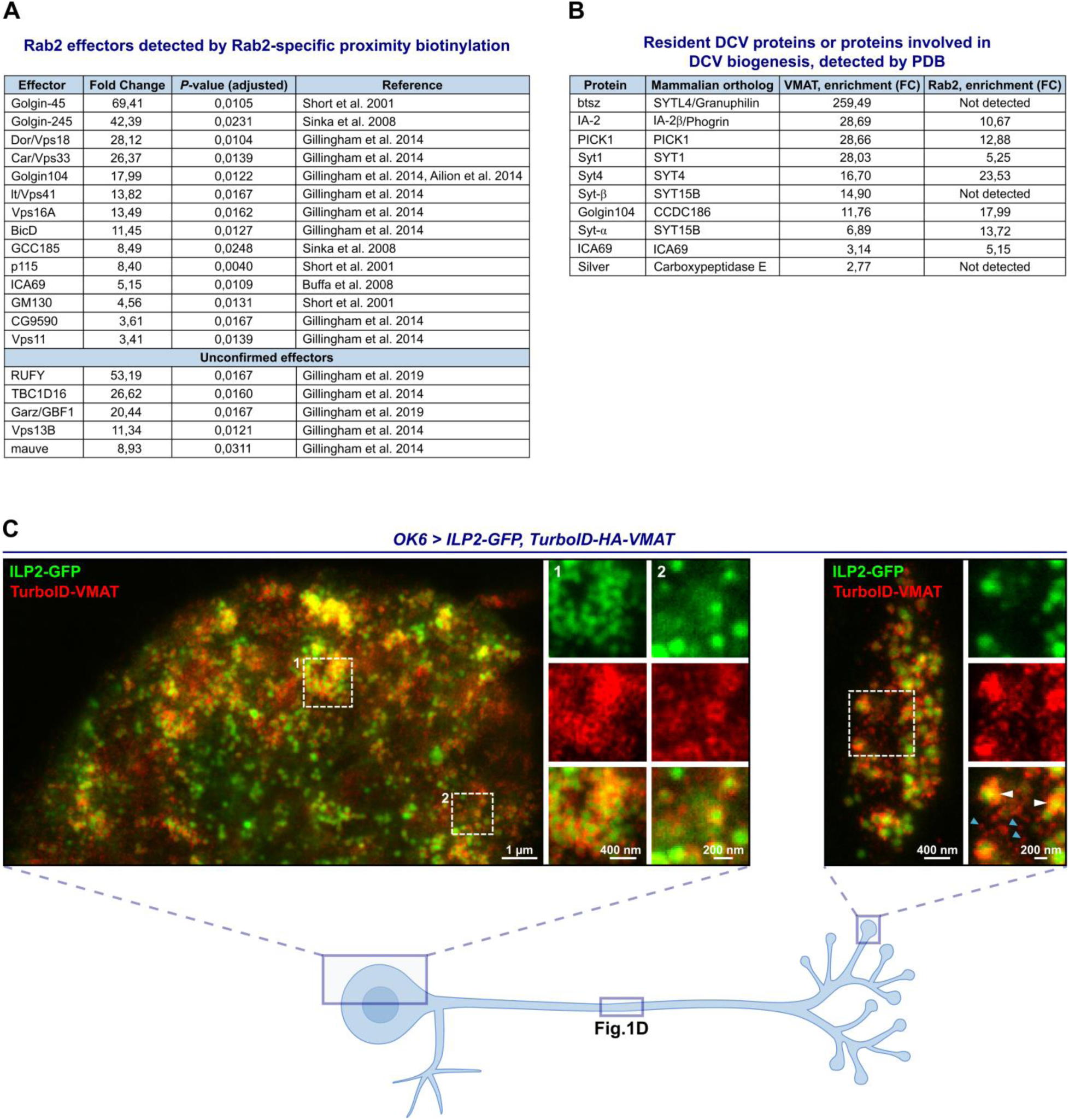
**A**, Previously identified Rab2 effectors (both Drosophila proteins and mammalian orthologs) found in our screen to be significantly enriched in TurboID-Rab2^Q65L^ samples relative to TurboID-Rab2^S20N^. *First 14 entries*, Confirmed effectors (also labeled in Fig. 1B). All subunits of the HOPS complex have been counted as effectors. *Last 5 entries*, Potential effectors detected by affinity proteomics in Drosophila S2 cells ^30^ and MitoID relocalization proximity protemics in HEK cells ^43^ but not confirmed using other methods. The list of unconfirmed effectors is not exhaustive. **B**, Proteins known to reside on DCVs or to be involved in DCV biogenesis that were significantly enriched in TurboID-VMAT samples relative to free TurboID. **C**, Representative STED images showing the distribution of ILP2-GFP and TurboID-HA-VMAT in motor neuron cell body located in the dorsomedial aspect ventral nerve cord (*left*) and in peripheral synaptic bouton (*right*) in a third instar larva. *White arrowheads*, TurboID-VMAT associated with ILP-GFP-positive DCVs. *Blue arrowheads*, small TurboID-VMAT positive vesicles not associated with ILP-GFP. The distribution of ILP2-GFP and TurboID-HA-VMAT in the mid-axon region in the same type of preparation is shown in Fig. 1D. Scale bars (left to right): 1µm, 400 nm (inset 1), 200 nm (inset 2), 400 nm, 200 nm (inset).

**Figure S2.**
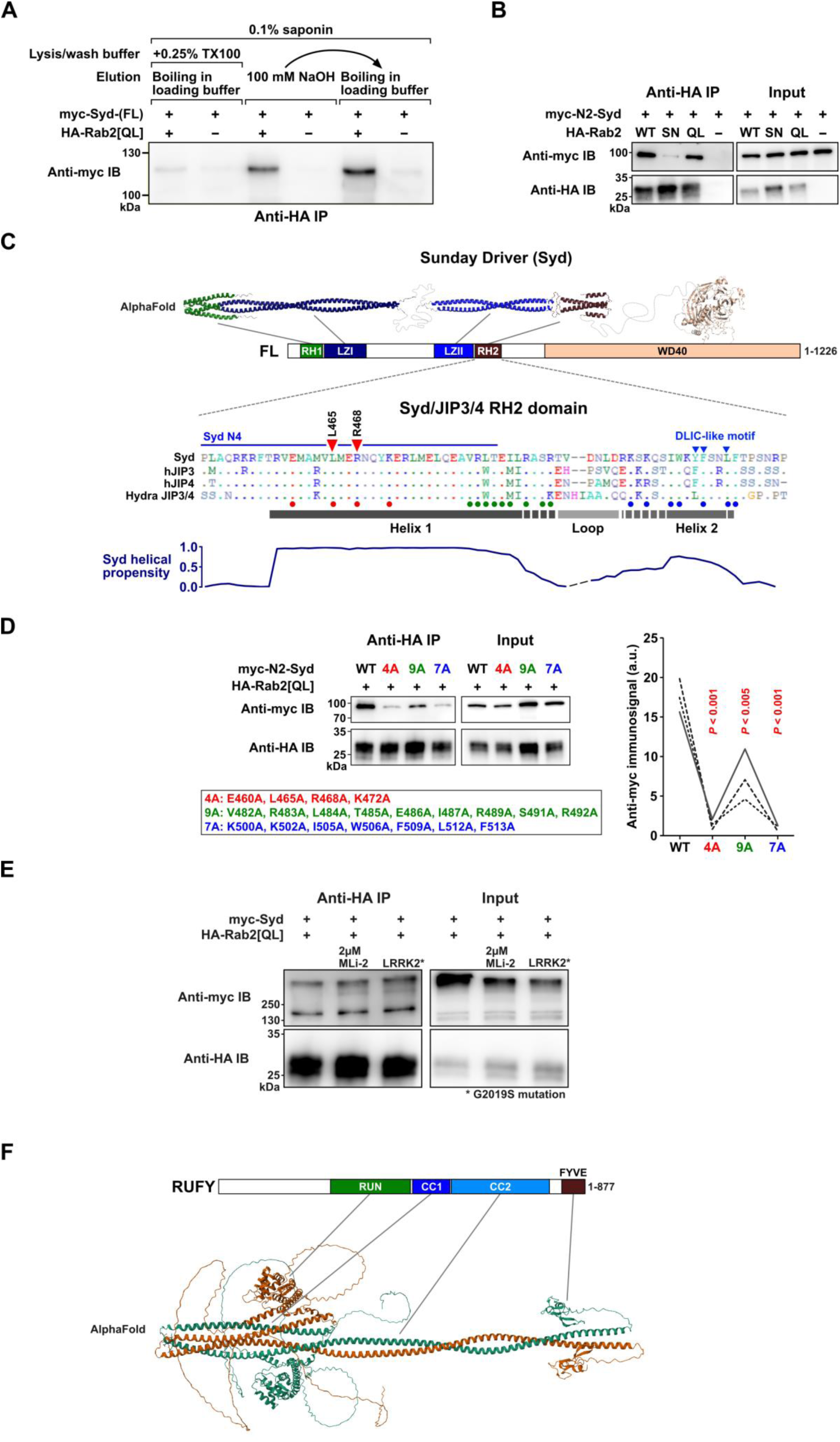
**A**,Co-IP experiment performed on lysates from HEK cells transfected with constructs encoding epitope-tagged Drosophila proteins, illustrating the detergent sensitivity of the Rab2^Q65L^:Syd interaction. Western blot of eluates probed against myc showing co-precipitation of myc-Syd in the presence (lanes 1, 3, and 5) or absence (lanes 2, 4 and 6) of HA-Rab2^Q65L^ when immunoprecipitating against HA. In lanes 1-2 the experiment was performed in the presence of 0.1% saponin and 0.25% Triton-X100, and proteins were eluted from the anti-HA beads by boiling in SDS-PAGE loading buffer. In lanes 3-4 the experiment was performed only in the presence of 0.1% saponin and proteins were eluted with 100 mM NaOH. In lanes 5-6 the same anti-HA beads that were eluted with 100 mM NaOH were boiled in SDS-PAGE loading buffer to elute the remaining protein. **B**, Co-IP of myc-N2-Syd by HA-Rab2, HA-Rab2^S20N^ and HA-Rab2^Q65L^. Compared to the experiment using full length myc-Syd shown in Fig. 2B, the amount of transfecting DNA encoding HA-Rab2^S20N^ was increased to match the higher expression levels of HA-Rab2 and HA-Rab2^Q65L^. **C**, *Top*, expected structure of Syd homodimer assembled from three separate AlphaFold predictions mapped onto the domain architecture of Syd. *Middle*, alignment of the RH2 domain from Drosophila Syd, human JIP3 and JIP4 and the cnidarian (Hydra vulgaris) JIP3/4 ortholog. Dots indicate residue identity to Syd-RH2. The predicted locations of Helix 1, Helix 2 and the intervening loop from the AlphaFold model in Fig. 2H is shown together with a helical propensity estimation (*bottom*). Also indicated are the location of the residues mutated to alanines in D and Fig. 2I, the C-terminal extent of the N4-Syd (Syd^1-485^) truncation and the partially conserved DLIC-like motif involved in autoinhibition ^35^. **D**, *Left*, Co-IP of wild type myc-N2-Syd and three different sets of myc-N2-Syd alanine substitution mutants by HA-Rab2^Q65L^. The position of the mutations is indicated in C. *Right*, Quantification of the anti-myc immunosignal from eluted wild type and mutated myc-N2-Syd (*n* = three independent experiments). **E**, The effect of different levels of LRRK2 activity on co-IP of full length myc-Syd by HA-Rab2^Q65L^. Endogenous HEK cell LRRK2 activity was inhibited by treatment of cells with 2µM MLi-2 for 2 h before lysis or increased by co-transfection with a constitutively active LRRK2^G2019S^ mutant. **F,** Structure of a RUFY dimer predicted using AlphaFold 3, and the RUFY domain architecture.

**Figure S3.**
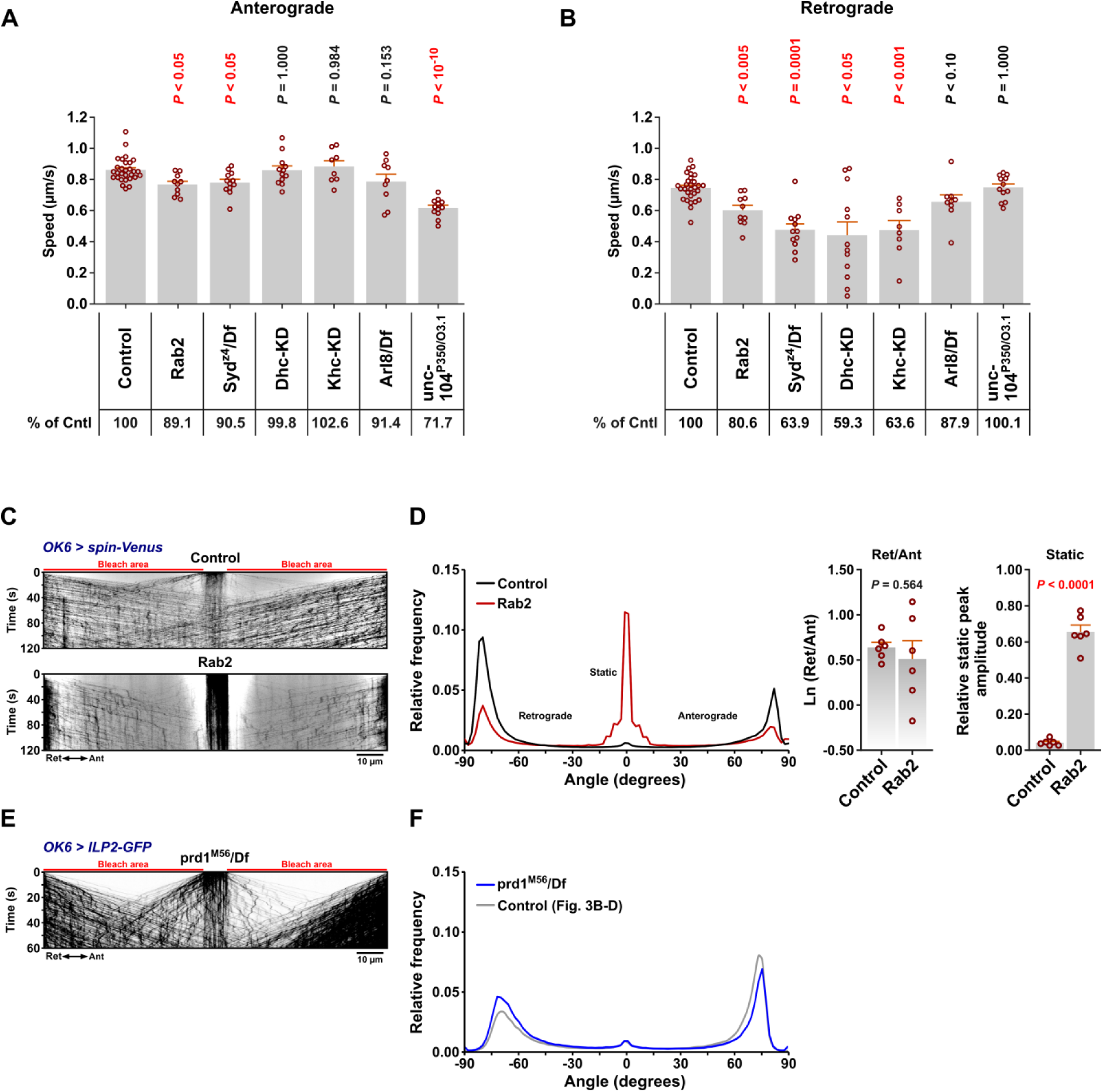
**A-B**, Anterograde (A) and retrograde (B) DCV movement speeds from the experiments in Fig. 3 (*OK6 > ILP2-GFP*). **C**, Representative kymographs showing transport of Spinster-positive organelles in motor axons (*OK6 > Spinster-GFP*) in control and *Rab2* larvae. Scale bar: 10 µm. **D**, *Left*, Directional distributions derived from C, averaged from *n* larvae, where *n* is equal to the number of data points in the bar graphs at the *right*, which depict the logarithmic ratio of retrograde to anterograde peak amplitude, and the relative static peak amplitude. **E**, Kymographs showing DCVs in motor axons of *prd1^M56^/Df* larvae. Scale bar: 10 µm. **F**, Directional distribution derived from E, averaged from four larvae (*blue curve*), shown together with a replica of the directional distribution of control larvae in Fig. 3B-D (*grey curve*). *Arl8/Df* results in A and B, and the results in C and D represent re-analysis of data published earlier ^16^. Bar graphs in A, B and C represent mean+SEM, and each data point represents one larva. Details of statistical tests of the quantitative data in B are contained in the Source File.

**Figure S4.**
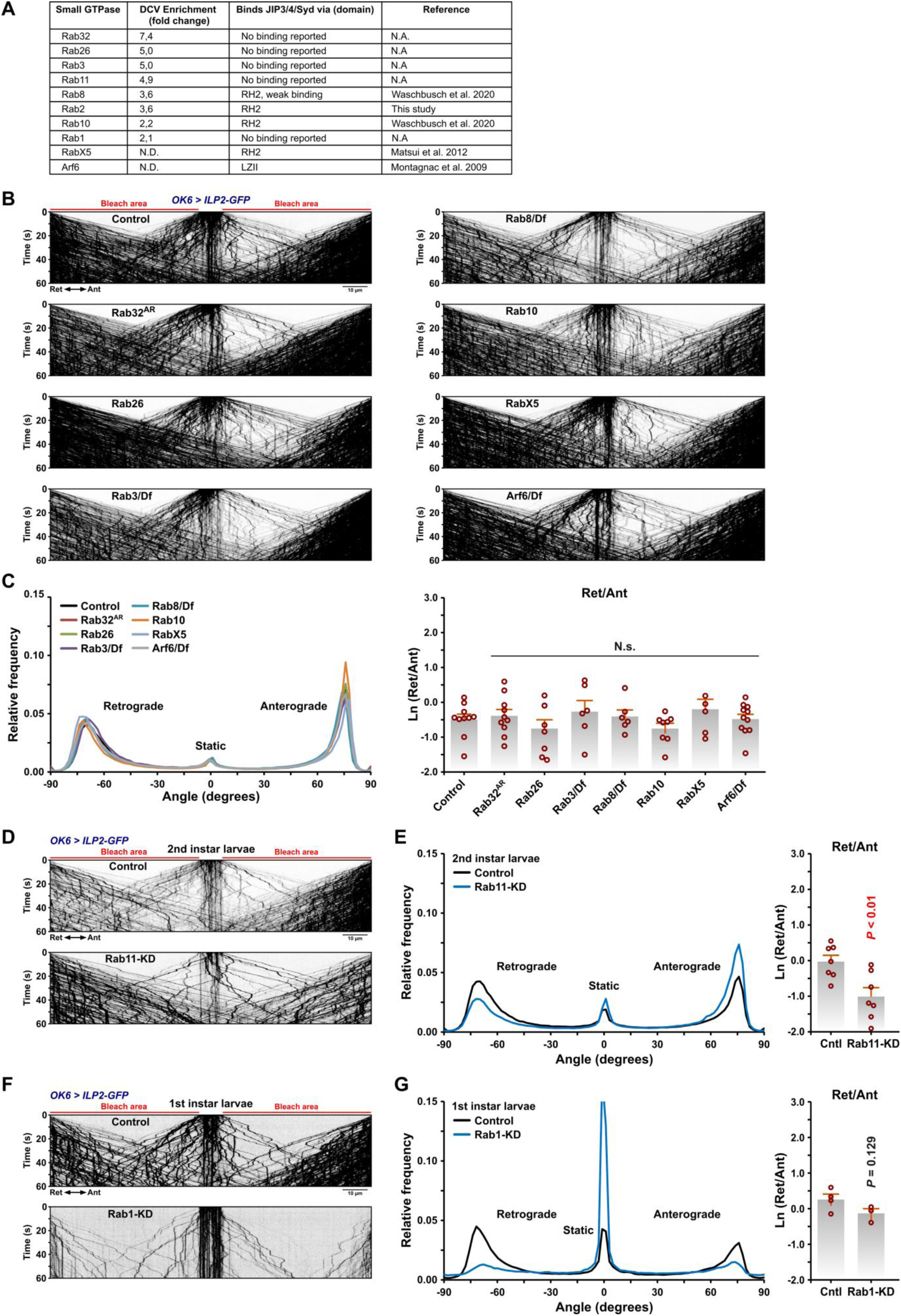
**A**, Small GTPases enriched in the VMAT-specific PDB dataset or known to bind JIP3/4/Syd. **B**, Representative kymographs showing transport of ILP2-GFP-positive DCVs in third instar larval motor axons in controls, Rab GTPase mutants, and *Arf6/Df*. Scale bar: 10 µm. **C**, *Left*, Directional distributions derived from B, averaged from *n* larvae, where *n* is equal to the number of data points in the bar graphs at the *right*, which depict the logarithmic ratio of retrograde to anterograde peak amplitude. N.s., not significant (ANOVA, *P* = 0.522). **D**, Representative kymographs showing transport of ILP2-GFP-positive DCVs in motor axons in control second instar larvae and larvae subjected to motor neuron-specific knockdown of Rab11. Scale bar: 10 µm. **E**, *Left*, Directional distributions derived from D, averaged from *n* larvae, where *n* is equal to the number of data points in the bar graphs at the *right*, which depict the logarithmic ratio of retrograde to anterograde peak amplitude. **F**, Representative kymographs showing transport of ILP2-GFP-positive DCVs in motor axons in control first instar larvae and larvae subjected to motor neuron-specific knockdown of Rab1. Scale bar: 10 µm. **G**, *Left*, Directional distributions derived from F, averaged from *n* larvae, where *n* is equal to the number of data points in the bar graphs at the *right*, which depict the logarithmic ratio of retrograde to anterograde peak amplitude. Bar graphs in C, E, and G represent mean+SEM, and each data point represents one larva. Details of statistical tests of the quantitative data in C, E, and G are contained in the Source File.

**Figure S5.**
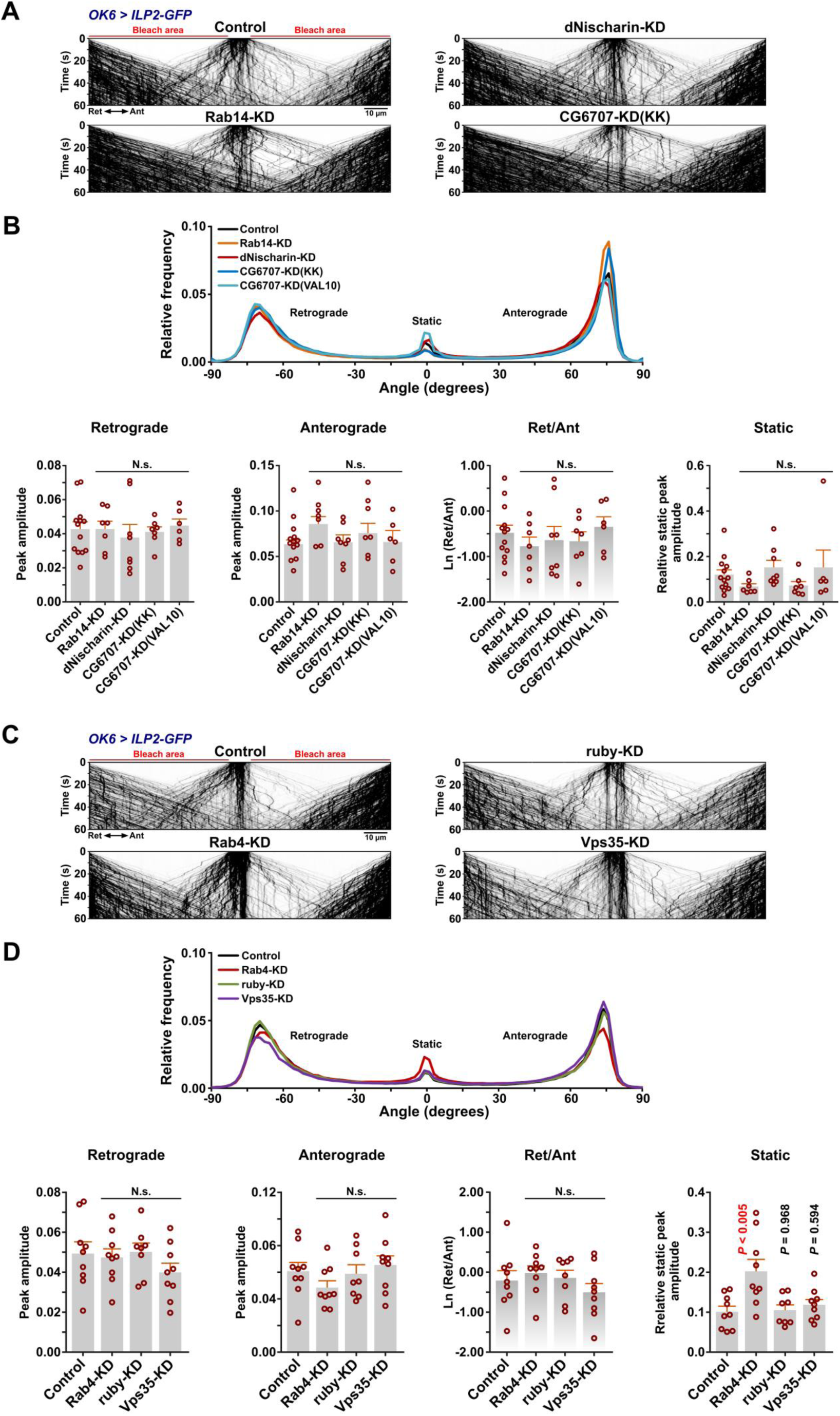
**A**, Representative kymographs showing transport of ILP2-GFP-positive DCVs in motor axons of control larvae and larvae subjected to motor neuron-targeted knockdown of *Rab14*, *dNischarin*, or *CG6707*. Scale bar: 10 µm. **B**, *Top*, Directional distributions derived from A, averaged from *n* larvae, where *n* is equal to the number of data points in the bar graphs at the *bottom*, which depict the retrograde peak amplitude, the anterograde peak amplitude, the logarithmic ratio of retrograde to anterograde peak amplitude, and the relative static peak amplitude. In A and B, “KK” and “VAL20” refer to the use of *UAS-RNAi* lines from the KK collection and the Valium20 vector-based collection, respectively. For simplicity, only KK line data are illustrated in A. **C**, Representative kymographs showing transport of ILP2-GFP-positive DCVs in motor axons of control larvae and larvae subjected to motor neuron-targeted knockdown of *Rab4*, *ruby*, or *Vps35*. Scale bar: 10 µm. **D**, *Top*, Directional distributions derived from C, averaged from *n* larvae, where *n* is equal to the number of data points in the bar graphs at the *bottom*, which depict the retrograde peak amplitude, the anterograde peak amplitude, the logarithmic ratio of retrograde to anterograde peak amplitude, and the relative static peak amplitude. Data in all panels are from third instar larvae. Bar graphs in B and D represent mean+SEM, and each data point represents one larva. N.s., not significant. Details of statistical tests of the quantitative data in B and D are contained in the Source File.

**Figure S6.**
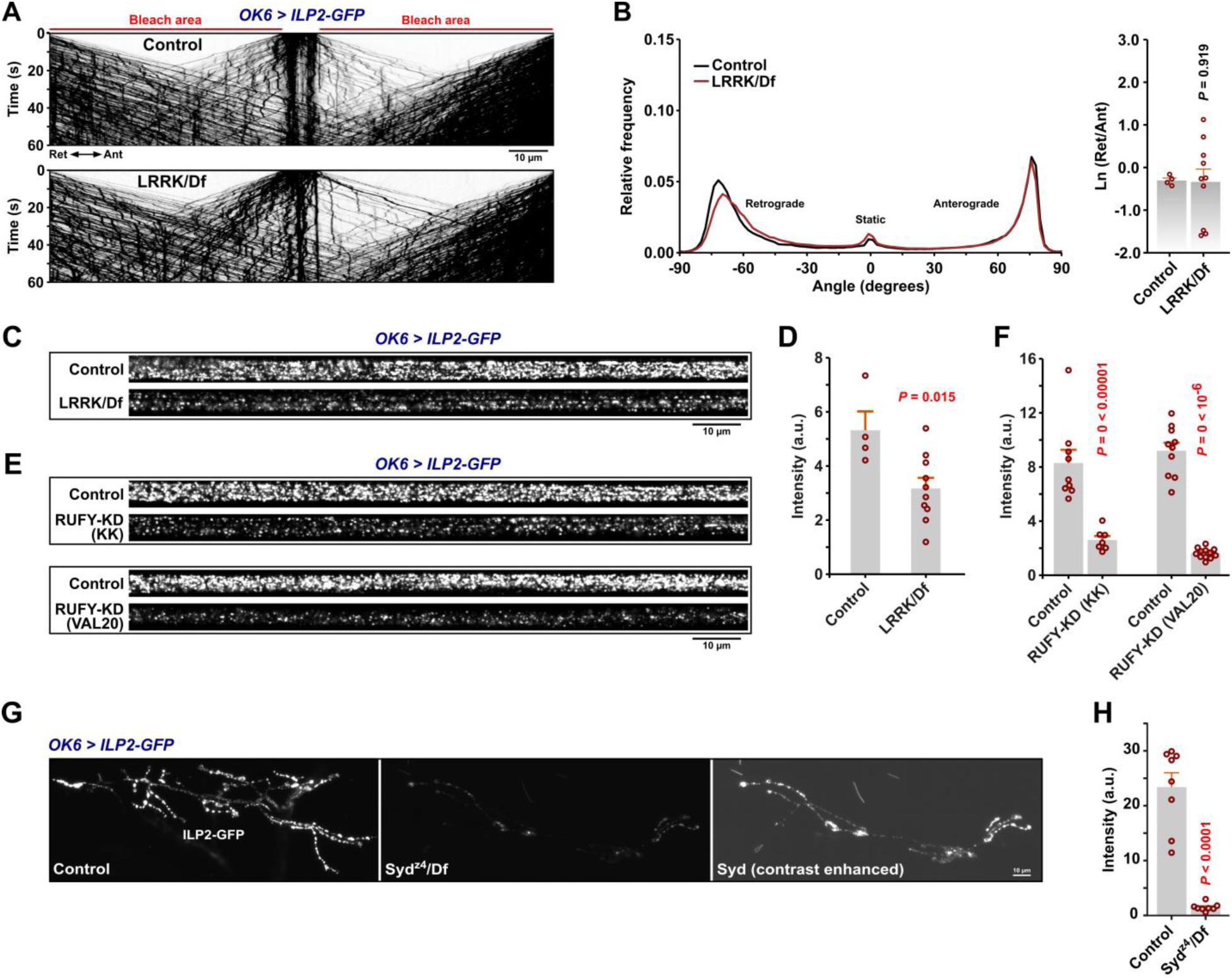
**A**, Representative kymographs showing transport of ILP2-GFP-positive DCVs in motor axons of control and *LRRK/Df* larvae. Scale bar: 10 µm. **B**, *Left*, Directional distributions derived from A, averaged from *n* larvae, where *n* is equal to the number of data points in the bar graphs at the *right*, which depict the logarithmic ratio of retrograde to anterograde peak amplitude. **C**, Representative confocal images showing the pre-bleach ILP2-GFP intensity in A7 nerves of control and *LRRK/Df* larvae. Scale bar: 10 µm. **D**, Quantification of C. **E**, Pre-bleach ILP2-GFP intensity in A7 nerves of control larvae and larvae subjected to motor neuron-targeted knockdown of *RUFY* (KK or Valium20 *UAS-RNAi* lines, cf. Fig. 5). Scale bar: 10 µm. **F**, Quantification of E. **G**, Confocal images showing the intensity of the presynaptic ILP2-GFP signal in the neuromuscular junction of muscle fiber 6/7 in controls and *Syd^z4^/Df*. The *Syd^z4^/Df* micrograph is shown with brightness and contrast settings that match those of the control, as well as with the contrast digitally enhanced for better visibility. Scale bar: 10 µm. Data in all panels are from third instar larvae. Bar graphs in B, D, F, and H represent mean+SEM, and each data point represents one larva. Details of statistical tests of the quantitative data in B, D, F, and H are contained in the Source File.

**Figure S7.**
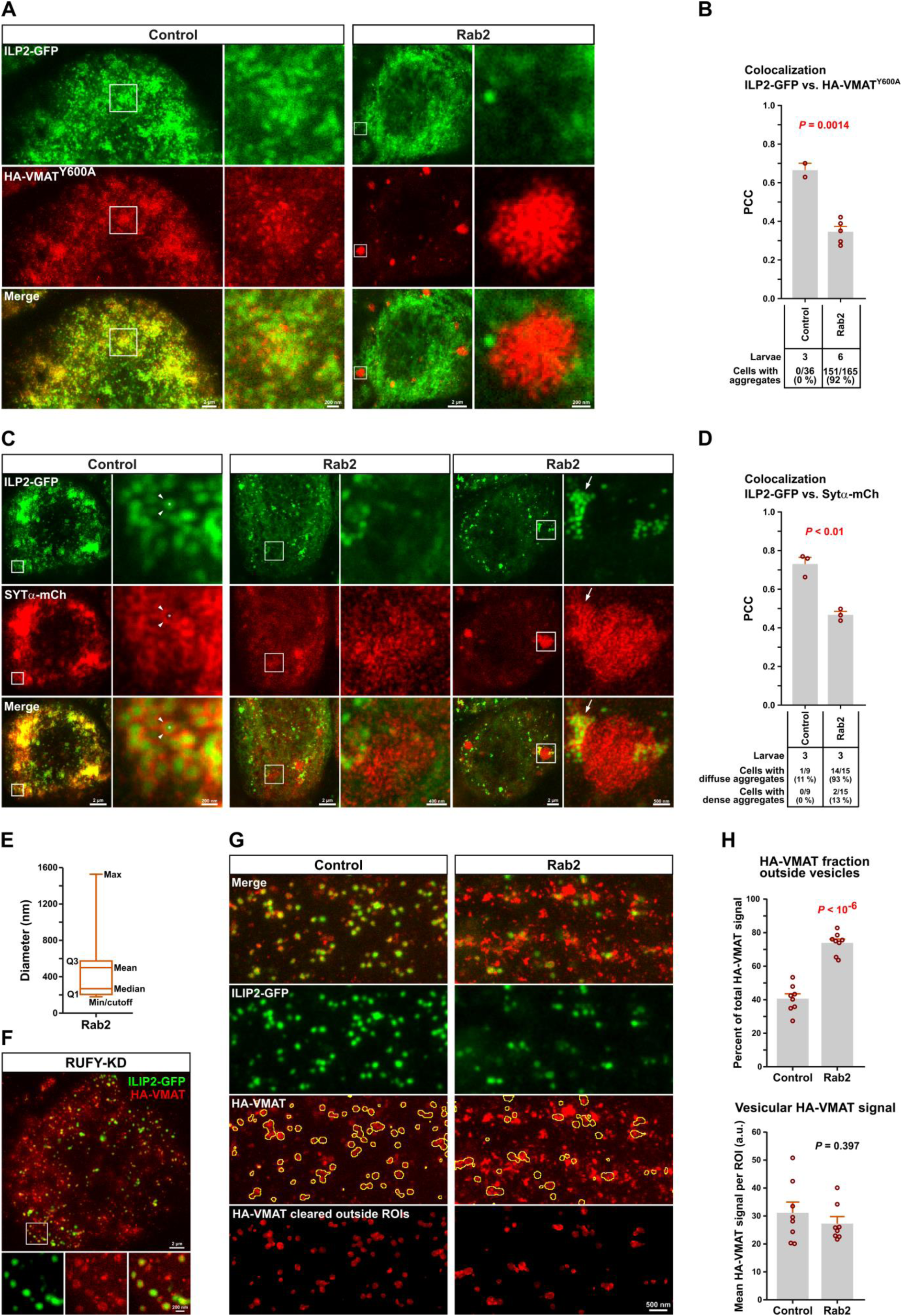
**A**, Spatial relationship between ILP2-GFP and HA-VMAT^Y600A^ in cell bodies. Representative STED images showing the distribution of ILP2-GFP and hemagglutinin (HA)-tagged VMAT^Y600A^ in motor neuron cell bodies in ventral nerve cords of control and *Rab2* third instar larvae. Scale bars (left to right): 2 µm, 200 nm (inset), 2 µm, 200 nm (inset). **B**, Quantification of A. Pearson’s correlation coefficient (PCC) between the ILP2-GFP and HA-VMAT^Y600A^ signals is shown. Counts of HA-VMAT^Y600A^ aggregates are also given (*numbers below graph*). **C**, Spatial relationship between ILP2-GFP and Sytα-mCherry in cell bodies. Representative STED images showing the distribution of ILP2-GFP and mCherry-tagged Sytα in motor neuron cell bodies in ventral nerve cords of control and *Rab2* third instar larvae. Examples of diffuse (*middle*) and dense (*right*) Sytα-mCherry aggregates in *Rab2* mutants are shown. Scale bars (left to right): 2 µm, 200 nm (inset), 2 µm, 400 nm (inset), 2 µm, 500 nm (inset). **D**, The PCC between ILP2-GFP and Sytα-mCherry. Counts of diffuse and dense Sytα-mCherry aggregates are also given (*numbers below graph*). **E,** Size distribution of VMAT aggregates in *Rab2* cell bodies (Fig. 6A). Q1 and Q3, first and third quartile. Diameters were calculated from the areas of the aggregates, assuming a circular shape. A lower diameter cutoff of 178.4 nm (area 0.025 µ^2^) was applied. **F**, Representative STED image showing the distribution of ILP2-GFP and HA-VMAT in a VNC motor neuron cell body from third instar larva subjected to motor neuron-targeted knockdown of *RUFY* (*OK6 > ILP2-GFP, HA-VMAT, RUFY-RNAi^VAL20^)*. Scale bars: 2 µm, 200 nm (inset). **G**, Spatial relationship between ILP2-GFP and HA-tagged wild type VMAT in motor axons. The multiple regions of interest (ROIs) defined by *yellow outlines* mark the DCV-associated ILP2-GFP2 signal superimposed on the VMAT image. In the *bottom row*, the VMAT signal outside the DCV-associated ROIs has been digitally erased. Scale bar: 500 nm. **H**, Quantification of E, showing the percentage of VMAT signal located outside the DCV-associated ROIs (*top*) and the VMAT signal intensity inside the ROIs (*bottom*). Bar graphs in B, D, and H represent mean+SEM, and each data point represents one larva. Details of statistical tests of the data in B, D, and H are contained in the Source File.

Supplementary Tables 1-3

**Supplementary Table 1.** Drosophila lines.

| Line | Source | Identifier |
| --- | --- | --- |
| elav-Gal4 (X) | BDSC | BDSC #458 |
| OK6-Gal4 (II) | BDSC | BDSC #64199 |
| UAS-ILP2-Gal4 (II) | Dr. Edwin S Levitan <sup>1</sup> | N/A |
| UAS-spinster-Venus | Dr. Alyssa Johnson | N/A |
| UAS-DCR2 (X) | BDSC | BDSC #24645 |
| UAS-TurboID-2xHA-Rab2[Q65L] | <sup>20</sup> | N/A |
| UAS-TurboID-2xHA-Rab2[S20N] | <sup>20</sup> | N/A |
| UAS-TurboID-HA-VMAT | this paper | N/A |
| UAS-TurboID | this paper | N/A |
| UAS-HA-VMAT (II) | Dr. David Krantz <sup>48</sup> | N/A |
| UAS-HA-VMAT (III) | Dr. David Krantz <sup>48</sup> | N/A |
| UAS-HA-VMAT[Y600A] (III) | Dr. David Krantz <sup>48</sup> | N/A |
| UAS-syt-α-mCherry (III) | Dr. Paul Taghert <sup>78</sup> | N/A |
| Rab2[Δ1] | <sup>20</sup> | N/A |
| syd[z4] Diap1[th-1] st[1] cu[1] sr[1]<br>e[s] ca[1] | BDSC | BDSC #32016 |
| unc-104[P350] | Dr. William Saxton <sup>13</sup> | N/A |
| unc-104[O3.1] | Dr. William Saxton <sup>13</sup> | N/A |
| PBac{w[+mC]=RB}Lrrk[e03680] | BDSC | BDSC #85160 |
| Rab3[rup] | Dr. Robin Hiesinger <sup>109</sup> | N/A |
| Arf6[GX16w-] | BDSC | BDSC #60585 |
| Rab8[1] | Dr. Robin Hiesinger <sup>110</sup> | N/A |
| Rab26[exon1-2delta] | Dr. Robin Hiesinger <sup>111</sup> | N/A |
| Rab32[AR] | Dr. Robin Hiesinger <sup>112</sup> | N/A |
| Rab10[KO] | Dr. Robin Hiesinger <sup>111</sup> | N/A |
| Prd1[M56] | BDSC | BDSC #37744 |
| RabX5[e04143] | BDSC | BDSC #18228 |
| Df(3L)BSC795 | BDSC | BDSC #27367 |
| Df(3R)BSC141 | BDSC | BDSC #9501 |
| Df(2R)BSC639 | BDSC | BDSC #25729 |
| Df(2R)BSC346 | BDSC | BDSC #24370 |
| Df(3L)BSC445 | BDSC | BDSC #24949 |
| Df(2R)BSC279 | BDSC | BDSC #23664 |
| Df(3R)Exel7310 | BDSC | BDSC #7965 |
| UAS-Dhc64C-RNAi[TRiP.HMS01587] | BDSC | BDSC #36698 |
| UAS-Khc-RNAi[TRiP.GL00330] | BDSC | BDSC #35409 |
| UAS-Rab11-RNAi[KK108297] | VDRC | VDRC id 108382 |
| UAS-Rab11-RNAi[GD11761] | VDRC | VDRC id 22198 |
| UAS-Rab1-RNAi[VSH330620] | VDRC | VDRC id 330620 |
| UAS-RUFY/CG31064-RNAi[TRiP.HMC03246] VAL20 | BDSC | BDSC #51494 |
| UAS-RUFY/CG31064-RNAi[KK100333] | VDRC | VDRC id 103379 |
| UAS-CG6707-RNAi[TRiP.JF02947]<br>VAL10 | BDSC | BDSC #28316 |
| UAS-CG6707-RNAi[KK108707] | VDRC | VDRC id 110291 |
| UAS-Vps35-RNAi[TRiP.HMS01858] VAL20 | BDSC | BDSC #38944 |
| UAS-ruby-RNAi[TRiP.HMS00479]<br>VAL20 | BDSC | BDSC #32477 |
| UAS-Rab4-RNAi[TRiP.HMS01100]<br>VAL20 | BDSC | BDSC #33757 |
| UAS-Rab14-RNAi[TRiP.JF03135]<br>VAL10 | BDSC | BDSC #28708 |
| UAS-dNischarin/CG11807-RNAi[GD7390] | VDRC | VDRC id 38566 |
| UAS-Hep.Act (II) | BDSC | BDSC #9306 |
| UAS-bsk.DN (X) | BDSC | BDSC #6409 |

**Supplementary Table 2.**
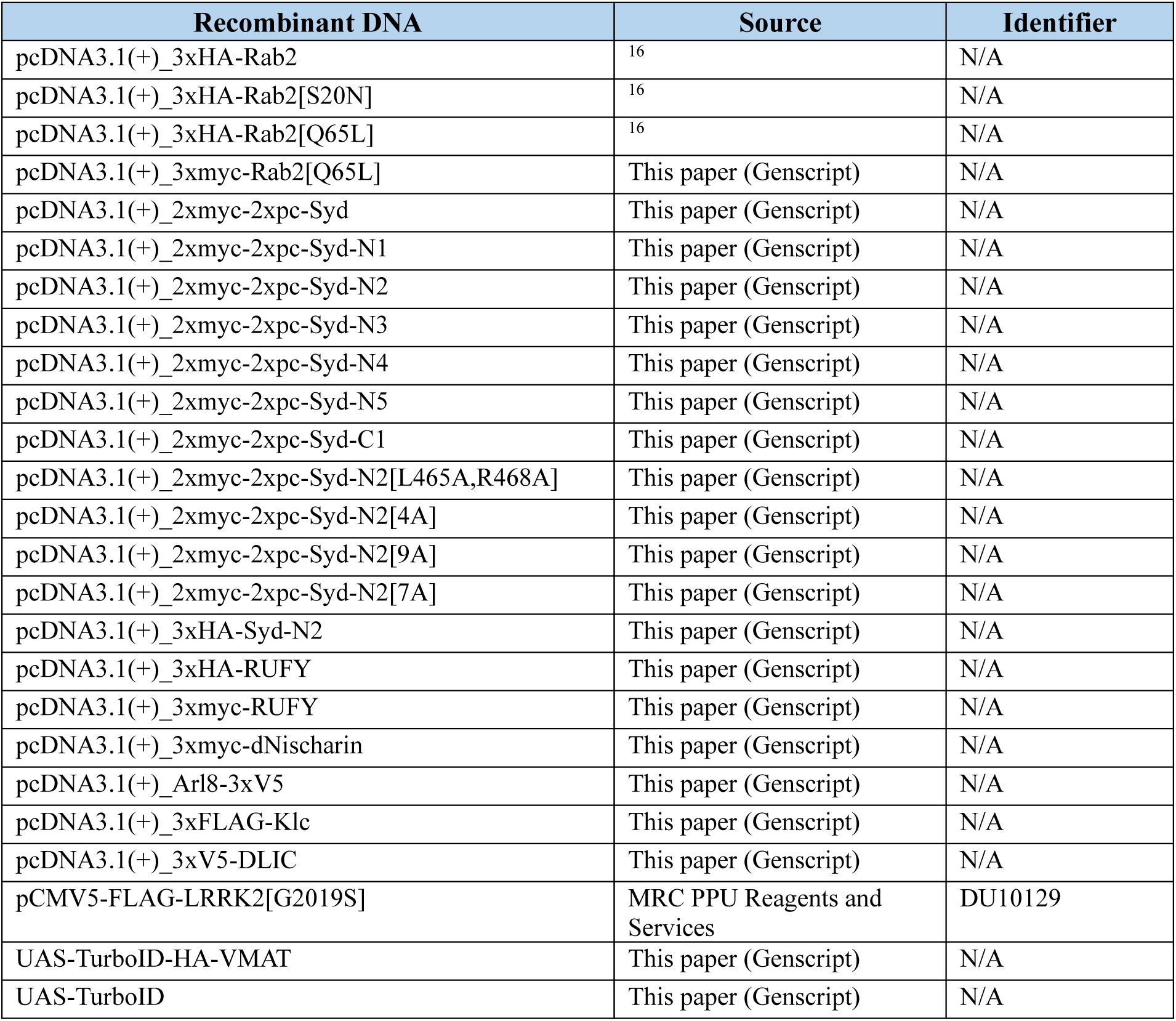
Recombinant DNAs.

| Recombinant DNA | Source | Identifier |
| --- | --- | --- |
| pcDNA3.1(+)_3xHA-Rab2 | 16 | N/A |
| pcDNA3.1(+)_3xHA-Rab2[S20N] | 16 | N/A |
| pcDNA3.1(+)_3xHA-Rab2[Q65L] | 16 | N/A |
| pcDNA3.1(+)_3xmyc-Rab2[Q65L] | This paper (Genscript) | N/A |
| pcDNA3.1(+)_2xmyc-2xpc-Syd | This paper (Genscript) | N/A |
| pcDNA3.1(+)_2xmyc-2xpc-Syd-N1 | This paper (Genscript) | N/A |
| pcDNA3.1(+)_2xmyc-2xpc-Syd-N2 | This paper (Genscript) | N/A |
| pcDNA3.1(+)_2xmyc-2xpc-Syd-N3 | This paper (Genscript) | N/A |
| pcDNA3.1(+)_2xmyc-2xpc-Syd-N4 | This paper (Genscript) | N/A |
| pcDNA3.1(+)_2xmyc-2xpc-Syd-N5 | This paper (Genscript) | N/A |
| pcDNA3.1(+)_2xmyc-2xpc-Syd-C1 | This paper (Genscript) | N/A |
| pcDNA3.1(+)_2xmyc-2xpc-Syd-N2[L465A,R468A] | This paper (Genscript) | N/A |
| pcDNA3.1(+)_2xmyc-2xpc-Syd-N2[4A] | This paper (Genscript) | N/A |
| pcDNA3.1(+)_2xmyc-2xpc-Syd-N2[9A] | This paper (Genscript) | N/A |
| pcDNA3.1(+)_2xmyc-2xpc-Syd-N2[7A] | This paper (Genscript) | N/A |
| pcDNA3.1(+)_3xHA-Syd-N2 | This paper (Genscript) | N/A |
| pcDNA3.1(+)_3xHA-RUFY | This paper (Genscript) | N/A |
| pcDNA3.1(+)_3xmyc-RUFY | This paper (Genscript) | N/A |
| pcDNA3.1(+)_3xmyc-dNischarin | This paper (Genscript) | N/A |
| pcDNA3.1(+)_Arl8-3xV5 | This paper (Genscript) | N/A |
| pcDNA3.1(+)_3xFLAG-Klc | This paper (Genscript) | N/A |
| pcDNA3.1(+)_3xV5-DLIC | This paper (Genscript) | N/A |
| pCMV5-FLAG-LRRK2[G2019S] | MRC PPU Reagents and Services | DU10129 |
| UAS-TurboID-HA-VMAT | This paper (Genscript) | N/A |
| UAS-TurboID | This paper (Genscript) | N/A |

**Supplementary Table 3.** Antibodies and reagents.

| Antibodies | Source | Identifier |
| --- | --- | --- |
| Rat anti-HA 3F10 (WB and IHC, 100 ng/ml) | Sigma-Aldrich | Ref: 11867423001, Lot: 47877600 |
| Mouse anti-HA 6E2 HRP (WB, 1:1000-5000) | Cell Signaling Technology | Ref: 2999S, Lot: 4 |
| Rabbit anti-myc (WB, 1:1000) | Sigma-Aldrich, EMD Millipore Corp. | Ref: 06-549, Lot: 3445807 |
| Rabbit anti-V5 (WB, 1:500) | GenScript | Cat: A00623-40, Lot: 19K002008 |
| Mouse anti-FLAG (WB, 1:1000) | Invitrogen | Ref: MA1-91878, Lot: WF324531 |
| Rabbit anti-mCherry (IHC, 1:100) | Invitrogen | Ref: PA5-34974, Lot: WI3386681C |
| Alpaca anti-GFP VHH single domain antibody/nanobody, custom conjugated to abberior STAR ORNANGE (IHC: 1:100 from 0.5 mg/ml stock) | Chromotech, this paper | Ref: gt-250, Lot: 71017001U |
| Goat anti-rat HRP (WB, 1:8300) | Cell Signaling Technology | Ref: #7077S 01/2012, Lot: 8 |
| Goat anti-rabbit HRP (WB, 1:8300) | Cell Signaling Technology | Ref: #7074S 12/2019, Lot: 28 |
| Goat anti-mouse HRP (WB, 1:2000) | ThermoFisher Scientific | Ref: 31430 |
| Goat anti-rat Abberior STAR RED (IHC, 1:500) | Abberior | Ref: STRED-1007-500UG, Lot: 00928PK-2 |
| Goat anti-rabbit Abberior STAR RED (IHC, 1:500) | Abberior | Ref: STRED-1002-500UG |
| Chemicals, peptides, and recombinant proteins | Source | Identifier |
| Flystuff Nutri-Fly Bloomington Formulation medium | Scientific Laboratory Supplies | FLY1034 |
| GppNHp/GMP-PNP | Sigma-Aldrich | Ref: 0635-25MG, Lot: SLCC2098, SLCN9946 |
| Mli-2 | Abcam | Ref: ab147471, Lot: GR3406958 |
| abberior STAR ORANGE, NHS ester | abberior | Ref: STORANGE-0002-1MG, Lot: 10319RK-1 |
| Pierce™ Anti-HA Magnetic Beads | Thermo Fisher Scientific | Ref: 88836, Lot: VJ307840, WF325730, UL291451, XD335925 |
| Dynabeads MyOne StreptavidinT1 | Thermo Fisher Scientific | Ref: 65601, Lot: 00804134 |
| SuperSignal ELISA Femto Maximum Sensitivity Substrate | Thermo Fisher Scientific | Ref: 37075 |
| Pierce™ Anti-c-Myc Magnetic Beads | Thermo Fisher Scientific | Ref: 88842, Lot: YL384976 |
| Cell lines | Source | Identifier |
| Human: HEK293 cells | ATCC | CRL-1573 |

Legend to Video S1

**Legend for Video S1**

Montage showing time lapse imaging of DCV transport in axons in the A7 nerve of live fillet preparations of third instar *OK6* > ILP2-GFP larvae with the indicated background genotypes. The first video frame is a pre-bleach image, while the subsequent frames shows DCV transport after photobleaching of the nerve, sparing the ∼10 µm wide central region. Ant, anterograde; Ret, retrograde. Scale bar: 10 µm.

Legends to Supplementary Data 1-3

**Legend for Supplementary Data 1**

Table containing fold change and Student’s *t*-test statistics of biotinylated protein label free quantification (LFQ) intensities from flies with pan-neuronal expression of TurboID-Rab2Q65L (*elav > ILP2-GFP, 2xHA-TurboID-Rab2Q65L*) relative to TurboID-Rab2S20N (*elav > ILP2-GFP, 2xHA-TurboID-Rab2S20N*). The Table comprises the Source Data from Fig. 1B.

**Legend for Supplementary Data 2**

Table containing annotation of the candidate proteins (hits) from Supplementary Table 1 that exhibit a fold change in biotinylation intensity of at least four for TurboID-Rab2Q65L relative to TurboID-Rab2S20N. Also included is the subcellular localization classification underlying Figure 1C.

**Legend for Supplementary Data 3**

Table containing fold change and Student’s *t*-test statistics of biotinylated protein LFQ intensities from flies with pan-neuronal expression of TurboID-VMAT (*elav > ILP2-GFP, TurboID-HA-VMAT*) relative to free cytosolic TurboID (*elav > ILP2-GFP, TurboID*). The Table comprises the Source Data from Fig. 1E.

