## Supplementary Data 1 for "Rab2 and Arl8/BORC control retrograde axonal transport of dense core vesicles via Syd/dJIP3/4 and RUFY dynein adaptors"

| identifier | T-statistics | pvalue | mean_group1 | mean_group2 | std(group1) | std(group2) | log2FC | test | correction | padj | rejected | group1 | group2 | FC | -LOG10 pvalue | Method |
|---|---|---|---|---|---|---|---|---|---|---|---|---|---|---|---|---|
| Atg9~Q7JQL5 | 9.930548614 | 0.002172301 | 19.95004086 | 12.1720971 | 0.027560275 | 1.340328009 | 7.777943766 | t-Test | FDR correction BH | 0.016678022 | TRUE | QL | SN | 219.4797102 | 2.663080047 | Paired t-test |
| syd~M9PEX9 | 18.85382881 | 0.000325755 | 19.82601281 | 12.19509202 | 0.240783688 | 0.902275283 | 7.630920791 | t-Test | FDR correction BH | 0.010357789 | TRUE | QL | SN | 198.2147882 | 3.487108856 | Paired t-test |
| Trpml~Q9VW35 | 21.87195205 | 0.000209194 | 18.61485726 | 11.31938598 | 0.195912326 | 0.495427882 | 7.295471287 | t-Test | FDR correction BH | 0.008237157 | TRUE | QL | SN | 157.0925863 | 3.679450489 | Paired t-test |
| Dmel\CG11807~Q7K490 | 19.2529723 | 0.000306038 | 18.54301058 | 11.83747099 | 0.186536041 | 0.585090835 | 6.705539586 | t-Test | FDR correction BH | 0.010357789 | TRUE | QL | SN | 104.3682863 | 3.514224454 | Paired t-test |
| Golgin-45~Q9VHE3 | 16.40942027 | 0.00049251 | 19.52463351 | 13.40761995 | 0.483088106 | 0.513568885 | 6.117013552 | t-Test | FDR correction BH | 0.010461282 | TRUE | QL | SN | 69.40720615 | 3.307585001 | Paired t-test |
| RUFY1~A0A0B4LHR8 | 9.933360262 | 0.0021705 | 17.48785644 | 11.75490517 | 0.331852346 | 0.955414324 | 5.732951271 | t-Test | FDR correction BH | 0.016678022 | TRUE | QL | SN | 53.1851387 | 2.663440158 | Paired t-test |
| Blos2~Q9VTE0 | 13.60280889 | 0.000859408 | 18.52143179 | 12.88855344 | 0.392177089 | 0.331862065 | 5.63287835 | t-Test | FDR correction BH | 0.012559972 | TRUE | QL | SN | 49.62098091 | 3.065800605 | Paired t-test |
| raptor~Q9W437 | 6.471884593 | 0.007486108 | 17.21972494 | 11.63104266 | 0.235284226 | 1.466518729 | 5.588682284 | t-Test | FDR correction BH | 0.029665099 | TRUE | QL | SN | 48.12392085 | 2.125743915 | Paired t-test |
| Dmel\CG17598~Q9VR62 | 8.404310818 | 0.003533952 | 19.17016588 | 13.65038884 | 0.279015428 | 0.995886027 | 5.519777038 | t-Test | FDR correction BH | 0.020315914 | TRUE | QL | SN | 45.87947692 | 2.451739337 | Paired t-test |
| Dmel\CG6707~Q9VT68 | 10.01292302 | 0.00212036 | 19.6239903 | 14.20274131 | 0.615632093 | 0.85143633 | 5.42124898 | t-Test | FDR correction BH | 0.016678022 | TRUE | QL | SN | 42.85076277 | 2.67359035 | Paired t-test |
| Golgin245~Q9W1R3 | 7.702432147 | 0.00454823 | 17.57972249 | 12.17401462 | 0.221316022 | 1.087466689 | 5.405707861 | t-Test | FDR correction BH | 0.023001049 | TRUE | QL | SN | 42.39163957 | 2.342157578 | Paired t-test |
| Dmel\CG11802~Q9VYT1 | 17.26634881 | 0.000423301 | 18.22632818 | 12.93165243 | 0.252637861 | 0.489546683 | 5.294675753 | t-Test | FDR correction BH | 0.010461282 | TRUE | QL | SN | 39.25149647 | 3.373350346 | Paired t-test |
| mon2~X2J564 | 10.04167522 | 0.002102619 | 17.86605654 | 12.76197932 | 1.061968851 | 0.564672554 | 5.104077221 | t-Test | FDR correction BH | 0.016678022 | TRUE | QL | SN | 34.39381442 | 2.677239495 | Paired t-test |
| CT25972~Q9VMK9 | 10.32136159 | 0.00193988 | 17.39759027 | 12.40988429 | 0.501611438 | 0.364594589 | 4.987705978 | t-Test | FDR correction BH | 0.016614137 | TRUE | QL | SN | 31.72846844 | 2.71222505 | Paired t-test |
| Start1~Q9W145 | 12.87514989 | 0.001011258 | 17.46597882 | 12.50976336 | 0.482165409 | 0.386284591 | 4.95621546 | t-Test | FDR correction BH | 0.013062707 | TRUE | QL | SN | 31.04341698 | 2.995138131 | Paired t-test |
| dor~Q24314 | 18.57362273 | 0.000340618 | 16.67110524 | 11.85749774 | 0.678584502 | 0.658378172 | 4.813607498 | t-Test | FDR correction BH | 0.010357789 | TRUE | QL | SN | 28.1216141 | 3.467732676 | Paired t-test |
| TBC1D16~Q9VKV6 | 10.98088128 | 0.001617121 | 18.56302532 | 13.82868086 | 0.204602177 | 0.542824167 | 4.734344464 | t-Test | FDR correction BH | 0.016000463 | TRUE | QL | SN | 26.61826187 | 2.79125736 | Paired t-test |
| car~A4V4S5 | 12.18905106 | 0.001188877 | 17.77341811 | 13.05259135 | 0.974776081 | 0.426621097 | 4.720826762 | t-Test | FDR correction BH | 0.013862839 | TRUE | QL | SN | 26.37002002 | 2.924863243 | Paired t-test |
| RagC-D~Q7K519 | 17.04175671 | 0.000440119 | 18.23216199 | 13.53363089 | 0.204447241 | 0.527719495 | 4.6985311 | t-Test | FDR correction BH | 0.010461282 | TRUE | QL | SN | 25.96562596 | 3.356429713 | Paired t-test |
| nSyb~M9PDW0 | 6.094453343 | 0.00887353 | 18.35405599 | 13.68306119 | 0.850615901 | 0.860460722 | 4.670994805 | t-Test | FDR correction BH | 0.033338521 | TRUE | QL | SN | 25.47472743 | 2.051903594 | Paired t-test |
| Dmel\CG14969~Q95SH7 | 23.02397772 | 0.000179468 | 17.92157745 | 13.25640615 | 0.302265919 | 0.627535582 | 4.665171292 | t-Test | FDR correction BH | 0.007523507 | TRUE | QL | SN | 25.37210463 | 3.746011832 | Paired t-test |
| Dmel\CG6398~Q9VX17 | 13.407881 | 0.000896935 | 17.22889669 | 12.597737 | 0.637151623 | 0.202849001 | 4.631159694 | t-Test | FDR correction BH | 0.012641521 | TRUE | QL | SN | 24.78095177 | 3.047238946 | Paired t-test |
| Atg4b~Q9VF80 | 16.36145827 | 0.000496815 | 18.11930313 | 13.48975896 | 0.160693558 | 0.35947948 | 4.629544176 | t-Test | FDR correction BH | 0.010461282 | TRUE | QL | SN | 24.75321781 | 3.30380505 | Paired t-test |
| Syt4~Q9U6P7 | 5.680093001 | 0.010813087 | 18.45801569 | 13.90134269 | 0.205600774 | 1.386400521 | 4.556672999 | t-Test | FDR correction BH | 0.036454821 | TRUE | QL | SN | 23.53397312 | 1.96605031 | Paired t-test |
| Dmel\CG44247~Q9W0Z5 | 9.668789381 | 0.002348974 | 17.46841759 | 12.97901138 | 0.448646519 | 0.371889779 | 4.489406213 | t-Test | FDR correction BH | 0.017164751 | TRUE | QL | SN | 22.46187121 | 2.629121778 | Paired t-test |
| ldlCp~Q9VF78 | 6.709653112 | 0.006755976 | 16.47818896 | 12.02708137 | 0.572877342 | 0.962130898 | 4.451107583 | t-Test | FDR correction BH | 0.028247427 | TRUE | QL | SN | 21.8734303 | 2.170311911 | Paired t-test |
| garz~A1Z8W8 | 9.954957645 | 0.002156736 | 17.52640135 | 13.17310824 | 0.520044685 | 0.673116813 | 4.353293108 | t-Test | FDR correction BH | 0.016678022 | TRUE | QL | SN | 20.43957237 | 2.666203078 | Paired t-test |
| Nhe3~D5SHT8 | 23.62770928 | 0.000166116 | 17.15050422 | 12.80920127 | 0.251147403 | 0.191290888 | 4.341302951 | t-Test | FDR correction BH | 0.007209085 | TRUE | QL | SN | 20.27040419 | 3.779587591 | Paired t-test |
| Syx13~Q9VU45 | 25.04894492 | 0.000139514 | 19.6739691 | 15.35189003 | 0.436279806 | 0.495416369 | 4.322079068 | t-Test | FDR correction BH | 0.007209085 | TRUE | QL | SN | 20.00209304 | 3.855383738 | Paired t-test |
| CT30701~Q8IQH6 | 12.69619505 | 0.001053986 | 17.11731277 | 12.91380675 | 0.159306774 | 0.504843812 | 4.203506022 | t-Test | FDR correction BH | 0.013092474 | TRUE | QL | SN | 18.42389287 | 2.977165207 | Paired t-test |
| Dmel\CG34126~M9PC65 | 46.71070911 | 2.16E-05 | 17.29402769 | 13.09843639 | 0.107608435 | 0.076458828 | 4.195591294 | t-Test | FDR correction BH | 0.003864721 | TRUE | QL | SN | 18.32309482 | 4.665494733 | Paired t-test |
| Rme-8~A0A0B4LFW6 | 5.136651841 | 0.014293394 | 16.96054579 | 12.77145067 | 0.443871099 | 1.154163424 | 4.189095119 | t-Test | FDR correction BH | 0.043787262 | TRUE | QL | SN | 18.24077497 | 1.844864639 | Paired t-test |
| dp18~Q9VW73 | 13.33121922 | 0.00091229 | 17.31653358 | 13.13880284 | 0.48601804 | 0.174412889 | 4.17773074 | t-Test | FDR correction BH | 0.012641521 | TRUE | QL | SN | 18.0976534 | 3.039866866 | Paired t-test |
| Golgin104~Q9VV40 | 14.08463956 | 0.00077519 | 17.03377496 | 12.86429561 | 0.320848 | 0.251361483 | 4.169479346 | t-Test | FDR correction BH | 0.012134178 | TRUE | QL | SN | 17.99444057 | 3.110592077 | Paired t-test |
| ClC-b~Q7JZ25 | 76.16041525 | 4.99E-06 | 19.59933373 | 15.49931941 | 0.358345617 | 0.304055018 | 4.100014316 | t-Test | FDR correction BH | 0.003100532 | TRUE | QL | SN | 17.14854556 | 5.301986641 | Paired t-test |
| Snx3~Q9VG51 | 13.64690587 | 0.000851209 | 16.66566534 | 12.68476026 | 0.249152624 | 0.364639395 | 3.980905083 | t-Test | FDR correction BH | 0.012555335 | TRUE | QL | SN | 15.78962588 | 3.069963705 | Paired t-test |
| Dmel\CG32850~Q8SX35 | 24.05645822 | 0.000157428 | 16.93651507 | 13.125331 | 0.276074348 | 0.250655338 | 3.81118407 | t-Test | FDR correction BH | 0.007209085 | TRUE | QL | SN | 14.03720764 | 3.802919351 | Paired t-test |
| Dmel\CG17180~Q9W0Q3 | 24.06586835 | 0.000157244 | 18.47433472 | 14.66605357 | 0.120590144 | 0.201577261 | 3.808281146 | t-Test | FDR correction BH | 0.007209085 | TRUE | QL | SN | 14.00899102 | 3.803426796 | Paired t-test |
| CG16823~Q9VK04 | 12.17022788 | 0.001194314 | 18.47078862 | 14.66490564 | 0.298330449 | 0.529061845 | 3.805882981 | t-Test | FDR correction BH | 0.013862839 | TRUE | QL | SN | 13.98572348 | 2.922881652 | Paired t-test |
| Dmel\CG1513~A1Z814 | 10.64187096 | 0.001773294 | 16.91804212 | 13.12321678 | 0.580355819 | 0.707449951 | 3.79482534 | t-Test | FDR correction BH | 0.016234815 | TRUE | QL | SN | 13.87893864 | 2.751219174 | Paired t-test |
| lt~Q7PL76 | 9.94364668 | 0.00216393 | 16.9457247 | 13.15657589 | 0.835035323 | 0.322395743 | 3.789148811 | t-Test | FDR correction BH | 0.016678022 | TRUE | QL | SN | 13.82443689 | 2.664756797 | Paired t-test |
| Sytalpha~Q9I7M6 | 14.19925995 | 0.000756786 | 16.10609095 | 12.32880881 | 0.39585823 | 0.151318104 | 3.777282142 | t-Test | FDR correction BH | 0.012134178 | TRUE | QL | SN | 13.71119245 | 3.121027036 | Paired t-test |
| Dmel\CG9205~Q9W0K9 | 3.547746571 | 0.038152001 | 18.50158062 | 14.73573776 | 0.630361127 | 1.472365825 | 3.765842856 | t-Test | FDR correction BH | 0.082008824 | FALSE | QL | SN | 13.6029048 | 1.418482681 | Paired t-test |
| Vps16A~Q9VHG1 | 10.8991006 | 0.001653071 | 17.45902232 | 13.70533827 | 0.250907803 | 0.479724327 | 3.753684055 | t-Test | FDR correction BH | 0.016191675 | TRUE | QL | SN | 13.48874345 | 2.78170856 | Paired t-test |
| Syb~H1UUB1 | 15.40298255 | 0.000594436 | 17.79432515 | 14.063012 | 0.499806405 | 0.226604733 | 3.731313151 | t-Test | FDR correction BH | 0.011237107 | TRUE | QL | SN | 13.28119587 | 3.225894635 | Paired t-test |
| rg~X2JAH8 | 4.978311889 | 0.015576884 | 17.72896183 | 14.01963189 | 0.200053114 | 1.166187348 | 3.709329937 | t-Test | FDR correction BH | 0.046408873 | TRUE | QL | SN | 13.08035633 | 1.807519406 | Paired t-test |
| PICK1~Q86PF5 | 16.80940088 | 0.000458467 | 16.48685894 | 12.79955626 | 0.191158166 | 0.334328611 | 3.68730268 | t-Test | FDR correction BH | 0.010461282 | TRUE | QL | SN | 12.88216062 | 3.338691727 | Paired t-test |
| spin~Q9GQQ0-4 | 26.64157737 | 0.000116036 | 16.59624041 | 12.94532361 | 0.106182736 | 0.237572731 | 3.650916796 | t-Test | FDR correction BH | 0.006717766 | TRUE | QL | SN | 12.56132543 | 3.935407439 | Paired t-test |
| Snap25~P36975-3 | 18.6816893 | 0.000334781 | 16.58937829 | 12.99193293 | 0.424900402 | 0.177656266 | 3.597445358 | t-Test | FDR correction BH | 0.010357789 | TRUE | QL | SN | 12.10427998 | 3.475239394 | Paired t-test |
| Dmel\CG7970~Q9W0B2 | 4.911680404 | 0.016161867 | 17.77564317 | 14.20339274 | 0.30813503 | 1.094834708 | 3.572250429 | t-Test | FDR correction BH | 0.04758938 | TRUE | QL | SN | 11.89472843 | 1.791508467 | Paired t-test |
| cbs~A0A0B4K864 | 3.972396786 | 0.028520267 | 18.23898521 | 14.67270302 | 0.333246382 | 1.599708329 | 3.566282191 | t-Test | FDR correction BH | 0.068013152 | FALSE | QL | SN | 11.84562316 | 1.54484641 | Paired t-test |
| NijA~M9PHZ8 | 16.67074372 | 0.000469903 | 16.57479999 | 13.0102946 | 0.170344004 | 0.308378149 | 3.564505387 | t-Test | FDR correction BH | 0.010461282 | TRUE | QL | SN | 11.83104323 | 3.327991453 | Paired t-test |
| Golgin84~Q8SZ63 | 16.73788497 | 0.000464319 | 17.50614703 | 13.94968246 | 0.34421781 | 0.1206702 | 3.55646457 | t-Test | FDR correction BH | 0.010461282 | TRUE | QL | SN | 11.76528668 | 3.333183674 | Paired t-test |
| Vamp7~Q7JYX5 | 10.60283379 | 0.001792543 | 16.38633007 | 12.86357108 | 0.410725946 | 0.448587509 | 3.522758982 | t-Test | FDR correction BH | 0.016234895 | TRUE | QL | SN | 11.49360112 | 2.746530425 | Paired t-test |
| BicD~M9PDL9 | 13.22743731 | 0.000933639 | 20.04396369 | 16.52632813 | 0.399276107 | 0.366640457 | 3.517635552 | t-Test | FDR correction BH | 0.012659318 | TRUE | QL | SN | 11.45285639 | 3.029820792 | Paired t-test |
| Syt7~Q9V4C4 | 8.106019389 | 0.0039242 | 17.39469988 | 13.88224631 | 0.546100307 | 0.822652786 | 3.512453567 | t-Test | FDR correction BH | 0.021213554 | TRUE | QL | SN | 11.41179292 | 2.406248902 | Paired t-test |
| Vps13B~Q9VAD3 | 14.15649744 | 0.000763583 | 16.23236044 | 12.72850779 | 0.152019894 | 0.324761748 | 3.50385265 | t-Test | FDR correction BH | 0.012134178 | TRUE | QL | SN | 11.34396161 | 3.117143618 | Paired t-test |
| IA-2~Q59E11 | 24.46175133 | 0.000149762 | 18.82142415 | 15.40604721 | 0.253608781 | 0.079507767 | 3.415376931 | t-Test | FDR correction BH | 0.007209085 | TRUE | QL | SN | 10.66917658 | 3.824598573 | Paired t-test |
| Scamp~Q9VXV6 | 11.18296624 | 0.001532629 | 16.37885111 | 12.99059328 | 0.781008748 | 0.314452784 | 3.388257836 | t-Test | FDR correction BH | 0.015838197 | TRUE | QL | SN | 10.47049567 | 2.814562879 | Paired t-test |
| Dmel\CG3652~Q9VQZ0 | 13.10916486 | 0.000958786 | 16.17263533 | 12.789509 | 0.506704364 | 0.143524923 | 3.383126328 | t-Test | FDR correction BH | 0.012773921 | TRUE | QL | SN | 10.43331942 | 3.0182781 | Paired t-test |
| Fas2~M9PJ19 | 10.75745608 | 0.001717885 | 17.04954078 | 13.68967032 | 0.371312114 | 0.883407643 | 3.35987046 | t-Test | FDR correction BH | 0.016207074 | TRUE | QL | SN | 10.26648531 | 2.765005988 | Paired t-test |
| RhoGAP68F~F0JAN9 | 3.930680737 | 0.029316816 | 16.74431704 | 13.41103871 | 0.812383851 | 0.974658844 | 3.333278325 | t-Test | FDR correction BH | 0.068933059 | FALSE | QL | SN | 10.07898409 | 1.532883201 | Paired t-test |
| Hsl~Q7JR83 | 16.39873867 | 0.000493464 | 16.59497976 | 13.26366725 | 0.430952373 | 0.110502026 | 3.33131251 | t-Test | FDR correction BH | 0.010461282 | TRUE | QL | SN | 10.06525983 | 3.306744111 | Paired t-test |
| Frq2~Q9VWX8 | 7.349403807 | 0.005205956 | 16.2998681 | 12.98505333 | 0.580019788 | 0.298793803 | 3.314814767 | t-Test | FDR correction BH | 0.024608571 | TRUE | QL | SN | 9.950815521 | 2.28349948 | Paired t-test |
| hook~Q24185 | 9.065410268 | 0.002835331 | 17.10373568 | 13.8271703 | 0.642322248 | 0.028054941 | 3.276565385 | t-Test | FDR correction BH | 0.018663977 | TRUE | QL | SN | 9.69046157 | 2.547396228 | Paired t-test |
| mtd~B7Z0T3 | 4.121715369 | 0.025887592 | 17.88277039 | 14.60852272 | 0.520106011 | 1.759161969 | 3.274247669 | t-Test | FDR correction BH | 0.063738692 | FALSE | QL | SN | 9.674906168 | 1.586908346 | Paired t-test |
| VAChT~O17444 | 7.936674585 | 0.004171337 | 18.0719437 | 14.82955744 | 0.846221745 | 0.181293757 | 3.242386258 | t-Test | FDR correction BH | 0.021858353 | TRUE | QL | SN | 9.46358138 | 2.379724763 | Paired t-test |
| CG14261~Q8IMP0 | 9.441475449 | 0.002518166 | 16.3796965 | 13.13946922 | 0.156871882 | 0.615393614 | 3.240227277 | t-Test | FDR correction BH | 0.017810152 | TRUE | QL | SN | 9.449429803 | 2.598915601 | Paired t-test |
| CG6327~M9PEU1 | 48.03002608 | 1.99E-05 | 18.09728692 | 14.87514314 | 0.288861696 | 0.401690887 | 3.222143778 | t-Test | FDR correction BH | 0.003864721 | TRUE | QL | SN | 9.331724905 | 4.701745043 | Paired t-test |
| Atg17~Q7KTS2 | 4.858429007 | 0.016650095 | 16.68848087 | 13.48340217 | 0.24189231 | 1.228101009 | 3.205078706 | t-Test | FDR correction BH | 0.048228926 | TRUE | QL | SN | 9.221993865 | 1.778583285 | Paired t-test |
| nrv2~Q1RL12 | 3.460840559 | 0.04061536 | 17.24508853 | 14.04206245 | 0.089321658 | 1.542531528 | 3.203026087 | t-Test | FDR correction BH | 0.084890988 | FALSE | QL | SN | 9.208882443 | 1.391309696 | Paired t-test |
| cue~Q95RU0 | 16.11545544 | 0.000519703 | 16.40262591 | 13.2027284 | 0.722391733 | 0.577992229 | 3.199897504 | t-Test | FDR correction BH | 0.01061394 | TRUE | QL | SN | 9.188933991 | 3.284244537 | Paired t-test |
| Dmel\CG8500~Q9VH66 | 4.329300944 | 0.02272538 | 16.40147975 | 13.20251418 | 0.698210525 | 0.705509123 | 3.198965566 | t-Test | FDR correction BH | 0.058768719 | FALSE | QL | SN | 9.183000128 | 1.643488836 | Paired t-test |
| Syx16~Q9VR90 | 8.67770701 | 0.003220131 | 16.53305618 | 13.34991334 | 0.292275622 | 0.426541291 | 3.183142835 | t-Test | FDR correction BH | 0.019559199 | TRUE | QL | SN | 9.082836023 | 2.4921264 | Paired t-test |
| Iml1~Q9W0E3-2 | 6.263871507 | 0.008212279 | 15.88373577 | 12.7075021 | 0.87979527 | 0.612499519 | 3.17623367 | t-Test | FDR correction BH | 0.03159942 | TRUE | QL | SN | 9.039441697 | 2.085536317 | Paired t-test |
| ergic53~Q9V3A8 | 10.38792284 | 0.00190363 | 16.02048092 | 12.85540953 | 0.209660988 | 0.432130673 | 3.165071393 | t-Test | FDR correction BH | 0.01648088 | TRUE | QL | SN | 8.969772492 | 2.720417556 | Paired t-test |
| mv~Q9W060 | 6.316046661 | 0.008021732 | 15.94577991 | 12.78775848 | 0.666890734 | 0.250047642 | 3.158021433 | t-Test | FDR correction BH | 0.031091532 | TRUE | QL | SN | 8.926047188 | 2.095731827 | Paired t-test |
| Dmel\CG18065~A1ZBV5 | 9.609101221 | 0.002391905 | 16.45140071 | 13.32462258 | 0.37475225 | 0.353681205 | 3.126778125 | t-Test | FDR correction BH | 0.017319568 | TRUE | QL | SN | 8.734820915 | 2.621256044 | Paired t-test |
| VGlut~Q9VQC0 | 6.833679027 | 0.006412114 | 16.53969802 | 13.42497771 | 0.819031978 | 0.594425709 | 3.114720314 | t-Test | FDR correction BH | 0.027532339 | TRUE | QL | SN | 8.662120932 | 2.192998752 | Paired t-test |
| Nrx-IV~Q8IQH0 | 6.633318221 | 0.00697975 | 15.48525705 | 12.37503157 | 0.351561438 | 0.525164179 | 3.110225485 | t-Test | FDR correction BH | 0.028582883 | TRUE | QL | SN | 8.635175414 | 2.156160148 | Paired t-test |
| htt~Q9V3N4 | 5.824857713 | 0.010077478 | 16.73003379 | 13.62564588 | 0.570019572 | 0.870687161 | 3.104387918 | t-Test | FDR correction BH | 0.034898746 | TRUE | QL | SN | 8.600305558 | 1.996648126 | Paired t-test |
| Syx8~Q9VV76 | 14.39978708 | 0.000725961 | 16.11806038 | 13.02296305 | 0.146079623 | 0.508128149 | 3.095097332 | t-Test | FDR correction BH | 0.012134178 | TRUE | QL | SN | 8.545099743 | 3.139086572 | Paired t-test |
| GCC185~Q9VGE4 | 7.323056013 | 0.005259949 | 18.16021962 | 15.07396038 | 0.607639923 | 0.257028044 | 3.086259241 | t-Test | FDR correction BH | 0.024737406 | TRUE | QL | SN | 8.492911644 | 2.279018478 | Paired t-test |
| p115~Q9W3N6 | 38.46372958 | 3.87E-05 | 15.81204174 | 12.74151038 | 0.337837923 | 0.203330296 | 3.070531366 | t-Test | FDR correction BH | 0.003902745 | TRUE | QL | SN | 8.400827046 | 4.412739202 | Paired t-test |
| Q9VP46~Q9VP46 | 42.14004266 | 2.94E-05 | 15.87016054 | 12.80373327 | 0.223757829 | 0.180623529 | 3.066427268 | t-Test | FDR correction BH | 0.003864721 | TRUE | QL | SN | 8.376962809 | 4.5314939 | Paired t-test |
| Pex11~Q7JZE1 | 13.95208086 | 0.000797223 | 16.22351211 | 13.16748395 | 0.478913657 | 0.447759613 | 3.056028153 | t-Test | FDR correction BH | 0.012329867 | TRUE | QL | SN | 8.316797776 | 3.098420156 | Paired t-test |
| Pex23~Q9VWB0 | 4.845751119 | 0.01676915 | 16.70740357 | 13.65398684 | 1.158636735 | 0.896571592 | 3.053416738 | t-Test | FDR correction BH | 0.048376646 | TRUE | QL | SN | 8.301757198 | 1.775488948 | Paired t-test |
| Dmel\CG8785~Q7K2W3 | 4.642019149 | 0.018843707 | 16.22186936 | 13.18551045 | 0.482436208 | 0.677152338 | 3.036358913 | t-Test | FDR correction BH | 0.05220526 | FALSE | QL | SN | 8.204178678 | 1.724833666 | Paired t-test |
| Dmel\CG15210~Q9VZ67 | 11.70564653 | 0.001339647 | 16.4761736 | 13.47243321 | 0.181176526 | 0.591351948 | 3.003740387 | t-Test | FDR correction BH | 0.014923481 | TRUE | QL | SN | 8.020768021 | 2.873009615 | Paired t-test |
| fws~M9NCV1 | 4.552868786 | 0.019856377 | 16.71290652 | 13.74947269 | 0.666803261 | 1.274528126 | 2.963433823 | t-Test | FDR correction BH | 0.053431096 | FALSE | QL | SN | 7.799782112 | 1.702099983 | Paired t-test |
| Vps20~Q9W236 | 6.443623137 | 0.007579676 | 16.66940159 | 13.73902818 | 0.238134092 | 0.721430314 | 2.930373411 | t-Test | FDR correction BH | 0.029813392 | TRUE | QL | SN | 7.6230768 | 2.120349364 | Paired t-test |
| MESK2~Q9I7V7 | 17.98191572 | 0.000375101 | 16.3896526 | 13.52739449 | 0.235157603 | 0.302414849 | 2.862258106 | t-Test | FDR correction BH | 0.010461282 | TRUE | QL | SN | 7.271525731 | 3.42585131 | Paired t-test |
| CG11976~Q9VHI9 | 13.88970646 | 0.000807879 | 15.80390144 | 12.95166423 | 0.255463468 | 0.366092004 | 2.852237211 | t-Test | FDR correction BH | 0.012374532 | TRUE | QL | SN | 7.221193047 | 3.092653663 | Paired t-test |
| Dmel\CG1486~Q9VRG8 | 5.2813117 | 0.013238855 | 15.88558057 | 13.03458588 | 0.406899446 | 0.697753006 | 2.850994695 | t-Test | FDR correction BH | 0.0419776 | TRUE | QL | SN | 7.214976496 | 1.878149563 | Paired t-test |
| Sema1b~Q7KK54 | 15.3520112 | 0.000600317 | 15.68432581 | 12.83992391 | 0.564140204 | 0.255288109 | 2.8444019 | t-Test | FDR correction BH | 0.011237107 | TRUE | QL | SN | 7.182080879 | 3.221619314 | Paired t-test |
| LpR1~A8JRD1 | 18.52192262 | 0.000343458 | 15.58388211 | 12.74216127 | 0.3281149 | 0.30090366 | 2.841720839 | t-Test | FDR correction BH | 0.010357789 | TRUE | QL | SN | 7.168746312 | 3.464126142 | Paired t-test |
| ZnT35C~Q9V471 | 9.150219808 | 0.002759377 | 15.70044801 | 12.8651915 | 0.280037201 | 0.288239538 | 2.835256517 | t-Test | FDR correction BH | 0.018547205 | TRUE | QL | SN | 7.136696977 | 2.559188957 | Paired t-test |
| w~P10090 | 16.33629185 | 0.000499094 | 18.70544918 | 15.87238596 | 0.26292334 | 0.299948842 | 2.833063221 | t-Test | FDR correction BH | 0.010461282 | TRUE | QL | SN | 7.125855464 | 3.301817294 | Paired t-test |
| Mdr50~Q5BI62 | 7.350646658 | 0.005203428 | 16.58321965 | 13.77254221 | 0.711629994 | 0.766603969 | 2.810677446 | t-Test | FDR correction BH | 0.024608571 | TRUE | QL | SN | 7.016139563 | 2.28371049 | Paired t-test |
| Dysb~M9PI95 | 7.18723462 | 0.005550313 | 15.8537204 | 13.05840613 | 0.360327272 | 0.706046573 | 2.795314275 | t-Test | FDR correction BH | 0.025267898 | TRUE | QL | SN | 6.94182153 | 2.255682527 | Paired t-test |
| Rab8~O18338 | 16.24154666 | 0.000507801 | 15.73323493 | 12.94437969 | 0.242587293 | 0.357196174 | 2.788855239 | t-Test | FDR correction BH | 0.010505536 | TRUE | QL | SN | 6.91081203 | 3.294306822 | Paired t-test |
| Piezo~M9PCC7 | 55.18730003 | 1.31E-05 | 15.97880052 | 13.19697498 | 0.173621158 | 0.17029784 | 2.781825535 | t-Test | FDR correction BH | 0.003864721 | TRUE | QL | SN | 6.877220179 | 4.882559801 | Paired t-test |
| c11.1~Q9W350 | 5.832230697 | 0.010041793 | 18.53926021 | 15.76085266 | 0.324609936 | 0.584841537 | 2.778407558 | t-Test | FDR correction BH | 0.034898746 | TRUE | QL | SN | 6.86094622 | 1.998188741 | Paired t-test |
| Dmel\CG7600~Q9XZ12 | 7.448144611 | 0.005009998 | 15.8385093 | 13.0614828 | 0.555745351 | 0.430247938 | 2.777026499 | t-Test | FDR correction BH | 0.024556696 | TRUE | QL | SN | 6.854381538 | 2.300162476 | Paired t-test |
| Snapin~Q9VQF9 | 7.165740424 | 0.005598186 | 16.04664572 | 13.29532684 | 0.519537426 | 0.237352393 | 2.751318879 | t-Test | FDR correction BH | 0.025407152 | TRUE | QL | SN | 6.733323962 | 2.25195267 | Paired t-test |
| Tret1-1~A1Z8N1 | 17.62469184 | 0.000398195 | 21.08476763 | 18.34197912 | 0.110532205 | 0.19689844 | 2.742788515 | t-Test | FDR correction BH | 0.010461282 | TRUE | QL | SN | 6.693628649 | 3.399904272 | Paired t-test |
| Dmel\CG3662~Q2PDW1 | 10.25440034 | 0.001977278 | 16.64309041 | 13.91566491 | 0.529771726 | 0.899086158 | 2.727425496 | t-Test | FDR correction BH | 0.016665632 | TRUE | QL | SN | 6.622727491 | 2.703932172 | Paired t-test |
| Csp~M9PG83 | 72.26753414 | 5.84E-06 | 17.00005595 | 14.27330318 | 0.102118024 | 0.1034761 | 2.726752771 | t-Test | FDR correction BH | 0.003100532 | TRUE | QL | SN | 6.619640051 | 5.233658246 | Paired t-test |
| Syx7~Q7K0X9 | 22.18649635 | 0.000200464 | 15.8542494 | 13.14665488 | 0.205601396 | 0.179573057 | 2.70759452 | t-Test | FDR correction BH | 0.008188203 | TRUE | QL | SN | 6.532315714 | 3.697962549 | Paired t-test |
| Gli~Q7KT70 | 5.863410073 | 0.009892702 | 16.45263606 | 13.77869785 | 0.162706631 | 0.887728757 | 2.673938215 | t-Test | FDR correction BH | 0.034635327 | TRUE | QL | SN | 6.381688609 | 2.004685089 | Paired t-test |
| fab1~O96838 | 6.456969942 | 0.007535297 | 16.22653011 | 13.55468297 | 1.004472454 | 0.469741406 | 2.671847135 | t-Test | FDR correction BH | 0.029712198 | TRUE | QL | SN | 6.372445524 | 2.122899632 | Paired t-test |
| Rab32~A1Z7S3 | 26.71584154 | 0.000115074 | 19.28543542 | 16.62592965 | 0.341866254 | 0.247654828 | 2.659505774 | t-Test | FDR correction BH | 0.006717766 | TRUE | QL | SN | 6.318165702 | 3.939022031 | Paired t-test |
| Dmel\CG3106~Q9W322 | 3.611399099 | 0.036467652 | 17.11806941 | 14.46356606 | 0.266729953 | 1.125853421 | 2.654503349 | t-Test | FDR correction BH | 0.079590314 | FALSE | QL | SN | 6.296295925 | 1.438092194 | Paired t-test |
| Mul1~Q9VZJ9 | 6.284689335 | 0.008135545 | 16.45841114 | 13.81241945 | 0.454845555 | 1.004103203 | 2.645991685 | t-Test | FDR correction BH | 0.031389509 | TRUE | QL | SN | 6.25925818 | 2.089613322 | Paired t-test |
| Dmel\CG12531~M9PHM0 | 14.09434714 | 0.000773608 | 19.51383276 | 16.87115609 | 0.340921199 | 0.161501295 | 2.642676674 | t-Test | FDR correction BH | 0.012134178 | TRUE | QL | SN | 6.244892227 | 3.111479056 | Paired t-test |
| mio~Q9VQ89 | 13.315161 | 0.000915551 | 15.7645947 | 13.12978974 | 0.375212255 | 0.219114091 | 2.634804952 | t-Test | FDR correction BH | 0.012641521 | TRUE | QL | SN | 6.210911247 | 3.038317421 | Paired t-test |
| Dmel\CG5439~Q9VK31 | 29.39778094 | 8.64E-05 | 15.87164948 | 13.25753982 | 0.261553336 | 0.173201253 | 2.614109658 | t-Test | FDR correction BH | 0.006557185 | TRUE | QL | SN | 6.122452422 | 4.063279028 | Paired t-test |
| EG:22E5.5~O77266 | 5.759037977 | 0.010403534 | 16.11961995 | 13.51999261 | 0.518461015 | 0.942857841 | 2.599627341 | t-Test | FDR correction BH | 0.035517536 | TRUE | QL | SN | 6.061300386 | 1.982819119 | Paired t-test |
| Nckx30C~M9PCP1 | 7.516593171 | 0.004879835 | 15.548845 | 12.96294381 | 0.449096981 | 0.265225238 | 2.585901186 | t-Test | FDR correction BH | 0.024066801 | TRUE | QL | SN | 6.003905154 | 2.311594896 | Paired t-test |
| Dmel\CG4658~Q9VL57 | 5.777719875 | 0.0103096 | 15.85625485 | 13.27237755 | 0.336402119 | 0.668599604 | 2.583877303 | t-Test | FDR correction BH | 0.03547126 | TRUE | QL | SN | 5.995488487 | 1.986758174 | Paired t-test |
| CT18912~Q5BIG2 | 3.846293779 | 0.031017599 | 16.56459633 | 13.99727709 | 0.456585296 | 1.171553411 | 2.567319241 | t-Test | FDR correction BH | 0.072238356 | FALSE | QL | SN | 5.927070597 | 1.508391823 | Paired t-test |
| Vps35~Q7KVL7 | 5.482196933 | 0.01193676 | 15.8167257 | 13.25954779 | 0.271505083 | 0.591984264 | 2.557177913 | t-Test | FDR correction BH | 0.03880665 | TRUE | QL | SN | 5.885552746 | 1.923113544 | Paired t-test |

| | | | | | | | | | | | | | | | | |
|---|---|---|---|---|---|---|---|---|---|---|---|---|---|---|---|---|
| Usp32~M9PD06-2 | 6.073719609 | 0.008959195 | 15.54870991 | 13.00096049 | 0.705928928 | 0.216183862 | 2.54774942 | t-Test | FDR correction BH | 0.033357703 | TRUE | QL | SN | 5.847214111 | 2.047731027 | Paired t-test |
| Dmel\CG8814~Q9VQI6 | 4.66182661 | 0.018627867 | 15.68451084 | 13.13826798 | 0.478876862 | 0.75980305 | 2.546242861 | t-Test | FDR correction BH | 0.051863962 | FALSE | QL | SN | 5.841111243 | 1.729836866 | Paired t-test |
| PRL-1~O61722 | 7.42547076 | 0.005054123 | 17.44984371 | 14.90733314 | 0.792814085 | 0.674249115 | 2.542510577 | t-Test | FDR correction BH | 0.024608571 | TRUE | QL | SN | 5.826019691 | 2.296354156 | Paired t-test |
| nudE~A8JNQ0 | 14.13050707 | 0.000767754 | 15.87888746 | 13.34854147 | 0.492703715 | 0.7228448 | 2.530345993 | t-Test | FDR correction BH | 0.012134178 | TRUE | QL | SN | 5.777102106 | 3.114777739 | Paired t-test |
| RyR~Q24498-5 | 10.66009343 | 0.001764403 | 15.07574958 | 12.55811412 | 0.489775021 | 0.14726685 | 2.517635462 | t-Test | FDR correction BH | 0.016234815 | TRUE | QL | SN | 5.726427836 | 2.753402231 | Paired t-test |
| jp~Q7JV09 | 5.894438715 | 0.009747199 | 16.63142822 | 14.12901716 | 1.105298721 | 0.735214384 | 2.502411058 | t-Test | FDR correction BH | 0.034358997 | TRUE | QL | SN | 5.666315989 | 2.011120176 | Paired t-test |
| jus~Q8MQS2 | 10.56392232 | 0.001812006 | 17.15923656 | 14.66294185 | 0.243383174 | 0.488454606 | 2.496294719 | t-Test | FDR correction BH | 0.016234895 | TRUE | QL | SN | 5.642344362 | 2.741840276 | Paired t-test |
| Nep3~Q9W5Y0 | 8.326995915 | 0.003629986 | 15.62630045 | 13.1406494 | 0.449743064 | 0.152140537 | 2.485651046 | t-Test | FDR correction BH | 0.020315914 | TRUE | QL | SN | 5.600870401 | 2.440095002 | Paired t-test |
| Dmel\CG4552~Q9VPW9 | 10.59850878 | 0.001794693 | 15.63516356 | 13.15001067 | 0.177252428 | 0.503689653 | 2.485152893 | t-Test | FDR correction BH | 0.016234895 | TRUE | QL | SN | 5.59893679 | 2.74600993 | Paired t-test |
| Wwox~X2J7F1 | 24.64472217 | 0.000146464 | 15.88827486 | 13.41063749 | 0.340406453 | 0.430252353 | 2.477637375 | t-Test | FDR correction BH | 0.007209085 | TRUE | QL | SN | 5.569845754 | 3.834269239 | Paired t-test |
| Vps26~Q9W552 | 6.744940036 | 0.006655722 | 15.96645971 | 13.5233444 | 0.452989852 | 0.674267404 | 2.443115307 | t-Test | FDR correction BH | 0.027994359 | TRUE | QL | SN | 5.438147591 | 2.176804847 | Paired t-test |
| sky~Q0E8N9 | 3.184190468 | 0.049933101 | 16.33104638 | 13.89008832 | 0.405155675 | 1.538657954 | 2.440958059 | t-Test | FDR correction BH | 0.097252442 | FALSE | QL | SN | 5.430022058 | 1.301611459 | Paired t-test |
| EG:8D8.3~M9PDG9 | 5.921399456 | 0.009623042 | 15.41784501 | 12.98460007 | 0.455693964 | 0.858288193 | 2.433244934 | t-Test | FDR correction BH | 0.034358997 | TRUE | QL | SN | 5.401068829 | 2.016687629 | Paired t-test |
| Syx6~Q7JY00 | 11.48176838 | 0.001418111 | 15.15047179 | 12.72376602 | 0.146135745 | 0.331369213 | 2.426705763 | t-Test | FDR correction BH | 0.015251006 | TRUE | QL | SN | 5.3766433 | 2.848289792 | Paired t-test |
| CNT2~A1Z7N2 | 5.320072834 | 0.012973665 | 16.86883027 | 14.443826 | 0.276035799 | 0.588518452 | 2.425004266 | t-Test | FDR correction BH | 0.041416931 | TRUE | QL | SN | 5.370305892 | 1.886937316 | Paired t-test |
| numb~M9PCN6 | 7.399212684 | 0.005105867 | 17.19109943 | 14.77406807 | 0.202733846 | 0.388729517 | 2.41703136 | t-Test | FDR correction BH | 0.024608571 | TRUE | QL | SN | 5.340709305 | 2.291930467 | Paired t-test |
| mGluR~L0MPZ9 | 8.060589075 | 0.003988557 | 16.79467387 | 14.39116529 | 0.103908303 | 0.614449315 | 2.403508571 | t-Test | FDR correction BH | 0.021213554 | TRUE | QL | SN | 5.290883205 | 2.399184155 | Paired t-test |
| Vps33B~Q9VBR1 | 5.273945828 | 0.013290048 | 15.96255108 | 13.56723429 | 0.332315595 | 1.029147341 | 2.395316793 | t-Test | FDR correction BH | 0.042006046 | TRUE | QL | SN | 5.260926132 | 1.876473435 | Paired t-test |
| Syt1~X2J979 | 15.67029124 | 0.000564819 | 19.05269593 | 16.66099212 | 0.465283669 | 0.271171057 | 2.391703816 | t-Test | FDR correction BH | 0.010886227 | TRUE | QL | SN | 5.247767548 | 3.248091044 | Paired t-test |
| Dmel\CG13743~Q8MRD1 | 4.604050064 | 0.019266611 | 15.72397661 | 13.33494075 | 0.046785875 | 0.932731691 | 2.389035858 | t-Test | FDR correction BH | 0.053008137 | FALSE | QL | SN | 5.238071882 | 1.715194674 | Paired t-test |
| MSBP~Q9VXM4 | 41.35253229 | 3.11E-05 | 18.53122026 | 16.14567194 | 0.241379955 | 0.201140853 | 2.385548314 | t-Test | FDR correction BH | 0.003864721 | TRUE | QL | SN | 5.225424757 | 4.506949114 | Paired t-test |
| Dmel\CG43066~A0A0B4K7V4 | 16.49709643 | 0.000484767 | 18.88733037 | 16.50396338 | 0.17706499 | 0.106494376 | 2.383366998 | t-Test | FDR correction BH | 0.010461282 | TRUE | QL | SN | 5.217530027 | 3.31446691 | Paired t-test |
| CG11935~Q9VBX8 | 10.78477385 | 0.001705126 | 15.17043632 | 12.79222293 | 0.106572924 | 0.376025454 | 2.378213387 | t-Test | FDR correction BH | 0.016207074 | TRUE | QL | SN | 5.198925158 | 2.76824364 | Paired t-test |
| AAF46211~Q9W3V2 | 6.698913303 | 0.006786883 | 17.02872393 | 14.65539662 | 0.607435338 | 0.494213778 | 2.373327316 | t-Test | FDR correction BH | 0.028302368 | TRUE | QL | SN | 5.181347395 | 2.168329637 | Paired t-test |
| Dmel\CG1951~Q9VAR0 | 3.240211986 | 0.047843249 | 15.96632532 | 13.60070843 | 0.777305037 | 1.25044133 | 2.365616885 | t-Test | FDR correction BH | 0.094558679 | FALSE | QL | SN | 5.153729741 | 1.320179337 | Paired t-test |
| ICA69~Q9VP78 | 15.78632351 | 0.000552573 | 15.34706268 | 12.98350296 | 0.357885564 | 0.253858686 | 2.363559717 | t-Test | FDR correction BH | 0.010867262 | TRUE | QL | SN | 5.146386171 | 3.257610636 | Paired t-test |
| ClC-c~Q9VUY1 | 8.852376147 | 0.003038725 | 17.47902821 | 15.1543452 | 0.220340723 | 0.674008215 | 2.324683003 | t-Test | FDR correction BH | 0.019235452 | TRUE | QL | SN | 5.009556907 | 2.51730856 | Paired t-test |
| out~Q9V3F9 | 4.167519886 | 0.025143235 | 15.71243674 | 13.40339194 | 0.495598092 | 0.812445922 | 2.309044799 | t-Test | FDR correction BH | 0.062779269 | FALSE | QL | SN | 4.955548671 | 1.599578845 | Paired t-test |
| Dmel\CG13077~Q9VIT5 | 8.556775054 | 0.003354199 | 17.78459174 | 15.48040187 | 0.370195356 | 0.122214181 | 2.304189867 | t-Test | FDR correction BH | 0.019789776 | TRUE | QL | SN | 4.938900373 | 2.474411143 | Paired t-test |
| mgl~A8JTM7 | 6.426828971 | 0.007636007 | 16.44658378 | 14.14661913 | 0.129762251 | 0.581100554 | 2.299964644 | t-Test | FDR correction BH | 0.029887369 | TRUE | QL | SN | 4.924456968 | 2.117133674 | Paired t-test |
| Wdfy2~Q9VKW1 | 6.157029886 | 0.008621416 | 17.18113281 | 14.89161455 | 0.200380853 | 0.737352161 | 2.289518261 | t-Test | FDR correction BH | 0.032762891 | TRUE | QL | SN | 4.888928347 | 2.064421375 | Paired t-test |
| Ykt6~Q9W3M8 | 20.63685808 | 0.000248818 | 18.28499899 | 16.00192582 | 0.45634922 | 0.295701176 | 2.28307317 | t-Test | FDR correction BH | 0.009437294 | TRUE | QL | SN | 4.867136278 | 3.60411898 | Paired t-test |
| Dmel\CG11857~A0A140SRF8 | 12.59910601 | 0.001078174 | 18.80280472 | 16.52055275 | 0.291019336 | 0.303740992 | 2.282251965 | t-Test | FDR correction BH | 0.01321178 | TRUE | QL | SN | 4.864366615 | 2.967311096 | Paired t-test |
| mmd~M9NDY9 | 9.11291226 | 0.002792453 | 15.91981478 | 13.65084326 | 0.410957314 | 0.392387256 | 2.268971522 | t-Test | FDR correction BH | 0.018612459 | TRUE | QL | SN | 4.819794118 | 2.55401413 | Paired t-test |
| Klc~P46824 | 16.02623416 | 0.000528351 | 18.9427771 | 16.68131579 | 0.166194449 | 0.114886616 | 2.261461311 | t-Test | FDR correction BH | 0.010653957 | TRUE | QL | SN | 4.794768998 | 3.277077726 | Paired t-test |
| Gorab~Q8IQQ4 | 4.338333792 | 0.022599423 | 16.4071767 | 14.15499196 | 0.24236814 | 0.919008972 | 2.252184737 | t-Test | FDR correction BH | 0.058633357 | FALSE | QL | SN | 4.764037392 | 1.645902646 | Paired t-test |
| Vps24~Q9VN02 | 4.554926927 | 0.019832213 | 16.01074194 | 13.75951742 | 0.551482568 | 1.023427842 | 2.251224517 | t-Test | FDR correction BH | 0.053431096 | FALSE | QL | SN | 4.76086763 | 1.702628827 | Paired t-test |
| red~Q9I7J4 | 8.448561951 | 0.003480495 | 15.97376079 | 13.72877531 | 0.585531967 | 0.148943148 | 2.244985476 | t-Test | FDR correction BH | 0.020309262 | TRUE | QL | SN | 4.740323362 | 2.458358981 | Paired t-test |
| Mitofilin~A0A0B4KHL7 | 3.626344292 | 0.036086017 | 15.84918509 | 13.61560043 | 0.227779219 | 1.257402288 | 2.233584658 | t-Test | FDR correction BH | 0.079118678 | FALSE | QL | SN | 4.703010843 | 1.442661055 | Paired t-test |
| para~X2JFE8 | 4.500697696 | 0.020481876 | 14.87684933 | 12.66284533 | 0.722160671 | 0.341683044 | 2.214003998 | t-Test | FDR correction BH | 0.054652643 | FALSE | QL | SN | 4.639611474 | 1.688630277 | Paired t-test |
| CG13600~Q7KS34 | 96.41155249 | 2.46E-06 | 15.74954827 | 13.53604045 | 0.386952721 | 0.421506218 | 2.213507818 | t-Test | FDR correction BH | 0.003100532 | TRUE | QL | SN | 4.638016066 | 5.609084658 | Paired t-test |
| Pi4KIIalpha~Q9VND3 | 16.94177347 | 0.000447892 | 19.78721222 | 17.57720893 | 0.077040382 | 0.20577015 | 2.210003295 | t-Test | FDR correction BH | 0.010461282 | TRUE | QL | SN | 4.626763304 | 3.348826352 | Paired t-test |
| Cog1~A0A0B4KH76 | 9.43049977 | 0.002526738 | 15.39773343 | 13.18954259 | 0.239164439 | 0.445183082 | 2.208190838 | t-Test | FDR correction BH | 0.017810152 | TRUE | QL | SN | 4.620954353 | 2.597439715 | Paired t-test |
| Ent2~Q9VMB6 | 3.763824032 | 0.032804052 | 16.54451854 | 14.34565566 | 0.287378336 | 1.228736259 | 2.198862882 | t-Test | FDR correction BH | 0.0752763 | FALSE | QL | SN | 4.591173277 | 1.484072513 | Paired t-test |
| Dmel\CG40006~Q8SYC3 | 4.788236984 | 0.01732338 | 15.1538972 | 12.95583171 | 0.315786929 | 0.527380885 | 2.198065493 | t-Test | FDR correction BH | 0.049366983 | TRUE | QL | SN | 4.588636399 | 1.761367378 | Paired t-test |
| bw~A0A0B4LHB3 | 13.66666735 | 0.000847569 | 15.16034235 | 12.96794972 | 0.385702414 | 0.313120693 | 2.192392625 | t-Test | FDR correction BH | 0.012555335 | TRUE | QL | SN | 4.5706287 | 3.07182509 | Paired t-test |
| GM130~Q9W289 | 12.82641765 | 0.001022662 | 19.51678237 | 17.32894703 | 0.553659451 | 0.280585945 | 2.187835342 | t-Test | FDR correction BH | 0.013062707 | TRUE | QL | SN | 4.556213469 | 2.990267977 | Paired t-test |
| Shaw~M9PEE2 | 3.814427051 | 0.031692735 | 23.59823536 | 21.41647736 | 0.558094469 | 0.829155235 | 2.181757993 | t-Test | FDR correction BH | 0.073444505 | FALSE | QL | SN | 4.537060801 | 1.49904028 | Paired t-test |
| nmd~Q9VL02 | 4.467677727 | 0.020891005 | 15.6811588 | 13.49975997 | 0.303156171 | 0.640332029 | 2.181398834 | t-Test | FDR correction BH | 0.055465618 | FALSE | QL | SN | 4.535931439 | 1.680040669 | Paired t-test |
| Q9VA71~Q9VA71 | 9.809824409 | 0.002251513 | 15.44610769 | 13.26582851 | 0.192983279 | 0.217467025 | 2.180279179 | t-Test | FDR correction BH | 0.016909887 | TRUE | QL | SN | 4.532412534 | 2.647525569 | Paired t-test |
| Spase25~B4F5U4 | 5.672118988 | 0.010855621 | 15.7340304 | 13.5574365 | 0.267594715 | 0.757397214 | 2.176593904 | t-Test | FDR correction BH | 0.036454821 | TRUE | QL | SN | 4.520849541 | 1.964345336 | Paired t-test |
| Rgk3~A8DYK6 | 3.202373972 | 0.049242349 | 15.73251477 | 13.55646842 | 1.503573938 | 0.36795832 | 2.176046353 | t-Test | FDR correction BH | 0.096367398 | FALSE | QL | SN | 4.519134053 | 1.307661238 | Paired t-test |
| Flo2~A4V4F2 | 3.324908225 | 0.044889324 | 16.36063272 | 14.19402146 | 0.223551412 | 1.348254594 | 2.166611263 | t-Test | FDR correction BH | 0.090402899 | FALSE | QL | SN | 4.489675774 | 1.347856936 | Paired t-test |
| Dmel\CG7956~E1JIS1 | 5.544797754 | 0.011565437 | 17.05206903 | 14.89636364 | 0.542319935 | 0.837532469 | 2.155705386 | t-Test | FDR correction BH | 0.037987094 | TRUE | QL | SN | 4.455864574 | 1.936837964 | Paired t-test |
| Dmel\CG6834~Q9VGJ7 | 19.22131552 | 0.000307543 | 14.98906864 | 12.83338056 | 0.085650688 | 0.223385161 | 2.15568808 | t-Test | FDR correction BH | 0.010357789 | TRUE | QL | SN | 4.455811125 | 3.512094219 | Paired t-test |
| mew~X2JDJ5 | 7.154856302 | 0.005622635 | 17.99876694 | 15.84804124 | 0.049791445 | 0.534523402 | 2.150725702 | t-Test | FDR correction BH | 0.02544562 | TRUE | QL | SN | 4.440510984 | 2.25006007 | Paired t-test |
| Vha100-1~Q8IML5 | 5.322495729 | 0.01295732 | 15.84093347 | 13.69215605 | 0.365664548 | 0.534031345 | 2.148777419 | t-Test | FDR correction BH | 0.041416931 | TRUE | QL | SN | 4.434518359 | 1.88748482 | Paired t-test |
| Dmel\CG5916~Q9VEY4 | 28.73805096 | 9.25E-05 | 15.15764128 | 13.0102491 | 0.086811247 | 0.114207754 | 2.147392176 | t-Test | FDR correction BH | 0.006567284 | TRUE | QL | SN | 4.430262476 | 4.033791045 | Paired t-test |
| Dmel\CG14232~Q9VWD5 | 10.81045192 | 0.001693247 | 15.43884187 | 13.31567262 | 0.068705303 | 0.35033557 | 2.123169246 | t-Test | FDR correction BH | 0.016207074 | TRUE | QL | SN | 4.356499103 | 2.771279788 | Paired t-test |
| wdp~Q9W266 | 10.42521665 | 0.001883711 | 16.97854654 | 14.85651767 | 0.287673379 | 0.334774314 | 2.122028874 | t-Test | FDR correction BH | 0.01648088 | TRUE | QL | SN | 4.353056887 | 2.724985835 | Paired t-test |
| Dmel\CG2930~Q9W4P6 | 18.04310959 | 0.000371326 | 15.57422984 | 13.45928797 | 0.080930175 | 0.226807495 | 2.114941864 | t-Test | FDR correction BH | 0.010461282 | TRUE | QL | SN | 4.331725624 | 3.430245138 | Paired t-test |
| Tango5~Q9W2S1 | 4.522563892 | 0.020216648 | 16.19755232 | 14.08353008 | 0.182737225 | 0.94342187 | 2.114022234 | t-Test | FDR correction BH | 0.054126252 | FALSE | QL | SN | 4.328965294 | 1.694290852 | Paired t-test |
| Wdr24~Q9XZ25 | 8.353160999 | 0.003597104 | 15.09419639 | 12.98600468 | 0.346750877 | 0.123014361 | 2.108191709 | t-Test | FDR correction BH | 0.020315914 | TRUE | QL | SN | 4.311505465 | 2.444046972 | Paired t-test |
| Dmel\CG10132~Q9VIU8 | 9.795712433 | 0.002261021 | 18.20711357 | 16.1037146 | 0.300248694 | 0.407287777 | 2.103398976 | t-Test | FDR correction BH | 0.016909887 | TRUE | QL | SN | 4.29720611 | 2.645695456 | Paired t-test |
| Dmel\CG10804~B7Z122 | 9.934799015 | 0.00216958 | 17.42359508 | 15.32595327 | 0.394481937 | 0.232845215 | 2.09764181 | t-Test | FDR correction BH | 0.016678022 | TRUE | QL | SN | 4.280092007 | 2.663624393 | Paired t-test |
| Arfip~Q9VGQ8 | 26.60616256 | 0.000116498 | 15.61933028 | 13.52795256 | 0.485634917 | 0.483196849 | 2.091377717 | t-Test | FDR correction BH | 0.006717766 | TRUE | QL | SN | 4.261548398 | 3.933680201 | Paired t-test |
| Dmel\CG1695~Q8IQ31 | 5.570185437 | 0.011419156 | 15.35410124 | 13.27448174 | 0.310951714 | 0.38132018 | 2.079619492 | t-Test | FDR correction BH | 0.037584122 | TRUE | QL | SN | 4.226957164 | 1.942366006 | Paired t-test |
| CG40016~M9PGG8 | 14.50941699 | 0.000709813 | 15.21394122 | 13.13489848 | 0.160834681 | 0.191316717 | 2.079042738 | t-Test | FDR correction BH | 0.012134178 | TRUE | QL | SN | 4.225267669 | 3.148856251 | Paired t-test |
| DCTN3-p24~Q9W1V8 | 3.45507428 | 0.040785878 | 16.06524433 | 13.98784173 | 0.686456677 | 1.451629408 | 2.077402609 | t-Test | FDR correction BH | 0.084890988 | FALSE | QL | SN | 4.220466898 | 1.38949018 | Paired t-test |
| CD98hc~Q9VHX9 | 9.670123208 | 0.002348026 | 17.81149012 | 15.73537316 | 0.173161961 | 0.334829027 | 2.076116966 | t-Test | FDR correction BH | 0.017164751 | TRUE | QL | SN | 4.216707546 | 2.629297023 | Paired t-test |
| Dmel\CG6420~Q9VBC4 | 8.765299324 | 0.003127421 | 15.07859352 | 13.00297022 | 0.563937982 | 0.433832751 | 2.075623305 | t-Test | FDR correction BH | 0.019235452 | TRUE | QL | SN | 4.215264923 | 2.504813608 | Paired t-test |
| FBgn0259166~Q9VY47 | 13.75417392 | 0.000831695 | 15.02813943 | 12.95264748 | 0.456006434 | 0.503688291 | 2.07549195 | t-Test | FDR correction BH | 0.012498956 | TRUE | QL | SN | 4.214881147 | 3.080036154 | Paired t-test |
| if~P12080-2 | 5.315254451 | 0.013006251 | 15.20914459 | 13.13856147 | 0.923468569 | 0.268172588 | 2.070583111 | t-Test | FDR correction BH | 0.041437915 | TRUE | QL | SN | 4.200564183 | 1.885847872 | Paired t-test |
| Ctl1~M9PHD2 | 8.656074909 | 0.003243586 | 15.15861272 | 13.09177502 | 0.033231847 | 0.409900414 | 2.066837701 | t-Test | FDR correction BH | 0.019572092 | TRUE | QL | SN | 4.189673155 | 2.488974606 | Paired t-test |
| Sema1a~M9PB77 | 6.924476422 | 0.006174862 | 15.29410961 | 13.23849347 | 0.67543769 | 0.3746071 | 2.055616142 | t-Test | FDR correction BH | 0.026875835 | TRUE | QL | SN | 4.157211484 | 2.209372722 | Paired t-test |
| Gos28~Q9VE50 | 17.21207561 | 0.000427286 | 19.00309115 | 16.94766363 | 0.090249318 | 0.128300858 | 2.055427521 | t-Test | FDR correction BH | 0.010461282 | TRUE | QL | SN | 4.156667997 | 3.369281363 | Paired t-test |
| bchs~Q9VML2 | 3.464762974 | 0.040499886 | 15.12188548 | 13.06836193 | 0.697038253 | 0.406444803 | 2.053523552 | t-Test | FDR correction BH | 0.084778341 | FALSE | QL | SN | 4.151185934 | 1.392546204 | Paired t-test |
| Dmel\CG18745~Q9I7L0 | 6.501152606 | 0.007390802 | 14.72057751 | 12.67130386 | 0.221959597 | 0.474954696 | 2.049273656 | t-Test | FDR correction BH | 0.029507637 | TRUE | QL | SN | 4.138975349 | 2.131308453 | Paired t-test |
| Pex3~Q9VUL8 | 16.97192241 | 0.000445529 | 17.43091053 | 15.38224069 | 0.390357442 | 0.184505244 | 2.048669835 | t-Test | FDR correction BH | 0.010461282 | TRUE | QL | SN | 4.137243396 | 3.351123706 | Paired t-test |
| OtopLa~M9PGH2 | 11.63598946 | 0.001363429 | 18.01222652 | 15.96935512 | 0.363591415 | 0.564230921 | 2.042871403 | t-Test | FDR correction BH | 0.014978909 | TRUE | QL | SN | 4.120648496 | 2.86536759 | Paired t-test |
| Snx16~Q7JR96 | 5.838600777 | 0.010011095 | 17.4718532 | 15.43565627 | 0.726208274 | 0.136058668 | 2.036196928 | t-Test | FDR correction BH | 0.034896442 | TRUE | QL | SN | 4.101628787 | 1.99951843 | Paired t-test |
| fra~A1Z921 | 7.267164845 | 0.005376951 | 16.41662796 | 14.38337233 | 0.248458145 | 0.400136006 | 2.033255627 | t-Test | FDR correction BH | 0.024813171 | TRUE | QL | SN | 4.09327509 | 2.26946393 | Paired t-test |
| Vps2~Q9VBI3 | 5.166250541 | 0.01406894 | 15.16552193 | 13.14211826 | 0.435494846 | 0.556097963 | 2.023403672 | t-Test | FDR correction BH | 0.04334975 | TRUE | QL | SN | 4.065417934 | 1.851738638 | Paired t-test |
| iPLA2-VIA~Q9VT60 | 7.805309135 | 0.004377386 | 17.56789298 | 15.55178033 | 0.590079502 | 0.355998309 | 2.016112647 | t-Test | FDR correction BH | 0.022531262 | TRUE | QL | SN | 4.044924142 | 2.358785121 | Paired t-test |
| Dmel\CG17660~Q9VQ34 | 11.97653488 | 0.001252228 | 18.86870444 | 16.86251051 | 0.061324019 | 0.293225587 | 2.006193931 | t-Test | FDR correction BH | 0.014351069 | TRUE | QL | SN | 4.017210141 | 2.902316714 | Paired t-test |
| Grasp65~Q9VW57 | 8.797407708 | 0.003094322 | 19.22088998 | 17.22943674 | 0.45800279 | 0.281745173 | 1.991453246 | t-Test | FDR correction BH | 0.019235452 | TRUE | QL | SN | 3.976373418 | 2.509434483 | Paired t-test |
| Muted~A0A0B4LHG2 | 45.18712908 | 2.39E-05 | 16.2576661 | 14.26779023 | 0.397843895 | 0.369858545 | 1.989875866 | t-Test | FDR correction BH | 0.003864721 | TRUE | QL | SN | 3.972028201 | 4.622338671 | Paired t-test |
| l(2)01289~Q9V9A9 | 5.700481382 | 0.010705322 | 14.92127092 | 12.93575569 | 0.730089617 | 0.641436165 | 1.985515225 | t-Test | FDR correction BH | 0.036439268 | TRUE | QL | SN | 3.96004061 | 1.970400279 | Paired t-test |
| cn~A1Z746 | 3.654324506 | 0.035385141 | 15.56035008 | 13.57589444 | 0.413376638 | 1.075751952 | 1.984455646 | t-Test | FDR correction BH | 0.078398494 | FALSE | QL | SN | 3.95713325 | 1.451179072 | Paired t-test |
| Dmel\CG11377~M9PDP6 | 7.138072613 | 0.005660614 | 15.22614607 | 13.24241022 | 0.143222217 | 0.392620133 | 1.983735846 | t-Test | FDR correction BH | 0.025472763 | TRUE | QL | SN | 3.955159419 | 2.247136464 | Paired t-test |
| plx~A0A0B4KG76 | 8.541790989 | 0.003371322 | 16.97827077 | 15.00638438 | 0.467399049 | 0.265695988 | 1.971886389 | t-Test | FDR correction BH | 0.0198174 | TRUE | QL | SN | 3.922807087 | 2.472199811 | Paired t-test |
| Blos4~M9PET7 | 26.4866174 | 0.000118077 | 16.44425422 | 14.48081755 | 0.152752028 | 0.118345182 | 1.963436667 | t-Test | FDR correction BH | 0.006717766 | TRUE | QL | SN | 3.899898744 | 3.927832877 | Paired t-test |
| MRP~Q7KTC0 | 3.976565552 | 0.028442204 | 15.53291115 | 13.57942538 | 0.214979968 | 0.957720745 | 1.953485779 | t-Test | FDR correction BH | 0.067928682 | FALSE | QL | SN | 3.873092021 | 1.546036751 | Paired t-test |
| Q9I7L1_DROME~Q9I7L1 | 8.809621497 | 0.003081853 | 15.15111443 | 13.19844226 | 0.481761119 | 0.468982686 | 1.952672179 | t-Test | FDR correction BH | 0.019235452 | TRUE | QL | SN | 3.870908428 | 2.511188066 | Paired t-test |
| CG15639~Q9VK03 | 41.16875842 | 3.15E-05 | 14.79988273 | 12.85448615 | 0.288219147 | 0.273791886 | 1.945396574 | t-Test | FDR correction BH | 0.003864721 | TRUE | QL | SN | 3.851436327 | 4.501154262 | Paired t-test |
| Rbcn-3B~Q9W543 | 12.5092394 | 0.001101224 | 17.74682761 | 15.81047821 | 0.302261122 | 0.140111888 | 1.936349404 | t-Test | FDR correction BH | 0.013258226 | TRUE | QL | SN | 3.827359463 | 2.958124139 | Paired t-test |
| EP(3)3622~X2JCD3 | 3.850663117 | 0.030926473 | 15.47686362 | 13.55560342 | 0.336720863 | 1.151914707 | 1.921260205 | t-Test | FDR correction BH | 0.072131584 | FALSE | QL | SN | 3.787537584 | 1.509669604 | Paired t-test |
| unc-104~A0A126GUN3 | 6.980313021 | 0.006034681 | 14.84414624 | 12.93215053 | 0.19756645 | 0.302387033 | 1.911995709 | t-Test | FDR correction BH | 0.026555929 | TRUE | QL | SN | 3.763293237 | 2.219345699 | Paired t-test |
| Rac1~M9PBH7 | 4.110811387 | 0.026068964 | 14.8978359 | 12.98614888 | 0.137543366 | 0.786394528 | 1.911687021 | t-Test | FDR correction BH | 0.064086203 | FALSE | QL | SN | 3.762488107 | 1.583876226 | Paired t-test |
| Nup44A~Q7K2X8 | 3.310596722 | 0.045371895 | 16.6299931 | 14.73479863 | 0.533036017 | 1.015531839 | 1.895194467 | t-Test | FDR correction BH | 0.091029507 | FALSE | QL | SN | 3.719721134 | 1.343213084 | Paired t-test |
| TM4SF~O76137 | 15.29786964 | 0.000606649 | 18.01855017 | 16.12541362 | 0.267457929 | 0.46807803 | 1.893136557 | t-Test | FDR correction BH | 0.011237107 | TRUE | QL | SN | 3.714418978 | 3.217062824 | Paired t-test |
| Rab3-GEF~A8JUX2 | 5.399004484 | 0.012454731 | 16.25507984 | 14.3693143 | 0.194983975 | 0.752577909 | 1.885765537 | t-Test | FDR correction BH | 0.040238085 | TRUE | QL | SN | 3.695489661 | 1.904665654 | Paired t-test |
| GluClalpha~E1JIQ1 | 13.00579323 | 0.000981507 | 16.73632104 | 14.86315838 | 0.555902442 | 0.415246381 | 1.873162662 | t-Test | FDR correction BH | 0.012817077 | TRUE | QL | SN | 3.663347759 | 3.008106388 | Paired t-test |
| Ccz1~Q9VZL5 | 3.515416213 | 0.039045563 | 14.87325012 | 13.00617596 | 0.644806109 | 0.322561403 | 1.867074161 | t-Test | FDR correction BH | 0.08299724 | FALSE | QL | SN | 3.647920176 | 1.408428314 | Paired t-test |
| Dmel\CG4829~A8JV25 | 7.650226463 | 0.0046383 | 14.96156553 | 13.09847571 | 0.640320433 | 0.240925871 | 1.863089826 | t-Test | FDR correction BH | 0.023308553 | TRUE | QL | SN | 3.637859504 | 2.333641198 | Paired t-test |
| mge~Q9VZL1 | 6.038076289 | 0.009109015 | 17.11792329 | 15.26012775 | 0.371971986 | 0.612820409 | 1.857795548 | t-Test | FDR correction BH | 0.033357703 | TRUE | QL | SN | 3.624534065 | 2.040528581 | Paired t-test |
| Dmel\CG9590~Q9VF39 | 9.978346302 | 0.00214196 | 19.35269119 | 17.50032827 | 0.207001501 | 0.252780825 | 1.852362917 | t-Test | FDR correction BH | 0.016678022 | TRUE | QL | SN | 3.610911137 | 2.669188717 | Paired t-test |
| TrpRS~Q0KI98 | 4.608132839 | 0.019220549 | 15.6651595 | 13.81424848 | 0.229683361 | 0.857445787 | 1.850911024 | t-Test | FDR correction BH | 0.052972898 | FALSE | QL | SN | 3.607279032 | 1.716234208 | Paired t-test |
| Tsc1~Q9VCC9 | 23.62339871 | 0.000166207 | 15.26467228 | 13.43539012 | 0.380462287 | 0.297207829 | 1.829282155 | t-Test | FDR correction BH | 0.007209085 | TRUE | QL | SN | 3.553602112 | 3.779350892 | Paired t-test |
| anchor~M9PFK5 | 6.993487409 | 0.006002218 | 14.86305816 | 13.05549845 | 0.23908756 | 0.246582601 | 1.807559715 | t-Test | FDR correction BH | 0.026555929 | TRUE | QL | SN | 3.500496867 | 2.221688221 | Paired t-test |
| Hk~Q9W2W9 | 10.46039255 | 0.001865175 | 18.66548655 | 16.86013669 | 0.311797601 | 0.162401675 | 1.805349861 | t-Test | FDR correction BH | 0.016415603 | TRUE | QL | SN | 3.49513907 | 2.72928037 | Paired t-test |
| gish~A0A0B4KHL4 | 9.298280517 | 0.002633108 | 18.20005792 | 16.39849001 | 0.195793718 | 0.424237171 | 1.801567906 | t-Test | FDR correction BH | 0.018079919 | TRUE | QL | SN | 3.485988731 | 2.57953132 | Paired t-test |
| didum~A1Z6Z8 | 9.325337767 | 0.002610865 | 15.46935026 | 13.68506924 | 0.154655463 | 0.179279238 | 1.784281013 | t-Test | FDR correction BH | 0.018079919 | TRUE | QL | SN | 3.444467612 | 2.583215622 | Paired t-test |
| GlcAT-P~A4V1U5 | 11.30371569 | 0.001484924 | 16.75276452 | 14.979135 | 0.402882084 | 0.342797515 | 1.773629518 | t-Test | FDR correction BH | 0.015562398 | TRUE | QL | SN | 3.419130571 | 2.828295657 | Paired t-test |
| Vps11~Q8SX81 | 12.14749495 | 0.001200924 | 15.21283287 | 13.44270284 | 0.37912782 | 0.443468144 | 1.770130036 | t-Test | FDR correction BH | 0.013862839 | TRUE | QL | SN | 3.410846987 | 2.920484507 | Paired t-test |
| santa-maria~Q9VM10 | 5.924097181 | 0.009610733 | 15.02833857 | 13.2635804 | 0.342463743 | 0.646045114 | 1.764758172 | t-Test | FDR correction BH | 0.034358997 | TRUE | QL | SN | 3.39817034 | 2.017243494 | Paired t-test |
| Ost48~Q24319 | 4.700059868 | 0.018220242 | 14.72473842 | 12.97142512 | 0.668498931 | 0.351930156 | 1.753313294 | t-Test | FDR correction BH | 0.051351 | FALSE | QL | SN | 3.371319351 | 1.739445861 | Paired t-test |
| VhaAC45~Q7JR49 | 8.244095182 | 0.00373683 | 15.17159377 | 13.42200162 | 0.964641094 | 0.66090951 | 1.749592142 | t-Test | FDR correction BH | 0.02059782 | TRUE | QL | SN | 3.362634891 | 2.427496681 | Paired t-test |
| Dmel\CG11665~Q7JWI7 | 9.831168114 | 0.002237233 | 15.02734683 | 13.29134274 | 0.349026651 | 0.318882182 | 1.736004088 | t-Test | FDR correction BH | 0.016890579 | TRUE | QL | SN | 3.331112521 | 2.650288774 | Paired t-test |
| Alp12~Q9VIW9 | 19.75194472 | 0.000283562 | 14.71797171 | 12.99012883 | 0.166897113 | 0.191870258 | 1.727842888 | t-Test | FDR correction BH | 0.010038098 | TRUE | QL | SN | 3.312321907 | 3.547351823 | Paired t-test |
| CLLD6~Q9VED7 | 40.11054398 | 3.41E-05 | 17.22801359 | 15.50290191 | 0.209507845 | 0.255197898 | 1.725111687 | t-Test | FDR correction BH | 0.003879823 | TRUE | QL | SN | 3.306057203 | 4.46727584 | Paired t-test |
| DmSNX21~M9PBU4 | 10.0759538 | 0.002081723 | 14.83301209 | 13.10791185 | 0.371655113 | 0.232995455 | 1.725100243 | t-Test | FDR correction BH | 0.016678022 | TRUE | QL | SN | 3.306030979 | 2.681577029 | Paired t-test |
| Dmel\CG10681~M9PFF6 | 16.6793545 | 0.000469182 | 15.27510376 | 13.55081504 | 0.19703854 | 0.353815145 | 1.724288715 | t-Test | FDR correction BH | 0.010461282 | TRUE | QL | SN | 3.30417183 | 3.328658497 | Paired t-test |

| | | | | | | | | | | | | | | | | |
|---|---|---|---|---|---|---|---|---|---|---|---|---|---|---|---|---|
| DS01368.1~Q7KTB0 | 3.338530858 | 0.044436013 | 15.51632821 | 13.79785654 | 1.0022762 | 0.358909343 | 1.718471674 | t-Test | FDR correction BH | 0.08983067 | FALSE | QL | SN | 3.290876017 | 1.352264918 | Paired t-test |
| path~Q9VT04 | 9.153749349 | 0.002756275 | 18.79250909 | 17.08250709 | 0.07675574 | 0.258422298 | 1.710001997 | t-Test | FDR correction BH | 0.018547205 | TRUE | QL | SN | 3.271612762 | 2.559677498 | Paired t-test |
| Dmel\CG11811~Q9VTB3 | 6.554754959 | 0.007220352 | 15.95224623 | 14.24234367 | 0.082380672 | 0.464916694 | 1.70990256 | t-Test | FDR correction BH | 0.029045505 | TRUE | QL | SN | 3.271387275 | 2.141441653 | Paired t-test |
| CT39634~Q7KUK9 | 3.913846086 | 0.02964636 | 14.93655198 | 13.22946236 | 0.586425621 | 0.174877058 | 1.707089624 | t-Test | FDR correction BH | 0.069348974 | FALSE | QL | SN | 3.265015009 | 1.528028623 | Paired t-test |
| EG:33C11.3~A0A0S0X7W8 | 4.126461642 | 0.025809153 | 16.01085466 | 14.31073517 | 0.695135219 | 1.111077414 | 1.700119491 | t-Test | FDR correction BH | 0.06368161 | FALSE | QL | SN | 3.249278696 | 1.588226252 | Paired t-test |
| RSG7~Q9VXA3 | 4.737493809 | 0.017832295 | 14.85126345 | 13.15915435 | 0.725370249 | 0.170092628 | 1.692109106 | t-Test | FDR correction BH | 0.050545989 | FALSE | QL | SN | 3.231287473 | 1.748792762 | Paired t-test |
| OtopLa~Q9W4A6 | 6.851718184 | 0.006364035 | 18.71363958 | 17.04122411 | 0.485738737 | 0.350553457 | 1.672415472 | t-Test | FDR correction BH | 0.027492254 | TRUE | QL | SN | 3.187478193 | 2.196267454 | Paired t-test |
| Dmel\CG8379~Q9VHK5 | 57.03504101 | 1.19E-05 | 14.95586171 | 13.28438388 | 0.43968059 | 0.460981783 | 1.671477823 | t-Test | FDR correction BH | 0.003864721 | TRUE | QL | SN | 3.185407233 | 4.925434941 | Paired t-test |
| Sec22~O77434 | 8.669345189 | 0.003229171 | 18.09030876 | 16.42198283 | 0.647595745 | 0.332808928 | 1.668325926 | t-Test | FDR correction BH | 0.019559199 | TRUE | QL | SN | 3.178455579 | 2.49090896 | Paired t-test |
| ATP8A~A0A0B4JD16 | 12.5236465 | 0.001097485 | 14.82343411 | 13.16320689 | 0.276879589 | 0.157724491 | 1.660227222 | t-Test | FDR correction BH | 0.013258226 | TRUE | QL | SN | 3.160663008 | 2.959601261 | Paired t-test |
| Dromel_CG7071_FBtr0084211_mORF~Q8SXS( | 3.184041964 | 0.049938793 | 15.30629619 | 13.64881084 | 0.123500538 | 0.977031565 | 1.657485355 | t-Test | FDR correction BH | 0.097252442 | FALSE | QL | SN | 3.154661818 | 1.301561959 | Paired t-test |
| Dmel\CG5065~E1JH83 | 9.889733369 | 0.002198662 | 16.57074025 | 14.91593726 | 0.435989803 | 0.281644104 | 1.654802988 | t-Test | FDR correction BH | 0.016758223 | TRUE | QL | SN | 3.148801882 | 2.657841533 | Paired t-test |
| galectin~Q9VPI6 | 5.279028341 | 0.013254697 | 17.61077608 | 15.96222117 | 0.317032184 | 0.233539461 | 1.648554912 | t-Test | FDR correction BH | 0.0419776 | TRUE | QL | SN | 3.135194423 | 1.877630189 | Paired t-test |
| Vps16B~Q9VAG4 | 5.909283653 | 0.009678578 | 14.54296291 | 12.89897798 | 0.266365262 | 0.309646108 | 1.643984925 | t-Test | FDR correction BH | 0.034358997 | TRUE | QL | SN | 3.125278865 | 2.014188436 | Paired t-test |
| nrv3~Q7JS69 | 8.423600909 | 0.003510516 | 18.24926914 | 16.60806352 | 0.475124621 | 0.270329873 | 1.64120562 | t-Test | FDR correction BH | 0.020315914 | TRUE | QL | SN | 3.119263911 | 2.454628991 | Paired t-test |
| CT14766~Q9VDT3 | 4.581742666 | 0.019520828 | 16.02889367 | 14.40183644 | 0.627255182 | 0.372049692 | 1.627057229 | t-Test | FDR correction BH | 0.053339071 | FALSE | QL | SN | 3.088823058 | 1.709501771 | Paired t-test |
| Amph~A0A0B4KFR7 | 5.145336841 | 0.014227052 | 19.69135336 | 18.06867105 | 0.378655275 | 0.535458917 | 1.622682316 | t-Test | FDR correction BH | 0.043752305 | TRUE | QL | SN | 3.079470518 | 1.846885079 | Paired t-test |
| anne~L0MLL1 | 4.132326399 | 0.025712652 | 14.87215135 | 13.25194727 | 0.457599084 | 0.67929638 | 1.620204083 | t-Test | FDR correction BH | 0.06368161 | FALSE | QL | SN | 3.074185204 | 1.589853132 | Paired t-test |
| Rab11~O18335 | 4.572052863 | 0.019632609 | 17.45710648 | 15.83801118 | 0.527004505 | 0.180171612 | 1.619095302 | t-Test | FDR correction BH | 0.053431096 | FALSE | QL | SN | 3.071823452 | 1.707021978 | Paired t-test |
| tweek~M9MSI8 | 9.497752787 | 0.002474813 | 15.29249821 | 13.67499549 | 0.199227766 | 0.250551771 | 1.617502711 | t-Test | FDR correction BH | 0.017758453 | TRUE | QL | SN | 3.068434337 | 2.606457677 | Paired t-test |
| Dg~A1ZA89 | 8.323291488 | 0.003634674 | 16.79300196 | 15.17840423 | 0.321768058 | 0.376843369 | 1.614597731 | t-Test | FDR correction BH | 0.020315914 | TRUE | QL | SN | 3.06226202 | 2.439534549 | Paired t-test |
| Dmel\CG3860~Q9W1D9 | 7.746731441 | 0.004473604 | 16.65031928 | 15.04415122 | 0.539535753 | 0.659387563 | 1.606168066 | t-Test | FDR correction BH | 0.022695703 | TRUE | QL | SN | 3.0444214 | 2.349342484 | Paired t-test |
| norpA~X2JCI4 | 6.054630288 | 0.009039028 | 14.90234671 | 13.30164026 | 0.631072607 | 0.691960636 | 1.600706451 | t-Test | FDR correction BH | 0.033357703 | TRUE | QL | SN | 3.032917911 | 2.043878287 | Paired t-test |
| Rbcn-3A~Q9W425 | 15.94520154 | 0.000536371 | 18.16319036 | 16.56699957 | 0.108818274 | 0.107090748 | 1.596190794 | t-Test | FDR correction BH | 0.010680489 | TRUE | QL | SN | 3.023439674 | 3.270534662 | Paired t-test |
| Ocrl~O46094 | 13.33852834 | 0.000910811 | 16.44659194 | 14.85079695 | 0.168858105 | 0.038440991 | 1.595794989 | t-Test | FDR correction BH | 0.012641521 | TRUE | QL | SN | 3.022610304 | 3.040571515 | Paired t-test |
| Vps13~A1Z713 | 9.414732162 | 0.002539121 | 18.96565226 | 17.37220371 | 0.083191939 | 0.336360281 | 1.593448544 | t-Test | FDR correction BH | 0.017818589 | TRUE | QL | SN | 3.01769823 | 2.595316614 | Paired t-test |
| scramb2~Q9VZW1 | 3.932156935 | 0.029288144 | 14.37609463 | 12.7827887 | 0.565155921 | 0.36221126 | 1.59330593 | t-Test | FDR correction BH | 0.068933059 | FALSE | QL | SN | 3.01739994 | 1.533308152 | Paired t-test |
| Dmel\CG5110~M9NEQ5 | 12.31498973 | 0.001153332 | 20.59330973 | 19.00584062 | 0.51702624 | 0.380973962 | 1.587469115 | t-Test | FDR correction BH | 0.013609321 | TRUE | QL | SN | 3.005216888 | 2.938045546 | Paired t-test |
| Rac2~M9PBU2 | 8.625424058 | 0.003277207 | 15.89754528 | 14.33491964 | 0.020138372 | 0.30449081 | 1.562625636 | t-Test | FDR correction BH | 0.019626282 | TRUE | QL | SN | 2.953909519 | 2.484496101 | Paired t-test |
| Past1~Q8T6I0 | 17.71937732 | 0.000391894 | 20.90645242 | 19.34803501 | 0.084363111 | 0.089660324 | 1.55841741 | t-Test | FDR correction BH | 0.010461282 | TRUE | QL | SN | 2.945305756 | 3.406831878 | Paired t-test |
| Sbf~Q7KSP6 | 5.482628642 | 0.011934146 | 14.99330928 | 13.43497871 | 0.173773802 | 0.530950886 | 1.558330572 | t-Test | FDR correction BH | 0.03880665 | TRUE | QL | SN | 2.945128479 | 1.923208649 | Paired t-test |
| Cpes~A0A0B4LGC8 | 3.481206239 | 0.040020327 | 14.58838479 | 13.03044013 | 0.419575324 | 0.40547277 | 1.557944662 | t-Test | FDR correction BH | 0.084542764 | FALSE | QL | SN | 2.944340786 | 1.397719365 | Paired t-test |
| Pxt~Q9VEG6 | 4.742963961 | 0.017776507 | 15.30851226 | 13.75627392 | 0.115123457 | 0.631193663 | 1.55223834 | t-Test | FDR correction BH | 0.050477674 | FALSE | QL | SN | 2.932717974 | 1.750153576 | Paired t-test |
| CG8020~Q9VWJ3 | 8.988422669 | 0.002906694 | 15.62299028 | 14.07337338 | 0.350071114 | 0.30347868 | 1.549616901 | t-Test | FDR correction BH | 0.018976903 | TRUE | QL | SN | 2.927393936 | 2.536600619 | Paired t-test |
| Cisd2~Q9VAM6 | 11.4178445 | 0.00144163 | 20.6419076 | 19.10131476 | 0.200334396 | 0.244107494 | 1.540592847 | t-Test | FDR correction BH | 0.015251006 | TRUE | QL | SN | 2.909140242 | 2.841146174 | Paired t-test |
| CG15684~Q8IN56 | 8.401239991 | 0.003537702 | 14.93701184 | 13.39750286 | 0.320010913 | 0.594392466 | 1.539508976 | t-Test | FDR correction BH | 0.020315914 | TRUE | QL | SN | 2.906955478 | 2.451278755 | Paired t-test |
| Dmel\CG32039~Q8IQC6 | 5.344896583 | 0.012807466 | 14.78964526 | 13.25118786 | 0.399537234 | 0.589552069 | 1.538457398 | t-Test | FDR correction BH | 0.041050893 | TRUE | QL | SN | 2.904837375 | 1.892536777 | Paired t-test |
| nrv1~B5RIR1 | 6.075595204 | 0.008951401 | 18.80047268 | 17.26832128 | 0.32605007 | 0.450204109 | 1.532151401 | t-Test | FDR correction BH | 0.033357703 | TRUE | QL | SN | 2.892168086 | 2.048108997 | Paired t-test |
| CadN~Q4ABE9 | 3.825259028 | 0.03146115 | 14.74945525 | 13.21864019 | 0.272173473 | 0.66454561 | 1.530815064 | t-Test | FDR correction BH | 0.073164397 | FALSE | QL | SN | 2.889490375 | 1.502225408 | Paired t-test |
| Syx1A~A0A0B4JCZ4 | 9.099276581 | 0.002804673 | 16.18593506 | 14.65603176 | 0.450698184 | 0.348229388 | 1.529903298 | t-Test | FDR correction BH | 0.018616017 | TRUE | QL | SN | 2.887664827 | 2.552117768 | Paired t-test |
| Snap25~P36975 | 13.50610025 | 0.00087776 | 20.17901787 | 18.65636499 | 0.130109247 | 0.177375093 | 1.522652875 | t-Test | FDR correction BH | 0.012597039 | TRUE | QL | SN | 2.873188956 | 3.056624318 | Paired t-test |
| Sply~Q9V7Y2 | 3.269478224 | 0.046795445 | 16.01766491 | 14.50260684 | 0.347677188 | 1.033636852 | 1.515058068 | t-Test | FDR correction BH | 0.092783534 | FALSE | QL | SN | 2.858103316 | 1.32979642 | Paired t-test |
| Tom20~Q95RF6 | 10.02120721 | 0.002115228 | 18.32550744 | 16.81327718 | 0.390305542 | 0.133285926 | 1.512230264 | t-Test | FDR correction BH | 0.016678022 | TRUE | QL | SN | 2.852506679 | 2.674642779 | Paired t-test |
| Mfe2~X2JFD6 | 5.677958123 | 0.010824453 | 15.75222933 | 14.26699056 | 0.379700609 | 0.382685487 | 1.48523877 | t-Test | FDR correction BH | 0.036454821 | TRUE | QL | SN | 2.79963504 | 1.965594041 | Paired t-test |
| uif~Q9VM55 | 4.219400663 | 0.024333149 | 17.14512281 | 15.66234801 | 0.700739993 | 0.362752988 | 1.482774798 | t-Test | FDR correction BH | 0.061625924 | FALSE | QL | SN | 2.794857637 | 1.613801686 | Paired t-test |
| Cyp9b2~Q9V4I1 | 7.098615632 | 0.005751239 | 15.14354607 | 13.67729243 | 0.273741605 | 0.568885625 | 1.466253641 | t-Test | FDR correction BH | 0.02573518 | TRUE | QL | SN | 2.763034624 | 2.240238572 | Paired t-test |
| OstDelta~A0A0B4LFR4 | 5.015754717 | 0.015260233 | 14.50430313 | 13.04188959 | 0.191251124 | 0.416020121 | 1.462413532 | t-Test | FDR correction BH | 0.045856543 | TRUE | QL | SN | 2.755689867 | 1.816438827 | Paired t-test |
| Rip11~X2JFP6 | 7.842699761 | 0.004317395 | 15.54915023 | 14.08925133 | 0.280732413 | 0.324462975 | 1.459898896 | t-Test | FDR correction BH | 0.022329903 | TRUE | QL | SN | 2.750890848 | 2.364778215 | Paired t-test |
| mys~P11584 | 38.28631882 | 3.92E-05 | 19.00729981 | 17.54959575 | 0.089083689 | 0.106305682 | 1.457704061 | t-Test | FDR correction BH | 0.003902745 | TRUE | QL | SN | 2.746708979 | 4.406725656 | Paired t-test |
| Dmel\CG6701~A1Z9K0 | 4.821168919 | 0.01700317 | 14.90476713 | 13.44887224 | 0.104451194 | 0.608364665 | 1.455894892 | t-Test | FDR correction BH | 0.048803692 | TRUE | QL | SN | 2.743266708 | 1.769470112 | Paired t-test |
| Dmel\CG8004~Q9VW58 | 7.301538419 | 0.005304592 | 19.24845332 | 17.79662244 | 0.635190357 | 0.323235762 | 1.451830877 | t-Test | FDR correction BH | 0.024737406 | TRUE | QL | SN | 2.735549908 | 2.275347989 | Paired t-test |
| Syx5~A4V0S1 | 8.385201393 | 0.003557373 | 17.75691317 | 16.3061829 | 0.164569621 | 0.245229705 | 1.45073027 | t-Test | FDR correction BH | 0.020315914 | TRUE | QL | SN | 2.733463801 | 2.448870625 | Paired t-test |
| Pex1~Q9VUC7 | 4.1842607 | 0.024878084 | 14.92054048 | 13.47132349 | 0.172201033 | 0.646818921 | 1.44921699 | t-Test | FDR correction BH | 0.062509129 | FALSE | QL | SN | 2.730598102 | 1.60418307 | Paired t-test |
| RpS5b~Q9VFE4 | 3.404129444 | 0.042332699 | 14.51313611 | 13.07196223 | 0.216865182 | 0.648745076 | 1.441173875 | t-Test | FDR correction BH | 0.086790205 | FALSE | QL | SN | 2.715417205 | 1.373324041 | Paired t-test |
| Pvr~B6IDV6 | 3.466423219 | 0.040451135 | 16.12599021 | 14.68889979 | 0.688058489 | 0.806145566 | 1.437090417 | t-Test | FDR correction BH | 0.084778341 | FALSE | QL | SN | 2.707742253 | 1.393069293 | Paired t-test |
| CG11578~H9XVM5 | 3.518468091 | 0.038960081 | 14.49249741 | 13.05632325 | 0.597258756 | 0.798037205 | 1.436174167 | t-Test | FDR correction BH | 0.082972471 | FALSE | QL | SN | 2.706023122 | 1.409380153 | Paired t-test |
| CG17338~Q8MT77 | 5.179543035 | 0.013969635 | 15.19644754 | 13.7651634 | 0.235773689 | 0.524653971 | 1.431284144 | t-Test | FDR correction BH | 0.043148094 | TRUE | QL | SN | 2.696866569 | 1.854814933 | Paired t-test |
| Hsc70-1~P29843 | 4.802847987 | 0.017180352 | 16.06183197 | 14.63159231 | 0.478025556 | 0.554213533 | 1.430239662 | t-Test | FDR correction BH | 0.049047133 | TRUE | QL | SN | 2.694914799 | 1.764967948 | Paired t-test |
| Mec2~Q9VWL0 | 4.946762519 | 0.015850369 | 16.5191224 | 15.08950166 | 0.531628213 | 0.385200115 | 1.42962074 | t-Test | FDR correction BH | 0.047107534 | TRUE | QL | SN | 2.693758917 | 1.799960612 | Paired t-test |
| ed~M9PCI6 | 10.75674161 | 0.00171822 | 14.33033775 | 12.90380178 | 0.302320813 | 0.315368693 | 1.426535973 | t-Test | FDR correction BH | 0.016207074 | TRUE | QL | SN | 2.688005281 | 2.764921204 | Paired t-test |
| Atg13~Q9VHR6 | 3.358693336 | 0.043775685 | 14.99907859 | 13.57849742 | 0.787367923 | 0.151164188 | 1.420581166 | t-Test | FDR correction BH | 0.088947278 | FALSE | QL | SN | 2.676933251 | 1.358767053 | Paired t-test |
| Dmel\CG6055~Q9VM07 | 3.341150934 | 0.044349493 | 14.83544833 | 13.42731344 | 0.518804618 | 0.869042831 | 1.408134891 | t-Test | FDR correction BH | 0.089769685 | FALSE | QL | SN | 2.653938411 | 1.353111341 | Paired t-test |
| Slc25A46a~Q9VXK8 | 3.678413942 | 0.034795611 | 15.54390747 | 14.13668211 | 0.750572181 | 0.549074215 | 1.407225356 | t-Test | FDR correction BH | 0.077523649 | FALSE | QL | SN | 2.652265786 | 1.458475528 | Paired t-test |
| Slh~Q24179 | 3.378237566 | 0.043147449 | 21.61802413 | 20.21419017 | 0.429383495 | 0.606988194 | 1.403833964 | t-Test | FDR correction BH | 0.087992357 | FALSE | QL | SN | 2.646038337 | 1.365044871 | Paired t-test |
| nrv2~A4V0B5 | 7.210431508 | 0.005499251 | 17.45935415 | 16.05599512 | 0.317633158 | 0.106667174 | 1.40335903 | t-Test | FDR correction BH | 0.025173296 | TRUE | QL | SN | 2.645167407 | 2.259696446 | Paired t-test |
| Nrt~M9MS15 | 4.659283511 | 0.018655399 | 14.89027745 | 13.49974889 | 0.206528292 | 0.460778432 | 1.390528557 | t-Test | FDR correction BH | 0.051863962 | FALSE | QL | SN | 2.621747155 | 1.729195462 | Paired t-test |
| RabX6~Q9W0N7 | 4.870590739 | 0.016536918 | 14.48465436 | 13.09546279 | 0.410797067 | 0.33373255 | 1.389191568 | t-Test | FDR correction BH | 0.048228926 | TRUE | QL | SN | 2.619318628 | 1.781545415 | Paired t-test |
| kirre~Q9W4T9 | 10.06798068 | 0.002086559 | 14.40206772 | 13.01788905 | 0.147406284 | 0.228440946 | 1.384178671 | t-Test | FDR correction BH | 0.016678022 | TRUE | QL | SN | 2.61023314 | 2.680569386 | Paired t-test |
| fax~Q59E30 | 4.563627546 | 0.019730479 | 15.7241191 | 14.34105422 | 0.575189791 | 0.174778415 | 1.383064883 | t-Test | FDR correction BH | 0.053431096 | FALSE | QL | SN | 2.608218769 | 1.704862363 | Paired t-test |
| Akap200~Q9VLL3-2 | 5.438594682 | 0.012204646 | 16.88278013 | 15.50926938 | 0.559677093 | 0.41096032 | 1.373510743 | t-Test | FDR correction BH | 0.039596744 | TRUE | QL | SN | 2.591003103 | 1.913474812 | Paired t-test |
| Synd~Q9VDI1 | 6.033164624 | 0.009129917 | 16.308394 | 14.94945391 | 0.067388865 | 0.370414871 | 1.358940086 | t-Test | FDR correction BH | 0.033357703 | TRUE | QL | SN | 2.564966681 | 2.03953315 | Paired t-test |
| Sec8~Q9VNH6 | 7.528134472 | 0.004858329 | 14.86389852 | 13.50729141 | 0.49146836 | 0.560101043 | 1.356607113 | t-Test | FDR correction BH | 0.024066801 | TRUE | QL | SN | 2.560822244 | 2.313513112 | Paired t-test |
| Shab~M9MRU9 | 3.949085485 | 0.028961905 | 15.23844954 | 13.90479545 | 0.292180135 | 0.666238751 | 1.333654098 | t-Test | FDR correction BH | 0.06865523 | FALSE | QL | SN | 2.520402417 | 1.538172879 | Paired t-test |
| Dmel\CG11151~Q9VY92 | 7.283732655 | 0.005341913 | 15.94803608 | 14.61757167 | 0.621852476 | 0.320754447 | 1.330464404 | t-Test | FDR correction BH | 0.024737406 | TRUE | QL | SN | 2.514836145 | 2.272303174 | Paired t-test |
| trio~Q8IRI5 | 5.020128837 | 0.015223791 | 15.0072649 | 13.69762875 | 0.15652271 | 0.598275183 | 1.309636144 | t-Test | FDR correction BH | 0.045856543 | TRUE | QL | SN | 2.478790155 | 1.817477195 | Paired t-test |
| Oatp58Da~Q9W271 | 8.918512393 | 0.002973571 | 14.9188081 | 13.61114197 | 0.513676671 | 0.353075872 | 1.307666127 | t-Test | FDR correction BH | 0.019035448 | TRUE | QL | SN | 2.475407649 | 2.526721627 | Paired t-test |
| kcc~Q8MLQ5 | 9.724737876 | 0.002309655 | 17.97881679 | 16.67183879 | 0.115759501 | 0.213588766 | 1.306978008 | t-Test | FDR correction BH | 0.017112931 | TRUE | QL | SN | 2.474227241 | 2.636452908 | Paired t-test |
| Dmel\CG5676~Q9VL16 | 5.93242653 | 0.009572859 | 17.94818219 | 16.6527522 | 0.714466936 | 0.541186658 | 1.295429982 | t-Test | FDR correction BH | 0.034358997 | TRUE | QL | SN | 2.454501387 | 2.018958358 | Paired t-test |
| Lkb1~Q8T9L5 | 4.593476728 | 0.019386566 | 14.61021204 | 13.3151179 | 0.229820833 | 0.685739292 | 1.295094137 | t-Test | FDR correction BH | 0.053154562 | FALSE | QL | SN | 2.453930071 | 1.712499101 | Paired t-test |
| Dmel\CG31321~Q8INF8 | 11.72604935 | 0.001332786 | 16.09635303 | 14.80496833 | 0.214831406 | 0.320164495 | 1.291384695 | t-Test | FDR correction BH | 0.014923481 | TRUE | QL | SN | 2.447628656 | 2.875239679 | Paired t-test |
| CG18590~Q9I7N9 | 4.021514713 | 0.027617708 | 14.13547421 | 12.84732667 | 0.523059239 | 0.616680063 | 1.288147538 | t-Test | FDR correction BH | 0.066584131 | FALSE | QL | SN | 2.44214276 | 1.558812368 | Paired t-test |
| wus~Q9VX95 | 6.343984867 | 0.007922085 | 14.97271121 | 13.69155747 | 0.354328813 | 0.681625991 | 1.281153744 | t-Test | FDR correction BH | 0.030931083 | TRUE | QL | SN | 2.430332564 | 2.101160486 | Paired t-test |
| nahoda~Q9W1X5 | 9.227780089 | 0.002692256 | 17.02296484 | 15.75498591 | 0.141433115 | 0.311797575 | 1.267978927 | t-Test | FDR correction BH | 0.018328051 | TRUE | QL | SN | 2.408239586 | 2.569883623 | Paired t-test |
| Khc~P17210 | 4.566374208 | 0.019698504 | 15.43955883 | 14.17199649 | 0.498188311 | 0.553850544 | 1.267562345 | t-Test | FDR correction BH | 0.053431096 | FALSE | QL | SN | 2.4075443 | 1.70556675 | Paired t-test |
| Vmat~Q86NW1 | 4.992994082 | 0.015451707 | 18.6273544 | 17.36106136 | 0.184518623 | 0.317934663 | 1.266293042 | t-Test | FDR correction BH | 0.046181181 | TRUE | QL | SN | 2.405427041 | 1.81102353 | Paired t-test |
| Droj2~Q9VFV9 | 5.032823808 | 0.015118664 | 17.61220904 | 16.35012234 | 0.66407949 | 0.288390646 | 1.262086701 | t-Test | FDR correction BH | 0.045787132 | TRUE | QL | SN | 2.398423959 | 1.820486586 | Paired t-test |
| Cerk~Q9VNA6 | 6.610881368 | 0.007047373 | 14.24883879 | 12.98858223 | 0.207482637 | 0.123152544 | 1.260256558 | t-Test | FDR correction BH | 0.028709057 | TRUE | QL | SN | 2.395383349 | 2.151972728 | Paired t-test |
| FBgn 30055~Q9W3C9 | 6.487641735 | 0.007434598 | 14.29360048 | 13.0385864 | 0.310296596 | 0.202412354 | 1.255014083 | t-Test | FDR correction BH | 0.02953445 | TRUE | QL | SN | 2.386694784 | 2.128742508 | Paired t-test |
| Crag~Q9W3D3-2 | 7.375528142 | 0.005153141 | 15.77607812 | 14.52169769 | 0.563587727 | 0.370578694 | 1.254380428 | t-Test | FDR correction BH | 0.024608571 | TRUE | QL | SN | 2.385646739 | 2.287927971 | Paired t-test |
| Dmel\CG33090~X2JAF7 | 8.101362127 | 0.003930734 | 14.39672435 | 13.14873428 | 0.108200115 | 0.237308276 | 1.247990066 | t-Test | FDR correction BH | 0.021213554 | TRUE | QL | SN | 2.375102978 | 2.40552636 | Paired t-test |
| STUB1~Q9XYW6 | 5.84626822 | 0.009974307 | 17.66550023 | 16.4187776 | 0.126547179 | 0.252823141 | 1.246722631 | t-Test | FDR correction BH | 0.034844455 | TRUE | QL | SN | 2.373017321 | 2.001117253 | Paired t-test |
| snu~A0A0B4JD13 | 3.508843173 | 0.039230485 | 14.29145422 | 13.04756286 | 0.211952753 | 0.482061874 | 1.243891359 | t-Test | FDR correction BH | 0.083214596 | FALSE | QL | SN | 2.368364869 | 1.406376327 | Paired t-test |
| Rab1~O18332 | 6.616838541 | 0.007029335 | 18.25966291 | 17.01617086 | 0.45013549 | 0.203101352 | 1.243492045 | t-Test | FDR correction BH | 0.028709057 | TRUE | QL | SN | 2.367709436 | 2.153085766 | Paired t-test |
| Diap2~Q24307 | 6.109204106 | 0.008813238 | 14.57746545 | 13.34279596 | 0.306937255 | 0.136654916 | 1.234669486 | t-Test | FDR correction BH | 0.033190279 | TRUE | QL | SN | 2.35327429 | 2.054864508 | Paired t-test |
| Inx2~E1JJF3 | 5.380094197 | 0.012576549 | 15.71866517 | 14.50296909 | 0.37864291 | 0.313652925 | 1.215696076 | t-Test | FDR correction BH | 0.040473622 | TRUE | QL | SN | 2.32252814 | 1.90043851 | Paired t-test |
| Atg16~Q86BR6 | 5.813097156 | 0.010134745 | 15.54674191 | 14.33730059 | 0.323193704 | 0.242324451 | 1.209441325 | t-Test | FDR correction BH | 0.035020929 | TRUE | QL | SN | 2.3124807 | 1.994187192 | Paired t-test |
| stnB~Q24212 | 10.21892774 | 0.001997478 | 19.03363086 | 17.82506591 | 0.253954106 | 0.170317663 | 1.208564954 | t-Test | FDR correction BH | 0.016678022 | TRUE | QL | SN | 2.311076402 | 2.699518039 | Paired t-test |
| RhoGAP100F~A0A0B4LJ24 | 3.453919104 | 0.040820148 | 18.34705663 | 17.14113037 | 0.545561376 | 0.594280475 | 1.205926251 | t-Test | FDR correction BH | 0.084890988 | FALSE | QL | SN | 2.306853283 | 1.389125421 | Paired t-test |
| koi~A1Z6Q0 | 3.743470662 | 0.033265112 | 16.05283451 | 14.87165599 | 0.363252864 | 0.658646638 | 1.181178521 | t-Test | FDR correction BH | 0.076027724 | FALSE | QL | SN | 2.267619406 | 1.47801101 | Paired t-test |
| Ras64B~P04388 | 3.211776413 | 0.048889905 | 14.1727178 | 12.9925942 | 0.028297269 | 0.61382867 | 1.180123596 | t-Test | FDR correction BH | 0.095913324 | FALSE | QL | SN | 2.265961888 | 1.310780802 | Paired t-test |
| Evi5~Q9VYY9-3 | 6.015262608 | 0.009206637 | 16.88105967 | 15.7051367 | 0.467330125 | 0.254389465 | 1.175922964 | t-Test | FDR correction BH | 0.033370926 | TRUE | QL | SN | 2.259373781 | 2.035898972 | Paired t-test |
| PMCA~E6EK15 | 11.45563691 | 0.001427663 | 18.7417001 | 17.56916453 | 0.269723064 | 0.11161077 | 1.172535571 | t-Test | FDR correction BH | 0.015251006 | TRUE | QL | SN | 2.254075081 | 2.845374196 | Paired t-test |
| Tomosyn~Q9VYK6 | 8.039124159 | 0.004019451 | 18.85701114 | 17.68737192 | 0.278489077 | 0.247265599 | 1.169639221 | t-Test | FDR correction BH | 0.021213554 | TRUE | QL | SN | 2.249554347 | 2.395833315 | Paired t-test |
| NTPase~O76268 | 9.19274291 | 0.002722307 | 15.75276198 | 14.58749029 | 0.23583219 | 0.232183529 | 1.165271691 | t-Test | FDR correction BH | 0.018453763 | TRUE | QL | SN | 2.242754476 | 2.565062965 | Paired t-test |
| Ehbp1~A1ZAN2 | 8.714745859 | 0.003180491 | 17.01389399 | 15.85088385 | 0.165051341 | 0.084847396 | 1.163010147 | t-Test | FDR correction BH | 0.019411961 | TRUE | QL | SN | 2.239241526 | 2.497505864 | Paired t-test |
| anon-EST:fe1H12~A1ZBH1 | 9.077468065 | 0.002824365 | 15.09324074 | 13.94877544 | 0.691521021 | 0.478417817 | 1.144465302 | t-Test | FDR correction BH | 0.018663977 | TRUE | QL | SN | 2.210641837 | 2.549079196 | Paired t-test |
| g~M9PJM7 | 3.472916359 | 0.040261188 | 13.98376023 | 12.83983759 | 0.609605481 | 0.324955154 | 1.143922639 | t-Test | FDR correction BH | 0.084581635 | FALSE | QL | SN | 2.20981047 | 1.395113418 | Paired t-test |
| bond~Q9VCY7 | 5.571205519 | 0.011413329 | 16.60423807 | 15.46404308 | 0.067865104 | 0.362173715 | 1.140194987 | t-Test | FDR correction BH | 0.037584122 | TRUE | QL | SN | 2.204108108 | 1.942587671 | Paired t-test |
| CG12441~Q0IGY6 | 3.294296729 | 0.045929556 | 14.66105849 | 13.52707274 | 0.131323974 | 0.470282636 | 1.133985751 | t-Test | FDR correction BH | 0.091801484 | FALSE | QL | SN | 2.194642199 | 1.337907753 | Paired t-test |
| Snap24~Q9VH76 | 8.862693427 | 0.003028438 | 19.0516277 | 17.92577788 | 0.076733079 | 0.158447327 | 1.12584982 | t-Test | FDR correction BH | 0.019235452 | TRUE | QL | SN | 2.182300571 | 2.518781352 | Paired t-test |
| PMCA~Q59DP9 | 13.80489192 | 0.000822675 | 16.88048268 | 15.75986223 | 0.131185366 | 0.227293626 | 1.120620445 | t-Test | FDR correction BH | 0.012481149 | TRUE | QL | SN | 2.174404648 | 3.084771926 | Paired t-test |
| Dmel\CG12576~Q8IQ25 | 7.287657087 | 0.005333658 | 16.43773041 | 15.32152613 | 0.537508277 | 0.416702745 | 1.116204282 | t-Test | FDR correction BH | 0.024737406 | TRUE | QL | SN | 2.167758861 | 2.272974843 | Paired t-test |
| pst~Q8IQ82 | 5.91491788 | 0.0096527 | 16.7449334 | 15.62922482 | 0.317557473 | 0.390447237 | 1.115708584 | t-Test | FDR correction BH | 0.034358997 | TRUE | QL | SN | 2.167014165 | 2.015351196 | Paired t-test |
| Alk~Q7KJ08 | 5.10560304 | 0.014533891 | 14.90385053 | 13.79327752 | 0.519725098 | 0.151591352 | 1.110573015 | t-Test | FDR correction BH | 0.044268621 | TRUE | QL | SN | 2.159313947 | 1.837618097 | Paired t-test |
| Dmel\CG30392~Q8MLW4 | 3.713366314 | 0.033962391 | 16.14188657 | 15.0350218 | 0.570624806 | 0.157030709 | 1.106864772 | t-Test | FDR correction BH | 0.07641514 | FALSE | QL | SN | 2.153770864 | 1.469001748 | Paired t-test |
| Arp1~P45889 | 3.267584024 | 0.046862377 | 14.75902066 | 13.66035394 | 0.302334829 | 0.761815755 | 1.098666722 | t-Test | FDR correction BH | 0.092783534 | FALSE | QL | SN | 2.141566864 | 1.329175684 | Paired t-test |
| Tspo~Q9VPR1 | 14.54670989 | 0.000704428 | 18.58661849 | 17.48971351 | 0.108783162 | 0.150994063 | 1.096904975 | t-Test | FDR correction BH | 0.012134178 | TRUE | QL | SN | 2.138953286 | 3.152163171 | Paired t-test |
| nrm~Q9VNP2 | 7.484219548 | 0.004940834 | 20.24868196 | 19.15944242 | 0.198712422 | 0.206966157 | 1.089239541 | t-Test | FDR correction BH | 0.024292432 | TRUE | QL | SN | 2.127618581 | 2.306199763 | Paired t-test |
| ClC-a~Q9VGH7-6 | 10.0383846 | 0.002104639 | 15.89165216 | 14.80308597 | 0.074713699 | 0.120329413 | 1.088566184 | t-Test | FDR correction BH | 0.016678022 | TRUE | QL | SN | 2.126625776 | 2.676822363 | Paired t-test |
| RpL28~D1Z3A1 | 9.47773166 | 0.002490122 | 14.552787 | 13.46452703 | 0.440097834 | 0.400793435 | 1.088259967 | t-Test | FDR correction BH | 0.01778818 | TRUE | QL | SN | 2.12617444 | 2.60377939 | Paired t-test |
| igl~Q7JPS2 | 6.964690896 | 0.006073475 | 18.9769406 | 17.89091176 | 0.218436832 | 0.1211352 | 1.086028839 | t-Test | FDR correction BH | 0.026651081 | TRUE | QL | SN | 2.122888853 | 2.216562742 | Paired t-test |
| Bem46~O76462 | 8.379502802 | 0.003564397 | 17.50891956 | 16.43424881 | 0.307569702 | 0.28462557 | 1.074670748 | t-Test | FDR correction BH | 0.020315914 | TRUE | QL | SN | 2.10624133 | 2.448013968 | Paired t-test |
| Nrg~M9NEZ5 | 8.341487032 | 0.003611726 | 18.41450663 | 17.34296338 | 0.149231451 | 0.175666044 | 1.071543254 | t-Test | FDR correction BH | 0.020315914 | TRUE | QL | SN | 2.101680338 | 2.442285169 | Paired t-test |

| | | | | | | | | | | | | | | | | |
|---|---|---|---|---|---|---|---|---|---|---|---|---|---|---|---|---|
| Dmel\CG1882~Q5U191 | 11.21274773 | 0.001520677 | 18.46621432 | 17.39666338 | 0.058731918 | 0.144048153 | 1.06955094 | t-Test | FDR correction BH | 0.015832927 | TRUE | QL | SN | 2.098779989 | 2.817963131 | Paired t-test |
| Dmel\CG11396~Q9VPB6 | 12.4145403 | 0.001126228 | 15.51228892 | 14.44386026 | 0.329499245 | 0.319446571 | 1.068428664 | t-Test | FDR correction BH | 0.013388667 | TRUE | QL | SN | 2.097147978 | 2.94837363 | Paired t-test |
| Dmel\CG30382~E1JGZ9 | 4.682415971 | 0.018406898 | 14.64136704 | 13.57703283 | 0.456345137 | 0.524941019 | 1.064334213 | t-Test | FDR correction BH | 0.051623572 | FALSE | QL | SN | 2.09120459 | 1.735019391 | Paired t-test |
| colt~Q9VQG4 | 3.667913834 | 0.035051019 | 14.66948099 | 13.60580906 | 0.08726042 | 0.461872566 | 1.063671929 | t-Test | FDR correction BH | 0.077874858 | FALSE | QL | SN | 2.090244822 | 1.45529935 | Paired t-test |
| subdued~Q9VDV4 | 10.67061162 | 0.001759298 | 16.44337212 | 15.38134656 | 0.111021604 | 0.08492912 | 1.062025553 | t-Test | FDR correction BH | 0.016234815 | TRUE | QL | SN | 2.087860835 | 2.754660683 | Paired t-test |
| Dmel\CG9485~Q9W2H8 | 7.076081953 | 0.005803854 | 20.41396998 | 19.35402333 | 0.23918569 | 0.477569597 | 1.059946647 | t-Test | FDR correction BH | 0.025897868 | TRUE | QL | SN | 2.084854419 | 2.236283541 | Paired t-test |
| Dmel\CG32544~Q8IQX3 | 6.057664973 | 0.009026274 | 19.38391151 | 18.33565812 | 0.397777798 | 0.249525139 | 1.048253396 | t-Test | FDR correction BH | 0.033357703 | TRUE | QL | SN | 2.068024671 | 2.044491482 | Paired t-test |
| trc~M9PG43 | 5.683783012 | 0.010793478 | 14.23950803 | 13.19643662 | 0.387798408 | 0.534034711 | 1.043071415 | t-Test | FDR correction BH | 0.036454821 | TRUE | QL | SN | 2.060609907 | 1.966838593 | Paired t-test |
| Dmel\CG17029~Q9VUW2 | 5.235116166 | 0.013564232 | 15.76923706 | 14.72798214 | 0.039193163 | 0.307848222 | 1.041254924 | t-Test | FDR correction BH | 0.042535083 | TRUE | QL | SN | 2.058017035 | 1.867604779 | Paired t-test |
| Trs31~Q7K2Q8 | 5.301508263 | 0.013099803 | 17.21041702 | 16.17018718 | 0.394678452 | 0.416800456 | 1.040229839 | t-Test | FDR correction BH | 0.041652668 | TRUE | QL | SN | 2.056555262 | 1.88273523 | Paired t-test |
| OtopLc~B7Z0W9-3 | 14.8081641 | 0.000668164 | 16.54267045 | 15.51569156 | 0.114256475 | 0.114466459 | 1.026978898 | t-Test | FDR correction BH | 0.012134178 | TRUE | QL | SN | 2.037752587 | 3.175117243 | Paired t-test |
| Amph~Q7KLE5 | 6.744293864 | 0.00665754 | 21.99006034 | 20.96552583 | 0.420548572 | 0.250933972 | 1.024534517 | t-Test | FDR correction BH | 0.027994359 | TRUE | QL | SN | 2.034302914 | 2.176686224 | Paired t-test |
| DCTN4-p62~Q7K130 | 3.476181755 | 0.040166093 | 16.53496069 | 15.52243738 | 0.146698748 | 0.446891868 | 1.01252331 | t-Test | FDR correction BH | 0.084548776 | FALSE | QL | SN | 2.017436563 | 1.396140415 | Paired t-test |
| Nhe2~M9NFD1 | 6.566772462 | 0.007182848 | 14.13979209 | 13.13058879 | 0.200544279 | 0.386309817 | 1.009203305 | t-Test | FDR correction BH | 0.029041313 | TRUE | QL | SN | 2.012799271 | 2.143703299 | Paired t-test |
| Dmel\CG7332~X2JL49 | 3.709290721 | 0.034058223 | 16.87305491 | 15.86609126 | 0.265290921 | 0.247998343 | 1.006963652 | t-Test | FDR correction BH | 0.07641514 | FALSE | QL | SN | 2.009677008 | 1.467778016 | Paired t-test |
| PIP5K59B~Q9W1Y2 | 4.86120576 | 0.016624166 | 14.7115313 | 13.71112857 | 0.208885788 | 0.409448437 | 1.000402737 | t-Test | FDR correction BH | 0.048228926 | TRUE | QL | SN | 2.000558391 | 1.779260132 | Paired t-test |
| Plc21C~P25455-6 | 6.437759358 | 0.007599282 | 16.2462657 | 15.25249603 | 0.264053638 | 0.288769674 | 0.993769667 | t-Test | FDR correction BH | 0.029816887 | TRUE | QL | SN | 1.991381548 | 2.119227438 | Paired t-test |
| Nup133~Q9VCW3 | 3.77609113 | 0.032530119 | 15.60684461 | 14.62318987 | 0.446519641 | 0.453027859 | 0.983654743 | t-Test | FDR correction BH | 0.075018876 | FALSE | QL | SN | 1.977468541 | 1.487714353 | Paired t-test |
| Snap29~Q9W1I8 | 9.303075741 | 0.002629148 | 17.85314235 | 16.86975833 | 0.130012605 | 0.134691151 | 0.983384017 | t-Test | FDR correction BH | 0.018079919 | TRUE | QL | SN | 1.977097497 | 2.58018501 | Paired t-test |
| Argk~P48610-2 | 9.360226821 | 0.002582549 | 21.69411905 | 20.71087462 | 0.25542503 | 0.130177066 | 0.983244429 | t-Test | FDR correction BH | 0.018043861 | TRUE | QL | SN | 1.976906213 | 2.587951446 | Paired t-test |
| SPoCk~Q7KTU2 | 8.098003005 | 0.003935456 | 16.68461873 | 15.70180682 | 0.243109543 | 0.177750171 | 0.982811916 | t-Test | FDR correction BH | 0.021213554 | TRUE | QL | SN | 1.976313635 | 2.405004976 | Paired t-test |
| Dmel\CG42258~Q8SWS2 | 7.410350567 | 0.005083834 | 18.58472582 | 17.61030616 | 0.269326839 | 0.081606063 | 0.974419663 | t-Test | FDR correction BH | 0.024608571 | TRUE | QL | SN | 1.96485066 | 2.293808619 | Paired t-test |
| Vps53~Q9VQY8 | 3.925305938 | 0.029421515 | 15.6436174 | 14.69191737 | 0.62233583 | 0.248559368 | 0.951700035 | t-Test | FDR correction BH | 0.068933059 | FALSE | QL | SN | 1.934150469 | 1.531334961 | Paired t-test |
| Use1~Q9VSU7 | 5.193078281 | 0.013869455 | 15.61052367 | 14.66231632 | 0.352575509 | 0.307839899 | 0.948207346 | t-Test | FDR correction BH | 0.043138422 | TRUE | QL | SN | 1.929473655 | 1.857940596 | Paired t-test |
| Pldn~Q9VTM0 | 4.176969567 | 0.02499312 | 15.41652609 | 14.46838292 | 0.59163581 | 0.264078447 | 0.948143167 | t-Test | FDR correction BH | 0.062600691 | FALSE | QL | SN | 1.929387824 | 1.602179534 | Paired t-test |
| anon-EST:Posey232~Q9I7R0 | 6.049649362 | 0.009060012 | 15.78664698 | 14.84417534 | 0.196627666 | 0.323184505 | 0.942471632 | t-Test | FDR correction BH | 0.033357703 | TRUE | QL | SN | 1.921817887 | 2.042871243 | Paired t-test |
| ScpX~Q9VJ43 | 3.47577516 | 0.040177918 | 14.79726579 | 13.86075033 | 0.596702607 | 0.653545189 | 0.936515465 | t-Test | FDR correction BH | 0.084548776 | FALSE | QL | SN | 1.913900017 | 1.396012573 | Paired t-test |
| Dmel\CG14696~Q9VGU7 | 5.190016523 | 0.013892034 | 15.68536588 | 14.75064452 | 0.232483252 | 0.11560057 | 0.934721362 | t-Test | FDR correction BH | 0.043138422 | TRUE | QL | SN | 1.911521414 | 1.857234152 | Paired t-test |
| Dmel\CG7888~Q7YTZ0 | 5.004734938 | 0.015352549 | 17.25839688 | 16.32590146 | 0.073807204 | 0.254729328 | 0.932495417 | t-Test | FDR correction BH | 0.045971072 | TRUE | QL | SN | 1.908574387 | 1.813819514 | Paired t-test |
| mtd~Q9VNA1 | 23.56489079 | 0.000167443 | 20.59062912 | 19.66295697 | 0.161337201 | 0.179452479 | 0.927672148 | t-Test | FDR correction BH | 0.007209085 | TRUE | QL | SN | 1.902204228 | 3.776133905 | Paired t-test |
| PlexA~Q9V491 | 3.650352786 | 0.035483562 | 16.78885922 | 15.86148301 | 1.018059042 | 0.701535708 | 0.927376211 | t-Test | FDR correction BH | 0.078398494 | FALSE | QL | SN | 1.901814073 | 1.449972789 | Paired t-test |
| Fas3~A8DZ14 | 5.5980602 | 0.01126132 | 15.70640114 | 14.7875912 | 0.375004281 | 0.348671692 | 0.918809934 | t-Test | FDR correction BH | 0.037218429 | TRUE | QL | SN | 1.890555148 | 1.948410694 | Paired t-test |
| Dap160~M9PBF6 | 6.088458169 | 0.008898189 | 17.34748021 | 16.43267423 | 0.290228029 | 0.122500482 | 0.914805978 | t-Test | FDR correction BH | 0.033352506 | TRUE | QL | SN | 1.885315506 | 2.05069838 | Paired t-test |
| RyR~Q24498-3 | 6.206088848 | 0.008430304 | 18.74354363 | 17.83008012 | 0.377051857 | 0.173157811 | 0.913463502 | t-Test | FDR correction BH | 0.032127929 | TRUE | QL | SN | 1.883561973 | 2.074156766 | Paired t-test |
| kek5~Q9VWI6 | 3.573207654 | 0.037466584 | 15.57375054 | 14.66079802 | 0.654805141 | 0.935006537 | 0.91295252 | t-Test | FDR correction BH | 0.081203087 | FALSE | QL | SN | 1.882894961 | 1.426355899 | Paired t-test |
| yata~Q9VAH7 | 5.129278384 | 0.014350034 | 17.00902364 | 16.09616244 | 0.378409747 | 0.139178131 | 0.9128612 | t-Test | FDR correction BH | 0.0438764 | TRUE | QL | SN | 1.882775781 | 1.843147061 | Paired t-test |
| Trap1~A1Z6L9 | 3.390530827 | 0.042758151 | 14.06274694 | 13.15178841 | 0.248993018 | 0.376245625 | 0.910958524 | t-Test | FDR correction BH | 0.08732638 | FALSE | QL | SN | 1.880294348 | 1.368981082 | Paired t-test |
| Dmel\CG3566~Q8SY77 | 3.266892421 | 0.046886846 | 14.20374606 | 13.29880606 | 0.23564062 | 0.390966859 | 0.90494 | t-Test | FDR correction BH | 0.092783534 | FALSE | QL | SN | 1.87246662 | 1.328948985 | Paired t-test |
| Dmel\CG1246~Q9VZZ0 | 6.493336325 | 0.007416097 | 18.01822664 | 17.11750947 | 0.337280904 | 0.114742777 | 0.900717172 | t-Test | FDR correction BH | 0.02953445 | TRUE | QL | SN | 1.866993846 | 2.12982459 | Paired t-test |
| RhoL~D3DMW7 | 4.092227092 | 0.02638188 | 14.26370763 | 13.36468586 | 0.537509828 | 0.451750825 | 0.89902177 | t-Test | FDR correction BH | 0.064556583 | FALSE | QL | SN | 1.864801112 | 1.578694253 | Paired t-test |
| Dmel\CG8888~Q7K3N4 | 3.471595871 | 0.040299724 | 15.09992437 | 14.20462981 | 0.361626599 | 0.39556821 | 0.895294561 | t-Test | FDR correction BH | 0.084581635 | FALSE | QL | SN | 1.859989609 | 1.394697924 | Paired t-test |
| Irk1~A8JR87 | 7.377237585 | 0.00514971 | 17.45147062 | 16.55894301 | 0.467274642 | 0.25953041 | 0.892527615 | t-Test | FDR correction BH | 0.024608571 | TRUE | QL | SN | 1.856425751 | 2.288217247 | Paired t-test |
| Cadps~Q9NHE5-2 | 6.04609341 | 0.009075032 | 17.05730013 | 16.16769766 | 0.202134097 | 0.161816723 | 0.889602471 | t-Test | FDR correction BH | 0.033357703 | TRUE | QL | SN | 1.852665559 | 2.042151854 | Paired t-test |
| Faf2~Q9VJ58 | 4.85684003 | 0.016664956 | 16.29454716 | 15.40958946 | 0.728869886 | 0.430891745 | 0.884957695 | t-Test | FDR correction BH | 0.048228926 | TRUE | QL | SN | 1.846710468 | 1.778195821 | Paired t-test |
| cpx~E1JJ37 | 6.777482379 | 0.006564994 | 21.31168693 | 20.42697055 | 0.052034678 | 0.187834443 | 0.884716381 | t-Test | FDR correction BH | 0.027962661 | TRUE | QL | SN | 1.846401602 | 2.182765682 | Paired t-test |
| Dmel\CG6126~Q961R9 | 12.76754766 | 0.001036667 | 17.25513278 | 16.37554204 | 0.165319004 | 0.100050101 | 0.879590736 | t-Test | FDR correction BH | 0.013062707 | TRUE | QL | SN | 1.839853297 | 2.984360658 | Paired t-test |
| Drp1~Q9VQE0 | 12.7478567 | 0.001041409 | 18.44375155 | 17.56695698 | 0.249001921 | 0.173615663 | 0.876794566 | t-Test | FDR correction BH | 0.013062707 | TRUE | QL | SN | 1.836290825 | 2.98237886 | Paired t-test |
| Dmel\CG1648~Q7K2P3 | 8.083246183 | 0.003956288 | 22.21131855 | 21.34935837 | 0.248765608 | 0.166309085 | 0.861960175 | t-Test | FDR correction BH | 0.021213554 | TRUE | QL | SN | 1.817506061 | 2.402712117 | Paired t-test |
| ND-B16.6~Q9W402 | 3.267554301 | 0.046863428 | 16.68471337 | 15.82602588 | 0.134015457 | 0.372764701 | 0.858687485 | t-Test | FDR correction BH | 0.092783534 | FALSE | QL | SN | 1.813387801 | 1.329165942 | Paired t-test |
| cg3702~Q9VQX6 | 4.131314817 | 0.025729263 | 17.81512753 | 16.95916248 | 0.255522032 | 0.188399218 | 0.855965053 | t-Test | FDR correction BH | 0.06368161 | FALSE | QL | SN | 1.809969081 | 1.589572646 | Paired t-test |
| Pen~P52295 | 3.538302582 | 0.038410298 | 16.22075006 | 15.36750388 | 0.355924668 | 0.549096821 | 0.853246179 | t-Test | FDR correction BH | 0.082131013 | FALSE | QL | SN | 1.806561263 | 1.415552324 | Paired t-test |
| ReepB~A1Z9M2 | 5.369496564 | 0.012645499 | 17.71391613 | 16.86886482 | 0.489593013 | 0.318912586 | 0.845051306 | t-Test | FDR correction BH | 0.040613469 | TRUE | QL | SN | 1.796328627 | 1.898064017 | Paired t-test |
| Gint3~Q7K0S5 | 4.635757891 | 0.01891261 | 16.81772802 | 15.97975237 | 0.505353013 | 0.318793356 | 0.83797565 | t-Test | FDR correction BH | 0.052305186 | FALSE | QL | SN | 1.787540154 | 1.72324854 | Paired t-test |
| CHMP2B~Q9VRJ5 | 8.285762706 | 0.003682616 | 17.32038095 | 16.48377772 | 0.183321526 | 0.132086953 | 0.836603227 | t-Test | FDR correction BH | 0.020440444 | TRUE | QL | SN | 1.785840492 | 2.433843547 | Paired t-test |
| GlyT~Q9W1J0 | 3.232209277 | 0.048134923 | 16.4170723 | 15.58733128 | 0.560486197 | 0.313419843 | 0.829741022 | t-Test | FDR correction BH | 0.095017264 | FALSE | QL | SN | 1.777366278 | 1.317539719 | Paired t-test |
| Dmel\CG10737~Q0E919 | 4.686429629 | 0.01836422 | 17.87230891 | 17.04600437 | 0.380848148 | 0.23559601 | 0.826304534 | t-Test | FDR correction BH | 0.051594713 | FALSE | QL | SN | 1.773137645 | 1.736027517 | Paired t-test |
| Syx18~Q9VC58 | 4.163875661 | 0.025201437 | 17.43906412 | 16.61425516 | 0.215267566 | 0.186515126 | 0.824808965 | t-Test | FDR correction BH | 0.062826118 | FALSE | QL | SN | 1.771300475 | 1.598574692 | Paired t-test |
| Dmel\CG5174~A0A0B4K7G1 | 7.056305558 | 0.005850554 | 20.53163392 | 19.7124624 | 0.279086587 | 0.153383375 | 0.819171517 | t-Test | FDR correction BH | 0.026033331 | TRUE | QL | SN | 1.764392481 | 2.232803004 | Paired t-test |
| Desat1~Q7K4Y0 | 3.440831573 | 0.041210981 | 16.36512596 | 15.54624063 | 0.207961775 | 0.410313749 | 0.818885334 | t-Test | FDR correction BH | 0.085447912 | FALSE | QL | SN | 1.764042518 | 1.38498705 | Paired t-test |
| Myo31DF~Q23978 | 6.853118189 | 0.006360323 | 14.44032379 | 13.6286912 | 0.352088855 | 0.460648047 | 0.811632589 | t-Test | FDR correction BH | 0.027492254 | TRUE | QL | SN | 1.755196543 | 2.19652081 | Paired t-test |
| rb~Q9W4K1 | 3.965134173 | 0.028656924 | 14.58146218 | 13.77288711 | 0.397951671 | 0.256534929 | 0.808575074 | t-Test | FDR correction BH | 0.068135045 | FALSE | QL | SN | 1.75148068 | 1.542770423 | Paired t-test |
| Tango11~Q961C9 | 16.88589084 | 0.000452317 | 18.66606869 | 17.86346424 | 0.094455904 | 0.058674263 | 0.802604453 | t-Test | FDR correction BH | 0.010461282 | TRUE | QL | SN | 1.744247122 | 3.344557433 | Paired t-test |
| Fitm~Q9VRJ2 | 3.41207192 | 0.042086695 | 17.03930128 | 16.2411923 | 0.630794538 | 0.564404581 | 0.798108978 | t-Test | FDR correction BH | 0.086508522 | FALSE | QL | SN | 1.738820462 | 1.37585518 | Paired t-test |
| FACL~Q9VCC6 | 5.013878104 | 0.015275903 | 15.76660008 | 14.98183242 | 0.081692471 | 0.216866925 | 0.784767661 | t-Test | FDR correction BH | 0.045856543 | TRUE | QL | SN | 1.722814846 | 1.81599311 | Paired t-test |
| zda~Q7K3D4 | 4.288020395 | 0.023312708 | 17.38610445 | 16.60310686 | 0.398241677 | 0.280340261 | 0.782997589 | t-Test | FDR correction BH | 0.059717089 | FALSE | QL | SN | 1.720702385 | 1.632407279 | Paired t-test |
| atl~A0A0B4LHV8 | 4.154837144 | 0.025346544 | 15.47264323 | 14.69886235 | 0.047255158 | 0.297808075 | 0.77378088 | t-Test | FDR correction BH | 0.062990709 | FALSE | QL | SN | 1.709744656 | 1.596081249 | Paired t-test |
| Gdi~Q9VLB7 | 8.925968481 | 0.002966342 | 15.45994261 | 14.68648454 | 0.159568688 | 0.021432226 | 0.773458075 | t-Test | FDR correction BH | 0.019035448 | TRUE | QL | SN | 1.709362141 | 2.527778713 | Paired t-test |
| Syn~E2QCY9 | 8.178242203 | 0.003824683 | 20.35086335 | 19.59339043 | 0.16204519 | 0.073294111 | 0.757472921 | t-Test | FDR correction BH | 0.020865478 | TRUE | QL | SN | 1.69052684 | 2.417404591 | Paired t-test |
| Vps8~Q9VRX2 | 4.597665725 | 0.019338926 | 13.75794521 | 13.0044615 | 0.260460805 | 0.18999387 | 0.753483711 | t-Test | FDR correction BH | 0.053115361 | FALSE | QL | SN | 1.685858803 | 1.713567642 | Paired t-test |
| Acsl~A1Z7H3 | 9.504455917 | 0.002469715 | 16.63280699 | 15.88274031 | 0.150023899 | 0.034733202 | 0.750066679 | t-Test | FDR correction BH | 0.017758453 | TRUE | QL | SN | 1.681870563 | 2.607353179 | Paired t-test |
| Ccdc128~Q9VC66 | 4.299116294 | 0.023152929 | 15.45616285 | 14.71003305 | 0.338918178 | 0.063311048 | 0.746129796 | t-Test | FDR correction BH | 0.059488089 | FALSE | QL | SN | 1.677287264 | 1.635394067 | Paired t-test |
| Sap47~Q960T2-5 | 4.85503927 | 0.016681819 | 21.9790113 | 21.24078798 | 0.245463406 | 0.166737539 | 0.738223323 | t-Test | FDR correction BH | 0.048228926 | TRUE | QL | SN | 1.668120285 | 1.77775659 | Paired t-test |
| Dmel\CG11980~Q9VHI8 | 4.083537193 | 0.026529859 | 14.35955789 | 13.62655753 | 0.120620892 | 0.38862857 | 0.733000364 | t-Test | FDR correction BH | 0.064719856 | FALSE | QL | SN | 1.662092142 | 1.576265051 | Paired t-test |
| Rab7~O76742 | 3.99514022 | 0.028097702 | 16.49183731 | 15.75993905 | 0.389192839 | 0.354654505 | 0.731898256 | t-Test | FDR correction BH | 0.067409096 | FALSE | QL | SN | 1.660822917 | 1.551329192 | Paired t-test |
| deltaCOP~Q7K7G0 | 6.593670034 | 0.007099833 | 19.80422296 | 19.09173071 | 0.188950697 | 0.170695674 | 0.712492247 | t-Test | FDR correction BH | 0.02877871 | TRUE | QL | SN | 1.638632401 | 2.148751896 | Paired t-test |
| ArfGAP1~Q9VTX5 | 8.9487552 | 0.002944393 | 18.99192354 | 18.28964314 | 0.203447645 | 0.083470996 | 0.702280403 | t-Test | FDR correction BH | 0.019035448 | TRUE | QL | SN | 1.627074604 | 2.53100415 | Paired t-test |
| Dmel\CG3164~Q9VPJ9 | 6.241640047 | 0.008295276 | 18.05060088 | 17.35583272 | 0.340752835 | 0.317371378 | 0.694768158 | t-Test | FDR correction BH | 0.031765323 | TRUE | QL | SN | 1.618624298 | 2.081169171 | Paired t-test |
| Rop~M9PEL1 | 3.710682648 | 0.034025455 | 16.55117668 | 15.8673795 | 0.413324787 | 0.202072042 | 0.683797178 | t-Test | FDR correction BH | 0.07641514 | FALSE | QL | SN | 1.606362146 | 1.468196063 | Paired t-test |
| alphaTub67C~P06606 | 3.214453148 | 0.048790154 | 16.35094701 | 15.66938138 | 0.177371647 | 0.540313149 | 0.681565623 | t-Test | FDR correction BH | 0.095835653 | FALSE | QL | SN | 1.603879353 | 1.311667815 | Paired t-test |
| Agpat3~D5AEK7 | 3.436010692 | 0.041356146 | 16.55672458 | 15.87560794 | 0.524464228 | 0.281527899 | 0.681116638 | t-Test | FDR correction BH | 0.085447912 | FALSE | QL | SN | 1.603380283 | 1.383459943 | Paired t-test |
| Liprin-beta~Q9VU88 | 4.809358945 | 0.01711711 | 16.14142491 | 15.46685093 | 0.165610139 | 0.31199731 | 0.674573978 | t-Test | FDR correction BH | 0.049042368 | TRUE | QL | SN | 1.596125373 | 1.76656955 | Paired t-test |
| Sccpdh1~Q9VN86 | 7.428568279 | 0.005048065 | 14.96973914 | 14.29702633 | 0.306512129 | 0.177818189 | 0.672712812 | t-Test | FDR correction BH | 0.024608571 | TRUE | QL | SN | 1.5940676 | 2.296875047 | Paired t-test |
| Atpalpha~P13607-4 | 4.71749799 | 0.018038172 | 18.5363229 | 17.86437849 | 0.258139531 | 0.040795786 | 0.671944403 | t-Test | FDR correction BH | 0.051038734 | FALSE | QL | SN | 1.593218794 | 1.743807488 | Paired t-test |
| Rpn3~M9PG62 | 4.018599469 | 0.027670241 | 15.19397031 | 14.52673604 | 0.491687686 | 0.312085296 | 0.667234269 | t-Test | FDR correction BH | 0.066584131 | FALSE | QL | SN | 1.588025709 | 1.557987051 | Paired t-test |
| Dmel\CG5745~Q95RC5 | 4.674262417 | 0.018493994 | 15.90093362 | 15.23449857 | 0.28671707 | 0.472372141 | 0.666435051 | t-Test | FDR correction BH | 0.051711753 | FALSE | QL | SN | 1.587146225 | 1.732969278 | Paired t-test |
| Marc~A1Z803 | 9.986926022 | 0.002136573 | 18.360247 | 17.69498197 | 0.18538444 | 0.08599701 | 0.665265026 | t-Test | FDR correction BH | 0.016678022 | TRUE | QL | SN | 1.585859572 | 2.670282275 | Paired t-test |
| ATP7~Q9VYT4 | 3.625496847 | 0.03610752 | 16.73853729 | 16.07613293 | 0.338204539 | 0.176359477 | 0.662404363 | t-Test | FDR correction BH | 0.079118678 | FALSE | QL | SN | 1.58271815 | 1.442402341 | Paired t-test |
| Dmel\CG3529~M9PF36 | 7.787550088 | 0.004406266 | 19.86683296 | 19.21027582 | 0.239229041 | 0.132868936 | 0.656557137 | t-Test | FDR correction BH | 0.022531262 | TRUE | QL | SN | 1.576316393 | 2.355929331 | Paired t-test |
| Dmel\CG5653~Q9VSR8 | 4.045525257 | 0.027189877 | 15.76118167 | 15.10988818 | 0.226937379 | 0.223995666 | 0.651293486 | t-Test | FDR correction BH | 0.066127442 | FALSE | QL | SN | 1.570575706 | 1.56559275 | Paired t-test |
| Hcs~Q9VNC3 | 13.29074903 | 0.000920538 | 17.73559386 | 17.08659903 | 0.026427054 | 0.082242578 | 0.648994825 | t-Test | FDR correction BH | 0.012641521 | TRUE | QL | SN | 1.568075284 | 3.035958442 | Paired t-test |
| Dmel\CG15602~Q4QQ70 | 6.045335443 | 0.009078237 | 17.09567551 | 16.44886787 | 0.138272932 | 0.070465203 | 0.646807637 | t-Test | FDR correction BH | 0.033357703 | TRUE | QL | SN | 1.565699815 | 2.041998465 | Paired t-test |
| Stim~P83094 | 6.754275162 | 0.006629527 | 18.49379687 | 17.85551345 | 0.13717529 | 0.102651774 | 0.638283422 | t-Test | FDR correction BH | 0.027994359 | TRUE | QL | SN | 1.556476098 | 2.178517426 | Paired t-test |
| RpL23~P48159 | 14.1532954 | 0.000764095 | 18.40414123 | 17.76630204 | 0.106274721 | 0.069689791 | 0.637839191 | t-Test | FDR correction BH | 0.012134178 | TRUE | QL | SN | 1.555996905 | 3.116852368 | Paired t-test |
| rdo~Q9VJA9 | 3.762142759 | 0.032841826 | 16.0368359 | 15.4074571 | 0.238253855 | 0.329442831 | 0.6293788 | t-Test | FDR correction BH | 0.0752763 | FALSE | QL | SN | 1.546898781 | 1.483572709 | Paired t-test |
| Sep2~P54359 | 4.353427225 | 0.022390963 | 17.71872689 | 17.08936704 | 0.388346157 | 0.280988397 | 0.62935985 | t-Test | FDR correction BH | 0.058187282 | FALSE | QL | SN | 1.546878463 | 1.649927231 | Paired t-test |
| betaggt-II~Q9XZ68 | 3.583529931 | 0.037193186 | 15.47127811 | 14.84345963 | 0.297266338 | 0.28622075 | 0.627818475 | t-Test | FDR correction BH | 0.080940909 | FALSE | QL | SN | 1.545226661 | 1.429536617 | Paired t-test |
| Dmel\CG13850~Q9VD14 | 4.881610987 | 0.016435228 | 16.17513715 | 15.55643573 | 0.190554886 | 0.333111048 | 0.618701417 | t-Test | FDR correction BH | 0.048039115 | TRUE | QL | SN | 1.535492448 | 1.784224276 | Paired t-test |
| Dmel\CG1309~Q9VZF1 | 14.07384678 | 0.000776953 | 13.7450563 | 13.13545671 | 0.209623977 | 0.237584316 | 0.609599585 | t-Test | FDR correction BH | 0.012134178 | TRUE | QL | SN | 1.52583566 | 3.109605243 | Paired t-test |
| poly~Q9VFW4 | 3.428640759 | 0.041579327 | 15.00829064 | 14.40244736 | 0.273939694 | 0.368637307 | 0.605843281 | t-Test | FDR correction BH | 0.085797756 | FALSE | QL | SN | 1.521868052 | 1.381122548 | Paired t-test |
| Dmel\CG2023~Q9VNI9 | 7.10786166 | 0.005729832 | 18.72095777 | 18.11751731 | 0.087693945 | 0.138858907 | 0.603440461 | t-Test | FDR correction BH | 0.025711613 | TRUE | QL | SN | 1.519335478 | 2.241858096 | Paired t-test |
| CG9798~Q95T98 | 4.022552617 | 0.027599036 | 16.20856008 | 15.60683004 | 0.182381459 | 0.440542539 | 0.60173004 | t-Test | FDR correction BH | 0.066584131 | FALSE | QL | SN | 1.517535262 | 1.559106094 | Paired t-test |
| Aos1~Q7KSM5 | 8.579042548 | 0.003328966 | 19.08940106 | 18.48794983 | 0.137876534 | 0.132663591 | 0.601451227 | t-Test | FDR correction BH | 0.019763199 | TRUE | QL | SN | 1.517242014 | 2.477690661 | Paired t-test |
| aralar1~A0A0B4KHW3 | 3.336311985 | 0.044509451 | 16.36285111 | 15.76358081 | 0.142923536 | 0.323909349 | 0.599270297 | t-Test | FDR correction BH | 0.08986509 | FALSE | QL | SN | 1.514950124 | 1.351547759 | Paired t-test |
| yl~P98163 | 4.031829182 | 0.027432866 | 15.33238603 | 14.74447377 | 0.170640149 | 0.308749295 | 0.587912258 | t-Test | FDR correction BH | 0.066515305 | FALSE | QL | SN | 1.503070061 | 1.561728822 | Paired t-test |
| Dmel\CG13907~Q9W0L6 | 14.29307394 | 0.000742153 | 19.13959427 | 18.55206316 | 0.1208831 | 0.126595406 | 0.587531104 | t-Test | FDR correction BH | 0.012134178 | TRUE | QL | SN | 1.502673009 | 3.12950676 | Paired t-test |
| Mpcp2~Q0E8E8 | 3.49644333 | 0.039582383 | 16.94732534 | 16.36098167 | 0.126944955 | 0.393349029 | 0.58634367 | t-Test | FDR correction BH | 0.08373803 | FALSE | QL | SN | 1.501436718 | 1.40249806 | Paired t-test |
| eIF3l~Q9VTU4 | 3.983318386 | 0.028316335 | 16.59288749 | 16.02339116 | 0.209903725 | 0.337935273 | 0.569496333 | t-Test | FDR correction BH | 0.067831461 | FALSE | QL | SN | 1.484005391 | 1.547962962 | Paired t-test |
| Dmel\CG7556~Q9VWL4 | 4.394108316 | 0.021841358 | 15.48593179 | 14.91922081 | 0.256951736 | 0.068867307 | 0.566710979 | t-Test | FDR correction BH | 0.057280709 | FALSE | QL | SN | 1.481143044 | 1.660720363 | Paired t-test |
| CG10097~Q95T70 | 3.437188075 | 0.041320633 | 14.78979752 | 14.22369574 | 0.372320662 | 0.091312487 | 0.566101782 | t-Test | FDR correction BH | 0.085447912 | FALSE | QL | SN | 1.480517745 | 1.383833037 | Paired t-test |
| RpS2~M9PB84 | 8.341715821 | 0.003611439 | 18.8606279 | 18.29856594 | 0.155283095 | 0.2173103 | 0.562061958 | t-Test | FDR correction BH | 0.020315914 | TRUE | QL | SN | 1.476377808 | 2.442319719 | Paired t-test |
| axo~M9PE65 | 3.872459364 | 0.030476994 | 17.82805323 | 17.27014644 | 0.421065853 | 0.278694013 | 0.557906785 | t-Test | FDR correction BH | 0.071187464 | FALSE | QL | SN | 1.472131742 | 1.516027877 | Paired t-test |
| uex~A0A0B7P9G0 | 5.486506193 | 0.011910703 | 16.61378084 | 16.06983496 | 0.186121608 | 0.06859754 | 0.543945887 | t-Test | FDR correction BH | 0.03880665 | TRUE | QL | SN | 1.457954692 | 1.924062587 | Paired t-test |
| Stam~Q9XTL2 | 4.652234575 | 0.018731987 | 17.14487088 | 16.60148982 | 0.175393329 | 0.047996228 | 0.543381059 | t-Test | FDR correction BH | 0.05198616 | FALSE | QL | SN | 1.457384001 | 1.727416145 | Paired t-test |
| RhoGDI~Q9VW59 | 4.389834206 | 0.021898275 | 16.1124218 | 15.57514855 | 0.260387758 | 0.144309016 | 0.53727325 | t-Test | FDR correction BH | 0.057280709 | FALSE | QL | SN | 1.451227047 | 1.659590098 | Paired t-test |
| Marf~Q7YU24 | 3.222157964 | 0.048504457 | 15.30639961 | 14.78192523 | 0.199182261 | 0.234532663 | 0.524474376 | t-Test | FDR correction BH | 0.095392098 | FALSE | QL | SN | 1.438409422 | 1.314218357 | Paired t-test |
| RpS15Aa~E1JJM9 | 4.196187851 | 0.024691373 | 18.53887715 | 18.02085991 | 0.11136973 | 0.18301157 | 0.51801724 | t-Test | FDR correction BH | 0.062138005 | FALSE | QL | SN | 1.431985853 | 1.60745476 | Paired t-test |
| RpL18~M9PHM6 | 11.40708801 | 0.001445638 | 19.46679011 | 18.95291779 | 0.343811297 | 0.34058048 | 0.513872326 | t-Test | FDR correction BH | 0.015251006 | TRUE | QL | SN | 1.427877611 | 2.839940327 | Paired t-test |
| kermit~Q7JX82 | 4.827183121 | 0.016945525 | 17.02008887 | 16.50625533 | 0.279213536 | 0.113701518 | 0.513833539 | t-Test | FDR correction BH | 0.04872603 | TRUE | QL | SN | 1.427839224 | 1.770944982 | Paired t-test |
| RpS4~X2JGM9 | 5.89403609 | 0.009749069 | 18.32537484 | 17.81431107 | 0.245845848 | 0.216930722 | 0.511063772 | t-Test | FDR correction BH | 0.034358997 | TRUE | QL | SN | 1.425100607 | 2.011036865 | Paired t-test |
| Sc2~Q9VZL3 | 4.125535667 | 0.025824432 | 17.01168745 | 16.50285224 | 0.243111318 | 0.200165188 | 0.508835208 | t-Test | FDR correction BH | 0.06368161 | FALSE | QL | SN | 1.422900921 | 1.587969226 | Paired t-test |
| RpS27~Q9VBU9 | 4.440365751 | 0.021237444 | 18.0595894 | 17.55556089 | 0.158279946 | 0.154826376 | 0.504028507 | t-Test | FDR correction BH | 0.056198087 | FALSE | QL | SN | 1.418168058 | 1.672897749 | Paired t-test |
| Dmel\CG2010~Q8T0K1 | 7.880449749 | 0.004257924 | 15.89078153 | 15.40460202 | 0.101041393 | 0.082279719 | 0.486179514 | t-Test | FDR correction BH | 0.022238926 | TRUE | QL | SN | 1.400730603 | 2.370802132 | Paired t-test |
| RpL3~O16797 | 3.390514046 | 0.04275868 | 19.09644323 | 18.62318895 | 0.197110969 | 0.152122487 | 0.473254279 | t-Test | FDR correction BH | 0.08732638 | FALSE | QL | SN | 1.388237378 | 1.368975715 | Paired t-test |

| | | | | | | | | | | | | | | | | |
|---|---|---|---|---|---|---|---|---|---|---|---|---|---|---|---|---|
| CG6544~A0A0B4LGZ7 | 4.024470665 | 0.027564572 | 17.41571769 | 16.95394713 | 0.260603017 | 0.15274386 | 0.46177056 | t-Test | FDR correction BH | 0.066584131 | FALSE | QL | SN | 1.377231 | 1.55964875 | Paired t-test |
| DnaJ-H~Q9VK35 | 6.556067687 | 0.007216242 | 18.9807031 | 18.52070547 | 0.141677137 | 0.140198513 | 0.459997621 | t-Test | FDR correction BH | 0.029045505 | TRUE | QL | SN | 1.37553955 | 2.141688884 | Paired t-test |
| Rab2~O18333 | 3.546032692 | 0.038198711 | 20.59682893 | 20.14294584 | 0.164856321 | 0.218528578 | 0.453883088 | t-Test | FDR correction BH | 0.082008824 | FALSE | QL | SN | 1.369721977 | 1.417951289 | Paired t-test |
| Fs(2)Ket~M9NDC3 | 3.559572575 | 0.037831671 | 14.98882916 | 14.53612014 | 0.269440202 | 0.21087103 | 0.452709022 | t-Test | FDR correction BH | 0.081661045 | FALSE | QL | SN | 1.36860775 | 1.422144481 | Paired t-test |
| UbcE2M~M9PBW0 | 4.17968627 | 0.024950177 | 15.63335547 | 15.19644099 | 0.147868809 | 0.164149939 | 0.436914487 | t-Test | FDR correction BH | 0.062591546 | FALSE | QL | SN | 1.353706038 | 1.602926373 | Paired t-test |
| ArfGAP3~M9PGG0 | 3.735386401 | 0.033450545 | 18.57057871 | 18.14526634 | 0.167860562 | 0.086642958 | 0.425312373 | t-Test | FDR correction BH | 0.076100982 | FALSE | QL | SN | 1.342863228 | 1.475596797 | Paired t-test |
| scu~O18404 | 5.245067236 | 0.013493267 | 16.82891091 | 16.40758405 | 0.140551424 | 0.116947181 | 0.421326856 | t-Test | FDR correction BH | 0.042396006 | TRUE | QL | SN | 1.339158622 | 1.869882873 | Paired t-test |
| RpL12~Q9W1B9 | 11.15333622 | 0.001544645 | 18.73714595 | 18.33040227 | 0.124021781 | 0.166031074 | 0.406743683 | t-Test | FDR correction BH | 0.015838197 | TRUE | QL | SN | 1.32569021 | 2.81117126 | Paired t-test |
| RpL11~A0A0B4LGZ5 | 3.690173996 | 0.034512376 | 18.84787392 | 18.44589096 | 0.282912924 | 0.213975899 | 0.401982963 | t-Test | FDR correction BH | 0.077108295 | FALSE | QL | SN | 1.321322802 | 1.462025143 | Paired t-test |
| RpL31~Q9V597 | 8.04785214 | 0.004006851 | 16.78989807 | 16.38830379 | 0.288972925 | 0.302416113 | 0.401594281 | t-Test | FDR correction BH | 0.021213554 | TRUE | QL | SN | 1.320966867 | 2.397196822 | Paired t-test |
| comt~M9PH10 | 5.190549585 | 0.0138881 | 14.8722569 | 14.47098778 | 0.417021166 | 0.352116198 | 0.401269123 | t-Test | FDR correction BH | 0.043138422 | TRUE | QL | SN | 1.320669178 | 1.857357171 | Paired t-test |
| RpL13A~Q9VNE9 | 4.272577976 | 0.023537457 | 17.24092188 | 16.84620096 | 0.367929094 | 0.276658509 | 0.394720921 | t-Test | FDR correction BH | 0.060069823 | FALSE | QL | SN | 1.314688419 | 1.628240469 | Paired t-test |
| CCT3~A4V303 | 4.159572851 | 0.025270382 | 17.76740122 | 17.38285684 | 0.101266367 | 0.235926004 | 0.384544386 | t-Test | FDR correction BH | 0.062899559 | FALSE | QL | SN | 1.305447452 | 1.5973882 | Paired t-test |
| Vap33~Q9W4N8 | 4.463975203 | 0.020937538 | 22.63665157 | 22.25774855 | 0.140263384 | 0.160934208 | 0.378903025 | t-Test | FDR correction BH | 0.05549667 | FALSE | QL | SN | 1.300352737 | 1.679074377 | Paired t-test |
| Dmel\CG5973~Q9VM11 | 3.683196732 | 0.034680063 | 15.20085336 | 14.82461645 | 0.163002342 | 0.282101929 | 0.376236905 | t-Test | FDR correction BH | 0.077374425 | FALSE | QL | SN | 1.297951886 | 1.459920127 | Paired t-test |
| CCT7~Q9VHL2 | 3.634308737 | 0.035884723 | 17.73047194 | 17.36352573 | 0.077026208 | 0.167430288 | 0.366946201 | t-Test | FDR correction BH | 0.078874358 | FALSE | QL | SN | 1.289620159 | 1.445090405 | Paired t-test |
| sgg~A4V3W2 | 4.17221116 | 0.025068566 | 16.04556323 | 15.68089589 | 0.411842381 | 0.313289662 | 0.364667342 | t-Test | FDR correction BH | 0.062691091 | FALSE | QL | SN | 1.287584703 | 1.600870517 | Paired t-test |
| RpL30~Q9VJ19 | 3.356642814 | 0.043842269 | 18.01989699 | 17.6599555 | 0.126722914 | 0.103647351 | 0.359941489 | t-Test | FDR correction BH | 0.088969088 | FALSE | QL | SN | 1.283373847 | 1.358106982 | Paired t-test |
| Rpt1~Q7KMQ0 | 4.926505768 | 0.016029267 | 15.87418483 | 15.51533968 | 0.493582807 | 0.594545737 | 0.358845144 | t-Test | FDR correction BH | 0.047390884 | TRUE | QL | SN | 1.282398945 | 1.795086325 | Paired t-test |
| fidipidine~A1ZAC7 | 3.514334682 | 0.039075913 | 16.85573368 | 16.50481667 | 0.210827876 | 0.23953247 | 0.350917012 | t-Test | FDR correction BH | 0.08299724 | FALSE | QL | SN | 1.275371027 | 1.408090862 | Paired t-test |
| Gapdh2~M9PJN8 | 7.417813142 | 0.005069142 | 19.92360654 | 19.58386047 | 0.285406835 | 0.268840232 | 0.339746075 | t-Test | FDR correction BH | 0.024608571 | TRUE | QL | SN | 1.265533831 | 2.295065564 | Paired t-test |
| RpL7~X2J5G6 | 4.272311102 | 0.023541365 | 18.10752586 | 17.77238188 | 0.437612431 | 0.480253408 | 0.335143984 | t-Test | FDR correction BH | 0.060069823 | FALSE | QL | SN | 1.261503303 | 1.628168356 | Paired t-test |
| RpL14~P55841 | 3.599727133 | 0.036769296 | 18.52270178 | 18.18844317 | 0.2783542 | 0.366601894 | 0.334258611 | t-Test | FDR correction BH | 0.080127891 | FALSE | QL | SN | 1.260729364 | 1.434514688 | Paired t-test |
| Rm62~E1JJ68 | 3.400375492 | 0.042449607 | 15.71609551 | 15.38311305 | 0.240187018 | 0.146603049 | 0.332982467 | t-Test | FDR correction BH | 0.086918026 | FALSE | QL | SN | 1.259614672 | 1.372126325 | Paired t-test |
| Dmel\CG6282~M9PET5 | 7.360559946 | 0.005183315 | 22.3876123 | 22.08687229 | 0.273166655 | 0.208747884 | 0.300740007 | t-Test | FDR correction BH | 0.024608571 | TRUE | QL | SN | 1.231776071 | 2.285392385 | Paired t-test |
| Adh~P00334 | 3.348176959 | 0.044118534 | 17.73281498 | 17.43914129 | 0.781011151 | 0.903536919 | 0.293673689 | t-Test | FDR correction BH | 0.089415807 | FALSE | QL | SN | 1.225757585 | 1.355378928 | Paired t-test |
| RpS20~P55828 | 5.757887652 | 0.010409354 | 18.65252375 | 18.35912279 | 0.108659046 | 0.152832502 | 0.293400961 | t-Test | FDR correction BH | 0.035517536 | TRUE | QL | SN | 1.225525889 | 1.982576212 | Paired t-test |
| Gapdh1~P07486 | 3.291361129 | 0.046030911 | 20.69622069 | 20.41101346 | 0.354645389 | 0.28427589 | 0.28520723 | t-Test | FDR correction BH | 0.091888774 | FALSE | QL | SN | 1.21858529 | 1.336950429 | Paired t-test |
| Rpt3~Q9V405 | 3.193828482 | 0.04956546 | 15.69818349 | 15.41811034 | 0.232044806 | 0.361540192 | 0.280073145 | t-Test | FDR correction BH | 0.096880709 | FALSE | QL | SN | 1.214256446 | 1.304820858 | Paired t-test |
| Gga~Q9W329 | 5.529543934 | 0.011654508 | 15.55748331 | 15.28364762 | 0.313623628 | 0.255955819 | 0.273835692 | t-Test | FDR correction BH | 0.038122446 | TRUE | QL | SN | 1.209017973 | 1.933506055 | Paired t-test |
| ine~E1JHT2 | 4.557938757 | 0.01979692 | 16.96111549 | 16.69031014 | 0.11841782 | 0.109288482 | 0.270805346 | t-Test | FDR correction BH | 0.053431096 | FALSE | QL | SN | 1.206481126 | 1.70340238 | Paired t-test |
| Mccc1~Q9V9T5 | 5.870006995 | 0.00986153 | 18.01425317 | 17.74671399 | 0.326047908 | 0.351055701 | 0.267539178 | t-Test | FDR correction BH | 0.034635327 | TRUE | QL | SN | 1.20375282 | 2.006055721 | Paired t-test |
| Hsc70-3~F3YDH0 | 3.751681712 | 0.033078116 | 17.31050609 | 17.06359465 | 0.228350247 | 0.14850051 | 0.246911436 | t-Test | FDR correction BH | 0.075708964 | FALSE | QL | SN | 1.186663949 | 1.480459233 | Paired t-test |
| CCT6~Q9VXQ5 | 3.519466112 | 0.038932178 | 17.22171708 | 16.97756735 | 0.181606895 | 0.260356209 | 0.244149738 | t-Test | FDR correction BH | 0.082972471 | FALSE | QL | SN | 1.184394535 | 1.409691297 | Paired t-test |
| Ran~A4V4A5 | 3.543414886 | 0.038270198 | 18.30118777 | 18.05854264 | 0.128129895 | 0.237249571 | 0.24264513 | t-Test | FDR correction BH | 0.082051717 | FALSE | QL | SN | 1.183159956 | 1.417139287 | Paired t-test |
| ThrRS~Q8IP94 | 4.323659759 | 0.022804504 | 16.2678744 | 16.02821624 | 0.212284236 | 0.200381769 | 0.239658158 | t-Test | FDR correction BH | 0.058791519 | FALSE | QL | SN | 1.180712862 | 1.641979368 | Paired t-test |
| CCT1~A4V391 | 6.019954027 | 0.00918645 | 17.17250935 | 16.93360224 | 0.216059401 | 0.275475563 | 0.23890711 | t-Test | FDR correction BH | 0.033370926 | TRUE | QL | SN | 1.180098358 | 2.036852266 | Paired t-test |
| RpL5~R9Q794 | 3.546052731 | 0.038198165 | 18.04444789 | 17.80663731 | 0.282589661 | 0.205389909 | 0.237810587 | t-Test | FDR correction BH | 0.082008824 | FALSE | QL | SN | 1.179201763 | 1.417957503 | Paired t-test |
| COX4~Q9VIQ8 | 3.43871237 | 0.041274714 | 18.19121211 | 17.95493627 | 0.27538761 | 0.389156196 | 0.236275839 | t-Test | FDR correction BH | 0.085447912 | FALSE | QL | SN | 1.177947988 | 1.384315932 | Paired t-test |
| CCT5~A1Z8U4 | 4.543029625 | 0.019972428 | 17.57970989 | 17.3457393 | 0.136511183 | 0.221569014 | 0.23397059 | t-Test | FDR correction BH | 0.053652745 | FALSE | QL | SN | 1.176067275 | 1.699569139 | Paired t-test |
| ND-B22~Q9VJZ4 | 10.26429789 | 0.001971691 | 16.54909852 | 16.36636554 | 0.186972749 | 0.194398261 | 0.182732984 | t-Test | FDR correction BH | 0.016665632 | TRUE | QL | SN | 1.135032009 | 2.705161197 | Paired t-test |
| CaMKI~Q7JMV3 | 4.618211639 | 0.019107451 | 15.54136567 | 15.35950929 | 0.223681238 | 0.242685788 | 0.181856373 | t-Test | FDR correction BH | 0.05275246 | FALSE | QL | SN | 1.13434255 | 1.718797249 | Paired t-test |
| Scsalpha1~Q94522 | 6.320558841 | 0.008005528 | 19.24216259 | 19.0675758 | 0.219415476 | 0.214066708 | 0.174586785 | t-Test | FDR correction BH | 0.031091532 | TRUE | QL | SN | 1.128641095 | 2.09661003 | Paired t-test |
| Pdha~Q7KVX1 | 3.406968194 | 0.042244564 | 19.44202038 | 19.2734237 | 0.152928464 | 0.09698343 | 0.168596681 | t-Test | FDR correction BH | 0.086721122 | FALSE | QL | SN | 1.123964665 | 1.374229164 | Paired t-test |
| Dmel\CG1907~Q9VAJ9 | 4.935820534 | 0.015946679 | 16.77581226 | 16.6092729 | 0.200343183 | 0.228190684 | 0.166539366 | t-Test | FDR correction BH | 0.047305512 | TRUE | QL | SN | 1.122363009 | 1.797329747 | Paired t-test |
| Kap-alpha3~Q9V455 | 3.725490374 | 0.033679342 | 15.66394015 | 15.54608808 | 0.040527532 | 0.040292941 | 0.117852075 | t-Test | FDR correction BH | 0.076100982 | FALSE | QL | SN | 1.085118105 | 1.472636396 | Paired t-test |
| Hsc70-4~C7LA75 | 3.437688934 | 0.041305537 | 19.39297689 | 19.29314418 | 0.19894166 | 0.15880827 | 0.099832705 | t-Test | FDR correction BH | 0.085447912 | FALSE | QL | SN | 1.071649187 | 1.383991725 | Paired t-test |
| Sod1~P61851 | -4.391318251 | 0.021878491 | 16.05263676 | 16.16044645 | 0.236925731 | 0.212623692 | -0.107809695 | t-Test | FDR correction BH | 0.057280709 | FALSE | QL | SN | 0.92799588 | 1.659982644 | Paired t-test |
| ATPsynbeta~X2JH42 | -3.552248109 | 0.038029662 | 17.90129583 | 18.1150911 | 0.195040677 | 0.111956089 | -0.213795268 | t-Test | FDR correction BH | 0.081977335 | FALSE | QL | SN | 0.8622659 | 1.419877539 | Paired t-test |
| CCT2~Q9W392 | -8.20237945 | 0.003792168 | 17.78968352 | 18.02728704 | 0.21345704 | 0.207130027 | -0.237603523 | t-Test | FDR correction BH | 0.020759188 | TRUE | QL | SN | 0.84815302 | 2.421112433 | Paired t-test |
| Ate1~A0A0B4KFD5 | -3.311651739 | 0.045336097 | 12.99557598 | 13.25320205 | 0.834316254 | 0.743647029 | -0.257626068 | t-Test | FDR correction BH | 0.091029507 | FALSE | QL | SN | 0.836463175 | 1.343555875 | Paired t-test |
| Mlp60A~B7YZP9 | -3.225826265 | 0.048369178 | 21.20479165 | 21.46733065 | 0.302791416 | 0.362401867 | -0.262538995 | t-Test | FDR correction BH | 0.095243635 | FALSE | QL | SN | 0.833619543 | 1.315431292 | Paired t-test |
| CG10517~Q961L8 | -5.395358037 | 0.0124781 | 16.29412824 | 16.58485077 | 0.073602568 | 0.150653695 | -0.290722523 | t-Test | FDR correction BH | 0.040238085 | TRUE | QL | SN | 0.817492544 | 1.903851522 | Paired t-test |
| Jafrac1~X2JF59 | -4.426349796 | 0.021418121 | 19.07926313 | 19.3833046 | 0.028549732 | 0.105139435 | -0.304041473 | t-Test | FDR correction BH | 0.0565822 | FALSE | QL | SN | 0.809980189 | 1.669218634 | Paired t-test |
| HBS1~Q9W074 | -4.08688991 | 0.02647264 | 15.76172228 | 16.06878984 | 0.165460823 | 0.151318719 | -0.307067566 | t-Test | FDR correction BH | 0.064679318 | FALSE | QL | SN | 0.808283014 | 1.577202745 | Paired t-test |
| RpS21~M9PEA6 | -3.376738038 | 0.043195244 | 18.05450161 | 18.37044864 | 0.299169431 | 0.154647282 | -0.315947038 | t-Test | FDR correction BH | 0.087992357 | FALSE | QL | SN | 0.803323487 | 1.364564073 | Paired t-test |
| CHORD~Q9VCC0 | -6.147960876 | 0.008657367 | 16.51292596 | 16.83007013 | 0.210095512 | 0.211030395 | -0.317144178 | t-Test | FDR correction BH | 0.032762891 | TRUE | QL | SN | 0.80265717 | 2.062614192 | Paired t-test |
| Mlp84B~Q24400 | -9.436999278 | 0.002521658 | 20.12607667 | 20.45900766 | 0.330620588 | 0.304505143 | -0.332930994 | t-Test | FDR correction BH | 0.017810152 | TRUE | QL | SN | 0.793921904 | 2.598313892 | Paired t-test |
| pug~A0A0B4K623 | -3.569518234 | 0.037564925 | 16.16403486 | 16.49780526 | 0.326359611 | 0.378628155 | -0.333770393 | t-Test | FDR correction BH | 0.081305605 | FALSE | QL | SN | 0.793460113 | 1.425217473 | Paired t-test |
| yrt~A0T1Z5 | -3.980555817 | 0.028367741 | 15.34137652 | 15.68442607 | 0.285262985 | 0.251979377 | -0.34304955 | t-Test | FDR correction BH | 0.067852569 | FALSE | QL | SN | 0.788373097 | 1.547175249 | Paired t-test |
| wrd~Q8IN90 | -6.605202114 | 0.007064627 | 16.72956031 | 17.08133664 | 0.139433356 | 0.121280995 | -0.351776339 | t-Test | FDR correction BH | 0.028709057 | TRUE | QL | SN | 0.783618662 | 2.150910777 | Paired t-test |
| Dmel\CG9705~M9PFS8 | -3.725672141 | 0.033675122 | 17.85684318 | 18.21906873 | 0.185370589 | 0.141633592 | -0.362225547 | t-Test | FDR correction BH | 0.076100982 | FALSE | QL | SN | 0.777963542 | 1.472690823 | Paired t-test |
| CRMP~Q9V3N7 | -3.269949425 | 0.046778814 | 21.54674199 | 21.91233635 | 0.193355076 | 0.120786202 | -0.365594362 | t-Test | FDR correction BH | 0.092783534 | FALSE | QL | SN | 0.776149051 | 1.329950798 | Paired t-test |
| Dmel\CG40045~M9PG40 | -8.32676085 | 0.003630284 | 17.72000255 | 18.08798435 | 0.1212263 | 0.066506819 | -0.367981798 | t-Test | FDR correction BH | 0.020315914 | TRUE | QL | SN | 0.774865707 | 2.440059445 | Paired t-test |
| Liprin-alpha~M9PC73 | -3.716636066 | 0.033885756 | 14.96728708 | 15.34236282 | 0.051788894 | 0.205956422 | -0.375075739 | t-Test | FDR correction BH | 0.076350791 | FALSE | QL | SN | 0.771064932 | 1.46998282 | Paired t-test |
| cib~O97428 | -3.412507154 | 0.042073267 | 19.48931174 | 19.86907366 | 0.185661205 | 0.086387935 | -0.379761918 | t-Test | FDR correction BH | 0.086508522 | FALSE | QL | SN | 0.768564413 | 1.375993766 | Paired t-test |
| CIAPIN1~Q9V3Y2 | -5.623980146 | 0.011117106 | 19.14589736 | 19.56427174 | 0.14917239 | 0.094879257 | -0.418374381 | t-Test | FDR correction BH | 0.036894895 | TRUE | QL | SN | 0.748267292 | 1.954008262 | Paired t-test |
| CRMP~Q8IPQ2 | -3.458844617 | 0.04067428 | 19.0367673 | 19.46701642 | 0.199428721 | 0.135763884 | -0.43024912 | t-Test | FDR correction BH | 0.084890988 | FALSE | QL | SN | 0.742133625 | 1.390680123 | Paired t-test |
| Dmel\CG9330~Q9VW13 | -7.186635014 | 0.005551641 | 14.84544341 | 15.29915328 | 0.050638781 | 0.077707633 | -0.453709869 | t-Test | FDR correction BH | 0.025267898 | TRUE | QL | SN | 0.730162829 | 2.255578617 | Paired t-test |
| Usp14~Q9VKZ8 | -3.417433626 | 0.041921652 | 16.691691 | 17.15151254 | 0.321386028 | 0.262961768 | -0.459821539 | t-Test | FDR correction BH | 0.086392228 | FALSE | QL | SN | 0.727076192 | 1.377561609 | Paired t-test |
| eRF3~Q9VK85 | -4.02058579 | 0.027634433 | 18.66966133 | 19.1327562 | 0.246498321 | 0.145002313 | -0.463094867 | t-Test | FDR correction BH | 0.066584131 | FALSE | QL | SN | 0.7254284 | 1.558549435 | Paired t-test |
| mop~Q9VUH6 | -5.629948896 | 0.01108424 | 16.05199293 | 16.52613664 | 0.065318962 | 0.153779517 | -0.474143712 | t-Test | FDR correction BH | 0.036862617 | TRUE | QL | SN | 0.719893944 | 1.955294097 | Paired t-test |
| bel~A0A0B4KGU4 | -3.79740201 | 0.032061165 | 17.4728113 | 17.94837423 | 0.35925289 | 0.308545514 | -0.475562928 | t-Test | FDR correction BH | 0.074126901 | FALSE | QL | SN | 0.719186114 | 1.494020707 | Paired t-test |
| nudC~Q9VVA6 | -4.270501941 | 0.023567884 | 18.67851806 | 19.16068395 | 0.228978636 | 0.12885764 | -0.482165887 | t-Test | FDR correction BH | 0.060069823 | FALSE | QL | SN | 0.715902049 | 1.627679403 | Paired t-test |
| Hex-A~Q9W330 | -3.300494466 | 0.045716499 | 19.59108429 | 20.07729746 | 0.103779183 | 0.177203566 | -0.486213168 | t-Test | FDR correction BH | 0.09149043 | FALSE | QL | SN | 0.713896499 | 1.339927039 | Paired t-test |
| eEF5~Q9GU68 | -6.642638064 | 0.006951911 | 17.71751941 | 18.22818131 | 0.186471695 | 0.167790498 | -0.510661896 | t-Test | FDR correction BH | 0.028582883 | TRUE | QL | SN | 0.701900338 | 2.157895783 | Paired t-test |
| Dmel\CG33217~Q8MS38 | -7.96997964 | 0.004121141 | 13.68734432 | 14.2180209 | 0.156732236 | 0.058941871 | -0.530676572 | t-Test | FDR correction BH | 0.021666593 | TRUE | QL | SN | 0.692230027 | 2.38498252 | Paired t-test |
| Dmel\CG7789~Q9VAG9 | -5.061471171 | 0.014884889 | 18.78339231 | 19.31501156 | 0.151741113 | 0.09656707 | -0.531619246 | t-Test | FDR correction BH | 0.045251199 | TRUE | QL | SN | 0.691777864 | 1.8272544 | Paired t-test |
| Uba1~Q8T0L3 | -3.653085119 | 0.035415816 | 15.88034444 | 16.41630863 | 0.218456186 | 0.09214139 | -0.535964187 | t-Test | FDR correction BH | 0.078398494 | FALSE | QL | SN | 0.689697581 | 1.450802748 | Paired t-test |
| Dmel\CG13901~Q9W0M5 | -4.201904773 | 0.02460252 | 15.15919053 | 15.70667668 | 0.273380018 | 0.239426968 | -0.54748615 | t-Test | FDR correction BH | 0.062110641 | FALSE | QL | SN | 0.684211307 | 1.609020408 | Paired t-test |
| Ank2~M9MRX4 | -5.643775518 | 0.011008592 | 19.42083707 | 19.97467146 | 0.24256766 | 0.216168109 | -0.553834388 | t-Test | FDR correction BH | 0.036764543 | TRUE | QL | SN | 0.681207211 | 1.958268227 | Paired t-test |
| p47~Q7K3Z3 | -3.579358643 | 0.03730336 | 17.50968282 | 18.07176116 | 0.267532288 | 0.185550669 | -0.56207834 | t-Test | FDR correction BH | 0.081043602 | FALSE | QL | SN | 0.677325708 | 1.428252047 | Paired t-test |
| Hrb27C~E1JHA4 | -8.818819097 | 0.003072507 | 17.72195179 | 18.29136164 | 0.205880943 | 0.112043343 | -0.569409848 | t-Test | FDR correction BH | 0.019235452 | TRUE | QL | SN | 0.673892396 | 2.512507094 | Paired t-test |
| Dmel\CG17883~A0A0B4KEH8 | -8.916620859 | 0.002975409 | 17.63546858 | 18.21109682 | 0.18287631 | 0.097048936 | -0.575628235 | t-Test | FDR correction BH | 0.019035448 | TRUE | QL | SN | 0.670993997 | 2.526453322 | Paired t-test |
| Hsc70Cb~Q9VUC1 | -5.268860204 | 0.013325545 | 16.72639072 | 17.3055987 | 0.230920536 | 0.042963888 | -0.579207983 | t-Test | FDR correction BH | 0.042034837 | TRUE | QL | SN | 0.669331129 | 1.875315029 | Paired t-test |
| Wdr62~M9PBP2 | -4.504417826 | 0.020436435 | 15.63593121 | 16.22059588 | 0.125583963 | 0.310692363 | -0.584664678 | t-Test | FDR correction BH | 0.054622887 | FALSE | QL | SN | 0.666804304 | 1.689594864 | Paired t-test |
| Mlc1~P06742-2 | -13.55252329 | 0.000868886 | 19.50292877 | 20.10531873 | 0.311157062 | 0.282594667 | -0.602389962 | t-Test | FDR correction BH | 0.01258305 | TRUE | QL | SN | 0.658661915 | 3.061037172 | Paired t-test |
| sta~P38979 | -7.298585132 | 0.005310759 | 19.3120823 | 19.92004557 | 0.171284786 | 0.128004306 | -0.60796327 | t-Test | FDR correction BH | 0.024737406 | TRUE | QL | SN | 0.656122332 | 2.274843442 | Paired t-test |
| Cortactin~Q9VDF4 | -11.48084378 | 0.001418447 | 17.82346856 | 18.44326496 | 0.141176668 | 0.230139533 | -0.619796398 | t-Test | FDR correction BH | 0.015251006 | TRUE | QL | SN | 0.650762761 | 2.84818674 | Paired t-test |
| Tps1~Q9Y119 | -3.933152368 | 0.02926883 | 18.34861972 | 18.97688618 | 0.155812704 | 0.190234925 | -0.628266454 | t-Test | FDR correction BH | 0.068933059 | FALSE | QL | SN | 0.646953329 | 1.533594639 | Paired t-test |
| capt~Q9VPX7 | -6.147646325 | 0.008658617 | 19.09403541 | 19.72289221 | 0.178042971 | 0.101382279 | -0.628856798 | t-Test | FDR correction BH | 0.032762891 | TRUE | QL | SN | 0.646688653 | 2.062551469 | Paired t-test |
| pAbp~P21187 | -4.321584661 | 0.022833699 | 16.7424211 | 17.37778682 | 0.306340503 | 0.094566993 | -0.635365719 | t-Test | FDR correction BH | 0.058791519 | FALSE | QL | SN | 0.643777598 | 1.641423726 | Paired t-test |
| UGP~E1JI91 | -6.650684583 | 0.006927994 | 17.52146939 | 18.16002783 | 0.334104172 | 0.227880141 | -0.638558442 | t-Test | FDR correction BH | 0.028582883 | TRUE | QL | SN | 0.642354476 | 2.159392528 | Paired t-test |
| Tm2~A0A0B4KHJ9 | -5.790623303 | 0.010245369 | 17.91897728 | 18.57179088 | 0.480234434 | 0.475824282 | -0.6528136 | t-Test | FDR correction BH | 0.035326565 | TRUE | QL | SN | 0.636038676 | 1.989472388 | Paired t-test |
| Gclm~Q9VCW6 | -4.102786008 | 0.026203503 | 16.28155893 | 16.93707836 | 0.276472396 | 0.090473327 | -0.655519428 | t-Test | FDR correction BH | 0.064317688 | FALSE | QL | SN | 0.63484688 | 1.581640652 | Paired t-test |
| Dmel\CG9674~Q9VVA4 | -7.625823367 | 0.00468121 | 17.69111439 | 18.36048892 | 0.245404382 | 0.137083439 | -0.669374531 | t-Test | FDR correction BH | 0.023450212 | TRUE | QL | SN | 0.62877923 | 2.329641887 | Paired t-test |
| Dmel21.3~Q8IQW5 | -4.97723047 | 0.015586156 | 20.1020559 | 20.78286194 | 0.123707723 | 0.192820273 | -0.680806037 | t-Test | FDR correction BH | 0.046408873 | TRUE | QL | SN | 0.623816649 | 1.80726097 | Paired t-test |
| 34F4T~Q9VTK9 | -10.75666605 | 0.001718256 | 16.84203396 | 17.53887294 | 0.146801829 | 0.062618136 | -0.696838984 | t-Test | FDR correction BH | 0.016207074 | TRUE | QL | SN | 0.616922435 | 2.764912238 | Paired t-test |
| AGO1~Q7KY08 | -3.566096489 | 0.037656426 | 15.83133137 | 16.529247 | 0.352536922 | 0.667824625 | -0.697915631 | t-Test | FDR correction BH | 0.081393061 | FALSE | QL | SN | 0.616462213 | 1.424160906 | Paired t-test |
| EG:100G10.2~Q9W4W4 | -4.913650862 | 0.016144161 | 18.04867986 | 18.75239433 | 0.136645273 | 0.225863367 | -0.703714472 | t-Test | FDR correction BH | 0.04758938 | TRUE | QL | SN | 0.613989347 | 1.791984518 | Paired t-test |
| Dmel\CG17816~A8JQV2 | -4.925866355 | 0.016034957 | 17.86735645 | 18.57275349 | 0.352200493 | 0.116602286 | -0.70539704 | t-Test | FDR correction BH | 0.047390884 | TRUE | QL | SN | 0.613273688 | 1.794932197 | Paired t-test |
| Dmel\CG9643~Q9VQJ8 | -8.571673597 | 0.003337288 | 15.12429187 | 15.83751321 | 0.060190194 | 0.093952798 | -0.713221342 | t-Test | FDR correction BH | 0.019763199 | TRUE | QL | SN | 0.609956667 | 2.476606259 | Paired t-test |
| Fim~O61604 | -4.497304583 | 0.020523436 | 17.50752939 | 18.2271133 | 0.389350132 | 0.144504437 | -0.719583912 | t-Test | FDR correction BH | 0.054671962 | FALSE | QL | SN | 0.60727256 | 1.687749928 | Paired t-test |
| DCTN2-p50~Q7K2D2 | -4.523268921 | 0.020208171 | 14.36675634 | 15.09448457 | 0.058037754 | 0.278571916 | -0.727728227 | t-Test | FDR correction BH | 0.054126252 | FALSE | QL | SN | 0.603854038 | 1.694473002 | Paired t-test |
| Dmel\CG15717~Q9VYG8 | -3.226661175 | 0.048338455 | 17.34716747 | 18.07758644 | 0.282385558 | 0.467215449 | -0.730418975 | t-Test | FDR correction BH | 0.095243635 | FALSE | QL | SN | 0.602728849 | 1.315707232 | Paired t-test |
| Dmel\CG5023~Q9I7J0 | -4.490356008 | 0.020608887 | 19.40070385 | 20.13528179 | 0.092849731 | 0.209113364 | -0.73457794 | t-Test | FDR correction BH | 0.054807942 | FALSE | QL | SN | 0.600993819 | 1.685945458 | Paired t-test |
| rhea~Q9VSL8 | -4.803835782 | 0.017170738 | 17.67920993 | 18.43200981 | 0.169755872 | 0.230311213 | -0.752799881 | t-Test | FDR correction BH | 0.049047133 | TRUE | QL | SN | 0.593450712 | 1.765211045 | Paired t-test |
| alt~Q9VEH0 | -5.024548551 | 0.015187083 | 15.84901203 | 16.63272499 | 0.262383562 | 0.144492949 | -0.78371296 | t-Test | FDR correction BH | 0.045856543 | TRUE | QL | SN | 0.580869925 | 1.818525622 | Paired t-test |
| futsch~M9NGG5 | -7.853017182 | 0.004301033 | 20.8864807 | 21.67623204 | 0.291410049 | 0.145185314 | -0.789751342 | t-Test | FDR correction BH | 0.022317736 | TRUE | QL | SN | 0.578443782 | 2.36642727 | Paired t-test |
| Idh~Q7KUB0 | -11.88600745 | 0.001280576 | 18.93272305 | 19.72254095 | 0.098486799 | 0.092411435 | -0.789817892 | t-Test | FDR correction BH | 0.014467786 | TRUE | QL | SN | 0.5784171 | 2.89259458 | Paired t-test |
| cact~M9PD95 | -4.55814651 | 0.019794488 | 18.7681902 | 19.5604246 | 0.441228613 | 0.335416628 | -0.792234403 | t-Test | FDR correction BH | 0.053431096 | FALSE | QL | SN | 0.577449063 | 1.703455724 | Paired t-test |
| twin~Q9VCB6 | -17.60541548 | 0.000399494 | 13.43757297 | 14.23663415 | 0.272227832 | 0.246471559 | -0.799061178 | t-Test | FDR correction BH | 0.010461282 | TRUE | QL | SN | 0.574723052 | 3.39848944 | Paired t-test |
| Cat~P17336 | -4.209948429 | 0.024478202 | 18.64076747 | 19.44555777 | 0.267730258 | 0.194467729 | -0.804790305 | t-Test | FDR correction BH | 0.061894883 | FALSE | QL | SN | 0.572445279 | 1.611220483 | Paired t-test |
| Pax~Q9VIX2 | -7.588431598 | 0.004747982 | 16.87269689 | 17.68021857 | 0.122359401 | 0.128772019 | -0.807521682 | t-Test | FDR correction BH | 0.02371014 | TRUE | QL | SN | 0.571362524 | 2.323490978 | Paired t-test |
| Lmpt~Q8IQQ3 | -4.666356603 | 0.018578956 | 14.49256576 | 15.30042839 | 0.10436617 | 0.353115273 | -0.807862639 | t-Test | FDR correction BH | 0.051832359 | FALSE | QL | SN | 0.571227508 | 1.730978691 | Paired t-test |
| Vha55~E1JIJ5 | -8.634185019 | 0.00326755 | 20.63069512 | 21.4531246 | 0.300000172 | 0.178664297 | -0.822429489 | t-Test | FDR correction BH | 0.019626282 | TRUE | QL | SN | 0.565488861 | 2.485777718 | Paired t-test |
| chico~M9PCQ8 | -3.615499814 | 0.036362427 | 15.3971344 | 16.22034097 | 0.392082475 | 0.266542861 | -0.823206575 | t-Test | FDR correction BH | 0.079567782 | FALSE | QL | SN | 0.565184351 | 1.439347143 | Paired t-test |
| Dmel21.3~Q9VWG1 | -27.68701252 | 0.000103421 | 19.55312709 | 20.37721443 | 0.206718253 | 0.165731059 | -0.824087346 | t-Test | FDR correction BH | 0.006717766 | TRUE | QL | SN | 0.564839409 | 3.985393343 | Paired t-test |
| Dmel\CG5381~Q9VKZ6 | -12.67089406 | 0.001060219 | 15.37543668 | 16.20820862 | 0.333782093 | 0.297416997 | -0.832771937 | t-Test | FDR correction BH | 0.013092474 | TRUE | QL | SN | 0.561449459 | 2.974604333 | Paired t-test |
| GstE12~Q9XYZ9 | -8.834337304 | 0.003056824 | 17.86219273 | 18.69657298 | 0.219183357 | 0.062946409 | -0.834380248 | t-Test | FDR correction BH | 0.019235452 | TRUE | QL | SN | 0.560823906 | 2.514729629 | Paired t-test |
| gw~Q8SY33-2 | -7.526071295 | 0.004862164 | 16.32083558 | 17.15675648 | 0.110368479 | 0.173114991 | -0.8359209 | t-Test | FDR correction BH | 0.024066801 | TRUE | QL | SN | 0.560225322 | 2.313170401 | Paired t-test |
| cher~B7Z0L2 | -3.926098626 | 0.029406044 | 13.47876513 | 14.31923632 | 0.437440903 | 0.226803705 | -0.840471195 | t-Test | FDR correction BH | 0.068933059 | FALSE | QL | SN | 0.558461142 | 1.531563399 | Paired t-test |
| Dmel\CG6106~Q9VWW1 | -3.633821616 | 0.035896993 | 15.39125117 | 16.23584236 | 0.13511102 | 0.440943755 | -0.844591192 | t-Test | FDR correction BH | 0.078874358 | FALSE | QL | SN | 0.556868583 | 1.44494193 | Paired t-test |

| | | | | | | | | | | | | | | | | |
|---|---|---|---|---|---|---|---|---|---|---|---|---|---|---|---|---|
| Dhx15~Q7K3M5 | -5.896702112 | 0.009736695 | 14.95352339 | 15.8063416 | 0.125798727 | 0.35682492 | -0.852818206 | t-Test | FDR correction BH | 0.034358997 | TRUE | QL | SN | 0.55370206 | 2.011588428 | Paired t-test |
| Zw~P12646 | -6.985670863 | 0.006021451 | 16.75048295 | 17.60658393 | 0.209069537 | 0.099271548 | -0.85610098 | t-Test | FDR correction BH | 0.026555929 | TRUE | QL | SN | 0.552443573 | 2.220298854 | Paired t-test |
| Zasp52~G3JX25 | -3.674012802 | 0.034902375 | 15.23893559 | 16.10623802 | 0.084428048 | 0.332903791 | -0.867302427 | t-Test | FDR correction BH | 0.07765291 | FALSE | QL | SN | 0.548170871 | 1.457145016 | Paired t-test |
| Rel~Q94527-2 | -5.963700261 | 0.009432389 | 16.59985612 | 17.47267201 | 0.483546653 | 0.448375699 | -0.872815894 | t-Test | FDR correction BH | 0.033995013 | TRUE | QL | SN | 0.546079955 | 2.025378299 | Paired t-test |
| Dp1~Q7KN75 | -7.774865861 | 0.004427046 | 17.90903295 | 18.78332503 | 0.192606785 | 0.138973267 | -0.874292081 | t-Test | FDR correction BH | 0.022531262 | TRUE | QL | SN | 0.545521484 | 2.35388592 | Paired t-test |
| san~Q9NHD5 | -4.096527624 | 0.026309041 | 13.98559852 | 14.86248316 | 0.498081285 | 0.192281431 | -0.876884643 | t-Test | FDR correction BH | 0.064477387 | FALSE | QL | SN | 0.544542047 | 1.579894987 | Paired t-test |
| Lmpt~M9MRX0 | -6.953878982 | 0.006100517 | 17.13915186 | 18.0221031 | 0.309715921 | 0.133216763 | -0.88295124 | t-Test | FDR correction BH | 0.026651081 | TRUE | QL | SN | 0.542257031 | 2.214633368 | Paired t-test |
| Msp300~M9MRD1 | -6.284013774 | 0.008138021 | 17.77365108 | 18.65913821 | 0.108333763 | 0.149924119 | -0.885487137 | t-Test | FDR correction BH | 0.031389509 | TRUE | QL | SN | 0.541304716 | 2.089481207 | Paired t-test |
| Reps~M9NDL7 | -7.294261332 | 0.005319803 | 14.9705751 | 15.85698574 | 0.333776466 | 0.296420615 | -0.886410641 | t-Test | FDR correction BH | 0.024737406 | TRUE | QL | SN | 0.540958325 | 2.274104417 | Paired t-test |
| ATPCL~E2QCF1 | -6.713943468 | 0.006743681 | 17.94216546 | 18.82941389 | 0.176929659 | 0.105665574 | -0.887248426 | t-Test | FDR correction BH | 0.028247427 | TRUE | QL | SN | 0.540644277 | 2.171102995 | Paired t-test |
| Abp1~Q9VU84 | -4.859041828 | 0.016644368 | 17.30249965 | 18.18999003 | 0.230886598 | 0.179264695 | -0.887490387 | t-Test | FDR correction BH | 0.048228926 | TRUE | QL | SN | 0.540553611 | 1.77873269 | Paired t-test |
| TTLL4B~Q960F9 | -3.505505898 | 0.039324801 | 13.75744062 | 14.6450152 | 0.07135608 | 0.47101996 | -0.887574582 | t-Test | FDR correction BH | 0.083303733 | FALSE | QL | SN | 0.540522065 | 1.40533347 | Paired t-test |
| Cen~Q9VIK6 | -3.203096301 | 0.049215159 | 13.59256243 | 14.48899971 | 0.735969074 | 0.352639254 | -0.896437274 | t-Test | FDR correction BH | 0.096367398 | FALSE | QL | SN | 0.537211736 | 1.307901104 | Paired t-test |
| mal~X2JFY7 | -5.246268744 | 0.013484732 | 16.26427269 | 17.16634797 | 0.144776372 | 0.319804557 | -0.902075279 | t-Test | FDR correction BH | 0.042396006 | TRUE | QL | SN | 0.535116426 | 1.870157687 | Paired t-test |
| CG14576~Q9VP02 | -5.866381876 | 0.009878643 | 16.55067815 | 17.45490802 | 0.281727785 | 0.106437197 | -0.904229864 | t-Test | FDR correction BH | 0.034635327 | TRUE | QL | SN | 0.534317856 | 2.005302702 | Paired t-test |
| Hil~Q0E908 | -3.77559481 | 0.032541145 | 13.82116708 | 14.72778775 | 0.649046 | 0.455249544 | -0.906620667 | t-Test | FDR correction BH | 0.075018876 | FALSE | QL | SN | 0.533433129 | 1.487567173 | Paired t-test |
| RpL22~P50887 | -8.305266359 | 0.003657597 | 16.68804427 | 17.59879493 | 0.344322775 | 0.167955476 | -0.910750655 | t-Test | FDR correction BH | 0.020372559 | TRUE | QL | SN | 0.53190826 | 2.436804149 | Paired t-test |
| mxt~M9PC59 | -9.132385363 | 0.002775124 | 14.95279906 | 15.86792983 | 0.296328918 | 0.201013255 | -0.915130779 | t-Test | FDR correction BH | 0.018574671 | TRUE | QL | SN | 0.530295798 | 2.556717682 | Paired t-test |
| eRF1~M9PD27 | -6.637809394 | 0.006966316 | 16.74329986 | 17.66288222 | 0.174535273 | 0.249438485 | -0.919582365 | t-Test | FDR correction BH | 0.028582883 | TRUE | QL | SN | 0.528662037 | 2.156996813 | Paired t-test |
| eIF4G1~O61380 | -5.185344999 | 0.013926577 | 17.35146655 | 18.27257005 | 0.305561843 | 0.211078882 | -0.921103491 | t-Test | FDR correction BH | 0.043148094 | TRUE | QL | SN | 0.528104928 | 1.856155607 | Paired t-test |
| anon-EST:fe2A9~Q59DP4 | -8.218768856 | 0.003770298 | 16.67609325 | 17.60269915 | 0.156429652 | 0.198944571 | -0.926605899 | t-Test | FDR correction BH | 0.020710638 | TRUE | QL | SN | 0.526094583 | 2.423624298 | Paired t-test |
| Ge-1~Q9VKK1 | -8.043255508 | 0.00401348 | 15.38497005 | 16.32062108 | 0.152601195 | 0.064436189 | -0.935651037 | t-Test | FDR correction BH | 0.021213554 | TRUE | QL | SN | 0.522806493 | 2.396478896 | Paired t-test |
| eIF3d1~Q9VCK0 | -3.186631462 | 0.049839665 | 15.6264213 | 16.56889577 | 0.67535094 | 0.201857645 | -0.942474473 | t-Test | FDR correction BH | 0.097252442 | FALSE | QL | SN | 0.520339642 | 1.302424885 | Paired t-test |
| vig2~Q9VBX3 | -6.120176623 | 0.008768738 | 15.04491634 | 15.99075313 | 0.292574031 | 0.555100374 | -0.945836791 | t-Test | FDR correction BH | 0.033100945 | TRUE | QL | SN | 0.51912836 | 2.05706293 | Paired t-test |
| Tm1~P06754-1 | -10.55983186 | 0.001814069 | 17.33081306 | 18.28395164 | 0.307828141 | 0.408310515 | -0.953138582 | t-Test | FDR correction BH | 0.016234895 | TRUE | QL | SN | 0.516507577 | 2.741346277 | Paired t-test |
| eIF4E1~M9PBZ9 | -3.732104565 | 0.033526201 | 17.42683254 | 18.38750201 | 0.284893221 | 0.231455627 | -0.960669475 | t-Test | FDR correction BH | 0.076100982 | FALSE | QL | SN | 0.513818423 | 1.474615662 | Paired t-test |
| Jupiter~Q9I7K0-5 | -14.6633458 | 0.000687935 | 15.17892505 | 16.14085268 | 0.163238178 | 0.125526836 | -0.961927634 | t-Test | FDR correction BH | 0.012134178 | TRUE | QL | SN | 0.513370523 | 3.162452527 | Paired t-test |
| hpo~A0A0B4KFW0 | -8.779846489 | 0.003112367 | 13.66615989 | 14.63479208 | 0.363934808 | 0.363049182 | -0.968632189 | t-Test | FDR correction BH | 0.019235452 | TRUE | QL | SN | 0.5109903 | 2.50690913 | Paired t-test |
| Dmel\CG11835~Q9VPR2 | -5.760407304 | 0.010396611 | 16.96776353 | 17.93849091 | 0.368216648 | 0.175903328 | -0.970727388 | t-Test | FDR correction BH | 0.035517536 | TRUE | QL | SN | 0.510248737 | 1.983108216 | Paired t-test |
| Strn-Mlck~A1ZA73 | -14.76131106 | 0.000674477 | 18.87536004 | 19.84997369 | 0.260404984 | 0.181859218 | -0.974613649 | t-Test | FDR correction BH | 0.012134178 | TRUE | QL | SN | 0.508876104 | 3.171033138 | Paired t-test |
| su(r)~Q9W374 | -3.479528741 | 0.040068918 | 15.58788102 | 16.56328028 | 0.59502902 | 0.161868911 | -0.975399259 | t-Test | FDR correction BH | 0.084542764 | FALSE | QL | SN | 0.508599074 | 1.397192384 | Paired t-test |
| Mp20~P14318 | -9.643890643 | 0.002366758 | 19.27339439 | 20.25347467 | 0.130882737 | 0.281189138 | -0.980080278 | t-Test | FDR correction BH | 0.017215729 | TRUE | QL | SN | 0.50695153 | 2.625846234 | Paired t-test |
| Dmel\CG8635~A0A0B4LEV3 | -5.609096999 | 0.011199616 | 16.74238184 | 17.72612627 | 0.328008144 | 0.033644837 | -0.98374443 | t-Test | FDR correction BH | 0.037091451 | TRUE | QL | SN | 0.50566561 | 1.950796882 | Paired t-test |
| NUAK~Q9VH03 | -3.577894747 | 0.037342124 | 14.54348103 | 15.55266725 | 0.717466163 | 0.524498662 | -1.009186225 | t-Test | FDR correction BH | 0.081043602 | FALSE | QL | SN | 0.496826411 | 1.427800983 | Paired t-test |
| kis~Q9VPL6 | -11.11134786 | 0.001561888 | 13.65424918 | 14.67127163 | 0.045080405 | 0.136602059 | -1.01702245 | t-Test | FDR correction BH | 0.015838197 | TRUE | QL | SN | 0.494135136 | 2.806350154 | Paired t-test |
| Unc-89~A8DYP0-3 | -10.66242568 | 0.001763269 | 18.02583515 | 19.0437592 | 0.192204751 | 0.0313633 | -1.017924057 | t-Test | FDR correction BH | 0.016234815 | TRUE | QL | SN | 0.493826425 | 2.753681376 | Paired t-test |
| fmt~M9PEB1 | -6.74331242 | 0.006660303 | 15.68753351 | 16.71702518 | 0.307469587 | 0.256624169 | -1.029491671 | t-Test | FDR correction BH | 0.027994359 | TRUE | QL | SN | 0.489882727 | 2.176506033 | Paired t-test |
| Dera~Q7K209 | -6.642794466 | 0.006951445 | 15.49314799 | 16.53919245 | 0.386834821 | 0.432252258 | -1.04604446 | t-Test | FDR correction BH | 0.028582883 | TRUE | QL | SN | 0.48429417 | 2.157924891 | Paired t-test |
| Pfrx~Q0KHQ6 | -4.408624485 | 0.021649475 | 18.13624775 | 19.18508816 | 0.266030041 | 0.152075014 | -1.048840407 | t-Test | FDR correction BH | 0.056910254 | FALSE | QL | SN | 0.483356515 | 1.664552629 | Paired t-test |
| CAP~A0A0B4K7L3 | -10.39925527 | 0.001897547 | 18.57517974 | 19.63413008 | 0.133619594 | 0.229365086 | -1.058950333 | t-Test | FDR correction BH | 0.01648088 | TRUE | QL | SN | 0.479981154 | 2.721807372 | Paired t-test |
| REG~Q9V3P3 | -3.362550997 | 0.043650767 | 15.24808798 | 16.32039243 | 0.237282615 | 0.531893957 | -1.072304448 | t-Test | FDR correction BH | 0.088806734 | FALSE | QL | SN | 0.475558772 | 1.360008116 | Paired t-test |
| Lasp~M9NDA6 | -4.673397598 | 0.018503264 | 15.5341699 | 16.61443627 | 0.109448653 | 0.373887598 | -1.080266362 | t-Test | FDR correction BH | 0.051711753 | FALSE | QL | SN | 0.472941497 | 1.73275166 | Paired t-test |
| Mtap~A0A0B4KG54 | -8.037602127 | 0.004021653 | 17.47638733 | 18.55866787 | 0.139471969 | 0.204957819 | -1.082280543 | t-Test | FDR correction BH | 0.021213554 | TRUE | QL | SN | 0.472281673 | 2.395595398 | Paired t-test |
| RanBPM~A0A0B4KFL0 | -10.51095568 | 0.001838953 | 14.32263964 | 15.41056911 | 0.158617724 | 0.208450296 | -1.087929477 | t-Test | FDR correction BH | 0.016365656 | TRUE | QL | SN | 0.47043605 | 2.735429333 | Paired t-test |
| NAT1~A1Z968 | -4.906997851 | 0.016204045 | 15.6839045 | 16.77426243 | 0.34952937 | 0.530522497 | -1.090357924 | t-Test | FDR correction BH | 0.047625542 | TRUE | QL | SN | 0.469644844 | 1.790376558 | Paired t-test |
| EMSY~Q9VQ58 | -3.726984946 | 0.03364466 | 13.23159382 | 14.32512587 | 0.413355003 | 0.227193074 | -1.093532047 | t-Test | FDR correction BH | 0.076100982 | FALSE | QL | SN | 0.468612698 | 1.473083861 | Paired t-test |
| eIF4B~Q7PLL1 | -4.331354478 | 0.022696666 | 16.6443582 | 17.7388155 | 0.455111263 | 0.189597632 | -1.094457303 | t-Test | FDR correction BH | 0.058768719 | FALSE | QL | SN | 0.468312255 | 1.644037939 | Paired t-test |
| Gclc~A4V448 | -9.727024801 | 0.002308066 | 15.86847217 | 16.96364298 | 0.281536833 | 0.16611928 | -1.095170809 | t-Test | FDR correction BH | 0.017112931 | TRUE | QL | SN | 0.468080702 | 2.636751718 | Paired t-test |
| Kank~Q7KMM8 | -4.886440116 | 0.016390923 | 14.71597952 | 15.82572471 | 0.470939721 | 0.436662757 | -1.109745194 | t-Test | FDR correction BH | 0.047997683 | TRUE | QL | SN | 0.463375864 | 1.785396599 | Paired t-test |
| CG6108~A8DZ00 | -4.035723169 | 0.027363497 | 15.24101543 | 16.35602798 | 0.442243864 | 0.371562952 | -1.115012547 | t-Test | FDR correction BH | 0.066448249 | FALSE | QL | SN | 0.46168714 | 1.562828398 | Paired t-test |
| AGBE~A1Z992 | -3.652325029 | 0.035434645 | 17.80462711 | 18.91997853 | 1.41913443 | 1.449720175 | -1.115351423 | t-Test | FDR correction BH | 0.078398494 | FALSE | QL | SN | 0.461578707 | 1.450571912 | Paired t-test |
| Dys~Q9VDW6-1 | -3.727558748 | 0.033631356 | 15.93816971 | 17.06420553 | 0.526059744 | 0.268715915 | -1.126035826 | t-Test | FDR correction BH | 0.076100982 | FALSE | QL | SN | 0.458172944 | 1.473255619 | Paired t-test |
| CysRS~Q7KN90 | -3.966360791 | 0.028633785 | 14.75234326 | 15.883959 | 0.6932753 | 0.307418751 | -1.131615737 | t-Test | FDR correction BH | 0.068135045 | FALSE | QL | SN | 0.456404292 | 1.543121246 | Paired t-test |
| Gart~X2J516 | -4.555995735 | 0.019819679 | 13.81229954 | 14.95026585 | 0.275632895 | 0.404506192 | -1.137966308 | t-Test | FDR correction BH | 0.053431096 | FALSE | QL | SN | 0.45439967 | 1.702903385 | Paired t-test |
| cno~A0A0B4KFK9 | -3.736423104 | 0.033426692 | 14.3422529 | 15.48918224 | 0.167571706 | 0.475474824 | -1.14692934 | t-Test | FDR correction BH | 0.076100982 | FALSE | QL | SN | 0.451585372 | 1.475906599 | Paired t-test |
| eIF3g1~Q9W4X7 | -6.635077964 | 0.006974482 | 17.46000901 | 18.61696412 | 0.3359372 | 0.259957035 | -1.156955104 | t-Test | FDR correction BH | 0.028582883 | TRUE | QL | SN | 0.448458035 | 2.156488033 | Paired t-test |
| Bx42~M9PGQ6 | -5.21657567 | 0.013697757 | 13.21734876 | 14.37978808 | 0.396434949 | 0.202144144 | -1.162439326 | t-Test | FDR correction BH | 0.042785346 | TRUE | QL | SN | 0.446756515 | 1.863350552 | Paired t-test |
| Mlc2~P18432 | -4.005737417 | 0.027903561 | 16.73785738 | 17.90384397 | 0.31225979 | 0.302704967 | -1.165986593 | t-Test | FDR correction BH | 0.067044304 | FALSE | QL | SN | 0.445659389 | 1.554340362 | Paired t-test |
| B52~A0A0C4DHG5 | -3.284275984 | 0.046276704 | 13.36336141 | 14.5334334 | 0.524462062 | 0.449542635 | -1.170071992 | t-Test | FDR correction BH | 0.092263817 | FALSE | QL | SN | 0.444399164 | 1.334637577 | Paired t-test |
| sqd~Q08473-2 | -14.72306365 | 0.000679689 | 17.15432953 | 18.33171122 | 0.299488602 | 0.313319161 | -1.177381688 | t-Test | FDR correction BH | 0.012134178 | TRUE | QL | SN | 0.442153223 | 3.167689759 | Paired t-test |
| Pu~P48596 | -8.244223932 | 0.003736661 | 18.02171336 | 19.20088797 | 0.414396132 | 0.240681089 | -1.179174613 | t-Test | FDR correction BH | 0.02059782 | TRUE | QL | SN | 0.441604074 | 2.427516339 | Paired t-test |
| eEF1delta~Q9VL18 | -7.669592102 | 0.004604616 | 14.49867217 | 15.67847252 | 0.394869368 | 0.462898557 | -1.179800344 | t-Test | FDR correction BH | 0.023212511 | TRUE | QL | SN | 0.441412582 | 2.336806579 | Paired t-test |
| eas~P54352-2 | -6.46541987 | 0.007507377 | 14.44466125 | 15.62556151 | 0.326643098 | 0.03806288 | -1.180900253 | t-Test | FDR correction BH | 0.029675561 | TRUE | QL | SN | 0.441076177 | 2.124511787 | Paired t-test |
| sls~M9PBJ0 | -8.771968018 | 0.003120508 | 17.12938034 | 18.31586331 | 0.151628408 | 0.231193619 | -1.186482961 | t-Test | FDR correction BH | 0.019235452 | TRUE | QL | SN | 0.43937267 | 2.50577464 | Paired t-test |
| cher~A0A0B4JCY6 | -4.247460877 | 0.023909027 | 16.1597316 | 17.35060003 | 0.560133969 | 0.121123579 | -1.190868431 | t-Test | FDR correction BH | 0.060821719 | FALSE | QL | SN | 0.438039103 | 1.621438094 | Paired t-test |
| RpS10b~M9NEQ9 | -5.22774064 | 0.013617145 | 16.29205372 | 17.48966181 | 0.342631663 | 0.216543418 | -1.197608091 | t-Test | FDR correction BH | 0.042617115 | TRUE | QL | SN | 0.435997543 | 1.865913944 | Paired t-test |
| Tlk~M9PDQ2 | -4.756508543 | 0.017639343 | 13.21944169 | 14.42007171 | 0.419564511 | 0.092085346 | -1.200630016 | t-Test | FDR correction BH | 0.050177632 | FALSE | QL | SN | 0.435085241 | 1.753517585 | Paired t-test |
| eIF3a~Q9VN25 | -4.886722082 | 0.016388341 | 16.02347603 | 17.22786932 | 0.538322931 | 0.414479551 | -1.204393287 | t-Test | FDR correction BH | 0.047997683 | TRUE | QL | SN | 0.4339518 | 1.78546502 | Paired t-test |
| Larp4B~Q9I7T7-2 | -7.312009951 | 0.005282804 | 14.32201067 | 15.53509613 | 0.276302217 | 0.414828872 | -1.213085462 | t-Test | FDR correction BH | 0.024737406 | TRUE | QL | SN | 0.43134512 | 2.277135466 | Paired t-test |
| Lsp1alpha~P11995 | -4.287721581 | 0.02331703 | 14.0829284 | 15.29689346 | 0.197187488 | 0.422817715 | -1.213965062 | t-Test | FDR correction BH | 0.059717089 | FALSE | QL | SN | 0.431082212 | 1.632326761 | Paired t-test |
| Msp300~M9MRJ4 | -5.178632653 | 0.013976407 | 16.20573203 | 17.42043971 | 0.326078659 | 0.150144143 | -1.214707678 | t-Test | FDR correction BH | 0.043148094 | TRUE | QL | SN | 0.430860373 | 1.854604454 | Paired t-test |
| Atx2~Q8SWR8 | -10.32571567 | 0.001937481 | 16.33021804 | 17.54845996 | 0.238273437 | 0.229156904 | -1.218241918 | t-Test | FDR correction BH | 0.016614137 | TRUE | QL | SN | 0.429806165 | 2.712762502 | Paired t-test |
| Pkcdelta~Q9VYN1 | -7.85696207 | 0.004294798 | 13.7692695 | 14.98907679 | 0.406824355 | 0.305717785 | -1.219807284 | t-Test | FDR correction BH | 0.022317736 | TRUE | QL | SN | 0.429340066 | 2.367057259 | Paired t-test |
| tral~M9PFB2 | -3.694425386 | 0.034410709 | 15.62474019 | 16.84619714 | 0.530582245 | 0.389359406 | -1.221456947 | t-Test | FDR correction BH | 0.076989129 | FALSE | QL | SN | 0.428849414 | 1.463306375 | Paired t-test |
| mbo~Q9GYU8 | -5.757318781 | 0.010412234 | 14.01417474 | 15.23806251 | 0.068300939 | 0.3601519 | -1.223887771 | t-Test | FDR correction BH | 0.035517536 | TRUE | QL | SN | 0.428127445 | 1.982456071 | Paired t-test |
| Dmel\CG10576~Q9VRP2 | -3.637199792 | 0.035812009 | 15.27254576 | 16.49831833 | 0.506072463 | 0.141802605 | -1.225772564 | t-Test | FDR correction BH | 0.078874358 | FALSE | QL | SN | 0.427568488 | 1.445971314 | Paired t-test |
| HIPP1~M9PID3 | -11.08965894 | 0.001570895 | 13.58903854 | 14.8157467 | 0.17363922 | 0.364552259 | -1.226708162 | t-Test | FDR correction BH | 0.015838197 | TRUE | QL | SN | 0.427291297 | 2.803852961 | Paired t-test |
| eEF1beta~A0A0B4LFL3 | -6.843634783 | 0.00638552 | 15.52057314 | 16.7505592 | 0.291711278 | 0.089889455 | -1.229986058 | t-Test | FDR correction BH | 0.027492254 | TRUE | QL | SN | 0.426321566 | 2.194803702 | Paired t-test |
| GlyS~Q9VFC8 | -7.143738229 | 0.005647756 | 18.9229427 | 20.15363575 | 0.295499765 | 0.192748338 | -1.23069305 | t-Test | FDR correction BH | 0.025472763 | TRUE | QL | SN | 0.426112698 | 2.248124076 | Paired t-test |
| shot~A1Z9J3 | -6.512096352 | 0.007355575 | 17.74515866 | 18.99809546 | 0.122744763 | 0.242768226 | -1.252936805 | t-Test | FDR correction BH | 0.029507637 | TRUE | QL | SN | 0.419593198 | 2.133383363 | Paired t-test |
| eIF3j~A0A0B4LF26 | -7.781187586 | 0.004416673 | 16.9579417 | 18.22207755 | 0.27978339 | 0.070639995 | -1.264135845 | t-Test | FDR correction BH | 0.022531262 | TRUE | QL | SN | 0.41634868 | 2.354904726 | Paired t-test |
| LamC~A0A0B4KFY9 | -4.843156669 | 0.01679365 | 13.61414672 | 14.90936019 | 0.565116468 | 0.129202455 | -1.295213462 | t-Test | FDR correction BH | 0.048376646 | TRUE | QL | SN | 0.407475871 | 1.774854893 | Paired t-test |
| Nup358~A0A0B4K7J2-2 | -12.77689666 | 0.001034426 | 18.0848218 | 19.38748953 | 0.10205605 | 0.252236473 | -1.302667734 | t-Test | FDR correction BH | 0.013062707 | TRUE | QL | SN | 0.405375911 | 2.98530055 | Paired t-test |
| grsm~Q9V3D8 | -19.81386932 | 0.000280928 | 18.03716358 | 19.34155679 | 0.225719425 | 0.20892212 | -1.304393213 | t-Test | FDR correction BH | 0.010038098 | TRUE | QL | SN | 0.404891367 | 3.551405292 | Paired t-test |
| pod1~Q9W3W1 | -3.315019907 | 0.04522205 | 15.67913503 | 16.98753512 | 0.462501029 | 0.301028613 | -1.308400096 | t-Test | FDR correction BH | 0.090957986 | FALSE | QL | SN | 0.403768398 | 1.344649758 | Paired t-test |
| Spat~Q9W3Z3 | -4.196635884 | 0.024684395 | 16.57524264 | 17.88486862 | 0.514357504 | 0.392100588 | -1.309625974 | t-Test | FDR correction BH | 0.062138005 | FALSE | QL | SN | 0.403425456 | 1.607577519 | Paired t-test |
| Tctp~A0A126GUT1 | -3.539486012 | 0.038377809 | 13.7539986 | 15.09188941 | 0.712699607 | 0.072926595 | -1.337890816 | t-Test | FDR correction BH | 0.082131013 | FALSE | QL | SN | 0.395598589 | 1.415919827 | Paired t-test |
| p23~A0A0B4K6D2 | -21.77438234 | 0.000212005 | 16.42712167 | 17.77000796 | 0.394196288 | 0.315829238 | -1.342886291 | t-Test | FDR correction BH | 0.008237157 | TRUE | QL | SN | 0.394231158 | 3.67365457 | Paired t-test |
| Dmel\CG9684~Q9VHR5 | -9.969718822 | 0.002147394 | 16.14753507 | 17.49152482 | 0.187301556 | 0.232421245 | -1.343989757 | t-Test | FDR correction BH | 0.016678022 | TRUE | QL | SN | 0.39392974 | 2.668088168 | Paired t-test |
| Men-b~Q9VB69 | -16.5161925 | 0.000483102 | 17.21247701 | 18.56357914 | 0.200610727 | 0.211010175 | -1.351102129 | t-Test | FDR correction BH | 0.010461282 | TRUE | QL | SN | 0.391992477 | 3.315961038 | Paired t-test |
| Ald1~C8VV14 | -6.327642274 | 0.007980176 | 17.95408235 | 19.31386459 | 0.358701794 | 0.324821507 | -1.359782231 | t-Test | FDR correction BH | 0.031081711 | TRUE | QL | SN | 0.3896411 | 2.097987552 | Paired t-test |
| scrib~E1JIZ0 | -7.235704593 | 0.005444322 | 16.92085503 | 18.28526845 | 0.179528883 | 0.209603321 | -1.364413411 | t-Test | FDR correction BH | 0.024993673 | TRUE | QL | SN | 0.388392323 | 2.264056224 | Paired t-test |
| BEST:LD19244~Q9VID9 | -3.524708899 | 0.038786019 | 13.49738068 | 14.86438736 | 0.427850075 | 0.292772048 | -1.367006682 | t-Test | FDR correction BH | 0.082823229 | FALSE | QL | SN | 0.387694808 | 1.411324792 | Paired t-test |
| lig~A0A126GUM8 | -9.26445852 | 0.002661267 | 18.77067695 | 20.13886738 | 0.32724215 | 0.130138264 | -1.368190428 | t-Test | FDR correction BH | 0.018194841 | TRUE | QL | SN | 0.387376831 | 2.574911586 | Paired t-test |
| Nup214~Q9W1X4 | -9.922182091 | 0.00217767 | 16.7001168 | 18.0914196 | 0.213330096 | 0.291040467 | -1.391302802 | t-Test | FDR correction BH | 0.016678022 | TRUE | QL | SN | 0.381220392 | 2.662007903 | Paired t-test |
| Slu7~Q9VAQ7 | -3.813162568 | 0.031719912 | 13.5073917 | 14.91029953 | 0.228407591 | 0.465977187 | -1.402907834 | t-Test | FDR correction BH | 0.073444505 | FALSE | QL | SN | 0.378166158 | 1.498668029 | Paired t-test |
| mask~A0A0B4LHK4 | -7.522659533 | 0.004868515 | 15.04363743 | 16.45705077 | 0.459072775 | 0.426474161 | -1.413413343 | t-Test | FDR correction BH | 0.024066801 | TRUE | QL | SN | 0.375422405 | 2.31260349 | Paired t-test |
| sm~A0A0B4K7I2 | -3.455909912 | 0.040761111 | 13.90844104 | 15.33066198 | 0.615590405 | 0.529966982 | -1.422220943 | t-Test | FDR correction BH | 0.084890988 | FALSE | QL | SN | 0.373137447 | 1.389753987 | Paired t-test |
| ps~Q0KI96 | -5.405543115 | 0.012412969 | 16.46845228 | 17.89220828 | 0.408994011 | 0.065501007 | -1.423756003 | t-Test | FDR correction BH | 0.04019077 | TRUE | QL | SN | 0.372740632 | 1.906124345 | Paired t-test |
| Hex-C~Q7JYW9 | -4.417559465 | 0.021532452 | 14.16265196 | 15.58643849 | 0.66544937 | 0.151130997 | -1.423786536 | t-Test | FDR correction BH | 0.056790059 | FALSE | QL | SN | 0.372732743 | 1.666906517 | Paired t-test |
| Dmel\CG12224~Q9VGF2 | -3.717833553 | 0.033857745 | 13.90030094 | 15.32513059 | 0.867279712 | 0.788366883 | -1.424829658 | t-Test | FDR correction BH | 0.076350791 | FALSE | QL | SN | 0.372463341 | 1.470341965 | Paired t-test |
| RpS19a~E2QD65 | -13.00539389 | 0.000981597 | 18.55469012 | 19.98662242 | 0.188972979 | 0.171273684 | -1.431932301 | t-Test | FDR correction BH | 0.012817077 | TRUE | QL | SN | 0.370634144 | 3.008066941 | Paired t-test |
| Cbp20~Q9V3L6 | -4.360440688 | 0.022294944 | 13.52429221 | 14.97343843 | 0.59054506 | 0.306331983 | -1.449146223 | t-Test | FDR correction BH | 0.058032428 | FALSE | QL | SN | 0.366238097 | 1.651793614 | Paired t-test |
| Dif~E1JHK1 | -5.686642276 | 0.010778316 | 14.52544686 | 15.97668224 | 0.211689256 | 0.577071125 | -1.451235388 | t-Test | FDR correction BH | 0.036454821 | TRUE | QL | SN | 0.365708132 | 1.967449103 | Paired t-test |
| Rpn8~A0A0B4KFH4 | -4.12761345 | 0.025790164 | 13.62335836 | 15.09970635 | 0.643690501 | 0.173446739 | -1.476347999 | t-Test | FDR correction BH | 0.06368161 | FALSE | QL | SN | 0.359397431 | 1.588545902 | Paired t-test |
| ISY1~A0A0B4LGT9 | -5.900563311 | 0.009718811 | 15.92190098 | 17.4074604 | 0.262782937 | 0.420580224 | -1.485559414 | t-Test | FDR correction BH | 0.034358997 | TRUE | QL | SN | 0.357110036 | 2.01238687 | Paired t-test |
| Sf3b2~Q9VQK7 | -8.340288474 | 0.003613232 | 13.42187935 | 14.92620992 | 0.187181638 | 0.259934106 | -1.504330576 | t-Test | FDR correction BH | 0.020315914 | TRUE | QL | SN | 0.352493711 | 2.442104155 | Paired t-test |
| Mf~Q95U15 | -6.99640225 | 0.005995067 | 18.2916534 | 19.79975865 | 0.278784899 | 0.312601382 | -1.508105244 | t-Test | FDR correction BH | 0.026555929 | TRUE | QL | SN | 0.351572652 | 2.222205963 | Paired t-test |
| mod~A0A0B4K7G4 | -7.261299794 | 0.005389427 | 15.81007912 | 17.33093306 | 0.24080433 | 0.122635966 | -1.520853933 | t-Test | FDR correction BH | 0.024813171 | TRUE | QL | SN | 0.34847959 | 2.26845741 | Paired t-test |
| Map205~Q8IMF5 | -30.02072139 | 8.12E-05 | 17.05204214 | 18.57494137 | 0.212425636 | 0.178117688 | -1.522899227 | t-Test | FDR correction BH | 0.006466378 | TRUE | QL | SN | 0.347985904 | 4.09052469 | Paired t-test |
| Nup62~Q7JXF5 | -11.66204218 | 0.001354469 | 13.76725306 | 15.29211973 | 0.330681894 | 0.118255699 | -1.524866669 | t-Test | FDR correction BH | 0.014978909 | TRUE | QL | SN | 0.34751167 | 2.868230972 | Paired t-test |
| by~Q9VHC3 | -5.669417252 | 0.010870082 | 15.15089348 | 16.68498224 | 0.296928407 | 0.245888286 | -1.53408876 | t-Test | FDR correction BH | 0.036454821 | TRUE | QL | SN | 0.345297367 | 1.963767191 | Paired t-test |
| vig~Q9V426 | -5.117326371 | 0.014442469 | 14.17118272 | 15.74310068 | 1.223556257 | 1.002165233 | -1.571917952 | t-Test | FDR correction BH | 0.044074432 | TRUE | QL | SN | 0.336360931 | 1.840358553 | Paired t-test |
| Papss~Q961A8 | -3.770341107 | 0.032658154 | 13.3912061 | 14.96350192 | 0.372137786 | 0.817313684 | -1.572295817 | t-Test | FDR correction BH | 0.075179826 | FALSE | QL | SN | 0.336272844 | 1.486008365 | Paired t-test |
| Zasp52~A1ZA47 | -6.816003218 | 0.00645969 | 15.59524134 | 17.1686676 | 0.142182059 | 0.535212207 | -1.57342626 | t-Test | FDR correction BH | 0.02766206 | TRUE | QL | SN | 0.336009456 | 2.189788314 | Paired t-test |
| SelD~O18373 | -5.825684347 | 0.010073469 | 16.57079385 | 18.14593598 | 0.491696227 | 0.160184833 | -1.575142125 | t-Test | FDR correction BH | 0.034898746 | TRUE | QL | SN | 0.335610062 | 1.99682094 | Paired t-test |
| brm~M9PFM5 | -3.950931528 | 0.028926612 | 13.09387734 | 14.67202537 | 0.473282385 | 0.369775524 | -1.578148031 | t-Test | FDR correction BH | 0.06865523 | FALSE | QL | SN | 0.334911534 | 1.538702433 | Paired t-test |
| qkr58E-1~Q9W255 | -6.012782974 | 0.00921733 | 13.16954052 | 14.7504842 | 0.383093562 | 0.485250778 | -1.580943673 | t-Test | FDR correction BH | 0.033370926 | TRUE | QL | SN | 0.334263174 | 2.035394848 | Paired t-test |
| Taf4~A4V201 | -6.244716077 | 0.008283727 | 13.37513692 | 14.96375134 | 0.236293219 | 0.233445219 | -1.588614425 | t-Test | FDR correction BH | 0.031765323 | TRUE | QL | SN | 0.332490627 | 2.081774247 | Paired t-test |
| Zasp66~Q9VSN2 | -8.486392436 | 0.003435639 | 16.28923808 | 17.88383828 | 0.243419353 | 0.350019286 | -1.594600191 | t-Test | FDR correction BH | 0.020121224 | TRUE | QL | SN | 0.331113975 | 2.463992472 | Paired t-test |
| ND-B12~Q9W2E8 | -4.080869617 | 0.0265755 | 13.4035098 | 15.00154015 | 0.365052243 | 0.321672539 | -1.598030351 | t-Test | FDR correction BH | 0.064732066 | FALSE | QL | SN | 0.330327652 | 1.575518561 | Paired t-test |
| Rnp4F~A8JUY0 | -5.660910215 | 0.01091578 | 13.0040707 | 14.6029625 | 0.318656209 | 0.76092908 | -1.5988918 | t-Test | FDR correction BH | 0.036531172 | TRUE | QL | SN | 0.330130469 | 1.961945212 | Paired t-test |
| hebe~Q7K568 | -6.951497778 | 0.006106494 | 17.811745 | 19.42024089 | 0.26389529 | 0.31494196 | -1.608495889 | t-Test | FDR correction BH | 0.026651081 | TRUE | QL | SN | 0.327940073 | 2.214208079 | Paired t-test |

| | | | | | | | | | | | | | | | |
|---|---|---|---|---|---|---|---|---|---|---|---|---|---|---|---|
| Dmel\CG11550~Q9V9V4 | -5.138066524 | 0.01428256 | 13.45858181 | 15.06841948 | 0.21817458 | 0.472988127 | -1.60983767 | t-Test | FDR correction BH | 0.043787262 | TRUE | QL | SN | 0.327635214 | 1.845193939 | Paired t-test |
| Nlp~A0A0B4KI24 | -4.709275849 | 0.018123722 | 14.29339591 | 15.92454045 | 0.66511414 | 0.232865766 | -1.631144536 | t-Test | FDR correction BH | 0.051189876 | FALSE | QL | SN | 0.322831993 | 1.741752597 | Paired t-test |
| nocte~M9PE74 | -7.383338245 | 0.005137489 | 15.79929505 | 17.44677798 | 0.509497916 | 0.392152509 | -1.647482931 | t-Test | FDR correction BH | 0.024608571 | TRUE | QL | SN | 0.319196573 | 2.289249116 | Paired t-test |
| Trh~Q9W0K2 | -3.305100879 | 0.045558957 | 15.72543037 | 17.37363162 | 0.802473475 | 1.057168358 | -1.64820125 | t-Test | FDR correction BH | 0.091289834 | FALSE | QL | SN | 0.319037685 | 1.341426231 | Paired t-test |
| l(3)72Ab~Q9VUV9 | -6.053133041 | 0.009045329 | 13.18421936 | 14.836735 | 0.414937104 | 0.434559287 | -1.652515634 | t-Test | FDR correction BH | 0.033357703 | TRUE | QL | SN | 0.318085027 | 2.04357565 | Paired t-test |
| fln~M9PD14 | -14.08595262 | 0.000774975 | 18.33319266 | 19.99273148 | 0.324468527 | 0.127493202 | -1.659538826 | t-Test | FDR correction BH | 0.012134178 | TRUE | QL | SN | 0.316540318 | 3.110712086 | Paired t-test |
| Dmel\CG1674~H9XVM9 | -13.09317992 | 0.000962254 | 21.60358728 | 23.26677865 | 0.256780166 | 0.064668975 | -1.663191367 | t-Test | FDR correction BH | 0.012773921 | TRUE | QL | SN | 0.315739931 | 3.0167103 | Paired t-test |
| Dmel\CG14480~Q7JWU9 | -10.27656556 | 0.001964794 | 13.34733091 | 15.02158198 | 0.249976774 | 0.179319378 | -1.674251071 | t-Test | FDR correction BH | 0.016665632 | TRUE | QL | SN | 0.313328722 | 2.706682961 | Paired t-test |
| nop5~Q9VM69 | -11.52960779 | 0.001400842 | 13.73083545 | 15.4084868 | 0.452657665 | 0.557485105 | -1.677651356 | t-Test | FDR correction BH | 0.015251006 | TRUE | QL | SN | 0.312591108 | 2.85361089 | Paired t-test |
| tyf~Q9W4M7 | -5.042406273 | 0.015039938 | 13.57001062 | 15.25372591 | 0.568934031 | 0.260977899 | -1.683715295 | t-Test | FDR correction BH | 0.045635468 | TRUE | QL | SN | 0.311279982 | 1.822753965 | Paired t-test |
| AOX1~Q9VF53 | -7.350153719 | 0.00520443 | 14.8384253 | 16.5353225 | 0.327256119 | 0.265928719 | -1.696897201 | t-Test | FDR correction BH | 0.024608571 | TRUE | QL | SN | 0.30844877 | 2.283626803 | Paired t-test |
| bic~Q7KM15 | -3.697391513 | 0.034340005 | 13.7112908 | 15.41372686 | 0.821432604 | 0.231174215 | -1.702436058 | t-Test | FDR correction BH | 0.076938999 | FALSE | QL | SN | 0.307266831 | 1.464199646 | Paired t-test |
| RpA-70~Q24492 | -11.93667863 | 0.001264605 | 14.45249303 | 16.16673483 | 0.189011882 | 0.35500603 | -1.714241803 | t-Test | FDR correction BH | 0.014389396 | TRUE | QL | SN | 0.304762689 | 2.898045165 | Paired t-test |
| Ns1~Q8MT06 | -3.642197295 | 0.035686759 | 13.53775137 | 15.26461471 | 0.666427479 | 0.179264217 | -1.726863339 | t-Test | FDR correction BH | 0.078738237 | FALSE | QL | SN | 0.302108077 | 1.44749289 | Paired t-test |
| CG13621~Q9VC53 | -4.410583402 | 0.021623748 | 13.25030715 | 14.97773948 | 0.240718158 | 0.505274772 | -1.727432334 | t-Test | FDR correction BH | 0.056910254 | FALSE | QL | SN | 0.30198895 | 1.66506902 | Paired t-test |
| CG9480~Q9W2J5 | -6.037203524 | 0.009112725 | 16.86231022 | 18.59258738 | 0.468733372 | 0.173566756 | -1.730277161 | t-Test | FDR correction BH | 0.033357703 | TRUE | QL | SN | 0.301394049 | 2.040351752 | Paired t-test |
| Cpsf5~Q9VSY2 | -10.75424131 | 0.001719395 | 13.23542607 | 14.97656406 | 0.281262146 | 0.211897597 | -1.74113799 | t-Test | FDR correction BH | 0.016207074 | TRUE | QL | SN | 0.299133628 | 2.764624461 | Paired t-test |
| Dmel\CG7966~Q9VFZ4 | -4.367116356 | 0.022204044 | 14.59140162 | 16.35855156 | 0.641822763 | 0.114924195 | -1.767149943 | t-Test | FDR correction BH | 0.057890413 | FALSE | QL | SN | 0.293788546 | 1.653567918 | Paired t-test |
| Patj~A4V1B2 | -8.076928042 | 0.003965252 | 13.49493134 | 15.26982786 | 0.651543515 | 0.595149439 | -1.774896525 | t-Test | FDR correction BH | 0.021213554 | TRUE | QL | SN | 0.29221527 | 2.401729238 | Paired t-test |
| Dmel\CG14712~Q9VGL0 | -5.692884362 | 0.010745311 | 13.47327925 | 15.25703357 | 0.277821964 | 0.751342912 | -1.783754315 | t-Test | FDR correction BH | 0.036454821 | TRUE | QL | SN | 0.290426637 | 1.96878099 | Paired t-test |
| RanGAP~M9PD65 | -20.12778888 | 0.000268063 | 13.30343193 | 15.09014099 | 0.443867106 | 0.334072473 | -1.786709064 | t-Test | FDR correction BH | 0.009930797 | TRUE | QL | SN | 0.28983243 | 3.571763228 | Paired t-test |
| bt~L0MN91 | -4.37093829 | 0.022152218 | 13.485785 | 15.27716353 | 0.612679026 | 0.481962165 | -1.791378524 | t-Test | FDR correction BH | 0.057849973 | FALSE | QL | SN | 0.288895868 | 1.654582778 | Paired t-test |
| BcDNA:GH23906~M9PDW8 | -3.934602847 | 0.029240716 | 16.07410871 | 17.90482018 | 0.804592978 | 0.180011341 | -1.830711465 | t-Test | FDR correction BH | 0.068933059 | FALSE | QL | SN | 0.28112595 | 1.534011992 | Paired t-test |
| Zasp66~Q95TZ7 | -8.922478056 | 0.002969724 | 17.4977616 | 19.35752603 | 0.233785202 | 0.272742033 | -1.85976443 | t-Test | FDR correction BH | 0.019035448 | TRUE | QL | SN | 0.275521264 | 2.527283962 | Paired t-test |
| D1~A4V2K7 | -5.634016871 | 0.011061913 | 13.38850166 | 15.2505718 | 0.279345261 | 0.52366527 | -1.862070141 | t-Test | FDR correction BH | 0.036862617 | TRUE | QL | SN | 0.275081278 | 1.956169779 | Paired t-test |
| Dmel\CG1674~L0MLJ3 | -4.558584337 | 0.019789365 | 13.73926938 | 15.60384689 | 0.577077249 | 0.415725753 | -1.86457751 | t-Test | FDR correction BH | 0.053431096 | FALSE | QL | SN | 0.274603609 | 1.703568137 | Paired t-test |
| Uba2~Q7KUA4 | -3.268776495 | 0.046820227 | 13.1585456 | 15.03623311 | 0.309260978 | 0.912317983 | -1.877687512 | t-Test | FDR correction BH | 0.092783534 | FALSE | QL | SN | 0.272119545 | 1.329566489 | Paired t-test |
| BEAF-32~Q94513 | -10.02403922 | 0.002113478 | 13.49342751 | 15.37119551 | 0.428382302 | 0.176738356 | -1.877768002 | t-Test | FDR correction BH | 0.016678022 | TRUE | QL | SN | 0.272104364 | 2.675002368 | Paired t-test |
| Zasp52~A0A126GUN6 | -18.50112951 | 0.000344609 | 19.011639 | 20.91656782 | 0.323748488 | 0.214218818 | -1.904928821 | t-Test | FDR correction BH | 0.010357789 | TRUE | QL | SN | 0.267029526 | 3.462672842 | Paired t-test |
| jbug~Q9W205 | -3.334350887 | 0.044574487 | 15.38552317 | 17.29211124 | 0.907299203 | 0.510278825 | -1.906588068 | t-Test | FDR correction BH | 0.089882478 | FALSE | QL | SN | 0.266722592 | 1.35091365 | Paired t-test |
| grh~A0A0B4LFT7 | -3.735102433 | 0.033457083 | 13.32591064 | 15.23748145 | 0.303228715 | 0.845122185 | -1.91157081 | t-Test | FDR correction BH | 0.076100982 | FALSE | QL | SN | 0.265802981 | 1.475511927 | Paired t-test |
| rin~Q9VFT4 | -3.76588158 | 0.032757899 | 14.20261002 | 16.13259553 | 0.777843738 | 0.37968185 | -1.929985517 | t-Test | FDR correction BH | 0.0752763 | FALSE | QL | SN | 0.262431805 | 1.484683956 | Paired t-test |
| pzg~Q9VP57 | -5.012728296 | 0.015285514 | 12.9722898 | 14.90840684 | 0.269748274 | 0.528981283 | -1.936117034 | t-Test | FDR correction BH | 0.045856543 | TRUE | QL | SN | 0.261318826 | 1.815719948 | Paired t-test |
| l(3)02640~Q9W0G4 | -3.925112854 | 0.029425286 | 14.76332531 | 16.71393214 | 0.633276568 | 0.259059583 | -1.950606832 | t-Test | FDR correction BH | 0.068933059 | FALSE | QL | SN | 0.25870739 | 1.531279313 | Paired t-test |
| sls~Q9I7U4-3 | -4.245440424 | 0.023939245 | 17.27577082 | 19.23777195 | 1.092989672 | 0.347326134 | -1.962001129 | t-Test | FDR correction BH | 0.060821719 | FALSE | QL | SN | 0.256672186 | 1.620889547 | Paired t-test |
| CAH1~Q9V396 | -3.929041021 | 0.029348706 | 13.76376 | 15.72995601 | 0.498002749 | 0.679957493 | -1.966196008 | t-Test | FDR correction BH | 0.068933059 | FALSE | QL | SN | 0.255926952 | 1.532411039 | Paired t-test |
| Zasp67~Q9VT49 | -11.09758431 | 0.001567595 | 17.48406414 | 19.45208696 | 0.325776585 | 0.254638084 | -1.968022813 | t-Test | FDR correction BH | 0.015838197 | TRUE | QL | SN | 0.255603091 | 2.804766008 | Paired t-test |
| Ars2~Q9V9K7 | -9.040639179 | 0.002858037 | 13.45358519 | 15.46591361 | 0.408260119 | 0.102189341 | -2.012328415 | t-Test | FDR correction BH | 0.01873602 | TRUE | QL | SN | 0.24787275 | 2.543932163 | Paired t-test |
| CG5340~Q86B83 | -4.590253442 | 0.019423328 | 13.79188309 | 15.81214205 | 0.80598865 | 0.360030373 | -2.020258955 | t-Test | FDR correction BH | 0.053163851 | FALSE | QL | SN | 0.246513924 | 1.711676363 | Paired t-test |
| Taldo~Q9W1G0 | -18.13258687 | 0.000365895 | 13.18763788 | 15.21044927 | 0.22080897 | 0.08213307 | -2.022811391 | t-Test | FDR correction BH | 0.010461282 | TRUE | QL | SN | 0.246078174 | 3.436643403 | Paired t-test |
| Dmel\CG6115~Q9VJG4 | -13.20792525 | 0.000937727 | 13.5591102 | 15.61191812 | 0.143103927 | 0.344839093 | -2.052807925 | t-Test | FDR correction BH | 0.012659318 | TRUE | QL | SN | 0.241014538 | 3.027923464 | Paired t-test |
| DmRH27~Q9VTC1 | -3.81755046 | 0.031625734 | 15.05366002 | 17.14504986 | 0.546364 | 0.623064099 | -2.091389839 | t-Test | FDR correction BH | 0.073439932 | FALSE | QL | SN | 0.234654521 | 1.499959392 | Paired t-test |
| Dmel\CG1674~Q9V4C1 | -8.748176509 | 0.003145264 | 20.19687641 | 22.30935838 | 0.363960367 | 0.06923046 | -2.112481964 | t-Test | FDR correction BH | 0.019270792 | TRUE | QL | SN | 0.23124884 | 2.502342869 | Paired t-test |
| Dmel\CG30069~A1Z9M5 | -10.71899785 | 0.001736063 | 13.84040193 | 16.01055307 | 0.618834066 | 0.415926941 | -2.17015114 | t-Test | FDR correction BH | 0.016234815 | TRUE | QL | SN | 0.222187392 | 2.760434591 | Paired t-test |
| Nup50~Q7K0D8 | -9.684192568 | 0.002338061 | 16.39323566 | 18.5812438 | 0.181139119 | 0.262632183 | -2.188008145 | t-Test | FDR correction BH | 0.017164751 | TRUE | QL | SN | 0.21945421 | 2.631144127 | Paired t-test |
| EndoGI~Q9V3V9 | -6.506737467 | 0.007372797 | 15.82745402 | 18.03686041 | 0.45121533 | 0.285683718 | -2.209406391 | t-Test | FDR correction BH | 0.029507637 | TRUE | QL | SN | 0.216223256 | 2.13236772 | Paired t-test |
| mor~A0A0B4JDA0 | -9.84831607 | 0.002225847 | 13.3234493 | 15.55746374 | 0.37411454 | 0.043436141 | -2.234014437 | t-Test | FDR correction BH | 0.016884641 | TRUE | QL | SN | 0.212566413 | 2.652504655 | Paired t-test |
| caz~Q27294 | -9.311866157 | 0.002621908 | 13.42852164 | 15.68245619 | 0.39267972 | 0.325185827 | -2.253934549 | t-Test | FDR correction BH | 0.018079919 | TRUE | QL | SN | 0.209651557 | 2.581382501 | Paired t-test |
| Drep2~Q7K304 | -5.983877631 | 0.009343192 | 13.11233568 | 15.36825769 | 0.595906668 | 0.201775651 | -2.255922009 | t-Test | FDR correction BH | 0.033749898 | TRUE | QL | SN | 0.20936294 | 2.029504716 | Paired t-test |
| Hcf~Q9V4C8-5 | -10.98246842 | 0.001616434 | 13.80333879 | 16.11384258 | 0.605139412 | 0.325218969 | -2.310503791 | t-Test | FDR correction BH | 0.016000463 | TRUE | QL | SN | 0.201590032 | 2.791441999 | Paired t-test |
| Hrb98DE~A4V3J6 | -4.320788347 | 0.022844915 | 13.57754469 | 15.89313025 | 0.635195959 | 0.618292878 | -2.315585562 | t-Test | FDR correction BH | 0.058791519 | FALSE | QL | SN | 0.200881197 | 1.641210444 | Paired t-test |
| Cp190~Q24478 | -15.64807278 | 0.000567205 | 15.59131527 | 17.93218761 | 0.249541641 | 0.130514608 | -2.34087234 | t-Test | FDR correction BH | 0.010886227 | TRUE | QL | SN | 0.197390937 | 3.246260299 | Paired t-test |
| Dmel\CG11267~Q9VU35 | -6.746560432 | 0.006651165 | 13.26639273 | 15.60749407 | 0.343940987 | 0.337189319 | -2.341101342 | t-Test | FDR correction BH | 0.027994359 | TRUE | QL | SN | 0.197359608 | 2.177102271 | Paired t-test |
| CtBP~A0A0B4KGG9 | -10.89080749 | 0.001656775 | 16.98137774 | 19.33934716 | 0.322071538 | 0.287156037 | -2.357969422 | t-Test | FDR correction BH | 0.016191675 | TRUE | QL | SN | 0.195065504 | 2.780736406 | Paired t-test |
| Dmel\CG10731~A1ZAA9 | -6.848491139 | 0.006372601 | 13.82077087 | 16.19857554 | 0.606771266 | 0.504787967 | -2.37780467 | t-Test | FDR correction BH | 0.027492254 | TRUE | QL | SN | 0.192401951 | 2.195683284 | Paired t-test |
| BcDNA:RE07178~Q5LJR0 | -8.821698424 | 0.003069589 | 13.34411311 | 15.73043514 | 0.223464085 | 0.586733319 | -2.386322028 | t-Test | FDR correction BH | 0.019235452 | TRUE | QL | SN | 0.191269398 | 2.512919752 | Paired t-test |
| BcDNA:RH19933~M9PF57 | -5.53476939 | 0.011623894 | 16.21612879 | 18.61564496 | 0.792284421 | 0.286738386 | -2.399516169 | t-Test | FDR correction BH | 0.038100543 | TRUE | QL | SN | 0.189528121 | 1.934648345 | Paired t-test |
| Sf3b1~Q9VPR5 | -28.50256284 | 9.48E-05 | 16.20088328 | 18.62697933 | 0.254019431 | 0.107502646 | -2.426096047 | t-Test | FDR correction BH | 0.006567284 | TRUE | QL | SN | 0.186068269 | 4.023102152 | Paired t-test |
| scrib~A0A0B4K6I1 | -4.697683389 | 0.018245239 | 14.34887459 | 16.83858718 | 1.151084498 | 0.52560916 | -2.489712582 | t-Test | FDR correction BH | 0.051351 | FALSE | QL | SN | 0.178041741 | 1.73885044 | Paired t-test |
| bt~D1YSG0 | -5.913355977 | 0.009659865 | 16.55884043 | 19.05020191 | 0.896970014 | 0.200547992 | -2.491361479 | t-Test | FDR correction BH | 0.034358997 | TRUE | QL | SN | 0.177838368 | 2.015028956 | Paired t-test |
| Sf3a1~Q9VEP9 | -7.794380196 | 0.004395129 | 14.38297591 | 16.94826188 | 0.595852524 | 0.380556701 | -2.565285968 | t-Test | FDR correction BH | 0.022531262 | TRUE | QL | SN | 0.168955361 | 2.357028376 | Paired t-test |
| Bacc~Q9VQF7 | -12.48736944 | 0.001106933 | 13.46559333 | 16.05141514 | 0.373802434 | 0.020956802 | -2.585821816 | t-Test | FDR correction BH | 0.013258226 | TRUE | QL | SN | 0.166567424 | 2.955878713 | Paired t-test |
| Dmel\CG1737~X2JEU3 | -8.608919376 | 0.003295503 | 13.30505799 | 15.89237007 | 0.372883526 | 0.235675807 | -2.587312083 | t-Test | FDR correction BH | 0.019661931 | TRUE | QL | SN | 0.166395453 | 2.482078354 | Paired t-test |
| Pep~M9NG39 | -46.68915555 | 2.16E-05 | 14.4625305 | 17.05399646 | 0.205504356 | 0.111526312 | -2.59146596 | t-Test | FDR correction BH | 0.003864721 | TRUE | QL | SN | 0.165917048 | 4.66489407 | Paired t-test |
| Bap111~Q9W384 | -4.237932043 | 0.024051974 | 12.9528709 | 15.5459422 | 0.446686291 | 0.650487305 | -2.593071305 | t-Test | FDR correction BH | 0.06101082 | FALSE | QL | SN | 0.165732529 | 1.618849271 | Paired t-test |
| zormin~M9PE03 | -33.81094244 | 5.69E-05 | 13.45617475 | 16.08882312 | 0.303634891 | 0.319234029 | -2.632648368 | t-Test | FDR correction BH | 0.005033571 | TRUE | QL | SN | 0.161247828 | 4.245067035 | Paired t-test |
| Dmel\CG10417~E2QC56 | -9.967636528 | 0.002148709 | 13.07830831 | 15.72020415 | 0.443746317 | 0.257034332 | -2.64189584 | t-Test | FDR correction BH | 0.016678022 | TRUE | QL | SN | 0.160217558 | 2.667822408 | Paired t-test |
| Dsp1~F2FBA6 | -6.808772827 | 0.006479285 | 14.29969575 | 16.99480079 | 0.889615493 | 0.397250412 | -2.695105042 | t-Test | FDR correction BH | 0.027671586 | TRUE | QL | SN | 0.154416086 | 2.188472898 | Paired t-test |
| Dmel\CG1674~Q8IMB9 | -10.19045262 | 0.002013891 | 14.80466331 | 17.53277942 | 0.596711401 | 0.35760311 | -2.728116104 | t-Test | FDR correction BH | 0.016678022 | TRUE | QL | SN | 0.150922928 | 2.695964051 | Paired t-test |
| how~E1JIT0 | -6.641123497 | 0.006956425 | 14.46812951 | 17.23107114 | 0.701102224 | 0.187077465 | -2.762941624 | t-Test | FDR correction BH | 0.028582883 | TRUE | QL | SN | 0.147323387 | 2.157613874 | Paired t-test |
| Nup153~A8JV18 | -18.59371368 | 0.000339522 | 14.39863199 | 17.20902426 | 0.462868519 | 0.207550446 | -2.810392265 | t-Test | FDR correction BH | 0.010357789 | TRUE | QL | SN | 0.142556698 | 3.469131525 | Paired t-test |
| Actn~M9MS06 | -6.225782692 | 0.008355151 | 12.75541335 | 15.66141114 | 0.332039774 | 0.756894404 | -2.905997794 | t-Test | FDR correction BH | 0.031917877 | TRUE | QL | SN | 0.133415871 | 2.078045721 | Paired t-test |
| mip120~E1JH59 | -17.969679 | 0.000375863 | 13.4735558 | 16.41752895 | 0.424650926 | 0.367515181 | -2.943973146 | t-Test | FDR correction BH | 0.010461282 | TRUE | QL | SN | 0.129949848 | 3.424970925 | Paired t-test |
| Mtor~A0A0B4KFE9 | -3.611204775 | 0.036472648 | 14.03233444 | 17.02876686 | 1.296815467 | 0.156576873 | -2.996432419 | t-Test | FDR correction BH | 0.079590314 | FALSE | QL | SN | 0.12530949 | 1.4380327 | Paired t-test |
| Dmel\CG1674~Q8IMB8 | -6.021219206 | 0.009181016 | 13.45656896 | 16.57901996 | 0.532283795 | 0.610230806 | -3.122451007 | t-Test | FDR correction BH | 0.033370926 | TRUE | QL | SN | 0.114828208 | 2.037109238 | Paired t-test |
| htk~R9PY26 | -10.46168721 | 0.001864498 | 13.65512435 | 16.94992786 | 0.428996709 | 0.26563664 | -3.294803518 | t-Test | FDR correction BH | 0.016415603 | TRUE | QL | SN | 0.101897918 | 2.729438169 | Paired t-test |
| cg~A8DYD3 | -23.97329989 | 0.000159065 | 13.2635518 | 16.59391272 | 0.221252617 | 0.196767083 | -3.330360922 | t-Test | FDR correction BH | 0.007209085 | TRUE | QL | SN | 0.099417186 | 3.798426427 | Paired t-test |
| Sin3A~A0A0B4LF82 | -30.03506505 | 8.11E-05 | 12.8695466 | 16.44495 | 0.354957421 | 0.242073716 | -3.575403395 | t-Test | FDR correction BH | 0.006466378 | TRUE | QL | SN | 0.083887322 | 4.091145398 | Paired t-test |
| CtBP~A0A0B4KHB2 | -42.26835368 | 2.91E-05 | 13.46266484 | 17.10539073 | 0.262524983 | 0.308041762 | -3.642725886 | t-Test | FDR correction BH | 0.003864721 | TRUE | QL | SN | 0.080062702 | 4.535449657 | Paired t-test |
| Anon1A3~Q9V9U3 | -37.0383011 | 4.33E-05 | 12.96581977 | 17.14078153 | 0.147439167 | 0.200187173 | -4.174961759 | t-Test | FDR correction BH | 0.004056448 | TRUE | QL | SN | 0.055361937 | 4.363620939 | Paired t-test |
| CG8728-RA~Q7K3W2 | -11.04575239 | 0.00158934 | 11.532161 | 17.16941349 | 0.701015831 | 0.233037992 | -5.63725249 | t-Test | FDR correction BH | 0.015923384 | TRUE | QL | SN | 0.020091757 | 2.798783287 | Paired t-test |
