## Supplementary Data 2 for "Rab2 and Arl8/BORC control retrograde axonal transport of dense core vesicles via Syd/dJIP3/4 and RUFY dynein adaptors"

| Dmel ID | UniProt ID | Name | Symbol | FlyBase ID | Fold change | *P*-value(adjusted) | Subcellular location: ER | Golgi | Autophagosomes | Early/recycling endosomes | Late endosomes/lysosomes | Presynapse | Function |
|---|---|---|---|---|---|---|---|---|---|---|---|---|---|
| Atg9 | Q7JQL5 | Autophagy-related 9 | Atg9 | FBgn0034110 | 219.4797102 | 0.016678022 | | | x | | | | Transmembrane autophagy-related protein |
| syd | M9PEX9 | sunday driver | syd | FBgn0024187 | 198.2147882 | 0.010357789 | | x | | | x | | Dynein activating adaptor |
| Trpml | Q9VW35 | Transient receptor potential cation channel, mucolipin | Trpml | FBgn0262516 | 157.0925863 | 0.008237157 |  |  |  |  | x |  | Cation channel thought to mediate calcium release from late endosomes and lysosomes |
| CG11807 | Q7K490 | dNischarin | CG11807 | FBgn0033996 | 104.3682863 | 0.010357789 | | | | x | | | Rab4 and Rab14 effector involved in early endosome maturation |
| Golgin-45 | Q9VHE3 | - | CG9356 | FBgn0037688 | 69.40720615 | 0.010461282 | | x | | | | | Golgi-located tethering protein maintaining Golgi structure and ER-Golgi protein transport |
| RUFY1 | A0A0B4LHR8 | RUFY1 | CG31064 | FBgn0051064 | 53.1851387 | 0.016678022 | | | | x | x | | Dynein (activating?) adaptor involved in cargo sorting and retrieval from endosomes to Golgi |
| Blos2 | Q9VTE0 | Biogenesis of lysosome-related organelles complex 1, subunit 2 | Blos2 | FBgn0036118 | 49.62098091 | 0.012559972 | | | | | x | | BLOC-1 and BORC complex member |
| raptor | Q9W437 | raptor | raptor | FBgn0029840 | 48.12392085 | 0.029665099 | | | | | x | | mTORC1 complex member |
| CG17598 | Q9VR62 | - | CG17598 | FBgn0031194 | 45.87947692 | 0.020315914 | | | | | | | Protein phosphatase with unclear cellular function |
| CG6707 | Q9VT68 | - | CG6707 | FBgn0036058 | 42.85076277 | 0.016678022 | | | | | x | | Catalyzes hydrolysis of PI(4,5)P2 to PI4P. Human ortholog recruits JIP4 to lysosomal membranes, promoting retrograde transport of lysosomes |
| Golgin245 | Q9W1R3 | Golgin-245 | Golgin245 | FBgn0034854 | 42.39163957 | 0.023001049 | | x | | | | | Golgi-located tethering protein |
| CG11802 | Q9VYT1 | BLOC-1 related complex subunit 5 | BORCS5 | FBgn0030346 | 39.25149647 | 0.010461282 | | | | | x | | BORC complex member |
| mon2 | X2J564 | monensin sensitivity 2 | mon2 | FBgn0031985 | 34.39381442 | 0.016678022 | | | | x | | | Involved in the SNX3-retromer pathway |
| CT25972 | Q9VMK9 | - | CG9044 | FBgn0031752 | 31.72846844 | 0.016614137 | | | | | x | | Serine/threonine-protein kinase related to Nischarin. The mammalian ortholog localizes to lysosomes |
| Start1 | Q9W145 | Start1 | Start1 | FBgn0035028 | 31.04341698 | 0.013062707 | x | | | | x | | Putative cholesterol transporter at ER-late endosome contact sites |
| dor | Q24314 | deep orange | dor | FBgn0000482 | 28.1216141 | 0.010357789 | | | | x | x | | HOPS and CORVET complex member |
| TBC1D16 | Q9VKV6 | TBC1 domain family member 16 | TBC1D16 | FBgn0032249 | 26.61826187 | 0.016000463 | | | | x | | | Rab GTPase-activating protein involved in early endosome maturation |
| car | A4V4S5 | carnation | car | FBgn0000257 | 26.37002002 | 0.013862839 | | | | x | x | | HOPS and CORVET complex member |
| RagC-D | Q7K519 | Ras-related GTP binding C/D | RagC-D | FBgn0033272 | 25.96562596 | 0.010461282 | | | | | x | | Guanine nucleotide-binding protein regulating mTORC1 signaling |
| nSyb | M9PDW0 | neuronal Synaptobrevin | nSyb | FBgn0013342 | 25.47472743 | 0.033338521 | | | | | | x | SNARE involved in synaptic vesicle release |
| CG14969 | Q95SH7 | - | Ostml | FBgn0035440 | 25.37210463 | 0.007523507 | | | | | x | | Transmembrane protein involved in lysosome biogenesis. Human ortholog stabilizes the ClC-7 lysosomal chloride transporter |
| CG6398 | Q9VX17 | - | CG6398 | FBgn0030870 | 24.78095177 | 0.012641521 | | | | | | | Related to human claudin domain-containing protein |
| Atg4b | Q9VF80 | Autophagy-related 4b | Atg4b | FBgn0038325 | 24.75321781 | 0.010461282 | | | x | | | | Autophagy-related protein |
| Syt4 | Q9U6P7 | Synaptotagmin 4 | Syt4 | FBgn0028400 | 23.53397312 | 0.036454821 | | | | | x | | Synaptotagmin family member. Human ortholog implicated in regulating dense core vesicle motility |
| CG44247 | Q9W0Z5 | - | CG44247 | FBgn0265182 | 22.46187121 | 0.017164751 | | | | | | | C. elegans ortholog implicated in regulating trafficking and secretion of DCVs |
| ldlCp | Q9VF78 | Component of oligomeric golgi complex 2 | Cog2 | FBgn0026634 | 21.8734303 | 0.028247427 | | x | | | | | Golgi-located tethering protein. Oligomeric golgi complex 2 member |
| garz | A1Z8W8 | gartenzwerg | garz | FBgn0264560 | 20.43957237 | 0.016678022 | x | x | | | | | Arf guanine-nucleotide exchange factor (GEF) involved in early secretory pathway trafficking |
| Nhe3 | D5SHT8 | Na[+]/H[+] hydrogen exchanger 3 | Nhe3 | FBgn0028703 | 20.27040419 | 0.007209085 | | | | x | x | | Endosomal Na+, K+/H+ antiporter |
| Syx13 | Q9VU45 | Syntaxin 13 | Syx13 | FBgn0036341 | 20.00209304 | 0.007209085 | | | | | x | | SNARE implicated in degradative fusion of glue granules and lysosomes (crinophagy) |
| CT30701 | Q8IQH6 | nebulosa | nebu | FBgn0036316 | 18.42389287 | 0.013092474 | | | | | | | Glucose transporter |
| CG34126 | M9PC65 | - | CG34126 | FBgn0083962 | 18.32309482 | 0.003864721 | | | | x | | | Lipid transport protein involved in early endosome-Golgi trafficking |
| Rme-8 | A0A0B4LFW6 | Receptor mediated endocytosis 8 | Rme-8 | FBgn0015477 | 18.24077497 | 0.043787262 | | | | x | | | Associated with the endosomal WASH actin-regulating and retromer complexes. Also functions as LRRK regulator linked to Arl8-mediated transport of lysosomes and DCVs |
| dp18 | Q9VW73 | Late endosomal/lysosomal adaptor, MAPK and MTOR activator 1 | Lamtor1 | FBgn0036932 | 18.0976534 | 0.012641521 | | | | | x | | Ragulator complex member |
| Golgin104 | Q9VV40 | Golgin 104 | Golgin104 | FBgn0036614 | 17.99444057 | 0.012134178 | | x | | | | | Trans-Golgi tethering protein homologous to C. elegans CCCP-1 involved in dense core vesicle biogenesis |
| ClC-b | Q7JZ25 | Chloride channel-b | ClC-b | FBgn0033755 | 17.14854556 | 0.003100532 | | | | | x | | Late endosome/lysosome chloride transporter |
| Snx3 | Q9VG51 | Sorting nexin 3 | Snx3 | FBgn0038065 | 15.78962588 | 0.012555335 | | | | x | | | Sorting nexin involved in a variant retromer pathway (denoted the SN3-retromer pathway) |
| CG32850 | Q8SX35 | Ring finger protein 11 | Rnf11 | FBgn0052850 | 14.03720764 | 0.007209085 | | | | x | | | Ubiquitin ligase regulating dopamine release and uptake |
| CG17180 | Q9W0Q3 | BLOC-1 related complex subunit 6 | BORCS6 | FBgn0035140 | 14.00899102 | 0.007209085 | | | | | x | | BORC complex member |
| CG16823 | Q9VK04 | - | CG31729 | FBgn0051729 | 13.98572348 | 0.013862839 | | x | | x | | | Lipid transporter involved in the SNX3-retromer pathway. Orthologous to human ATP9A/B. |
| CG1513 | A1Z814 | - | CG1513 | FBgn0033463 | 13.87893864 | 0.016234815 | | x | | | | | Golgi-located sterol binding protein |
| lt | Q7PL76 | light | lt | FBgn0002566 | 13.82443689 | 0.016678022 | | | | | x | | HOPS complex member |
| Sytalpha | Q9I7M6 | Synaptotagmin alpha | Sytalpha | FBgn0261089 | 13.71119245 | 0.012134178 | | | | | | | Synaptotagmin family member specifically associated with dense core vesicles |
| Vps16A | Q9VHG1 | Vacuolar protein sorting 16A | Vps16A | FBgn0285911 | 13.48874345 | 0.016191675 | | | | x | x | | Member of HOPS and CORVET |
| Syb | H1UUB1 | Synaptobrevin | Syb | FBgn0003660 | 13.28119587 | 0.011237107 | | | | | | | SNARE involved in maturation of glue granules and their protection against early degradation. Also involved in fusion of constitutive secretory vesicles with the plasma membrane |
| rg | X2JAH8 | rugose | rg | FBgn0266098 | 13.08035633 | 0.046408873 | | | | x | | | Protein kinase A binding protein. Human ortholog NBEA located in tubular vesicles extending from recycling endosomes |
| PICK1 | Q86PF5 | Protein interacting with C kinase 1 | PICK1 | FBgn0032447 | 12.88216062 | 0.010461282 | | x | | | | | BAR domain protein involved in early stages of dense core vesicle biogenesis |
| spin | Q9GQQ0-4 | spinster | spin | FBgn0086676 | 12.56132543 | 0.006717766 | | | | | x | | Putative sphingosine-1-phosphate transporter at the late-endosomal/lysosomal membrane |
| Snap25 | P36975-3 | Synaptosomal-associated protein 25kDa | Snap25 | FBgn0011288 | 12.10427998 | 0.010357789 | | | | | | x | SNARE involved in synaptic vesicle release |
| CG7970 | Q9W0B2 | - | CG7970 | FBgn0035252 | 11.89472843 | 0.04758938 | | | | | | | Peroxisomal transmembrane protein |
| NijA | M9PHZ8 | Ninjurin A | NijA | FBgn0036101 | 11.83104323 | 0.010461282 | | x | | | | | Plasma membrane protein involved in programmed cell death, cell adhesion, axon guidance and repair. Also located in Golgi. |
| Golgin84 | Q8SZ63 | Golgin-84 | Golgin84 | FBgn0039188 | 11.76528668 | 0.010461282 | | x | | | | | Golgi-located tethering protein |
| Vamp7 | Q7JYX5 | Vesicle-associated membrane protein 7 | Vamp7 | FBgn0266186 | 11.49360112 | 0.016234895 | | | | | x | | Late endosome/lysosome SNARE mediating fusion of these organelles with autophagosomes |
| BicD | M9PDL9 | Bicaudal D | BicD | FBgn0000183 | 11.45285639 | 0.012659318 | | x | | | | | Dynein activating adaptor |
| Syt7 | Q9V4C4 | Synaptotagmin 7 | Syt7 | FBgn0039900 | 11.41179292 | 0.021213554 | | | | | x | x | Synaptotagmin family member mediating exocytosis of dense core vesicles and lysosomes, and asynchronous release of synaptic vesicles |
| Vps13B | Q9VAD3 | Vacuolar protein sorting 13B | Vps13B | FBgn0039727 | 11.34396161 | 0.012134178 | | x | | x | | | Lipid transfer protein maintaining Golgi structure and functioning as a tethering factor in trafficking between early and recycling endosomes |
| IA-2 | Q59E11 | IA-2 protein tyrosine phosphatase | IA-2 | FBgn0031294 | 10.66917658 | 0.007209085 | | | | | | | Transmembrane protein primarily detected in the membrane of dense core vesicles |
| Scamp | Q9VXV6 | Scamp | Scamp | FBgn0040285 | 10.47049567 | 0.015838197 | | x | | x | | | Transmembrane protein in Golgi and recycling endosomes. Also implicated in neurotransmitter release. |
| CG3652 | Q9VQZ0 | - | CG3652 | FBgn0031600 | 10.43331942 | 0.012773921 | | x | | | | | Golgi transmembrane protein. Human ortholog YIPF6 implicated in glycan synthesis |
| Fas2 | M9PJ19 | Fasciclin 2 | Fas2 | FBgn0000635 | 10.26648531 | 0.016207074 | | | | | | | Cell adhesion molecule |
| Hsl | Q7JR83 | Hormone-sensitive lipase | Hsl | FBgn0034491 | 10.06525983 | 0.010461282 | | | | | | | Cytosolic lipase |
| Frq2 | Q9VWX8 | Frequenin 2 | Frq2 | FBgn0083228 | 9.950815521 | 0.024608571 | | | | | | x | Calcium sensor regulating neurotransmitter secretion |
| hook | Q24185 | hook | hook | FBgn0001202 | 9.69046157 | 0.018663977 | | | | x | x | | Dynein activating adaptor involved in endosomal motility and maturation and regulation of synapse size. |
| VAChT | O17444 | Vesicular acetylcholine transporter | VAChT | FBgn0270928 | 9.46358138 | 0.021858353 | | | | | | x | Vesicular acetylcholine transporter |
| CG6327 | M9PEU1 | - | CG43693 | FBgn0263776 | 9.331724905 | 0.003864721 | | | | | x | | Transmembrane amino acid transporter. Human ortholog SLC36A1 may be involved in efflux from lysosomes of neutral amino acids |
| Atg17 | Q7KTS2 | Autophagy-related 17 | Atg17 | FBgn0037363 | 9.221993865 | 0.048228926 | | | x | | | | Autophagy-related protein |
| cue | Q95RU0 | cueball | cue | FBgn0011204 | 9.188933991 | 0.01061394 | | | | | | | Involved in triglyceride homeostasis |
| Syx16 | Q9VR90 | Syntaxin 16 | Syx16 | FBgn0031106 | 9.082836023 | 0.019559199 | | x | | | x | | SNARE involved in late endosome-to-Golgi trafficking |
| Iml1 | Q9W0E3-2 | Increased minichromosome loss 1 | Iml1 | FBgn0035227 | 9.039441697 | 0.03159942 | | | | | x | | Member of the GATOR1 complex, which inhibits amino acid-sensing TORC1 signaling |
| ergic53 | Q9V3A8 | ergic53 | ergic53 | FBgn0035909 | 8.969772492 | 0.01648088 | x | x | | | | | Lectin involved in ER-to-Golgi transport |
| mv | Q9W060 | mauve | mv | FBgn0288310 | 8.926047188 | 0.031091532 | | | | | | | Adaptor protein negatively regulating the size of lysosome-related organelles, phagosomes and autophagosomes |
| CG18065 | A1ZBV5 | BLOC-1 related complex subunit 7 | BORCS7 | FBgn0034519 | 8.734820915 | 0.017319568 | | | | | x | | BORC complex member |
| VGlut | Q9VQC0 | Vesicular glutamate transporter 1 | VGlut1 | FBgn0031424 | 8.662120932 | 0.027532339 | | | | | | x | Vesicular glutamate transporter |
| Nrx-IV | Q8IQH0 | Neurexin IV | Nrx-IV | FBgn0013997 | 8.635175414 | 0.028582883 | | | | | | x | Transmembrane protein that influences active zone formation and appears to cluster around T-bars in axon terminals |
| htt | Q9V3N4 | huntingtin | htt | FBgn0027655 | 8.600305558 | 0.034898746 | | | | | | | Involved in axonal transport of several organelles |
| Syx8 | Q9VV76 | Syntaxin 8 | Syx8 | FBgn0036643 | 8.545099743 | 0.012134178 | | | | x | x | | SNARE involved in endosomal fusion |
| GCC185 | Q9VGE4 | GRIP and coiled-coil domain containing 185 kDa | GCC185 | FBgn0037979 | 8.492911644 | 0.024737406 | | x | | | | | Golgi-located tethering protein |
| p115 | Q9W3N6 | p115 | p115 | FBgn0040087 | 8.400827046 | 0.003902745 | | x | | | | | Tethering protein regulating ER-Golgi transport |
| Q9VP46 | Q9VP46 | TBC1 domain family member 8/9 | Tbc1d8-9 | FBgn0037074 | 8.376962809 | 0.003864721 | | | | | | | Predicted Rab GTPase activator |
| Pex11 | Q7JZE1 | Peroxin 11a/b | Pex11ab | FBgn0034058 | 8.316797776 | 0.012329867 | | | | | | | Peroxisomal membrane protein |
| Pex23 | Q9VWB0 | Peroxin 23 | Pex23 | FBgn0288469 | 8.301757198 | 0.048376646 | | | x | | | | Tethering factor involved in autophagy |
| CG15210 | Q9VZ67 | - | CG15210 | FBgn0040850 | 8.020768021 | 0.014923481 | | | | | | | Unknown function |
| Vps20 | Q9W236 | Vacuolar protein sorting 20 | Vps20 | FBgn0034744 | 7.6230768 | 0.029813392 | | | | | x | | ESCRT-III component |
| MESK2 | Q9I7V7 | Misexpression suppressor of KSR 2 | MESK2 | FBgn0043070 | 7.271525731 | 0.010461282 | | | | | | | Serine hydrolase |
| CG11976 | Q9VHI9 | Glucose uptake by glia | Glug | FBgn0051100 | 7.221193047 | 0.012374532 | | | | | | | Glucose transporter |
| CG1486 | Q9VRG8 | - | CG1486 | FBgn0031174 | 7.214976496 | 0.0419776 | | x | | | | | Pyridoxal dependent decarboxylase domain containing protein |
| Sema1b | Q7KK54 | Semaphorin 1b | Sema1b | FBgn0016059 | 7.182080879 | 0.011237107 | | | | | | | transmembrane protein mediating axonal and dendritic guidance |
| LpR1 | A8JRD1 | Lipophorin receptor 1 | LpR1 | FBgn0066101 | 7.168746312 | 0.010357789 | | | | | | | Lipoprotein receptor |
| ZnT35C | Q9V471 | Zinc transporter 35C | ZnT35C | FBgn0028516 | 7.136696977 | 0.018547205 | | | | | | | Zinc transporter |
| w | P10090 | white | w | FBgn0003996 | 7.125855464 | 0.010461282 | | | | | | | ABC transporter |
| Mdr50 | Q5BI62 | Multi drug resistance 50 | Mdr50 | FBgn0010241 | 7.016139563 | 0.024608571 | | | | | | | ABC transporter |
| Dysb | M9PI95 | Dysbindin | Dysb | FBgn0036819 | 6.94182153 | 0.025267898 | | | | x | | | BLOC-1 complex member also involved in homeostatic modulation of neurotransmission |
| Rab8 | O18338 | Rab8 | Rab8 | FBgn0262518 | 6.91081203 | 0.010505536 |  |  |  | x |  |  | Rab GTPase regulating recycling endosomes and synaptic secretory autophagy |
| Piezo | M9PCC7 | Piezo | Piezo | FBgn0264953 | 6.877220179 | 0.003864721 | | | | | | | Mechanosensitive channel |
| c11.1 | Q9W350 | c11.1 | c11.1 | FBgn0040236 | 6.86094622 | 0.034898746 | | | | | x | | Late endosome/lysosome membrane protein |
| CG7600 | Q9XZ12 | - | CG7600 | FBgn0064766 | 6.854381538 | 0.024556696 | | x | | | | | Protein facilitating Golgin-mediated vesicle capture |
| Snapin | Q9VQF9 | SNAP associated protein | Snapin | FBgn0031455 | 6.733323962 | 0.025407152 | | | | x | x | | BORC and BLOC-1 complex member |
| Tret1-1 | A1Z8N1 | Trehalose transporter 1 | Tret1 | FBgn0050035 | 6.693628649 | 0.010461282 | | | | | | | Trehalose transporter |
| CG3662 | Q2PDW1 | - | CG3662 | FBgn0031285 | 6.622727491 | 0.016665632 | | x | | | | | May play a role in axon guidance and repair |
| Csp | M9PG83 | Cysteine string protein | Csp | FBgn0004179 | 6.619640051 | 0.003100532 | | | | | | x | Synaptic vesicle-associated co-chaperone of Hsc70 |
| Syx7 | Q7K0X9 | Syntaxin 7 | Syx7 | FBgn0267849 | 6.532315714 | 0.008188203 | | | | x | | | SNARE mediating trafficking from early endosomes to late endosomes/lysosomes |
| Gli | Q7KT70 | Gliotactin | Gli | FBgn0001987 | 6.381688609 | 0.034635327 | | | | | | | Unknown neuronal function |
| fab1 | O96838 | fab1 kinase | fab1 | FBgn0028741 | 6.372445524 | 0.029712198 | | | | | x | | Kinase with multiple functions related to the endocytic pathway |
| Rab32 | A1Z7S3 | Rab32 | Rab32 | FBgn0002567 | 6.318165702 | 0.006717766 | | | | | x | | Rab GTPase functioning as an A-kinase anchoring protein |
| Mul1 | Q9VZJ9 | Mitochondrial E3 ubiquitin protein ligase 1 | Mul1 | FBgn0035483 | 6.25925818 | 0.031389509 | | | | | | | E3 ubiquitin-protein ligase |
| CG12531 | M9PHM0 | - | CG12531 | FBgn0031064 | 6.244892227 | 0.012134178 | | | | | x | | Lysosomal amino acid transporter |
| mio | Q9VQ89 | missing oocyte | mio | FBgn0031399 | 6.210911247 | 0.012641521 | | | | | x | | GATOR complex member |
| CG5439 | Q9VK31 | - | CG5439 | FBgn0032476 | 6.122452422 | 0.006557185 | | | | | | | Sorting nexin involved in substance transport and lipid metabolism |
| EG:22E5.5 | O77266 | - | CG4199 | FBgn0289578 | 6.061300386 | 0.035517536 | | | | | | | Unknown cellular function |
| Nckx30C | M9PCP1 | Nckx30C | Nckx30C | FBgn0028704 | 6.003905154 | 0.024066801 | | | | | | | Possible function in calcium homeostasis |
| CG4658 | Q9VL57 | - | CG4658 | FBgn0032170 | 5.995488487 | 0.03547126 | | | | | | | Unknown function |
| Vps35 | Q7KVL7 | Vacuolar protein sorting 35 | Vps35 | FBgn0034708 | 5.885552746 | 0.03880665 | | | | x | x | | Retromer complex member |
| Usp32 | M9PD06-2 | Ubiquitin specific protease 32 | Usp32 | FBgn0036913 | 5.847214111 | 0.033357703 | | x | | | x | | Deubiquitinase acting as a positive regulator of mTORC1 signaling |
| PRL-1 | O61722 | PRL-1 phosphatase | PRL-1 | FBgn0024734 | 5.826019691 | 0.024608571 | | | | x | | | Protein tyrosine phosphatase |
| nudE | A8JNQ0 | nudE | nudE | FBgn0036059 | 5.777102106 | 0.012134178 | | | | | | | Positive regulator of dynein activity |
| RyR | Q24498-5 | Ryanodine receptor | RyR | FBgn0011286 | 5.726427836 | 0.016234815 | | | | | | | Cytosolic calcium-activated calcium channel |
| jp | Q7JV09 | junctophilin | jp | FBgn0032129 | 5.666315989 | 0.034358997 | | | | | | | Member of junctional complex mediating cross-talk between cell surface and intracellular calcium release channels |
| jus | Q8MQS2 | julius seizure | jus | FBgn0039647 | 5.642344362 | 0.016234895 | | | | | | | Transmembrane protein regulating neuronal excitability |
| Nep3 | Q9W5Y0 | Neprilysin 3 | Nep3 | FBgn0031081 | 5.600870401 | 0.020315914 | | | | | | | Peptide hormone processing protein |
| CG4552 | Q9VPW9 | TBC1 domain family member 23 | TBC1D23 | FBgn0031304 | 5.59893679 | 0.016234895 | | x | | x | | | Rab GTPase-activating protein involved in endosome-to-Golgi trafficking |
| Wwox | X2J7F1 | WW domain containing oxidoreductase | Wwox | FBgn0031972 | 5.569845754 | 0.007209085 | | | | | | | Putative oxidoreductase required for normal brain development |
| Vps26 | Q9W552 | Vacuolar protein sorting 26 | Vps26 | FBgn0014411 | 5.438147591 | 0.027994359 | | | | x | x | | Retromer complex member |
| EG:8D8.3 | M9PDG9 | - | CG12773 | FBgn0024365 | 5.401068829 | 0.034358997 | | | | | | | Putative amino acid transporter |
| Syx6 | Q7JY00 | Syntaxin 6 | Syx6 | FBgn0037084 | 5.3766433 | 0.015251006 | | x | | x | | | SNARE implicated in endosome-to-Golgi transport and constitutive secretion |
| CNT2 | A1Z7N2 | Concentrative nucleoside transporter 2 | CNT2 | FBgn0025709 | 5.370305892 | 0.041416931 | | | | | | | Cyclic-di-GMP transporter |
| numb | M9PCN6 | numb | numb | FBgn0002973 | 5.340709305 | 0.024608571 | | | | x | x | | Membrane-associated inhibitor of Notch signaling |
| mGluR | L0MPZ9 | metabotropic Glutamate Receptor | mGluR | FBgn0019985 | 5.290883205 | 0.021213554 | | | | | | | G-protein coupled receptor for glutamate |
| Vps33B | Q9VBR1 | Vacuolar protein sorting 33B | Vps33B | FBgn0039335 | 5.260926132 | 0.042006046 | | | | | x | | CHEVI complex member, SM protein required for phagosome-lysosome fusion |
| Syt1 | X2J979 | Synaptotagmin 1 | Syt1 | FBgn0004242 | 5.247767548 | 0.010886227 | | | | | | x | Fast calcium sensor for neurotransmitter release |
| MSBP | Q9VXM4 | membrane steroid binding protein | MSBP | FBgn0030703 | 5.225424757 | 0.003864721 | | | | | | | Membrane protein able to bind ecdysteroid |
| CG43066 | A0A0B4K7V4 | - | CG43066 | FBgn0262476 | 5.217530027 | 0.010461282 | | | | | | | Putative neurotransmitter transporter |
| CG11935 | Q9VBX8 | - | CG33096 | FBgn0053096 | 5.198925158 | 0.016207074 | | | | | | | Depalmitoylating enzyme that may regulate MAP and organelle trafficking |
| AAF46211 | Q9W3V2 | TATA element modulatory factor | Tmf | FBgn0029912 | 5.181347395 | 0.028302368 | | x | | | | | Golgi-located cargo receptor |
| ICA69 | Q9VP78 | Islet cell autoantigen 69 kDa | ICA69 | FBgn0037050 | 5.146386171 | 0.010867262 | | x | | | | x | BAR domain protein involved in early stages of secretory vesicle biogenesis |
| ClC-c | Q9VUY1 | Chloride channel-c | ClC-c | FBgn0036566 | 5.009556907 | 0.019235452 | | | | x | | | Chloride transporter involved in endosomal lumen acidification |
| CG13077 | Q9VIT5 | - | CG13077 | FBgn0032810 | 4.938900373 | 0.019789776 | | | | | | | Putative heme binding and transmembrane ascorbate ferrireductase activity |
| mgl | A8JTM7 | Megalin | mgl | FBgn0261260 | 4.924456968 | 0.029887369 | | | | x | | | Involved in endocytosis regulation |
| Wdfy2 | Q9VKW1 | WD repeat and FYVE domain containing 2 | Wdfy2 | FBgn0032246 | 4.888928347 | 0.032762891 | | | | x | | | Endosomal protein kinase |
| Ykt6 | Q9W3M8 | YKT6 v-SNARE | Ykt6 | FBgn0260858 | 4.867136278 | 0.009437294 | | x | x | x | | | SNARE involved in Golgi-related vesicle transport and autophagosome-lysosome fusion. Also implicated in fusion of constitutive secretory vesicles with the plasma membrane. |
| CG11857 | A0A140SRF8 | Retention in endoplasmic reticulum 1 | Rer1 | FBgn0039303 | 4.864366615 | 0.01321178 | | x | | | | | Involved in the retrieval of endoplasmic reticulum membrane proteins from the early Golgi compartment |
| mmd | M9NDY9 | mind-meld | mmd | FBgn0259110 | 4.819794118 | 0.018612459 | | | | | | | ADAM metalloproteinase ortholog |
| Klc | P46824 | Kinesin light chain | Klc | FBgn0010235 | 4.794768998 | 0.010653957 | | | | | | | Kinesin motor protein complex 1 member |
| red | Q9I7J4 | red Malpighian tubules | red | FBgn0285913 | 4.740323362 | 0.020309262 | | | | | | | Putative regulator of lysosome-related organelles |
| CG13600 | Q7KS34 | - | CG31140 | FBgn0051140 | 4.638016066 | 0.003100532 | | | | | | | Diacylglycerol kinase regulating synaptic vesicle endocytosis |
| Pi4KIIalpha | Q9VND3 | Phosphatidylinositol 4-kinase II alpha | Pi4KIIalpha | FBgn0037339 | 4.626763304 | 0.010461282 | | x | x | | x | | Phosphatidylinositol 4-kinase regulating autophagy and LE-to-Golgi protein retrieval |
| Cog1 | A0A0B4KH76 | Component of oligomeric golgi complex 1 | Cog1 | FBgn0037998 | 4.620954353 | 0.017810152 | | x | | | | | Conserved oligomeric Golgi complex member required for normal Golgi function |
| CG40006 | Q8SYC3 | - | CG40006 | FBgn0058006 | 4.588636399 | 0.049366983 | | | | | | | Membrane-located scavenger receptor |
| bw | A0A0B4LHB3 | brown | bw | FBgn0000241 | 4.5706287 | 0.012555335 | | | | | | | ABC transporter |
| GM130 | Q9W289 | Golgi matrix protein 130 kD | GM130 | FBgn0034697 | 4.556213469 | 0.013062707 | | x | | | | | Membrane-located cis-Golgi stack structural component |
| Q9VA71 | Q9VA71 | - | CG2224 | FBgn0039773 | 4.532412534 | 0.016909887 | | | | x | | | Deubiquitinase regulating receptor-mediated endocytosis and sorting of receptors to lysosomes |
| Spase25 | B4F5U4 | Spase 25-subunit | Spase25 | FBgn0030306 | 4.520849541 | 0.036454821 | | | | | | | Signal peptidase complex member |
| CG7956 | E1JIS1 | spermathreecae | sp3 | FBgn0038890 | 4.455864574 | 0.037987094 | | | | x | | | Inositol 4-phosphatase associated with clathrin-coated pits, EE and RE |
| CG6834 | Q9VGJ7 | - | CG6834 | FBgn0037935 | 4.455811125 | 0.010357789 | | | | | | | Myosin binding protein |
| mew | X2JDJ5 | multiple edematous wings | mew | FBgn0004456 | 4.440510984 | 0.02544562 | | | | | | | Integrin alpha subunit |
| Vha100-1 | Q8IML5 | Vacuolar H[+] ATPase 100kD subunit 1 | Vha100-1 | FBgn0028671 | 4.434518359 | 0.041416931 | | | x | | x | | Subunit of Vacuolar H[+] ATPase which acidifies autophagosomes and lysosomes in neurons |
| CG5916 | Q9VEY4 | - | CG5916 | FBgn0038401 | 4.430262476 | 0.006567284 | | | | | | | Putative Rab GTPase-activating protein |
| CG14232 | Q9VWD5 | - | CG14232 | FBgn0031061 | 4.356499103 | 0.016207074 | | x | | | | | Golgi-located fatty-acyl-CoA binding protein |
| wdp | Q9W266 | windpipe | wdp | FBgn0034718 | 4.353056887 | 0.01648088 | | | | | | | Regulator of the JAK/STAT pathway involved in synaptic target recognition |
| CG2930 | Q9W4P6 | - | CG2930 | FBgn0028491 | 4.331725624 | 0.010461282 | | | | | | | Dipeptide transporter |
| Wdr24 | Q9XZ25 | WD repeat domain 24 | Wdr24 | FBgn0027518 | 4.311505465 | 0.020315914 | | | x | | x | | GATOR2 complex member |
| CG10132 | Q9VIU8 | - | CG10132 | FBgn0032798 | 4.29720611 | 0.016909887 | | | | | | | Putative regulator of ER calcium ATPase |
| CG10804 | B7Z122 | - | CG10804 | FBgn0029663 | 4.280092007 | 0.016678022 | | | | | | | Putative neurotransmitter transporter |
| Arfip | Q9VGQ8 | Arfaptin | Arfip | FBgn0037884 | 4.261548398 | 0.006717766 | | x | | | | | BAR domain protein implicated in secretory vesicle biogenesis |
| CG1695 | Q8IQ31 | - | CG1695 | FBgn0031116 | 4.226957164 | 0.037584122 | | | | | | | Rab GTPase activator for some Rabs, effector for others |
| CG40016 | M9PGG8 | - | CG17493 | FBgn0040010 | 4.225267669 | 0.012134178 | | | | | | | Involved in microtubule organizing center structure and function |
| CD98hc | Q9VHX9 | CD98 heavy chain | CD98hc | FBgn0037533 | 4.216707546 | 0.017164751 | | | | | | | L-leucine transporter |
| CG6420 | Q9VBC4 | WD repeat domain 20 | Wdr20 | FBgn0039451 | 4.215264923 | 0.019235452 | | | | | | | Regulates ubiquitin-dependent protein catabolic process |
| FBgn0259166 | Q9VY47 | - | CG42271 | FBgn0259166 | 4.214881147 | 0.012498956 | | | | x | | | Inositol polyphosphate-4-phosphatase |
| if | P12080-2 | inflated | if | FBgn0001250 | 4.200564183 | 0.041437915 | | | | | | | Integrin complex component |
| Ctl1 | M9PHD2 | Choline transporter-like 1 | Ctl1 | FBgn0035523 | 4.189673155 | 0.019572092 | | | | | | | Choline transporter |
| Sema1a | M9PB77 | Semaphorin 1a | Sema1a | FBgn0011259 | 4.157211484 | 0.026875835 | | | | | | | Axon guidance protein also required for adult neuron survival |
| Gos28 | Q9VE50 | Golgi SNARE, 28 kDa | Gos28 | FBgn0044871 | 4.156667997 | 0.010461282 | | x | | | | | Golgi SNARE protein |
| CG18745 | Q9I7L0 | - | CG18745 | FBgn0042102 | 4.138975349 | 0.029507637 | | | | | | | Adaptor protein regulating the trafficking of GPCRs and extracellular vesicle biogenesis |
| Pex3 | Q9VUL8 | Peroxin 3 | Pex3 | FBgn0036484 | 4.137243396 | 0.010461282 | | | | | | | Membrane protein involved in peroxisome biogenesis |
| OtopLa | M9PGH2 | Otopetrin-like a | OtopLa | FBgn0259994 | 4.120648496 | 0.014978909 | | | | | | | Proton channel mediating acid taste |
| Snx16 | Q7JR96 | Sorting nexin 16 | Snx16 | FBgn0034265 | 4.101628787 | 0.034896442 | | | | x | | | Sorting nexin involved in early-to-late endosome trafficking |
| fra | A1Z921 | frazzled | fra | FBgn0011592 | 4.09327509 | 0.024813171 | | | | | | | Netrin receptor mediating axon guidance |
| Vps2 | Q9VBI3 | Vacuolar protein sorting 2 | Vps2 | FBgn0039402 | 4.065417934 | 0.04334975 | | | | x | x | | ESCRT-III complex member |
| iPLA2-VIA | Q9VT60 | calcium-independent phospholipase A2 VIA | iPLA2-VIA | FBgn0036053 | 4.044924142 | 0.022531262 | | | | x | x | | Calcium-independent phospholipase regulating retromer and maintaining mitochondrial integrity |
| CG17660 | Q9VQ34 | - | CG17660 | FBgn0031356 | 4.017210141 | 0.014351069 | | x | | | | | Membrane protein possibly involved in endosome-to-Golgi trafficking |
| Grasp65 | Q9VW57 | Grasp65 | Grasp65 | FBgn0036919 | 3.976373418 | 0.019235452 | | | | | | | |
| Muted | A0A0B4LHG2 | Muted | Muted | FBgn0083967 | 3.972028201 | 0.003864721 | | | | | | | |
| l(2)01289 | Q9V9A9 | hulk | hlk | FBgn0010482 | 3.96004061 | 0.036439268 | | | | | | | |
| CG11377 | M9PDP6 | Cysteine rich with EGF like domains | Creld | FBgn0031217 | 3.955159419 | 0.025472763 | | | | | | | |
| plx | A0A0B4KG76 | pollux | plx | FBgn0261261 | 3.922807087 | 0.0198174 | | | | | | | |
| Blos4 | M9PET7 | Biogenesis of lysosome-related organelles complex 1, subunit 4 | Blos4 | FBgn0036105 | 3.899898744 | 0.006717766 | | | | | | | BLOC-1 complex member |
| Q9I7L1 | Q9I7L1 | - | CG18747 | FBgn0042104 | 3.870908428 | 0.019235452 | | | | | | | |
| CG15639 | Q9VK03 | headbutt | hbt | FBgn0264308 | 3.851436327 | 0.003864721 | | | | | | | |
| Rbcn-3B | Q9W543 | Rabconnectin-3B | Rbcn-3B | FBgn0023510 | 3.827359463 | 0.013258226 | | | | | | | |
| unc-104 | A0A126GUN3 | uncoordinated-104 | unc-104 | FBgn0267002 | 3.763293237 | 0.026555929 | | | | | | | Kinesin-3, fast anterograde DCV motor |
| TM4SF | O76137 | Transmembrane 4 superfamily | TM4SF | FBgn0020372 | 3.714418978 | 0.011237107 | | | | | | | |
| Rab3-GEF | A8JUX2 | Rab3 GDP-GTP exchange factor | Rab3-GEF | FBgn0030613 | 3.695489661 | 0.040238085 | | | | | | | |
| GluClalpha | E1JIQ1 | Glutamate-gated chloride channel subunit alpha | GluClalpha | FBgn0024963 | 3.663347759 | 0.012817077 | | | | | | | |
| CG4829 | A8JV25 | - | CG4829 | FBgn0030796 | 3.637859504 | 0.023308553 | | | | | | | |
| mge | Q9VZL1 | maggie | mge | FBgn0035473 | 3.624534065 | 0.033357703 | | | | | | | |
| CG9590 | Q9VF39 | - | CG9590 | FBgn0038360 | 3.610911137 | 0.016678022 | | | | | | | |
| Tsc1 | Q9VCC9 | Tsc1 | Tsc1 | FBgn0026317 | 3.553602112 | 0.007209085 | | | | | | | |
| anchor | M9PFK5 | anchor | anchor | FBgn0036741 | 3.500496867 | 0.026555929 | | | | | | | |
| Hk | Q9W2W9 | Hyperkinetic | Hk | FBgn0263220 | 3.49513907 | 0.016415603 | | | | | | | |
| gish | A0A0B4KHL4 | gilgamesh | gish | FBgn0250823 | 3.485988731 | 0.018079919 | | | | | | | |
| didum | A1Z6Z8 | dilute class unconventional myosin | didum | FBgn0261397 | 3.444467612 | 0.018079919 | | | | | | | |

| | | | | | | | | |
|---|---|---|---|---|---|---|---|---|
| GlcAT-P | A4V1U5 | Glucuronyltransferase P | GlcAT-P | FBgn0036144 | 3.419130571 | 0.015562398 | | |
| Vps11 | Q8SX81 | Vacuolar protein sorting 11 | Vps11 | FBgn0052350 | 3.410846987 | 0.013862839 | | HOPS and CORVET complex member |
| santa-maria | Q9VM10 | scavenger receptor acting in neural tissue and majority of rhodopsin i | santa-maria | FBgn0025697 | 3.39817034 | 0.034358997 | | |
| VhaAC45 | Q7JR49 | Vacuolar H[+] ATPase AC45 accessory subunit | VhaAC45 | FBgn0262515 | 3.362634891 | 0.02059782 | | |
| CG11665 | Q7JWI7 | hermes | hrm | FBgn0033028 | 3.331112521 | 0.016890579 | | |
| Alp12 | Q9VIW9 | Alkaline phosphatase 12 | Alp12 | FBgn0032779 | 3.312321907 | 0.010038098 | | |
| CLLD6 | Q9VED7 | - | CG7785 | FBgn0038564 | 3.306057203 | 0.003879823 | | |
| DmSNX21 | M9PBU4 | Sorting nexin 21 | Snx21 | FBgn0031457 | 3.306030979 | 0.016678022 | | |
| CG10681 | M9PFF6 | - | CG10681 | FBgn0036291 | 3.30417183 | 0.010461282 | | |
| path | Q9VT04 | pathetic | path | FBgn0036007 | 3.271612762 | 0.018547205 | | |
| CG11811 | Q9VTB3 | oya | oya | FBgn0036099 | 3.271387275 | 0.029045505 | | |
| CG8379 | Q9VHK5 | - | CG8379 | FBgn0037638 | 3.185407233 | 0.003864721 | | |
| Sec22 | O77434 | Secretory 22 | Sec22 | FBgn0260855 | 3.178455579 | 0.019559199 | | |
| ATP8A | A0A0B4JD16 | ATPase 8A | ATP8A | FBgn0259221 | 3.160663008 | 0.013258226 | | |
| CG5065 | E1JH83 | - | CG5065 | FBgn0034145 | 3.148801882 | 0.016758223 | | |
| galectin | Q9VPI6 | galectin | galectin | FBgn0031213 | 3.135194423 | 0.0419776 | | |
| Vps16B | Q9VAG4 | Vacuolar protein sorting 16B | Vps16B | FBgn0039702 | 3.125278865 | 0.034358997 | | CHEVI complex member |
| nrv3 | Q7JS69 | nervana 3 | nrv3 | FBgn0032946 | 3.119263911 | 0.020315914 | | |
| Amph | A0A0B4KFR7 | Amphiphysin | Amph | FBgn0027356 | 3.079470518 | 0.043752305 | | |
| tweek | M9MSI8 | tweek | tweek | FBgn0261671 | 3.068434337 | 0.017758453 | | |
| Dg | A1ZA89 | Dystroglycan | Dg | FBgn0034072 | 3.06226202 | 0.020315914 | | |
| CG3860 | Q9W1D9 | - | CG3860 | FBgn0034951 | 3.0444214 | 0.022695703 | | |
| norpA | X2JCI4 | no receptor potential A | norpA | FBgn0262738 | 3.032917911 | 0.033357703 | | |
| Rbcn-3A | Q9W425 | Rabconnectin-3A | Rbcn-3A | FBgn0023458 | 3.023439674 | 0.010680489 | | |
| Ocrl | O46094 | Oculocerebrorenal syndrome of Lowe | Ocrl | FBgn0023508 | 3.022610304 | 0.012641521 | | |
| Vps13 | A1Z713 | Vacuolar protein sorting 13 | Vps13 | FBgn0033194 | 3.01769823 | 0.017818589 | | |
| CG5110 | M9NEQ5 | Late endosomal/lysosomal adaptor, MAPK and MTOR activator 3 | Lamtor3 | FBgn0032642 | 3.005216888 | 0.013609321 | | |
| Rac2 | M9PBU2 | Rac2 | Rac2 | FBgn0014011 | 2.953909519 | 0.019626282 | | |
| Past1 | Q8T6I0 | Putative Achaete Scute Target 1 | Past1 | FBgn0016693 | 2.945305756 | 0.010461282 | | |
| Sbf | Q7KSP6 | SET domain binding factor | Sbf | FBgn0025802 | 2.945128479 | 0.03880665 | | |
| CG8020 | Q9VWJ3 | - | CG32536 | FBgn0052536 | 2.927393936 | 0.018976903 | | |
| Cisd2 | Q9VAM6 | CDGSH iron sulfur domain | Cisd | FBgn0062442 | 2.909140242 | 0.015251006 | | |
| CG15684 | Q8IN56 | Oxysterol-binding protein-related protein 8 | Orp8 | FBgn0261550 | 2.906955478 | 0.020315914 | | |
| CG32039 | Q8IQC6 | small VCP interacting protein | Svip | FBgn0052039 | 2.904837375 | 0.041050893 | | |
| nrv1 | B5RIR1 | nervana 1 | nrv1 | FBgn0015776 | 2.892168086 | 0.033357703 | | |
| Syx1A | A0A0B4JCZ4 | Syntaxin 1A | Syx1A | FBgn0013343 | 2.887664827 | 0.018616017 | | |
| Tom20 | Q95RF6 | Translocase of outer membrane 20 | Tom20 | FBgn0036928 | 2.852506679 | 0.016678022 | | |
| Mfe2 | X2JFD6 | peroxisomal Multifunctional enzyme type 2 | Mfe2 | FBgn0030731 | 2.79963504 | 0.036454821 | | |
| Cyp9b2 | Q9V4I1 | Cytochrome P450 9b2 | Cyp9b2 | FBgn0015039 | 2.763034624 | 0.02573518 | | |
| OstDelta | A0A0B4LFR4 | Oligosaccharide transferase Delta subunit | OstDelta | FBgn0034277 | 2.755689867 | 0.045856543 | | |
| Rip11 | X2JFP6 | Rab11 interacting protein | Rip11 | FBgn0027335 | 2.750890848 | 0.022329903 | | |
| mys | P11584 | myospheroid | mys | FBgn0004657 | 2.746708979 | 0.003902745 | | |
| CG6701 | A1Z9K0 | - | CG6701 | FBgn0033889 | 2.743266708 | 0.048803692 | | |
| CG8004 | Q9VW58 | - | CG8004 | FBgn0036920 | 2.735549908 | 0.024737406 | | |
| Syx5 | A4V0S1 | Syntaxin 5 | Syx5 | FBgn0011708 | 2.733463801 | 0.020315914 | | |
| CG17338 | Q8MT77 | - | CG31793 | FBgn0051793 | 2.696866569 | 0.043148094 | | |
| Hsc70-1 | P29843 | Heat shock protein 70 cognate 1 | Hsc70-1 | FBgn0001216 | 2.694914799 | 0.049047133 | | |
| Mec2 | Q9VWL0 | Mec2 | Mec2 | FBgn0030993 | 2.693758917 | 0.047107534 | | |
| ed | M9PCI6 | echinoid | ed | FBgn0000547 | 2.688005281 | 0.016207074 | | |
| nrv2 | A4V0B5 | nervana 2 | nrv2 | FBgn0015777 | 2.645167407 | 0.025173296 | | |
| RabX6 | Q9W0N7 | RabX6 | RabX6 | FBgn0035155 | 2.619318628 | 0.048228926 | | |
| kirre | Q9W4T9 | kin of irre | kirre | FBgn0028369 | 2.61023314 | 0.016678022 | | |
| Akap200 | Q9VLL3-2 | A kinase anchor protein 200 | Akap200 | FBgn0027932 | 2.591003103 | 0.039596744 | | |
| Synd | Q9VDI1 | Syndapin | Synd | FBgn0053094 | 2.564966681 | 0.033357703 | | |
| Sec8 | Q9VNH6 | Secretory 8 | Sec8 | FBgn0266672 | 2.560822244 | 0.024066801 | | |
| CG11151 | Q9VY92 | - | CG11151 | FBgn0030519 | 2.514836145 | 0.024737406 | | |
| trio | Q8IRI5 | trio | trio | FBgn0024277 | 2.478790155 | 0.045856543 | | |
| Oatp58Da | Q9W271 | Organic anion transporting polypeptide 58Da | Oatp58Da | FBgn0050277 | 2.475407649 | 0.019035448 | | |
| kcc | Q8MLQ5 | kazachoc | kcc | FBgn0261794 | 2.474227241 | 0.017112931 | | |
| CG5676 | Q9VL16 | FUN14 domain containing 1 | Fundc1 | FBgn0032200 | 2.454501387 | 0.034358997 | | |
| CG31321 | Q8INF8 | - | CG31321 | FBgn0051321 | 2.447628656 | 0.014923481 | | |
| wus | Q9VX95 | wurst | wus | FBgn0030805 | 2.430332564 | 0.030931083 | | |
| nahoda | Q9W1X5 | nahoda | nahoda | FBgn0034797 | 2.408239586 | 0.018328051 | | |
| Vmat | Q86NW1 | Vesicular monoamine transporter | Vmat | FBgn0260964 | 2.405427041 | 0.046181181 | | Vesicular monoamine transporter. Transmembrane protein located in DCVs and SVs |
| Droj2 | Q9VFV9 | DnaJ-like-2 | Droj2 | FBgn0038145 | 2.398423959 | 0.045787132 | | |
| Cerk | Q9VNA6 | Ceramide kinase | Cerk | FBgn0037315 | 2.395383349 | 0.028709057 | | |
| FBgn 30055 | Q9W3C9 | Rubicon | Rubicon | FBgn0030055 | 2.386694784 | 0.02953445 | | |
| Crag | Q9W3D3-2 | Calmodulin-binding protein related to a Rab3 GDP/GTP exchange pr | Crag | FBgn0025864 | 2.385646739 | 0.024608571 | | |
| CG33090 | X2JAF7 | - | CG33090 | FBgn0028916 | 2.375102978 | 0.021213554 | | |
| STUB1 | Q9XYW6 | STIP1 homology and U-box containing protein 1 | STUB1 | FBgn0027052 | 2.373017321 | 0.034844455 | | |
| Rab1 | O18332 | Rab1 | Rab1 | FBgn0285937 | 2.367709436 | 0.028709057 | | |
| Diap2 | Q24307 | Death-associated inhibitor of apoptosis 2 | Diap2 | FBgn0015247 | 2.35327429 | 0.033190279 | | |
| Inx2 | E1JJF3 | Innexin 2 | Inx2 | FBgn0027108 | 2.32252814 | 0.040473622 | | |
| Atg16 | Q86BR6 | Autophagy-related 16 | Atg16 | FBgn0039705 | 2.3124807 | 0.035020929 | | Autphagy-related protein |
| stnB | Q24212 | stoned B | stnB | FBgn0016975 | 2.311076402 | 0.016678022 | | |
| Evi5 | Q9VYY9-3 | Ecotropic viral integration site 5 | Evi5 | FBgn0262740 | 2.259373781 | 0.033370926 | | |
| PMCA | E6EK15 | plasma membrane calcium ATPase | PMCA | FBgn0259214 | 2.254075081 | 0.015251006 | | |
| Tomosyn | Q9VYK6 | Tomosyn | Tomosyn | FBgn0030412 | 2.249554347 | 0.021213554 | | |
| NTPase | O76268 | NTPase | NTPase | FBgn0024947 | 2.242754476 | 0.018453763 | | |
| Ehbp1 | A1ZAN2 | Eps15 homology domain containing protein-binding protein 1 | Ehbp1 | FBgn0034180 | 2.239241526 | 0.019411961 | | |
| anon-[illegible] | A1ZBH1 | - | CG15118 | FBgn0034418 | 2.210641837 | 0.018663977 | | |
| bond | Q9VCY7 | james bond | bond | FBgn0260942 | 2.204108108 | 0.037584122 | | |
| Snap24 | Q9VH76 | Synaptosomal-associated protein 24kDa | Snap24 | FBgn0266720 | 2.182300571 | 0.019235452 | | |
| CG12576 | Q8IQ25 | - | CG12576 | FBgn0031190 | 2.167758861 | 0.024737406 | | |
| pst | Q8IQ82 | pastrel | pst | FBgn0035770 | 2.167014165 | 0.034358997 | | |
| Alk | Q7KJ08 | Anaplastic lymphoma kinase | Alk | FBgn0040505 | 2.159313947 | 0.044268621 | | |
| Tspo | Q9VPR1 | Translocator protein | Tspo | FBgn0031263 | 2.138953286 | 0.012134178 | | |
| nrm | Q9VNP2 | neuromusculin | nrm | FBgn0262509 | 2.127618581 | 0.024292432 | | |
| ClC-a | Q9VGH7-6 | Chloride channel-a | ClC-a | FBgn0051116 | 2.126625776 | 0.016678022 | | |
| RpL28 | D1Z3A1 | Ribosomal protein L28 | RpL28 | FBgn0035422 | 2.12617444 | 0.01778818 | | |
| igl | Q7JPS2 | igloo | igl | FBgn0013467 | 2.122888853 | 0.026651081 | | |
| Bem46 | O76462 | Bem46 | Bem46 | FBgn0025109 | 2.10624133 | 0.020315914 | | |
| Nrg | M9NEZ5 | Neuroglian | Nrg | FBgn0264975 | 2.101680338 | 0.020315914 | | |
| CG1882 | Q5U191 | pummelig | puml | FBgn0033226 | 2.098779989 | 0.015832927 | | |
| CG11396 | Q9VPB6 | - | CG11396 | FBgn0037022 | 2.097147978 | 0.013388667 | | |
| subdued | Q9VDV4 | subdued | subdued | FBgn0038721 | 2.087860835 | 0.016234815 | | |
| CG9485 | Q9W2H8 | Glycogen debranching enzyme | Agl | FBgn0034618 | 2.084854419 | 0.025897868 | | |
| CG32544 | Q8IQX3 | - | CG32544 | FBgn0052544 | 2.068024671 | 0.033357703 | | |
| trc | M9PG43 | tricornered | trc | FBgn0003744 | 2.060609907 | 0.036454821 | | |
| CG17029 | Q9VUW2 | - | CG17029 | FBgn0036551 | 2.058017035 | 0.042535083 | | |
| Trs31 | Q7K2Q8 | TRAPP subunit 31 | Trs31 | FBgn0266723 | 2.056555262 | 0.041652668 | | |
| OtopLc | B7Z0W9-3 | Otopetrin-like c | OtopLc | FBgn0259150 | 2.037752587 | 0.012134178 | | |
| Nhe2 | M9NFD1 | Na[+]/H[+] hydrogen exchanger 2 | Nhe2 | FBgn0040297 | 2.012799271 | 0.029041313 | | |
| PIP5K59B | Q9W1Y2 | Phosphatidylinositol 4-phosphate 5-kinase 59B | PIP5K59B | FBgn0034789 | 2.000558391 | 0.048228926 | | |
