## Supplementary Data 3 for "Rab2 and Arl8/BORC control retrograde axonal transport of dense core vesicles via Syd/dJIP3/4 and RUFY dynein adaptors"

| identifier | comparison | group1 | group2 | mean(group1) | std(group1) | mean(group2) | std(group2) | Fold-change | log2FC | T-statistics | pvalue | -Log10 pvalue | test | padj1 | correction1 | s0 | padj2 | correction2 |
|---|---|---|---|---|---|---|---|---|---|---|---|---|---|---|---|---|---|---|
| btsz~A0A0B4K657 | VMAT~Control | VMAT | Control | 21.62536654 | 0.198393671 | 13.60585743 | 0.486116646 | 259.4853183 | 8.019509103 | 6.494549489 | 1.10E-11 | 10.96027659 | Unpaired t-test | 0 | permutation FDR (250 perm) | 1 | 4.03E-10 | FDR correction BH |
| Atf-2~Q9W0Z5 | VMAT~Control | VMAT | Control | 19.4517689 | 0.201402002 | 11.5469864 | 0.392005967 | 239.6495625 | 7.904782496 | 6.603306346 | 2.22E-12 | 11.65338785 | Unpaired t-test | 0 | permutation FDR (250 perm) | 1 | 1.59E-10 | FDR correction BH |
| VGlut~Q9VQC0 | VMAT~Control | VMAT | Control | 20.14714691 | 0.132547094 | 12.67552831 | 0.463553651 | 177.4930346 | 7.471618599 | 6.146365501 | 9.49E-12 | 11.02275075 | Unpaired t-test | 0 | permutation FDR (250 perm) | 1 | 3.60E-10 | FDR correction BH |
| Dmel\CG30338~Q4QQ93 | VMAT~Control | VMAT | Control | 20.39909032 | 0.197986401 | 13.23769811 | 0.501259632 | 143.1508284 | 7.161392209 | 5.770556744 | 4.36E-11 | 10.36071528 | Unpaired t-test | 0 | permutation FDR (250 perm) | 1 | 1.14E-09 | FDR correction BH |
| Dmel\CG1695~Q8IQ31 | VMAT~Control | VMAT | Control | 19.1593824 | 0.112349473 | 12.44753295 | 0.414954961 | 104.8257582 | 6.711849454 | 5.629541638 | 8.81E-12 | 11.05480995 | Unpaired t-test | 0 | permutation FDR (250 perm) | 1 | 3.41E-10 | FDR correction BH |
| ChT~Q9VE46 | VMAT~Control | VMAT | Control | 19.81394889 | 0.336708603 | 13.43443822 | 0.460373015 | 83.25763438 | 6.379510662 | 5.082972112 | 2.39E-10 | 9.621669245 | Unpaired t-test | 0 | permutation FDR (250 perm) | 1 | 4.14E-09 | FDR correction BH |
| nSyb~Q9W0C1-2 | VMAT~Control | VMAT | Control | 21.13242816 | 0.077394296 | 14.77660157 | 0.553344283 | 81.90198957 | 6.355826593 | 5.085182501 | 2.02E-10 | 9.693978467 | Unpaired t-test | 0 | permutation FDR (250 perm) | 1 | 3.66E-09 | FDR correction BH |
| VAChT~O17444 | VMAT~Control | VMAT | Control | 20.94584735 | 0.10486712 | 14.97879149 | 0.206451447 | 62.55511106 | 5.967055859 | 5.407116091 | 6.02E-14 | 13.22073046 | Unpaired t-test | 0 | permutation FDR (250 perm) | 1 | 1.42E-11 | FDR correction BH |
| AP-2mu~O62530 | VMAT~Control | VMAT | Control | 18.6881511 | 0.11092731 | 12.76229974 | 0.376467157 | 60.79376062 | 5.92585136 | 5.041056506 | 1.23E-11 | 10.91058397 | Unpaired t-test | 0 | permutation FDR (250 perm) | 1 | 4.32E-10 | FDR correction BH |
| TBC1D16~Q9VKV6 | VMAT~Control | VMAT | Control | 18.15162954 | 0.093436827 | 12.32302689 | 1.436829039 | 56.83086026 | 5.828602649 | 3.545536771 | 3.93E-06 | 5.405564772 | Unpaired t-test | 0 | permutation FDR (250 perm) | 1 | 1.12E-05 | FDR correction BH |
| Atg9~Q7JQL5 | VMAT~Control | VMAT | Control | 19.00239845 | 0.171205495 | 13.19653807 | 0.377399437 | 55.94201764 | 5.805860383 | 4.898083699 | 2.58E-11 | 10.58821893 | Unpaired t-test | 0 | permutation FDR (250 perm) | 1 | 7.79E-10 | FDR correction BH |
| Rbsn-5~Q9VLS5 | VMAT~Control | VMAT | Control | 17.16701646 | 0.21395649 | 11.86708067 | 0.481269962 | 39.39486777 | 5.299935787 | 4.289566403 | 6.76E-10 | 9.169873974 | Unpaired t-test | 0 | permutation FDR (250 perm) | 1 | 9.20E-09 | FDR correction BH |
| bchs~Q9VML2 | VMAT~Control | VMAT | Control | 18.20698946 | 0.105012824 | 13.00139381 | 1.067203132 | 36.90119563 | 5.205595657 | 3.518309985 | 7.46E-07 | 6.127290867 | Unpaired t-test | 0 | permutation FDR (250 perm) | 1 | 2.60E-06 | FDR correction BH |
| Nos~Q27571 | VMAT~Control | VMAT | Control | 19.41170481 | 0.194353861 | 14.23623337 | 1.146258493 | 36.13866825 | 5.175471437 | 3.405052441 | 1.66E-06 | 5.780379117 | Unpaired t-test | 0 | permutation FDR (250 perm) | 1 | 5.24E-06 | FDR correction BH |
| Nhe3~D5SHT8 | VMAT~Control | VMAT | Control | 18.62135076 | 0.067383884 | 13.47337328 | 0.39420427 | 35.45648169 | 5.147977481 | 4.366946672 | 5.97E-11 | 10.22424643 | Unpaired t-test | 0 | permutation FDR (250 perm) | 1 | 1.45E-09 | FDR correction BH |
| Syx16~Q9VR90 | VMAT~Control | VMAT | Control | 17.73623324 | 0.132688347 | 12.6252856 | 0.450267163 | 34.55799581 | 5.110947646 | 4.22417854 | 3.12E-10 | 9.506535505 | Unpaired t-test | 0 | permutation FDR (250 perm) | 1 | 5.16E-09 | FDR correction BH |
| VGAT~A1Z9M6 | VMAT~Control | VMAT | Control | 18.30641892 | 0.174915593 | 13.36902787 | 0.254701726 | 30.64099099 | 4.937391052 | 4.337970667 | 7.00E-12 | 11.1549562 | Unpaired t-test | 0 | permutation FDR (250 perm) | 1 | 3.13E-10 | FDR correction BH |
| CG31653~M9PC10 | VMAT~Control | VMAT | Control | 17.75876674 | 0.080190231 | 12.82753013 | 0.450895212 | 30.51055683 | 4.931236605 | 4.092955767 | 3.48E-10 | 9.458980368 | Unpaired t-test | 0 | permutation FDR (250 perm) | 1 | 5.61E-09 | FDR correction BH |
| Vps26~Q9W552 | VMAT~Control | VMAT | Control | 18.82436894 | 0.211733149 | 13.89345542 | 1.118969016 | 30.50372483 | 4.930913517 | 3.267024345 | 2.14E-06 | 5.670347025 | Unpaired t-test | 0 | permutation FDR (250 perm) | 1 | 6.56E-06 | FDR correction BH |
| Blos2~Q9VTE0 | VMAT~Control | VMAT | Control | 18.11846602 | 0.05452621 | 13.22324577 | 0.23584016 | 29.75830089 | 4.89522025 | 4.417059755 | 6.73E-13 | 12.17192841 | Unpaired t-test | 0 | permutation FDR (250 perm) | 1 | 7.51E-11 | FDR correction BH |
| IA-2~Q59E11 | VMAT~Control | VMAT | Control | 20.84610783 | 0.064539252 | 16.00371292 | 0.210417742 | 28.68838617 | 4.842394908 | 4.408474918 | 2.91E-13 | 12.53649597 | Unpaired t-test | 0 | permutation FDR (250 perm) | 1 | 5.00E-11 | FDR correction BH |
| PICK1~Q86PF5 | VMAT~Control | VMAT | Control | 18.27959587 | 0.021507741 | 13.43877551 | 1.037375144 | 28.6570929 | 4.840820359 | 3.306508118 | 1.08E-06 | 5.967719866 | Unpaired t-test | 0 | permutation FDR (250 perm) | 1 | 3.61E-06 | FDR correction BH |
| Syt1~P21521 | VMAT~Control | VMAT | Control | 23.03029487 | 0.031788507 | 18.22137118 | 0.228363446 | 28.03046336 | 4.808923688 | 4.359415816 | 4.95E-13 | 12.3054189 | Unpaired t-test | 0 | permutation FDR (250 perm) | 1 | 6.76E-11 | FDR correction BH |
| Dmel\CG17493~M9PGG8 | VMAT~Control | VMAT | Control | 18.15081023 | 0.184065068 | 13.40859544 | 0.498050177 | 26.76386905 | 4.742214785 | 3.832219664 | 2.18E-09 | 8.662125994 | Unpaired t-test | 0 | permutation FDR (250 perm) | 1 | 2.29E-08 | FDR correction BH |
| Scamp~Q9VXV6 | VMAT~Control | VMAT | Control | 18.676471 | 0.238892446 | 14.0395872 | 0.976791046 | 24.87946916 | 4.636883799 | 3.198493082 | 1.20E-06 | 5.920884656 | Unpaired t-test | 0 | permutation FDR (250 perm) | 1 | 3.95E-06 | FDR correction BH |
| Snx3~Q9VG51 | VMAT~Control | VMAT | Control | 17.25969885 | 0.103806794 | 12.68083072 | 0.649064577 | 23.89883066 | 4.578868126 | 3.538648962 | 2.43E-08 | 7.614020675 | Unpaired t-test | 0 | permutation FDR (250 perm) | 1 | 1.53E-07 | FDR correction BH |
| Syb~H1UUB1 | VMAT~Control | VMAT | Control | 18.56700384 | 0.135657684 | 14.02195237 | 0.294040198 | 23.34515828 | 4.545051465 | 3.970105015 | 2.53E-11 | 10.59614149 | Unpaired t-test | 0 | permutation FDR (250 perm) | 1 | 7.75E-10 | FDR correction BH |
| Dmel\CG6870~Q9VJC0 | VMAT~Control | VMAT | Control | 17.81686826 | 0.142379881 | 13.29848868 | 0.424969803 | 22.9175288 | 4.518379581 | 3.763951231 | 6.64E-10 | 9.177780049 | Unpaired t-test | 0 | permutation FDR (250 perm) | 1 | 9.14E-09 | FDR correction BH |
| Tmep~A1A714 | VMAT~Control | VMAT | Control | 17.14134386 | 0.047728253 | 12.66370769 | 0.110068419 | 22.27936439 | 4.477636169 | 4.249632077 | 1.49E-15 | 14.82633023 | Unpaired t-test | 0 | permutation FDR (250 perm) | 1 | 1.18E-12 | FDR correction BH |
| SMC2~Q7KK96 | VMAT~Control | VMAT | Control | 18.66394144 | 0.889783724 | 14.22073491 | 1.941064452 | 21.75396594 | 4.443206536 | 2.272822543 | 0.000904013 | 3.043825141 | Unpaired t-test | 2.56E-05 | permutation FDR (250 perm) | 1 | 0.001453584 | FDR correction BH |
| tbc~Q8IQ30 | VMAT~Control | VMAT | Control | 17.48414024 | 0.203750087 | 13.07921649 | 0.751280123 | 21.18430288 | 4.404923749 | 3.267457935 | 1.77E-07 | 6.751637159 | Unpaired t-test | 0 | permutation FDR (250 perm) | 1 | 7.85E-07 | FDR correction BH |
| Syn~Q24546 | VMAT~Control | VMAT | Control | 18.03929227 | 0.448809602 | 13.68098894 | 0.329137961 | 20.51067875 | 4.35830333 | 3.489707094 | 7.84E-09 | 8.105681821 | Unpaired t-test | 0 | permutation FDR (250 perm) | 1 | 6.21E-08 | FDR correction BH |
| Dmel\CG3662~Q2PDW1 | VMAT~Control | VMAT | Control | 18.13862375 | 0.186914921 | 13.80317826 | 0.660589639 | 20.18827147 | 4.335445487 | 3.317038201 | 6.24E-08 | 7.204957795 | Unpaired t-test | 0 | permutation FDR (250 perm) | 1 | 3.21E-07 | FDR correction BH |
| Dmel\CG1090~A0A0B4KGG6 | VMAT~Control | VMAT | Control | 17.84446015 | 0.138649332 | 13.54992729 | 0.236907514 | 19.6238045 | 4.294532861 | 3.824982054 | 8.64E-12 | 11.063726 | Unpaired t-test | 0 | permutation FDR (250 perm) | 1 | 3.41E-10 | FDR correction BH |
| Sh~A0A0S0X7Z4 | VMAT~Control | VMAT | Control | 17.82234937 | 0.057768715 | 13.54270746 | 0.215574899 | 19.42229686 | 4.279641917 | 3.891257173 | 1.14E-12 | 11.94204915 | Unpaired t-test | 0 | permutation FDR (250 perm) | 1 | 1.04E-10 | FDR correction BH |
| BcDNA:RH51268~Q9VW17 | VMAT~Control | VMAT | Control | 17.81582016 | 0.090280444 | 13.5428919 | 0.332476913 | 19.33212408 | 4.272928254 | 3.702478643 | 8.62E-11 | 10.06462332 | Unpaired t-test | 0 | permutation FDR (250 perm) | 1 | 1.90E-09 | FDR correction BH |
| Csp~M9PG83 | VMAT~Control | VMAT | Control | 18.13766955 | 0.311679597 | 13.96459814 | 0.241735956 | 18.03929951 | 4.173071413 | 3.547330988 | 4.13E-10 | 9.383788628 | Unpaired t-test | 0 | permutation FDR (250 perm) | 1 | 6.32E-09 | FDR correction BH |
| EMC7~A1ZA83 | VMAT~Control | VMAT | Control | 18.56404259 | 0.152534077 | 14.46447668 | 1.234666184 | 17.14321642 | 4.099565909 | 2.634077604 | 2.40E-05 | 4.619577023 | Unpaired t-test | 0 | permutation FDR (250 perm) | 1 | 5.48E-05 | FDR correction BH |
| Dmel\CG5850~A0A6H2EIL3 | VMAT~Control | VMAT | Control | 17.54291661 | 0.092306108 | 13.44930832 | 0.211450672 | 17.07256937 | 4.09360829 | 3.710730635 | 2.47E-12 | 11.60656284 | Unpaired t-test | 0 | permutation FDR (250 perm) | 1 | 1.59E-10 | FDR correction BH |
| Rab3-GEF~Q9VXY2 | VMAT~Control | VMAT | Control | 17.56171899 | 0.51737637 | 13.47152653 | 0.395227653 | 17.03219489 | 4.090192458 | 3.16783271 | 6.56E-08 | 7.183380684 | Unpaired t-test | 0 | permutation FDR (250 perm) | 1 | 3.33E-07 | FDR correction BH |
| w~P10090 | VMAT~Control | VMAT | Control | 20.01987782 | 0.119402235 | 15.95307316 | 0.196996389 | 16.75830887 | 4.066804665 | 3.686976404 | 2.60E-12 | 11.58499605 | Unpaired t-test | 0 | permutation FDR (250 perm) | 1 | 1.59E-10 | FDR correction BH |
| Syt4~Q9U6P7 | VMAT~Control | VMAT | Control | 20.25415737 | 0.071224978 | 16.19250021 | 0.334942201 | 16.69862206 | 4.061657154 | 3.522258534 | 1.34E-10 | 9.873145883 | Unpaired t-test | 0 | permutation FDR (250 perm) | 1 | 2.65E-09 | FDR correction BH |
| Dmel\CG13743~Q8MRD1 | VMAT~Control | VMAT | Control | 19.14462357 | 0.219868016 | 15.11148997 | 0.459328756 | 16.37171557 | 4.033133602 | 3.285009431 | 7.02E-09 | 8.153443711 | Unpaired t-test | 0 | permutation FDR (250 perm) | 1 | 5.68E-08 | FDR correction BH |
| Syx13~Q9VU45 | VMAT~Control | VMAT | Control | 18.01427705 | 0.069004415 | 14.01921953 | 0.869483731 | 15.94527991 | 3.995057518 | 2.874002378 | 1.28E-06 | 5.894080191 | Unpaired t-test | 0 | permutation FDR (250 perm) | 1 | 4.15E-06 | FDR correction BH |
| mtd~Q9VNA2 | VMAT~Control | VMAT | Control | 20.29142389 | 0.121944245 | 16.30565197 | 0.206208834 | 15.84298115 | 3.985771926 | 3.600068686 | 4.69E-12 | 11.32892997 | Unpaired t-test | 0 | permutation FDR (250 perm) | 1 | 2.37E-10 | FDR correction BH |
| Rgk3~A8DYK6 | VMAT~Control | VMAT | Control | 18.02591686 | 0.267088413 | 14.07998104 | 0.722578171 | 15.41150466 | 3.945935817 | 2.934837976 | 4.52E-07 | 6.344602783 | Unpaired t-test | 0 | permutation FDR (250 perm) | 1 | 1.71E-06 | FDR correction BH |
| Dmel\CG6707~Q9VT68 | VMAT~Control | VMAT | Control | 19.44826182 | 0.094766458 | 15.54185602 | 0.577586152 | 14.99496033 | 3.9064058 | 3.096002223 | 3.67E-08 | 7.435120788 | Unpaired t-test | 0 | permutation FDR (250 perm) | 1 | 2.12E-07 | FDR correction BH |
| Sytbeta~Q9VUJ6 | VMAT~Control | VMAT | Control | 16.83065721 | 0.212414514 | 12.93343835 | 0.260603048 | 14.89977727 | 3.89721886 | 3.387838697 | 1.68E-10 | 9.774666077 | Unpaired t-test | 0 | permutation FDR (250 perm) | 1 | 3.22E-09 | FDR correction BH |
| Dmel\CG10804~B7Z122 | VMAT~Control | VMAT | Control | 18.15275466 | 0.523259243 | 14.28099254 | 0.43831645 | 14.63917271 | 3.871762121 | 2.966274155 | 1.73E-07 | 6.761356693 | Unpaired t-test | 0 | permutation FDR (250 perm) | 1 | 7.72E-07 | FDR correction BH |
| Dmel\CG1909~Q7K4Y8 | VMAT~Control | VMAT | Control | 18.19325919 | 0.20797903 | 14.32720285 | 1.58204947 | 14.5813899 | 3.866056339 | 2.256099942 | 0.000293975 | 3.531689593 | Unpaired t-test | 2.77E-05 | permutation FDR (250 perm) | 1 | 0.000525849 | FDR correction BH |
| Dmel\CG6282~M9PET5 | VMAT~Control | VMAT | Control | 18.30626257 | 0.106914796 | 14.47879879 | 1.11744227 | 14.19650388 | 3.827463782 | 2.548215121 | 1.79E-05 | 4.747103428 | Unpaired t-test | 2.19E-06 | permutation FDR (250 perm) | 1 | 4.18E-05 | FDR correction BH |
| CG17202~Q9VG76 | VMAT~Control | VMAT | Control | 17.31272755 | 0.348751526 | 13.52227218 | 0.405777957 | 13.8369625 | 3.790455371 | 3.058585745 | 2.07E-08 | 7.684485843 | Unpaired t-test | 0 | permutation FDR (250 perm) | 1 | 1.33E-07 | FDR correction BH |
| nebu~Q9VU17 | VMAT~Control | VMAT | Control | 17.12868542 | 0.446938346 | 13.37877234 | 0.414006255 | 13.45353211 | 3.749913085 | 2.946993761 | 7.97E-08 | 7.098613775 | Unpaired t-test | 0 | permutation FDR (250 perm) | 1 | 3.92E-07 | FDR correction BH |
| ine~Q9VR07-2 | VMAT~Control | VMAT | Control | 17.48557625 | 0.378569156 | 13.73934126 | 0.399182383 | 13.41927648 | 3.746234983 | 3.006529514 | 3.03E-08 | 7.518991661 | Unpaired t-test | 0 | permutation FDR (250 perm) | 1 | 1.82E-07 | FDR correction BH |
| Dmel\CG7600~Q9XZ12 | VMAT~Control | VMAT | Control | 16.84683736 | 0.113604105 | 13.10406425 | 0.087290377 | 13.38711431 | 3.742773106 | 3.517408989 | 5.25E-14 | 13.28004016 | Unpaired t-test | 0 | permutation FDR (250 perm) | 1 | 1.37E-11 | FDR correction BH |
| Nckx30C~Q9U6A0 | VMAT~Control | VMAT | Control | 18.08449084 | 0.136966207 | 14.34314148 | 0.778525646 | 13.37390952 | 3.741349357 | 2.764174166 | 9.43E-07 | 6.025423543 | Unpaired t-test | 0 | permutation FDR (250 perm) | 1 | 3.20E-06 | FDR correction BH |
| Dmel\CG12531~M9PHM0 | VMAT~Control | VMAT | Control | 20.83267533 | 0.133764031 | 17.09193476 | 0.16532762 | 13.36826718 | 3.740740568 | 3.415869839 | 2.70E-12 | 11.56920846 | Unpaired t-test | 0 | permutation FDR (250 perm) | 1 | 1.59E-10 | FDR correction BH |
| rb~Q9W4K1 | VMAT~Control | VMAT | Control | 18.96133333 | 0.118741821 | 15.23164369 | 0.21872701 | 13.26625853 | 3.729689642 | 3.356142873 | 1.32E-11 | 10.87823016 | Unpaired t-test | 0 | permutation FDR (250 perm) | 1 | 4.58E-10 | FDR correction BH |
| st~P45843 | VMAT~Control | VMAT | Control | 17.14117802 | 0.153246473 | 13.45885802 | 0.306641229 | 12.83774583 | 3.682319998 | 3.192839257 | 3.56E-10 | 9.448738691 | Unpaired t-test | 0 | permutation FDR (250 perm) | 1 | 5.66E-09 | FDR correction BH |
| Dmel\CG13077~Q9VIT5 | VMAT~Control | VMAT | Control | 17.89923321 | 0.217376139 | 14.22945051 | 0.738461838 | 12.72666675 | 3.669782706 | 2.729962881 | 8.82E-07 | 6.054396601 | Unpaired t-test | 0 | permutation FDR (250 perm) | 1 | 3.01E-06 | FDR correction BH |
| Dap160~M9PDC7 | VMAT~Control | VMAT | Control | 17.1460032 | 0.059360818 | 13.51208164 | 0.292768545 | 12.41421867 | 3.633921559 | 3.205663346 | 1.04E-10 | 9.981501326 | Unpaired t-test | 0 | permutation FDR (250 perm) | 1 | 2.17E-09 | FDR correction BH |
| Vps35~Q7KVL7 | VMAT~Control | VMAT | Control | 17.79730066 | 0.138718402 | 14.17272196 | 0.540344618 | 12.33408426 | 3.624578702 | 2.900856928 | 4.75E-08 | 7.323187581 | Unpaired t-test | 0 | permutation FDR (250 perm) | 1 | 2.61E-07 | FDR correction BH |
| Syt1~X2J4C1 | VMAT~Control | VMAT | Control | 17.24638517 | 0.079410928 | 13.6895375 | 0.233599296 | 11.76841131 | 3.55684767 | 3.203385665 | 1.94E-11 | 10.71128237 | Unpaired t-test | 0 | permutation FDR (250 perm) | 1 | 6.36E-10 | FDR correction BH |
| Golgin104~Q9VV40 | VMAT~Control | VMAT | Control | 17.12346928 | 0.032520637 | 13.56744058 | 0.348014798 | 11.7617327 | 3.556028703 | 3.07531141 | 6.07E-10 | 9.216645413 | Unpaired t-test | 0 | permutation FDR (250 perm) | 1 | 8.56E-09 | FDR correction BH |
| Dmel\CG7785~Q9VED7 | VMAT~Control | VMAT | Control | 20.14598296 | 0.127424812 | 16.62190266 | 0.285101203 | 11.50413261 | 3.524080306 | 3.09223008 | 2.19E-10 | 9.65976439 | Unpaired t-test | 0 | permutation FDR (250 perm) | 1 | 3.83E-09 | FDR correction BH |
| RhoGAP68F~F0JAN9 | VMAT~Control | VMAT | Control | 17.11760055 | 0.074457229 | 13.61522289 | 0.504939026 | 11.33236963 | 3.502377659 | 2.851501273 | 2.81E-08 | 7.551099445 | Unpaired t-test | 0 | permutation FDR (250 perm) | 1 | 1.70E-07 | FDR correction BH |
| Dmel\CG17560~Q9VES6 | VMAT~Control | VMAT | Control | 16.68635406 | 0.295929281 | 13.23076938 | 0.247338196 | 10.97070763 | 3.45558468 | 2.947238826 | 2.11E-09 | 8.675666643 | Unpaired t-test | 0 | permutation FDR (250 perm) | 1 | 2.24E-08 | FDR correction BH |
| Stt3B~Q9XZ53 | VMAT~Control | VMAT | Control | 18.04423627 | 0.145237479 | 14.59113592 | 0.131453291 | 10.9518323 | 3.453100356 | 3.174955772 | 2.64E-12 | 11.57843346 | Unpaired t-test | 0 | permutation FDR (250 perm) | 1 | 1.59E-10 | FDR correction BH |
| Hrs~Q960X8 | VMAT~Control | VMAT | Control | 20.81164204 | 0.149328365 | 17.36303202 | 0.519495594 | 10.91779813 | 3.448610022 | 2.77725529 | 5.66E-08 | 7.24748092 | Unpaired t-test | 0 | permutation FDR (250 perm) | 1 | 3.01E-07 | FDR correction BH |
| Vps8~Q9VRX2 | VMAT~Control | VMAT | Control | 16.12103426 | 0.278875047 | 12.71377955 | 0.382558796 | 10.60927901 | 3.407254711 | 2.811920559 | 1.78E-08 | 7.750793409 | Unpaired t-test | 0 | permutation FDR (250 perm) | 1 | 1.19E-07 | FDR correction BH |
| Dmel\CG17660~Q9VQ34 | VMAT~Control | VMAT | Control | 19.39318834 | 0.092151937 | 15.98718594 | 0.61038881 | 10.60007373 | 3.406002394 | 2.669139558 | 2.25E-07 | 6.647896912 | Unpaired t-test | 0 | permutation FDR (250 perm) | 1 | 9.54E-07 | FDR correction BH |
| Sfxn2~Q9VVW3 | VMAT~Control | VMAT | Control | 17.20317786 | 0.189874102 | 13.79875692 | 0.89135549 | 10.58846047 | 3.404420936 | 2.418651138 | 8.05E-06 | 5.094134541 | Unpaired t-test | 1.52E-05 | permutation FDR (250 perm) | 1 | 2.08E-05 | FDR correction BH |
| PP2C~Q9VR62 | VMAT~Control | VMAT | Control | 18.91935799 | 0.122735142 | 15.53612437 | 0.236173458 | 10.43409538 | 3.383233621 | 3.023360227 | 6.76E-11 | 10.17011032 | Unpaired t-test | 0 | permutation FDR (250 perm) | 1 | 1.58E-09 | FDR correction BH |
| Ykt6~Q9W3M8 | VMAT~Control | VMAT | Control | 20.03446147 | 0.151265916 | 16.66651031 | 0.546995193 | 10.32415036 | 3.367951153 | 2.686183984 | 1.13E-07 | 6.947537904 | Unpaired t-test | 0 | permutation FDR (250 perm) | 1 | 5.32E-07 | FDR correction BH |
| bw~A0A0B4LHB3 | VMAT~Control | VMAT | Control | 16.94174064 | 0.117535928 | 13.62201019 | 0.367001523 | 9.984778673 | 3.319730449 | 2.831713819 | 3.10E-09 | 8.509018459 | Unpaired t-test | 0 | permutation FDR (250 perm) | 1 | 3.01E-08 | FDR correction BH |
| Syt7~Q9V4C4 | VMAT~Control | VMAT | Control | 19.45767771 | 0.09976299 | 16.14789555 | 0.183938036 | 9.916164223 | 3.309782164 | 3.026557635 | 7.75E-12 | 11.11092408 | Unpaired t-test | 0 | permutation FDR (250 perm) | 1 | 3.24E-10 | FDR correction BH |
| Dmel\CG1951~Q9VAR0 | VMAT~Control | VMAT | Control | 17.40370602 | 0.068720636 | 14.10747716 | 0.440019965 | 9.82344366 | 3.296228858 | 2.748762461 | 1.35E-08 | 7.868263287 | Unpaired t-test | 0 | permutation FDR (250 perm) | 1 | 9.46E-08 | FDR correction BH |
| Pex23~Q9VWB0 | VMAT~Control | VMAT | Control | 18.45938969 | 0.093844308 | 15.17861455 | 0.48102581 | 9.71877941 | 3.280775136 | 2.690975457 | 3.57E-08 | 7.447652181 | Unpaired t-test | 0 | permutation FDR (250 perm) | 1 | 2.06E-07 | FDR correction BH |
| Syx1A~A0A0B4JCZ4 | VMAT~Control | VMAT | Control | 18.95539162 | 0.166620573 | 15.67999079 | 0.216124438 | 9.682642519 | 3.275400832 | 2.919140803 | 1.19E-10 | 9.923870691 | Unpaired t-test | 0 | permutation FDR (250 perm) | 1 | 2.42E-09 | FDR correction BH |
| CG7469~X2JAF7 | VMAT~Control | VMAT | Control | 18.57353963 | 0.296138772 | 15.30529005 | 0.83295533 | 9.634765705 | 3.268249584 | 2.342241612 | 8.83E-06 | 5.054283632 | Unpaired t-test | 1.93E-05 | permutation FDR (250 perm) | 1 | 2.26E-05 | FDR correction BH |
| Eps-15~Q8MMD2 | VMAT~Control | VMAT | Control | 22.06835208 | 0.086130692 | 18.82634287 | 0.149261364 | 9.461108446 | 3.242009217 | 3.010031821 | 1.38E-12 | 11.85951304 | Unpaired t-test | 0 | permutation FDR (250 perm) | 1 | 1.20E-10 | FDR correction BH |
| Dscam1~A0A0B4KEF4 | VMAT~Control | VMAT | Control | 18.10423422 | 0.198314689 | 14.88533142 | 1.866163849 | 9.310784965 | 3.218902802 | 1.750095131 | 0.003289997 | 2.482804547 | Unpaired t-test | 0.000549975 | permutation FDR (250 perm) | 1 | 0.004852539 | FDR correction BH |
| Dmel\CG6184~Q8IN69 | VMAT~Control | VMAT | Control | 17.01964677 | 0.204951451 | 13.81191574 | 0.884543478 | 9.23896368 | 3.207731036 | 2.28136196 | 1.31E-05 | 4.881099848 | Unpaired t-test | 2.35E-05 | permutation FDR (250 perm) | 1 | 3.20E-05 | FDR correction BH |
| mon2~X2J564 | VMAT~Control | VMAT | Control | 16.57708974 | 0.181963993 | 13.37618787 | 0.431690111 | 9.195333264 | 3.200901863 | 2.646450315 | 2.93E-08 | 7.53314417 | Unpaired t-test | 0 | permutation FDR (250 perm) | 1 | 1.76E-07 | FDR correction BH |
| CG10220~A1Z9W2 | VMAT~Control | VMAT | Control | 16.83149201 | 0.091495859 | 13.63236843 | 0.186483622 | 9.184005968 | 3.19912358 | 2.927200563 | 1.01E-11 | 10.99598898 | Unpaired t-test | 0 | permutation FDR (250 perm) | 1 | 3.77E-10 | FDR correction BH |
| Dap160~M9PBF6 | VMAT~Control | VMAT | Control | 21.50045149 | 0.094159909 | 18.30843711 | 0.143719518 | 9.138861072 | 3.192014381 | 2.964243957 | 1.57E-12 | 11.80521566 | Unpaired t-test | 0 | permutation FDR (250 perm) | 1 | 1.23E-10 | FDR correction BH |
| Vamp7~Q7JYX5 | VMAT~Control | VMAT | Control | 17.07197939 | 0.196351795 | 13.88358266 | 0.427971164 | 9.115973484 | 3.188396728 | 2.633782695 | 3.20E-08 | 7.495519878 | Unpaired t-test | 0 | permutation FDR (250 perm) | 1 | 1.89E-07 | FDR correction BH |
| Dmel\CG12773~M9PDG9 | VMAT~Control | VMAT | Control | 16.9583017 | 0.15052649 | 13.79105033 | 0.421108567 | 8.9833365 | 3.167251376 | 2.63938643 | 2.07E-08 | 7.683370156 | Unpaired t-test | 0 | permutation FDR (250 perm) | 1 | 1.33E-07 | FDR correction BH |
| Dmel\CG43066~A0A0B4K7V4 | VMAT~Control | VMAT | Control | 19.21544532 | 0.233290738 | 16.05208919 | 0.288705178 | 8.959114361 | 3.163356124 | 2.713005052 | 3.44E-09 | 8.463453379 | Unpaired t-test | 0 | permutation FDR (250 perm) | 1 | 3.24E-08 | FDR correction BH |
| Wwox~X2J7F1 | VMAT~Control | VMAT | Control | 17.53425821 | 0.189760834 | 14.37849956 | 1.595694429 | 8.912058162 | 3.155758648 | 1.836190423 | 0.001353386 | 2.868578416 | Unpaired t-test | 0.000320735 | permutation FDR (250 perm) | 1 | 0.002114048 | FDR correction BH |
| Cnx99A~Q0KHZ9 | VMAT~Control | VMAT | Control | 17.99937428 | 0.111179525 | 14.89524712 | 0.192183756 | 8.598751244 | 3.104127159 | 2.823748724 | 2.63E-11 | 10.57937344 | Unpaired t-test | 0 | permutation FDR (250 perm) | 1 | 7.84E-10 | FDR correction BH |
| CenB1A~Q7KS50 | VMAT~Control | VMAT | Control | 16.83651508 | 0.211367699 | 13.74951065 | 0.253086293 | 8.49729958 | 3.087004429 | 2.690283037 | 1.37E-09 | 8.862313064 | Unpaired t-test | 0 | permutation FDR (250 perm) | 1 | 1.57E-08 | FDR correction BH |
| GH10711~Q9Y110 | VMAT~Control | VMAT | Control | 16.6202165 | 0.38258295 | 13.53705356 | 0.242724572 | 8.474703747 | 3.083162936 | 2.563693863 | 3.05E-08 | 7.51611788 | Unpaired t-test | 2.19E-06 | permutation FDR (250 perm) | 1 | 1.82E-07 | FDR correction BH |
| Dmel\CG31729~Q9VK04 | VMAT~Control | VMAT | Control | 18.41183537 | 0.036454645 | 15.3634624 | 0.237524096 | 8.272784369 | 3.048372979 | 2.752561281 | 6.90E-11 | 10.16138554 | Unpaired t-test | 0 | permutation FDR (250 perm) | 1 | 1.59E-09 | FDR correction BH |
| Nep3~Q9W5Y0 | VMAT~Control | VMAT | Control | 16.68944205 | 0.456107304 | 13.65310568 | 0.255656189 | 8.204050501 | 3.036336373 | 2.460893436 | 1.39E-07 | 6.858030156 | Unpaired t-test | 1.30E-05 | permutation FDR (250 perm) | 1 | 6.39E-07 | FDR correction BH |
| Syx6~Q7JY00 | VMAT~Control | VMAT | Control | 16.65641488 | 0.13880746 | 13.63034668 | 0.361714241 | 8.145866658 | 3.026068199 | 2.57918446 | 8.04E-09 | 8.094788365 | Unpaired t-test | 2.19E-06 | permutation FDR (250 perm) | 1 | 6.33E-08 | FDR correction BH |
| Syx7~Q7K0X9 | VMAT~Control | VMAT | Control | 16.42512154 | 0.317038619 | 13.40094447 | 0.487484189 | 8.135195768 | 3.024177064 | 2.400027483 | 3.92E-07 | 6.406315653 | Unpaired t-test | 1.73E-05 | permutation FDR (250 perm) | 1 | 1.51E-06 | FDR correction BH |
| jef~A1Z9U2 | VMAT~Control | VMAT | Control | 16.71121125 | 0.109727973 | 13.68965636 | 0.379220533 | 8.120423034 | 3.021554887 | 2.568149629 | 9.79E-09 | 8.009378589 | Unpaired t-test | 2.19E-06 | permutation FDR (250 perm) | 1 | 7.38E-08 | FDR correction BH |
| Dmel\CG12024~Q8IRG7 | VMAT~Control | VMAT | Control | 17.69191494 | 0.144389144 | 14.67651658 | 0.525813368 | 8.085843994 | 3.015398369 | 2.424234948 | 2.20E-07 | 6.657165312 | Unpaired t-test | 1.30E-05 | permutation FDR (250 perm) | 1 | 9.42E-07 | FDR correction BH |
| htt~Q9V3N4 | VMAT~Control | VMAT | Control | 17.0093042 | 0.222307214 | 13.99476695 | 0.426068222 | 8.081019102 | 3.014537244 | 2.481262471 | 6.65E-08 | 7.17704136 | Unpaired t-test | 1.31E-05 | permutation FDR (250 perm) | 1 | 3.37E-07 | FDR correction BH |
| Hip14~Q9VUW9 | VMAT~Control | VMAT | Control | 16.55339931 | 0.215276611 | 13.5409922 | 0.15591002 | 8.069096279 | 3.012407104 | 2.692362428 | 2.10E-10 | 9.678024254 | Unpaired t-test | 0 | permutation FDR (250 perm) | 1 | 3.74E-09 | FDR correction BH |
| Lamtor1~Q9VW73 | VMAT~Control | VMAT | Control | 16.38915551 | 0.161408793 | 13.37849535 | 0.264542917 | 8.059331395 | 3.010660157 | 2.644200734 | 9.55E-10 | 9.019946457 | Unpaired t-test | 0 | permutation FDR (250 perm) | 1 | 1.18E-08 | FDR correction BH |
| Snapin~Q9VQF9 | VMAT~Control | VMAT | Control | 15.87172186 | 0.268333174 | 12.86647194 | 0.579538239 | 8.029164814 | 3.005249928 | 2.337605005 | 9.95E-07 | 6.002174621 | Unpaired t-test | 1.93E-05 | permutation FDR (250 perm) | 1 | 3.37E-06 | FDR correction BH |
| kairos~Q9VWW2 | VMAT~Control | VMAT | Control | 16.97555594 | 0.129101216 | 13.97646699 | 0.78472664 | 7.994949666 | 2.999088951 | 2.212275497 | 7.40E-06 | 5.130594479 | Unpaired t-test | 2.75E-05 | permutation FDR (250 perm) | 1 | 1.94E-05 | FDR correction BH |
| Rab3-GEF~M9NGF2 | VMAT~Control | VMAT | Control | 20.04121796 | 0.132125928 | 17.06567422 | 0.161912868 | 7.865528702 | 2.975543743 | 2.721220833 | 2.20E-11 | 10.6578878 | Unpaired t-test | 0 | permutation FDR (250 perm) | 1 | 6.98E-10 | FDR correction BH |
| Snap25~P36975-3 | VMAT~Control | VMAT | Control | 17.54739009 | 0.166028339 | 14.57259228 | 0.491152059 | 7.861462959 | 2.974797812 | 2.414882646 | 1.55E-07 | 6.808644285 | Unpaired t-test | 1.51E-05 | permutation FDR (250 perm) | 1 | 7.01E-07 | FDR correction BH |
| Vha100-1~Q8IML5 | VMAT~Control | VMAT | Control | 16.69273108 | 0.247472354 | 13.74602603 | 0.524827839 | 7.709862053 | 2.946705047 | 2.33959353 | 4.90E-07 | 6.309724152 | Unpaired t-test | 1.93E-05 | permutation FDR (250 perm) | 1 | 1.82E-06 | FDR correction BH |
| HPS4~A1ZAX6 | VMAT~Control | VMAT | Control | 16.02160391 | 0.115913022 | 13.10228395 | 0.282397937 | 7.564894469 | 2.919319957 | 2.568654951 | 1.11E-09 | 8.952914926 | Unpaired t-test | 2.19E-06 | permutation FDR (250 perm) | 1 | 1.31E-08 | FDR correction BH |
| Dmel\CG14971~Q9VZP2 | VMAT~Control | VMAT | Control | 16.31862334 | 0.177693316 | 13.41416943 | 0.377939913 | 7.487343339 | 2.904453912 | 2.447362255 | 2.47E-08 | 7.607166159 | Unpaired t-test | 1.30E-05 | permutation FDR (250 perm) | 1 | 1.53E-07 | FDR correction BH |
| ChAT~P07668-2 | VMAT~Control | VMAT | Control | 16.6210856 | 0.119167048 | 13.73200593 | 0.10968097 | 7.407977265 | 2.889079671 | 2.693955798 | 2.35E-12 | 11.62976735 | Unpaired t-test | 0 | permutation FDR (250 perm) | 1 | 1.59E-10 | FDR correction BH |
| Rab32~A1Z7S3 | VMAT~Control | VMAT | Control | 20.41471615 | 0.07845714 | 17.52633958 | 0.070221705 | 7.404367861 | 2.888376572 | 2.758483843 | 3.22E-14 | 13.49157702 | Unpaired t-test | 0 | permutation FDR (250 perm) | 1 | 9.49E-12 | FDR correction BH |
| Dmel\CG3652~Q9VQZ0 | VMAT~Control | VMAT | Control | 16.27374219 | 0.187797808 | 13.38580727 | 0.464949332 | 7.402101534 | 2.887934925 | 2.358937349 | 1.50E-07 | 6.823985546 | Unpaired t-test | 1.72E-05 | permutation FDR (250 perm) | 1 | 6.80E-07 | FDR correction BH |
| cbs~A0A0B4K864 | VMAT~Control | VMAT | Control | 17.94949394 | 0.080906953 | 15.08656823 | 0.419548714 | 7.274891396 | 2.862925711 | 2.403628727 | 3.54E-08 | 7.451516741 | Unpaired t-test | 1.73E-05 | permutation FDR (250 perm) | 1 | 2.05E-07 | FDR correction BH |
| Vmat~Q86NW1 | VMAT~Control | VMAT | Control | 20.03628242 | 0.098787331 | 17.17775068 | 0.390636633 | 7.252768198 | 2.85853174 | 2.422078884 | 2.04E-08 | 7.690426141 | Unpaired t-test | 1.30E-05 | permutation FDR (250 perm) | 1 | 1.32E-07 | FDR correction BH |
| Ost48~Q24319 | VMAT~Control | VMAT | Control | 17.09113565 | 0.066401789 | 14.27248776 | 0.055895764 | 7.055008863 | 2.818647895 | 2.71332654 | 5.99E-15 | 14.22293536 | Unpaired t-test | 0 | permutation FDR (250 perm) | 1 | 2.35E-12 | FDR correction BH |
| TBC1D23~Q9VPW9 | VMAT~Control | VMAT | Control | 16.35816691 | 0.191708053 | 13.55749263 | 0.256479739 | 6.967660253 | 2.800674279 | 2.449851096 | 2.67E-09 | 8.573418816 | Unpaired t-test | 1.30E-05 | permutation FDR (250 perm) | 1 | 2.67E-08 | FDR correction BH |
| retm~M9PB50 | VMAT~Control | VMAT | Control | 17.12408814 | 0.120004574 | 14.32761862 | 0.266309986 | 6.947382433 | 2.796469516 | 2.473370945 | 1.10E-09 | 8.95751565 | Unpaired t-test | 1.31E-05 | permutation FDR (250 perm) | 1 | 1.30E-08 | FDR correction BH |
| Tomosyn~Q9VYK6 | VMAT~Control | VMAT | Control | 20.37948961 | 0.071394862 | 17.59668727 | 0.143646152 | 6.881878136 | 2.782802345 | 2.596533188 | 3.09E-12 | 11.5094371 | Unpaired t-test | 0 | permutation FDR (250 perm) | 1 | 1.77E-10 | FDR correction BH |
| Golgin245~Q9W1R3 | VMAT~Control | VMAT | Control | 18.21544079 | 0.171676335 | 15.43328328 | 0.2656417 | 6.878802871 | 2.782157512 | 2.437392233 | 2.53E-09 | 8.59754114 | Unpaired t-test | 1.30E-05 | permutation FDR (250 perm) | 1 | 2.60E-08 | FDR correction BH |
| CG9818~Q0KID3 | VMAT~Control | VMAT | Control | 15.83113103 | 0.609538478 | 13.05847776 | 0.115631188 | 6.833635348 | 2.772653266 | 2.170450008 | 1.60E-06 | 5.795996 | Unpaired t-test | 3.17E-05 | permutation FDR (250 perm) | 1 | 5.07E-06 | FDR correction BH |
| Rnf11~Q8SX35 | VMAT~Control | VMAT | Control | 16.40021762 | 0.301350386 | 13.62939891 | 0.505305689 | 6.824951143 | 2.770818718 | 2.193640381 | 9.87E-07 | 6.005476674 | Unpaired t-test | 2.96E-05 | permutation FDR (250 perm) | 1 | 3.34E-06 | FDR correction BH |
| Sytalpha~Q9I7M6 | VMAT~Control | VMAT | Control | 20.36428181 | 0.098831217 | 17.59538901 | 0.242824975 | 6.81584629 | 2.7688928 | 2.478322648 | 4.20E-10 | 9.376406298 | Unpaired t-test | 1.31E-05 | permutation FDR (250 perm) | 1 | 6.34E-09 | FDR correction BH |
| numb~M9PCN6 | VMAT~Control | VMAT | Control | 19.07871331 | 0.123897512 | 16.31252506 | 0.174601093 | 6.803080945 | 2.766188255 | 2.524480282 | 5.75E-11 | 10.24023414 | Unpaired t-test | 1.31E-05 | permutation FDR (250 perm) | 1 | 1.41E-09 | FDR correction BH |
| ND-B17~Q9V3W2 | VMAT~Control | VMAT | Control | 16.49908776 | 0.476775666 | 13.7403005 | 0.204583569 | 6.768270646 | 2.758787259 | 2.239236616 | 3.18E-07 | 6.496896362 | Unpaired t-test | 2.76E-05 | permutation FDR (250 perm) | 1 | 1.26E-06 | FDR correction BH |
| dCat-4~Q9VPG2 | VMAT~Control | VMAT | Control | 16.63528956 | 0.214969203 | 13.88974709 | 0.282400676 | 6.706418301 | 2.745542471 | 2.369459336 | 8.83E-09 | 8.05428452 | Unpaired t-test | 1.72E-05 | permutation FDR (250 perm) | 1 | 6.81E-08 | FDR correction BH |
| cv-d~Q9VF24 | VMAT~Control | VMAT | Control | 16.66435412 | 0.23620474 | 13.93501779 | 0.784305686 | 6.631504991 | 2.729336321 | 1.997591489 | 2.18E-05 | 4.660955111 | Unpaired t-test | 0.000113813 | permutation FDR (250 perm) | 1 | 5.01E-05 | FDR correction BH |
| syd~E1JI64 | VMAT~Control | VMAT | Control | 17.90404463 | 0.146651101 | 15.1761638 | 0.424466337 | 6.624818074 | 2.727880837 | 2.271648883 | 9.04E-08 | 7.043814658 | Unpaired t-test | 2.56E-05 | permutation FDR (250 perm) | 1 | 4.38E-07 | FDR correction BH |
| car~A4V4S5 | VMAT~Control | VMAT | Control | 16.7446396 | 0.243395893 | 14.03358341 | 0.224478412 | 6.548008478 | 2.71105619 | 2.361391417 | 5.08E-09 | 8.293969579 | Unpaired t-test | 1.72E-05 | permutation FDR (250 perm) | 1 | 4.45E-08 | FDR correction BH |
| mv~Q9W060 | VMAT~Control | VMAT | Control | 16.43352455 | 0.067837079 | 13.72503201 | 0.228688992 | 6.536383103 | 2.708492544 | 2.447408899 | 2.06E-10 | 9.686088834 | Unpaired t-test | 1.30E-05 | permutation FDR (250 perm) | 1 | 3.70E-09 | FDR correction BH |
| CG4953~Q95TN1 | VMAT~Control | VMAT | Control | 16.99651991 | 0.080512268 | 14.29137737 | 0.538176325 | 6.521222928 | 2.705142539 | 2.175674408 | 5.98E-07 | 6.223320482 | Unpaired t-test | 3.17E-05 | permutation FDR (250 perm) | 1 | 2.16E-06 | FDR correction BH |
| fid~Q9VBJ3 | VMAT~Control | VMAT | Control | 16.38595305 | 0.174290182 | 13.6867058 | 0.083570331 | 6.494629612 | 2.699247251 | 2.484483603 | 2.66E-11 | 10.57439106 | Unpaired t-test | 1.31E-05 | permutation FDR (250 perm) | 1 | 7.84E-10 | FDR correction BH |
| Dmel\CG6420~Q9VBC4 | VMAT~Control | VMAT | Control | 15.97769578 | 0.177092992 | 13.30914011 | 0.310625511 | 6.357923553 | 2.66855567 | 2.300664849 | 1.25E-08 | 7.903241064 | Unpaired t-test | 1.92E-05 | permutation FDR (250 perm) | 1 | 8.81E-08 | FDR correction BH |
| Arfip~Q9VGQ8 | VMAT~Control | VMAT | Control | 16.27756618 | 0.407465764 | 13.62245863 | 0.417699184 | 6.298933374 | 2.655107551 | 2.105623707 | 1.36E-06 | 5.867638228 | Unpaired t-test | 5.03E-05 | permutation FDR (250 perm) | 1 | 4.39E-06 | FDR correction BH |
| Sap47~A0A4D6K1Y5 | VMAT~Control | VMAT | Control | 22.22879533 | 0.093993644 | 19.58099457 | 0.488044351 | 6.267111929 | 2.64780076 | 2.1662959 | 3.13E-07 | 6.504525985 | Unpaired t-test | 3.17E-05 | permutation FDR (250 perm) | 1 | 1.24E-06 | FDR correction BH |
| Dmel\CG5554~Q9W1I7 | VMAT~Control | VMAT | Control | 16.42512602 | 0.104355868 | 13.78458128 | 0.187646455 | 6.235670707 | 2.640544744 | 2.409207297 | 9.37E-11 | 10.02826876 | Unpaired t-test | 1.51E-05 | permutation FDR (250 perm) | 1 | 2.02E-09 | FDR correction BH |
| Dmel\CG5021~E1JI85 | VMAT~Control | VMAT | Control | 16.73143256 | 0.949628222 | 14.10085328 | 0.538089639 | 6.192746027 | 2.63057928 | 1.767712914 | 0.000306095 | 3.514143493 | Unpaired t-test | 0.000497715 | permutation FDR (250 perm) | 1 | 0.00054381 | FDR correction BH |
| unc-13~X2J9E1 | VMAT~Control | VMAT | Control | 19.19964558 | 0.12123658 | 16.57550776 | 0.123548084 | 6.165157813 | 2.624137825 | 2.435595551 | 1.18E-11 | 10.92797816 | Unpaired t-test | 1.30E-05 | permutation FDR (250 perm) | 1 | 4.21E-10 | FDR correction BH |
| spir~E1JHM3 | VMAT~Control | VMAT | Control | 16.27957068 | 0.122911115 | 13.69706104 | 0.14428334 | 5.989807514 | 2.582509642 | 2.380710673 | 3.40E-11 | 10.4685264 | Unpaired t-test | 1.73E-05 | permutation FDR (250 perm) | 1 | 9.31E-10 | FDR correction BH |
| Kr-h2~X2J4W8 | VMAT~Control | VMAT | Control | 16.35151585 | 0.082680538 | 13.77695014 | 0.717403502 | 5.956916355 | 2.574565701 | 1.946070103 | 1.21E-05 | 4.915647678 | Unpaired t-test | 0.000166838 | permutation FDR (250 perm) | 1 | 2.98E-05 | FDR correction BH |
| gammaSnap1~Q9U6R9 | VMAT~Control | VMAT | Control | 16.14245264 | 0.130794782 | 13.60797491 | 0.416646252 | 5.793670862 | 2.534477727 | 2.120377797 | 1.39E-07 | 6.855716897 | Unpaired t-test | 5.04E-05 | permutation FDR (250 perm) | 1 | 6.39E-07 | FDR correction BH |
| Rbcn-3B~Q9W543 | VMAT~Control | VMAT | Control | 19.86463274 | 0.04121996 | 17.33105378 | 0.117266845 | 5.790062617 | 2.53357895 | 2.400156997 | 6.22E-13 | 12.20592702 | Unpaired t-test | 1.73E-05 | permutation FDR (250 perm) | 1 | 7.36E-11 | FDR correction BH |

| | | | | | | | | | | | | | | | | | |
|---|---|---|---|---|---|---|---|---|---|---|---|---|---|---|---|---|---|
| CG14261~A0A0B4LHR8 | VMAT~Control | VMAT | Control | 18.74668303 | 0.080914174 | 16.21991697 | 0.140195699 | 5.76278448 | 2.526766066 | 2.356199553 | 8.83E-12 | 11.05400487 | Unpaired t-test | 1.72E-05 | permutation FDR (250 perm) | 1 | 3.41E-10 | FDR correction BH |
| Syn~E2QCY9 | VMAT~Control | VMAT | Control | 23.41174657 | 0.100967069 | 20.91543438 | 0.129491944 | 5.642412701 | 2.496312192 | 2.325539327 | 1.15E-11 | 10.94008837 | Unpaired t-test | 1.93E-05 | permutation FDR (250 perm) | 1 | 4.16E-10 | FDR correction BH |
| Dmel\CG8500~Q9VH66 | VMAT~Control | VMAT | Control | 16.96454631 | 0.091420943 | 14.4695686 | 0.444249552 | 5.63719592 | 2.494977709 | 2.074243203 | 2.31E-07 | 6.63552699 | Unpaired t-test | 6.68E-05 | permutation FDR (250 perm) | 1 | 9.71E-07 | FDR correction BH |
| CG32590~Q8IR45 | VMAT~Control | VMAT | Control | 18.32992177 | 0.047493817 | 15.86823379 | 0.283167964 | 5.508608678 | 2.46168798 | 2.181563415 | 3.24E-09 | 8.488839306 | Unpaired t-test | 3.17E-05 | permutation FDR (250 perm) | 1 | 3.12E-08 | FDR correction BH |
| DS01368.1~Q7KTB0 | VMAT~Control | VMAT | Control | 18.14500457 | 0.092041766 | 15.69038832 | 0.385720039 | 5.481672899 | 2.454616243 | 2.084878827 | 7.55E-08 | 7.121921525 | Unpaired t-test | 6.06E-05 | permutation FDR (250 perm) | 1 | 3.73E-07 | FDR correction BH |
| 6959~Q9VGH2 | VMAT~Control | VMAT | Control | 16.16580028 | 0.084123338 | 13.71526726 | 0.467758059 | 5.466180185 | 2.450533017 | 2.020985151 | 4.25E-07 | 6.371459911 | Unpaired t-test | 9.96E-05 | permutation FDR (250 perm) | 1 | 1.62E-06 | FDR correction BH |
| loaf~Q5BIG2 | VMAT~Control | VMAT | Control | 16.51720088 | 0.117564791 | 14.06938623 | 0.236313918 | 5.455890323 | 2.447814643 | 2.189382626 | 1.51E-09 | 8.822201399 | Unpaired t-test | 3.17E-05 | permutation FDR (250 perm) | 1 | 1.70E-08 | FDR correction BH |
| Vps20~Q9W236 | VMAT~Control | VMAT | Control | 16.16904942 | 0.232021441 | 13.72691386 | 0.693615621 | 5.434455761 | 2.442135562 | 1.840219666 | 2.14E-05 | 4.668691651 | Unpaired t-test | 0.000312564 | permutation FDR (250 perm) | 1 | 4.93E-05 | FDR correction BH |
| CG12441~Q0IGY6 | VMAT~Control | VMAT | Control | 17.05744003 | 0.498076832 | 14.67436292 | 0.441618001 | 5.216481748 | 2.38307711 | 1.836394478 | 1.17E-05 | 4.9314835 | Unpaired t-test | 0.000320735 | permutation FDR (250 perm) | 1 | 2.88E-05 | FDR correction BH |
| Crag~Q9W3D3-2 | VMAT~Control | VMAT | Control | 17.7318384 | 0.07407069 | 15.35456074 | 0.314175357 | 5.195554265 | 2.377277665 | 2.077394275 | 1.42E-08 | 7.84745461 | Unpaired t-test | 6.69E-05 | permutation FDR (250 perm) | 1 | 9.87E-08 | FDR correction BH |
| CT11439~Q9VT05 | VMAT~Control | VMAT | Control | 15.83627031 | 0.118075026 | 13.46263962 | 0.673036503 | 5.182437063 | 2.373630691 | 1.818055052 | 1.52E-05 | 4.817223796 | Unpaired t-test | 0.000367347 | permutation FDR (250 perm) | 1 | 3.64E-05 | FDR correction BH |
| ZnT35C~Q9V471 | VMAT~Control | VMAT | Control | 15.67888836 | 0.641984118 | 13.31236703 | 0.421655163 | 5.156961671 | 2.366521323 | 1.761469237 | 4.25E-05 | 4.371815378 | Unpaired t-test | 0.000511675 | permutation FDR (250 perm) | 1 | 9.12E-05 | FDR correction BH |
| Dmel\CG1513~A1Z814 | VMAT~Control | VMAT | Control | 17.03959687 | 0.156280651 | 14.69543461 | 0.77829953 | 5.07765462 | 2.344162267 | 1.729991502 | 6.06E-05 | 4.217870801 | Unpaired t-test | 0.000618807 | permutation FDR (250 perm) | 1 | 0.000125035 | FDR correction BH |
| Pdzd8~Q9VYR9 | VMAT~Control | VMAT | Control | 16.36573019 | 0.135387854 | 14.03576761 | 0.522741951 | 5.027923095 | 2.329962583 | 1.876745759 | 2.20E-06 | 5.656614567 | Unpaired t-test | 0.000260113 | permutation FDR (250 perm) | 1 | 6.73E-06 | FDR correction BH |
| Ctl1~M9PHD2 | VMAT~Control | VMAT | Control | 15.84930486 | 0.144506814 | 13.52355839 | 0.208265938 | 5.013251012 | 2.325746471 | 2.0889364 | 1.67E-09 | 8.776519465 | Unpaired t-test | 5.65E-05 | permutation FDR (250 perm) | 1 | 1.85E-08 | FDR correction BH |
| Rab26~M9PG39 | VMAT~Control | VMAT | Control | 15.95707127 | 0.080187136 | 13.63358757 | 0.433221219 | 5.00539422 | 2.323483699 | 1.941035144 | 3.44E-07 | 6.46318608 | Unpaired t-test | 0.000168811 | permutation FDR (250 perm) | 1 | 1.35E-06 | FDR correction BH |
| Rbcn-3A~Q9W425 | VMAT~Control | VMAT | Control | 20.10460352 | 0.082563627 | 17.78684961 | 0.136298987 | 4.985554313 | 2.317753918 | 2.16356541 | 1.78E-11 | 10.74948446 | Unpaired t-test | 3.17E-05 | permutation FDR (250 perm) | 1 | 5.90E-10 | FDR correction BH |
| CCG31673~X2J9A5 | VMAT~Control | VMAT | Control | 15.85598559 | 0.423031666 | 13.53922234 | 0.334256178 | 4.982132023 | 2.316763253 | 1.866678478 | 2.29E-06 | 5.640300394 | Unpaired t-test | 0.000270077 | permutation FDR (250 perm) | 1 | 6.94E-06 | FDR correction BH |
| Dmel\CG5149~Q9VM22 | VMAT~Control | VMAT | Control | 15.80730587 | 0.152224017 | 13.49080399 | 0.586426208 | 4.98122948 | 2.316501876 | 1.822654703 | 6.55E-06 | 5.183774903 | Unpaired t-test | 0.00035102 | permutation FDR (250 perm) | 1 | 1.75E-05 | FDR correction BH |
| Rab3~A0A4P7VAW6 | VMAT~Control | VMAT | Control | 18.69615712 | 0.119478242 | 16.38460566 | 0.136574378 | 4.964166343 | 2.311551459 | 2.138046366 | 6.62E-11 | 10.17936872 | Unpaired t-test | 3.58E-05 | permutation FDR (250 perm) | 1 | 1.57E-09 | FDR correction BH |
| dor~Q24314 | VMAT~Control | VMAT | Control | 17.13634105 | 0.193338983 | 14.83996408 | 0.73551995 | 4.912226115 | 2.296376971 | 1.713575183 | 5.03E-05 | 4.298397371 | Unpaired t-test | 0.000680464 | permutation FDR (250 perm) | 1 | 0.000106366 | FDR correction BH |
| BORCS7~A1ZBV5 | VMAT~Control | VMAT | Control | 15.72609907 | 0.107881908 | 13.43341362 | 0.260583013 | 4.899672948 | 2.292685453 | 2.035900361 | 5.45E-09 | 8.263570809 | Unpaired t-test | 8.74E-05 | permutation FDR (250 perm) | 1 | 4.65E-08 | FDR correction BH |
| mRpL37~Q9VGW9 | VMAT~Control | VMAT | Control | 15.90119596 | 0.07513652 | 13.61195473 | 0.285835614 | 4.887989681 | 2.28924124 | 2.021990272 | 8.72E-09 | 8.059697004 | Unpaired t-test | 9.34E-05 | permutation FDR (250 perm) | 1 | 6.75E-08 | FDR correction BH |
| Rab11~O18335 | VMAT~Control | VMAT | Control | 18.03547542 | 0.074748381 | 15.75220942 | 0.767133111 | 4.867786875 | 2.283266004 | 1.697977906 | 5.90E-05 | 4.229280533 | Unpaired t-test | 0.000726172 | permutation FDR (250 perm) | 1 | 0.000122509 | FDR correction BH |
| ClC-c~Q9VUY1 | VMAT~Control | VMAT | Control | 18.06028269 | 0.142968214 | 15.77999067 | 0.233737006 | 4.857762715 | 2.280292021 | 2.031379035 | 4.34E-09 | 8.362911142 | Unpaired t-test | 8.94E-05 | permutation FDR (250 perm) | 1 | 3.88E-08 | FDR correction BH |
| Dmel\CG7806~Q9VLP5 | VMAT~Control | VMAT | Control | 16.0642911 | 0.262750778 | 13.79860917 | 0.368994204 | 4.808816684 | 2.265681931 | 1.884016546 | 5.65E-07 | 6.248072242 | Unpaired t-test | 0.000254098 | permutation FDR (250 perm) | 1 | 2.06E-06 | FDR correction BH |
| didum~A1Z6Z8 | VMAT~Control | VMAT | Control | 16.48151745 | 0.113205152 | 14.21603124 | 0.378384296 | 4.808164345 | 2.265486209 | 1.925403036 | 1.56E-07 | 6.807381134 | Unpaired t-test | 0.000178737 | permutation FDR (250 perm) | 1 | 7.01E-07 | FDR correction BH |
| Mtmr6~Q7KK51 | VMAT~Control | VMAT | Control | 19.15252607 | 0.068292 | 16.90438762 | 0.245012018 | 4.750694534 | 2.248138446 | 2.018531588 | 2.41E-09 | 8.617961821 | Unpaired t-test | 0.000103734 | permutation FDR (250 perm) | 1 | 2.49E-08 | FDR correction BH |
| Ack-like~A0A0B4KFU0 | VMAT~Control | VMAT | Control | 16.90727686 | 0.158290832 | 14.66638228 | 0.333194519 | 4.726900784 | 2.240894583 | 1.923565128 | 9.03E-08 | 7.044167541 | Unpaired t-test | 0.000182845 | permutation FDR (250 perm) | 1 | 4.38E-07 | FDR correction BH |
| Orco~Q9VNB5 | VMAT~Control | VMAT | Control | 15.87302336 | 0.194950242 | 13.64255954 | 0.324250099 | 4.69284829 | 2.230463822 | 1.907683229 | 1.20E-07 | 6.920247934 | Unpaired t-test | 0.000215606 | permutation FDR (250 perm) | 1 | 5.63E-07 | FDR correction BH |
| Hsl~Q7JR83 | VMAT~Control | VMAT | Control | 17.98742492 | 0.687815536 | 15.75929163 | 0.058580762 | 4.685273577 | 2.22813329 | 1.702536348 | 2.87E-05 | 4.542668249 | Unpaired t-test | 0.000712412 | permutation FDR (250 perm) | 1 | 6.40E-05 | FDR correction BH |
| BORCS6~Q9W0Q3 | VMAT~Control | VMAT | Control | 20.7332742 | 0.138866343 | 18.51434198 | 0.234340525 | 4.655487403 | 2.218932218 | 1.977977034 | 5.34E-09 | 8.272307633 | Unpaired t-test | 0.000128033 | permutation FDR (250 perm) | 1 | 4.59E-08 | FDR correction BH |
| Dmel\CG3702~Q9VQX6 | VMAT~Control | VMAT | Control | 20.18183603 | 0.172666761 | 17.96734178 | 0.197342943 | 4.641188388 | 2.214494258 | 1.98206315 | 3.76E-09 | 8.424935635 | Unpaired t-test | 0.000119959 | permutation FDR (250 perm) | 1 | 3.47E-08 | FDR correction BH |
| cpx~E1JJ35 | VMAT~Control | VMAT | Control | 23.19072698 | 0.078747964 | 20.99665435 | 0.08072517 | 4.575954271 | 2.194072635 | 2.088730223 | 9.89E-13 | 12.00469033 | Unpaired t-test | 5.65E-05 | permutation FDR (250 perm) | 1 | 9.70E-11 | FDR correction BH |
| Dmel\CG9205~Q9W0K9 | VMAT~Control | VMAT | Control | 15.58391519 | 0.276554766 | 13.39408815 | 0.422173244 | 4.562507849 | 2.189827041 | 1.786586325 | 2.10E-06 | 5.678680662 | Unpaired t-test | 0.00042908 | permutation FDR (250 perm) | 1 | 6.46E-06 | FDR correction BH |
| X11L~M9PF12 | VMAT~Control | VMAT | Control | 16.38642185 | 0.41979041 | 14.20216723 | 0.349311973 | 4.544919145 | 2.184254629 | 1.755506328 | 4.38E-06 | 5.358590622 | Unpaired t-test | 0.00053171 | permutation FDR (250 perm) | 1 | 1.23E-05 | FDR correction BH |
| Stam~Q9XTL2 | VMAT~Control | VMAT | Control | 19.18333375 | 0.10612286 | 17.0012457 | 0.176543888 | 4.538098915 | 2.182088055 | 1.998030894 | 4.08E-10 | 9.389283227 | Unpaired t-test | 0.000113813 | permutation FDR (250 perm) | 1 | 6.28E-09 | FDR correction BH |
| BORCS5~Q9VYT1 | VMAT~Control | VMAT | Control | 17.56225274 | 0.182767336 | 15.39964326 | 0.541839154 | 4.477239468 | 2.162609484 | 1.722190692 | 7.22E-06 | 5.141565209 | Unpaired t-test | 0.000638384 | permutation FDR (250 perm) | 1 | 1.91E-05 | FDR correction BH |
| EndoA~B5RIU6 | VMAT~Control | VMAT | Control | 20.85692189 | 0.096111585 | 18.69505172 | 0.266673463 | 4.474945663 | 2.161870165 | 1.918645035 | 1.01E-08 | 7.994364393 | Unpaired t-test | 0.000190965 | permutation FDR (250 perm) | 1 | 7.55E-08 | FDR correction BH |
| rdo~Q9VJA9 | VMAT~Control | VMAT | Control | 17.51644529 | 0.110075857 | 15.35501815 | 0.490982165 | 4.473571676 | 2.161427132 | 1.76439499 | 2.30E-06 | 5.638970427 | Unpaired t-test | 0.000501778 | permutation FDR (250 perm) | 1 | 6.95E-06 | FDR correction BH |
| Mob2~D6W4U9 | VMAT~Control | VMAT | Control | 15.40672156 | 0.071541599 | 13.266655 | 0.421446564 | 4.407823837 | 2.140066566 | 1.796604464 | 5.60E-07 | 6.251801277 | Unpaired t-test | 0.000396947 | permutation FDR (250 perm) | 1 | 2.05E-06 | FDR correction BH |
| CG34239~M9PEU1 | VMAT~Control | VMAT | Control | 15.69780265 | 0.138740825 | 13.57372732 | 0.165903144 | 4.359236051 | 2.124075327 | 1.936754298 | 8.58E-10 | 9.066332923 | Unpaired t-test | 0.000168638 | permutation FDR (250 perm) | 1 | 1.10E-08 | FDR correction BH |
| Trp1~X2J9A4 | VMAT~Control | VMAT | Control | 17.16192009 | 0.218718603 | 15.05453747 | 0.25232598 | 4.309088177 | 2.107382621 | 1.833565287 | 6.28E-08 | 7.202243505 | Unpaired t-test | 0.000328907 | permutation FDR (250 perm) | 1 | 3.23E-07 | FDR correction BH |
| rg~X2JAH8 | VMAT~Control | VMAT | Control | 18.10160407 | 0.105086413 | 16.00981161 | 0.080187676 | 4.262773678 | 2.091792461 | 1.975037133 | 7.71E-12 | 11.11290613 | Unpaired t-test | 0.000132163 | permutation FDR (250 perm) | 1 | 3.24E-10 | FDR correction BH |
| Dmel\CG8245~A1Z6G9 | VMAT~Control | VMAT | Control | 16.72269136 | 0.151517987 | 14.63854018 | 0.218161373 | 4.240255456 | 2.084151183 | 1.862866361 | 7.69E-09 | 8.114138166 | Unpaired t-test | 0.000274169 | permutation FDR (250 perm) | 1 | 6.14E-08 | FDR correction BH |
| hook~Q24185 | VMAT~Control | VMAT | Control | 16.21046738 | 0.263657643 | 14.1601539 | 0.586826566 | 4.141959584 | 2.050313476 | 1.592218697 | 3.19E-05 | 4.495747664 | Unpaired t-test | 0.001216958 | permutation FDR (250 perm) | 1 | 7.07E-05 | FDR correction BH |
| lap~B7Z0U7 | VMAT~Control | VMAT | Control | 21.28868537 | 0.11080055 | 19.25137786 | 0.160537835 | 4.104787426 | 2.037307511 | 1.873843954 | 4.69E-10 | 9.32875795 | Unpaired t-test | 0.000262161 | permutation FDR (250 perm) | 1 | 6.86E-09 | FDR correction BH |
| Dmel\CG11883~A8DY95 | VMAT~Control | VMAT | Control | 17.70133034 | 0.164560036 | 15.67225304 | 0.135431891 | 4.081437316 | 2.0290773 | 1.852510941 | 1.16E-09 | 8.933861738 | Unpaired t-test | 0.000294479 | permutation FDR (250 perm) | 1 | 1.36E-08 | FDR correction BH |
| Mvl~A0A0B4K7L4 | VMAT~Control | VMAT | Control | 15.4584618 | 0.171165874 | 13.43637086 | 0.220596474 | 4.06172043 | 2.022090942 | 1.797624486 | 1.67E-08 | 7.776891524 | Unpaired t-test | 0.00039084 | permutation FDR (250 perm) | 1 | 1.13E-07 | FDR correction BH |
| OstDelta~A0A0B4LFR4 | VMAT~Control | VMAT | Control | 15.83094712 | 0.084731265 | 13.81098537 | 0.249493266 | 4.055730386 | 2.019961749 | 1.807029176 | 9.63E-09 | 8.016185909 | Unpaired t-test | 0.000384929 | permutation FDR (250 perm) | 1 | 7.29E-08 | FDR correction BH |
| Snx16~Q7JR96 | VMAT~Control | VMAT | Control | 15.76051046 | 0.06523807 | 13.74601311 | 0.302468716 | 4.040397829 | 2.014497352 | 1.769619673 | 4.66E-08 | 7.331673362 | Unpaired t-test | 0.000497715 | permutation FDR (250 perm) | 1 | 2.57E-07 | FDR correction BH |
| MCU~Q8IQ70-3 | VMAT~Control | VMAT | Control | 15.91170626 | 0.105056358 | 13.89905761 | 0.714495372 | 4.035223683 | 2.012648648 | 1.521313591 | 9.73E-05 | 4.011713479 | Unpaired t-test | 0.00176813 | permutation FDR (250 perm) | 1 | 0.000191906 | FDR correction BH |
| mRpL3~Q9VXQ0 | VMAT~Control | VMAT | Control | 16.3997484 | 0.166678525 | 14.39301059 | 1.087817053 | 4.018724871 | 2.006737811 | 1.344850587 | 0.00222371 | 2.652921919 | Unpaired t-test | 0.003908738 | permutation FDR (250 perm) | 1 | 0.003364147 | FDR correction BH |
| Vps45~Q9VHB5 | VMAT~Control | VMAT | Control | 15.35290184 | 0.326940843 | 13.34885384 | 0.334139733 | 4.011239207 | 2.004048003 | 1.657519613 | 2.34E-06 | 5.630936456 | Unpaired t-test | 0.00087707 | permutation FDR (250 perm) | 1 | 7.04E-06 | FDR correction BH |
| Frq2~Q9VWX8 | VMAT~Control | VMAT | Control | 15.37974065 | 0.071597436 | 13.39938912 | 0.261637291 | 3.945892176 | 1.980351534 | 1.76610571 | 1.55E-08 | 7.810706231 | Unpaired t-test | 0.000497715 | permutation FDR (250 perm) | 1 | 1.06E-07 | FDR correction BH |
| unc-13~Q8IM87 | VMAT~Control | VMAT | Control | 15.72969574 | 0.19470355 | 13.75355355 | 0.680715393 | 3.934396038 | 1.976142187 | 1.500905633 | 9.62E-05 | 4.017013926 | Unpaired t-test | 0.001958661 | permutation FDR (250 perm) | 1 | 0.000189896 | FDR correction BH |
| Dmel\CG11396~Q9VPB6 | VMAT~Control | VMAT | Control | 18.21512128 | 0.08358154 | 16.24790376 | 0.162475698 | 3.910132563 | 1.967217519 | 1.81861492 | 3.48E-10 | 9.458619805 | Unpaired t-test | 0.000365306 | permutation FDR (250 perm) | 1 | 5.61E-09 | FDR correction BH |
| Snap25~P36975 | VMAT~Control | VMAT | Control | 21.93472427 | 0.12409176 | 19.96788156 | 0.161905091 | 3.909116845 | 1.966842708 | 1.802413541 | 1.03E-09 | 8.987725636 | Unpaired t-test | 0.000389002 | permutation FDR (250 perm) | 1 | 1.25E-08 | FDR correction BH |
| mRpS25~Q9VY28 | VMAT~Control | VMAT | Control | 16.43639431 | 0.402519288 | 14.47519795 | 0.887636439 | 3.893847423 | 1.961196354 | 1.365857758 | 0.001144021 | 2.941566142 | Unpaired t-test | 0.003599417 | permutation FDR (250 perm) | 1 | 0.00181593 | FDR correction BH |
| veli~Q8IMT8 | VMAT~Control | VMAT | Control | 15.37785648 | 1.89835673 | 13.41840826 | 0.483532534 | 3.889132058 | 1.959448223 | 1.044438654 | 0.049286504 | 1.307271984 | Unpaired t-test | 0.011988658 | permutation FDR (250 perm) | 1 | 0.06158197 | FDR correction BH |
| Usp32~M9PD06-3 | VMAT~Control | VMAT | Control | 18.13676674 | 0.333151087 | 16.18019995 | 0.352053256 | 3.881372207 | 1.956566788 | 1.608009675 | 4.03E-06 | 5.394598466 | Unpaired t-test | 0.001118881 | permutation FDR (250 perm) | 1 | 1.14E-05 | FDR correction BH |
| nwk~X2JAU8-3 | VMAT~Control | VMAT | Control | 18.3571544 | 0.207791349 | 16.40342376 | 0.475268368 | 3.873749455 | 1.953730647 | 1.585855139 | 7.48E-06 | 5.125899964 | Unpaired t-test | 0.001267564 | permutation FDR (250 perm) | 1 | 1.95E-05 | FDR correction BH |
| Dmel\CG8199~Q9VHB8 | VMAT~Control | VMAT | Control | 15.88087337 | 0.119130661 | 13.9278172 | 0.629351246 | 3.871938863 | 1.953056173 | 1.518172035 | 4.64E-05 | 4.33381654 | Unpaired t-test | 0.001817464 | permutation FDR (250 perm) | 1 | 9.84E-05 | FDR correction BH |
| mgl~A8JTM7 | VMAT~Control | VMAT | Control | 16.43660555 | 0.258033654 | 14.48433407 | 0.815051132 | 3.869833454 | 1.952271478 | 1.412302643 | 0.000459836 | 3.337396712 | Unpaired t-test | 0.002881951 | permutation FDR (250 perm) | 1 | 0.000780429 | FDR correction BH |
| Dysb~M9PI95 | VMAT~Control | VMAT | Control | 15.50567055 | 0.320179387 | 13.56395841 | 0.371914696 | 3.841612857 | 1.941712137 | 1.592259058 | 4.83E-06 | 5.316489609 | Unpaired t-test | 0.001216958 | permutation FDR (250 perm) | 1 | 1.34E-05 | FDR correction BH |
| brp~A0A0B4K843 | VMAT~Control | VMAT | Control | 17.89060313 | 0.36392749 | 15.95093903 | 0.114392906 | 3.836163209 | 1.939664101 | 1.656976894 | 4.85E-07 | 6.314657938 | Unpaired t-test | 0.000887105 | permutation FDR (250 perm) | 1 | 1.81E-06 | FDR correction BH |
| Lrp4~B5X533 | VMAT~Control | VMAT | Control | 16.89656684 | 0.086128089 | 14.96136607 | 0.399230513 | 3.824313438 | 1.93520077 | 1.636327415 | 9.33E-07 | 6.029935055 | Unpaired t-test | 0.000985478 | permutation FDR (250 perm) | 1 | 3.18E-06 | FDR correction BH |
| lap~Q9VI75 | VMAT~Control | VMAT | Control | 19.03367462 | 0.139067836 | 17.10239529 | 0.141369239 | 3.813932544 | 1.931279327 | 1.773956076 | 9.32E-10 | 9.030392116 | Unpaired t-test | 0.000483494 | permutation FDR (250 perm) | 1 | 1.16E-08 | FDR correction BH |
| Irk1~A8JR87 | VMAT~Control | VMAT | Control | 19.2809905 | 0.095190876 | 17.36444646 | 0.123203876 | 3.775176377 | 1.916544049 | 1.791785105 | 9.28E-11 | 10.03233237 | Unpaired t-test | 0.000410778 | permutation FDR (250 perm) | 1 | 2.02E-09 | FDR correction BH |
| bb8~Q9VCN3 | VMAT~Control | VMAT | Control | 15.44179843 | 0.410275048 | 13.52599114 | 0.383892907 | 3.773248958 | 1.915807292 | 1.531082301 | 1.79E-05 | 4.747170503 | Unpaired t-test | 0.001683482 | permutation FDR (250 perm) | 1 | 4.18E-05 | FDR correction BH |
| Rabex-5~Q9W0H9 | VMAT~Control | VMAT | Control | 15.79497032 | 0.29490962 | 13.88154199 | 0.625799436 | 3.767032124 | 1.913428336 | 1.461317914 | 0.000103516 | 3.984991111 | Unpaired t-test | 0.002337255 | permutation FDR (250 perm) | 1 | 0.000203234 | FDR correction BH |
| Nrx-1~A0A0B4KH61 | VMAT~Control | VMAT | Control | 15.41507748 | 0.273149117 | 13.50237807 | 0.446073304 | 3.765129304 | 1.912699412 | 1.550100553 | 9.72E-06 | 5.01238681 | Unpaired t-test | 0.00150297 | permutation FDR (250 perm) | 1 | 2.45E-05 | FDR correction BH |
| Rim~A0A0B4KG75 | VMAT~Control | VMAT | Control | 15.64643461 | 0.153349259 | 13.73560901 | 0.359758586 | 3.760242223 | 1.910825599 | 1.626379152 | 7.02E-07 | 6.153538307 | Unpaired t-test | 0.001032516 | permutation FDR (250 perm) | 1 | 2.47E-06 | FDR correction BH |
| GAPcenA~A8JNN5 | VMAT~Control | VMAT | Control | 17.09382035 | 0.064040936 | 15.18857064 | 0.215652544 | 3.745737273 | 1.905249712 | 1.731092418 | 3.66E-09 | 8.436955136 | Unpaired t-test | 0.000616785 | permutation FDR (250 perm) | 1 | 3.42E-08 | FDR correction BH |
| Teh4~Q4V5U6 | VMAT~Control | VMAT | Control | 17.07588574 | 0.142225479 | 15.17522089 | 0.717351932 | 3.733852277 | 1.900664851 | 1.432243696 | 0.000170281 | 3.768833391 | Unpaired t-test | 0.002636675 | permutation FDR (250 perm) | 1 | 0.000317373 | FDR correction BH |
| x16~M9PC85 | VMAT~Control | VMAT | Control | 16.46710966 | 0.169013237 | 14.56703254 | 1.354389112 | 3.732331491 | 1.900077126 | 1.179879646 | 0.011007252 | 1.958321082 | Unpaired t-test | 0.007452107 | permutation FDR (250 perm) | 1 | 0.01510948 | FDR correction BH |
| Dmel\CG10616~M9NE87 | VMAT~Control | VMAT | Control | 16.21052518 | 0.390320697 | 14.31068624 | 0.82157269 | 3.731715348 | 1.899838943 | 1.350491825 | 0.000880158 | 3.055439217 | Unpaired t-test | 0.003811559 | permutation FDR (250 perm) | 1 | 0.001418133 | FDR correction BH |
| Pldn~Q9VTM0 | VMAT~Control | VMAT | Control | 15.71659026 | 0.49274552 | 13.82784751 | 0.545316589 | 3.703123713 | 1.888742747 | 1.421513075 | 0.000186091 | 3.730275648 | Unpaired t-test | 0.002765625 | permutation FDR (250 perm) | 1 | 0.000344927 | FDR correction BH |
| SPoCk~Q7KTU2 | VMAT~Control | VMAT | Control | 18.35975757 | 0.053986738 | 16.47554434 | 0.106347365 | 3.691515535 | 1.88421323 | 1.788803318 | 7.84E-12 | 11.10577829 | Unpaired t-test | 0.000425013 | permutation FDR (250 perm) | 1 | 3.24E-10 | FDR correction BH |
| Vha100-2~Q9VE75 | VMAT~Control | VMAT | Control | 16.07168974 | 0.513705078 | 14.18825139 | 1.007218969 | 3.689533349 | 1.883438356 | 1.250918175 | 0.003943808 | 2.404084256 | Unpaired t-test | 0.005889961 | permutation FDR (250 perm) | 1 | 0.005755563 | FDR correction BH |
| CT39634~Q7KUK9 | VMAT~Control | VMAT | Control | 15.2727951 | 2.918953112 | 13.39123723 | 0.315050774 | 3.68472735 | 1.881557872 | 0.813478817 | 0.182358255 | 0.739074573 | Unpaired t-test | 0.02506938 | permutation FDR (250 perm) | 1 | 0.209912632 | FDR correction BH |
| Flo1~O61491 | VMAT~Control | VMAT | Control | 15.60257082 | 0.408799415 | 13.73223016 | 0.315626552 | 3.656189033 | 1.870340663 | 1.519403199 | 1.06E-05 | 4.97530521 | Unpaired t-test | 0.001809571 | permutation FDR (250 perm) | 1 | 2.63E-05 | FDR correction BH |
| Rab8~O18338 | VMAT~Control | VMAT | Control | 16.25418409 | 0.088723793 | 14.39403398 | 1.104801498 | 3.630454345 | 1.86015011 | 1.243687757 | 0.003766585 | 2.424052197 | Unpaired t-test | 0.006042471 | permutation FDR (250 perm) | 1 | 0.005507169 | FDR correction BH |
| Dmel\CG11309~Q9VP52 | VMAT~Control | VMAT | Control | 15.59352359 | 0.316572899 | 13.7335821 | 0.131025012 | 3.629929402 | 1.85994149 | 1.612820764 | 2.62E-07 | 6.581413586 | Unpaired t-test | 0.001086913 | permutation FDR (250 perm) | 1 | 1.08E-06 | FDR correction BH |
| gammaSnap2~Q9VH25 | VMAT~Control | VMAT | Control | 16.13180039 | 0.182136801 | 14.2727329 | 0.972878967 | 3.627731004 | 1.859067484 | 1.288653229 | 0.001828933 | 2.737802132 | Unpaired t-test | 0.005002419 | permutation FDR (250 perm) | 1 | 0.002795655 | FDR correction BH |
| porin~M9PD75 | VMAT~Control | VMAT | Control | 15.59165167 | 0.133435503 | 13.73652089 | 0.352025089 | 3.617845463 | 1.855130784 | 1.587806504 | 6.49E-07 | 6.187854257 | Unpaired t-test | 0.001258853 | permutation FDR (250 perm) | 1 | 2.31E-06 | FDR correction BH |
| Rab2~O18333 | VMAT~Control | VMAT | Control | 15.65648644 | 0.244703437 | 13.81200968 | 0.106748856 | 3.59122677 | 1.844476756 | 1.647745234 | 2.63E-08 | 7.579516526 | Unpaired t-test | 0.000942828 | permutation FDR (250 perm) | 1 | 1.62E-07 | FDR correction BH |
| stnB~X2JGF5 | VMAT~Control | VMAT | Control | 22.01815745 | 0.128638967 | 20.18288063 | 0.128777976 | 3.568398712 | 1.835276822 | 1.697126646 | 6.63E-10 | 9.178327081 | Unpaired t-test | 0.000725806 | permutation FDR (250 perm) | 1 | 9.14E-09 | FDR correction BH |
| c11.1~Q9W350 | VMAT~Control | VMAT | Control | 17.96839336 | 0.06576207 | 16.13501713 | 0.228424581 | 3.56370082 | 1.833376225 | 1.657208719 | 9.08E-09 | 8.041762684 | Unpaired t-test | 0.000881084 | permutation FDR (250 perm) | 1 | 6.99E-08 | FDR correction BH |
| Vps53~Q9VQY8 | VMAT~Control | VMAT | Control | 16.85615304 | 0.072768378 | 15.02328234 | 0.084450988 | 3.562452298 | 1.832870696 | 1.745833561 | 5.27E-12 | 11.27787334 | Unpaired t-test | 0.000561581 | permutation FDR (250 perm) | 1 | 2.55E-10 | FDR correction BH |
| Dmel\CG15099~A0A0B4KG81 | VMAT~Control | VMAT | Control | 16.26453174 | 0.391253108 | 14.44004282 | 0.399782503 | 3.54181515 | 1.824488919 | 1.459402037 | 2.62E-05 | 4.581369216 | Unpaired t-test | 0.002357668 | permutation FDR (250 perm) | 1 | 5.91E-05 | FDR correction BH |
| nwk~X2JAU8-7 | VMAT~Control | VMAT | Control | 18.09227158 | 0.181557914 | 16.26860027 | 0.162206098 | 3.539808468 | 1.823671301 | 1.644606783 | 1.21E-08 | 7.918622229 | Unpaired t-test | 0.000950853 | permutation FDR (250 perm) | 1 | 8.60E-08 | FDR correction BH |
| Dmel\CG31140~Q7KS34 | VMAT~Control | VMAT | Control | 17.53435333 | 0.123561204 | 15.71786473 | 0.480184369 | 3.522228718 | 1.816488595 | 1.486803861 | 9.56E-06 | 5.019497381 | Unpaired t-test | 0.002047198 | permutation FDR (250 perm) | 1 | 2.42E-05 | FDR correction BH |
| trc~M9PG43 | VMAT~Control | VMAT | Control | 16.01076983 | 0.054267681 | 14.20022145 | 0.239688266 | 3.507755956 | 1.810548379 | 1.631264411 | 1.42E-08 | 7.848872928 | Unpaired t-test | 0.001007007 | permutation FDR (250 perm) | 1 | 9.86E-08 | FDR correction BH |
| Tret1-1~A1Z8N1 | VMAT~Control | VMAT | Control | 18.60481958 | 0.060605767 | 16.79785713 | 0.171622615 | 3.49904799 | 1.806962451 | 1.67095193 | 7.72E-10 | 9.112296654 | Unpaired t-test | 0.000819689 | permutation FDR (250 perm) | 1 | 1.00E-08 | FDR correction BH |
| sicily~X2JJK8 | VMAT~Control | VMAT | Control | 15.9015013 | 0.223952394 | 14.09565336 | 0.317780977 | 3.496345942 | 1.805847938 | 1.538382216 | 1.12E-06 | 5.950215952 | Unpaired t-test | 0.001610287 | permutation FDR (250 perm) | 1 | 3.74E-06 | FDR correction BH |
| Spase25~B4F5U4 | VMAT~Control | VMAT | Control | 16.61709286 | 0.223690606 | 14.81404647 | 0.219409622 | 3.489563043 | 1.803046396 | 1.58144328 | 1.51E-07 | 6.820521671 | Unpaired t-test | 0.001287494 | permutation FDR (250 perm) | 1 | 6.84E-07 | FDR correction BH |
| Pi4KIIalpha~Q9VND3 | VMAT~Control | VMAT | Control | 19.68697002 | 0.048627975 | 17.89991488 | 0.177171015 | 3.451097279 | 1.787055141 | 1.651372506 | 9.44E-10 | 9.02513074 | Unpaired t-test | 0.000922768 | permutation FDR (250 perm) | 1 | 1.17E-08 | FDR correction BH |
| X11Lbeta~Q9W2S5 | VMAT~Control | VMAT | Control | 15.67435873 | 0.210780138 | 13.88982011 | 0.529570172 | 3.445082709 | 1.784538619 | 1.422055197 | 3.71E-05 | 4.430446568 | Unpaired t-test | 0.002759766 | permutation FDR (250 perm) | 1 | 8.11E-05 | FDR correction BH |
| Sirt2~R4HZM7 | VMAT~Control | VMAT | Control | 15.10987076 | 0.262992368 | 13.33070555 | 0.303719271 | 3.432275148 | 1.779165211 | 1.508186179 | 1.74E-06 | 5.759670214 | Unpaired t-test | 0.001893596 | permutation FDR (250 perm) | 1 | 5.47E-06 | FDR correction BH |
| CtsB~Q9VY87 | VMAT~Control | VMAT | Control | 15.41194936 | 0.252164662 | 13.64345928 | 0.782209621 | 3.406971968 | 1.768490077 | 1.293188009 | 0.000711074 | 3.148085337 | Unpaired t-test | 0.004890179 | permutation FDR (250 perm) | 1 | 0.001161601 | FDR correction BH |
| LpR2~Q86B77 | VMAT~Control | VMAT | Control | 15.46861944 | 0.173159942 | 13.70085215 | 0.343896337 | 3.405265505 | 1.767767288 | 1.508087827 | 1.25E-06 | 5.903567986 | Unpaired t-test | 0.001895567 | permutation FDR (250 perm) | 1 | 4.08E-06 | FDR correction BH |
| slo~A0A0B4KHT2 | VMAT~Control | VMAT | Control | 15.29321406 | 0.369902071 | 13.52975917 | 0.431914788 | 3.39510193 | 1.763454889 | 1.405911884 | 4.02E-05 | 4.395374456 | Unpaired t-test | 0.002976608 | permutation FDR (250 perm) | 1 | 8.70E-05 | FDR correction BH |
| smal~A8DYR2 | VMAT~Control | VMAT | Control | 15.6578787 | 0.12481532 | 13.89772031 | 0.211235412 | 3.387353113 | 1.760158388 | 1.586119431 | 1.83E-08 | 7.737217279 | Unpaired t-test | 0.001267564 | permutation FDR (250 perm) | 1 | 1.22E-07 | FDR correction BH |
| ca~Q9VAG5 | VMAT~Control | VMAT | Control | 15.66741059 | 0.329563869 | 13.91633972 | 0.157949838 | 3.36608328 | 1.751070871 | 1.505082627 | 8.42E-07 | 6.074745649 | Unpaired t-test | 0.001915271 | permutation FDR (250 perm) | 1 | 2.90E-06 | FDR correction BH |
| unc-104~A0A126GUN3 | VMAT~Control | VMAT | Control | 16.50130964 | 0.144737439 | 14.75774025 | 0.306254595 | 3.348626317 | 1.743569391 | 1.514190037 | 4.32E-07 | 6.364471869 | Unpaired t-test | 0.001845244 | permutation FDR (250 perm) | 1 | 1.64E-06 | FDR correction BH |
| CG5741~M9PEU6 | VMAT~Control | VMAT | Control | 19.3358068 | 0.214809501 | 17.59848824 | 0.31702368 | 3.334148962 | 1.737318562 | 1.483292343 | 1.39E-06 | 5.855873196 | Unpaired t-test | 0.002093366 | permutation FDR (250 perm) | 1 | 4.50E-06 | FDR correction BH |
| CG6087~Q9VKM8 | VMAT~Control | VMAT | Control | 16.61859654 | 0.393374957 | 14.88806881 | 0.151221239 | 3.318491863 | 1.730527737 | 1.456092553 | 3.46E-06 | 5.461450121 | Unpaired t-test | 0.002389025 | permutation FDR (250 perm) | 1 | 9.99E-06 | FDR correction BH |
| aux~Q9VMY8 | VMAT~Control | VMAT | Control | 18.23869692 | 0.153949317 | 16.50823905 | 0.206382716 | 3.318331159 | 1.73045787 | 1.551775496 | 3.43E-08 | 7.464243591 | Unpaired t-test | 0.00148662 | permutation FDR (250 perm) | 1 | 2.00E-07 | FDR correction BH |
| Sap47~Q960T2-5 | VMAT~Control | VMAT | Control | 24.2033317 | 0.110431032 | 22.48821414 | 0.199917426 | 3.28323395 | 1.715117554 | 1.556171573 | 1.18E-08 | 7.929302188 | Unpaired t-test | 0.00146356 | permutation FDR (250 perm) | 1 | 8.42E-08 | FDR correction BH |
| Dmel\CG4658~Q9VL57 | VMAT~Control | VMAT | Control | 16.29409568 | 0.137367657 | 14.58232364 | 0.200514261 | 3.275629186 | 1.711772048 | 1.543948545 | 2.19E-08 | 7.659864493 | Unpaired t-test | 0.001551707 | permutation FDR (250 perm) | 1 | 1.39E-07 | FDR correction BH |
| l(3)76BDm~Q9VW22 | VMAT~Control | VMAT | Control | 17.81488883 | 0.073341126 | 16.11498562 | 0.061605228 | 3.248791613 | 1.699903208 | 1.630079007 | 2.47E-12 | 11.60776765 | Unpaired t-test | 0.001019019 | permutation FDR (250 perm) | 1 | 1.59E-10 | FDR correction BH |
| Dmel\CG34130~Q0KI07 | VMAT~Control | VMAT | Control | 15.15119135 | 0.239107181 | 13.45476847 | 0.194497244 | 3.240963736 | 1.696422878 | 1.490913145 | 2.30E-07 | 6.637751481 | Unpaired t-test | 0.002019666 | permutation FDR (250 perm) | 1 | 9.70E-07 | FDR correction BH |
| Galphas~A0A0B4LGI1 | VMAT~Control | VMAT | Control | 15.05627314 | 0.130755393 | 13.36307873 | 0.415542897 | 3.233719203 | 1.693194409 | 1.417113329 | 5.66E-06 | 5.2473126 | Unpaired t-test | 0.00281308 | permutation FDR (250 perm) | 1 | 1.55E-05 | FDR correction BH |
| Naprt~M9PEF6 | VMAT~Control | VMAT | Control | 16.49627178 | 0.229751751 | 14.80377006 | 1.158676584 | 3.232166947 | 1.692501718 | 1.107466435 | 0.009429257 | 2.025522517 | Unpaired t-test | 0.009743346 | permutation FDR (250 perm) | 1 | 0.013095263 | FDR correction BH |
| None~TurboID-HA-VMAT_bait | VMAT~Control | VMAT | Control | 23.16403155 | 0.133709537 | 21.48760474 | 0.088581513 | 3.196353153 | 1.676426815 | 1.564227074 | 4.66E-10 | 9.331939426 | Unpaired t-test | 0.001392254 | permutation FDR (250 perm) | 1 | 6.85E-09 | FDR correction BH |
| unc-13-4A~Q9VS41 | VMAT~Control | VMAT | Control | 18.8022238 | 0.138984614 | 17.1290668 | 0.251344994 | 3.189116956 | 1.673157007 | 1.482709836 | 1.35E-07 | 6.871103728 | Unpaired t-test | 0.002093366 | permutation FDR (250 perm) | 1 | 6.22E-07 | FDR correction BH |
| Dmel\CG32544~Q8IQX3 | VMAT~Control | VMAT | Control | 21.54723677 | 0.099017151 | 19.87660257 | 0.200475896 | 3.183545107 | 1.670634206 | 1.518765299 | 1.24E-08 | 7.908009639 | Unpaired t-test | 0.001813518 | permutation FDR (250 perm) | 1 | 8.76E-08 | FDR correction BH |
| stnA~A4V4W0 | VMAT~Control | VMAT | Control | 20.39185299 | 0.083467693 | 18.72471776 | 0.187766844 | 3.175833402 | 1.667135233 | 1.526827749 | 5.56E-09 | 8.255289004 | Unpaired t-test | 0.001708354 | permutation FDR (250 perm) | 1 | 4.70E-08 | FDR correction BH |
| Ifrd1~Q9VQI5 | VMAT~Control | VMAT | Control | 16.02893016 | 0.333161008 | 14.36655488 | 0.904190568 | 3.165372505 | 1.662375288 | 1.161734579 | 0.003172983 | 2.498532304 | Unpaired t-test | 0.007952267 | permutation FDR (250 perm) | 1 | 0.004685955 | FDR correction BH |
| Dmel\CG30392~Q8MLW4 | VMAT~Control | VMAT | Control | 15.00369481 | 2.045002204 | 13.34205647 | 0.273754453 | 3.163756006 | 1.661638341 | 0.864216499 | 0.101915585 | 0.991759397 | Unpaired t-test | 0.020508113 | permutation FDR (250 perm) | 1 | 0.122091241 | FDR correction BH |
| ICA69~Q9VP78 | VMAT~Control | VMAT | Control | 15.41004564 | 0.101692603 | 13.75963547 | 0.309846397 | 3.13922877 | 1.650410169 | 1.440349889 | 5.06E-07 | 6.2957864 | Unpaired t-test | 0.002545988 | permutation FDR (250 perm) | 1 | 1.87E-06 | FDR correction BH |
| ValRS-m~Q9VSR7 | VMAT~Control | VMAT | Control | 15.52902032 | 0.416605575 | 13.87969147 | 0.386751663 | 3.13687676 | 1.64932885 | 1.315024381 | 7.00E-05 | 4.154729795 | Unpaired t-test | 0.004465116 | permutation FDR (250 perm) | 1 | 0.000142231 | FDR correction BH |
| Dmel\CG13833~Q9VCS2 | VMAT~Control | VMAT | Control | 15.74867647 | 0.148839104 | 14.11390845 | 0.814794975 | 3.105376129 | 1.634768021 | 1.192898198 | 0.001307647 | 2.883509492 | Unpaired t-test | 0.007112176 | permutation FDR (250 perm) | 1 | 0.002048038 | FDR correction BH |
| Lnpk~Q7K2N0 | VMAT~Control | VMAT | Control | 15.32310733 | 0.869669333 | 13.69217575 | 0.414293229 | 3.097129242 | 1.630931587 | 1.13987013 | 0.003567986 | 2.447576831 | Unpaired t-test | 0.008734382 | permutation FDR (250 perm) | 1 | 0.005239576 | FDR correction BH |
| Pde1c~B7YZV1 | VMAT~Control | VMAT | Control | 16.30133818 | 0.425726535 | 14.67395832 | 0.443227767 | 3.089513878 | 1.627379854 | 1.27653334 | 0.000146824 | 3.833202972 | Unpaired t-test | 0.005243113 | permutation FDR (250 perm) | 1 | 0.000278056 | FDR correction BH |
| CG6001~A0A0B4LHR2 | VMAT~Control | VMAT | Control | 17.01808614 | 0.139447869 | 15.40301773 | 0.701796015 | 3.063261239 | 1.615068407 | 1.223547287 | 0.000501251 | 3.29994475 | Unpaired t-test | 0.006464802 | permutation FDR (250 perm) | 1 | 0.000845233 | FDR correction BH |
| CG6550~Q7K4W1 | VMAT~Control | VMAT | Control | 16.09099194 | 0.281110426 | 14.47747164 | 0.103050169 | 3.059975919 | 1.613520299 | 1.422986303 | 2.81E-07 | 6.551541151 | Unpaired t-test | 0.002742188 | permutation FDR (250 perm) | 1 | 1.14E-06 | FDR correction BH |
| nwk~X2JAU8-5 | VMAT~Control | VMAT | Control | 20.11805869 | 0.153830964 | 18.50882323 | 0.201478334 | 3.050901213 | 1.609235467 | 1.44538069 | 5.93E-08 | 7.226764404 | Unpaired t-test | 0.002484581 | permutation FDR (250 perm) | 1 | 3.11E-07 | FDR correction BH |
| Rbp~E1JIL7 | VMAT~Control | VMAT | Control | 20.78834348 | 0.073584484 | 19.18327565 | 0.108866713 | 3.042100527 | 1.605067828 | 1.515981166 | 1.00E-10 | 9.999151241 | Unpaired t-test | 0.001841145 | permutation FDR (250 perm) | 1 | 2.11E-09 | FDR correction BH |
| gry~Q8IRE3 | VMAT~Control | VMAT | Control | 19.66806951 | 0.034110958 | 18.06451954 | 0.074460526 | 3.038901614 | 1.603549968 | 1.546890971 | 9.29E-13 | 12.03199129 | Unpaired t-test | 0.001523998 | permutation FDR (250 perm) | 1 | 9.51E-11 | FDR correction BH |
| RagC-D~Q7K519 | VMAT~Control | VMAT | Control | 16.98187239 | 0.279667626 | 15.37907927 | 0.104631388 | 3.037307802 | 1.602793119 | 1.413974136 | 2.92E-07 | 6.535273191 | Unpaired t-test | 0.002852123 | permutation FDR (250 perm) | 1 | 1.17E-06 | FDR correction BH |
| Atg4b~Q9VF80 | VMAT~Control | VMAT | Control | 15.72767705 | 0.365147402 | 14.12776728 | 0.576909625 | 3.031243546 | 1.599909771 | 1.225666779 | 0.000378911 | 3.421462926 | Unpaired t-test | 0.006416586 | permutation FDR (250 perm) | 1 | 0.000653927 | FDR correction BH |
| Dmel\CG7071~Q8SXS0 | VMAT~Control | VMAT | Control | 15.71433074 | 0.082962232 | 14.11469118 | 0.703455001 | 3.030675863 | 1.599639561 | 1.214816266 | 0.000499417 | 3.301536274 | Unpaired t-test | 0.006613976 | permutation FDR (250 perm) | 1 | 0.000842745 | FDR correction BH |
| Dmel\CG7556~Q9VWL4 | VMAT~Control | VMAT | Control | 16.63968087 | 0.350988766 | 15.04131409 | 0.705281214 | 3.028003291 | 1.598366773 | 1.181952184 | 0.001080002 | 2.966575605 | Unpaired t-test | 0.007415429 | permutation FDR (250 perm) | 1 | 0.001720111 | FDR correction BH |
| Dmel\CG6607~Q9VC66 | VMAT~Control | VMAT | Control | 17.56003913 | 0.142930953 | 15.97871248 | 0.176912691 | 2.992448981 | 1.58132665 | 1.435334877 | 2.48E-08 | 7.606077251 | Unpaired t-test | 0.002604697 | permutation FDR (250 perm) | 1 | 1.53E-07 | FDR correction BH |
| brp~A0A0B4K7K9 | VMAT~Control | VMAT | Control | 21.10630752 | 0.171202132 | 19.52778079 | 0.194358464 | 2.986646988 | 1.578526726 | 1.414663298 | 8.76E-08 | 7.057557562 | Unpaired t-test | 0.002848219 | permutation FDR (250 perm) | 1 | 4.27E-07 | FDR correction BH |
| Vps16A~Q9VHG1 | VMAT~Control | VMAT | Control | 17.28320706 | 0.111433712 | 15.70619324 | 0.304398352 | 2.983516632 | 1.577013819 | 1.377345443 | 7.31E-07 | 6.136139774 | Unpaired t-test | 0.003433577 | permutation FDR (250 perm) | 1 | 2.56E-06 | FDR correction BH |
| l(2)37Bb~Q9VJ10 | VMAT~Control | VMAT | Control | 15.36016694 | 0.10364115 | 13.79178835 | 0.394791987 | 2.965712186 | 1.568378595 | 1.326280793 | 6.27E-06 | 5.202919149 | Unpaired t-test | 0.004262009 | permutation FDR (250 perm) | 1 | 1.69E-05 | FDR correction BH |
| Pen~P52295 | VMAT~Control | VMAT | Control | 16.31131583 | 0.338898141 | 14.74485908 | 0.727106318 | 2.961764124 | 1.566456748 | 1.152859493 | 0.001407135 | 2.851664295 | Unpaired t-test | 0.008282443 | permutation FDR (250 perm) | 1 | 0.002193639 | FDR correction BH |
| Gos28~Q9VE50 | VMAT~Control | VMAT | Control | 17.9733159 | 0.139843785 | 16.40699382 | 0.461546689 | 2.961487675 | 1.566322082 | 1.288436654 | 2.72E-05 | 4.565640103 | Unpaired t-test | 0.005006289 | permutation FDR (250 perm) | 1 | 6.09E-05 | FDR correction BH |
| CG13811~M9PFQ9 | VMAT~Control | VMAT | Control | 16.77114788 | 0.073357003 | 15.20531547 | 0.164232437 | 2.960482662 | 1.565832405 | 1.449253423 | 2.80E-09 | 8.553452164 | Unpaired t-test | 0.002431718 | permutation FDR (250 perm) | 1 | 2.79E-08 | FDR correction BH |

| | | | | | | | | | | | | | | | | | | |
|---|---|---|---|---|---|---|---|---|---|---|---|---|---|---|---|---|---|---|
| Dmel\CG14408~Q9VY16 | VMAT~Control | VMAT | Control | 17.516874 | 0.120516601 | 15.95660098 | 0.167519562 | 2.949096463 | 1.560273013 | 1.428442327 | 1.10E-08 | 7.957833012 | Unpaired t-test | 0.002687531 | permutation FDR (250 perm) | 1 | 7.96E-08 | FDR correction BH |
| Dmel\CG1486~Q9VRG8 | VMAT~Control | VMAT | Control | 14.92023262 | 0.276298236 | 13.36543175 | 0.416287305 | 2.937931743 | 1.554800878 | 1.270839557 | 3.91E-05 | 4.408110738 | Unpaired t-test | 0.005386177 | permutation FDR (250 perm) | 1 | 8.46E-05 | FDR correction BH |
| Dmel\CG7910~Q9VHW0 | VMAT~Control | VMAT | Control | 14.85029611 | 1.065393241 | 13.2978143 | 0.62613177 | 2.933212949 | 1.552481813 | 0.999892419 | 0.018501371 | 1.732796085 | Unpaired t-test | 0.013615819 | permutation FDR (250 perm) | 1 | 0.024550297 | FDR correction BH |
| ATP8A~Q8T0I4 | VMAT~Control | VMAT | Control | 16.7242787 | 0.293245781 | 15.1756385 | 0.519400935 | 2.925412776 | 1.548640203 | 1.222532932 | 0.000171635 | 3.765395263 | Unpaired t-test | 0.00648023 | permutation FDR (250 perm) | 1 | 0.000319642 | FDR correction BH |
| ergic53~Q9V3A8 | VMAT~Control | VMAT | Control | 15.61369374 | 0.476513689 | 14.06706249 | 0.297325081 | 2.921341984 | 1.546631256 | 1.236133945 | 0.000107281 | 3.969479107 | Unpaired t-test | 0.006199711 | permutation FDR (250 perm) | 1 | 0.000209575 | FDR correction BH |
| Dmel\CG8034~Q7K0I7 | VMAT~Control | VMAT | Control | 15.2994108 | 0.296879135 | 13.75468775 | 0.554847394 | 2.917480579 | 1.544723052 | 1.205475295 | 0.000265867 | 3.575335503 | Unpaired t-test | 0.006838214 | permutation FDR (250 perm) | 1 | 0.000481054 | FDR correction BH |
| Rph~Q9W2R9 | VMAT~Control | VMAT | Control | 15.06399613 | 0.140080273 | 13.52675745 | 0.349413217 | 2.902384557 | 1.537238685 | 1.315732422 | 3.63E-06 | 5.439753705 | Unpaired t-test | 0.004447891 | permutation FDR (250 perm) | 1 | 1.05E-05 | FDR correction BH |
| bbc~A0A0B4KFL2 | VMAT~Control | VMAT | Control | 15.68573799 | 0.160529108 | 14.15394377 | 0.611491833 | 2.891452131 | 1.531794218 | 1.19416381 | 0.000293921 | 3.531769118 | Unpaired t-test | 0.007096403 | permutation FDR (250 perm) | 1 | 0.000525849 | FDR correction BH |
| Dmel\CG11200~Q7K0F7 | VMAT~Control | VMAT | Control | 15.63466234 | 0.138975225 | 14.10928745 | 0.830697452 | 2.878615085 | 1.525374891 | 1.108024032 | 0.002324641 | 2.633644097 | Unpaired t-test | 0.009727057 | permutation FDR (250 perm) | 1 | 0.003503332 | FDR correction BH |
| Hk~Q9W2W9 | VMAT~Control | VMAT | Control | 20.93030992 | 0.094664564 | 19.40647562 | 0.160996394 | 2.875542782 | 1.523834302 | 1.406368873 | 5.26E-09 | 8.279099824 | Unpaired t-test | 0.002976608 | permutation FDR (250 perm) | 1 | 4.53E-08 | FDR correction BH |
| Cadps~Q9NHE5-2 | VMAT~Control | VMAT | Control | 19.75223324 | 0.070198604 | 18.2453859 | 0.156878875 | 2.841883349 | 1.506847337 | 1.399294761 | 2.61E-09 | 8.58361197 | Unpaired t-test | 0.003074525 | permutation FDR (250 perm) | 1 | 2.64E-08 | FDR correction BH |
| Fife~M9PE32 | VMAT~Control | VMAT | Control | 16.51488729 | 0.182727476 | 15.00892908 | 0.178064299 | 2.840132457 | 1.505958215 | 1.351723937 | 1.19E-07 | 6.925245622 | Unpaired t-test | 0.003796017 | permutation FDR (250 perm) | 1 | 5.58E-07 | FDR correction BH |
| Dmel\CG5853~Q9VL61 | VMAT~Control | VMAT | Control | 15.35198597 | 0.332437185 | 13.84926118 | 0.542963063 | 2.833774157 | 1.502724785 | 1.16969179 | 0.000358707 | 3.445260166 | Unpaired t-test | 0.007715377 | permutation FDR (250 perm) | 1 | 0.000624192 | FDR correction BH |
| g~P54362 | VMAT~Control | VMAT | Control | 18.69756689 | 0.16892346 | 17.19512548 | 0.686934192 | 2.833217594 | 1.502441407 | 1.141361816 | 0.00078121 | 3.107232263 | Unpaired t-test | 0.008673343 | permutation FDR (250 perm) | 1 | 0.001270883 | FDR correction BH |
| Past1~Q8T6I0 | VMAT~Control | VMAT | Control | 21.10108886 | 0.080654125 | 19.60052039 | 0.124505187 | 2.829541826 | 1.500568463 | 1.407210608 | 6.43E-10 | 9.19207689 | Unpaired t-test | 0.002964408 | permutation FDR (250 perm) | 1 | 8.95E-09 | FDR correction BH |
| Nepl16~Q9VBX6 | VMAT~Control | VMAT | Control | 15.07181308 | 0.083031851 | 13.57271759 | 0.206772456 | 2.826654381 | 1.499095493 | 1.363249682 | 3.40E-08 | 7.468571114 | Unpaired t-test | 0.003646087 | permutation FDR (250 perm) | 1 | 1.99E-07 | FDR correction BH |
| Hdc~A0A0B4KEQ1 | VMAT~Control | VMAT | Control | 15.16548525 | 0.318933509 | 13.67125423 | 0.380390015 | 2.817139528 | 1.49423102 | 1.222777162 | 5.16E-05 | 4.287081206 | Unpaired t-test | 0.006478303 | permutation FDR (250 perm) | 1 | 0.000108713 | FDR correction BH |
| comt~M9PH10 | VMAT~Control | VMAT | Control | 18.17106716 | 0.053943143 | 16.68043492 | 0.133589431 | 2.810120973 | 1.490632238 | 1.400404453 | 5.15E-10 | 9.288584919 | Unpaired t-test | 0.00305699 | permutation FDR (250 perm) | 1 | 7.34E-09 | FDR correction BH |
| MESK2~Q9I7V7 | VMAT~Control | VMAT | Control | 17.80970503 | 0.124202595 | 16.3357826 | 0.213390931 | 2.777760909 | 1.473922427 | 1.327356783 | 1.07E-07 | 6.971927393 | Unpaired t-test | 0.004227074 | permutation FDR (250 perm) | 1 | 5.06E-07 | FDR correction BH |
| svr~X2J9Z1 | VMAT~Control | VMAT | Control | 15.24459216 | 0.142293833 | 13.77645777 | 0.40159891 | 2.766638948 | 1.468134382 | 1.233165825 | 1.63E-05 | 4.786830648 | Unpaired t-test | 0.006234443 | permutation FDR (250 perm) | 1 | 3.87E-05 | FDR correction BH |
| Lrpprc2~Q9W592 | VMAT~Control | VMAT | Control | 15.09054834 | 0.336065512 | 13.6280729 | 0.561659966 | 2.755808118 | 1.462475439 | 1.13132337 | 0.000540278 | 3.267382376 | Unpaired t-test | 0.00898142 | permutation FDR (250 perm) | 1 | 0.000903278 | FDR correction BH |
| Dmel\CG6230~Q9VKJ6 | VMAT~Control | VMAT | Control | 15.23402976 | 0.396397169 | 13.78322968 | 0.338978632 | 2.733596079 | 1.450800084 | 1.176400028 | 9.89E-05 | 4.004973786 | Unpaired t-test | 0.007538977 | permutation FDR (250 perm) | 1 | 0.000194744 | FDR correction BH |
| red~Q9I7J4 | VMAT~Control | VMAT | Control | 16.25029946 | 0.174251497 | 14.81895724 | 0.231560011 | 2.69697512 | 1.431342212 | 1.26712062 | 6.35E-07 | 6.197134055 | Unpaired t-test | 0.005484775 | permutation FDR (250 perm) | 1 | 2.28E-06 | FDR correction BH |
| nmd~X2J7P4 | VMAT~Control | VMAT | Control | 15.66034403 | 0.181908579 | 14.23039777 | 0.9871176 | 2.694366792 | 1.429946262 | 0.986928269 | 0.009727411 | 2.012002734 | Unpaired t-test | 0.014190028 | permutation FDR (250 perm) | 1 | 0.013461685 | FDR correction BH |
| lace~Q9V3F2 | VMAT~Control | VMAT | Control | 15.83638002 | 0.268171414 | 14.41191829 | 0.164928585 | 2.684143363 | 1.42446173 | 1.248656138 | 1.43E-06 | 5.845156668 | Unpaired t-test | 0.005934363 | permutation FDR (250 perm) | 1 | 4.61E-06 | FDR correction BH |
| ND-20~Q9VXK7 | VMAT~Control | VMAT | Control | 18.01578514 | 0.279139601 | 16.59168433 | 0.634396233 | 2.683471951 | 1.424100808 | 1.08713265 | 0.000988473 | 3.005035335 | Unpaired t-test | 0.010420503 | permutation FDR (250 perm) | 1 | 0.001581825 | FDR correction BH |
| Dmel\CG10089~Q9VU80 | VMAT~Control | VMAT | Control | 15.52969907 | 0.590381497 | 14.10742929 | 0.805729987 | 2.68006834 | 1.422269789 | 0.983106046 | 0.00975507 | 2.010769608 | Unpaired t-test | 0.014337248 | permutation FDR (250 perm) | 1 | 0.01349203 | FDR correction BH |
| Ocrl~O46094 | VMAT~Control | VMAT | Control | 17.3268264 | 0.055646846 | 15.90487032 | 0.163356545 | 2.679485637 | 1.421956083 | 1.320075765 | 4.78E-09 | 8.320647467 | Unpaired t-test | 0.004358701 | permutation FDR (250 perm) | 1 | 4.21E-08 | FDR correction BH |
| Timp~Q9VH14 | VMAT~Control | VMAT | Control | 15.06237388 | 0.110701181 | 13.64316968 | 0.344415984 | 2.674379496 | 1.419204199 | 1.221568775 | 5.21E-06 | 5.283262201 | Unpaired t-test | 0.00648216 | permutation FDR (250 perm) | 1 | 1.43E-05 | FDR correction BH |
| GstO2~Q9VSL4 | VMAT~Control | VMAT | Control | 15.25882491 | 0.51648059 | 13.84396885 | 0.530814353 | 2.66633132 | 1.414856062 | 1.062830812 | 0.001632569 | 2.787128551 | Unpaired t-test | 0.011329863 | permutation FDR (250 perm) | 1 | 0.002516743 | FDR correction BH |
| Nt5c~Q7K3S2 | VMAT~Control | VMAT | Control | 16.48892823 | 0.094382898 | 15.07971874 | 0.059123182 | 2.655915946 | 1.409209489 | 1.342351023 | 7.07E-11 | 10.1504651 | Unpaired t-test | 0.003935922 | permutation FDR (250 perm) | 1 | 1.62E-09 | FDR correction BH |
| Tdc2~A1Z6N4 | VMAT~Control | VMAT | Control | 18.67391907 | 0.141089726 | 17.26751991 | 0.166818352 | 2.650747317 | 1.406399151 | 1.281213782 | 5.19E-08 | 7.284520976 | Unpaired t-test | 0.005136781 | permutation FDR (250 perm) | 1 | 2.80E-07 | FDR correction BH |
| Gpat4~A0A0B4LHH1 | VMAT~Control | VMAT | Control | 15.42157066 | 0.228899219 | 14.01699221 | 0.089779034 | 2.647404141 | 1.404578447 | 1.265432502 | 1.62E-07 | 6.790487264 | Unpaired t-test | 0.005523441 | permutation FDR (250 perm) | 1 | 7.26E-07 | FDR correction BH |
| Mhc~M9NEP1 | VMAT~Control | VMAT | Control | 15.30215333 | 0.031387994 | 13.89977511 | 0.95136991 | 2.643369708 | 1.402378218 | 0.983643905 | 0.008090886 | 2.092003922 | Unpaired t-test | 0.014304511 | permutation FDR (250 perm) | 1 | 0.011411591 | FDR correction BH |
| RpS7~Q8IMI7 | VMAT~Control | VMAT | Control | 15.85160826 | 0.287460307 | 14.45326445 | 0.853503713 | 2.635988019 | 1.398343813 | 0.996847566 | 0.006002469 | 2.221670058 | Unpaired t-test | 0.01373258 | permutation FDR (250 perm) | 1 | 0.008584333 | FDR correction BH |
| Osbp~Q9VC05 | VMAT~Control | VMAT | Control | 17.67159074 | 0.200282134 | 16.27936922 | 0.225985053 | 2.624825494 | 1.392221511 | 1.226581118 | 1.20E-06 | 5.920365213 | Unpaired t-test | 0.006385728 | permutation FDR (250 perm) | 1 | 3.95E-06 | FDR correction BH |
| brp~A0A0B4K6V0 | VMAT~Control | VMAT | Control | 18.67649989 | 0.159041126 | 17.28567215 | 0.12122721 | 2.622290896 | 1.390827735 | 1.276654403 | 2.47E-08 | 7.607383535 | Unpaired t-test | 0.005243113 | permutation FDR (250 perm) | 1 | 1.53E-07 | FDR correction BH |
| AP-2alpha~P91926 | VMAT~Control | VMAT | Control | 19.40177305 | 0.033433592 | 18.01145886 | 0.053785094 | 2.621357635 | 1.390314196 | 1.352022394 | 2.97E-13 | 12.52705301 | Unpaired t-test | 0.003790189 | permutation FDR (250 perm) | 1 | 5.00E-11 | FDR correction BH |
| Hmgs~Q7K4Q9 | VMAT~Control | VMAT | Control | 16.15775512 | 0.31800043 | 14.77060853 | 0.446836058 | 2.615608452 | 1.38714659 | 1.113932439 | 0.000210819 | 3.676089354 | Unpaired t-test | 0.009529971 | permutation FDR (250 perm) | 1 | 0.000386502 | FDR correction BH |
| Spt-I~Q6NR46 | VMAT~Control | VMAT | Control | 16.08767473 | 0.391494897 | 14.7058338 | 0.134814114 | 2.606006941 | 1.381840927 | 1.165941385 | 2.15E-05 | 4.666794212 | Unpaired t-test | 0.007810888 | permutation FDR (250 perm) | 1 | 4.95E-05 | FDR correction BH |
| Sccpdh1~Q9VN86 | VMAT~Control | VMAT | Control | 15.80243469 | 0.241067684 | 14.4312352 | 1.106617152 | 2.586855542 | 1.371199492 | 0.910188378 | 0.02203924 | 1.656803391 | Unpaired t-test | 0.017701493 | permutation FDR (250 perm) | 1 | 0.028870546 | FDR correction BH |
| CG30023~Q86PD3 | VMAT~Control | VMAT | Control | 15.41009971 | 0.040604114 | 14.04027482 | 0.66129673 | 2.584391947 | 1.369824885 | 1.056720793 | 0.000946043 | 3.024088948 | Unpaired t-test | 0.011553453 | permutation FDR (250 perm) | 1 | 0.001515988 | FDR correction BH |
| alphaTub67C~P06606 | VMAT~Control | VMAT | Control | 16.07639167 | 0.23108527 | 14.70692198 | 0.507280902 | 2.583755743 | 1.369469691 | 1.096196001 | 0.000264216 | 3.578040241 | Unpaired t-test | 0.010190974 | permutation FDR (250 perm) | 1 | 0.000478803 | FDR correction BH |
| wdp~Q9W266 | VMAT~Control | VMAT | Control | 17.41307541 | 0.161065059 | 16.05092228 | 0.223963091 | 2.570685524 | 1.362153134 | 1.212559094 | 6.37E-07 | 6.196048938 | Unpaired t-test | 0.006669234 | permutation FDR (250 perm) | 1 | 2.28E-06 | FDR correction BH |
| Trhn~Q9W0K2 | VMAT~Control | VMAT | Control | 18.07649497 | 0.068808623 | 16.7226446 | 0.197925526 | 2.555933618 | 1.353850368 | 1.237849726 | 5.01E-08 | 7.299862805 | Unpaired t-test | 0.006168837 | permutation FDR (250 perm) | 1 | 2.72E-07 | FDR correction BH |
| mew~X2JDJ5 | VMAT~Control | VMAT | Control | 16.82774236 | 0.136565874 | 15.47442969 | 0.149965651 | 2.554981203 | 1.353312677 | 1.240765049 | 3.68E-08 | 7.433911261 | Unpaired t-test | 0.006107091 | permutation FDR (250 perm) | 1 | 2.12E-07 | FDR correction BH |
| l(2)k05819~Q7KU45 | VMAT~Control | VMAT | Control | 15.08553685 | 0.34802924 | 13.73321843 | 0.360557592 | 2.553221018 | 1.352318429 | 1.104736169 | 0.000126233 | 3.89882695 | Unpaired t-test | 0.009856532 | permutation FDR (250 perm) | 1 | 0.000242178 | FDR correction BH |
| Mctp~A1ZBD6-3 | VMAT~Control | VMAT | Control | 14.68302793 | 0.197579695 | 13.33658311 | 0.42707174 | 2.542847297 | 1.346444828 | 1.112358499 | 7.85E-05 | 4.105094417 | Unpaired t-test | 0.009607992 | permutation FDR (250 perm) | 1 | 0.000157683 | FDR correction BH |
| ArfGAP1~Q9VTX5 | VMAT~Control | VMAT | Control | 19.95043983 | 0.039499379 | 18.60470704 | 0.086118962 | 2.5415926 | 1.345732795 | 1.291030048 | 2.25E-11 | 10.64739692 | Unpaired t-test | 0.004944364 | permutation FDR (250 perm) | 1 | 6.98E-10 | FDR correction BH |
| Sfxn1-3~Q9VN13 | VMAT~Control | VMAT | Control | 15.71437387 | 0.151406711 | 14.37794179 | 0.225185752 | 2.525260246 | 1.336432075 | 1.191803304 | 6.52E-07 | 6.185650541 | Unpaired t-test | 0.007133269 | permutation FDR (250 perm) | 1 | 2.32E-06 | FDR correction BH |
| mys~X2JE30 | VMAT~Control | VMAT | Control | 18.57376954 | 0.050843432 | 17.23914688 | 0.112706319 | 2.522095072 | 1.33462266 | 1.26469136 | 3.39E-10 | 9.469495599 | Unpaired t-test | 0.005542774 | permutation FDR (250 perm) | 1 | 5.55E-09 | FDR correction BH |
| Ih~A1Z9N8 | VMAT~Control | VMAT | Control | 19.66614222 | 0.179232433 | 18.33912326 | 0.411128931 | 2.50883737 | 1.327018954 | 1.105319675 | 5.96E-05 | 4.224984085 | Unpaired t-test | 0.009826128 | permutation FDR (250 perm) | 1 | 0.000123437 | FDR correction BH |
| NTPase~O76268 | VMAT~Control | VMAT | Control | 16.33155814 | 0.120433509 | 15.0083417 | 0.173409242 | 2.502233531 | 1.323216441 | 1.209058059 | 6.71E-08 | 7.173485739 | Unpaired t-test | 0.006766715 | permutation FDR (250 perm) | 1 | 3.40E-07 | FDR correction BH |
| Rgk1~A1ZBJ6 | VMAT~Control | VMAT | Control | 15.54522929 | 0.529639259 | 14.22643458 | 0.466724588 | 2.494576158 | 1.318794715 | 1.002348008 | 0.001895845 | 2.722197276 | Unpaired t-test | 0.013512953 | permutation FDR (250 perm) | 1 | 0.002886687 | FDR correction BH |
| Wdfy2~Q9VKW1 | VMAT~Control | VMAT | Control | 17.87007972 | 0.132764684 | 16.55717042 | 0.516767396 | 2.484420375 | 1.312909305 | 1.059985555 | 0.000260942 | 3.583456666 | Unpaired t-test | 0.011445601 | permutation FDR (250 perm) | 1 | 0.000473233 | FDR correction BH |
| Sac1~M9MRM4 | VMAT~Control | VMAT | Control | 16.90534587 | 0.106483779 | 15.59333362 | 0.241528012 | 2.482876076 | 1.312012256 | 1.173486576 | 5.99E-07 | 6.222775637 | Unpaired t-test | 0.007608031 | permutation FDR (250 perm) | 1 | 2.16E-06 | FDR correction BH |
| dare~Q9V3T9 | VMAT~Control | VMAT | Control | 14.91278226 | 0.20707695 | 13.60220425 | 0.231996759 | 2.480408977 | 1.310578016 | 1.150567708 | 2.73E-06 | 5.563698073 | Unpaired t-test | 0.008366412 | permutation FDR (250 perm) | 1 | 8.11E-06 | FDR correction BH |
| Dg~A1ZA89 | VMAT~Control | VMAT | Control | 16.80042812 | 0.163002916 | 15.49131344 | 0.212936641 | 2.477894344 | 1.309114673 | 1.168928979 | 7.08E-07 | 6.15000603 | Unpaired t-test | 0.0077383 | permutation FDR (250 perm) | 1 | 2.49E-06 | FDR correction BH |
| hrm~Q7JWI7 | VMAT~Control | VMAT | Control | 16.18868075 | 0.128670139 | 14.88546369 | 0.28845563 | 2.467785607 | 1.303217063 | 1.141917326 | 3.31E-06 | 5.480324604 | Unpaired t-test | 0.008661898 | permutation FDR (250 perm) | 1 | 9.63E-06 | FDR correction BH |
| Dmel\CG1344~A1Z6G6 | VMAT~Control | VMAT | Control | 16.4542956 | 0.191271853 | 15.1513388 | 0.401901988 | 2.467340448 | 1.302956795 | 1.086655043 | 6.51E-05 | 4.186591313 | Unpaired t-test | 0.010437589 | permutation FDR (250 perm) | 1 | 0.00013332 | FDR correction BH |
| POLDIP2~Q9VNC0 | VMAT~Control | VMAT | Control | 15.15477879 | 0.155490353 | 13.85406141 | 0.346622963 | 2.463513498 | 1.300717376 | 1.111822257 | 1.73E-05 | 4.76273181 | Unpaired t-test | 0.009627022 | permutation FDR (250 perm) | 1 | 4.05E-05 | FDR correction BH |
| DmTOM1~M9PF36 | VMAT~Control | VMAT | Control | 21.99592715 | 0.038067926 | 20.69936375 | 0.058362787 | 2.45643048 | 1.29656341 | 1.257380786 | 1.54E-12 | 11.81203563 | Unpaired t-test | 0.005718976 | permutation FDR (250 perm) | 1 | 1.23E-10 | FDR correction BH |
| path~Q9VT04 | VMAT~Control | VMAT | Control | 18.75367294 | 0.088385509 | 17.46197154 | 0.1266866 | 2.448166029 | 1.291701402 | 1.208234366 | 4.14E-09 | 8.383026781 | Unpaired t-test | 0.006780769 | permutation FDR (250 perm) | 1 | 3.76E-08 | FDR correction BH |
| Pu~P48596-2 | VMAT~Control | VMAT | Control | 15.96029858 | 0.408213396 | 14.68053571 | 1.419676782 | 2.427990661 | 1.279762873 | 0.770651799 | 0.081457384 | 1.089069543 | Unpaired t-test | 0.030336707 | permutation FDR (250 perm) | 1 | 0.098586468 | FDR correction BH |
| Rme-8~A0A0B4LFW6 | VMAT~Control | VMAT | Control | 17.21247837 | 0.087979909 | 15.93370992 | 0.140891243 | 2.426317668 | 1.278768449 | 1.190344507 | 9.25E-09 | 8.03392137 | Unpaired t-test | 0.007190796 | permutation FDR (250 perm) | 1 | 7.07E-08 | FDR correction BH |
| Ppox~Q9VC52 | VMAT~Control | VMAT | Control | 14.77479486 | 1.038086814 | 13.49881199 | 0.245357186 | 2.421637403 | 1.275982864 | 0.863879741 | 0.023299551 | 1.632652449 | Unpaired t-test | 0.020532221 | permutation FDR (250 perm) | 1 | 0.030386229 | FDR correction BH |
| RhoL~D3DMW7 | VMAT~Control | VMAT | Control | 15.57116701 | 0.325661671 | 14.29705353 | 0.632596504 | 2.418501595 | 1.274113489 | 0.96656067 | 0.002501267 | 2.601839947 | Unpaired t-test | 0.015084428 | permutation FDR (250 perm) | 1 | 0.003752698 | FDR correction BH |
| Pka-R2~P81900 | VMAT~Control | VMAT | Control | 19.4521105 | 0.056613951 | 18.18134418 | 0.211559216 | 2.412896979 | 1.27076632 | 1.157408305 | 1.40E-07 | 6.854801406 | Unpaired t-test | 0.008114504 | permutation FDR (250 perm) | 1 | 6.39E-07 | FDR correction BH |
| muc~Q7KTK9 | VMAT~Control | VMAT | Control | 15.13048013 | 0.547209155 | 13.86190376 | 0.469741304 | 2.409237078 | 1.268576367 | 0.959211705 | 0.002805041 | 2.55206084 | Unpaired t-test | 0.015428304 | permutation FDR (250 perm) | 1 | 0.004168602 | FDR correction BH |
| pch2~Q8SX76 | VMAT~Control | VMAT | Control | 15.17228048 | 0.358058785 | 13.9113346 | 0.461775736 | 2.396528142 | 1.260945881 | 0.99970272 | 0.000696691 | 3.156959616 | Unpaired t-test | 0.013623352 | permutation FDR (250 perm) | 1 | 0.001140481 | FDR correction BH |
| Dmel\CG2004~Q9W3C3 | VMAT~Control | VMAT | Control | 15.59514438 | 0.324125502 | 14.33621557 | 0.983573261 | 2.393179831 | 1.25892881 | 0.860431982 | 0.021624388 | 1.665056178 | Unpaired t-test | 0.020754752 | permutation FDR (250 perm) | 1 | 0.028374475 | FDR correction BH |
| prt~Q9VCJ5 | VMAT~Control | VMAT | Control | 15.03979768 | 0.320919958 | 13.78295802 | 0.301586561 | 2.389716794 | 1.256839654 | 1.050036304 | 8.02E-05 | 4.095832315 | Unpaired t-test | 0.011780615 | permutation FDR (250 perm) | 1 | 0.000160807 | FDR correction BH |
| ReepB~A1Z9M2 | VMAT~Control | VMAT | Control | 16.47889558 | 0.344797736 | 15.22404075 | 0.198477378 | 2.386431341 | 1.25485483 | 1.065313571 | 3.49E-05 | 4.457604362 | Unpaired t-test | 0.011237915 | permutation FDR (250 perm) | 1 | 7.67E-05 | FDR correction BH |
| unc-13~Q8IM86 | VMAT~Control | VMAT | Control | 19.00201033 | 0.125810225 | 17.75015647 | 0.275909311 | 2.381472458 | 1.251853864 | 1.102359767 | 3.29E-06 | 5.482449597 | Unpaired t-test | 0.009951544 | permutation FDR (250 perm) | 1 | 9.59E-06 | FDR correction BH |
| Cyp9b2~Q9V4I1 | VMAT~Control | VMAT | Control | 16.62237357 | 0.178916158 | 15.37231462 | 0.662247473 | 2.378511417 | 1.25005895 | 0.956591729 | 0.002233204 | 2.651071604 | Unpaired t-test | 0.015537453 | permutation FDR (250 perm) | 1 | 0.003376341 | FDR correction BH |
| drpr~Q9W0A0-3 | VMAT~Control | VMAT | Control | 15.85002767 | 0.434410307 | 14.60651023 | 0.137791518 | 2.36775112 | 1.243517444 | 1.032982184 | 0.000115501 | 3.937415081 | Unpaired t-test | 0.012410014 | permutation FDR (250 perm) | 1 | 0.000223043 | FDR correction BH |
| Gbeta13F~A4V4I0 | VMAT~Control | VMAT | Control | 16.03711695 | 1.252180474 | 14.80054116 | 1.21227023 | 2.356385842 | 1.23657579 | 0.694928062 | 0.143706393 | 0.842523912 | Unpaired t-test | 0.043810478 | permutation FDR (250 perm) | 1 | 0.168620375 | FDR correction BH |
| BcDNA:RE18604~Q7JRI6 | VMAT~Control | VMAT | Control | 18.56388679 | 0.048380438 | 17.33550204 | 0.207118592 | 2.343045138 | 1.228384747 | 1.121689947 | 1.46E-07 | 6.835492578 | Unpaired t-test | 0.009275583 | permutation FDR (250 perm) | 1 | 6.66E-07 | FDR correction BH |
| Dmel\CG6638~Q9VSL9 | VMAT~Control | VMAT | Control | 18.73026617 | 0.284626617 | 17.51159069 | 0.292734686 | 2.327329503 | 1.218675482 | 1.030508994 | 5.54E-05 | 4.256509544 | Unpaired t-test | 0.012476145 | permutation FDR (250 perm) | 1 | 0.000115506 | FDR correction BH |
| VhaAC39-1~Q9W4P5 | VMAT~Control | VMAT | Control | 22.92833248 | 0.715102259 | 21.72561952 | 0.587557773 | 2.301720973 | 1.202712953 | 0.850630827 | 0.01567827 | 1.804701861 | Unpaired t-test | 0.021513664 | permutation FDR (250 perm) | 1 | 0.020969686 | FDR correction BH |
| retm~B5RIN0 | VMAT~Control | VMAT | Control | 14.65921607 | 0.091565465 | 13.46060008 | 0.298731662 | 2.295193827 | 1.198615993 | 1.051664982 | 6.36E-06 | 5.196592227 | Unpaired t-test | 0.011718203 | permutation FDR (250 perm) | 1 | 1.71E-05 | FDR correction BH |
| Trs31~Q7K2Q8 | VMAT~Control | VMAT | Control | 18.18213156 | 0.072732504 | 16.984303 | 0.293259185 | 2.293941439 | 1.197828562 | 1.055241314 | 4.74E-06 | 5.324161516 | Unpaired t-test | 0.01158562 | permutation FDR (250 perm) | 1 | 1.32E-05 | FDR correction BH |
| lt~Q7PL76 | VMAT~Control | VMAT | Control | 17.15622864 | 0.129777201 | 15.95848263 | 0.14139973 | 2.293810169 | 1.197746002 | 1.103067104 | 6.99E-08 | 7.155746527 | Unpaired t-test | 0.009930641 | permutation FDR (250 perm) | 1 | 3.50E-07 | FDR correction BH |
| if~P12080-2 | VMAT~Control | VMAT | Control | 14.48963516 | 0.203659488 | 13.29635015 | 0.491767706 | 2.286728387 | 1.193285015 | 0.963850962 | 0.000527156 | 3.27806125 | Unpaired t-test | 0.015210502 | permutation FDR (250 perm) | 1 | 0.00088511 | FDR correction BH |
| exu~E1JGN9 | VMAT~Control | VMAT | Control | 16.05268357 | 0.488569521 | 14.85946915 | 0.131301507 | 2.286616481 | 1.193214412 | 0.973061435 | 0.000360772 | 3.442767007 | Unpaired t-test | 0.01479061 | permutation FDR (250 perm) | 1 | 0.000627221 | FDR correction BH |
| Nup205~Q8IQV9-2 | VMAT~Control | VMAT | Control | 16.08868437 | 2.684474534 | 14.89569068 | 2.042062857 | 2.286266673 | 1.192993691 | 0.475598564 | 0.447355262 | 0.34934745 | Unpaired t-test | 0.137878896 | permutation FDR (250 perm) | 1 | 0.48528769 | FDR correction BH |
| RapGAP1~Q9VM02 | VMAT~Control | VMAT | Control | 15.70893145 | 0.219926056 | 14.51810599 | 0.227304711 | 2.282833211 | 1.190825457 | 1.04326041 | 7.51E-06 | 5.124392963 | Unpaired t-test | 0.012037807 | permutation FDR (250 perm) | 1 | 1.95E-05 | FDR correction BH |
| Apl~M9PFW8 | VMAT~Control | VMAT | Control | 15.35477284 | 0.30367321 | 14.16525406 | 0.539308171 | 2.280766546 | 1.189518783 | 0.931646049 | 0.001567571 | 2.804772837 | Unpaired t-test | 0.016689397 | permutation FDR (250 perm) | 1 | 0.0024213 | FDR correction BH |
| Sp7~Q9V3Z2 | VMAT~Control | VMAT | Control | 19.12913996 | 0.367541375 | 17.93993394 | 0.39131156 | 2.280272145 | 1.189206017 | 0.958969003 | 0.000575792 | 3.239734517 | Unpaired t-test | 0.015437676 | permutation FDR (250 perm) | 1 | 0.00095789 | FDR correction BH |
| BG:DS04929.1~Q9V416 | VMAT~Control | VMAT | Control | 15.07291229 | 0.481001496 | 13.88522095 | 0.669044988 | 2.277879356 | 1.187691339 | 0.867874364 | 0.009128763 | 2.039588064 | Unpaired t-test | 0.020235512 | permutation FDR (250 perm) | 1 | 0.012707929 | FDR correction BH |
| mtd~A0A0B4K620 | VMAT~Control | VMAT | Control | 21.6205102 | 0.114521157 | 20.43835493 | 0.160191531 | 2.269155176 | 1.182155271 | 1.086475769 | 1.01E-07 | 6.995197071 | Unpaired t-test | 0.010452776 | permutation FDR (250 perm) | 1 | 4.85E-07 | FDR correction BH |
| Src64B~M9MSK3 | VMAT~Control | VMAT | Control | 14.98028173 | 0.128471837 | 13.80410203 | 0.358757819 | 2.259775895 | 1.176179706 | 1.004922417 | 4.19E-05 | 4.378239199 | Unpaired t-test | 0.013413095 | permutation FDR (250 perm) | 1 | 9.01E-05 | FDR correction BH |
| Dmel\CG8155~Q5BI03 | VMAT~Control | VMAT | Control | 14.84305434 | 0.110396757 | 13.66746751 | 0.479254673 | 2.258847427 | 1.175586828 | 0.963641548 | 0.000325923 | 3.486884763 | Unpaired t-test | 0.015212377 | permutation FDR (250 perm) | 1 | 0.000572555 | FDR correction BH |
| Nup154~Q9V463 | VMAT~Control | VMAT | Control | 16.95009042 | 0.128103492 | 15.77718356 | 0.103020704 | 2.254655271 | 1.172906867 | 1.09258338 | 1.93E-08 | 7.71437708 | Unpaired t-test | 0.010285986 | permutation FDR (250 perm) | 1 | 1.26E-07 | FDR correction BH |
| Rac2~M9PBU2 | VMAT~Control | VMAT | Control | 15.57320988 | 0.649309857 | 14.40126477 | 0.927568022 | 2.253152727 | 1.171945108 | 0.777999712 | 0.043177855 | 1.364738932 | Unpaired t-test | 0.029266452 | permutation FDR (250 perm) | 1 | 0.05432425 | FDR correction BH |
| Snap24~Q9VH76 | VMAT~Control | VMAT | Control | 19.36062632 | 0.064683399 | 18.18918146 | 0.106733382 | 2.252371597 | 1.171444863 | 1.109518319 | 1.34E-09 | 8.873381311 | Unpaired t-test | 0.00968331 | permutation FDR (250 perm) | 1 | 1.54E-08 | FDR correction BH |
| Akap200~Q9VLL3-2 | VMAT~Control | VMAT | Control | 18.43771792 | 0.181208877 | 17.26636545 | 0.24904641 | 2.25222735 | 1.171352467 | 1.029543692 | 6.87E-06 | 5.163192862 | Unpaired t-test | 0.012510156 | permutation FDR (250 perm) | 1 | 1.82E-05 | FDR correction BH |
| Scox~Q9VMX4 | VMAT~Control | VMAT | Control | 15.12573299 | 0.13506289 | 13.95456659 | 0.393439115 | 2.251936905 | 1.171166407 | 0.987467545 | 8.96E-05 | 4.04770934 | Unpaired t-test | 0.014161806 | permutation FDR (250 perm) | 1 | 0.000177983 | FDR correction BH |
| colt~Q9VQG4 | VMAT~Control | VMAT | Control | 15.71418206 | 0.211375725 | 14.54409248 | 0.183747175 | 2.250256683 | 1.170089577 | 1.039844924 | 2.96E-06 | 5.529262142 | Unpaired t-test | 0.012177694 | permutation FDR (250 perm) | 1 | 8.69E-06 | FDR correction BH |
| PPO1~Q7K2W6 | VMAT~Control | VMAT | Control | 15.71407736 | 0.27385432 | 14.54455405 | 0.102049143 | 2.249373624 | 1.169523315 | 1.034337209 | 4.36E-06 | 5.360707497 | Unpaired t-test | 0.012376181 | permutation FDR (250 perm) | 1 | 1.23E-05 | FDR correction BH |
| SMAP~A1Z7K6 | VMAT~Control | VMAT | Control | 17.25748485 | 0.151791269 | 16.09379191 | 0.215695674 | 2.240301567 | 1.163692946 | 1.040913493 | 1.80E-06 | 5.744958669 | Unpaired t-test | 0.012130435 | permutation FDR (250 perm) | 1 | 5.62E-06 | FDR correction BH |
| CD98hc~Q9VHX9 | VMAT~Control | VMAT | Control | 16.79743354 | 0.108687012 | 15.63942823 | 0.104803518 | 2.231486861 | 1.158005312 | 1.084759323 | 9.59E-09 | 8.018066277 | Unpaired t-test | 0.010511628 | permutation FDR (250 perm) | 1 | 7.28E-08 | FDR correction BH |
| Pka-C1~P12370 | VMAT~Control | VMAT | Control | 17.074072 | 0.045768999 | 15.91844678 | 0.263268937 | 2.227808494 | 1.155625221 | 1.032265916 | 2.16E-06 | 5.665619653 | Unpaired t-test | 0.012421351 | permutation FDR (250 perm) | 1 | 6.62E-06 | FDR correction BH |
| Dmel\CG6055~Q9VM07 | VMAT~Control | VMAT | Control | 16.66664246 | 0.228896589 | 15.52096375 | 0.378117884 | 2.212501923 | 1.145678709 | 0.956589782 | 0.000173921 | 3.759648072 | Unpaired t-test | 0.015537453 | permutation FDR (250 perm) | 1 | 0.000323644 | FDR correction BH |
| CkIalpha~E1JJI5 | VMAT~Control | VMAT | Control | 15.73913221 | 0.199574599 | 14.60009176 | 0.148267773 | 2.202344947 | 1.139040452 | 1.02506589 | 1.27E-06 | 5.89499862 | Unpaired t-test | 0.012660066 | permutation FDR (250 perm) | 1 | 4.15E-06 | FDR correction BH |
| Dmel\CG2145~X2JJD6 | VMAT~Control | VMAT | Control | 16.09456626 | 0.22898036 | 14.95646741 | 0.234932448 | 2.200908014 | 1.138098849 | 0.992486937 | 1.54E-05 | 4.81227671 | Unpaired t-test | 0.013893647 | permutation FDR (250 perm) | 1 | 3.67E-05 | FDR correction BH |
| Dhpr~Q9VSU6 | VMAT~Control | VMAT | Control | 14.43799274 | 0.258348149 | 13.30097431 | 0.397780594 | 2.199260394 | 1.137018431 | 0.938041619 | 0.000318906 | 3.496337834 | Unpaired t-test | 0.01639102 | permutation FDR (250 perm) | 1 | 0.000562746 | FDR correction BH |
| Snap29~Q9W1I8 | VMAT~Control | VMAT | Control | 18.64037849 | 0.0386229 | 17.50602236 | 0.109596465 | 2.195205704 | 1.134356136 | 1.078318597 | 9.11E-10 | 9.040423238 | Unpaired t-test | 0.010732448 | permutation FDR (250 perm) | 1 | 1.14E-08 | FDR correction BH |
| Mitofilin~A0A0B4KHL7 | VMAT~Control | VMAT | Control | 17.0232431 | 0.080750828 | 15.89293079 | 0.209457978 | 2.189061228 | 1.130312307 | 1.027190058 | 5.31E-07 | 6.275077366 | Unpaired t-test | 0.012567925 | permutation FDR (250 perm) | 1 | 1.96E-06 | FDR correction BH |
| Usp20-33~Q7JW61 | VMAT~Control | VMAT | Control | 16.68474114 | 0.157401715 | 15.55710101 | 0.578537879 | 2.185010371 | 1.127640127 | 0.889211557 | 0.001812819 | 2.741645472 | Unpaired t-test | 0.018849442 | permutation FDR (250 perm) | 1 | 0.002776432 | FDR correction BH |
| primo-1~P82890 | VMAT~Control | VMAT | Control | 15.25590994 | 0.259051218 | 14.12998615 | 0.600533802 | 2.18241246 | 1.125923786 | 0.871128492 | 0.003215054 | 2.492811756 | Unpaired t-test | 0.020077886 | permutation FDR (250 perm) | 1 | 0.004744976 | FDR correction BH |
| yl~P98163-2 | VMAT~Control | VMAT | Control | 14.71888117 | 0.325733131 | 13.59309494 | 0.384317839 | 2.182204378 | 1.125786226 | 0.918783549 | 0.000539863 | 3.267716198 | Unpaired t-test | 0.017297852 | permutation FDR (250 perm) | 1 | 0.000903225 | FDR correction BH |
| Dmc1~Q0E9B5 | VMAT~Control | VMAT | Control | 15.18403216 | 0.163019075 | 14.06305273 | 0.556260395 | 2.17494576 | 1.120979423 | 0.890210258 | 0.001503017 | 2.82303617 | Unpaired t-test | 0.018802974 | permutation FDR (250 perm) | 1 | 0.002326168 | FDR correction BH |
| Arp1~P45889 | VMAT~Control | VMAT | Control | 16.43409982 | 0.091689135 | 15.31436362 | 0.281900305 | 2.173072352 | 1.119736209 | 0.988668004 | 7.30E-06 | 5.136910262 | Unpaired t-test | 0.014099718 | permutation FDR (250 perm) | 1 | 1.92E-05 | FDR correction BH |
| Rab10~O15971 | VMAT~Control | VMAT | Control | 15.84966717 | 0.130388852 | 14.73098893 | 0.069926847 | 2.171479357 | 1.118678238 | 1.04925138 | 1.10E-08 | 7.957934082 | Unpaired t-test | 0.011797732 | permutation FDR (250 perm) | 1 | 7.96E-08 | FDR correction BH |
| Fas2~X2JCI0 | VMAT~Control | VMAT | Control | 17.64738365 | 0.149506571 | 16.53002134 | 0.147090091 | 2.169499586 | 1.117362311 | 1.02154617 | 3.13E-07 | 6.50462877 | Unpaired t-test | 0.012797737 | permutation FDR (250 perm) | 1 | 1.24E-06 | FDR correction BH |
| CG8306~Q960W6 | VMAT~Control | VMAT | Control | 16.08719506 | 0.332363231 | 14.97070047 | 0.257658785 | 2.168195116 | 1.116494591 | 0.939754156 | 0.00014381 | 3.84220982 | Unpaired t-test | 0.0163087 | permutation FDR (250 perm) | 1 | 0.000272568 | FDR correction BH |
| PCB~Q0E9E2 | VMAT~Control | VMAT | Control | 20.7707125 | 0.185908538 | 19.65808664 | 0.161466265 | 2.16238867 | 1.112625858 | 1.002256247 | 1.45E-06 | 5.839807567 | Unpaired t-test | 0.013516722 | permutation FDR (250 perm) | 1 | 4.66E-06 | FDR correction BH |
| mRpL2~Q9VTF8 | VMAT~Control | VMAT | Control | 16.10037026 | 0.204484577 | 14.989802 | 0.248491193 | 2.159306835 | 1.110568263 | 0.970846124 | 1.61E-05 | 4.792496268 | Unpaired t-test | 0.014890141 | permutation FDR (250 perm) | 1 | 3.83E-05 | FDR correction BH |
| Dmel\CG8768~Q8T0N5 | VMAT~Control | VMAT | Control | 16.98027429 | 0.115999706 | 15.87065169 | 0.17223122 | 2.157891918 | 1.109622607 | 1.015333663 | 3.04E-07 | 6.51693959 | Unpaired t-test | 0.013002355 | permutation FDR (250 perm) | 1 | 1.21E-06 | FDR correction BH |
| Desat1~Q7K4Y0 | VMAT~Control | VMAT | Control | 18.34244404 | 0.116400325 | 17.23495531 | 0.259165514 | 2.154702558 | 1.107488729 | 0.98263883 | 5.51E-06 | 5.258655505 | Unpaired t-test | 0.014354157 | permutation FDR (250 perm) | 1 | 1.51E-05 | FDR correction BH |
| EndoB~Q8SYD9 | VMAT~Control | VMAT | Control | 21.47596844 | 0.064258907 | 20.36889743 | 0.161451671 | 2.154078775 | 1.10707101 | 1.02724189 | 5.73E-08 | 7.241538244 | Unpaired t-test | 0.012566038 | permutation FDR (250 perm) | 1 | 3.05E-07 | FDR correction BH |
| mfrn~Q9VAY3 | VMAT~Control | VMAT | Control | 15.78197124 | 0.338231845 | 14.6774781 | 0.192943412 | 2.150233216 | 1.104493144 | 0.940680712 | 8.43E-05 | 4.074009219 | Unpaired t-test | 0.01626754 | permutation FDR (250 perm) | 1 | 0.000168235 | FDR correction BH |
| side-IV~Q9VFU7 | VMAT~Control | VMAT | Control | 14.59332982 | 0.070770894 | 13.48994121 | 0.463796794 | 2.14858762 | 1.103388612 | 0.912029307 | 0.000368687 | 3.433342016 | Unpaired t-test | 0.017608209 | permutation FDR (250 perm) | 1 | 0.000638624 | FDR correction BH |
| Rala~P48555 | VMAT~Control | VMAT | Control | 18.40029314 | 0.120283833 | 17.29794398 | 0.060587262 | 2.147040143 | 1.102349165 | 1.039725217 | 5.10E-09 | 8.292438297 | Unpaired t-test | 0.012183365 | permutation FDR (250 perm) | 1 | 4.45E-08 | FDR correction BH |
| ClpP~Q9VKY3 | VMAT~Control | VMAT | Control | 15.95824338 | 0.189194589 | 14.8592188 | 0.333886011 | 2.142098146 | 1.099024582 | 0.938035004 | 7.80E-05 | 4.108138676 | Unpaired t-test | 0.01639102 | permutation FDR (250 perm) | 1 | 0.000156849 | FDR correction BH |
| nrv2~A4V0B5 | VMAT~Control | VMAT | Control | 17.38893466 | 0.057644987 | 16.29303422 | 0.098204409 | 2.137464468 | 1.095900437 | 1.042795518 | 1.05E-09 | 8.979785146 | Unpaired t-test | 0.01205482 | permutation FDR (250 perm) | 1 | 1.26E-08 | FDR correction BH |
| CT19169~Q961R9 | VMAT~Control | VMAT | Control | 17.44932145 | 0.078020991 | 16.35475592 | 0.067493657 | 2.135487612 | 1.094565529 | 1.046293687 | 4.02E-10 | 9.395971696 | Unpaired t-test | 0.011905482 | permutation FDR (250 perm) | 1 | 6.26E-09 | FDR correction BH |
| dysc~M9PFJ0 | VMAT~Control | VMAT | Control | 14.84669033 | 0.324787173 | 13.75242262 | 0.241279131 | 2.135046826 | 1.094267711 | 0.926604811 | 0.000124007 | 3.906553639 | Unpaired t-test | 0.016905696 | permutation FDR (250 perm) | 1 | 0.000238102 | FDR correction BH |
| Dmel\CG13928~Q9W0C3 | VMAT~Control | VMAT | Control | 15.49911978 | 0.267566822 | 14.40713107 | 0.213716848 | 2.131676784 | 1.091988705 | 0.946965683 | 3.18E-05 | 4.497851245 | Unpaired t-test | 0.015949462 | permutation FDR (250 perm) | 1 | 7.04E-05 | FDR correction BH |
| Dmel\CG4199~O77266 | VMAT~Control | VMAT | Control | 17.36416591 | 0.121417853 | 16.27405684 | 0.186577213 | 2.128901305 | 1.090109069 | 0.991411705 | 6.88E-07 | 6.162608455 | Unpaired t-test | 0.013952963 | permutation FDR (250 perm) | 1 | 2.43E-06 | FDR correction BH |
| Abcd3~M9PF96 | VMAT~Control | VMAT | Control | 16.92817652 | 0.346866989 | 15.83869526 | 0.209802464 | 2.127975077 | 1.089481254 | 0.922279451 | 0.000130439 | 3.884591272 | Unpaired t-test | 0.01712605 | permutation FDR (250 perm) | 1 | 0.000249232 | FDR correction BH |
| ND-B17.2~Q9VQD7 | VMAT~Control | VMAT | Control | 17.36883949 | 0.33317652 | 16.28343428 | 0.511639399 | 2.121971396 | 1.085405209 | 0.852602194 | 0.002621575 | 2.581437711 | Unpaired t-test | 0.021339509 | permutation FDR (250 perm) | 1 | 0.003918214 | FDR correction BH |
| yata~Q9VAH7 | VMAT~Control | VMAT | Control | 17.80395348 | 0.1327614 | 16.71995593 | 0.177719471 | 2.119901961 | 1.083997546 | 0.98616356 | 7.02E-07 | 6.153957944 | Unpaired t-test | 0.014208647 | permutation FDR (250 perm) | 1 | 2.47E-06 | FDR correction BH |
| Dmel\CG1648~Q7K2P3 | VMAT~Control | VMAT | Control | 22.59180449 | 0.105482273 | 21.51002616 | 0.116769679 | 2.116643548 | 1.081778334 | 1.010655647 | 2.76E-08 | 7.558755847 | Unpaired t-test | 0.013173811 | permutation FDR (250 perm) | 1 | 1.68E-07 | FDR correction BH |
| ND-18~Q9VWI0 | VMAT~Control | VMAT | Control | 18.03555136 | 0.470758588 | 16.95420647 | 0.662131549 | 2.116007712 | 1.081344886 | 0.793166306 | 0.01389693 | 1.857081133 | Unpaired t-test | 0.027297372 | permutation FDR (250 perm) | 1 | 0.018757668 | FDR correction BH |
| BcDNA:HL07910~Q8MZI3 | VMAT~Control | VMAT | Control | 15.9934012 | 0.188906641 | 14.91333095 | 0.32169298 | 2.114139032 | 1.080070256 | 0.925639756 | 7.13E-05 | 4.147008109 | Unpaired t-test | 0.016963585 | permutation FDR (250 perm) | 1 | 0.000144408 | FDR correction BH |
| Rab1~O18332 | VMAT~Control | VMAT | Control | 18.76954499 | 0.047413191 | 17.69389738 | 0.130185485 | 2.107667978 | 1.075647616 | 1.012887376 | 8.52E-09 | 8.069355708 | Unpaired t-test | 0.013094677 | permutation FDR (250 perm) | 1 | 6.63E-08 | FDR correction BH |
| Dmel\CG5937~Q9W444 | VMAT~Control | VMAT | Control | 14.67699844 | 0.209852579 | 13.60293353 | 0.326312546 | 2.105357044 | 1.074064918 | 0.915263202 | 0.00010273 | 3.988301996 | Unpaired t-test | 0.017454036 | permutation FDR (250 perm) | 1 | 0.000202027 | FDR correction BH |
| Gp93~Q9VAY2 | VMAT~Control | VMAT | Control | 16.06872681 | 0.131408099 | 14.99849305 | 0.319501091 | 2.099773553 | 1.070233751 | 0.927011762 | 4.06E-05 | 4.391671224 | Unpaired t-test | 0.016883287 | permutation FDR (250 perm) | 1 | 8.75E-05 | FDR correction BH |
| Sfp24Bb~B4ZJ91 | VMAT~Control | VMAT | Control | 17.59480168 | 0.572488965 | 16.52642039 | 0.285645989 | 2.09707911 | 1.068381286 | 0.830697936 | 0.003884901 | 2.410620038 | Unpaired t-test | 0.023270708 | permutation FDR (250 perm) | 1 | 0.005673112 | FDR correction BH |
| kdn~Q9W401 | VMAT~Control | VMAT | Control | 18.15950229 | 0.388331593 | 17.09421523 | 0.48672552 | 2.092586205 | 1.065287056 | 0.833257254 | 0.003342697 | 2.475903017 | Unpaired t-test | 0.023048589 | permutation FDR (250 perm) | 1 | 0.004927181 | FDR correction BH |

| | | | | | | | | | | | | | | | | | |
|---|---|---|---|---|---|---|---|---|---|---|---|---|---|---|---|---|---|
| CadN~O15943-8 | VMAT~Control | VMAT | Control | 16.64198231 | 0.130825372 | 15.57679547 | 0.322423224 | 2.092440855 | 1.065186844 | 0.921753056 | 4.48E-05 | 4.348284368 | Unpaired t-test | 0.017163399 | permutation FDR (250 perm) | 1 | 9.56E-05 FDR correction BH |
| Mccc1~Q9V9T5 | VMAT~Control | VMAT | Control | 21.22347564 | 0.184658015 | 20.16252183 | 0.213100137 | 2.086310393 | 1.060953812 | 0.942146056 | 7.55E-06 | 5.121845237 | Unpaired t-test | 0.016203929 | permutation FDR (250 perm) | 1 | 1.96E-05 FDR correction BH |
| Cyp6d5~Q9VFP1 | VMAT~Control | VMAT | Control | 16.99600114 | 0.162815399 | 15.9362952 | 0.23582872 | 2.084506607 | 1.059705945 | 0.939322965 | 8.81E-06 | 5.055193286 | Unpaired t-test | 0.016314312 | permutation FDR (250 perm) | 1 | 2.26E-05 FDR correction BH |
| Dmel\CG8043~Q9VHN4 | VMAT~Control | VMAT | Control | 15.78207303 | 0.232174179 | 14.72339861 | 0.223428916 | 2.083016722 | 1.058674421 | 0.9253331 | 2.46E-05 | 4.608539036 | Unpaired t-test | 0.016978525 | permutation FDR (250 perm) | 1 | 5.60E-05 FDR correction BH |
| TyrRS-m~Q9W107 | VMAT~Control | VMAT | Control | 15.66917357 | 0.286045757 | 14.61051904 | 0.113206812 | 2.082988005 | 1.058654531 | 0.930621829 | 1.65E-05 | 4.781877852 | Unpaired t-test | 0.016713685 | permutation FDR (250 perm) | 1 | 3.90E-05 FDR correction BH |
| Dmel\CG9399~Q9VHB1 | VMAT~Control | VMAT | Control | 17.5360802 | 0.164153926 | 16.47897409 | 0.238474607 | 2.080753557 | 1.057106103 | 0.935928386 | 9.85E-06 | 5.006737177 | Unpaired t-test | 0.016491819 | permutation FDR (250 perm) | 1 | 2.47E-05 FDR correction BH |
| MSBP~Q9VXM4 | VMAT~Control | VMAT | Control | 17.12103733 | 0.2213385 | 16.06657189 | 0.198374272 | 2.076948503 | 1.054465446 | 0.930747386 | 1.27E-05 | 4.897025896 | Unpaired t-test | 0.01670808 | permutation FDR (250 perm) | 1 | 3.09E-05 FDR correction BH |
| Dmel\CG41099~Q7PLP4 | VMAT~Control | VMAT | Control | 16.70504864 | 0.092255291 | 15.65570972 | 0.072014388 | 2.069581286 | 1.049338914 | 0.997148662 | 2.10E-09 | 8.678685024 | Unpaired t-test | 0.013715631 | permutation FDR (250 perm) | 1 | 2.23E-08 FDR correction BH |
| Orp8~Q8IN56 | VMAT~Control | VMAT | Control | 15.85956293 | 0.251480734 | 14.81548774 | 0.210284494 | 2.062044104 | 1.04407519 | 0.910581286 | 3.21E-05 | 4.493383105 | Unpaired t-test | 0.017671642 | permutation FDR (250 perm) | 1 | 7.10E-05 FDR correction BH |
| Sfp33A3~E1JHG2 | VMAT~Control | VMAT | Control | 17.56212563 | 0.616179582 | 16.51823648 | 0.147562113 | 2.061778216 | 1.043889151 | 0.813406033 | 0.004218243 | 2.374868422 | Unpaired t-test | 0.025074931 | permutation FDR (250 perm) | 1 | 0.006137048 FDR correction BH |
| TyrRS~Q9VV60 | VMAT~Control | VMAT | Control | 16.3150225 | 0.288800233 | 15.27189747 | 0.405546681 | 2.060686488 | 1.04312503 | 0.853164639 | 0.000860998 | 3.06499808 | Unpaired t-test | 0.021291339 | permutation FDR (250 perm) | 1 | 0.001390115 FDR correction BH |
| RpL10Ab~Q9VTP4 | VMAT~Control | VMAT | Control | 16.98456451 | 0.068926875 | 15.94415314 | 0.137863888 | 2.056814053 | 1.040411372 | 0.97331955 | 3.29E-08 | 7.482341939 | Unpaired t-test | 0.014783099 | permutation FDR (250 perm) | 1 | 1.93E-07 FDR correction BH |
| Surf4~O18405 | VMAT~Control | VMAT | Control | 16.48257731 | 0.274391483 | 15.44590311 | 0.157596246 | 2.051492945 | 1.036674193 | 0.908159366 | 2.52E-05 | 4.597884056 | Unpaired t-test | 0.017809036 | permutation FDR (250 perm) | 1 | 5.72E-05 FDR correction BH |
| Mic26-27~Q9VEY5 | VMAT~Control | VMAT | Control | 16.92470946 | 0.138463977 | 15.88867255 | 0.304114431 | 2.050586942 | 1.036036913 | 0.901342523 | 4.03E-05 | 4.394712896 | Unpaired t-test | 0.01819637 | permutation FDR (250 perm) | 1 | 8.70E-05 FDR correction BH |
| garz~A1Z8W8 | VMAT~Control | VMAT | Control | 17.2881277 | 0.107555291 | 16.25236199 | 0.092818006 | 2.050201505 | 1.035765712 | 0.973889863 | 1.57E-08 | 7.804940112 | Unpaired t-test | 0.014768075 | permutation FDR (250 perm) | 1 | 1.06E-07 FDR correction BH |
| Arp2~P45888-2 | VMAT~Control | VMAT | Control | 15.14240672 | 0.098434824 | 14.10786783 | 0.824170545 | 2.048458826 | 1.034538895 | 0.754477127 | 0.019217994 | 1.716291958 | Unpaired t-test | 0.032662689 | permutation FDR (250 perm) | 1 | 0.025458164 FDR correction BH |
| zda~Q7K3D4 | VMAT~Control | VMAT | Control | 18.22633832 | 0.143139475 | 17.19243138 | 0.054040076 | 2.047561722 | 1.033906941 | 0.967693456 | 3.26E-08 | 7.486974836 | Unpaired t-test | 0.015031895 | permutation FDR (250 perm) | 1 | 1.92E-07 FDR correction BH |
| Hsdl2~Q9VB10 | VMAT~Control | VMAT | Control | 18.71370593 | 0.08996709 | 17.68221608 | 0.075326054 | 2.044134107 | 1.031489849 | 0.980061205 | 2.54E-09 | 8.594898383 | Unpaired t-test | 0.014448098 | permutation FDR (250 perm) | 1 | 2.60E-08 FDR correction BH |
| dsd~Q9VB20 | VMAT~Control | VMAT | Control | 14.86325158 | 0.124062684 | 13.83206 | 0.194464831 | 2.04371153 | 1.031191574 | 0.934763056 | 1.60E-06 | 5.797216321 | Unpaired t-test | 0.01654792 | permutation FDR (250 perm) | 1 | 5.07E-06 FDR correction BH |
| Fad2~Q9VTD4 | VMAT~Control | VMAT | Control | 15.70520029 | 0.256705176 | 14.67450492 | 0.445942292 | 2.043008738 | 1.030695375 | 0.837886118 | 0.001180726 | 2.927850808 | Unpaired t-test | 0.022585185 | permutation FDR (250 perm) | 1 | 0.001869152 FDR correction BH |
| Sara~Q7K9H6 | VMAT~Control | VMAT | Control | 14.80972835 | 0.276370474 | 13.78337057 | 0.317958265 | 2.036875476 | 1.026357784 | 0.863644695 | 0.000281777 | 3.550093746 | Unpaired t-test | 0.020560037 | permutation FDR (250 perm) | 1 | 0.000506339 FDR correction BH |
| Nup133~Q9VCW3 | VMAT~Control | VMAT | Control | 16.85519553 | 0.137823817 | 15.82969027 | 0.450349384 | 2.035672195 | 1.025505262 | 0.847088954 | 0.000652665 | 3.185309967 | Unpaired t-test | 0.021780454 | permutation FDR (250 perm) | 1 | 0.001076645 FDR correction BH |
| QIL1~Q9VVH3 | VMAT~Control | VMAT | Control | 16.33071445 | 0.361251363 | 15.31114458 | 0.318197251 | 2.027314434 | 1.019569867 | 0.83895091 | 0.000797196 | 3.09843506 | Unpaired t-test | 0.022462963 | permutation FDR (250 perm) | 1 | 0.001295994 FDR correction BH |
| Arc42~Q9VDT1 | VMAT~Control | VMAT | Control | 16.80631966 | 0.194275385 | 15.78793136 | 0.152851967 | 2.02565475 | 1.018388304 | 0.917012463 | 3.35E-06 | 5.474329946 | Unpaired t-test | 0.017393091 | permutation FDR (250 perm) | 1 | 9.74E-06 FDR correction BH |
| Sec22~O77434 | VMAT~Control | VMAT | Control | 19.60908228 | 0.258176848 | 18.59243047 | 0.540209379 | 2.023218038 | 1.016651804 | 0.801926623 | 0.003503815 | 2.455458809 | Unpaired t-test | 0.026358264 | permutation FDR (250 perm) | 1 | 0.005148552 FDR correction BH |
| Dmel\CG14696~Q9VGU7 | VMAT~Control | VMAT | Control | 19.37048031 | 0.07369068 | 18.35469454 | 0.246258145 | 2.022003894 | 1.015785776 | 0.91105531 | 4.88E-06 | 5.311688809 | Unpaired t-test | 0.017654851 | permutation FDR (250 perm) | 1 | 1.35E-05 FDR correction BH |
| Dmel\CG17691~Q7PLE6 | VMAT~Control | VMAT | Control | 17.15967044 | 0.136447503 | 16.14485418 | 0.341637366 | 2.020645519 | 1.014816253 | 0.871446027 | 0.000105745 | 3.975739849 | Unpaired t-test | 0.020051924 | permutation FDR (250 perm) | 1 | 0.000207091 FDR correction BH |
| Stoml2~Q9W1F7 | VMAT~Control | VMAT | Control | 18.17600273 | 0.258100089 | 17.16426138 | 0.173286143 | 2.016343379 | 1.011741348 | 0.888249861 | 2.67E-05 | 4.573232961 | Unpaired t-test | 0.018923792 | permutation FDR (250 perm) | 1 | 6.00E-05 FDR correction BH |
| Gk1~Q9W0U0 | VMAT~Control | VMAT | Control | 15.79150783 | 0.075543638 | 14.78462693 | 0.230344548 | 2.009561739 | 1.006880902 | 0.908399765 | 3.12E-06 | 5.506400823 | Unpaired t-test | 0.017790405 | permutation FDR (250 perm) | 1 | 9.09E-06 FDR correction BH |
| Dmel\CG4020~Q9W459 | VMAT~Control | VMAT | Control | 16.39328231 | 0.310969416 | 15.38759265 | 0.169613117 | 2.007903107 | 1.005689652 | 0.868162901 | 8.37E-05 | 4.077438007 | Unpaired t-test | 0.020216968 | permutation FDR (250 perm) | 1 | 0.000167053 FDR correction BH |
| Sec8~Q9VNH6 | VMAT~Control | VMAT | Control | 16.24029459 | 0.091194855 | 15.23526723 | 0.09191539 | 2.006981558 | 1.00502736 | 0.950016653 | 8.54E-09 | 8.068533609 | Unpaired t-test | 0.015816479 | permutation FDR (250 perm) | 1 | 6.63E-08 FDR correction BH |
| Pvr~B6IDV6 | VMAT~Control | VMAT | Control | 17.8154473 | 0.102668951 | 16.81184273 | 0.186225644 | 2.005003236 | 1.003604565 | 0.916449404 | 9.68E-07 | 6.013928798 | Unpaired t-test | 0.017411765 | permutation FDR (250 perm) | 1 | 3.28E-06 FDR correction BH |
| Dhc64C~M9PE73 | VMAT~Control | VMAT | Control | 18.47171308 | 0.140239037 | 17.47068432 | 0.307071631 | 2.001426676 | 1.001028762 | 0.869726064 | 5.85E-05 | 4.232901884 | Unpaired t-test | 0.020152063 | permutation FDR (250 perm) | 1 | 0.000121635 FDR correction BH |
| Mrgn1~Q9VY98 | VMAT~Control | VMAT | Control | 15.70564619 | 0.18109606 | 14.70510116 | 0.377619722 | 2.000755716 | 1.000545031 | 0.842711476 | 0.000327241 | 3.485131642 | Unpaired t-test | 0.022060213 | permutation FDR (250 perm) | 1 | 0.000574442 FDR correction BH |
| Pcyt2~Q9W0D9 | VMAT~Control | VMAT | Control | 14.76941012 | 0.185146737 | 13.77111184 | 0.336431887 | 1.997642313 | 0.998298285 | 0.851982488 | 0.000169929 | 3.769732464 | Unpaired t-test | 0.021402501 | permutation FDR (250 perm) | 1 | 0.000316998 FDR correction BH |
| Dmel\CG12106~Q9I7W2 | VMAT~Control | VMAT | Control | 16.20394536 | 0.234141711 | 15.20674165 | 0.132213644 | 1.996127266 | 0.997203704 | 0.890160117 | 8.58E-06 | 5.066286695 | Unpaired t-test | 0.018804833 | permutation FDR (250 perm) | 1 | 2.20E-05 FDR correction BH |
| Dmel\CG1907~Q9VAJ9 | VMAT~Control | VMAT | Control | 18.52775856 | 0.104435614 | 17.54548032 | 0.168837605 | 1.975582698 | 0.982278239 | 0.902179219 | 6.25E-07 | 6.20428992 | Unpaired t-test | 0.018147114 | permutation FDR (250 perm) | 1 | 2.25E-06 FDR correction BH |
| E3~Q9VVL7 | VMAT~Control | VMAT | Control | 19.20089652 | 0.193956579 | 18.22030558 | 0.230095057 | 1.97327351 | 0.980590938 | 0.864274325 | 2.64E-05 | 4.577761396 | Unpaired t-test | 0.020500695 | permutation FDR (250 perm) | 1 | 5.95E-05 FDR correction BH |
| slgA~M9NFJ2 | VMAT~Control | VMAT | Control | 15.18041868 | 0.152280526 | 14.20084165 | 0.63285498 | 1.971887201 | 0.979577027 | 0.758715325 | 0.007180491 | 2.143845886 | Unpaired t-test | 0.032080845 | permutation FDR (250 perm) | 1 | 0.010178869 FDR correction BH |
| CalpA~A8DYJ3 | VMAT~Control | VMAT | Control | 17.80383498 | 0.132559182 | 16.82516133 | 0.186525997 | 1.970652855 | 0.978673657 | 0.887817451 | 2.39E-06 | 5.62164814 | Unpaired t-test | 0.018964684 | permutation FDR (250 perm) | 1 | 7.17E-06 FDR correction BH |
| jus~Q8MQS2 | VMAT~Control | VMAT | Control | 17.13113617 | 0.161656777 | 16.15340064 | 0.306674242 | 1.969371837 | 0.977735532 | 0.846497319 | 8.83E-05 | 4.053836162 | Unpaired t-test | 0.021826772 | permutation FDR (250 perm) | 1 | 0.000175935 FDR correction BH |
| Tim17b~Q7PLT4 | VMAT~Control | VMAT | Control | 16.57979597 | 0.134297286 | 15.60493559 | 0.142206853 | 1.965450989 | 0.974860388 | 0.896444734 | 5.84E-07 | 6.23374305 | Unpaired t-test | 0.018426778 | permutation FDR (250 perm) | 1 | 2.12E-06 FDR correction BH |
| rdgB~E1JJK8 | VMAT~Control | VMAT | Control | 16.82780165 | 0.113885254 | 15.85701568 | 0.258627273 | 1.959908056 | 0.970785975 | 0.861864619 | 1.68E-05 | 4.775313195 | Unpaired t-test | 0.020654613 | permutation FDR (250 perm) | 1 | 3.95E-05 FDR correction BH |
| Cyt-b5-r~M9PD26 | VMAT~Control | VMAT | Control | 16.06079813 | 0.19275925 | 15.09074382 | 0.308630126 | 1.958914333 | 0.970054307 | 0.834288747 | 0.000139166 | 3.856466359 | Unpaired t-test | 0.022926423 | permutation FDR (250 perm) | 1 | 0.000264618 FDR correction BH |
| Rac1~M9PBH7 | VMAT~Control | VMAT | Control | 16.70077381 | 0.068647531 | 15.73128108 | 0.229442793 | 1.95815197 | 0.969492735 | 0.875701485 | 3.93E-06 | 5.405657679 | Unpaired t-test | 0.019760446 | permutation FDR (250 perm) | 1 | 1.12E-05 FDR correction BH |
| Dmel\CG8888~Q7K3N4 | VMAT~Control | VMAT | Control | 16.8067334 | 0.139996533 | 15.83739301 | 0.134439326 | 1.957945208 | 0.969340393 | 0.891919932 | 5.73E-07 | 6.241972878 | Unpaired t-test | 0.018715016 | permutation FDR (250 perm) | 1 | 2.08E-06 FDR correction BH |
| Nrx-IV~Q8IQH0 | VMAT~Control | VMAT | Control | 14.5428325 | 0.251420872 | 13.57665063 | 0.283437858 | 1.953663317 | 0.966181863 | 0.826191945 | 0.000197707 | 3.703978713 | Unpaired t-test | 0.023742712 | permutation FDR (250 perm) | 1 | 0.000364449 FDR correction BH |
| Shab~M9MRU9 | VMAT~Control | VMAT | Control | 17.1157117 | 0.160485093 | 16.14970173 | 0.313699528 | 1.953430556 | 0.966009969 | 0.83450562 | 0.000111187 | 3.953946137 | Unpaired t-test | 0.022900509 | permutation FDR (250 perm) | 1 | 0.000215419 FDR correction BH |
| Dmel\CG6180~Q9VK60 | VMAT~Control | VMAT | Control | 17.87824472 | 0.19587647 | 16.92672664 | 0.148387261 | 1.933906545 | 0.951518079 | 0.857303314 | 5.85E-06 | 5.232659261 | Unpaired t-test | 0.020984245 | permutation FDR (250 perm) | 1 | 1.60E-05 FDR correction BH |
| Dmel\CG8728~Q7K3W2 | VMAT~Control | VMAT | Control | 16.49376783 | 0.250378905 | 15.54369599 | 0.376630732 | 1.931968853 | 0.950071835 | 0.790239948 | 0.000844969 | 3.073159079 | Unpaired t-test | 0.027646083 | permutation FDR (250 perm) | 1 | 0.00136799 FDR correction BH |
| nrv3~Q7JS69 | VMAT~Control | VMAT | Control | 18.5680806 | 0.162293245 | 17.62090026 | 0.269567406 | 1.928100607 | 0.947180332 | 0.830338057 | 5.16E-05 | 4.287235967 | Unpaired t-test | 0.023307728 | permutation FDR (250 perm) | 1 | 0.000108713 FDR correction BH |
| nrv1~B5RIR1 | VMAT~Control | VMAT | Control | 16.55020867 | 0.24623545 | 15.60483421 | 0.480839139 | 1.925688645 | 0.94537446 | 0.761419998 | 0.002898783 | 2.537784265 | Unpaired t-test | 0.031678309 | permutation FDR (250 perm) | 1 | 0.004305196 FDR correction BH |
| alphaTub85E~P06604 | VMAT~Control | VMAT | Control | 17.13139747 | 0.112008328 | 16.18698079 | 0.586954149 | 1.924410644 | 0.944416684 | 0.745260306 | 0.005409802 | 2.2668186 | Unpaired t-test | 0.034189995 | permutation FDR (250 perm) | 1 | 0.007774527 FDR correction BH |
| anon-WO0140519.147~Q9VB43 | VMAT~Control | VMAT | Control | 15.41196193 | 0.722550028 | 14.47218022 | 1.175709314 | 1.918237974 | 0.93978171 | 0.581134514 | 0.158790214 | 0.799176266 | Unpaired t-test | 0.081261261 | permutation FDR (250 perm) | 1 | 0.184588723 FDR correction BH |
| Mpcp2~Q0E8E8 | VMAT~Control | VMAT | Control | 18.65033685 | 0.11024137 | 17.71570132 | 0.09718103 | 1.911407695 | 0.934635532 | 0.876996994 | 5.83E-08 | 7.234271684 | Unpaired t-test | 0.019675023 | permutation FDR (250 perm) | 1 | 3.08E-07 FDR correction BH |
| Ras85D~P08646 | VMAT~Control | VMAT | Control | 16.49244848 | 0.115236022 | 15.5596582 | 0.162181258 | 1.908964506 | 0.932790278 | 0.85657697 | 1.03E-06 | 5.987518891 | Unpaired t-test | 0.021030584 | permutation FDR (250 perm) | 1 | 3.47E-06 FDR correction BH |
| Rop~M9PEL1 | VMAT~Control | VMAT | Control | 18.29895286 | 0.0488451 | 17.36775923 | 0.124473612 | 1.906853 | 0.93119363 | 0.878651277 | 2.44E-08 | 7.61282816 | Unpaired t-test | 0.019537604 | permutation FDR (250 perm) | 1 | 1.53E-07 FDR correction BH |
| RpL37A~M9MRF2 | VMAT~Control | VMAT | Control | 18.32167615 | 0.163687427 | 17.3920815 | 0.192763145 | 1.904740763 | 0.929594659 | 0.835144833 | 9.28E-06 | 5.032423297 | Unpaired t-test | 0.022835724 | permutation FDR (250 perm) | 1 | 2.36E-05 FDR correction BH |
| Rpn3~M9PG62 | VMAT~Control | VMAT | Control | 16.16825901 | 0.147823195 | 15.23890199 | 0.20490607 | 1.904427047 | 0.929357024 | 0.835006337 | 9.23E-06 | 5.034830554 | Unpaired t-test | 0.022857936 | permutation FDR (250 perm) | 1 | 2.35E-05 FDR correction BH |
| ben~E1JJN1 | VMAT~Control | VMAT | Control | 16.17988912 | 0.192071458 | 15.25470074 | 0.247363548 | 1.898932174 | 0.925188376 | 0.811527995 | 6.03E-05 | 4.219379383 | Unpaired t-test | 0.025358002 | permutation FDR (250 perm) | 1 | 0.000124711 FDR correction BH |
| scpr-B~Q9VGQ4 | VMAT~Control | VMAT | Control | 16.16170364 | 0.366266212 | 15.23697919 | 0.209320559 | 1.898321637 | 0.924724452 | 0.777954328 | 0.000621729 | 3.206399056 | Unpaired t-test | 0.029266452 | permutation FDR (250 perm) | 1 | 0.001028496 FDR correction BH |
| Egm~Q5U117 | VMAT~Control | VMAT | Control | 15.3536855 | 0.243379401 | 14.42976034 | 0.803231221 | 1.897270213 | 0.923925165 | 0.671777725 | 0.033587502 | 1.473822295 | Unpaired t-test | 0.049610909 | permutation FDR (250 perm) | 1 | 0.042946757 FDR correction BH |
| TSG101~Q9VVA7 | VMAT~Control | VMAT | Control | 14.86971348 | 0.1870662 | 13.94799714 | 0.63827951 | 1.894367634 | 0.921716337 | 0.71040385 | 0.011275886 | 1.947849325 | Unpaired t-test | 0.040551346 | permutation FDR (250 perm) | 1 | 0.01543223 FDR correction BH |
| fab1~O96838 | VMAT~Control | VMAT | Control | 16.3489672 | 0.125864799 | 15.4279086 | 0.096273134 | 1.893504177 | 0.921058604 | 0.860105728 | 1.37E-07 | 6.861867332 | Unpaired t-test | 0.020773296 | permutation FDR (250 perm) | 1 | 6.34E-07 FDR correction BH |
| CG17093~Q9VKT1 | VMAT~Control | VMAT | Control | 18.65762958 | 0.313715991 | 17.73706148 | 0.139530338 | 1.892860506 | 0.920568096 | 0.79803122 | 0.000132944 | 3.87633007 | Unpaired t-test | 0.026754613 | permutation FDR (250 perm) | 1 | 0.000253813 FDR correction BH |
| Abl~M9PCU1 | VMAT~Control | VMAT | Control | 16.47096985 | 0.150403132 | 15.55070845 | 0.209320505 | 1.892458155 | 0.9202614 | 0.825146536 | 1.20E-05 | 4.921194263 | Unpaired t-test | 0.023840814 | permutation FDR (250 perm) | 1 | 2.95E-05 FDR correction BH |
| iPLA2-VIA~Q9VT60 | VMAT~Control | VMAT | Control | 17.61250479 | 0.175914236 | 16.69265202 | 0.201128688 | 1.891922207 | 0.919852769 | 0.821665577 | 1.65E-05 | 4.783196121 | Unpaired t-test | 0.024236927 | permutation FDR (250 perm) | 1 | 3.89E-05 FDR correction BH |
| lost~Q9VN21 | VMAT~Control | VMAT | Control | 18.33059546 | 0.085029843 | 17.41120214 | 0.097944165 | 1.891319803 | 0.919393329 | 0.868987367 | 2.06E-08 | 7.686942209 | Unpaired t-test | 0.020189152 | permutation FDR (250 perm) | 1 | 1.32E-07 FDR correction BH |
| Agpat3~D5AEK7 | VMAT~Control | VMAT | Control | 17.09329701 | 0.085985303 | 16.17399985 | 0.119122644 | 1.891193738 | 0.919297163 | 0.862621332 | 6.81E-08 | 7.166873449 | Unpaired t-test | 0.020617524 | permutation FDR (250 perm) | 1 | 3.44E-07 FDR correction BH |
| Pten~Q7KMQ6 | VMAT~Control | VMAT | Control | 15.36638494 | 0.104480859 | 14.45074687 | 0.973371937 | 1.886403195 | 0.915638067 | 0.63683018 | 0.062978065 | 1.20081069 | Unpaired t-test | 0.060252154 | permutation FDR (250 perm) | 1 | 0.077536801 FDR correction BH |
| Lon~Q7KUT2-2 | VMAT~Control | VMAT | Control | 15.97784708 | 0.099301714 | 15.06229243 | 0.238164226 | 1.886294127 | 0.915554651 | 0.820832546 | 1.27E-05 | 4.897519089 | Unpaired t-test | 0.024344285 | permutation FDR (250 perm) | 1 | 3.09E-05 FDR correction BH |
| Itpr~A0A0B4LGN1 | VMAT~Control | VMAT | Control | 14.74848533 | 0.299128846 | 13.83605351 | 0.363268836 | 1.882215505 | 0.912431819 | 0.753796641 | 0.001476389 | 2.830799186 | Unpaired t-test | 0.032783846 | permutation FDR (250 perm) | 1 | 0.002289473 FDR correction BH |
| EMC8-9~Q9W1Y1 | VMAT~Control | VMAT | Control | 15.72379854 | 0.386967657 | 14.81233518 | 0.107445675 | 1.880952427 | 0.911463361 | 0.772685682 | 0.000481405 | 3.317489243 | Unpaired t-test | 0.030075437 | permutation FDR (250 perm) | 1 | 0.000813516 FDR correction BH |
| Nedd8~X2J935 | VMAT~Control | VMAT | Control | 16.06685332 | 0.371381634 | 15.15806131 | 0.234566061 | 1.877472808 | 0.908792013 | 0.759579392 | 0.000941506 | 3.026177085 | Unpaired t-test | 0.031944878 | permutation FDR (250 perm) | 1 | 0.001509744 FDR correction BH |
| L2HGDH~Q9VJ28 | VMAT~Control | VMAT | Control | 17.62896923 | 0.098765036 | 16.72311521 | 0.107353254 | 1.873653293 | 0.905854016 | 0.850378007 | 7.32E-08 | 7.13522918 | Unpaired t-test | 0.021532191 | permutation FDR (250 perm) | 1 | 3.63E-07 FDR correction BH |
| Dmel\CG8778~A1Z934 | VMAT~Control | VMAT | Control | 16.50048783 | 0.119757297 | 15.59876313 | 0.207217534 | 1.868298154 | 0.901724707 | 0.814541436 | 7.46E-06 | 5.127022947 | Unpaired t-test | 0.024963922 | permutation FDR (250 perm) | 1 | 1.95E-05 FDR correction BH |
| FAM21~A1ZBW7 | VMAT~Control | VMAT | Control | 16.48453931 | 0.223882025 | 15.58395476 | 0.189573965 | 1.866822231 | 0.900584553 | 0.796135116 | 4.38E-05 | 4.358485893 | Unpaired t-test | 0.02696679 | permutation FDR (250 perm) | 1 | 9.37E-05 FDR correction BH |
| Mtpalpha~Q8IPE8 | VMAT~Control | VMAT | Control | 18.94658749 | 0.241706316 | 18.04780043 | 0.305324217 | 1.86449776 | 0.898787064 | 0.765477524 | 0.000424654 | 3.371964646 | Unpaired t-test | 0.031152184 | permutation FDR (250 perm) | 1 | 0.000727008 FDR correction BH |
| Dmel\CG31051~Q8IMM3 | VMAT~Control | VMAT | Control | 14.20616476 | 1.053717085 | 13.31048031 | 0.329392556 | 1.860492332 | 0.895684444 | 0.599631643 | 0.099726694 | 1.001188577 | Unpaired t-test | 0.073734177 | permutation FDR (250 perm) | 1 | 0.119651701 FDR correction BH |
| Ndg~A1Z877 | VMAT~Control | VMAT | Control | 16.66840373 | 0.177627248 | 15.77475852 | 0.34270547 | 1.857864395 | 0.893645203 | 0.762088929 | 0.000415012 | 3.381939728 | Unpaired t-test | 0.031606618 | permutation FDR (250 perm) | 1 | 0.000711017 FDR correction BH |
| Sccpdh2~Q9VG81 | VMAT~Control | VMAT | Control | 17.29179683 | 0.204868578 | 16.39897025 | 0.139492197 | 1.856810499 | 0.892826586 | 0.803738943 | 1.11E-05 | 4.955118367 | Unpaired t-test | 0.026176443 | permutation FDR (250 perm) | 1 | 2.75E-05 FDR correction BH |
| ACC~A1Z784 | VMAT~Control | VMAT | Control | 21.57141156 | 0.263659859 | 20.67995152 | 0.07534495 | 1.855052535 | 0.891460044 | 0.79408018 | 2.70E-05 | 4.569214619 | Unpaired t-test | 0.027202952 | permutation FDR (250 perm) | 1 | 6.05E-05 FDR correction BH |
| Usp8~Q9VDD8 | VMAT~Control | VMAT | Control | 15.42788916 | 0.541314606 | 14.53759598 | 0.255685538 | 1.853552753 | 0.890293175 | 0.702273528 | 0.007677972 | 2.114753503 | Unpaired t-test | 0.042238833 | permutation FDR (250 perm) | 1 | 0.010848706 FDR correction BH |
| RpL23~P48159 | VMAT~Control | VMAT | Control | 20.24940698 | 0.171004894 | 19.36124846 | 0.105638108 | 1.850812203 | 0.888158517 | 0.814905714 | 1.77E-06 | 5.75086943 | Unpaired t-test | 0.02493802 | permutation FDR (250 perm) | 1 | 5.56E-06 FDR correction BH |
| RhoGAP100F~A0A0B4K7G3 | VMAT~Control | VMAT | Control | 19.55358912 | 0.155406895 | 18.66822328 | 0.139762 | 1.84723299 | 0.885365844 | 0.809683422 | 2.61E-06 | 5.583807999 | Unpaired t-test | 0.025557097 | permutation FDR (250 perm) | 1 | 7.76E-06 FDR correction BH |
| Ent2~Q9VMB6 | VMAT~Control | VMAT | Control | 16.22823292 | 0.112011164 | 15.34307572 | 0.217100372 | 1.846965858 | 0.885157197 | 0.797977261 | 1.05E-05 | 4.977329003 | Unpaired t-test | 0.026756458 | permutation FDR (250 perm) | 1 | 2.62E-05 FDR correction BH |
| RpS2~M9PB84 | VMAT~Control | VMAT | Control | 19.93888781 | 0.122764679 | 19.05520356 | 0.090148021 | 1.845081116 | 0.883684243 | 0.827331112 | 1.40E-07 | 6.85440639 | Unpaired t-test | 0.023639056 | permutation FDR (250 perm) | 1 | 6.39E-07 FDR correction BH |
| FBgn 33734~Q7K4T8 | VMAT~Control | VMAT | Control | 16.60661829 | 0.077845373 | 15.72358747 | 0.058177115 | 1.844245642 | 0.883030827 | 0.846251527 | 1.84E-09 | 8.735269566 | Unpaired t-test | 0.021839741 | permutation FDR (250 perm) | 1 | 2.00E-08 FDR correction BH |
| mEFTu2~Q7K3V6 | VMAT~Control | VMAT | Control | 16.44476109 | 0.076926594 | 15.56335005 | 0.148669699 | 1.842176185 | 0.881411047 | 0.820023759 | 3.49E-07 | 6.456865959 | Unpaired t-test | 0.024431282 | permutation FDR (250 perm) | 1 | 1.37E-06 FDR correction BH |
| Tango5~Q9W2S1 | VMAT~Control | VMAT | Control | 17.72553055 | 0.10700707 | 16.84464366 | 0.241418023 | 1.841507016 | 0.880886894 | 0.787845583 | 2.16E-05 | 4.665094468 | Unpaired t-test | 0.027902349 | permutation FDR (250 perm) | 1 | 4.97E-05 FDR correction BH |
| Use1~Q9VSU7 | VMAT~Control | VMAT | Control | 16.73947067 | 0.140167593 | 15.85880374 | 0.186042185 | 1.841226269 | 0.880666931 | 0.797581555 | 7.24E-06 | 5.140400172 | Unpaired t-test | 0.026797048 | permutation FDR (250 perm) | 1 | 1.91E-05 FDR correction BH |
| RpL18A~P41093 | VMAT~Control | VMAT | Control | 19.19644933 | 0.187828241 | 18.31704641 | 0.322399513 | 1.839613791 | 0.879402919 | 0.753645325 | 0.000362748 | 3.440395396 | Unpaired t-test | 0.032802203 | permutation FDR (250 perm) | 1 | 0.000629726 FDR correction BH |
| Irp-1A~Q9VCV4 | VMAT~Control | VMAT | Control | 15.42098629 | 0.326693653 | 14.54216891 | 0.79604651 | 1.838867301 | 0.878817374 | 0.634609248 | 0.04549619 | 1.342024973 | Unpaired t-test | 0.061008617 | permutation FDR (250 perm) | 1 | 0.057088503 FDR correction BH |
| vas~M9PBB5 | VMAT~Control | VMAT | Control | 15.31172678 | 0.23770407 | 14.43356603 | 0.187661062 | 1.838030547 | 0.878160744 | 0.773410117 | 7.04E-05 | 4.152492915 | Unpaired t-test | 0.029972401 | permutation FDR (250 perm) | 1 | 0.000142842 FDR correction BH |
| AP-1gamma~Q9W388 | VMAT~Control | VMAT | Control | 14.65269053 | 0.097680571 | 13.7751051 | 0.37070106 | 1.837297726 | 0.877585428 | 0.749150087 | 0.000451436 | 3.345404289 | Unpaired t-test | 0.033529142 | permutation FDR (250 perm) | 1 | 0.0007695 FDR correction BH |
| anon-EST:Posey113~Q9W4R9 | VMAT~Control | VMAT | Control | 16.39966076 | 0.269114939 | 15.52214217 | 0.254954683 | 1.837212616 | 0.877518596 | 0.752727094 | 0.000351001 | 3.454692012 | Unpaired t-test | 0.032961909 | permutation FDR (250 perm) | 1 | 0.000612495 FDR correction BH |
| PMCA~Q59DP9 | VMAT~Control | VMAT | Control | 19.87422698 | 0.027333914 | 18.99876865 | 0.075678917 | 1.834590821 | 0.875458326 | 0.845049592 | 3.14E-10 | 9.503445123 | Unpaired t-test | 0.021943492 | permutation FDR (250 perm) | 1 | 5.16E-09 FDR correction BH |
| Mdh1~Q9VKX2 | VMAT~Control | VMAT | Control | 16.74312075 | 0.50527137 | 15.86813229 | 0.577077796 | 1.833993419 | 0.874988462 | 0.651507527 | 0.02882297 | 1.540261265 | Unpaired t-test | 0.05551317 | permutation FDR (250 perm) | 1 | 0.03711667 FDR correction BH |
| Faf2~Q9VJ58 | VMAT~Control | VMAT | Control | 17.58990192 | 0.260980195 | 16.71625164 | 0.467615056 | 1.832293078 | 0.873650283 | 0.704847241 | 0.00447725 | 2.348988686 | Unpaired t-test | 0.041620784 | permutation FDR (250 perm) | 1 | 0.006481824 FDR correction BH |
| Dmel\CG1774~Q9V9W4 | VMAT~Control | VMAT | Control | 15.42030256 | 0.321215262 | 14.55095667 | 0.162296668 | 1.826834435 | 0.869345889 | 0.748824879 | 0.000300803 | 3.521718367 | Unpaired t-test | 0.033591556 | permutation FDR (250 perm) | 1 | 0.000536431 FDR correction BH |
| Pde9~Q9I7S6 | VMAT~Control | VMAT | Control | 15.06619137 | 0.317653563 | 14.19709707 | 0.170358248 | 1.82651589 | 0.869094304 | 0.748445567 | 0.000305101 | 3.515555766 | Unpaired t-test | 0.033659477 | permutation FDR (250 perm) | 1 | 0.000542864 FDR correction BH |
| Flo2~A4V4F2 | VMAT~Control | VMAT | Control | 17.44306533 | 0.05333203 | 16.57969375 | 0.118785607 | 1.819285018 | 0.86337158 | 0.815862954 | 3.91E-08 | 7.407897153 | Unpaired t-test | 0.024838113 | permutation FDR (250 perm) | 1 | 2.23E-07 FDR correction BH |
| SK~X2JIE0 | VMAT~Control | VMAT | Control | 15.33160757 | 0.194875513 | 14.46955692 | 0.129240453 | 1.817620049 | 0.862050653 | 0.780436568 | 9.05E-06 | 5.043456223 | Unpaired t-test | 0.028941068 | permutation FDR (250 perm) | 1 | 2.31E-05 FDR correction BH |
| Mdh2~Q9VEB1 | VMAT~Control | VMAT | Control | 17.8616505 | 0.157125397 | 17.00031048 | 0.289456384 | 1.816724962 | 0.861340024 | 0.750759767 | 0.000162075 | 3.790283653 | Unpaired t-test | 0.033248279 | permutation FDR (250 perm) | 1 | 0.000304732 FDR correction BH |
| Blos4~M9PET7 | VMAT~Control | VMAT | Control | 16.05478182 | 0.194879491 | 15.19456315 | 0.338284643 | 1.815313435 | 0.860218668 | 0.732354334 | 0.000598623 | 3.22284678 | Unpaired t-test | 0.036364302 | permutation FDR (250 perm) | 1 | 0.000992365 FDR correction BH |
| Ucp4A~Q9VX14 | VMAT~Control | VMAT | Control | 16.15246728 | 0.09373669 | 15.29625957 | 0.34095754 | 1.810273533 | 0.856207705 | 0.739296598 | 0.00029538 | 3.529618604 | Unpaired t-test | 0.035189363 | permutation FDR (250 perm) | 1 | 0.000527561 FDR correction BH |
| ND-39~Q9VPE2 | VMAT~Control | VMAT | Control | 19.2290405 | 0.190914354 | 18.37283303 | 0.194499935 | 1.810273236 | 0.856207469 | 0.763187182 | 3.61E-05 | 4.442933264 | Unpaired t-test | 0.031470588 | permutation FDR (250 perm) | 1 | 7.90E-05 FDR correction BH |
| Cyp12a4~Q9VE00 | VMAT~Control | VMAT | Control | 15.656653 | 0.099556447 | 14.80523038 | 0.182222835 | 1.804279219 | 0.851422619 | 0.77907614 | 3.50E-06 | 5.455853789 | Unpaired t-test | 0.029117864 | permutation FDR (250 perm) | 1 | 1.01E-05 FDR correction BH |
| msk~Q9VSD6 | VMAT~Control | VMAT | Control | 18.04435494 | 0.20658859 | 17.19296874 | 0.336392898 | 1.804233671 | 0.851386198 | 0.723633076 | 0.000699585 | 3.15515954 | Unpaired t-test | 0.037906679 | permutation FDR (250 perm) | 1 | 0.001144422 FDR correction BH |
| Tsf2~Q9VTZ5 | VMAT~Control | VMAT | Control | 17.12364233 | 0.309728985 | 16.27250131 | 0.11799955 | 1.803927072 | 0.851141015 | 0.741265582 | 0.000187166 | 3.727771929 | Unpaired t-test | 0.034863699 | permutation FDR (250 perm) | 1 | 0.000346376 FDR correction BH |
| Eaat1~O77062 | VMAT~Control | VMAT | Control | 17.51267202 | 0.071661228 | 16.66185164 | 0.114350418 | 1.80352619 | 0.850820374 | 0.802394863 | 6.34E-08 | 7.198160754 | Unpaired t-test | 0.026297322 | permutation FDR (250 perm) | 1 | 3.24E-07 FDR correction BH |
| Dmel\CG17544~Q9VIX4 | VMAT~Control | VMAT | Control | 16.87551877 | 0.146434337 | 16.02531952 | 0.175678628 | 1.802749886 | 0.850199251 | 0.771309693 | 8.40E-06 | 5.075464734 | Unpaired t-test | 0.03025391 | permutation FDR (250 perm) | 1 | 2.16E-05 FDR correction BH |
| inaC~P13677 | VMAT~Control | VMAT | Control | 17.95021993 | 0.09822514 | 17.10154612 | 0.219947675 | 1.800844745 | 0.848673809 | 0.766140101 | 1.35E-05 | 4.870708776 | Unpaired t-test | 0.031008736 | permutation FDR (250 perm) | 1 | 3.26E-05 FDR correction BH |
| Start1~Q9W145 | VMAT~Control | VMAT | Control | 16.22302483 | 0.132476438 | 15.37448271 | 0.146177928 | 1.800680371 | 0.848542119 | 0.779748994 | 2.27E-06 | 5.644181661 | Unpaired t-test | 0.029025783 | permutation FDR (250 perm) | 1 | 6.90E-06 FDR correction BH |
| AP-1-2beta~Q24253 | VMAT~Control | VMAT | Control | 17.62502357 | 0.091044419 | 16.7769743 | 0.043070831 | 1.800065327 | 0.848049265 | 0.81149731 | 3.87E-09 | 8.412259382 | Unpaired t-test | 0.025361702 | permutation FDR (250 perm) | 1 | 3.53E-08 FDR correction BH |
| betaTub85D~P61857 | VMAT~Control | VMAT | Control | 19.36488706 | 0.246035514 | 18.51688094 | 0.110807163 | 1.8000115 | 0.848006124 | 0.756692605 | 3.60E-05 | 4.444209512 | Unpaired t-test | 0.032336088 | permutation FDR (250 perm) | 1 | 7.90E-05 FDR correction BH |
| galectin~Q9VPI6 | VMAT~Control | VMAT | Control | 17.09097817 | 0.102310314 | 16.24497475 | 0.116014379 | 1.797514522 | 0.846003426 | 0.791266605 | 2.44E-07 | 6.61192996 | Unpaired t-test | 0.027555556 | permutation FDR (250 perm) | 1 | 1.02E-06 FDR correction BH |
| Idi~Q9VDC2 | VMAT~Control | VMAT | Control | 15.20927093 | 0.109184949 | 14.36409714 | 0.48248495 | 1.796481132 | 0.845173783 | 0.69206778 | 0.003371666 | 2.472155499 | Unpaired t-test | 0.044411658 | permutation FDR (250 perm) | 1 | 0.004963916 FDR correction BH |
| alt~Q960Y8 | VMAT~Control | VMAT | Control | 18.98835057 | 0.208813512 | 18.14362588 | 0.138014536 | 1.795922 | 0.844724692 | 0.759686549 | 1.96E-05 | 4.708600668 | Unpaired t-test | 0.031920992 | permutation FDR (250 perm) | 1 | 4.54E-05 FDR correction BH |
| PlexA~Q9V491 | VMAT~Control | VMAT | Control | 17.64147345 | 0.392039163 | 16.79822463 | 0.393958261 | 1.794085724 | 0.843248826 | 0.675379826 | 0.006855746 | 2.163945309 | Unpaired t-test | 0.04864211 | permutation FDR (250 perm) | 1 | 0.00973367 FDR correction BH |
| chp~A0A0B4KHG5 | VMAT~Control | VMAT | Control | 16.46888361 | 0.196664762 | 15.62709856 | 0.287577802 | 1.792266352 | 0.841785055 | 0.728309798 | 0.000300241 | 3.522530085 | Unpaired t-test | 0.03705764 | permutation FDR (250 perm) | 1 | 0.000535836 FDR correction BH |
| gprs~A0A0B4KF89 | VMAT~Control | VMAT | Control | 15.16809305 | 0.576338055 | 14.32679225 | 0.36034886 | 1.791664863 | 0.841300802 | 0.645179604 | 0.019868076 | 1.70184418 | Unpaired t-test | 0.057508167 | permutation FDR (250 perm) | 1 | 0.026245484 FDR correction BH |
| wal~Q7KN94 | VMAT~Control | VMAT | Control | 18.84715669 | 0.125224542 | 18.00615529 | 0.186342222 | 1.791293074 | 0.841001397 | 0.764266173 | 7.86E-06 | 5.104806424 | Unpaired t-test | 0.031323529 | permutation FDR (250 perm) | 1 | 2.03E-05 FDR correction BH |
| Sc2~Q9VZL3 | VMAT~Control | VMAT | Control | 18.10950173 | 0.082680272 | 17.26969916 | 0.096189861 | 1.789805183 | 0.839802562 | 0.794722106 | 3.96E-08 | 7.401769307 | Unpaired t-test | 0.027136531 | permutation FDR (250 perm) | 1 | 2.26E-07 FDR correction BH |
| Phb2~Q0E924 | VMAT~Control | VMAT | Control | 18.92334352 | 0.205143022 | 18.08931024 | 0.229906126 | 1.782662119 | 0.834033284 | 0.733024486 | 0.000123181 | 3.909455433 | Unpaired t-test | 0.036267277 | permutation FDR (250 perm) | 1 | 0.000236902 FDR correction BH |
| ND-49~Q9V4E0 | VMAT~Control | VMAT | Control | 15.60679919 | 0.406409335 | 14.77392735 | 0.20666819 | 1.78122756 | 0.832871839 | 0.691810284 | 0.002197881 | 2.657995903 | Unpaired t-test | 0.044453552 | permutation FDR (250 perm) | 1 | 0.003329351 FDR correction BH |
| GstS1~A4UZL5 | VMAT~Control | VMAT | Control | 18.86205572 | 0.275912639 | 18.03040804 | 0.363905874 | 1.779716787 | 0.831647678 | 0.690603718 | 0.002242754 | 2.649218446 | Unpaired t-test | 0.044697632 | permutation FDR (250 perm) | 1 | 0.003388602 FDR correction BH |
| Vha68-1~M9PD18 | VMAT~Control | VMAT | Control | 18.80181417 | 0.068487661 | 17.97148676 | 0.075867631 | 1.778088844 | 0.830327411 | 0.794033135 | 5.48E-09 | 8.26083341 | Unpaired t-test | 0.027214022 | permutation FDR (250 perm) | 1 | 4.66E-08 FDR correction BH |
| mRpS5~Q8SYP9 | VMAT~Control | VMAT | Control | 16.9109061 | 0.183711546 | 16.08091123 | 0.232829463 | 1.777679044 | 0.829994872 | 0.732800304 | 9.41E-05 | 4.026418361 | Unpaired t-test | 0.036298398 | permutation FDR (250 perm) | 1 | 0.000186141 FDR correction BH |
| COX4~Q9VIQ8 | VMAT~Control | VMAT | Control | 19.40330785 | 0.214224106 | 18.57368674 | 0.361799359 | 1.777218551 | 0.829621105 | 0.698312147 | 0.001310343 | 2.882615038 | Unpaired t-test | 0.0430866 | permutation FDR (250 perm) | 1 | 0.002050896 FDR correction BH |
| sea~Q7KSQ0 | VMAT~Control | VMAT | Control | 18.0026547 | 0.099683105 | 17.17509141 | 0.136226126 | 1.774685387 | 0.827563289 | 0.76947503 | 6.81E-07 | 6.167152912 | Unpaired t-test | 0.03049126 | permutation FDR (250 perm) | 1 | 2.41E-06 FDR correction BH |
| ND-B16.6~Q9W402 | VMAT~Control | VMAT | Control | 18.09874146 | 0.243039531 | 17.27149977 | 0.339660588 | 1.774289829 | 0.827241692 | 0.697046054 | 0.001276283 | 2.894053086 | Unpaired t-test | 0.043305695 | permutation FDR (250 perm) | 1 | 0.002002913 FDR correction BH |
| Aos1~Q7KSM5 | VMAT~Control | VMAT | Control | 16.19664163 | 0.160636776 | 15.37082685 | 0.176476943 | 1.772535814 | 0.825814777 | 0.746180637 | 1.57E-05 | 4.80288836 | Unpaired t-test | 0.034002754 | permutation FDR (250 perm) | 1 | 3.75E-05 FDR correction BH |
| Liprin-alpha~Q9VM93 | VMAT~Control | VMAT | Control | 17.26120877 | 0.078192039 | 16.43597224 | 0.157779155 | 1.771825508 | 0.825236532 | 0.764992909 | 1.03E-06 | 5.985719842 | Unpaired t-test | 0.031216552 | permutation FDR (250 perm) | 1 | 3.47E-06 FDR correction BH |
| yip2~Q9VL70 | VMAT~Control | VMAT | Control | 19.20449452 | 0.30577487 | 18.38043268 | 0.342198842 | 1.770383414 | 0.82406184 | 0.683737811 | 0.002457004 | 2.609594143 | Unpaired t-test | 0.046480437 | permutation FDR (250 perm) | 1 | 0.003693351 FDR correction BH |
| RpL21~Q9V9M7 | VMAT~Control | VMAT | Control | 18.35489556 | 0.214147934 | 17.53613103 | 0.196366198 | 1.76389481 | 0.818764529 | 0.724610333 | 8.89E-05 | 4.050878548 | Unpaired t-test | 0.037714547 | permutation FDR (250 perm) | 1 | 0.000176838 FDR correction BH |
| spidey~Q9W3N1 | VMAT~Control | VMAT | Control | 17.22896576 | 0.087797839 | 16.41027008 | 0.141184587 | 1.763810633 | 0.818695678 | 0.762036296 | 6.53E-07 | 6.184995167 | Unpaired t-test | 0.031608456 | permutation FDR (250 perm) | 1 | 2.32E-06 FDR correction BH |
| Dmel\CG6330~Q8IMQ8 | VMAT~Control | VMAT | Control | 16.40640225 | 0.168947364 | 15.58821129 | 0.181446395 | 1.763193687 | 0.818190963 | 0.736528572 | 2.37E-05 | 4.625047543 | Unpaired t-test | 0.035670027 | permutation FDR (250 perm) | 1 | 5.41E-05 FDR correction BH |
| vnc~Q9VT75 | VMAT~Control | VMAT | Control | 16.32734922 | 0.116767729 | 15.51020398 | 0.276119201 | 1.761916122 | 0.817145245 | 0.720540883 | 0.000116487 | 3.933721626 | Unpaired t-test | 0.038511203 | permutation FDR (250 perm) | 1 | 0.000224763 FDR correction BH |
| Dmel\CG11771~Q9VC06 | VMAT~Control | VMAT | Control | 15.57358667 | 0.261667053 | 14.75892674 | 0.201170265 | 1.758883501 | 0.81465993 | 0.709877105 | 0.000254869 | 3.593682146 | Unpaired t-test | 0.040660885 | permutation FDR (250 perm) | 1 | 0.000463264 FDR correction BH |
| Afg3l2~Q8T4G5 | VMAT~Control | VMAT | Control | 17.07636933 | 0.088316102 | 16.26216724 | 0.090568411 | 1.758325408 | 0.814202091 | 0.77060676 | 5.20E-08 | 7.284027059 | Unpaired t-test | 0.030344066 | permutation FDR (250 perm) | 1 | 2.80E-07 FDR correction BH |
| CG12163~Q9VN93-2 | VMAT~Control | VMAT | Control | 18.45550093 | 0.055482174 | 17.64202569 | 0.098767679 | 1.757439766 | 0.813475244 | 0.774249985 | 1.81E-08 | 7.741259475 | Unpaired t-test | 0.029831569 | permutation FDR (250 perm) | 1 | 1.21E-07 FDR correction BH |

| | | | | | | | | | | | | | | | | | | |
|---|---|---|---|---|---|---|---|---|---|---|---|---|---|---|---|---|---|---|
| mfas~D3DMM4 | VMAT~Control | VMAT | Control | 14.41718981 | 0.567270711 | 13.60418274 | 0.144276531 | 1.756869547 | 0.813007071 | 0.644339713 | 0.011139431 | 1.953137002 | Unpaired t-test | 0.057803993 | permutation FDR (250 perm) | 1 | 0.015263225 | FDR correction BH |
| Epp~A8JR97 | VMAT~Control | VMAT | Control | 16.19947869 | 0.197658007 | 15.3865956 | 0.463062867 | 1.756718574 | 0.812883091 | 0.663488781 | 0.004766763 | 2.321776448 | Unpaired t-test | 0.051852794 | permutation FDR (250 perm) | 1 | 0.006884024 | FDR correction BH |
| UQCR-Q~Q9VVH5 | VMAT~Control | VMAT | Control | 16.12967636 | 0.397178353 | 15.31681792 | 0.09420154 | 1.756688553 | 0.812858436 | 0.68737699 | 0.001229847 | 2.91014907 | Unpaired t-test | 0.045511152 | permutation FDR (250 perm) | 1 | 0.001939088 | FDR correction BH |
| CG16935-RA~H8F4T6 | VMAT~Control | VMAT | Control | 17.74006276 | 0.071591788 | 16.92800298 | 0.130057589 | 1.755716348 | 0.812059783 | 0.761501195 | 2.44E-07 | 6.612384879 | Unpaired t-test | 0.031661765 | permutation FDR (250 perm) | 1 | 1.02E-06 | FDR correction BH |
| Dmel\CG6028~Q95SI7 | VMAT~Control | VMAT | Control | 16.60341537 | 0.178221041 | 15.791595 | 0.202956425 | 1.755425009 | 0.811820366 | 0.724327135 | 5.23E-05 | 4.281672508 | Unpaired t-test | 0.037780421 | permutation FDR (250 perm) | 1 | 0.000109978 | FDR correction BH |
| Dmel\CG17097~E2QCS7 | VMAT~Control | VMAT | Control | 17.39357541 | 0.36124535 | 16.58276346 | 0.236410715 | 1.754198428 | 0.810811949 | 0.679598931 | 0.001830195 | 2.737502593 | Unpaired t-test | 0.047512506 | permutation FDR (250 perm) | 1 | 0.002795769 | FDR correction BH |
| ClpX~Q9VDS7 | VMAT~Control | VMAT | Control | 16.21595402 | 0.236444754 | 15.40557547 | 0.127091995 | 1.753671525 | 0.810378546 | 0.723520789 | 5.04E-05 | 4.297603323 | Unpaired t-test | 0.037937786 | permutation FDR (250 perm) | 1 | 0.000106397 | FDR correction BH |
| Sec20~Q9VNI9 | VMAT~Control | VMAT | Control | 20.42253972 | 0.052629939 | 19.6131243 | 0.061240203 | 1.752501182 | 0.809415417 | 0.781204787 | 7.02E-10 | 9.153371134 | Unpaired t-test | 0.028830571 | permutation FDR (250 perm) | 1 | 9.37E-09 | FDR correction BH |
| Cyp9f2~Q9VG82 | VMAT~Control | VMAT | Control | 16.21499231 | 0.05922058 | 15.40601331 | 0.210900142 | 1.751971118 | 0.808978992 | 0.736798359 | 8.91E-06 | 5.05016177 | Unpaired t-test | 0.03563703 | permutation FDR (250 perm) | 1 | 2.28E-05 | FDR correction BH |
| mAcon1~Q9VIE8 | VMAT~Control | VMAT | Control | 21.8381514 | 0.04782499 | 21.02980789 | 0.148734517 | 1.751199584 | 0.808343517 | 0.755552924 | 4.12E-07 | 6.385427098 | Unpaired t-test | 0.032501147 | permutation FDR (250 perm) | 1 | 1.58E-06 | FDR correction BH |
| Nep1~Q9W436 | VMAT~Control | VMAT | Control | 15.77419089 | 0.244550692 | 14.96731547 | 0.252890009 | 1.749418453 | 0.806875417 | 0.697188997 | 0.000445041 | 3.351600375 | Unpaired t-test | 0.043280182 | permutation FDR (250 perm) | 1 | 0.000759149 | FDR correction BH |
| fwe~Q95T12 | VMAT~Control | VMAT | Control | 14.46862856 | 1.083778193 | 13.66201474 | 0.437452298 | 1.749101267 | 0.806613819 | 0.529735128 | 0.153801343 | 0.813039872 | Unpaired t-test | 0.106774483 | permutation FDR (250 perm) | 1 | 0.179054581 | FDR correction BH |
| cu~A8JQX3-5 | VMAT~Control | VMAT | Control | 16.74497428 | 0.136583028 | 15.93986322 | 0.342920495 | 1.747280288 | 0.805111055 | 0.691037744 | 0.000644616 | 3.19069924 | Unpaired t-test | 0.044584699 | permutation FDR (250 perm) | 1 | 0.001064113 | FDR correction BH |
| pepCG10602~Q9VJ39 | VMAT~Control | VMAT | Control | 17.32280226 | 0.208691765 | 16.5200045 | 0.216343876 | 1.744480847 | 0.802797758 | 0.707666284 | 0.000137166 | 3.862754259 | Unpaired t-test | 0.041088919 | permutation FDR (250 perm) | 1 | 0.000261025 | FDR correction BH |
| Nrt~M9MS15 | VMAT~Control | VMAT | Control | 15.10983055 | 0.267649616 | 14.31028945 | 0.145789566 | 1.740547388 | 0.799541093 | 0.703634446 | 0.000158157 | 3.800911295 | Unpaired t-test | 0.041899727 | permutation FDR (250 perm) | 1 | 0.000297841 | FDR correction BH |
| Acp76A~M9NEE3 | VMAT~Control | VMAT | Control | 17.21401073 | 0.460080356 | 16.41491453 | 0.231830443 | 1.740010736 | 0.799096208 | 0.649460853 | 0.006038166 | 2.219094969 | Unpaired t-test | 0.056190649 | permutation FDR (250 perm) | 1 | 0.008624904 | FDR correction BH |
| Evi5~Q9VYY9 | VMAT~Control | VMAT | Control | 17.21944671 | 0.095118763 | 16.42276206 | 0.089145708 | 1.737104628 | 0.796684652 | 0.752796504 | 8.53E-08 | 7.068937558 | Unpaired t-test | 0.032945388 | permutation FDR (250 perm) | 1 | 4.17E-07 | FDR correction BH |
| P5CDh1~Q9VNX4 | VMAT~Control | VMAT | Control | 16.54936794 | 0.266348317 | 15.75586207 | 0.606276266 | 1.733281376 | 0.793505876 | 0.612204074 | 0.023114882 | 1.636108324 | Unpaired t-test | 0.068967391 | permutation FDR (250 perm) | 1 | 0.030162102 | FDR correction BH |
| Glut1~Q8IRI6-16 | VMAT~Control | VMAT | Control | 16.34033152 | 0.217965928 | 15.54697637 | 0.173371892 | 1.733100306 | 0.793355155 | 0.705485042 | 8.14E-05 | 4.089183079 | Unpaired t-test | 0.041474932 | permutation FDR (250 perm) | 1 | 0.000163011 | FDR correction BH |
| Arpc1~O97182 | VMAT~Control | VMAT | Control | 15.51223035 | 0.474917614 | 14.71925715 | 0.267489794 | 1.732641527 | 0.7929732 | 0.637560669 | 0.008675226 | 2.061719201 | Unpaired t-test | 0.059989116 | permutation FDR (250 perm) | 1 | 0.012148413 | FDR correction BH |
| MagR~Q8T3X9 | VMAT~Control | VMAT | Control | 16.84086187 | 0.228080296 | 16.04828463 | 0.165232425 | 1.732166056 | 0.792577242 | 0.703916022 | 8.99E-05 | 4.046029143 | Unpaired t-test | 0.041843209 | permutation FDR (250 perm) | 1 | 0.000178372 | FDR correction BH |
| l(2)k09913~A0A0B4LGC6 | VMAT~Control | VMAT | Control | 17.25444736 | 0.112070539 | 16.46250023 | 0.101216072 | 1.731409673 | 0.791947124 | 0.741846913 | 3.63E-07 | 6.440398156 | Unpaired t-test | 0.034760899 | permutation FDR (250 perm) | 1 | 1.41E-06 | FDR correction BH |
| Dmel\CG9062~A0A0B4KET0 | VMAT~Control | VMAT | Control | 15.63933634 | 0.136870605 | 14.84753936 | 0.118554934 | 1.73122949 | 0.79179698 | 0.732480666 | 1.95E-06 | 5.709398426 | Unpaired t-test | 0.036344165 | permutation FDR (250 perm) | 1 | 6.07E-06 | FDR correction BH |
| Cyp1~P25007 | VMAT~Control | VMAT | Control | 17.51875667 | 0.058672155 | 16.72704419 | 0.252473059 | 1.73112809 | 0.791712477 | 0.709471828 | 4.57E-05 | 4.340131401 | Unpaired t-test | 0.040744865 | permutation FDR (250 perm) | 1 | 9.72E-05 | FDR correction BH |
| Tina-1~Q2PE12 | VMAT~Control | VMAT | Control | 14.34963505 | 1.104621855 | 13.5607395 | 0.266029412 | 1.72775128 | 0.788895548 | 0.523096513 | 0.151577231 | 0.81936603 | Unpaired t-test | 0.110409336 | permutation FDR (250 perm) | 1 | 0.176815066 | FDR correction BH |
| scu~O18404 | VMAT~Control | VMAT | Control | 18.55393338 | 0.136614878 | 17.76542969 | 0.247579987 | 1.727282068 | 0.788503697 | 0.699984333 | 9.69E-05 | 4.013700946 | Unpaired t-test | 0.042723792 | permutation FDR (250 perm) | 1 | 0.00019119 | FDR correction BH |
| ND-24~Q9VX36 | VMAT~Control | VMAT | Control | 17.83393014 | 0.346642116 | 17.04571221 | 0.463397916 | 1.726939965 | 0.78821793 | 0.626163985 | 0.012343758 | 1.908552597 | Unpaired t-test | 0.063956482 | permutation FDR (250 perm) | 1 | 0.016815513 | FDR correction BH |
| lqf~M9PEM7 | VMAT~Control | VMAT | Control | 19.20056417 | 0.155550729 | 18.41288088 | 0.261735928 | 1.726300109 | 0.787683292 | 0.693283742 | 0.000176562 | 3.753103616 | Unpaired t-test | 0.044153005 | permutation FDR (250 perm) | 1 | 0.000327781 | FDR correction BH |
| SdhC~Q9VGS3 | VMAT~Control | VMAT | Control | 16.25793835 | 0.211019019 | 15.47196907 | 0.293440678 | 1.724250369 | 0.785969275 | 0.6766035 | 0.000658962 | 3.181139558 | Unpaired t-test | 0.048372897 | permutation FDR (250 perm) | 1 | 0.001083995 | FDR correction BH |
| Mpcp1~Q7JUS9 | VMAT~Control | VMAT | Control | 20.81892156 | 0.091756903 | 20.0332545 | 0.123406921 | 1.723889205 | 0.785667055 | 0.735111279 | 4.63E-07 | 6.334131307 | Unpaired t-test | 0.035952337 | permutation FDR (250 perm) | 1 | 1.74E-06 | FDR correction BH |
| BVR~Q9VH72 | VMAT~Control | VMAT | Control | 17.0632883 | 0.114435798 | 16.27827183 | 0.082199839 | 1.72311199 | 0.78501647 | 0.738483384 | 2.05E-07 | 6.687808208 | Unpaired t-test | 0.035347088 | permutation FDR (250 perm) | 1 | 8.88E-07 | FDR correction BH |
| bs33f06.y1~A1Z9T6 | VMAT~Control | VMAT | Control | 15.06097769 | 0.804073123 | 14.27721207 | 0.41058545 | 1.721618657 | 0.783765618 | 0.558332735 | 0.080924882 | 1.091917928 | Unpaired t-test | 0.091857722 | permutation FDR (250 perm) | 1 | 0.098042806 | FDR correction BH |
| sp3~E1JIS1 | VMAT~Control | VMAT | Control | 18.15006839 | 0.062342284 | 17.36639775 | 0.125860978 | 1.721505324 | 0.783670643 | 0.737354953 | 2.02E-07 | 6.693706009 | Unpaired t-test | 0.035517874 | permutation FDR (250 perm) | 1 | 8.79E-07 | FDR correction BH |
| bsf~Q9VJ86 | VMAT~Control | VMAT | Control | 14.93336016 | 0.411358087 | 14.15030495 | 0.076995332 | 1.720771107 | 0.783055206 | 0.659603935 | 0.001876083 | 2.726747986 | Unpaired t-test | 0.053014085 | permutation FDR (250 perm) | 1 | 0.002860297 | FDR correction BH |
| Vps16B~Q9VAG4 | VMAT~Control | VMAT | Control | 14.19056819 | 0.739504535 | 13.40770301 | 0.35807589 | 1.720544468 | 0.78286518 | 0.572501431 | 0.058966158 | 1.229397171 | Unpaired t-test | 0.08518018 | permutation FDR (250 perm) | 1 | 0.072902487 | FDR correction BH |
| CG9798~Q95T98 | VMAT~Control | VMAT | Control | 17.84676517 | 0.073379481 | 17.06443051 | 0.27510148 | 1.719911892 | 0.78233466 | 0.693970943 | 0.000109162 | 3.961929542 | Unpaired t-test | 0.043985428 | permutation FDR (250 perm) | 1 | 0.000212703 | FDR correction BH |
| RpS24~Q9W229 | VMAT~Control | VMAT | Control | 18.08244213 | 0.115922919 | 17.30079929 | 0.205660146 | 1.719087335 | 0.78164284 | 0.706998878 | 2.31E-05 | 4.637145479 | Unpaired t-test | 0.041197812 | permutation FDR (250 perm) | 1 | 5.28E-05 | FDR correction BH |
| AsnRS~Q9V434 | VMAT~Control | VMAT | Control | 17.36188474 | 0.147040439 | 16.58030032 | 0.295403022 | 1.719017717 | 0.781584414 | 0.681078045 | 0.000349348 | 3.456741191 | Unpaired t-test | 0.047157416 | permutation FDR (250 perm) | 1 | 0.000610064 | FDR correction BH |
| Rpn7~Q9V3G7 | VMAT~Control | VMAT | Control | 17.4115544 | 0.172164058 | 16.63111686 | 0.181557575 | 1.71765172 | 0.780437538 | 0.70189794 | 3.83E-05 | 4.416685417 | Unpaired t-test | 0.042309936 | permutation FDR (250 perm) | 1 | 8.33E-05 | FDR correction BH |
| anon-EST:fe1H12~A1ZBH1 | VMAT~Control | VMAT | Control | 16.84574948 | 0.196465009 | 16.0653227 | 0.17045474 | 1.717638911 | 0.78042678 | 0.699105825 | 5.30E-05 | 4.275443308 | Unpaired t-test | 0.042909754 | permutation FDR (250 perm) | 1 | 0.000111367 | FDR correction BH |
| Dmel\CG13850~Q9VD14 | VMAT~Control | VMAT | Control | 17.61853805 | 0.065723831 | 16.83943048 | 0.12262741 | 1.716069002 | 0.779107564 | 0.733470486 | 1.96E-07 | 6.708635813 | Unpaired t-test | 0.036201373 | permutation FDR (250 perm) | 1 | 8.53E-07 | FDR correction BH |
| Gga~Q9W329 | VMAT~Control | VMAT | Control | 16.69166585 | 0.110607563 | 15.91332292 | 0.277479755 | 1.715159713 | 0.778342925 | 0.686618755 | 0.000166855 | 3.777660474 | Unpaired t-test | 0.045694268 | permutation FDR (250 perm) | 1 | 0.000312223 | FDR correction BH |
| BBS9~B7YZM8 | VMAT~Control | VMAT | Control | 14.31053348 | 0.265062143 | 13.53266435 | 0.31649938 | 1.714596533 | 0.777869131 | 0.656638091 | 0.00179063 | 2.74699418 | Unpaired t-test | 0.053887375 | permutation FDR (250 perm) | 1 | 0.002746021 | FDR correction BH |
| whd~Q7JQH9 | VMAT~Control | VMAT | Control | 16.8975035 | 0.052370397 | 16.11972815 | 0.093076199 | 1.714485077 | 0.777775347 | 0.742320935 | 1.58E-08 | 7.800352032 | Unpaired t-test | 0.034652593 | permutation FDR (250 perm) | 1 | 1.07E-07 | FDR correction BH |
| Myo31DF~Q23978 | VMAT~Control | VMAT | Control | 16.12090275 | 0.056842219 | 15.34318928 | 0.139741234 | 1.714411541 | 0.777713467 | 0.728560062 | 4.26E-07 | 6.370704303 | Unpaired t-test | 0.037021043 | permutation FDR (250 perm) | 1 | 1.62E-06 | FDR correction BH |
| sau~Q9VQ93 | VMAT~Control | VMAT | Control | 15.88844354 | 0.369335748 | 15.11120911 | 0.647580787 | 1.713842375 | 0.777234428 | 0.582897729 | 0.041960455 | 1.377159815 | Unpaired t-test | 0.080571171 | permutation FDR (250 perm) | 1 | 0.052849069 | FDR correction BH |
| Dmel\CG9646~A1ZAP1 | VMAT~Control | VMAT | Control | 15.8673372 | 0.171502097 | 15.09218543 | 0.379063788 | 1.711370081 | 0.775151774 | 0.653548735 | 0.001930181 | 2.714402028 | Unpaired t-test | 0.05479564 | permutation FDR (250 perm) | 1 | 0.002937069 | FDR correction BH |
| ATP7~Q9VYT4 | VMAT~Control | VMAT | Control | 17.18647694 | 0.173792922 | 16.41250643 | 0.178023836 | 1.709969406 | 0.773970513 | 0.69647871 | 3.92E-05 | 4.407016728 | Unpaired t-test | 0.04342779 | permutation FDR (250 perm) | 1 | 8.48E-05 | FDR correction BH |
| PyK~O62619-2 | VMAT~Control | VMAT | Control | 21.12787182 | 0.09218474 | 20.35393855 | 0.154984173 | 1.709925261 | 0.773933268 | 0.716177128 | 2.31E-06 | 5.63546674 | Unpaired t-test | 0.039327854 | permutation FDR (250 perm) | 1 | 7.00E-06 | FDR correction BH |
| Idh3g~A8JRB8 | VMAT~Control | VMAT | Control | 20.96168334 | 0.053109692 | 20.18806476 | 0.138183267 | 1.70955233 | 0.773618585 | 0.725581727 | 3.75E-07 | 6.42603142 | Unpaired t-test | 0.037522415 | permutation FDR (250 perm) | 1 | 1.45E-06 | FDR correction BH |
| Dmel\CG8187~Q7K1W5 | VMAT~Control | VMAT | Control | 16.13450504 | 0.155234402 | 15.36223571 | 0.15381452 | 1.707954242 | 0.772269324 | 0.703514255 | 1.31E-05 | 4.882225176 | Unpaired t-test | 0.041921604 | permutation FDR (250 perm) | 1 | 3.19E-05 | FDR correction BH |
| MKP-like~X2JEB8 | VMAT~Control | VMAT | Control | 16.75167602 | 0.165185609 | 15.98070971 | 0.267761473 | 1.706412351 | 0.770966313 | 0.675871213 | 0.000269432 | 3.569551433 | Unpaired t-test | 0.048529332 | permutation FDR (250 perm) | 1 | 0.000486382 | FDR correction BH |
| Arf4~L0MNA8 | VMAT~Control | VMAT | Control | 16.79447572 | 0.092317591 | 16.02436545 | 0.050003319 | 1.705400124 | 0.770110267 | 0.735572989 | 1.48E-08 | 7.830470985 | Unpaired t-test | 0.035866178 | permutation FDR (250 perm) | 1 | 1.02E-07 | FDR correction BH |
| ND-30~Q9VZU4 | VMAT~Control | VMAT | Control | 17.71235768 | 0.321675846 | 16.94229644 | 0.326277381 | 1.705342174 | 0.770061243 | 0.639104846 | 0.003733666 | 2.427864479 | Unpaired t-test | 0.059479129 | permutation FDR (250 perm) | 1 | 0.005465828 | FDR correction BH |
| CG32919~Q9VG87 | VMAT~Control | VMAT | Control | 16.82199555 | 0.244246345 | 16.05526326 | 0.179293521 | 1.701411715 | 0.766732293 | 0.675237075 | 0.000209973 | 3.677835827 | Unpaired t-test | 0.048674852 | permutation FDR (250 perm) | 1 | 0.000385251 | FDR correction BH |
| Cg17746~Q9VZS1 | VMAT~Control | VMAT | Control | 17.27537192 | 0.074156246 | 16.5088784 | 0.114004774 | 1.701130142 | 0.766493516 | 0.722547158 | 1.84E-07 | 6.735006385 | Unpaired t-test | 0.038120769 | permutation FDR (250 perm) | 1 | 8.10E-07 | FDR correction BH |
| Cbp53E~A0A0B4LFM0 | VMAT~Control | VMAT | Control | 18.21656741 | 0.185910752 | 17.45010809 | 0.220213458 | 1.701089831 | 0.766459329 | 0.678952505 | 0.00014184 | 3.848200506 | Unpaired t-test | 0.047694407 | permutation FDR (250 perm) | 1 | 0.000269268 | FDR correction BH |
| Gbeta76C~P29829 | VMAT~Control | VMAT | Control | 18.02363343 | 0.216176017 | 17.25784623 | 0.233743279 | 1.700297499 | 0.765787195 | 0.670340504 | 0.000310854 | 3.507444086 | Unpaired t-test | 0.049996364 | permutation FDR (250 perm) | 1 | 0.000550602 | FDR correction BH |
| RpL3~O16797 | VMAT~Control | VMAT | Control | 20.83121997 | 0.114981583 | 20.06647015 | 0.137870363 | 1.699075337 | 0.764749823 | 0.707914391 | 2.48E-06 | 5.605955068 | Unpaired t-test | 0.041037848 | permutation FDR (250 perm) | 1 | 7.41E-06 | FDR correction BH |
| Trap1~A1Z6L9 | VMAT~Control | VMAT | Control | 16.27994755 | 0.124053347 | 15.51523355 | 0.0772374 | 1.699033148 | 0.764714 | 0.717803626 | 3.70E-07 | 6.431629626 | Unpaired t-test | 0.039030137 | permutation FDR (250 perm) | 1 | 1.44E-06 | FDR correction BH |
| Dmel\CG9297~Q9VFR2 | VMAT~Control | VMAT | Control | 17.83319175 | 0.091187396 | 17.07047097 | 0.159373624 | 1.696687398 | 0.762720783 | 0.704842054 | 3.11E-06 | 5.506729525 | Unpaired t-test | 0.041620784 | permutation FDR (250 perm) | 1 | 9.09E-06 | FDR correction BH |
| EMC1~A0A0B4LGV1 | VMAT~Control | VMAT | Control | 15.29158949 | 0.208674947 | 14.53044206 | 0.163849898 | 1.694838057 | 0.76114743 | 0.680414572 | 7.68E-05 | 4.114398771 | Unpaired t-test | 0.047279673 | permutation FDR (250 perm) | 1 | 0.000155002 | FDR correction BH |
| Fatp1~Q9VKU1 | VMAT~Control | VMAT | Control | 16.21028 | 0.134460234 | 15.4496665 | 0.215006167 | 1.694210923 | 0.760613496 | 0.683139779 | 5.32E-05 | 4.274165561 | Unpaired t-test | 0.046642402 | permutation FDR (250 perm) | 1 | 0.000111496 | FDR correction BH |
| RpL30~Q9VJ19 | VMAT~Control | VMAT | Control | 18.25424627 | 0.195947515 | 17.49448817 | 0.275361909 | 1.693206705 | 0.759758107 | 0.660003777 | 0.000516562 | 3.286877218 | Unpaired t-test | 0.052939573 | permutation FDR (250 perm) | 1 | 0.000868563 | FDR correction BH |
| emb~A0A0S0WNN6 | VMAT~Control | VMAT | Control | 15.11214451 | 0.352649575 | 14.35330916 | 0.074734317 | 1.692124071 | 0.758835355 | 0.653485491 | 0.000832607 | 3.079559964 | Unpaired t-test | 0.054817439 | permutation FDR (250 perm) | 1 | 0.001349832 | FDR correction BH |
| Had1~Q9VXI1 | VMAT~Control | VMAT | Control | 18.46579849 | 0.152177947 | 17.70843477 | 0.163421409 | 1.690398891 | 0.757363726 | 0.688597243 | 1.87E-05 | 4.727224763 | Unpaired t-test | 0.045187073 | permutation FDR (250 perm) | 1 | 4.37E-05 | FDR correction BH |
| Nipsnap~M9PJP0 | VMAT~Control | VMAT | Control | 17.29754218 | 0.058183329 | 16.54052024 | 0.065701494 | 1.689998477 | 0.757021947 | 0.728432504 | 3.06E-09 | 8.514625835 | Unpaired t-test | 0.037033852 | permutation FDR (250 perm) | 1 | 2.99E-08 | FDR correction BH |
| Prosalpha4~P22769 | VMAT~Control | VMAT | Control | 14.23863782 | 0.694021835 | 13.4838864 | 0.504625558 | 1.687340839 | 0.754751425 | 0.545439875 | 0.077557124 | 1.110378306 | Unpaired t-test | 0.098612061 | permutation FDR (250 perm) | 1 | 0.094107974 | FDR correction BH |
| RpS14b~X2JCX8 | VMAT~Control | VMAT | Control | 18.69121057 | 0.180093721 | 17.93746954 | 0.140259216 | 1.686159514 | 0.753741025 | 0.683922976 | 2.36E-05 | 4.627101 | Unpaired t-test | 0.0464495 | permutation FDR (250 perm) | 1 | 5.39E-05 | FDR correction BH |
| Rab7~O76742 | VMAT~Control | VMAT | Control | 17.69354341 | 0.096719795 | 16.94082922 | 0.106047798 | 1.684959821 | 0.75271419 | 0.707312802 | 3.63E-07 | 6.440412879 | Unpaired t-test | 0.041138167 | permutation FDR (250 perm) | 1 | 1.41E-06 | FDR correction BH |
| RFeSP~Q9VQ29 | VMAT~Control | VMAT | Control | 19.3639097 | 0.222913838 | 18.61247966 | 0.228764695 | 1.683460696 | 0.751430039 | 0.657508111 | 0.000367822 | 3.434361877 | Unpaired t-test | 0.053640327 | permutation FDR (250 perm) | 1 | 0.000637595 | FDR correction BH |
| fidipidine~A1ZAC7 | VMAT~Control | VMAT | Control | 18.63169574 | 0.129224056 | 17.88050391 | 0.138503138 | 1.683182757 | 0.75119183 | 0.692525497 | 4.73E-06 | 5.325344035 | Unpaired t-test | 0.044320583 | permutation FDR (250 perm) | 1 | 1.32E-05 | FDR correction BH |
| CG17896~Q7KW39-2 | VMAT~Control | VMAT | Control | 17.53298232 | 0.11151229 | 16.78181134 | 0.210449389 | 1.683158425 | 0.751170975 | 0.678863885 | 3.49E-05 | 4.457379578 | Unpaired t-test | 0.04772533 | permutation FDR (250 perm) | 1 | 7.67E-05 | FDR correction BH |
| Apt1~Q9I7R0 | VMAT~Control | VMAT | Control | 17.35772423 | 0.084663107 | 16.60801318 | 0.103456854 | 1.681456035 | 0.749711057 | 0.707418145 | 1.93E-07 | 6.714686986 | Unpaired t-test | 0.041123575 | permutation FDR (250 perm) | 1 | 8.42E-07 | FDR correction BH |
| Arl1~P25160 | VMAT~Control | VMAT | Control | 15.33795379 | 0.156928984 | 14.58944173 | 0.174070156 | 1.680059186 | 0.748512058 | 0.677502126 | 3.14E-05 | 4.503582789 | Unpaired t-test | 0.048092769 | permutation FDR (250 perm) | 1 | 6.97E-05 | FDR correction BH |
| alpha-KGDHC~Q9VGQ1 | VMAT~Control | VMAT | Control | 16.15813417 | 0.388917707 | 15.4101501 | 0.430049845 | 1.679444438 | 0.747984067 | 0.593964965 | 0.01625818 | 1.788928069 | Unpaired t-test | 0.076081301 | permutation FDR (250 perm) | 1 | 0.021653184 | FDR correction BH |
| Dmel\CG1635~Q95TK5 | VMAT~Control | VMAT | Control | 16.95150479 | 0.186304481 | 16.20521763 | 0.18770908 | 1.677420721 | 0.746287155 | 0.667356078 | 8.80E-05 | 4.055702401 | Unpaired t-test | 0.050854545 | permutation FDR (250 perm) | 1 | 0.000175329 | FDR correction BH |
| Sec63~Q9VS57 | VMAT~Control | VMAT | Control | 18.09586001 | 0.113221931 | 17.35246386 | 0.072066101 | 1.674112115 | 0.743396148 | 0.701303088 | 2.17E-07 | 6.663806707 | Unpaired t-test | 0.042450319 | permutation FDR (250 perm) | 1 | 9.30E-07 | FDR correction BH |
| Dmel\CG8507~Q9VH64 | VMAT~Control | VMAT | Control | 16.39189015 | 0.492812196 | 15.64898979 | 1.033355062 | 1.673536905 | 0.742900366 | 0.491338344 | 0.177416421 | 0.751006187 | Unpaired t-test | 0.128484145 | permutation FDR (250 perm) | 1 | 0.204524121 | FDR correction BH |
| flr~Q9VU68-2 | VMAT~Control | VMAT | Control | 16.89650869 | 0.309765016 | 16.15392477 | 0.305308962 | 1.673169858 | 0.742583914 | 0.621664783 | 0.003386032 | 2.470308965 | Unpaired t-test | 0.065551427 | permutation FDR (250 perm) | 1 | 0.004981699 | FDR correction BH |
| Atpalpha~P13607-4 | VMAT~Control | VMAT | Control | 20.20211146 | 0.031514348 | 19.46060306 | 0.096180169 | 1.671922993 | 0.7415084 | 0.709398754 | 1.49E-08 | 7.825445947 | Unpaired t-test | 0.040740876 | permutation FDR (250 perm) | 1 | 1.03E-07 | FDR correction BH |
| Sply~Q9V7Y2 | VMAT~Control | VMAT | Control | 16.75700463 | 0.188554051 | 16.01668389 | 0.143530542 | 1.670547192 | 0.740320739 | 0.669382805 | 3.78E-05 | 4.422526872 | Unpaired t-test | 0.050258182 | permutation FDR (250 perm) | 1 | 8.23E-05 | FDR correction BH |
| Fit1~Q9VZI3 | VMAT~Control | VMAT | Control | 17.44664864 | 0.068267341 | 16.7070284 | 0.078242167 | 1.669736261 | 0.739620243 | 0.706798211 | 1.96E-08 | 7.7072649 | Unpaired t-test | 0.041236098 | permutation FDR (250 perm) | 1 | 1.27E-07 | FDR correction BH |
| Dmel\CG10237~Q9I7M3 | VMAT~Control | VMAT | Control | 16.1399992 | 0.107629393 | 15.40398426 | 0.188480031 | 1.665568783 | 0.736014934 | 0.67089409 | 1.88E-05 | 4.726599312 | Unpaired t-test | 0.049849091 | permutation FDR (250 perm) | 1 | 4.37E-05 | FDR correction BH |
| RpL10~M9PIM0 | VMAT~Control | VMAT | Control | 19.71276123 | 0.190565001 | 18.97680741 | 0.215904206 | 1.665498224 | 0.735953816 | 0.651986802 | 0.000194388 | 3.711329545 | Unpaired t-test | 0.055355132 | permutation FDR (250 perm) | 1 | 0.000358894 | FDR correction BH |
| CCT3~A4V303 | VMAT~Control | VMAT | Control | 18.93694412 | 0.139454561 | 18.20139898 | 0.148926855 | 1.665026499 | 0.735545138 | 0.674043154 | 1.10E-05 | 4.958130215 | Unpaired t-test | 0.048960437 | permutation FDR (250 perm) | 1 | 2.73E-05 | FDR correction BH |
| Dmel\CG9175~Q9VMH4 | VMAT~Control | VMAT | Control | 15.21897351 | 0.312219674 | 14.48673216 | 0.125867637 | 1.66121793 | 0.732241349 | 0.636428231 | 0.000657637 | 3.182013654 | Unpaired t-test | 0.060386395 | permutation FDR (250 perm) | 1 | 0.001082572 | FDR correction BH |
| Echs1~Q7JR58 | VMAT~Control | VMAT | Control | 15.82726938 | 0.142491595 | 15.09537596 | 0.283659789 | 1.660817343 | 0.731893414 | 0.640908459 | 0.000427989 | 3.368567826 | Unpaired t-test | 0.058898367 | permutation FDR (250 perm) | 1 | 0.000731652 | FDR correction BH |
| CG9951~Q9VVB4 | VMAT~Control | VMAT | Control | 17.87216112 | 0.102873278 | 17.14062876 | 0.089170732 | 1.660401752 | 0.73153236 | 0.689549784 | 2.89E-07 | 6.539569542 | Unpaired t-test | 0.044969035 | permutation FDR (250 perm) | 1 | 1.16E-06 | FDR correction BH |
| Bem46~O76462 | VMAT~Control | VMAT | Control | 18.02887064 | 0.091550429 | 17.30161437 | 0.10570398 | 1.655487685 | 0.727256279 | 0.684452219 | 3.92E-07 | 6.406417839 | Unpaired t-test | 0.046282075 | permutation FDR (250 perm) | 1 | 1.51E-06 | FDR correction BH |
| Dmel\CG13887~Q9W0M4 | VMAT~Control | VMAT | Control | 15.52914123 | 0.291361365 | 14.80206188 | 0.226389294 | 1.655284678 | 0.727079355 | 0.624096426 | 0.001322143 | 2.878721549 | Unpaired t-test | 0.06469778 | permutation FDR (250 perm) | 1 | 0.00206799 | FDR correction BH |
| slgA~A4V4U5 | VMAT~Control | VMAT | Control | 18.79611322 | 0.04671279 | 18.06956386 | 0.110240792 | 1.654676696 | 0.726549359 | 0.689623737 | 9.13E-08 | 7.039704353 | Unpaired t-test | 0.04495082 | permutation FDR (250 perm) | 1 | 4.41E-07 | FDR correction BH |
| Dmel\CG13332~A1Z9E9 | VMAT~Control | VMAT | Control | 16.54248678 | 0.272394441 | 15.8174991 | 0.212476797 | 1.652886518 | 0.724987677 | 0.627968963 | 0.000851073 | 3.070033078 | Unpaired t-test | 0.063309157 | permutation FDR (250 perm) | 1 | 0.00137598 | FDR correction BH |
| ScsbetaA~A0A0B4JCW4 | VMAT~Control | VMAT | Control | 20.8540717 | 0.15100317 | 20.13091372 | 0.19431429 | 1.650791573 | 0.723157979 | 0.65146167 | 6.30E-05 | 4.20034295 | Unpaired t-test | 0.055529519 | permutation FDR (250 perm) | 1 | 0.000129616 | FDR correction BH |
| Gabat~Q9VW68 | VMAT~Control | VMAT | Control | 17.15407463 | 0.122917301 | 16.43233949 | 0.214768408 | 1.64916431 | 0.721735145 | 0.649822311 | 6.71E-05 | 4.173418449 | Unpaired t-test | 0.056045393 | permutation FDR (250 perm) | 1 | 0.00013683 | FDR correction BH |
| MP1~A0A126GUP6-2 | VMAT~Control | VMAT | Control | 16.24178102 | 0.119247896 | 15.52103826 | 0.135133705 | 1.648030302 | 0.72074277 | 0.666984345 | 4.38E-06 | 5.358107086 | Unpaired t-test | 0.050963636 | permutation FDR (250 perm) | 1 | 1.23E-05 | FDR correction BH |
| ND-75~A4V449 | VMAT~Control | VMAT | Control | 20.60944476 | 0.154621697 | 19.88878741 | 0.275088504 | 1.64793273 | 0.720657352 | 0.63153226 | 0.000459639 | 3.33758357 | Unpaired t-test | 0.062096102 | permutation FDR (250 perm) | 1 | 0.000780429 | FDR correction BH |
| ninaC~P10676 | VMAT~Control | VMAT | Control | 19.30993743 | 0.043161467 | 18.59338726 | 0.060877997 | 1.643247937 | 0.716550174 | 0.693408517 | 1.07E-09 | 8.970036552 | Unpaired t-test | 0.044136612 | permutation FDR (250 perm) | 1 | 1.27E-08 | FDR correction BH |
| UQCR-C2~Q9VV75 | VMAT~Control | VMAT | Control | 22.79518727 | 0.072077317 | 22.07876903 | 0.147481742 | 1.643097679 | 0.716418248 | 0.667422013 | 1.99E-06 | 5.701808291 | Unpaired t-test | 0.050834545 | permutation FDR (250 perm) | 1 | 6.16E-06 | FDR correction BH |
| FeCh~Q9V9S8 | VMAT~Control | VMAT | Control | 18.09880576 | 0.068921495 | 17.38422465 | 0.102031479 | 1.641006679 | 0.71458111 | 0.677286543 | 1.39E-07 | 6.855544612 | Unpaired t-test | 0.048163711 | permutation FDR (250 perm) | 1 | 6.39E-07 | FDR correction BH |
| Pdhb~Q7K5K3 | VMAT~Control | VMAT | Control | 20.79421816 | 0.109011923 | 20.07993268 | 0.143740439 | 1.640670441 | 0.714285476 | 0.660960341 | 4.79E-06 | 5.319227649 | Unpaired t-test | 0.052663335 | permutation FDR (250 perm) | 1 | 1.33E-05 | FDR correction BH |
| pug~A0A0B4K623 | VMAT~Control | VMAT | Control | 16.76003998 | 0.138632676 | 16.04665933 | 0.135544393 | 1.639641773 | 0.713380651 | 0.656460351 | 9.20E-06 | 5.036024112 | Unpaired t-test | 0.053934605 | permutation FDR (250 perm) | 1 | 2.34E-05 | FDR correction BH |
| Dmel\CG30427~Q9W0Y5 | VMAT~Control | VMAT | Control | 14.20461262 | 0.282316804 | 13.49447401 | 0.398318152 | 1.635961291 | 0.710138612 | 0.582874282 | 0.008684323 | 2.061264027 | Unpaired t-test | 0.080572973 | permutation FDR (250 perm) | 1 | 0.012153922 | FDR correction BH |
| RH61753p~Q8SY69 | VMAT~Control | VMAT | Control | 17.49864031 | 0.132479904 | 16.78931471 | 0.162391954 | 1.635039624 | 0.709325598 | 0.648540963 | 1.91E-05 | 4.719418999 | Unpaired t-test | 0.056524739 | permutation FDR (250 perm) | 1 | 4.43E-05 | FDR correction BH |
| betaTub97EF~Q8MST5 | VMAT~Control | VMAT | Control | 19.10714839 | 0.068404481 | 18.39970866 | 0.06574721 | 1.632903717 | 0.707439726 | 0.678644298 | 1.26E-08 | 7.89934353 | Unpaired t-test | 0.047765348 | permutation FDR (250 perm) | 1 | 8.86E-08 | FDR correction BH |
| cindr~A0A0B4KI34 | VMAT~Control | VMAT | Control | 19.37503785 | 0.157830859 | 18.66878722 | 0.071526193 | 1.631558402 | 0.70625063 | 0.655456689 | 3.70E-06 | 5.432253613 | Unpaired t-test | 0.054212534 | permutation FDR (250 perm) | 1 | 1.06E-05 | FDR correction BH |
| Act57B~P53501 | VMAT~Control | VMAT | Control | 22.23158117 | 0.221273658 | 21.52604273 | 0.237155986 | 1.630753179 | 0.705538441 | 0.616161096 | 0.00065754 | 3.182077536 | Unpaired t-test | 0.067514493 | permutation FDR (250 perm) | 1 | 0.001082572 | FDR correction BH |
| betaTub60D~A0A0B4LGH1 | VMAT~Control | VMAT | Control | 16.63204639 | 0.163633663 | 15.93154467 | 0.194058645 | 1.625069839 | 0.700501721 | 0.629087298 | 0.00010542 | 3.977076273 | Unpaired t-test | 0.06294107 | permutation FDR (250 perm) | 1 | 0.000206627 | FDR correction BH |
| RpL31~Q9V597 | VMAT~Control | VMAT | Control | 18.55709445 | 0.084785589 | 17.85737313 | 0.263803455 | 1.624191029 | 0.699721325 | 0.622572168 | 0.000213467 | 3.670668536 | Unpaired t-test | 0.065217943 | permutation FDR (250 perm) | 1 | 0.000391052 | FDR correction BH |
| Idh3b~Q9VD58 | VMAT~Control | VMAT | Control | 21.25846584 | 0.046590263 | 20.55890119 | 0.121347008 | 1.624014652 | 0.699564649 | 0.661132704 | 2.84E-07 | 6.546098088 | Unpaired t-test | 0.052614266 | permutation FDR (250 perm) | 1 | 1.14E-06 | FDR correction BH |
| r~X2JFG0 | VMAT~Control | VMAT | Control | 15.55191656 | 0.154957291 | 14.85421874 | 0.130455717 | 1.621914558 | 0.69769782 | 0.63974494 | 1.64E-05 | 4.785283396 | Unpaired t-test | 0.059241379 | permutation FDR (250 perm) | 1 | 3.87E-05 | FDR correction BH |
| PMCA~E6EK18 | VMAT~Control | VMAT | Control | 16.98784287 | 0.037814023 | 16.29024406 | 0.05420054 | 1.621803246 | 0.697598805 | 0.677572889 | 4.23E-10 | 9.374009409 | Unpaired t-test | 0.048069122 | permutation FDR (250 perm) | 1 | 6.34E-09 | FDR correction BH |
| RpS16~A0A0B4LG52 | VMAT~Control | VMAT | Control | 20.66324863 | 0.154064938 | 19.96711339 | 0.1392351 | 1.620158813 | 0.696135237 | 0.636980106 | 2.07E-05 | 4.683526363 | Unpaired t-test | 0.060195918 | permutation FDR (250 perm) | 1 | 4.78E-05 | FDR correction BH |
| levy~Q9W1N3 | VMAT~Control | VMAT | Control | 17.57119004 | 0.287682724 | 16.8763845 | 0.49141951 | 1.61866624 | 0.69480554 | 0.55378065 | 0.021252326 | 1.672593539 | Unpaired t-test | 0.094177398 | permutation FDR (250 perm) | 1 | 0.027948589 | FDR correction BH |
| ND-51~Q9VMI3 | VMAT~Control | VMAT | Control | 19.48859649 | 0.208329218 | 18.79451165 | 0.254380631 | 1.617857837 | 0.694084842 | 0.605107111 | 0.000816152 | 3.088229195 | Unpaired t-test | 0.07169579 | permutation FDR (250 perm) | 1 | 0.001324067 | FDR correction BH |
| CG45076~C0HK92 | VMAT~Control | VMAT | Control | 18.56591879 | 0.18966578 | 17.87205686 | 0.139410308 | 1.617607879 | 0.69386193 | 0.627776237 | 6.15E-05 | 4.211476247 | Unpaired t-test | 0.063356301 | permutation FDR (250 perm) | 1 | 0.000126556 | FDR correction BH |
| Pfas~P35421 | VMAT~Control | VMAT | Control | 16.2420591 | 0.113883238 | 15.54917069 | 0.201296537 | 1.616516704 | 0.692888415 | 0.627940052 | 5.37E-05 | 4.269952605 | Unpaired t-test | 0.063318223 | permutation FDR (250 perm) | 1 | 0.000112483 | FDR correction BH |
| Rap1~M9MRT6 | VMAT~Control | VMAT | Control | 17.78929312 | 0.15910222 | 17.09688515 | 0.15119192 | 1.615978467 | 0.692407974 | 0.630519059 | 3.48E-05 | 4.458283268 | Unpaired t-test | 0.062417044 | permutation FDR (250 perm) | 1 | 7.67E-05 | FDR correction BH |
| Synj~Q5U0V7 | VMAT~Control | VMAT | Control | 16.50383261 | 0.113955061 | 15.81148404 | 0.189574593 | 1.615911931 | 0.692348572 | 0.630027262 | 3.72E-05 | 4.429545866 | Unpaired t-test | 0.062605621 | permutation FDR (250 perm) | 1 | 8.12E-05 | FDR correction BH |
| Arf1~M9PG22 | VMAT~Control | VMAT | Control | 17.79959296 | 0.089237991 | 17.10738974 | 0.103002996 | 1.615749138 | 0.692203222 | 0.652438665 | 4.89E-07 | 6.310364455 | Unpaired t-test | 0.055209809 | permutation FDR (250 perm) | 1 | 1.82E-06 | FDR correction BH |
| Vps13~A1Z713 | VMAT~Control | VMAT | Control | 17.68511527 | 0.096502152 | 16.99437382 | 0.0827832 | 1.61411285 | 0.690741448 | 0.653578486 | 2.60E-07 | 6.584490507 | Unpaired t-test | 0.054784741 | permutation FDR (250 perm) | 1 | 1.07E-06 | FDR correction BH |
| RpS15Aa~E1JJM9 | VMAT~Control | VMAT | Control | 20.75118933 | 0.131035422 | 20.06057189 | 0.108394698 | 1.613974119 | 0.690617444 | 0.641806632 | 3.82E-06 | 5.418150267 | Unpaired t-test | 0.058578947 | permutation FDR (250 perm) | 1 | 1.09E-05 | FDR correction BH |
| Cyp4g1~Q9V3S0 | VMAT~Control | VMAT | Control | 18.19749576 | 0.11008816 | 17.50693809 | 0.163769945 | 1.613907246 | 0.690557667 | 0.634558138 | 1.43E-05 | 4.845158833 | Unpaired t-test | 0.061014059 | permutation FDR (250 perm) | 1 | 3.44E-05 | FDR correction BH |
| Dmel\CG14419~Q9W4V3 | VMAT~Control | VMAT | Control | 19.38550933 | 0.196056358 | 18.69713212 | 0.769332298 | 1.61146987 | 0.688377215 | 0.508007899 | 0.081241488 | 1.090222132 | Unpaired t-test | 0.118750783 | permutation FDR (250 perm) | 1 | 0.098375752 | FDR correction BH |
| stck~Q8INR0 | VMAT~Control | VMAT | Control | 16.09200291 | 0.089962313 | 15.40403269 | 0.10194864 | 1.611015331 | 0.687970224 | 0.648535487 | 5.07E-07 | 6.294932108 | Unpaired t-test | 0.056524739 | permutation FDR (250 perm) | 1 | 1.87E-06 | FDR correction BH |
| Acat1~Q9W3N9 | VMAT~Control | VMAT | Control | 17.34617752 | 0.062045298 | 16.65886875 | 0.06643965 | 1.610276867 | 0.687308763 | 0.660458316 | 1.10E-08 | 7.957823111 | Unpaired t-test | 0.052796002 | permutation FDR (250 perm) | 1 | 7.96E-08 | FDR correction BH |
| RpS4~X2JGM9 | VMAT~Control | VMAT | Control | 19.20340271 | 0.146414487 | 18.51674177 | 0.12807321 | 1.609553964 | 0.686660947 | 0.631706177 | 1.32E-05 | 4.877991565 | Unpaired t-test | 0.062036265 | permutation FDR (250 perm) | 1 | 3.21E-05 | FDR correction BH |
| Arr1~P15372 | VMAT~Control | VMAT | Control | 18.75723247 | 0.129282414 | 18.07088 | 0.099326122 | 1.60920985 | 0.686352474 | 0.639710906 | 2.76E-06 | 5.559441564 | Unpaired t-test | 0.059246824 | permutation FDR (250 perm) | 1 | 8.18E-06 | FDR correction BH |
| krz~Q9V393 | VMAT~Control | VMAT | Control | 17.54844889 | 0.054343189 | 16.86451068 | 0.37799615 | 1.606519192 | 0.683938215 | 0.584171601 | 0.002499239 | 2.602192197 | Unpaired t-test | 0.080028842 | permutation FDR (250 perm) | 1 | 0.003752046 | FDR correction BH |
| Pdha~Q7KVX1 | VMAT~Control | VMAT | Control | 20.73916931 | 0.075395309 | 20.05547995 | 0.139803845 | 1.606242096 | 0.683689356 | 0.638344792 | 2.25E-06 | 5.646965379 | Unpaired t-test | 0.059735027 | permutation FDR (250 perm) | 1 | 6.87E-06 | FDR correction BH |
| kcc~Q8MLQ5 | VMAT~Control | VMAT | Control | 18.56565387 | 0.05526304 | 17.8843311 | 0.186441455 | 1.603609389 | 0.681322769 | 0.626812149 | 1.41E-05 | 4.849708212 | Unpaired t-test | 0.063711695 | permutation FDR (250 perm) | 1 | 3.41E-05 | FDR correction BH |
| Pgm1~Q9VUY9 | VMAT~Control | VMAT | Control | 18.73729709 | 0.184670824 | 18.05667842 | 0.301165815 | 1.602826954 | 0.680618676 | 0.587758695 | 0.001541952 | 2.811929175 | Unpaired t-test | 0.078499549 | permutation FDR (250 perm) | 1 | 0.002383293 | FDR correction BH |
| mib2~Q9VJ14 | VMAT~Control | VMAT | Control | 16.10969565 | 0.162754251 | 15.42947041 | 0.311466786 | 1.602389909 | 0.680225241 | 0.587838915 | 0.001493858 | 2.825690588 | Unpaired t-test | 0.078472498 | permutation FDR (250 perm) | 1 | 0.002313515 | FDR correction BH |
| Wdr81~Q9VKD0 | VMAT~Control | VMAT | Control | 13.91188207 | 0.892236571 | 13.23244216 | 0.293226692 | 1.601517889 | 0.679439913 | 0.478473357 | 0.136808528 | 0.863886829 | Unpaired t-test | 0.136178253 | permutation FDR (250 perm) | 1 | 0.160943166 | FDR correction BH |
| Gdh~P54385 | VMAT~Control | VMAT | Control | 19.98712103 | 0.24406021 | 19.3103494 | 0.313447107 | 1.598558599 | 0.676771631 | 0.574675138 | 0.003432713 | 2.464362478 | Unpaired t-test | 0.08416036 | permutation FDR (250 perm) | 1 | 0.005047225 | FDR correction BH |
| Acp~Q9VKS9 | VMAT~Control | VMAT | Control | 16.70490859 | 0.324970946 | 16.02926269 | 0.154163447 | 1.597311734 | 0.675645899 | 0.582024111 | 0.001827745 | 2.73808428 | Unpaired t-test | 0.080893694 | permutation FDR (250 perm) | 1 | 0.002795655 | FDR correction BH |
| igl~Q7JPS2 | VMAT~Control | VMAT | Control | 20.66392589 | 0.126108991 | 19.98865807 | 0.157442992 | 1.596893191 | 0.675267821 | 0.619390822 | 2.10E-05 | 4.67819367 | Unpaired t-test | 0.066342391 | permutation FDR (250 perm) | 1 | 4.83E-05 | FDR correction BH |
| uex~A0A0B7P9G0 | VMAT~Control | VMAT | Control | 18.24351988 | 0.038806757 | 17.56891691 | 0.089769095 | 1.596157451 | 0.674602972 | 0.646334493 | 2.67E-08 | 7.572908387 | Unpaired t-test | 0.057201452 | permutation FDR (250 perm) | 1 | 1.63E-07 | FDR correction BH |
| gish~Q59DW8 | VMAT~Control | VMAT | Control | 18.28266156 | 0.046212603 | 17.60908004 | 0.075381181 | 1.595027747 | 0.673581521 | 0.647959768 | 1.02E-08 | 7.989648419 | Unpaired t-test | 0.056655172 | permutation FDR (250 perm) | 1 | 7.60E-08 | FDR correction BH |
| Cyp312a1~M9PI92 | VMAT~Control | VMAT | Control | 15.17748173 | 0.460776917 | 14.50491348 | 0.206877834 | 1.593907869 | 0.672568242 | 0.548640196 | 0.01386641 | 1.858035947 | Unpaired t-test | 0.096965331 | permutation FDR (250 perm) | 1 | 0.018727212 | FDR correction BH |
| Dmel\CG10132~Q9VIU8 | VMAT~Control | VMAT | Control | 17.94725485 | 0.197026221 | 17.27656934 | 0.319519779 | 1.591829161 | 0.67068551 | 0.574277878 | 0.002538246 | 2.595466227 | Unpaired t-test | 0.084353153 | permutation FDR (250 perm) | 1 | 0.003805753 | FDR correction BH |
| Ldh~B5RJQ0 | VMAT~Control | VMAT | Control | 17.81976828 | 0.097436813 | 17.14923302 | 0.084696058 | 1.591663393 | 0.670535265 | 0.633934295 | 3.97E-07 | 6.40105614 | Unpaired t-test | 0.06124263 | permutation FDR (250 perm) | 1 | 1.53E-06 | FDR correction BH |

| | | | | | | | | | | | | | | | | | | |
|---|---|---|---|---|---|---|---|---|---|---|---|---|---|---|---|---|---|---|
| RpS18~P41094 | VMAT~Control | VMAT | Control | 18.5106954 | 0.123989479 | 17.84019391 | 0.210412388 | 1.591626124 | 0.670501484 | 0.604479302 | 0.000109912 | 3.958953857 | Unpaired t-test | 0.071960163 | permutation FDR (250 perm) | 1 | 0.000213435 | FDR correction BH |
| Idh3a~Q9VWH4 | VMAT~Control | VMAT | Control | 20.83895025 | 0.08479492 | 20.17116125 | 0.156647375 | 1.588636434 | 0.667788996 | 0.618517862 | 7.80E-06 | 5.107926718 | Unpaired t-test | 0.06670471 | permutation FDR (250 perm) | 1 | 2.02E-05 | FDR correction BH |
| for~A0A1W5Q0S1 | VMAT~Control | VMAT | Control | 16.06409449 | 0.045786791 | 15.3963741 | 0.150584549 | 1.588560892 | 0.667720392 | 0.623811724 | 2.57E-06 | 5.589755212 | Unpaired t-test | 0.064790213 | permutation FDR (250 perm) | 1 | 7.67E-06 | FDR correction BH |
| Ran~A4V4A5 | VMAT~Control | VMAT | Control | 19.81127097 | 0.098081953 | 19.14392435 | 0.111800307 | 1.588149382 | 0.667346619 | 0.625728056 | 1.54E-06 | 5.812018959 | Unpaired t-test | 0.064097915 | permutation FDR (250 perm) | 1 | 4.92E-06 | FDR correction BH |
| Dmel\CG4928~X2JFT8 | VMAT~Control | VMAT | Control | 15.98521467 | 0.230565492 | 15.31818453 | 0.07120491 | 1.587801027 | 0.667030135 | 0.602057761 | 0.000104014 | 3.982906885 | Unpaired t-test | 0.072883258 | permutation FDR (250 perm) | 1 | 0.000204041 | FDR correction BH |
| SP191~Q8T429 | VMAT~Control | VMAT | Control | 14.73347802 | 0.403438308 | 14.07131769 | 0.056956267 | 1.582450461 | 0.662160336 | 0.560102763 | 0.004582054 | 2.338939757 | Unpaired t-test | 0.090973435 | permutation FDR (250 perm) | 1 | 0.006629475 | FDR correction BH |
| fau~C0HK95 | VMAT~Control | VMAT | Control | 18.56999239 | 0.177412484 | 17.90803196 | 0.096540178 | 1.582231211 | 0.661960436 | 0.607120793 | 2.52E-05 | 4.599265334 | Unpaired t-test | 0.070908103 | permutation FDR (250 perm) | 1 | 5.71E-05 | FDR correction BH |
| Act88F~P83967 | VMAT~Control | VMAT | Control | 23.09490334 | 0.182992341 | 22.43595404 | 0.231636249 | 1.578932287 | 0.658949302 | 0.582102297 | 0.000544173 | 3.264263269 | Unpaired t-test | 0.080855856 | permutation FDR (250 perm) | 1 | 0.000907854 | FDR correction BH |
| alphaTub84D~P06605 | VMAT~Control | VMAT | Control | 22.23508037 | 0.053219705 | 21.57769448 | 0.065139225 | 1.577222173 | 0.657385898 | 0.633553089 | 7.99E-09 | 8.097328413 | Unpaired t-test | 0.061385941 | permutation FDR (250 perm) | 1 | 6.31E-08 | FDR correction BH |
| Mtpbeta~Q9W1H8 | VMAT~Control | VMAT | Control | 19.06098858 | 0.02841315 | 18.40496639 | 0.035782664 | 1.575732016 | 0.656022197 | 0.642885585 | 2.02E-11 | 10.69380804 | Unpaired t-test | 0.058235935 | permutation FDR (250 perm) | 1 | 6.53E-10 | FDR correction BH |
| AsnRS-m~Q7JZR5 | VMAT~Control | VMAT | Control | 18.6222119 | 0.149115428 | 17.96671957 | 0.208231806 | 1.575153397 | 0.655492333 | 0.588128672 | 0.000192176 | 3.716301123 | Unpaired t-test | 0.078353472 | permutation FDR (250 perm) | 1 | 0.000355088 | FDR correction BH |
| Cand1~M9PB90 | VMAT~Control | VMAT | Control | 16.7288494 | 0.205918797 | 16.0735669 | 0.220545199 | 1.574924314 | 0.655282499 | 0.577372414 | 0.000665243 | 3.177019521 | Unpaired t-test | 0.082904505 | permutation FDR (250 perm) | 1 | 0.001092038 | FDR correction BH |
| rl~P40417 | VMAT~Control | VMAT | Control | 17.34952568 | 0.178716402 | 16.69437375 | 0.266960423 | 1.574781792 | 0.655151937 | 0.572849782 | 0.001042032 | 2.98211886 | Unpaired t-test | 0.084985586 | permutation FDR (250 perm) | 1 | 0.001665271 | FDR correction BH |
| mil~Q9VAZ1 | VMAT~Control | VMAT | Control | 14.44580971 | 0.794214213 | 13.79197535 | 0.325345396 | 1.573344234 | 0.653834354 | 0.472481836 | 0.119306552 | 0.923335707 | Unpaired t-test | 0.139839786 | permutation FDR (250 perm) | 1 | 0.141699103 | FDR correction BH |
| anon-WO0153538.36~Q9VAC1 | VMAT~Control | VMAT | Control | 18.79956276 | 0.108760921 | 18.14977459 | 0.173441909 | 1.568937811 | 0.649788169 | 0.595287027 | 3.31E-05 | 4.480689354 | Unpaired t-test | 0.075533665 | permutation FDR (250 perm) | 1 | 7.30E-05 | FDR correction BH |
| para~X2JFE8 | VMAT~Control | VMAT | Control | 15.87409955 | 0.121991027 | 15.2265422 | 0.44302828 | 1.56651367 | 0.647557359 | 0.537168106 | 0.010313721 | 1.986584628 | Unpaired t-test | 0.102746403 | permutation FDR (250 perm) | 1 | 0.014231242 | FDR correction BH |
| Dmel\CG11857~A0A140SRF8 | VMAT~Control | VMAT | Control | 18.59479937 | 0.072190805 | 17.94727032 | 0.137786078 | 1.566482927 | 0.647529046 | 0.605413402 | 3.05E-06 | 5.515139492 | Unpaired t-test | 0.071561793 | permutation FDR (250 perm) | 1 | 8.95E-06 | FDR correction BH |
| bs29f09.y1~Q86BK2 | VMAT~Control | VMAT | Control | 15.06580403 | 1.8531267 | 14.41869432 | 1.599173351 | 1.566027675 | 0.647109708 | 0.30893263 | 0.567541034 | 0.246002733 | Unpaired t-test | 0.269039052 | permutation FDR (250 perm) | 1 | 0.602612356 | FDR correction BH |
| Cftr~Q9VC63 | VMAT~Control | VMAT | Control | 14.75148664 | 0.225955934 | 14.10505502 | 0.283406271 | 1.565291798 | 0.646431626 | 0.556263641 | 0.002567737 | 2.590449385 | Unpaired t-test | 0.092911301 | permutation FDR (250 perm) | 1 | 0.003842628 | FDR correction BH |
| Pgk~M9PCE0 | VMAT~Control | VMAT | Control | 18.47587347 | 0.100722543 | 17.83039847 | 0.25452856 | 1.564254225 | 0.645475001 | 0.575075843 | 0.000361383 | 3.442032356 | Unpaired t-test | 0.083983784 | permutation FDR (250 perm) | 1 | 0.00062782 | FDR correction BH |
| Sod2~A0A0B4LGQ1 | VMAT~Control | VMAT | Control | 16.31620156 | 0.084007636 | 15.67089939 | 0.355765217 | 1.564066841 | 0.645302169 | 0.554631773 | 0.00274216 | 2.561907198 | Unpaired t-test | 0.093757767 | permutation FDR (250 perm) | 1 | 0.004080306 | FDR correction BH |
| BcDNA:RH09070~Q9W4U2 | VMAT~Control | VMAT | Control | 15.86836915 | 0.12338865 | 15.22321445 | 0.127916387 | 1.563906969 | 0.645154694 | 0.597651817 | 1.04E-05 | 4.984340222 | Unpaired t-test | 0.074632911 | permutation FDR (250 perm) | 1 | 2.59E-05 | FDR correction BH |
| slf~Q8SZK9 | VMAT~Control | VMAT | Control | 18.3194876 | 0.14716005 | 17.67483303 | 0.152080963 | 1.56336492 | 0.644654571 | 0.588918655 | 4.67E-05 | 4.330342854 | Unpaired t-test | 0.078022553 | permutation FDR (250 perm) | 1 | 9.91E-05 | FDR correction BH |
| PCB~Q7KN97 | VMAT~Control | VMAT | Control | 24.9833389 | 0.214583457 | 24.33870692 | 0.301492948 | 1.563340434 | 0.644631975 | 0.55309685 | 0.002984403 | 2.525142463 | Unpaired t-test | 0.09458421 | permutation FDR (250 perm) | 1 | 0.004429562 | FDR correction BH |
| Dmel\CG6178~Q9VCC6 | VMAT~Control | VMAT | Control | 16.77092964 | 0.225285192 | 16.12841554 | 0.146331245 | 1.561047136 | 0.6425141 | 0.573602391 | 0.000324474 | 3.488819922 | Unpaired t-test | 0.084652252 | permutation FDR (250 perm) | 1 | 0.000571288 | FDR correction BH |
| Mcad~Q9VSA3 | VMAT~Control | VMAT | Control | 18.44464389 | 0.3273391 | 17.8022615 | 0.338886554 | 1.56090462 | 0.642382383 | 0.530582893 | 0.012280043 | 1.910800118 | Unpaired t-test | 0.106318059 | permutation FDR (250 perm) | 1 | 0.016738402 | FDR correction BH |
| RpS11~Q0E9B6 | VMAT~Control | VMAT | Control | 18.91192619 | 0.158074611 | 18.27024912 | 0.115365375 | 1.560141704 | 0.641677071 | 0.59003827 | 2.51E-05 | 4.60104574 | Unpaired t-test | 0.077573297 | permutation FDR (250 perm) | 1 | 5.69E-05 | FDR correction BH |
| blw~P35381 | VMAT~Control | VMAT | Control | 22.94406752 | 0.113598768 | 22.30288763 | 0.179887037 | 1.559604138 | 0.641179888 | 0.585474223 | 5.11E-05 | 4.291411793 | Unpaired t-test | 0.079511271 | permutation FDR (250 perm) | 1 | 0.000107828 | FDR correction BH |
| cype~Q9VMS1 | VMAT~Control | VMAT | Control | 18.92523223 | 0.314319812 | 18.2861596 | 0.450183835 | 1.557327782 | 0.639072631 | 0.51308663 | 0.026370536 | 1.578881038 | Unpaired t-test | 0.115711599 | permutation FDR (250 perm) | 1 | 0.034201786 | FDR correction BH |
| Ras64B~P04388 | VMAT~Control | VMAT | Control | 16.83553646 | 0.10440235 | 16.19718214 | 0.114442163 | 1.556552589 | 0.63835432 | 0.596995935 | 3.35E-06 | 5.475372574 | Unpaired t-test | 0.074866184 | permutation FDR (250 perm) | 1 | 9.73E-06 | FDR correction BH |
| RpS6~P29327 | VMAT~Control | VMAT | Control | 19.13453779 | 0.166037833 | 18.49643983 | 0.182331343 | 1.556276018 | 0.638097956 | 0.574715755 | 0.000176304 | 3.753737626 | Unpaired t-test | 0.08414955 | permutation FDR (250 perm) | 1 | 0.000327561 | FDR correction BH |
| RpL12~Q9W1B9 | VMAT~Control | VMAT | Control | 19.6992721 | 0.125585894 | 19.06195632 | 0.104971889 | 1.555432489 | 0.637315779 | 0.59384645 | 5.57E-06 | 5.254197642 | Unpaired t-test | 0.076117435 | permutation FDR (250 perm) | 1 | 1.53E-05 | FDR correction BH |
| apolpp~L0MPS3 | VMAT~Control | VMAT | Control | 19.57647812 | 0.175147843 | 18.93924074 | 0.341065969 | 1.555347962 | 0.637237376 | 0.543965795 | 0.003998857 | 2.398064094 | Unpaired t-test | 0.099351935 | permutation FDR (250 perm) | 1 | 0.005828675 | FDR correction BH |
| GLS~A1Z942 | VMAT~Control | VMAT | Control | 14.5233539 | 0.248162975 | 13.88621936 | 0.077186127 | 1.5552371 | 0.63713454 | 0.570793431 | 0.000268543 | 3.570985411 | Unpaired t-test | 0.085931532 | permutation FDR (250 perm) | 1 | 0.000485151 | FDR correction BH |
| Sgp~Q9VLJ7 | VMAT~Control | VMAT | Control | 17.56494068 | 0.290115406 | 16.92810332 | 0.245114932 | 1.554916779 | 0.636837367 | 0.544374366 | 0.003787741 | 2.421619744 | Unpaired t-test | 0.099148515 | permutation FDR (250 perm) | 1 | 0.005534663 | FDR correction BH |
| Gale~M9ND19 | VMAT~Control | VMAT | Control | 17.73195937 | 0.082350582 | 17.09642546 | 0.121386043 | 1.553512562 | 0.635533907 | 0.596409968 | 2.12E-06 | 5.67295127 | Unpaired t-test | 0.075104882 | permutation FDR (250 perm) | 1 | 6.53E-06 | FDR correction BH |
| GC1~Q9VGF7 | VMAT~Control | VMAT | Control | 16.66837368 | 0.206541955 | 16.03326739 | 0.08036667 | 1.553052171 | 0.635106295 | 0.577834537 | 7.75E-05 | 4.110440203 | Unpaired t-test | 0.082679279 | permutation FDR (250 perm) | 1 | 0.000156154 | FDR correction BH |
| ATPsynB~Q94516 | VMAT~Control | VMAT | Control | 17.25899335 | 0.242171774 | 16.62605856 | 0.40770606 | 1.550716317 | 0.632934788 | 0.522192659 | 0.013701023 | 1.863247012 | Unpaired t-test | 0.110947935 | permutation FDR (250 perm) | 1 | 0.018535751 | FDR correction BH |
| RpS3A~P55830 | VMAT~Control | VMAT | Control | 20.94928802 | 0.132790875 | 20.31855937 | 0.081394119 | 1.548346806 | 0.630728649 | 0.589656685 | 3.92E-06 | 5.407079903 | Unpaired t-test | 0.077690573 | permutation FDR (250 perm) | 1 | 1.12E-05 | FDR correction BH |
| CG4831~Q8IPA8 | VMAT~Control | VMAT | Control | 15.14693668 | 0.319428555 | 14.51635369 | 1.133935342 | 1.548190492 | 0.630582994 | 0.41299658 | 0.258950433 | 0.586783359 | Unpaired t-test | 0.179447788 | permutation FDR (250 perm) | 1 | 0.292359386 | FDR correction BH |
| 10737~Q0E919 | VMAT~Control | VMAT | Control | 18.21974814 | 0.084383575 | 17.59016179 | 0.124569742 | 1.547121335 | 0.629586347 | 0.589893724 | 2.91E-06 | 5.535525449 | Unpaired t-test | 0.077612991 | permutation FDR (250 perm) | 1 | 8.58E-06 | FDR correction BH |
| Uev1A~Q9VRL1 | VMAT~Control | VMAT | Control | 17.50448939 | 0.104614906 | 16.87500633 | 0.255922768 | 1.547010579 | 0.629483063 | 0.560215123 | 0.000470137 | 3.327775421 | Unpaired t-test | 0.090928411 | permutation FDR (250 perm) | 1 | 0.000796189 | FDR correction BH |
| BcDNA:RH27395~A0A0B4K7G1 | VMAT~Control | VMAT | Control | 22.22347512 | 0.047774222 | 21.5941858 | 0.049887011 | 1.546802836 | 0.629289315 | 0.610432789 | 1.79E-09 | 8.74656658 | Unpaired t-test | 0.069635129 | permutation FDR (250 perm) | 1 | 1.95E-08 | FDR correction BH |
| Mlp60A~B7YZP9 | VMAT~Control | VMAT | Control | 21.53828438 | 0.059608424 | 20.91178241 | 0.112087009 | 1.543817232 | 0.626501966 | 0.592843626 | 6.40E-07 | 6.193772582 | Unpaired t-test | 0.07648057 | permutation FDR (250 perm) | 1 | 2.29E-06 | FDR correction BH |
| TrpRS~Q0KI98 | VMAT~Control | VMAT | Control | 16.57647918 | 0.038684427 | 15.95326088 | 0.139066423 | 1.540307398 | 0.623218297 | 0.585426783 | 2.19E-06 | 5.659075478 | Unpaired t-test | 0.079532913 | permutation FDR (250 perm) | 1 | 6.70E-06 | FDR correction BH |
| SrpRalpha~Q9U5L1 | VMAT~Control | VMAT | Control | 18.73995583 | 0.0886215 | 18.11857904 | 0.042188822 | 1.538342544 | 0.621376785 | 0.595248603 | 6.09E-08 | 7.215300247 | Unpaired t-test | 0.075564392 | permutation FDR (250 perm) | 1 | 3.16E-07 | FDR correction BH |
| Cul1~Q24311 | VMAT~Control | VMAT | Control | 15.06948154 | 0.328028931 | 14.44816049 | 0.123398561 | 1.538283113 | 0.621321048 | 0.537133177 | 0.002668531 | 2.573727797 | Unpaired t-test | 0.102760791 | permutation FDR (250 perm) | 1 | 0.003978291 | FDR correction BH |
| Kap-alpha1~O76521 | VMAT~Control | VMAT | Control | 14.63189053 | 0.185869767 | 14.01266636 | 0.434454732 | 1.536048928 | 0.619224171 | 0.511194294 | 0.015036873 | 1.822842468 | Unpaired t-test | 0.116820054 | permutation FDR (250 perm) | 1 | 0.020203652 | FDR correction BH |
| RpL9~X2JDU0 | VMAT~Control | VMAT | Control | 19.47183086 | 0.13390209 | 18.8530915 | 0.135262553 | 1.535532833 | 0.618739361 | 0.570204458 | 2.70E-05 | 4.569104585 | Unpaired t-test | 0.086198198 | permutation FDR (250 perm) | 1 | 6.05E-05 | FDR correction BH |
| Dmel\CG1309~Q9VZF1 | VMAT~Control | VMAT | Control | 15.5273672 | 0.309612104 | 14.90990679 | 0.362294679 | 1.534172181 | 0.617460406 | 0.508982199 | 0.015911407 | 1.798291405 | Unpaired t-test | 0.118207699 | permutation FDR (250 perm) | 1 | 0.021269422 | FDR correction BH |
| alpha-PheRS~Q9W3J5 | VMAT~Control | VMAT | Control | 16.72329328 | 0.143378321 | 16.1070694 | 0.422513079 | 1.532857814 | 0.61622388 | 0.513718202 | 0.011477712 | 1.940144662 | Unpaired t-test | 0.115381997 | permutation FDR (250 perm) | 1 | 0.015690206 | FDR correction BH |
| Slh~Q24179 | VMAT~Control | VMAT | Control | 17.11473683 | 0.110144057 | 16.49971022 | 0.135381373 | 1.531586244 | 0.615026607 | 0.570498595 | 1.34E-05 | 4.871559858 | Unpaired t-test | 0.086055856 | permutation FDR (250 perm) | 1 | 3.26E-05 | FDR correction BH |
| lqfR~Q9VD13 | VMAT~Control | VMAT | Control | 19.82166166 | 0.069032099 | 19.20708825 | 0.15330551 | 1.5311052 | 0.614573412 | 0.571594938 | 9.75E-06 | 5.010898673 | Unpaired t-test | 0.085581982 | permutation FDR (250 perm) | 1 | 2.46E-05 | FDR correction BH |
| Galphao~P16378-2 | VMAT~Control | VMAT | Control | 18.372285 | 0.035042397 | 17.75819948 | 0.070097697 | 1.530587498 | 0.61408552 | 0.593292082 | 7.79E-09 | 8.108426055 | Unpaired t-test | 0.07632159 | permutation FDR (250 perm) | 1 | 6.20E-08 | FDR correction BH |
| ATPsynbeta~X2JH42 | VMAT~Control | VMAT | Control | 19.40353864 | 0.079765524 | 18.79015852 | 0.070266242 | 1.529839303 | 0.613380118 | 0.585543826 | 1.47E-07 | 6.831830448 | Unpaired t-test | 0.079469793 | permutation FDR (250 perm) | 1 | 6.71E-07 | FDR correction BH |
| aspr~Q9W5X1 | VMAT~Control | VMAT | Control | 16.90511008 | 0.2375028 | 16.29177586 | 0.386627258 | 1.529790627 | 0.613334214 | 0.509869992 | 0.012840505 | 1.891417885 | Unpaired t-test | 0.11765085 | permutation FDR (250 perm) | 1 | 0.017471994 | FDR correction BH |
| lectin-29Ca~Q9VLM0 | VMAT~Control | VMAT | Control | 14.11465689 | 1.003163035 | 13.50332702 | 0.293501454 | 1.527666755 | 0.611329868 | 0.416597471 | 0.220193905 | 0.657194707 | Unpaired t-test | 0.176821239 | permutation FDR (250 perm) | 1 | 0.250888892 | FDR correction BH |
| RpL8~Q9V3G1 | VMAT~Control | VMAT | Control | 18.48220313 | 0.294099661 | 17.87107428 | 0.537996826 | 1.527453919 | 0.611128857 | 0.479616727 | 0.049948243 | 1.301479781 | Unpaired t-test | 0.135529412 | permutation FDR (250 perm) | 1 | 0.062375684 | FDR correction BH |
| ine~E1JHT2 | VMAT~Control | VMAT | Control | 18.23190221 | 0.11533463 | 17.6212061 | 0.100064301 | 1.52699582 | 0.610696112 | 0.571659714 | 4.38E-06 | 5.358498518 | Unpaired t-test | 0.085553153 | permutation FDR (250 perm) | 1 | 1.23E-05 | FDR correction BH |
| Mrp4~Q9VGM1 | VMAT~Control | VMAT | Control | 16.73851101 | 0.093349513 | 16.12872759 | 0.216659982 | 1.526030097 | 0.609783416 | 0.551588512 | 0.000177806 | 3.750054133 | Unpaired t-test | 0.095366051 | permutation FDR (250 perm) | 1 | 0.00032983 | FDR correction BH |
| Pkc98E~P13678 | VMAT~Control | VMAT | Control | 20.09351644 | 0.12535826 | 19.48395968 | 0.104590145 | 1.525790371 | 0.609556763 | 0.568080076 | 8.09E-06 | 5.092176194 | Unpaired t-test | 0.087209365 | permutation FDR (250 perm) | 1 | 2.09E-05 | FDR correction BH |
| AGBE~A1Z992 | VMAT~Control | VMAT | Control | 19.3688114 | 0.124758969 | 18.76006464 | 0.214164336 | 1.524933957 | 0.608746763 | 0.548004182 | 0.000264759 | 3.577149599 | Unpaired t-test | 0.097339937 | permutation FDR (250 perm) | 1 | 0.000479417 | FDR correction BH |
| mt:CoII~P00408 | VMAT~Control | VMAT | Control | 21.2371756 | 0.173899843 | 20.63010683 | 0.235273559 | 1.523161342 | 0.607068768 | 0.536830202 | 0.000922277 | 3.035138399 | Unpaired t-test | 0.10289618 | permutation FDR (250 perm) | 1 | 0.001481939 | FDR correction BH |
| CG45076~C0HK92-2 | VMAT~Control | VMAT | Control | 16.72822318 | 0.164997116 | 16.12182793 | 0.159656734 | 1.522450423 | 0.606395249 | 0.549929355 | 0.000149888 | 3.824233973 | Unpaired t-test | 0.09625574 | permutation FDR (250 perm) | 1 | 0.000283462 | FDR correction BH |
| ATPsynD~A0A0B4LHL7 | VMAT~Control | VMAT | Control | 17.07052246 | 0.335367686 | 16.46429953 | 0.460811018 | 1.522268584 | 0.606222926 | 0.483092468 | 0.03871361 | 1.412136329 | Unpaired t-test | 0.133333928 | permutation FDR (250 perm) | 1 | 0.049127675 | FDR correction BH |
| CCT1~A4V391 | VMAT~Control | VMAT | Control | 18.40915102 | 0.095381881 | 17.80515505 | 0.099752903 | 1.519920611 | 0.60399597 | 0.568883075 | 1.94E-06 | 5.712636181 | Unpaired t-test | 0.08678018 | permutation FDR (250 perm) | 1 | 6.03E-06 | FDR correction BH |
| Acsl~A1Z7H3 | VMAT~Control | VMAT | Control | 17.74431459 | 0.036719654 | 17.14080467 | 0.19035507 | 1.51940863 | 0.603509921 | 0.55536082 | 3.89E-05 | 4.409502585 | Unpaired t-test | 0.093359748 | permutation FDR (250 perm) | 1 | 8.44E-05 | FDR correction BH |
| atl~A0A0B4LHV8 | VMAT~Control | VMAT | Control | 16.35351319 | 0.073396841 | 15.75011525 | 0.123209775 | 1.519290704 | 0.603397944 | 0.567030344 | 2.77E-06 | 5.557062605 | Unpaired t-test | 0.087695633 | permutation FDR (250 perm) | 1 | 8.20E-06 | FDR correction BH |
| Pgd~P41572 | VMAT~Control | VMAT | Control | 16.83083716 | 0.097022846 | 16.2276955 | 0.232556415 | 1.519020835 | 0.603141658 | 0.542056909 | 0.000322597 | 3.491340073 | Unpaired t-test | 0.100278902 | permutation FDR (250 perm) | 1 | 0.000568408 | FDR correction BH |
| RpL11~A0A0B4LGZ5 | VMAT~Control | VMAT | Control | 20.45460029 | 0.114428178 | 19.85220783 | 0.069362709 | 1.518232208 | 0.602392462 | 0.568379793 | 1.50E-06 | 5.825113211 | Unpaired t-test | 0.087061261 | permutation FDR (250 perm) | 1 | 4.80E-06 | FDR correction BH |
| poly~Q9VFW4 | VMAT~Control | VMAT | Control | 16.75889037 | 0.115492986 | 16.15718005 | 0.223665958 | 1.517514526 | 0.601710326 | 0.540826997 | 0.000325918 | 3.486892052 | Unpaired t-test | 0.100948268 | permutation FDR (250 perm) | 1 | 0.000572555 | FDR correction BH |
| mRpS11~Q9VEN6 | VMAT~Control | VMAT | Control | 16.34953595 | 0.253659644 | 15.74800843 | 0.151573212 | 1.517322249 | 0.601527518 | 0.531314492 | 0.00105526 | 2.976640598 | Unpaired t-test | 0.105899371 | permutation FDR (250 perm) | 1 | 0.001682982 | FDR correction BH |
| Sec23~Q9VNF8 | VMAT~Control | VMAT | Control | 15.27427072 | 0.295400105 | 14.67432931 | 0.282565228 | 1.515655021 | 0.599941418 | 0.507215459 | 0.00826504 | 2.082755038 | Unpaired t-test | 0.119253691 | permutation FDR (250 perm) | 1 | 0.011636306 | FDR correction BH |
| Nup43~Q9VE85 | VMAT~Control | VMAT | Control | 15.19328204 | 0.096447224 | 14.59471426 | 0.170597907 | 1.514212604 | 0.598567782 | 0.550335232 | 4.57E-05 | 4.339999965 | Unpaired t-test | 0.096057632 | permutation FDR (250 perm) | 1 | 9.72E-05 | FDR correction BH |
| betaTub56D~Q24560 | VMAT~Control | VMAT | Control | 21.98424853 | 0.057795038 | 21.38572923 | 0.06458221 | 1.514161722 | 0.598519302 | 0.576187122 | 2.64E-08 | 7.577795239 | Unpaired t-test | 0.083475676 | permutation FDR (250 perm) | 1 | 1.62E-07 | FDR correction BH |
| eIF3k~Q9W2D9 | VMAT~Control | VMAT | Control | 17.0348831 | 0.095933047 | 16.43802832 | 0.195521115 | 1.512415744 | 0.596854772 | 0.543881875 | 0.0001115 | 3.95272616 | Unpaired t-test | 0.09939694 | permutation FDR (250 perm) | 1 | 0.000215847 | FDR correction BH |
| RpL13A~Q9VNE9 | VMAT~Control | VMAT | Control | 18.42189616 | 0.115706992 | 17.82544078 | 0.222017013 | 1.511997109 | 0.596455381 | 0.536398114 | 0.00033429 | 3.475876684 | Unpaired t-test | 0.103104719 | permutation FDR (250 perm) | 1 | 0.000585505 | FDR correction BH |
| mEFTu1~A1Z9E3 | VMAT~Control | VMAT | Control | 20.14324723 | 0.109837755 | 19.54790154 | 0.068655291 | 1.510834565 | 0.595345696 | 0.562747247 | 1.24E-06 | 5.907908699 | Unpaired t-test | 0.089680324 | permutation FDR (250 perm) | 1 | 4.05E-06 | FDR correction BH |
| shi~E1JJA4 | VMAT~Control | VMAT | Control | 20.62292386 | 0.087997727 | 20.02765788 | 0.117604255 | 1.51075108 | 0.595265973 | 0.558574506 | 3.89E-06 | 5.410057841 | Unpaired t-test | 0.091733453 | permutation FDR (250 perm) | 1 | 1.11E-05 | FDR correction BH |
| RpL15~A8Y560 | VMAT~Control | VMAT | Control | 18.48866121 | 0.16918491 | 17.89429078 | 0.208441693 | 1.509813582 | 0.594370429 | 0.53065967 | 0.000577994 | 3.238077045 | Unpaired t-test | 0.106260557 | permutation FDR (250 perm) | 1 | 0.000960195 | FDR correction BH |
| Adh~P00334 | VMAT~Control | VMAT | Control | 19.79023131 | 0.90629147 | 19.19622981 | 0.767890676 | 1.509427537 | 0.594001499 | 0.387924711 | 0.289638258 | 0.538144073 | Unpaired t-test | 0.197972627 | permutation FDR (250 perm) | 1 | 0.324361779 | FDR correction BH |
| SmydA-8~X2JDK3 | VMAT~Control | VMAT | Control | 14.34539259 | 0.40390412 | 13.75196479 | 0.136312353 | 1.508827419 | 0.593427799 | 0.49841043 | 0.011007969 | 1.958292817 | Unpaired t-test | 0.124549441 | permutation FDR (250 perm) | 1 | 0.01510948 | FDR correction BH |
| Tim23~Q8MRW1 | VMAT~Control | VMAT | Control | 17.00327045 | 0.174633492 | 16.41165903 | 0.154190903 | 1.506928971 | 0.591611417 | 0.535790533 | 0.000204258 | 3.689820947 | Unpaired t-test | 0.103430112 | permutation FDR (250 perm) | 1 | 0.00037535 | FDR correction BH |
| Fitm2~Q9VRJ2 | VMAT~Control | VMAT | Control | 16.82856994 | 0.404079298 | 16.23709452 | 0.355238495 | 1.506786933 | 0.591475427 | 0.476760401 | 0.033780586 | 1.471332815 | Unpaired t-test | 0.137270379 | permutation FDR (250 perm) | 1 | 0.043170196 | FDR correction BH |
| Rab5~Q9V3I2 | VMAT~Control | VMAT | Control | 15.8203173 | 0.124194466 | 15.22922955 | 0.242881073 | 1.506382091 | 0.591087753 | 0.526817918 | 0.000676285 | 3.169870277 | Unpaired t-test | 0.108454627 | permutation FDR (250 perm) | 1 | 0.00110939 | FDR correction BH |
| COX5B~Q9VMB9 | VMAT~Control | VMAT | Control | 19.01966207 | 0.178798024 | 18.42921124 | 0.308766725 | 1.505717194 | 0.590450827 | 0.509200097 | 0.004106002 | 2.386580797 | Unpaired t-test | 0.118089526 | permutation FDR (250 perm) | 1 | 0.005977446 | FDR correction BH |
| D2hgdh~Q7K511 | VMAT~Control | VMAT | Control | 16.62366978 | 0.163937155 | 16.03512403 | 0.287385525 | 1.50373021 | 0.58854575 | 0.512686862 | 0.002610883 | 2.583212508 | Unpaired t-test | 0.115974922 | permutation FDR (250 perm) | 1 | 0.003904714 | FDR correction BH |
| FASN1~Q9VQL7 | VMAT~Control | VMAT | Control | 19.35895234 | 0.094062098 | 18.77111602 | 0.14582913 | 1.502990944 | 0.587836317 | 0.545501849 | 1.89E-05 | 4.722608286 | Unpaired t-test | 0.098579658 | permutation FDR (250 perm) | 1 | 4.41E-05 | FDR correction BH |
| PKD~A0A0B4KHC3 | VMAT~Control | VMAT | Control | 15.7801485 | 0.358209331 | 15.19305247 | 0.656240086 | 1.502219915 | 0.58709603 | 0.439985017 | 0.109627691 | 0.960079732 | Unpaired t-test | 0.161116733 | permutation FDR (250 perm) | 1 | 0.130930282 | FDR correction BH |
| bocks~Q709R6-2 | VMAT~Control | VMAT | Control | 18.25346408 | 0.286349884 | 17.66692833 | 0.536771787 | 1.501636632 | 0.58653575 | 0.461086334 | 0.056508903 | 1.247883124 | Unpaired t-test | 0.147377501 | permutation FDR (250 perm) | 1 | 0.070196284 | FDR correction BH |
| Dmel\CG15602~Q4QQ70 | VMAT~Control | VMAT | Control | 17.45446395 | 0.157315166 | 16.86794375 | 0.211336243 | 1.501620448 | 0.586520201 | 0.524698724 | 0.000555075 | 3.25564866 | Unpaired t-test | 0.109638814 | permutation FDR (250 perm) | 1 | 0.000924731 | FDR correction BH |
| ninaE~P06002 | VMAT~Control | VMAT | Control | 19.43074052 | 0.081514551 | 18.84463741 | 0.121160106 | 1.501186381 | 0.586103107 | 0.550173458 | 4.24E-06 | 5.372221727 | Unpaired t-test | 0.096129671 | permutation FDR (250 perm) | 1 | 1.20E-05 | FDR correction BH |
| anon-WO0140519.58~Q9VG86 | VMAT~Control | VMAT | Control | 18.76147524 | 0.179510731 | 18.17643742 | 0.167555418 | 1.500078311 | 0.585037818 | 0.527147983 | 0.000334205 | 3.475987175 | Unpaired t-test | 0.108226415 | permutation FDR (250 perm) | 1 | 0.000585505 | FDR correction BH |
| ScsbetaG~Q9V470 | VMAT~Control | VMAT | Control | 16.90881581 | 0.135680828 | 16.3244606 | 0.144431582 | 1.499368717 | 0.584355207 | 0.536784044 | 6.13E-05 | 4.212819936 | Unpaired t-test | 0.102915955 | permutation FDR (250 perm) | 1 | 0.000126276 | FDR correction BH |
| Pfk~P52034 | VMAT~Control | VMAT | Control | 20.37124844 | 0.068953557 | 19.78717864 | 0.096940698 | 1.499072122 | 0.584069795 | 0.554565956 | 6.71E-07 | 6.173028431 | Unpaired t-test | 0.093790185 | permutation FDR (250 perm) | 1 | 2.38E-06 | FDR correction BH |
| Alg2~Q9VZU8 | VMAT~Control | VMAT | Control | 20.03972566 | 0.18048167 | 19.45574294 | 0.058031557 | 1.498981641 | 0.583982714 | 0.538340286 | 4.26E-05 | 4.370936757 | Unpaired t-test | 0.102162843 | permutation FDR (250 perm) | 1 | 9.13E-05 | FDR correction BH |
| 21430192~Q9VM18 | VMAT~Control | VMAT | Control | 18.08504847 | 0.174320292 | 17.50274507 | 0.233566197 | 1.497237826 | 0.582303402 | 0.515158387 | 0.001201846 | 2.920151244 | Unpaired t-test | 0.11470372 | permutation FDR (250 perm) | 1 | 0.001897482 | FDR correction BH |
| RpS27~Q9VBU9 | VMAT~Control | VMAT | Control | 18.70040969 | 0.131988716 | 18.11873302 | 0.160396906 | 1.496587534 | 0.581676664 | 0.532234309 | 9.36E-05 | 4.02858365 | Unpaired t-test | 0.105405211 | permutation FDR (250 perm) | 1 | 0.000185371 | FDR correction BH |
| mtd~Q9VNA1 | VMAT~Control | VMAT | Control | 18.03809698 | 0.142398932 | 17.45814306 | 0.294029959 | 1.494801502 | 0.579953918 | 0.506022318 | 0.002645588 | 2.577477721 | Unpaired t-test | 0.119957047 | permutation FDR (250 perm) | 1 | 0.00394909 | FDR correction BH |
| ND-42~A4V383 | VMAT~Control | VMAT | Control | 20.76256242 | 0.06479105 | 20.18273556 | 0.112381304 | 1.494669861 | 0.579826861 | 0.54803387 | 1.60E-06 | 5.796656686 | Unpaired t-test | 0.097318325 | permutation FDR (250 perm) | 1 | 5.07E-06 | FDR correction BH |
| rtp~Q9VN91 | VMAT~Control | VMAT | Control | 16.24813443 | 0.162318328 | 15.66847063 | 0.147731273 | 1.494500936 | 0.5796638 | 0.527852567 | 0.00014992 | 3.824141769 | Unpaired t-test | 0.107825696 | permutation FDR (250 perm) | 1 | 0.000283462 | FDR correction BH |
| FASN2~Q9VQL6 | VMAT~Control | VMAT | Control | 17.23127418 | 0.06639588 | 16.65186066 | 0.065390425 | 1.494241689 | 0.579413518 | 0.556232153 | 7.24E-08 | 7.140391752 | Unpaired t-test | 0.092929311 | permutation FDR (250 perm) | 1 | 3.60E-07 | FDR correction BH |
| Rack1~O18640 | VMAT~Control | VMAT | Control | 20.29978474 | 0.149720764 | 19.72264905 | 0.131236749 | 1.491884332 | 0.577135685 | 0.529949689 | 7.06E-05 | 4.151440702 | Unpaired t-test | 0.106659479 | permutation FDR (250 perm) | 1 | 0.000143065 | FDR correction BH |
| ATPsyngamma~O01666 | VMAT~Control | VMAT | Control | 22.15693778 | 0.13248036 | 21.57999617 | 0.164320742 | 1.491683656 | 0.576941613 | 0.527178451 | 0.00011388 | 3.943551064 | Unpaired t-test | 0.10821743 | permutation FDR (250 perm) | 1 | 0.000220275 | FDR correction BH |
| Gs1~E1JHQ1 | VMAT~Control | VMAT | Control | 18.74093604 | 0.147002817 | 18.16442921 | 0.124729819 | 1.491234174 | 0.576506827 | 0.530747096 | 5.45E-05 | 4.263255602 | Unpaired t-test | 0.106204852 | permutation FDR (250 perm) | 1 | 0.00011413 | FDR correction BH |
| Nfs1~Q9VKD3 | VMAT~Control | VMAT | Control | 16.99384701 | 0.125222515 | 16.4209316 | 0.082113354 | 1.487526543 | 0.572915411 | 0.536956721 | 6.51E-06 | 5.186278532 | Unpaired t-test | 0.102874101 | permutation FDR (250 perm) | 1 | 1.75E-05 | FDR correction BH |
| Galk~Q95U34 | VMAT~Control | VMAT | Control | 16.79159824 | 0.086982856 | 16.22001627 | 0.082051118 | 1.486152292 | 0.571581962 | 0.542567625 | 8.61E-07 | 6.065204988 | Unpaired t-test | 0.100012601 | permutation FDR (250 perm) | 1 | 2.95E-06 | FDR correction BH |
| Rrp40~Q8IPX7 | VMAT~Control | VMAT | Control | 15.74509731 | 0.090295904 | 15.17369914 | 0.199692094 | 1.48596298 | 0.571398174 | 0.520394093 | 0.000166057 | 3.779743734 | Unpaired t-test | 0.11186541 | permutation FDR (250 perm) | 1 | 0.000311224 | FDR correction BH |
| Mhc~P05661-25 | VMAT~Control | VMAT | Control | 19.64594104 | 0.303198249 | 19.07689759 | 0.278228171 | 1.483539616 | 0.569043453 | 0.480597769 | 0.011403507 | 1.942961549 | Unpaired t-test | 0.134905526 | permutation FDR (250 perm) | 1 | 0.015597825 | FDR correction BH |
| serp~Q9VW34 | VMAT~Control | VMAT | Control | 14.29864888 | 1.158974617 | 13.73117666 | 0.234289568 | 1.481924786 | 0.567472227 | 0.371189543 | 0.308419512 | 0.510858154 | Unpaired t-test | 0.211211292 | permutation FDR (250 perm) | 1 | 0.344248237 | FDR correction BH |
| sesB~X2JB48 | VMAT~Control | VMAT | Control | 23.10308648 | 0.115148128 | 22.53563144 | 0.177970991 | 1.481907138 | 0.567455045 | 0.518319672 | 0.000134605 | 3.870939762 | Unpaired t-test | 0.112947085 | permutation FDR (250 perm) | 1 | 0.000256359 | FDR correction BH |
| Yp3~X2JEX8 | VMAT~Control | VMAT | Control | 20.36916292 | 0.272802134 | 19.80204877 | 0.527711379 | 1.481557013 | 0.567114145 | 0.448074582 | 0.058559308 | 1.232404067 | Unpaired t-test | 0.1556751 | permutation FDR (250 perm) | 1 | 0.072513735 | FDR correction BH |
| Sec61alpha~Q8STG9 | VMAT~Control | VMAT | Control | 17.6196965 | 0.120035399 | 17.0557216 | 0.086524641 | 1.478336701 | 0.563974891 | 0.528970764 | 6.74E-06 | 5.171572404 | Unpaired t-test | 0.107227314 | permutation FDR (250 perm) | 1 | 1.80E-05 | FDR correction BH |
| Hsp83~M9PBL3 | VMAT~Control | VMAT | Control | 19.1926757 | 0.091931235 | 18.62899643 | 0.077660483 | 1.478033809 | 0.56367927 | 0.534891848 | 1.04E-06 | 5.983600586 | Unpaired t-test | 0.103868764 | permutation FDR (250 perm) | 1 | 3.48E-06 | FDR correction BH |
| Usp5~Q9VZU7 | VMAT~Control | VMAT | Control | 16.64114971 | 0.226971236 | 16.07902738 | 0.178185559 | 1.476439589 | 0.562122327 | 0.49787314 | 0.001430211 | 2.844599824 | Unpaired t-test | 0.124817531 | permutation FDR (250 perm) | 1 | 0.002223723 | FDR correction BH |
| Tom40~M9PGL7 | VMAT~Control | VMAT | Control | 15.59096735 | 0.298420678 | 15.02968228 | 0.128816736 | 1.475583004 | 0.561285077 | 0.490050935 | 0.003153425 | 2.501217474 | Unpaired t-test | 0.129372041 | permutation FDR (250 perm) | 1 | 0.004665721 | FDR correction BH |
| Neurochondrin~Q9VI25 | VMAT~Control | VMAT | Control | 17.43265465 | 0.112371752 | 16.87272023 | 0.122551945 | 1.474202207 | 0.559934424 | 0.521179944 | 1.99E-05 | 4.7005539 | Unpaired t-test | 0.111502693 | permutation FDR (250 perm) | 1 | 4.62E-05 | FDR correction BH |
| aralar1~A0A0B4KHW3 | VMAT~Control | VMAT | Control | 17.1107437 | 0.092375313 | 16.55219529 | 0.056220024 | 1.472786602 | 0.558548408 | 0.532782557 | 4.18E-07 | 6.378518687 | Unpaired t-test | 0.105081761 | permutation FDR (250 perm) | 1 | 1.60E-06 | FDR correction BH |
| Dmel\CG4836~Q8MRL0 | VMAT~Control | VMAT | Control | 18.7507344 | 0.295484437 | 18.19442607 | 0.435856471 | 1.470501567 | 0.556308322 | 0.450272779 | 0.03979502 | 1.40017127 | Unpaired t-test | 0.154312611 | permutation FDR (250 perm) | 1 | 0.050322151 | FDR correction BH |
| eIF4A~C9QP42 | VMAT~Control | VMAT | Control | 19.73702192 | 0.113203154 | 19.18083426 | 0.095245958 | 1.470378577 | 0.556187652 | 0.521672966 | 7.61E-06 | 5.118706285 | Unpaired t-test | 0.111222621 | permutation FDR (250 perm) | 1 | 1.97E-05 | FDR correction BH |
| Dmel\CG14969~Q95SH7 | VMAT~Control | VMAT | Control | 16.5046957 | 0.212479279 | 15.94963768 | 0.182038708 | 1.469227726 | 0.555058027 | 0.493328624 | 0.001262481 | 2.89877502 | Unpaired t-test | 0.127440322 | permutation FDR (250 perm) | 1 | 0.001986552 | FDR correction BH |
| Dmel\CG2846~A0A0B4LHX1 | VMAT~Control | VMAT | Control | 16.00127321 | 0.119711947 | 15.4468254 | 0.138728687 | 1.468606418 | 0.55444781 | 0.512453735 | 4.94E-05 | 4.305927595 | Unpaired t-test | 0.116148679 | permutation FDR (250 perm) | 1 | 0.000104658 | FDR correction BH |
| SmydA-9~Q9W373 | VMAT~Control | VMAT | Control | 16.91999501 | 0.123067051 | 16.36665365 | 0.180461952 | 1.467480526 | 0.553341359 | 0.504098394 | 0.000208233 | 3.681450144 | Unpaired t-test | 0.121018345 | permutation FDR (250 perm) | 1 | 0.000382356 | FDR correction BH |
| MRP~Q7KTC0 | VMAT~Control | VMAT | Control | 16.25016732 | 0.107882604 | 15.69691307 | 0.216725993 | 1.467391921 | 0.553254248 | 0.499206602 | 0.000457248 | 3.339847791 | Unpaired t-test | 0.124132438 | permutation FDR (250 perm) | 1 | 0.000778281 | FDR correction BH |
| GstZ2~Q9VHD2-3 | VMAT~Control | VMAT | Control | 16.65077882 | 0.09550426 | 16.10099867 | 0.271296606 | 1.463862606 | 0.549780153 | 0.487123526 | 0.001626033 | 2.788870743 | Unpaired t-test | 0.131098617 | permutation FDR (250 perm) | 1 | 0.00250831 | FDR correction BH |
| Naa15-16~Q9VWI2 | VMAT~Control | VMAT | Control | 18.46333736 | 0.188602534 | 17.91413273 | 0.130066599 | 1.463278754 | 0.549204629 | 0.498163754 | 0.0003189 | 3.496345895 | Unpaired t-test | 0.124649374 | permutation FDR (250 perm) | 1 | 0.000562746 | FDR correction BH |
| CG16760~Q7KV94 | VMAT~Control | VMAT | Control | 17.21022027 | 0.12581432 | 16.66184931 | 0.204506506 | 1.462433436 | 0.548370961 | 0.495196805 | 0.000459423 | 3.337786977 | Unpaired t-test | 0.12634347 | permutation FDR (250 perm) | 1 | 0.000780429 | FDR correction BH |
| Dmel\CG8586~A1Z7H7 | VMAT~Control | VMAT | Control | 16.47916897 | 0.268986248 | 15.93428647 | 0.07806958 | 1.458901516 | 0.544882497 | 0.484228732 | 0.001443431 | 2.840603884 | Unpaired t-test | 0.132735058 | permutation FDR (250 perm) | 1 | 0.002242797 | FDR correction BH |
| Dmel\CG9356~Q9VHE3 | VMAT~Control | VMAT | Control | 16.34124132 | 0.045639242 | 15.79721446 | 0.094532956 | 1.458036529 | 0.544026864 | 0.519632437 | 4.05E-07 | 6.39218416 | Unpaired t-test | 0.112262001 | permutation FDR (250 perm) | 1 | 1.56E-06 | FDR correction BH |
| Lst8~Q9W328 | VMAT~Control | VMAT | Control | 15.87141794 | 0.09302633 | 15.3279481 | 0.056301755 | 1.457473692 | 0.543469842 | 0.518267149 | 5.69E-07 | 6.245101832 | Unpaired t-test | 0.112973991 | permutation FDR (250 perm) | 1 | 2.07E-06 | FDR correction BH |
| Vha44~Q9V7N5-2 | VMAT~Control | VMAT | Control | 17.42909192 | 0.031905019 | 16.8863504 | 0.060142462 | 1.456738094 | 0.542741519 | 0.526705023 | 6.59E-09 | 8.18109734 | Unpaired t-test | 0.108522911 | permutation FDR (250 perm) | 1 | 5.44E-08 | FDR correction BH |
| Ppcs~Q7KN99 | VMAT~Control | VMAT | Control | 15.82443655 | 0.123360889 | 15.2817081 | 0.209453045 | 1.456724891 | 0.542728443 | 0.489513776 | 0.000543326 | 3.264939115 | Unpaired t-test | 0.129704332 | permutation FDR (250 perm) | 1 | 0.000907086 | FDR correction BH |
| pn~O18399 | VMAT~Control | VMAT | Control | 15.26320842 | 0.070529237 | 14.7223306 | 0.802335589 | 1.454857471 | 0.540877823 | 0.397646083 | 0.164097871 | 0.784897052 | Unpaired t-test | 0.1903117 | permutation FDR (250 perm) | 1 | 0.19010157 | FDR correction BH |
| Dmel\CG4115~Q9VGA3 | VMAT~Control | VMAT | Control | 18.36035487 | 0.123812218 | 17.81949474 | 0.144507298 | 1.454839623 | 0.540860124 | 0.498441696 | 8.29E-05 | 4.081253315 | Unpaired t-test | 0.124536913 | permutation FDR (250 perm) | 1 | 0.000165733 | FDR correction BH |
| CaMKII~D1YSG7 | VMAT~Control | VMAT | Control | 20.63497865 | 0.06194757 | 20.09503816 | 0.111983816 | 1.453912541 | 0.539940488 | 0.51071114 | 2.70E-06 | 5.568007269 | Unpaired t-test | 0.11715667 | permutation FDR (250 perm) | 1 | 8.04E-06 | FDR correction BH |
| Hel25E~Q27268 | VMAT~Control | VMAT | Control | 17.46628862 | 0.127980009 | 16.92743823 | 0.075098081 | 1.452814374 | 0.538850382 | 0.505317257 | 1.03E-05 | 4.985800364 | Unpaired t-test | 0.120350783 | permutation FDR (250 perm) | 1 | 2.58E-05 | FDR correction BH |
| Ilk~Q9V400 | VMAT~Control | VMAT | Control | 16.59555438 | 0.175449654 | 16.05714409 | 0.146885431 | 1.452371261 | 0.538410288 | 0.488428984 | 0.000367282 | 3.435000715 | Unpaired t-test | 0.13035105 | permutation FDR (250 perm) | 1 | 0.000637127 | FDR correction BH |
| Dmel\CG7409~Q9VSA9 | VMAT~Control | VMAT | Control | 17.51449151 | 0.388261082 | 16.97685159 | 0.518122588 | 1.45159593 | 0.537639916 | 0.41692038 | 0.093000842 | 1.031513121 | Unpaired t-test | 0.176561062 | permutation FDR (250 perm) | 1 | 0.111924326 | FDR correction BH |
| RtcB~Q9VIW7 | VMAT~Control | VMAT | Control | 14.94032726 | 0.329218586 | 14.4032788 | 0.299872984 | 1.45100094 | 0.537048454 | 0.44785666 | 0.022441325 | 1.648951496 | Unpaired t-test | 0.155831336 | permutation FDR (250 perm) | 1 | 0.02936458 | FDR correction BH |
| UQCR-C1~Q9VFF0 | VMAT~Control | VMAT | Control | 21.21828837 | 0.121882692 | 20.68190395 | 0.199113997 | 1.450333238 | 0.536384421 | 0.485677388 | 0.000439325 | 3.357213885 | Unpaired t-test | 0.131844781 | permutation FDR (250 perm) | 1 | 0.000750487 | FDR correction BH |
| Syx5~A4V0S1 | VMAT~Control | VMAT | Control | 18.07077427 | 0.069813563 | 17.5349308 | 0.11920556 | 1.449789523 | 0.535843468 | 0.504665205 | 5.76E-06 | 5.239410847 | Unpaired t-test | 0.120715884 | permutation FDR (250 perm) | 1 | 1.58E-05 | FDR correction BH |
| Cpr~Q8IPJ7 | VMAT~Control | VMAT | Control | 16.14693017 | 0.254622586 | 15.6121989 | 0.275214669 | 1.448672285 | 0.534731269 | 0.457945 | 0.009669245 | 2.014607449 | Unpaired t-test | 0.149385505 | permutation FDR (250 perm) | 1 | 0.01338906 | FDR correction BH |

| | | | | | | | | | | | | | | | | | | |
|---|---|---|---|---|---|---|---|---|---|---|---|---|---|---|---|---|---|---|
| Dmel\CG43346~M9NEW0 | VMAT~Control | VMAT | Control | 15.62662352 | 0.149712158 | 15.09283066 | 0.136149906 | 1.447730291 | 0.533792857 | 0.489494037 | 0.0001514 | 3.819874188 | Unpaired t-test | 0.129729343 | permutation FDR (250 perm) | 1 | 0.000285802 | FDR correction BH |
| fax~Q59E30 | VMAT~Control | VMAT | Control | 16.61805568 | 0.045966649 | 16.08916567 | 0.132726103 | 1.442818679 | 0.528890006 | 0.497630901 | 7.50E-06 | 5.124737134 | Unpaired t-test | 0.124966011 | permutation FDR (250 perm) | 1 | 1.95E-05 | FDR correction BH |
| Etfb~Q0KHZ6 | VMAT~Control | VMAT | Control | 16.55309224 | 0.200207205 | 16.02699707 | 0.308834352 | 1.440026311 | 0.526095171 | 0.451739936 | 0.009552224 | 2.019895495 | Unpaired t-test | 0.153432889 | permutation FDR (250 perm) | 1 | 0.013242601 | FDR correction BH |
| CG32410~Q8I935 | VMAT~Control | VMAT | Control | 17.58297104 | 0.067278245 | 17.0569234 | 0.112174219 | 1.439978869 | 0.526047641 | 0.496976096 | 4.17E-06 | 5.380057302 | Unpaired t-test | 0.125372093 | permutation FDR (250 perm) | 1 | 1.18E-05 | FDR correction BH |
| Dmel\CG5087~M9PEV2 | VMAT~Control | VMAT | Control | 19.14177117 | 0.263137256 | 18.61707644 | 0.329490699 | 1.438629139 | 0.524694731 | 0.441448039 | 0.019370778 | 1.712852934 | Unpaired t-test | 0.160111851 | permutation FDR (250 perm) | 1 | 0.025646126 | FDR correction BH |
| Scsalpha1~Q94522 | VMAT~Control | VMAT | Control | 20.33707872 | 0.052570124 | 19.81275065 | 0.145488877 | 1.438263557 | 0.524328069 | 0.490401213 | 1.88E-05 | 4.72478918 | Unpaired t-test | 0.129114783 | permutation FDR (250 perm) | 1 | 4.39E-05 | FDR correction BH |
| 26-29-p~Q9V3U6 | VMAT~Control | VMAT | Control | 18.48435825 | 0.163746476 | 17.96036988 | 0.155593287 | 1.437924947 | 0.523988375 | 0.475913062 | 0.000408835 | 3.388451548 | Unpaired t-test | 0.137684773 | permutation FDR (250 perm) | 1 | 0.000700946 | FDR correction BH |
| ND-23~Q9VF27 | VMAT~Control | VMAT | Control | 17.27079892 | 0.270137194 | 16.74700259 | 0.405573714 | 1.437733556 | 0.523796338 | 0.430071503 | 0.037091629 | 1.430724092 | Unpaired t-test | 0.16762766 | permutation FDR (250 perm) | 1 | 0.047171094 | FDR correction BH |
| LBR~Q8MLV1-2 | VMAT~Control | VMAT | Control | 18.15497844 | 0.163893214 | 17.6361887 | 0.321510572 | 1.43275282 | 0.518789735 | 0.446698128 | 0.009260246 | 2.033377482 | Unpaired t-test | 0.156553928 | permutation FDR (250 perm) | 1 | 0.012868134 | FDR correction BH |
| Act79B~M9PFZ6 | VMAT~Control | VMAT | Control | 21.58183705 | 0.161242323 | 21.06488641 | 0.219578422 | 1.430927562 | 0.51695064 | 0.460809795 | 0.001707805 | 2.767561739 | Unpaired t-test | 0.147546465 | permutation FDR (250 perm) | 1 | 0.002629282 | FDR correction BH |
| P5CS~Q9VNW6 | VMAT~Control | VMAT | Control | 19.11188845 | 0.087188464 | 18.5953516 | 0.045788346 | 1.430517214 | 0.516536858 | 0.494747296 | 3.62E-07 | 6.441010845 | Unpaired t-test | 0.126599284 | permutation FDR (250 perm) | 1 | 1.41E-06 | FDR correction BH |
| Dmel\CG5958~Q9VM12 | VMAT~Control | VMAT | Control | 16.96244717 | 0.154093716 | 16.44627243 | 0.170572069 | 1.430158192 | 0.516174735 | 0.46805813 | 0.000520872 | 3.283268646 | Unpaired t-test | 0.142662517 | permutation FDR (250 perm) | 1 | 0.000875185 | FDR correction BH |
| Dmel\CG3609~Q9VQB4 | VMAT~Control | VMAT | Control | 16.77651484 | 0.31942831 | 16.26080784 | 0.317461224 | 1.429694591 | 0.515706994 | 0.429253918 | 0.028345395 | 1.547517485 | Unpaired t-test | 0.168102791 | permutation FDR (250 perm) | 1 | 0.036541654 | FDR correction BH |
| ND-13A~Q9VMU0 | VMAT~Control | VMAT | Control | 18.26522466 | 0.298972188 | 17.74984986 | 0.489500876 | 1.42936543 | 0.515374802 | 0.410162617 | 0.0722766 | 1.141002283 | Unpaired t-test | 0.181436533 | permutation FDR (250 perm) | 1 | 0.088154983 | FDR correction BH |
| ClC-a~Q9VGH7-6 | VMAT~Control | VMAT | Control | 16.16599809 | 0.12728713 | 15.65416006 | 0.24906354 | 1.425865629 | 0.511838032 | 0.454931676 | 0.002172794 | 2.66298144 | Unpaired t-test | 0.151340151 | permutation FDR (250 perm) | 1 | 0.003293469 | FDR correction BH |
| Nsf2~A0A0B4LH53 | VMAT~Control | VMAT | Control | 15.29221473 | 0.276213714 | 14.78045613 | 0.2130991 | 1.425787128 | 0.511758601 | 0.442691556 | 0.008286652 | 2.081620876 | Unpaired t-test | 0.159218819 | permutation FDR (250 perm) | 1 | 0.011652796 | FDR correction BH |
| ND-B14.7~Q7JYH3 | VMAT~Control | VMAT | Control | 15.83787781 | 0.121495469 | 15.32935618 | 0.172124348 | 1.422591674 | 0.508521625 | 0.464733949 | 0.000302653 | 3.519054837 | Unpaired t-test | 0.144793241 | permutation FDR (250 perm) | 1 | 0.000539323 | FDR correction BH |
| Irp-1B~Q9VGZ3 | VMAT~Control | VMAT | Control | 18.24053715 | 0.141148868 | 17.73262982 | 0.154297774 | 1.421986067 | 0.507907329 | 0.464469699 | 0.000288508 | 3.539841401 | Unpaired t-test | 0.144988884 | permutation FDR (250 perm) | 1 | 0.000516856 | FDR correction BH |
| CG9132~Q9VXB0 | VMAT~Control | VMAT | Control | 18.82079703 | 0.162247801 | 18.31489701 | 0.108960208 | 1.420008941 | 0.505900014 | 0.465236788 | 0.000175781 | 3.755028989 | Unpaired t-test | 0.144465985 | permutation FDR (250 perm) | 1 | 0.000326846 | FDR correction BH |
| SdhB~P21914 | VMAT~Control | VMAT | Control | 16.42693897 | 0.300144561 | 15.92190116 | 0.479904498 | 1.419160549 | 0.50503781 | 0.403018403 | 0.073988837 | 1.1308338 | Unpaired t-test | 0.186379143 | permutation FDR (250 perm) | 1 | 0.090056733 | FDR correction BH |
| RpS29~A0A0C4DHE8 | VMAT~Control | VMAT | Control | 17.40816461 | 0.105333383 | 16.90510239 | 0.174565333 | 1.417218521 | 0.503062225 | 0.461026249 | 0.00025552 | 3.592575801 | Unpaired t-test | 0.147407737 | permutation FDR (250 perm) | 1 | 0.000463757 | FDR correction BH |
| Lkb1~Q8T9L5 | VMAT~Control | VMAT | Control | 15.71276113 | 0.224120087 | 15.21007131 | 0.323169152 | 1.416852741 | 0.502689821 | 0.42750111 | 0.017010819 | 1.76927477 | Unpaired t-test | 0.169254207 | permutation FDR (250 perm) | 1 | 0.022636217 | FDR correction BH |
| Eno~E1JHR5 | VMAT~Control | VMAT | Control | 19.64801545 | 0.256412907 | 19.14632365 | 0.301494609 | 1.415872932 | 0.501691796 | 0.426245893 | 0.017717634 | 1.751594268 | Unpaired t-test | 0.169965471 | permutation FDR (250 perm) | 1 | 0.023536857 | FDR correction BH |
| Grasp65~Q9VW57 | VMAT~Control | VMAT | Control | 18.90301623 | 0.128094427 | 18.40140247 | 0.349671161 | 1.41579635 | 0.501613761 | 0.430001284 | 0.013073566 | 1.883605944 | Unpaired t-test | 0.167666667 | permutation FDR (250 perm) | 1 | 0.017778841 | FDR correction BH |
| Lam~M9NE89 | VMAT~Control | VMAT | Control | 20.95683456 | 0.119433057 | 20.45538893 | 0.36159303 | 1.415631366 | 0.501445633 | 0.428475431 | 0.014673341 | 1.833471003 | Unpaired t-test | 0.168648649 | permutation FDR (250 perm) | 1 | 0.019760322 | FDR correction BH |
| Galphai~P20353 | VMAT~Control | VMAT | Control | 15.94162307 | 0.148103543 | 15.44026266 | 0.156813661 | 1.41554775 | 0.501360416 | 0.45725263 | 0.000402774 | 3.394938943 | Unpaired t-test | 0.149888839 | permutation FDR (250 perm) | 1 | 0.00069156 | FDR correction BH |
| Tapdelta~Q7KLX3 | VMAT~Control | VMAT | Control | 16.8095516 | 0.188951305 | 16.3085809 | 0.18895375 | 1.415165414 | 0.500970694 | 0.44749341 | 0.001851829 | 2.732399229 | Unpaired t-test | 0.156067466 | permutation FDR (250 perm) | 1 | 0.002826981 | FDR correction BH |
| GramD1B~A8DYU2 | VMAT~Control | VMAT | Control | 13.93391157 | 0.926605107 | 13.43513121 | 0.258414085 | 1.413018508 | 0.498780362 | 0.348747849 | 0.273225517 | 0.563478744 | Unpaired t-test | 0.230440141 | permutation FDR (250 perm) | 1 | 0.306857284 | FDR correction BH |
| PhKgamma~X2JEJ4 | VMAT~Control | VMAT | Control | 15.66305848 | 0.162139142 | 15.16620321 | 0.080087889 | 1.411134274 | 0.496855271 | 0.459679097 | 0.000109243 | 3.961607251 | Unpaired t-test | 0.148288128 | permutation FDR (250 perm) | 1 | 0.000212703 | FDR correction BH |
| Mhc~P05661-24 | VMAT~Control | VMAT | Control | 16.55192907 | 0.430805801 | 16.05511161 | 0.471889352 | 1.411097286 | 0.496817456 | 0.386402014 | 0.112734338 | 0.94794378 | Unpaired t-test | 0.199251214 | permutation FDR (250 perm) | 1 | 0.134299915 | FDR correction BH |
| Hsc70-4~C7LA75 | VMAT~Control | VMAT | Control | 20.93163969 | 0.050067879 | 20.43494248 | 0.066833109 | 1.410979681 | 0.496697212 | 0.478815573 | 1.10E-07 | 6.956924378 | Unpaired t-test | 0.135951872 | permutation FDR (250 perm) | 1 | 5.22E-07 | FDR correction BH |
| CaMKII~A4V134 | VMAT~Control | VMAT | Control | 16.43269268 | 0.226480969 | 15.93637435 | 0.174832456 | 1.410609168 | 0.496318322 | 0.440016795 | 0.00306438 | 2.51365732 | Unpaired t-test | 0.161095428 | permutation FDR (250 perm) | 1 | 0.00453968 | FDR correction BH |
| GH07925p~Q9VVU1 | VMAT~Control | VMAT | Control | 17.16183626 | 0.190472679 | 16.66990929 | 0.20938887 | 1.406322016 | 0.491926977 | 0.43665178 | 0.003029082 | 2.518688937 | Unpaired t-test | 0.163155654 | permutation FDR (250 perm) | 1 | 0.004490214 | FDR correction BH |
| Aldh~Q9VLC5 | VMAT~Control | VMAT | Control | 19.58735495 | 0.09061324 | 19.09671136 | 0.106254385 | 1.40507154 | 0.490643588 | 0.461803417 | 1.38E-05 | 4.860309638 | Unpaired t-test | 0.146906181 | permutation FDR (250 perm) | 1 | 3.34E-05 | FDR correction BH |
| ScpX~Q9VJ43 | VMAT~Control | VMAT | Control | 13.88723819 | 0.785928485 | 13.39668333 | 0.506552231 | 1.404985137 | 0.490554869 | 0.34590999 | 0.26792773 | 0.571982335 | Unpaired t-test | 0.232867194 | permutation FDR (250 perm) | 1 | 0.301194784 | FDR correction BH |
| Pgm2a~A1Z7P1 | VMAT~Control | VMAT | Control | 15.55953028 | 0.166221212 | 15.07140018 | 0.209441206 | 1.402625728 | 0.488130097 | 0.435994518 | 0.002206975 | 2.656202587 | Unpaired t-test | 0.16356541 | permutation FDR (250 perm) | 1 | 0.003340977 | FDR correction BH |
| l(2)tid~Q27237 | VMAT~Control | VMAT | Control | 15.2281746 | 0.199919712 | 14.74126027 | 0.258679524 | 1.401444227 | 0.486914331 | 0.424804743 | 0.007614022 | 2.118385867 | Unpaired t-test | 0.170958831 | permutation FDR (250 perm) | 1 | 0.01076481 | FDR correction BH |
| Art1~Q9VGW7 | VMAT~Control | VMAT | Control | 15.72208884 | 0.234055822 | 15.23527286 | 0.347518927 | 1.401348692 | 0.48681598 | 0.409992612 | 0.026580282 | 1.57544042 | Unpaired t-test | 0.181570986 | permutation FDR (250 perm) | 1 | 0.034435874 | FDR correction BH |
| Dmel\CG5439~Q9VK31 | VMAT~Control | VMAT | Control | 14.12201614 | 0.921110477 | 13.63604622 | 0.42620351 | 1.400527113 | 0.485969913 | 0.334254266 | 0.309478073 | 0.509370116 | Unpaired t-test | 0.243701099 | permutation FDR (250 perm) | 1 | 0.345266058 | FDR correction BH |
| Arp3~P32392 | VMAT~Control | VMAT | Control | 16.41479139 | 0.201927018 | 15.93066212 | 0.298332444 | 1.398741402 | 0.484129263 | 0.416955033 | 0.013229614 | 1.878452826 | Unpaired t-test | 0.176529204 | permutation FDR (250 perm) | 1 | 0.017959926 | FDR correction BH |
| Dmel\CG15531~Q9VA93 | VMAT~Control | VMAT | Control | 16.01267616 | 0.139858065 | 15.52972817 | 0.13376953 | 1.397596581 | 0.482947984 | 0.444478355 | 0.000234112 | 3.630576406 | Unpaired t-test | 0.157984909 | permutation FDR (250 perm) | 1 | 0.000426548 | FDR correction BH |
| RpL18~M9PHM6 | VMAT~Control | VMAT | Control | 21.59528269 | 0.06002451 | 21.11256665 | 0.388211574 | 1.397371902 | 0.482716036 | 0.410585765 | 0.020557292 | 1.687034091 | Unpaired t-test | 0.181066785 | permutation FDR (250 perm) | 1 | 0.027079947 | FDR correction BH |
| ThrRS~Q8IP94 | VMAT~Control | VMAT | Control | 17.5138978 | 0.096404063 | 17.03175844 | 0.131550548 | 1.396813454 | 0.48213936 | 0.449363941 | 6.00E-05 | 4.221849073 | Unpaired t-test | 0.154936057 | permutation FDR (250 perm) | 1 | 0.000124112 | FDR correction BH |
| Drat~Q0E9G4 | VMAT~Control | VMAT | Control | 16.24531014 | 0.060480688 | 15.76549188 | 0.246804081 | 1.394567975 | 0.479818257 | 0.430855831 | 0.00176527 | 2.753188925 | Unpaired t-test | 0.167015957 | permutation FDR (250 perm) | 1 | 0.002708895 | FDR correction BH |
| UQCR-14~Q9VXI6 | VMAT~Control | VMAT | Control | 13.87131329 | 0.9553266 | 13.3939605 | 0.2418033 | 1.39218679 | 0.477352791 | 0.331332086 | 0.30416698 | 0.516887934 | Unpaired t-test | 0.246535385 | permutation FDR (250 perm) | 1 | 0.339662747 | FDR correction BH |
| RpS20~P55828 | VMAT~Control | VMAT | Control | 19.83760328 | 0.123929626 | 19.36050953 | 0.132564722 | 1.391936845 | 0.477093754 | 0.441280903 | 0.000155463 | 3.808374337 | Unpaired t-test | 0.160232579 | permutation FDR (250 perm) | 1 | 0.000293001 | FDR correction BH |
| Vps2~Q9VBI3 | VMAT~Control | VMAT | Control | 15.98128673 | 0.239629227 | 15.50470281 | 0.276641536 | 1.391445035 | 0.47658392 | 0.409549595 | 0.01551912 | 1.809132897 | Unpaired t-test | 0.181891199 | permutation FDR (250 perm) | 1 | 0.020780438 | FDR correction BH |
| CCT5~A1Z8U4 | VMAT~Control | VMAT | Control | 19.86978002 | 0.130059287 | 19.39410958 | 0.101094482 | 1.390564282 | 0.475670439 | 0.443032719 | 7.28E-05 | 4.137640877 | Unpaired t-test | 0.158989791 | permutation FDR (250 perm) | 1 | 0.00014743 | FDR correction BH |
| Dmel\CG9953~Q9VS02 | VMAT~Control | VMAT | Control | 15.71080082 | 0.358216187 | 15.23550537 | 0.125727382 | 1.390202888 | 0.475295447 | 0.406311832 | 0.018811363 | 1.725579733 | Unpaired t-test | 0.184160991 | permutation FDR (250 perm) | 1 | 0.024947577 | FDR correction BH |
| cg17026~Q9VUW1 | VMAT~Control | VMAT | Control | 14.83840339 | 0.230294767 | 14.36345511 | 0.214931774 | 1.389868386 | 0.474948273 | 0.416301109 | 0.007104983 | 2.148436947 | Unpaired t-test | 0.177030088 | permutation FDR (250 perm) | 1 | 0.010081453 | FDR correction BH |
| SERCA~P22700-2 | VMAT~Control | VMAT | Control | 21.85774633 | 0.045748478 | 21.38339463 | 0.071912644 | 1.38929378 | 0.474351704 | 0.456934928 | 2.07E-07 | 6.683363791 | Unpaired t-test | 0.150082703 | permutation FDR (250 perm) | 1 | 8.94E-07 | FDR correction BH |
| RpS17~P17704 | VMAT~Control | VMAT | Control | 18.92599845 | 0.216020729 | 18.45262311 | 0.261332201 | 1.388353879 | 0.473375345 | 0.411047838 | 0.010838966 | 1.965012139 | Unpaired t-test | 0.180810619 | permutation FDR (250 perm) | 1 | 0.014894878 | FDR correction BH |
| p24-1~Q9VYT6 | VMAT~Control | VMAT | Control | 15.51093788 | 0.317186672 | 15.03949483 | 0.212572778 | 1.386495613 | 0.471443052 | 0.402681273 | 0.020100289 | 1.696797704 | Unpaired t-test | 0.186628369 | permutation FDR (250 perm) | 1 | 0.026522466 | FDR correction BH |
| RpL35A~Q9VNB9 | VMAT~Control | VMAT | Control | 19.20765222 | 0.114724579 | 18.73755923 | 0.235720786 | 1.385198748 | 0.470092989 | 0.420762785 | 0.002479388 | 2.605655418 | Unpaired t-test | 0.173831858 | permutation FDR (250 perm) | 1 | 0.003724621 | FDR correction BH |
| ctp~A0A4D6K1L3 | VMAT~Control | VMAT | Control | 16.34422457 | 0.256530455 | 15.87464686 | 0.266217969 | 1.384704095 | 0.469577712 | 0.402954797 | 0.017544099 | 1.75586893 | Unpaired t-test | 0.186419797 | permutation FDR (250 perm) | 1 | 0.023319486 | FDR correction BH |
| Dmel\CG11178~Q9VYB2 | VMAT~Control | VMAT | Control | 14.02994148 | 0.818521168 | 13.56174126 | 0.259707233 | 1.383382601 | 0.468200216 | 0.338285741 | 0.25076304 | 0.600736474 | Unpaired t-test | 0.239950748 | permutation FDR (250 perm) | 1 | 0.284480095 | FDR correction BH |
| Cp1~A0A0B4JD48 | VMAT~Control | VMAT | Control | 19.43359291 | 0.144259405 | 18.96566191 | 0.130275471 | 1.383124473 | 0.467930996 | 0.430507773 | 0.000308789 | 3.510338322 | Unpaired t-test | 0.167317376 | permutation FDR (250 perm) | 1 | 0.000547769 | FDR correction BH |
| Psa~Q9W0E4 | VMAT~Control | VMAT | Control | 16.8508434 | 0.086745157 | 16.38487974 | 0.086173901 | 1.381239654 | 0.465963658 | 0.441804839 | 6.75E-06 | 5.17099131 | Unpaired t-test | 0.159829561 | permutation FDR (250 perm) | 1 | 1.80E-05 | FDR correction BH |
| Dmel\CG17597~Q9VJ44 | VMAT~Control | VMAT | Control | 17.4073098 | 0.116600047 | 16.94218812 | 0.170701103 | 1.380433778 | 0.46512168 | 0.425760396 | 0.000513297 | 3.28963114 | Unpaired t-test | 0.170296591 | permutation FDR (250 perm) | 1 | 0.000864307 | FDR correction BH |
| Vps29~Q9VPX5 | VMAT~Control | VMAT | Control | 16.0204554 | 0.233730426 | 15.55533644 | 0.075874547 | 1.380431171 | 0.465118956 | 0.419064936 | 0.00173742 | 2.760095073 | Unpaired t-test | 0.175012389 | permutation FDR (250 perm) | 1 | 0.00266964 | FDR correction BH |
| Pdh~Q7KNR7 | VMAT~Control | VMAT | Control | 19.67743304 | 0.101572202 | 19.21456738 | 0.052261218 | 1.378276809 | 0.462865663 | 0.440369589 | 3.89E-06 | 5.409503717 | Unpaired t-test | 0.160903684 | permutation FDR (250 perm) | 1 | 1.11E-05 | FDR correction BH |
| 128up~P32234 | VMAT~Control | VMAT | Control | 15.93542587 | 0.210333588 | 15.47319222 | 0.186144188 | 1.377673149 | 0.462233652 | 0.410651542 | 0.004247086 | 2.371908963 | Unpaired t-test | 0.181099115 | permutation FDR (250 perm) | 1 | 0.006171383 | FDR correction BH |
| RpL5~R9Q794 | VMAT~Control | VMAT | Control | 19.05727879 | 0.095607366 | 18.59591046 | 0.144575904 | 1.376847079 | 0.461368334 | 0.428178065 | 0.000140867 | 3.851189618 | Unpaired t-test | 0.168854231 | permutation FDR (250 perm) | 1 | 0.000267637 | FDR correction BH |
| Rpt1~Q7KMQ0 | VMAT~Control | VMAT | Control | 17.05480051 | 0.120460038 | 16.59496401 | 0.089252636 | 1.37538594 | 0.459836503 | 0.430942971 | 4.41E-05 | 4.3552762 | Unpaired t-test | 0.166962766 | permutation FDR (250 perm) | 1 | 9.43E-05 | FDR correction BH |
| GCC185~Q9VGE4 | VMAT~Control | VMAT | Control | 16.17303597 | 0.129580068 | 15.71329356 | 0.159632352 | 1.375296242 | 0.459742412 | 0.421029035 | 0.000537374 | 3.269723259 | Unpaired t-test | 0.173631858 | permutation FDR (250 perm) | 1 | 0.000900982 | FDR correction BH |
| Cdc42~M9NFF8 | VMAT~Control | VMAT | Control | 15.99881417 | 0.097312735 | 15.54012504 | 0.319327244 | 1.374292529 | 0.458689126 | 0.399106017 | 0.01179164 | 1.928425804 | Unpaired t-test | 0.189215011 | permutation FDR (250 perm) | 1 | 0.016100649 | FDR correction BH |
| AcCoAS~Q59E09 | VMAT~Control | VMAT | Control | 18.63618544 | 0.121613955 | 18.17899402 | 0.160498282 | 1.37286658 | 0.457191426 | 0.419420453 | 0.000480032 | 3.318729784 | Unpaired t-test | 0.17476991 | permutation FDR (250 perm) | 1 | 0.000811778 | FDR correction BH |
| hoip~D2NUK9 | VMAT~Control | VMAT | Control | 16.73272029 | 0.291809657 | 16.28181637 | 0.308411465 | 1.366896417 | 0.45090392 | 0.378949332 | 0.038964142 | 1.409334885 | Unpaired t-test | 0.205183223 | permutation FDR (250 perm) | 1 | 0.049392348 | FDR correction BH |
| Rho1~P48148 | VMAT~Control | VMAT | Control | 19.15115965 | 0.162651129 | 18.70277381 | 0.110513774 | 1.364512721 | 0.448385844 | 0.412141439 | 0.000464926 | 3.332615938 | Unpaired t-test | 0.180021239 | permutation FDR (250 perm) | 1 | 0.000788499 | FDR correction BH |
| Sam-S~P40320 | VMAT~Control | VMAT | Control | 16.72014331 | 0.135973828 | 16.27466075 | 0.13284349 | 1.361769533 | 0.445482561 | 0.410577986 | 0.000378715 | 3.421687644 | Unpaired t-test | 0.181072092 | permutation FDR (250 perm) | 1 | 0.000653927 | FDR correction BH |
| alphaSnap~Q23983 | VMAT~Control | VMAT | Control | 18.50843255 | 0.155481506 | 18.06744448 | 0.33141826 | 1.357533754 | 0.44098807 | 0.378948602 | 0.022558416 | 1.64669139 | Unpaired t-test | 0.205186755 | permutation FDR (250 perm) | 1 | 0.029485015 | FDR correction BH |
| Muted~A0A0B4LHG2 | VMAT~Control | VMAT | Control | 17.98306714 | 0.261289823 | 17.54217918 | 0.570977274 | 1.357439558 | 0.440887961 | 0.344224345 | 0.147481989 | 0.831261013 | Unpaired t-test | 0.234379947 | permutation FDR (250 perm) | 1 | 0.172722688 | FDR correction BH |
| bgm~B5RIV0 | VMAT~Control | VMAT | Control | 15.9673504 | 0.219173976 | 15.52995996 | 0.204811678 | 1.354152706 | 0.437390438 | 0.385653842 | 0.008568529 | 2.067093719 | Unpaired t-test | 0.199848124 | permutation FDR (250 perm) | 1 | 0.012027619 | FDR correction BH |
| Aldh7A1~Q8SXQ1 | VMAT~Control | VMAT | Control | 16.57716941 | 0.230440983 | 16.14064529 | 0.23397738 | 1.353339795 | 0.436524115 | 0.380623389 | 0.013991726 | 1.8541287 | Unpaired t-test | 0.203821634 | permutation FDR (250 perm) | 1 | 0.018874799 | FDR correction BH |
| GlyP~A4UZZ4 | VMAT~Control | VMAT | Control | 21.19746135 | 0.083015195 | 20.76335625 | 0.095992203 | 1.351072503 | 0.434105096 | 0.410790414 | 1.74E-05 | 4.759164822 | Unpaired t-test | 0.180969912 | permutation FDR (250 perm) | 1 | 4.08E-05 | FDR correction BH |
| Dmel\CG18528~Q9VC87 | VMAT~Control | VMAT | Control | 13.91558699 | 0.269035283 | 13.48166558 | 0.315721627 | 1.350900497 | 0.433921415 | 0.366022491 | 0.04140031 | 1.382996402 | Unpaired t-test | 0.215733569 | permutation FDR (250 perm) | 1 | 0.052227401 | FDR correction BH |
| Dmel\CG11474~Q9W299 | VMAT~Control | VMAT | Control | 15.96733979 | 0.403175952 | 15.53419294 | 0.210235244 | 1.350175412 | 0.433146851 | 0.359951789 | 0.059012437 | 1.229056449 | Unpaired t-test | 0.220766858 | permutation FDR (250 perm) | 1 | 0.072921405 | FDR correction BH |
| Srrm234~Q7YZ99 | VMAT~Control | VMAT | Control | 19.56048399 | 0.322682314 | 19.12788418 | 0.510781526 | 1.349663554 | 0.432599815 | 0.340577983 | 0.140442836 | 0.85250041 | Unpaired t-test | 0.237854002 | permutation FDR (250 perm) | 1 | 0.165053637 | FDR correction BH |
| Dmel\CG3626~Q9W4K8 | VMAT~Control | VMAT | Control | 15.62199976 | 0.228787867 | 15.19124528 | 0.350434839 | 1.347938318 | 0.43075448 | 0.362843827 | 0.044141224 | 1.355155625 | Unpaired t-test | 0.218447287 | permutation FDR (250 perm) | 1 | 0.055417836 | FDR correction BH |
| RnrL~P48591 | VMAT~Control | VMAT | Control | 15.81219623 | 0.233119921 | 15.38396872 | 0.218989719 | 1.345579386 | 0.42822751 | 0.374639315 | 0.013486421 | 1.8701033 | Unpaired t-test | 0.208474636 | permutation FDR (250 perm) | 1 | 0.018287462 | FDR correction BH |
| GstD1~P20432 | VMAT~Control | VMAT | Control | 19.91102778 | 0.108100886 | 19.48758431 | 0.184383822 | 1.341124769 | 0.423443462 | 0.3864996 | 0.001274164 | 2.894774636 | Unpaired t-test | 0.199168212 | permutation FDR (250 perm) | 1 | 0.002000922 | FDR correction BH |
| GstD4~Q9VG96 | VMAT~Control | VMAT | Control | 19.99719728 | 0.307005363 | 19.57407235 | 0.319520123 | 1.34082869 | 0.423124924 | 0.353144333 | 0.058504137 | 1.232813419 | Unpaired t-test | 0.226489437 | permutation FDR (250 perm) | 1 | 0.072483547 | FDR correction BH |
| Ogdh~Q9VVC5 | VMAT~Control | VMAT | Control | 19.42062629 | 0.054741045 | 18.99779138 | 0.215420647 | 1.340559174 | 0.422834903 | 0.384604856 | 0.001679298 | 2.774872108 | Unpaired t-test | 0.200692274 | permutation FDR (250 perm) | 1 | 0.002587087 | FDR correction BH |
| Dmel\CG17593~Q9VQR9 | VMAT~Control | VMAT | Control | 17.45824818 | 0.145105457 | 17.03715114 | 0.393463077 | 1.33894531 | 0.421097034 | 0.354594089 | 0.048566193 | 1.313665942 | Unpaired t-test | 0.225292254 | permutation FDR (250 perm) | 1 | 0.060746449 | FDR correction BH |
| jar~A0A0B4LHV4 | VMAT~Control | VMAT | Control | 17.00492725 | 0.112749547 | 16.58440363 | 0.094181018 | 1.338413243 | 0.420523626 | 0.394598462 | 7.83E-05 | 4.106463094 | Unpaired t-test | 0.192639294 | permutation FDR (250 perm) | 1 | 0.000157321 | FDR correction BH |
| SC35~Q9V3T8 | VMAT~Control | VMAT | Control | 17.54258521 | 0.18408366 | 17.12280387 | 0.546640711 | 1.337724788 | 0.419781338 | 0.333701503 | 0.134722019 | 0.870561416 | Unpaired t-test | 0.244269011 | permutation FDR (250 perm) | 1 | 0.15864714 | FDR correction BH |
| RpL7~X2J5G6 | VMAT~Control | VMAT | Control | 19.39512477 | 0.062935833 | 18.98171195 | 0.219543684 | 1.331832654 | 0.413412818 | 0.375100944 | 0.002332515 | 2.632175491 | Unpaired t-test | 0.208088222 | permutation FDR (250 perm) | 1 | 0.00351295 | FDR correction BH |
| Mlp60A~A0A0B4LGF8 | VMAT~Control | VMAT | Control | 19.75169027 | 0.138923388 | 19.33917906 | 0.329976976 | 1.331000594 | 0.412511216 | 0.35557778 | 0.027589039 | 1.559263426 | Unpaired t-test | 0.224390845 | permutation FDR (250 perm) | 1 | 0.035605591 | FDR correction BH |
| bur~Q9VIE7 | VMAT~Control | VMAT | Control | 16.51122356 | 0.075670811 | 16.10006646 | 0.213733365 | 1.329751901 | 0.4111571 | 0.373304662 | 0.002305458 | 2.637242837 | Unpaired t-test | 0.209552713 | permutation FDR (250 perm) | 1 | 0.003476648 | FDR correction BH |
| DnaJ-H~Q9VK35 | VMAT~Control | VMAT | Control | 19.91470123 | 0.145869837 | 19.50405463 | 0.143583841 | 1.329281445 | 0.410646595 | 0.3762098 | 0.001167816 | 2.932625416 | Unpaired t-test | 0.207207764 | permutation FDR (250 perm) | 1 | 0.00184996 | FDR correction BH |
| Imp~Q0KHU2 | VMAT~Control | VMAT | Control | 16.95850877 | 0.081115547 | 16.54983102 | 0.114731026 | 1.327468613 | 0.40867775 | 0.384515636 | 6.86E-05 | 4.163473156 | Unpaired t-test | 0.20076468 | permutation FDR (250 perm) | 1 | 0.000139758 | FDR correction BH |
| tsr~P45594 | VMAT~Control | VMAT | Control | 19.87331065 | 0.15166126 | 19.4660185 | 0.191848263 | 1.326194289 | 0.407292148 | 0.367138871 | 0.003948851 | 2.403529205 | Unpaired t-test | 0.21482135 | permutation FDR (250 perm) | 1 | 0.005759353 | FDR correction BH |
| Dmel\CG13907~Q9W0L6 | VMAT~Control | VMAT | Control | 19.78886808 | 0.067670932 | 19.38361223 | 0.051375807 | 1.324323755 | 0.405255858 | 0.390421062 | 8.78E-07 | 6.056511625 | Unpaired t-test | 0.196072406 | permutation FDR (250 perm) | 1 | 3.00E-06 | FDR correction BH |
| DmCG8814~Q9VQI6 | VMAT~Control | VMAT | Control | 14.1930317 | 0.792750099 | 13.78838122 | 0.566140868 | 1.323768165 | 0.404650481 | 0.281858114 | 0.374850569 | 0.426141825 | Unpaired t-test | 0.299335083 | permutation FDR (250 perm) | 1 | 0.411950626 | FDR correction BH |
| dock~Q8IPW2 | VMAT~Control | VMAT | Control | 17.55235441 | 0.098091912 | 17.14906995 | 0.151186659 | 1.32251534 | 0.403284457 | 0.373205214 | 0.000534402 | 3.272131632 | Unpaired t-test | 0.209619762 | permutation FDR (250 perm) | 1 | 0.000896638 | FDR correction BH |
| Nop56~Q95WY3 | VMAT~Control | VMAT | Control | 16.59446613 | 0.132343367 | 16.19533132 | 0.123636599 | 1.318716841 | 0.399134817 | 0.36922918 | 0.000597798 | 3.223445526 | Unpaired t-test | 0.212952801 | permutation FDR (250 perm) | 1 | 0.000991696 | FDR correction BH |
| Dmel\CG6972~Q9VCV9 | VMAT~Control | VMAT | Control | 14.4878904 | 0.894501559 | 14.08977263 | 0.849281339 | 1.317787517 | 0.398117766 | 0.256582341 | 0.486994647 | 0.312475812 | Unpaired t-test | 0.331416339 | permutation FDR (250 perm) | 1 | 0.52418171 | FDR correction BH |
| Vha68-2~A4V0N4 | VMAT~Control | VMAT | Control | 20.03700734 | 0.0588322 | 19.63891505 | 0.053200995 | 1.31776425 | 0.398092293 | 0.384454637 | 5.47E-07 | 6.261947095 | Unpaired t-test | 0.200812362 | permutation FDR (250 perm) | 1 | 2.01E-06 | FDR correction BH |
| chic~M9NEU5 | VMAT~Control | VMAT | Control | 19.33620739 | 0.154716829 | 18.93869751 | 0.236768847 | 1.31723238 | 0.397509882 | 0.352875204 | 0.010462827 | 1.980350973 | Unpaired t-test | 0.226758803 | permutation FDR (250 perm) | 1 | 0.014420078 | FDR correction BH |
| Chc~X2JC31 | VMAT~Control | VMAT | Control | 18.6215971 | 0.127966978 | 18.22457771 | 0.055699927 | 1.316784623 | 0.397019394 | 0.373695282 | 8.23E-05 | 4.084356186 | Unpaired t-test | 0.209201588 | permutation FDR (250 perm) | 1 | 0.000164692 | FDR correction BH |
| Prx3~Q9VEJ0 | VMAT~Control | VMAT | Control | 18.67352257 | 0.130702603 | 18.27683997 | 0.136338044 | 1.316477254 | 0.396682595 | 0.365786623 | 0.00084612 | 3.072568191 | Unpaired t-test | 0.215938244 | permutation FDR (250 perm) | 1 | 0.001368911 | FDR correction BH |
| RpL17~X2JIQ5 | VMAT~Control | VMAT | Control | 20.38505136 | 0.157327829 | 19.98870572 | 0.292474566 | 1.316169817 | 0.396345642 | 0.345092005 | 0.02354887 | 1.62802992 | Unpaired t-test | 0.233525066 | permutation FDR (250 perm) | 1 | 0.030694375 | FDR correction BH |
| CASK~E1JIS7 | VMAT~Control | VMAT | Control | 15.86613838 | 0.204404181 | 15.4710652 | 0.524712967 | 1.315009469 | 0.395073188 | 0.315595213 | 0.147771528 | 0.830409235 | Unpaired t-test | 0.262017559 | permutation FDR (250 perm) | 1 | 0.172889751 | FDR correction BH |
| Rpn5~Q9V3Z4 | VMAT~Control | VMAT | Control | 16.48358197 | 0.123966585 | 16.08961924 | 0.08716998 | 1.313997679 | 0.393962728 | 0.368957164 | 0.000169946 | 3.769690265 | Unpaired t-test | 0.213235112 | permutation FDR (250 perm) | 1 | 0.000316998 | FDR correction BH |
| Pka-R1~M9MS76 | VMAT~Control | VMAT | Control | 16.2971878 | 0.177934179 | 15.90544875 | 0.116534293 | 1.311973928 | 0.391739051 | 0.357712758 | 0.002082782 | 2.681356184 | Unpaired t-test | 0.222718943 | permutation FDR (250 perm) | 1 | 0.003161102 | FDR correction BH |
| Sema1a~M9PB77 | VMAT~Control | VMAT | Control | 15.19754116 | 0.122353645 | 14.80830579 | 0.175295462 | 1.30969908 | 0.389235373 | 0.355270708 | 0.002245096 | 2.648765054 | Unpaired t-test | 0.224679577 | permutation FDR (250 perm) | 1 | 0.003389966 | FDR correction BH |
| Dmel\CG5162~Q9VX69 | VMAT~Control | VMAT | Control | 17.69384374 | 0.432028196 | 17.30492239 | 0.122695877 | 1.309414034 | 0.388921347 | 0.323871844 | 0.081568237 | 1.088478926 | Unpaired t-test | 0.25389011 | permutation FDR (250 perm) | 1 | 0.098669902 | FDR correction BH |
| Dys~Q9VDW3 | VMAT~Control | VMAT | Control | 16.40748522 | 0.220638266 | 16.01906238 | 0.194477729 | 1.308961666 | 0.388422847 | 0.343271758 | 0.014458371 | 1.839880641 | Unpaired t-test | 0.235359719 | permutation FDR (250 perm) | 1 | 0.019481972 | FDR correction BH |
| RpL27A~M9MRC9 | VMAT~Control | VMAT | Control | 21.83754619 | 0.205000205 | 21.454507 | 0.465244794 | 1.304086158 | 0.383039188 | 0.31208214 | 0.12295396 | 0.910257478 | Unpaired t-test | 0.265583151 | permutation FDR (250 perm) | 1 | 0.145590353 | FDR correction BH |
| ACSL1~E2QCY2 | VMAT~Control | VMAT | Control | 15.48464795 | 0.184393065 | 15.10653273 | 0.167149493 | 1.299642854 | 0.378115221 | 0.340245498 | 0.00680271 | 2.167318019 | Unpaired t-test | 0.238182938 | permutation FDR (250 perm) | 1 | 0.009670036 | FDR correction BH |
| Prx5~Q960M4 | VMAT~Control | VMAT | Control | 19.10578906 | 0.219815864 | 18.72770617 | 0.323469085 | 1.299613728 | 0.378082889 | 0.321799869 | 0.055948697 | 1.252210021 | Unpaired t-test | 0.255806678 | permutation FDR (250 perm) | 1 | 0.069537082 | FDR correction BH |
| Epac~A1Z6P8 | VMAT~Control | VMAT | Control | 15.73038158 | 0.212949972 | 15.3543784 | 0.290341092 | 1.297741626 | 0.376003178 | 0.323854439 | 0.041689331 | 1.379975072 | Unpaired t-test | 0.253897143 | permutation FDR (250 perm) | 1 | 0.052535699 | FDR correction BH |
| Fas1~B7Z0L1 | VMAT~Control | VMAT | Control | 15.75683333 | 0.178862856 | 15.38103976 | 0.225126784 | 1.297553096 | 0.375793575 | 0.332976847 | 0.015235933 | 1.817130945 | Unpaired t-test | 0.244894945 | permutation FDR (250 perm) | 1 | 0.020447769 | FDR correction BH |
| Rpt3~Q9V405 | VMAT~Control | VMAT | Control | 16.26019383 | 0.102100514 | 15.88483778 | 0.129511192 | 1.297159647 | 0.375356049 | 0.349573862 | 0.000471298 | 3.326704648 | Unpaired t-test | 0.22968662 | permutation FDR (250 perm) | 1 | 0.000797581 | FDR correction BH |
| eIF2gamma~A0A0B4KGP8 | VMAT~Control | VMAT | Control | 17.5616841 | 0.093583583 | 17.186399 | 0.046986971 | 1.297095855 | 0.375285099 | 0.358496393 | 1.16E-05 | 4.93551462 | Unpaired t-test | 0.221936564 | permutation FDR (250 perm) | 1 | 2.86E-05 | FDR correction BH |
| Tg~Q8IPH0 | VMAT~Control | VMAT | Control | 15.34358098 | 0.166271472 | 14.96862774 | 0.036763817 | 1.296797517 | 0.374953234 | 0.348419417 | 0.000601664 | 3.220645916 | Unpaired t-test | 0.230801056 | permutation FDR (250 perm) | 1 | 0.000996705 | FDR correction BH |
| verm~Q8IQU1 | VMAT~Control | VMAT | Control | 13.54036312 | 0.976347578 | 13.16746726 | 0.441242793 | 1.294949524 | 0.372895864 | 0.252100522 | 0.4544497 | 0.342514179 | Unpaired t-test | 0.33750393 | permutation FDR (250 perm) | 1 | 0.492303081 | FDR correction BH |
| Act5C~X2JCP8 | VMAT~Control | VMAT | Control | 19.24300043 | 0.134215947 | 18.8701775 | 0.645655277 | 1.294884064 | 0.372822933 | 0.287912273 | 0.23484488 | 0.629218903 | Unpaired t-test | 0.292289093 | permutation FDR (250 perm) | 1 | 0.267065144 | FDR correction BH |
| Lectin-galC1~Q9VIX1 | VMAT~Control | VMAT | Control | 16.44439396 | 0.100297364 | 16.07224138 | 0.110484804 | 1.294282533 | 0.372152583 | 0.348871357 | 0.000235171 | 3.628616597 | Unpaired t-test | 0.230352113 | permutation FDR (250 perm) | 1 | 0.000428145 | FDR correction BH |
| Dmel\CG2747~Q32KD3 | VMAT~Control | VMAT | Control | 13.89319699 | 0.759791398 | 13.52107202 | 0.409727482 | 1.294257761 | 0.372124969 | 0.268479423 | 0.357807857 | 0.446350127 | Unpaired t-test | 0.315510703 | permutation FDR (250 perm) | 1 | 0.394880307 | FDR correction BH |
| nst~Q9VTZ4 | VMAT~Control | VMAT | Control | 16.26135393 | 0.120034582 | 15.88972537 | 0.100733681 | 1.293812495 | 0.371628551 | 0.347290648 | 0.000346071 | 3.460835221 | Unpaired t-test | 0.23184507 | permutation FDR (250 perm) | 1 | 0.000604789 | FDR correction BH |
| Hsp26~X2JGG6 | VMAT~Control | VMAT | Control | 17.51163347 | 0.169914592 | 17.1407211 | 0.304593875 | 1.293170387 | 0.370912376 | 0.32086408 | 0.038747647 | 1.411754662 | Unpaired t-test | 0.256775044 | permutation FDR (250 perm) | 1 | 0.049144376 | FDR correction BH |
| Kap-alpha3~Q9V455 | VMAT~Control | VMAT | Control | 17.14676706 | 0.097301015 | 16.77620309 | 0.141220296 | 1.292858125 | 0.370563966 | 0.344167983 | 0.000690067 | 3.161108775 | Unpaired t-test | 0.234436236 | permutation FDR (250 perm) | 1 | 0.001130423 | FDR correction BH |
| Dmel\CG3860~Q9W1D9 | VMAT~Control | VMAT | Control | 18.13226556 | 0.133894821 | 17.76301712 | 0.434489582 | 1.291679766 | 0.36924844 | 0.306856315 | 0.099418628 | 1.002532236 | Unpaired t-test | 0.271149123 | permutation FDR (250 perm) | 1 | 0.119342912 | FDR correction BH |
| Mlp84B~Q24400 | VMAT~Control | VMAT | Control | 20.50643336 | 0.092341209 | 20.13821904 | 0.022769371 | 1.290754226 | 0.368214322 | 0.353191956 | 5.86E-06 | 5.232129224 | Unpaired t-test | 0.226463028 | permutation FDR (250 perm) | 1 | 1.60E-05 | FDR correction BH |
| CCT4~Q9VK69 | VMAT~Control | VMAT | Control | 18.38818616 | 0.09179493 | 18.02219988 | 0.146700276 | 1.288762372 | 0.365986277 | 0.33969662 | 0.000805371 | 3.094004136 | Unpaired t-test | 0.238643799 | permutation FDR (250 perm) | 1 | 0.00130838 | FDR correction BH |
| Gpdh1~M9PC43 | VMAT~Control | VMAT | Control | 20.43842726 | 0.059825616 | 20.07283202 | 0.15695711 | 1.288413105 | 0.36559524 | 0.340050819 | 0.000654678 | 3.183972223 | Unpaired t-test | 0.238362357 | permutation FDR (250 perm) | 1 | 0.00107921 | FDR correction BH |
| RpL14~P55841 | VMAT~Control | VMAT | Control | 19.87688802 | 0.11025969 | 19.51185198 | 0.296228676 | 1.287913806 | 0.365036043 | 0.319826425 | 0.027304843 | 1.563760311 | Unpaired t-test | 0.257850615 | permutation FDR (250 perm) | 1 | 0.035258147 | FDR correction BH |
| par-1~Q7YU80 | VMAT~Control | VMAT | Control | 17.10396818 | 0.119923329 | 16.74057395 | 0.186551021 | 1.28644897 | 0.363394231 | 0.330604986 | 0.004360116 | 2.360501958 | Unpaired t-test | 0.247171868 | permutation FDR (250 perm) | 1 | 0.006331717 | FDR correction BH |
| cathD~Q7K485 | VMAT~Control | VMAT | Control | 16.32263729 | 0.205371313 | 15.95936998 | 0.368984621 | 1.2863358 | 0.36326731 | 0.305561201 | 0.083319007 | 1.079255916 | Unpaired t-test | 0.272542105 | permutation FDR (250 perm) | 1 | 0.100684262 | FDR correction BH |
| Dmel\CG9590~Q9VF39 | VMAT~Control | VMAT | Control | 19.29362954 | 0.098221027 | 18.93417883 | 0.140441756 | 1.282937342 | 0.359450712 | 0.333862283 | 0.000854573 | 3.068251002 | Unpaired t-test | 0.244096703 | permutation FDR (250 perm) | 1 | 0.001380689 | FDR correction BH |
| STUB1~Q9XYW6 | VMAT~Control | VMAT | Control | 17.57275456 | 0.107480953 | 17.21451978 | 0.177546971 | 1.281856514 | 0.358234781 | 0.327808539 | 0.003162433 | 2.499978626 | Unpaired t-test | 0.249905934 | permutation FDR (250 perm) | 1 | 0.004676111 | FDR correction BH |
| Dmel\CG5973~Q9VM11 | VMAT~Control | VMAT | Control | 16.45339198 | 0.33337269 | 16.09559128 | 0.135220274 | 1.281470876 | 0.35780069 | 0.308213402 | 0.050357482 | 1.297935995 | Unpaired t-test | 0.269870996 | permutation FDR (250 perm) | 1 | 0.062835724 | FDR correction BH |
| Baldspot~Q9VV87 | VMAT~Control | VMAT | Control | 15.94695339 | 0.164822554 | 15.58921396 | 0.181859088 | 1.28141647 | 0.357739439 | 0.322356711 | 0.008585619 | 2.06622837 | Unpaired t-test | 0.255344176 | permutation FDR (250 perm) | 1 | 0.012044427 | FDR correction BH |
| BG:DS09217.4~Q9V3K7 | VMAT~Control | VMAT | Control | 15.62163888 | 0.248114191 | 15.26519264 | 0.155092416 | 1.280268348 | 0.356446235 | 0.315200833 | 0.02141309 | 1.669320652 | Unpaired t-test | 0.262410887 | permutation FDR (250 perm) | 1 | 0.028144285 | FDR correction BH |
| fra~A1Z921 | VMAT~Control | VMAT | Control | 15.45496048 | 0.100771859 | 15.10072173 | 0.105111371 | 1.27831089 | 0.354238748 | 0.332580927 | 0.000284948 | 3.545234982 | Unpaired t-test | 0.245278242 | permutation FDR (250 perm) | 1 | 0.000511645 | FDR correction BH |
| RpL4~P09180 | VMAT~Control | VMAT | Control | 19.72735631 | 0.100716325 | 19.37331925 | 0.149750697 | 1.278132193 | 0.354037057 | 0.327597278 | 0.001363209 | 2.865437671 | Unpaired t-test | 0.250127473 | permutation FDR (250 perm) | 1 | 0.002126569 | FDR correction BH |
| Gpo1~Q7K569 | VMAT~Control | VMAT | Control | 18.38688812 | 0.315331215 | 18.03412142 | 0.443581603 | 1.277007239 | 0.352766703 | 0.283713176 | 0.177857482 | 0.74992786 | Unpaired t-test | 0.297225919 | permutation FDR (250 perm) | 1 | 0.204932214 | FDR correction BH |
| RpS5a~X2JKU5 | VMAT~Control | VMAT | Control | 16.76736043 | 0.159327174 | 16.41485836 | 0.173039591 | 1.276773018 | 0.352502069 | 0.318950686 | 0.007352905 | 2.133541073 | Unpaired t-test | 0.258681898 | permutation FDR (250 perm) | 1 | 0.010414403 | FDR correction BH |
| ncm~Q9VJ87 | VMAT~Control | VMAT | Control | 15.4399109 | 0.752938673 | 15.09045568 | 0.917990569 | 1.274079425 | 0.349455216 | 0.228258018 | 0.525302851 | 0.279590242 | Unpaired t-test | 0.371942484 | permutation FDR (250 perm) | 1 | 0.561563538 | FDR correction BH |
| RpL24~X2JE06 | VMAT~Control | VMAT | Control | 19.00569444 | 0.141260838 | 18.65827649 | 0.232915123 | 1.272281537 | 0.347417953 | 0.309690555 | 0.017196107 | 1.764569863 | Unpaired t-test | 0.268284335 | permutation FDR (250 perm) | 1 | 0.022869851 | FDR correction BH |
| Dmel\CG18428~Q9VCD4 | VMAT~Control | VMAT | Control | 19.27497991 | 0.336070571 | 18.93093483 | 0.55926146 | 1.269310542 | 0.344045074 | 0.266331354 | 0.265662011 | 0.575670544 | Unpaired t-test | 0.318353866 | permutation FDR (250 perm) | 1 | 0.299362553 | FDR correction BH |

| | | | | | | | | | | | | | | | | | | |
|---|---|---|---|---|---|---|---|---|---|---|---|---|---|---|---|---|---|---|
| FipoQ~Q8T0K1 | VMAT~Control | VMAT | Control | 17.71279097 | 0.077941957 | 17.36887189 | 0.092027103 | 1.269199691 | 0.343919075 | 0.326319622 | 8.07E-05 | 4.093160644 | Unpaired t-test | 0.251432088 | permutation FDR (250 perm) | 1 | 0.000161662 | FDR correction BH |
| RpL37a~M9PH59 | VMAT~Control | VMAT | Control | 20.13630853 | 0.143071288 | 19.79251 | 0.520702426 | 1.269093649 | 0.343798532 | 0.276922896 | 0.18499987 | 0.732828576 | Unpaired t-test | 0.304956294 | permutation FDR (250 perm) | 1 | 0.212745332 | FDR correction BH |
| CG11899~Q9VAN0 | VMAT~Control | VMAT | Control | 20.12812006 | 0.133255151 | 19.78515585 | 0.272279878 | 1.26835993 | 0.342964205 | 0.302019992 | 0.029879424 | 1.524627778 | Unpaired t-test | 0.276698246 | permutation FDR (250 perm) | 1 | 0.038414071 | FDR correction BH |
| RpS23~A0A0B4LFD9 | VMAT~Control | VMAT | Control | 19.11157945 | 0.134517691 | 18.76939578 | 0.259216665 | 1.267673905 | 0.342183676 | 0.30265538 | 0.02559817 | 1.591791089 | Unpaired t-test | 0.275896491 | permutation FDR (250 perm) | 1 | 0.033236675 | FDR correction BH |
| Bka~Q9VL75 | VMAT~Control | VMAT | Control | 19.46069126 | 0.173818715 | 19.12110011 | 0.691952826 | 1.265397931 | 0.339591142 | 0.257448419 | 0.312202486 | 0.505563643 | Unpaired t-test | 0.330259502 | permutation FDR (250 perm) | 1 | 0.347975687 | FDR correction BH |
| CG31123~A0A0B4KFZ2 | VMAT~Control | VMAT | Control | 15.58725322 | 0.433820387 | 15.25024561 | 0.158711676 | 1.263133925 | 0.337007611 | 0.279306675 | 0.133875677 | 0.873298321 | Unpaired t-test | 0.302111888 | permutation FDR (250 perm) | 1 | 0.157729401 | FDR correction BH |
| Dmel\CG15019~Q9VZC8 | VMAT~Control | VMAT | Control | 16.98740824 | 0.266454296 | 16.6507237 | 0.860299139 | 1.262851098 | 0.336684542 | 0.240014323 | 0.422720732 | 0.373946452 | Unpaired t-test | 0.354067161 | permutation FDR (250 perm) | 1 | 0.460474134 | FDR correction BH |
| Tom70~Q9VKC8 | VMAT~Control | VMAT | Control | 16.85177629 | 0.106815656 | 16.51745517 | 0.068975557 | 1.260783984 | 0.334321113 | 0.316333335 | 0.000155317 | 3.808781333 | Unpaired t-test | 0.26123266 | permutation FDR (250 perm) | 1 | 0.000292961 | FDR correction BH |
| RpS12~P80455 | VMAT~Control | VMAT | Control | 17.71716637 | 0.166244365 | 17.38525903 | 0.116547094 | 1.258676333 | 0.331907343 | 0.304279682 | 0.004421944 | 2.354386785 | Unpaired t-test | 0.274019298 | permutation FDR (250 perm) | 1 | 0.006413589 | FDR correction BH |
| Syx18~Q9VC58 | VMAT~Control | VMAT | Control | 18.36441437 | 0.03070298 | 18.03532633 | 0.054943912 | 1.256219039 | 0.329088039 | 0.320078518 | 3.73E-07 | 6.428171397 | Unpaired t-test | 0.257608084 | permutation FDR (250 perm) | 1 | 1.45E-06 | FDR correction BH |
| Dmel\CG32536~Q9VWJ3 | VMAT~Control | VMAT | Control | 15.34484993 | 0.540823379 | 15.01687588 | 0.805651319 | 1.255249413 | 0.32797405 | 0.228720666 | 0.46720637 | 0.330491245 | Unpaired t-test | 0.371180828 | permutation FDR (250 perm) | 1 | 0.50472868 | FDR correction BH |
| Scp1~Q8MSI2 | VMAT~Control | VMAT | Control | 22.70450982 | 0.088908423 | 22.37711117 | 0.127642127 | 1.254748881 | 0.327398659 | 0.306104181 | 0.00083357 | 3.079057841 | Unpaired t-test | 0.271949123 | permutation FDR (250 perm) | 1 | 0.001350464 | FDR correction BH |
| dPR72~A0A0B4LHF0 | VMAT~Control | VMAT | Control | 15.47269691 | 0.270724681 | 15.14574142 | 0.235798266 | 1.254363504 | 0.326955489 | 0.281722916 | 0.069063788 | 1.160749606 | Unpaired t-test | 0.299483815 | permutation FDR (250 perm) | 1 | 0.084542983 | FDR correction BH |
| Cam~A0A0B4LF57 | VMAT~Control | VMAT | Control | 19.13248405 | 0.082209586 | 18.80651854 | 0.369522521 | 1.25350306 | 0.325965518 | 0.278770798 | 0.083061283 | 1.080601366 | Unpaired t-test | 0.302741259 | permutation FDR (250 perm) | 1 | 0.100424376 | FDR correction BH |
| RpS8~A0A0B4K6N1 | VMAT~Control | VMAT | Control | 21.6995207 | 0.105248658 | 21.37429997 | 0.630125467 | 1.252856103 | 0.325220723 | 0.252951371 | 0.281526777 | 0.550480291 | Unpaired t-test | 0.336272489 | permutation FDR (250 perm) | 1 | 0.315878948 | FDR correction BH |
| Fit2~Q9VVD2 | VMAT~Control | VMAT | Control | 15.64634866 | 0.334671477 | 15.32126659 | 0.150375043 | 1.252735699 | 0.325082068 | 0.279259995 | 0.075722118 | 1.12077725 | Unpaired t-test | 0.302181818 | permutation FDR (250 perm) | 1 | 0.092118793 | FDR correction BH |
| EloA~Q9VCP0 | VMAT~Control | VMAT | Control | 18.71995206 | 0.330474032 | 18.39777096 | 0.682862452 | 1.250219231 | 0.3221811 | 0.240565052 | 0.364683039 | 0.438084435 | Unpaired t-test | 0.353252508 | permutation FDR (250 perm) | 1 | 0.401714494 | FDR correction BH |
| CG2967~X2JB38 | VMAT~Control | VMAT | Control | 15.22176257 | 0.269053063 | 14.89976325 | 0.084121195 | 1.250061717 | 0.321999324 | 0.285950114 | 0.028658375 | 1.542748446 | Unpaired t-test | 0.294635129 | permutation FDR (250 perm) | 1 | 0.036924912 | FDR correction BH |
| Dmel\CG4557~Q9W3V2 | VMAT~Control | VMAT | Control | 16.73456205 | 0.083621513 | 16.413169 | 0.121978858 | 1.249536505 | 0.32139305 | 0.301455298 | 0.000662001 | 3.179141277 | Unpaired t-test | 0.277378947 | permutation FDR (250 perm) | 1 | 0.001088234 | FDR correction BH |
| 14-3-3epsilon~P92177-4 | VMAT~Control | VMAT | Control | 15.78355062 | 0.097077232 | 15.46489028 | 0.181058746 | 1.24717191 | 0.31866034 | 0.291846562 | 0.00603762 | 2.219134213 | Unpaired t-test | 0.287812527 | permutation FDR (250 perm) | 1 | 0.008624904 | FDR correction BH |
| GM130~Q9W289 | VMAT~Control | VMAT | Control | 18.59874156 | 0.088386728 | 18.28063302 | 0.13833841 | 1.246694989 | 0.318108545 | 0.296351527 | 0.001482688 | 2.828950227 | Unpaired t-test | 0.282898729 | permutation FDR (250 perm) | 1 | 0.002297727 | FDR correction BH |
| Droj2~Q9VFV9 | VMAT~Control | VMAT | Control | 18.2729062 | 0.132760971 | 17.9549306 | 0.364481959 | 1.246580113 | 0.317975602 | 0.270968592 | 0.096708365 | 1.014535961 | Unpaired t-test | 0.312286588 | permutation FDR (250 perm) | 1 | 0.116148719 | FDR correction BH |
| Rpt6~O18413 | VMAT~Control | VMAT | Control | 16.71001883 | 0.147184923 | 16.39459318 | 0.216900466 | 1.244378741 | 0.315425654 | 0.282329417 | 0.02267021 | 1.644544456 | Unpaired t-test | 0.298684164 | permutation FDR (250 perm) | 1 | 0.029614692 | FDR correction BH |
| RpS28b~X2JEM4 | VMAT~Control | VMAT | Control | 18.48813197 | 0.160893398 | 18.17353618 | 0.228825365 | 1.243663154 | 0.314595785 | 0.279616303 | 0.030659801 | 1.513430668 | Unpaired t-test | 0.301688811 | permutation FDR (250 perm) | 1 | 0.039352874 | FDR correction BH |
| Fip1~Q9VN31 | VMAT~Control | VMAT | Control | 20.16317228 | 0.30221253 | 19.8505451 | 0.608573357 | 1.241967295 | 0.312627183 | 0.239768127 | 0.327811078 | 0.484376374 | Unpaired t-test | 0.354464893 | permutation FDR (250 perm) | 1 | 0.364337714 | FDR correction BH |
| Khc-73~A4UZI0 | VMAT~Control | VMAT | Control | 15.55982451 | 0.285628725 | 15.2480318 | 0.278406675 | 1.241249135 | 0.311792712 | 0.264594756 | 0.111052154 | 0.954473011 | Unpaired t-test | 0.320746177 | permutation FDR (250 perm) | 1 | 0.1325571 | FDR correction BH |
| Rpn10~M9PIG8 | VMAT~Control | VMAT | Control | 15.56921295 | 0.154820594 | 15.2605106 | 0.121140783 | 1.238593132 | 0.308702352 | 0.283756164 | 0.00561845 | 2.250383449 | Unpaired t-test | 0.297176883 | permutation FDR (250 perm) | 1 | 0.008054709 | FDR correction BH |
| Prm~A4V1N9 | VMAT~Control | VMAT | Control | 17.6457373 | 0.196260769 | 17.33838043 | 0.168845027 | 1.237438535 | 0.307356867 | 0.275463279 | 0.024120789 | 1.617608491 | Unpaired t-test | 0.306840909 | permutation FDR (250 perm) | 1 | 0.031387693 | FDR correction BH |
| Gprk1~E1JGX2 | VMAT~Control | VMAT | Control | 15.42348729 | 0.29313061 | 15.11718951 | 0.228290916 | 1.236530455 | 0.306297773 | 0.26265548 | 0.095065688 | 1.021976202 | Unpaired t-test | 0.323405854 | permutation FDR (250 perm) | 1 | 0.114292457 | FDR correction BH |
| Hsc70-5~A0A0B4LFD2 | VMAT~Control | VMAT | Control | 17.63299112 | 0.115739998 | 17.32820059 | 0.154141866 | 1.235239275 | 0.30479053 | 0.280601609 | 0.005395308 | 2.267983792 | Unpaired t-test | 0.300845144 | permutation FDR (250 perm) | 1 | 0.007758433 | FDR correction BH |
| SLIRP1~Q0KHQ1 | VMAT~Control | VMAT | Control | 16.27735108 | 0.149692789 | 15.97669149 | 0.062917223 | 1.231707414 | 0.300659591 | 0.280304573 | 0.002010814 | 2.696628113 | Unpaired t-test | 0.301086139 | permutation FDR (250 perm) | 1 | 0.003053843 | FDR correction BH |
| poe~M9PB68 | VMAT~Control | VMAT | Control | 16.49984958 | 0.123239518 | 16.20092426 | 0.082790856 | 1.230227663 | 0.298925322 | 0.280313555 | 0.001139328 | 2.943351388 | Unpaired t-test | 0.301079143 | permutation FDR (250 perm) | 1 | 0.001809912 | FDR correction BH |
| AspRS~Q7K0E6 | VMAT~Control | VMAT | Control | 16.36641498 | 0.208889114 | 16.07065241 | 0.187497259 | 1.227533645 | 0.295762568 | 0.262776081 | 0.040220247 | 1.395555271 | Unpaired t-test | 0.323239843 | permutation FDR (250 perm) | 1 | 0.050793165 | FDR correction BH |
| Dmel\CG5377~Q9VD00 | VMAT~Control | VMAT | Control | 13.68761122 | 0.396184916 | 13.39273615 | 0.249871065 | 1.226778742 | 0.294875073 | 0.243804271 | 0.189540536 | 0.722297895 | Unpaired t-test | 0.349005674 | permutation FDR (250 perm) | 1 | 0.217541893 | FDR correction BH |
| CG7483~Q9VHS8 | VMAT~Control | VMAT | Control | 14.93753055 | 0.116189768 | 14.64422582 | 0.070734771 | 1.225444149 | 0.293304733 | 0.276485217 | 0.000700704 | 3.154465324 | Unpaired t-test | 0.305498252 | permutation FDR (250 perm) | 1 | 0.001145457 | FDR correction BH |
| ND-B22~Q9VJZ4 | VMAT~Control | VMAT | Control | 17.75387915 | 0.22320345 | 17.46108104 | 0.314533552 | 1.225013896 | 0.292798115 | 0.249724925 | 0.120441691 | 0.919223154 | Unpaired t-test | 0.340745852 | permutation FDR (250 perm) | 1 | 0.142903095 | FDR correction BH |
| AIF~M9PBQ4 | VMAT~Control | VMAT | Control | 17.49827519 | 0.200269627 | 17.20885012 | 0.100199349 | 1.222153141 | 0.289425072 | 0.263078372 | 0.016107682 | 1.792966962 | Unpaired t-test | 0.3228519 | permutation FDR (250 perm) | 1 | 0.021495172 | FDR correction BH |
| or~Q9W1E8 | VMAT~Control | VMAT | Control | 15.81715476 | 0.078447573 | 15.52774079 | 0.094148754 | 1.222143735 | 0.289413969 | 0.274376718 | 0.000357243 | 3.44703668 | Unpaired t-test | 0.308071678 | permutation FDR (250 perm) | 1 | 0.000622464 | FDR correction BH |
| Dmel\CG15561~Q9V9Z8 | VMAT~Control | VMAT | Control | 16.56985958 | 0.257624361 | 16.28075317 | 0.642696132 | 1.221883222 | 0.28910641 | 0.220750187 | 0.372483188 | 0.428893325 | Unpaired t-test | 0.383500218 | permutation FDR (250 perm) | 1 | 0.409598436 | FDR correction BH |
| ND-15~Q7K1C0 | VMAT~Control | VMAT | Control | 18.68540691 | 0.345522014 | 18.39762399 | 0.413608786 | 1.220762816 | 0.287782924 | 0.231891905 | 0.260027613 | 0.584980531 | Unpaired t-test | 0.366323312 | permutation FDR (250 perm) | 1 | 0.293434804 | FDR correction BH |
| ALT~Q7KV27 | VMAT~Control | VMAT | Control | 16.68321257 | 0.329739358 | 16.39670366 | 0.249734189 | 1.219685259 | 0.286508907 | 0.241782954 | 0.152463802 | 0.816833254 | Unpaired t-test | 0.351689354 | permutation FDR (250 perm) | 1 | 0.17758525 | FDR correction BH |
| CG4196~E1JIL1 | VMAT~Control | VMAT | Control | 14.99161565 | 0.13982446 | 14.7054571 | 0.138547157 | 1.219389095 | 0.286158549 | 0.263006178 | 0.008710131 | 2.059975334 | Unpaired t-test | 0.322934032 | permutation FDR (250 perm) | 1 | 0.012168372 | FDR correction BH |
| Dmel\CG12321~Q9VEC2 | VMAT~Control | VMAT | Control | 15.34636085 | 0.117226471 | 15.06104714 | 0.20455227 | 1.218675237 | 0.285313715 | 0.258100634 | 0.02208326 | 1.655936811 | Unpaired t-test | 0.329375273 | permutation FDR (250 perm) | 1 | 0.028912121 | FDR correction BH |
| eIF3l~Q9VTU4 | VMAT~Control | VMAT | Control | 17.77277357 | 0.102326901 | 17.48851813 | 0.083912936 | 1.217781624 | 0.284255449 | 0.268372779 | 0.000720243 | 3.14252076 | Unpaired t-test | 0.315688947 | permutation FDR (250 perm) | 1 | 0.001175765 | FDR correction BH |
| eIF2alpha~X2JFR6 | VMAT~Control | VMAT | Control | 17.86910629 | 0.116598667 | 17.58705844 | 0.06832297 | 1.215919609 | 0.282047848 | 0.265973173 | 0.000885116 | 3.052999578 | Unpaired t-test | 0.318830931 | permutation FDR (250 perm) | 1 | 0.001425146 | FDR correction BH |
| Ptp99A~A0A0B4JDF4 | VMAT~Control | VMAT | Control | 14.52602628 | 1.849165566 | 14.24572241 | 1.613890276 | 1.214450655 | 0.280303872 | 0.133628281 | 0.803613054 | 0.094953017 | Unpaired t-test | 0.552221645 | permutation FDR (250 perm) | 1 | 0.826072109 | FDR correction BH |
| Dmel\CG14367~Q9VFS4 | VMAT~Control | VMAT | Control | 18.37402626 | 0.180419459 | 18.0946081 | 0.191806094 | 1.213705303 | 0.279418166 | 0.249979769 | 0.039095721 | 1.407870776 | Unpaired t-test | 0.340420961 | permutation FDR (250 perm) | 1 | 0.049532469 | FDR correction BH |
| CaMKII~L0MLR4 | VMAT~Control | VMAT | Control | 14.73445147 | 0.187990161 | 14.45618123 | 0.140566552 | 1.212739961 | 0.278270237 | 0.251833857 | 0.024279202 | 1.61476559 | Unpaired t-test | 0.337882969 | permutation FDR (250 perm) | 1 | 0.031576377 | FDR correction BH |
| Ote~P20240 | VMAT~Control | VMAT | Control | 18.63886691 | 0.119983133 | 18.36089897 | 0.229009425 | 1.212485875 | 0.27796794 | 0.249159795 | 0.037053434 | 1.43117154 | Unpaired t-test | 0.341573799 | permutation FDR (250 perm) | 1 | 0.047147991 | FDR correction BH |
| RpS27A~P15357 | VMAT~Control | VMAT | Control | 22.0559157 | 0.113926853 | 21.77798426 | 0.403893654 | 1.212455196 | 0.277931436 | 0.234013052 | 0.169433664 | 0.771000298 | Unpaired t-test | 0.363015251 | permutation FDR (250 perm) | 1 | 0.195897272 | FDR correction BH |
| RpL27~Q9VBN5 | VMAT~Control | VMAT | Control | 18.21790995 | 0.146467728 | 17.94002476 | 0.672050932 | 1.212416338 | 0.277885198 | 0.212514578 | 0.387566936 | 0.41165328 | Unpaired t-test | 0.396438261 | permutation FDR (250 perm) | 1 | 0.424735832 | FDR correction BH |
| Hem~P55162 | VMAT~Control | VMAT | Control | 15.26836481 | 0.159339486 | 14.99101307 | 0.104351505 | 1.211968107 | 0.277351735 | 0.255581278 | 0.008631499 | 2.063913752 | Unpaired t-test | 0.332816077 | permutation FDR (250 perm) | 1 | 0.012094375 | FDR correction BH |
| Fs(2)Ket~M9NDC3 | VMAT~Control | VMAT | Control | 15.63313094 | 0.189918897 | 15.35758945 | 0.081524127 | 1.210448325 | 0.275541491 | 0.252228341 | 0.013781201 | 1.86071294 | Unpaired t-test | 0.337320524 | permutation FDR (250 perm) | 1 | 0.01862904 | FDR correction BH |
| Sgs5~P07701 | VMAT~Control | VMAT | Control | 17.81585694 | 0.380742621 | 17.54154157 | 0.160262232 | 1.209420023 | 0.274315369 | 0.23154008 | 0.168407794 | 0.773637812 | Unpaired t-test | 0.366828758 | permutation FDR (250 perm) | 1 | 0.194806854 | FDR correction BH |
| UbcE2M~M9PBW0 | VMAT~Control | VMAT | Control | 15.73923637 | 0.148075608 | 15.46562387 | 0.115524217 | 1.208830945 | 0.273612498 | 0.252412207 | 0.008608296 | 2.065082807 | Unpaired t-test | 0.337044541 | permutation FDR (250 perm) | 1 | 0.012069046 | FDR correction BH |
| RpL29~Q24154 | VMAT~Control | VMAT | Control | 17.9781262 | 0.161072682 | 17.70464961 | 0.578722603 | 1.208717076 | 0.273476592 | 0.215565034 | 0.332690086 | 0.477960141 | Unpaired t-test | 0.391935596 | permutation FDR (250 perm) | 1 | 0.368889525 | FDR correction BH |
| trio~Q8IRI5 | VMAT~Control | VMAT | Control | 16.28425669 | 0.117782334 | 16.01123256 | 0.150453007 | 1.208338055 | 0.273024132 | 0.251530749 | 0.009570599 | 2.019060872 | Unpaired t-test | 0.338263755 | permutation FDR (250 perm) | 1 | 0.013260265 | FDR correction BH |
| Acox3~Q9VMD7 | VMAT~Control | VMAT | Control | 15.88863627 | 0.254021985 | 15.61624153 | 0.240160323 | 1.207811021 | 0.272394743 | 0.235567196 | 0.112054478 | 0.950570783 | Unpaired t-test | 0.360760244 | permutation FDR (250 perm) | 1 | 0.133557591 | FDR correction BH |
| RpS26~M9PG47 | VMAT~Control | VMAT | Control | 22.17976043 | 0.15041177 | 21.90860848 | 0.371460443 | 1.206771018 | 0.271151954 | 0.229940969 | 0.161242576 | 0.792520273 | Unpaired t-test | 0.369347277 | permutation FDR (250 perm) | 1 | 0.186977844 | FDR correction BH |
| Dab~M9NDT7 | VMAT~Control | VMAT | Control | 17.8047949 | 0.117277259 | 17.53384851 | 0.187732589 | 1.206599082 | 0.270946389 | 0.24654073 | 0.020938779 | 1.679048644 | Unpaired t-test | 0.345275109 | permutation FDR (250 perm) | 1 | 0.027551641 | FDR correction BH |
| Dyb~Q0E9B2 | VMAT~Control | VMAT | Control | 17.05698754 | 0.13338711 | 16.78929634 | 0.158976775 | 1.203879676 | 0.267691206 | 0.244957431 | 0.016262884 | 1.788802435 | Unpaired t-test | 0.347292885 | permutation FDR (250 perm) | 1 | 0.021653184 | FDR correction BH |
| Vap33~Q9W4N8 | VMAT~Control | VMAT | Control | 22.64623213 | 0.151390524 | 22.37999134 | 0.120861009 | 1.202669963 | 0.266240792 | 0.245014431 | 0.011776408 | 1.928987173 | Unpaired t-test | 0.347181144 | permutation FDR (250 perm) | 1 | 0.016089184 | FDR correction BH |
| Dmel\CG11807~Q7K490 | VMAT~Control | VMAT | Control | 14.49014101 | 1.510941342 | 14.22714141 | 1.311932724 | 1.199971049 | 0.262999599 | 0.138794365 | 0.77485026 | 0.110782217 | Unpaired t-test | 0.540585535 | permutation FDR (250 perm) | 1 | 0.798947662 | FDR correction BH |
| RpLP0~M9PG76 | VMAT~Control | VMAT | Control | 16.87212748 | 0.117837124 | 16.60947597 | 0.18459454 | 1.19968156 | 0.262651512 | 0.239222207 | 0.023022549 | 1.637846596 | Unpaired t-test | 0.355277802 | permutation FDR (250 perm) | 1 | 0.030058281 | FDR correction BH |
| Atg16~Q86BR6 | VMAT~Control | VMAT | Control | 15.36180361 | 0.710415772 | 15.10048055 | 0.707039208 | 1.198577383 | 0.261323056 | 0.18044183 | 0.572806083 | 0.241992378 | Unpaired t-test | 0.454166015 | permutation FDR (250 perm) | 1 | 0.607380865 | FDR correction BH |
| Dmel\CG6983~Q9VSK4 | VMAT~Control | VMAT | Control | 16.95883844 | 0.060448667 | 16.69769993 | 0.113928769 | 1.198424071 | 0.261138506 | 0.246897908 | 0.001095574 | 2.960358379 | Unpaired t-test | 0.34481048 | permutation FDR (250 perm) | 1 | 0.001743733 | FDR correction BH |
| Dmel\CG5171~Q9VM19 | VMAT~Control | VMAT | Control | 14.44775648 | 1.146505769 | 14.18891221 | 0.593596275 | 1.196519806 | 0.258844278 | 0.164097717 | 0.663487038 | 0.178167557 | Unpaired t-test | 0.486660877 | permutation FDR (250 perm) | 1 | 0.693230576 | FDR correction BH |
| Nop60B~E1JGV6 | VMAT~Control | VMAT | Control | 18.52229781 | 0.217261139 | 18.26506931 | 0.413736066 | 1.195180484 | 0.257228496 | 0.21276348 | 0.246551122 | 0.608093017 | Unpaired t-test | 0.396008696 | permutation FDR (250 perm) | 1 | 0.279836712 | FDR correction BH |
| Gapdh2~M9PJN8 | VMAT~Control | VMAT | Control | 20.75262103 | 0.114933521 | 20.49617907 | 0.239987106 | 1.194529072 | 0.256441966 | 0.229170911 | 0.056585361 | 1.247295909 | Unpaired t-test | 0.370530719 | permutation FDR (250 perm) | 1 | 0.070254188 | FDR correction BH |
| 5PtaseI~Q8IMU3 | VMAT~Control | VMAT | Control | 15.35492565 | 0.278368307 | 15.09880972 | 0.292085254 | 1.194259151 | 0.256115932 | 0.216965505 | 0.186207115 | 0.73000373 | Unpaired t-test | 0.389784937 | permutation FDR (250 perm) | 1 | 0.213924621 | FDR correction BH |
| 152078_at~Q9VMC6 | VMAT~Control | VMAT | Control | 15.54019899 | 0.252824061 | 15.28765295 | 0.119169595 | 1.191307661 | 0.252546044 | 0.224485947 | 0.07093114 | 1.149163063 | Unpaired t-test | 0.377864924 | permutation FDR (250 perm) | 1 | 0.086603684 | FDR correction BH |
| PRL-1~O61722 | VMAT~Control | VMAT | Control | 17.26836237 | 0.022013211 | 17.01782418 | 0.314562058 | 1.189650827 | 0.250538192 | 0.219573792 | 0.106009309 | 0.974655998 | Unpaired t-test | 0.385443622 | permutation FDR (250 perm) | 1 | 0.126737386 | FDR correction BH |
| EG:BACR7A4.14~Q9U1L2 | VMAT~Control | VMAT | Control | 13.66896914 | 0.646343916 | 13.42105831 | 0.482400016 | 1.187486267 | 0.24791083 | 0.182195537 | 0.507500309 | 0.294563689 | Unpaired t-test | 0.450883478 | permutation FDR (250 perm) | 1 | 0.54351944 | FDR correction BH |
| Dmel\CG5065~E1JH83 | VMAT~Control | VMAT | Control | 15.3343533 | 0.071169941 | 15.0877406 | 0.189716356 | 1.186418256 | 0.246612702 | 0.22612213 | 0.021505792 | 1.667444556 | Unpaired t-test | 0.375274946 | permutation FDR (250 perm) | 1 | 0.028250354 | FDR correction BH |
| Gapdh1~P07486 | VMAT~Control | VMAT | Control | 21.02015183 | 0.160748474 | 20.77596753 | 0.282268605 | 1.184422909 | 0.2441843 | 0.213211278 | 0.123697918 | 0.90763761 | Unpaired t-test | 0.395290435 | permutation FDR (250 perm) | 1 | 0.14632407 | FDR correction BH |
| Ankle2~Q8MQX9-2 | VMAT~Control | VMAT | Control | 13.88721645 | 0.622657874 | 13.64472863 | 0.653136089 | 1.183030953 | 0.242487821 | 0.172766687 | 0.561293564 | 0.250809938 | Unpaired t-test | 0.46916073 | permutation FDR (250 perm) | 1 | 0.597596132 | FDR correction BH |
| Dmel\CG15210~Q9VZ67 | VMAT~Control | VMAT | Control | 14.22858869 | 0.917188506 | 13.98614329 | 0.436347996 | 1.182996168 | 0.242445401 | 0.166717121 | 0.60517643 | 0.218117995 | Unpaired t-test | 0.481175858 | permutation FDR (250 perm) | 1 | 0.639975435 | FDR correction BH |
| p120ctn~Q7PLI0 | VMAT~Control | VMAT | Control | 17.1292295 | 0.103740623 | 16.88746933 | 0.295420882 | 1.182434417 | 0.241760168 | 0.212065591 | 0.114953692 | 0.939477074 | Unpaired t-test | 0.397224348 | permutation FDR (250 perm) | 1 | 0.136736226 | FDR correction BH |
| hbt~Q9VK03 | VMAT~Control | VMAT | Control | 14.75461534 | 0.270642275 | 14.51315802 | 0.216065761 | 1.182186224 | 0.241457313 | 0.20907654 | 0.150045062 | 0.823778294 | Unpaired t-test | 0.402234783 | permutation FDR (250 perm) | 1 | 0.175114564 | FDR correction BH |
| kcc~A0A0B4LGD3 | VMAT~Control | VMAT | Control | 15.68591743 | 0.29269441 | 15.44492342 | 0.336303888 | 1.181806644 | 0.240994015 | 0.200931455 | 0.254585043 | 0.594167116 | Unpaired t-test | 0.41628 | permutation FDR (250 perm) | 1 | 0.288121726 | FDR correction BH |
| inaD~Q24008 | VMAT~Control | VMAT | Control | 21.24333549 | 0.092061722 | 21.00243758 | 0.257696881 | 1.181727922 | 0.240897911 | 0.214631529 | 0.077342241 | 1.111583249 | Unpaired t-test | 0.393347258 | permutation FDR (250 perm) | 1 | 0.093944084 | FDR correction BH |
| GlcAT-P~A4V1U5 | VMAT~Control | VMAT | Control | 16.47328819 | 0.096767372 | 16.23353356 | 0.150296509 | 1.180791818 | 0.23975463 | 0.222007118 | 0.013363328 | 1.87408536 | Unpaired t-test | 0.381682927 | permutation FDR (250 perm) | 1 | 0.018130994 | FDR correction BH |
| RpL6~Q9V9W2 | VMAT~Control | VMAT | Control | 22.35417654 | 0.134295666 | 22.1152642 | 0.639003016 | 1.180102644 | 0.238912349 | 0.18491471 | 0.432332004 | 0.364182614 | Unpaired t-test | 0.445794783 | permutation FDR (250 perm) | 1 | 0.470725965 | FDR correction BH |
| Srp72~Q9VDK7 | VMAT~Control | VMAT | Control | 17.00265398 | 0.086190595 | 16.76465726 | 0.20973176 | 1.179353912 | 0.237996721 | 0.216084394 | 0.040851814 | 1.388788651 | Unpaired t-test | 0.391251197 | permutation FDR (250 perm) | 1 | 0.051563094 | FDR correction BH |
| SdhA~Q94523 | VMAT~Control | VMAT | Control | 17.92637098 | 0.114104016 | 17.6894479 | 0.228605875 | 1.178476572 | 0.236923078 | 0.212627575 | 0.064903262 | 1.187733478 | Unpaired t-test | 0.396252174 | permutation FDR (250 perm) | 1 | 0.079740228 | FDR correction BH |
| SelR~D3DMP0 | VMAT~Control | VMAT | Control | 17.25911505 | 0.195809782 | 17.02317131 | 0.285735537 | 1.177676865 | 0.235943742 | 0.204296133 | 0.158710952 | 0.799393104 | Unpaired t-test | 0.410422609 | permutation FDR (250 perm) | 1 | 0.184587737 | FDR correction BH |
| Ube4B~Q9VK44 | VMAT~Control | VMAT | Control | 16.18753957 | 0.087884476 | 15.95256557 | 0.076979238 | 1.176885521 | 0.234973993 | 0.223306578 | 0.00114798 | 2.940065549 | Unpaired t-test | 0.379694989 | permutation FDR (250 perm) | 1 | 0.001820988 | FDR correction BH |
| RpL32~P04359 | VMAT~Control | VMAT | Control | 20.97261857 | 0.10061739 | 20.73903847 | 0.655491048 | 1.175748996 | 0.233580101 | 0.180151222 | 0.449204689 | 0.347555719 | Unpaired t-test | 0.454755324 | permutation FDR (250 perm) | 1 | 0.486845229 | FDR correction BH |
| Acp26Aa~P10333 | VMAT~Control | VMAT | Control | 16.37813735 | 0.381876264 | 16.14556394 | 0.311937281 | 1.174928859 | 0.232573406 | 0.190553499 | 0.316385643 | 0.499783232 | Unpaired t-test | 0.43508 | permutation FDR (250 perm) | 1 | 0.352137969 | FDR correction BH |
| Atg17~Q7KTS2 | VMAT~Control | VMAT | Control | 16.03194994 | 0.291044539 | 15.80455721 | 0.380681552 | 1.170717285 | 0.227392724 | 0.187262183 | 0.313604146 | 0.503618204 | Unpaired t-test | 0.441168696 | permutation FDR (250 perm) | 1 | 0.349207266 | FDR correction BH |
| Marf~Q7YU24 | VMAT~Control | VMAT | Control | 16.56409277 | 0.088170392 | 16.33723066 | 0.099286256 | 1.170286789 | 0.226862118 | 0.21414549 | 0.003371836 | 2.472133598 | Unpaired t-test | 0.393964332 | permutation FDR (250 perm) | 1 | 0.004963916 | FDR correction BH |
| Cmpk~Q9VBI2 | VMAT~Control | VMAT | Control | 17.30032805 | 0.143865274 | 17.07504172 | 0.150732639 | 1.169009236 | 0.225286328 | 0.206082462 | 0.036208929 | 1.441184321 | Unpaired t-test | 0.407387826 | permutation FDR (250 perm) | 1 | 0.046098334 | FDR correction BH |
| Tctp~A0A126GUT1 | VMAT~Control | VMAT | Control | 15.81784687 | 0.20702472 | 15.5946958 | 0.317974241 | 1.167280318 | 0.223151061 | 0.190778587 | 0.217839102 | 0.661864162 | Unpaired t-test | 0.434688696 | permutation FDR (250 perm) | 1 | 0.24844634 | FDR correction BH |
| Ebp~A0A0B4KGF3 | VMAT~Control | VMAT | Control | 16.80442483 | 0.718488645 | 16.58142365 | 0.764106936 | 1.167159051 | 0.223001174 | 0.151798571 | 0.644700753 | 0.190641822 | Unpaired t-test | 0.512093628 | permutation FDR (250 perm) | 1 | 0.675400789 | FDR correction BH |
| rdgA~Q09103-3 | VMAT~Control | VMAT | Control | 15.71087511 | 0.094693518 | 15.48943321 | 0.135788659 | 1.165898261 | 0.221441901 | 0.206177672 | 0.01354886 | 1.868097256 | Unpaired t-test | 0.407222609 | permutation FDR (250 perm) | 1 | 0.018350987 | FDR correction BH |
| Rpt2~A0A0B4LIJ0 | VMAT~Control | VMAT | Control | 15.86269318 | 0.121345354 | 15.64134198 | 0.197789213 | 1.16582496 | 0.221351194 | 0.200540314 | 0.058722615 | 1.231194615 | Unpaired t-test | 0.416853913 | permutation FDR (250 perm) | 1 | 0.072677726 | FDR correction BH |
| CG6066~Q9VB74 | VMAT~Control | VMAT | Control | 15.418027 | 0.662070433 | 15.19773675 | 0.76610625 | 1.164967938 | 0.22029025 | 0.151628811 | 0.63711275 | 0.195783704 | Unpaired t-test | 0.512449068 | permutation FDR (250 perm) | 1 | 0.66893997 | FDR correction BH |
| PIP5K59B~Q9W1Y3 | VMAT~Control | VMAT | Control | 15.97792148 | 0.409448472 | 15.75830628 | 0.349248001 | 1.164422967 | 0.2196152 | 0.177012678 | 0.382978784 | 0.416825285 | Unpaired t-test | 0.461103868 | permutation FDR (250 perm) | 1 | 0.420490698 | FDR correction BH |
| Rbp1-like~Q9VYD8 | VMAT~Control | VMAT | Control | 17.47038551 | 0.12160142 | 17.25177286 | 0.442636746 | 1.163614068 | 0.218612644 | 0.18137805 | 0.311953159 | 0.505910613 | Unpaired t-test | 0.452490435 | permutation FDR (250 perm) | 1 | 0.347862499 | FDR correction BH |
| RpL35~Q9W499 | VMAT~Control | VMAT | Control | 19.27719929 | 0.141779351 | 19.06115049 | 0.72806979 | 1.161548023 | 0.216048803 | 0.162233043 | 0.529534245 | 0.276105949 | Unpaired t-test | 0.490457664 | permutation FDR (250 perm) | 1 | 0.565830055 | FDR correction BH |
| Rm62~E1JJ68 | VMAT~Control | VMAT | Control | 16.69452949 | 0.255744575 | 16.47964267 | 0.192867122 | 1.160612857 | 0.214886815 | 0.187961329 | 0.16448668 | 0.783869266 | Unpaired t-test | 0.439848696 | permutation FDR (250 perm) | 1 | 0.190458261 | FDR correction BH |
| CG10167~Q9VP69 | VMAT~Control | VMAT | Control | 16.81021632 | 0.095502524 | 16.59554172 | 0.225542288 | 1.160442142 | 0.214674594 | 0.193481513 | 0.078453898 | 1.105385472 | Unpaired t-test | 0.429883478 | permutation FDR (250 perm) | 1 | 0.095098083 | FDR correction BH |
| Dip-B~Q9VFQ9 | VMAT~Control | VMAT | Control | 17.40245041 | 0.164589429 | 17.19028741 | 0.085066577 | 1.158423685 | 0.212163006 | 0.195929012 | 0.028343616 | 1.54754475 | Unpaired t-test | 0.425257391 | permutation FDR (250 perm) | 1 | 0.036541654 | FDR correction BH |
| bor~Q9VEX6 | VMAT~Control | VMAT | Control | 13.80785431 | 0.897807602 | 13.59702263 | 0.334216833 | 1.15735518 | 0.210831681 | 0.147596835 | 0.633269733 | 0.198411268 | Unpaired t-test | 0.520930273 | permutation FDR (250 perm) | 1 | 0.665900463 | FDR correction BH |
| Uch~P35122 | VMAT~Control | VMAT | Control | 16.95045252 | 0.07564417 | 16.73974521 | 0.067812416 | 1.157255411 | 0.210707308 | 0.201550369 | 0.000925082 | 3.033819789 | Unpaired t-test | 0.415182609 | permutation FDR (250 perm) | 1 | 0.001485432 | FDR correction BH |
| miple2~Q9W0S4 | VMAT~Control | VMAT | Control | 16.18620475 | 0.191422076 | 15.97718083 | 0.947795418 | 1.155905874 | 0.209023923 | 0.14592308 | 0.639237982 | 0.194337428 | Unpaired t-test | 0.524739714 | permutation FDR (250 perm) | 1 | 0.67087214 | FDR correction BH |
| Syn2~Q0E952 | VMAT~Control | VMAT | Control | 15.58009433 | 0.110813306 | 15.37179572 | 0.199048549 | 1.155324885 | 0.208298604 | 0.189038905 | 0.068132044 | 1.166648579 | Unpaired t-test | 0.437855652 | permutation FDR (250 perm) | 1 | 0.083489241 | FDR correction BH |
| ric8a~X2JEL4 | VMAT~Control | VMAT | Control | 18.03749162 | 0.19090597 | 17.8298908 | 0.333477268 | 1.154766231 | 0.207600824 | 0.177157357 | 0.254811521 | 0.593780941 | Unpaired t-test | 0.460794437 | permutation FDR (250 perm) | 1 | 0.288239462 | FDR correction BH |
| Rlip~A0A0B4KHU1 | VMAT~Control | VMAT | Control | 17.69673955 | 0.104408412 | 17.48945178 | 0.519399349 | 1.154515681 | 0.207287768 | 0.167582589 | 0.402155826 | 0.395605635 | Unpaired t-test | 0.479284412 | permutation FDR (250 perm) | 1 | 0.439088504 | FDR correction BH |
| Hao~A1Z8D3 | VMAT~Control | VMAT | Control | 14.5481655 | 0.367407425 | 14.34329968 | 0.277645239 | 1.152579145 | 0.20486582 | 0.16987933 | 0.343322818 | 0.464297332 | Unpaired t-test | 0.474872285 | permutation FDR (250 perm) | 1 | 0.3803209 | FDR correction BH |
| FASN3~Q7PLB8 | VMAT~Control | VMAT | Control | 17.32984189 | 0.144070538 | 17.128961 | 0.097361502 | 1.149399949 | 0.200880891 | 0.186386851 | 0.027264421 | 1.564403722 | Unpaired t-test | 0.442895652 | permutation FDR (250 perm) | 1 | 0.035225273 | FDR correction BH |
| unc-45~Q9VHW4 | VMAT~Control | VMAT | Control | 14.33474595 | 0.674220779 | 14.13485423 | 0.190678839 | 1.148612146 | 0.199891723 | 0.152200222 | 0.537855736 | 0.269334195 | Unpaired t-test | 0.511424111 | permutation FDR (250 perm) | 1 | 0.574200636 | FDR correction BH |
| MED19~X2J969 | VMAT~Control | VMAT | Control | 16.78978012 | 0.270678909 | 16.5903186 | 0.537974413 | 1.148269684 | 0.199461515 | 0.157139657 | 0.475969237 | 0.322421115 | Unpaired t-test | 0.501081597 | permutation FDR (250 perm) | 1 | 0.513959443 | FDR correction BH |
| Hn~Q59E23 | VMAT~Control | VMAT | Control | 16.79251153 | 0.144369564 | 16.59440633 | 0.164267266 | 1.147190677 | 0.198105205 | 0.1804562 | 0.070323024 | 1.152902462 | Unpaired t-test | 0.454152108 | permutation FDR (250 perm) | 1 | 0.08590576 | FDR correction BH |
| lark~Q94901 | VMAT~Control | VMAT | Control | 15.65707833 | 0.150686904 | 15.46008487 | 0.259459251 | 1.146306993 | 0.196993464 | 0.173687519 | 0.172816447 | 0.762414927 | Unpaired t-test | 0.467400521 | permutation FDR (250 perm) | 1 | 0.199612324 | FDR correction BH |
| eIF3h~Q9U9Q4 | VMAT~Control | VMAT | Control | 16.83357799 | 0.100460966 | 16.6383303 | 0.140447122 | 1.144920712 | 0.195247693 | 0.181250767 | 0.02995673 | 1.5235056 | Unpaired t-test | 0.452686957 | permutation FDR (250 perm) | 1 | 0.038492435 | FDR correction BH |
| CCT6~Q9VXQ5 | VMAT~Control | VMAT | Control | 18.05060729 | 0.092249942 | 17.85548321 | 0.038468263 | 1.144822622 | 0.195124085 | 0.186775468 | 0.00140937 | 2.850975032 | Unpaired t-test | 0.442092174 | permutation FDR (250 perm) | 1 | 0.002194217 | FDR correction BH |
| BcDNA:SD01241~Q9VIJ3 | VMAT~Control | VMAT | Control | 17.51353284 | 0.179104309 | 17.32108108 | 0.397807202 | 1.142704015 | 0.192451763 | 0.161033412 | 0.347200095 | 0.459420164 | Unpaired t-test | 0.492988276 | permutation FDR (250 perm) | 1 | 0.38407379 | FDR correction BH |
| Mkk4~O61444 | VMAT~Control | VMAT | Control | 16.8232067 | 0.114927504 | 16.63346233 | 0.066310448 | 1.140561607 | 0.189744375 | 0.179115872 | 0.009529624 | 2.020924246 | Unpaired t-test | 0.456818774 | permutation FDR (250 perm) | 1 | 0.013226848 | FDR correction BH |
| Pax~Q966T5 | VMAT~Control | VMAT | Control | 16.41570661 | 0.195818286 | 16.22756432 | 0.359390179 | 1.139295742 | 0.188142295 | 0.1590338 | 0.32821235 | 0.483845081 | Unpaired t-test | 0.497005208 | permutation FDR (250 perm) | 1 | 0.364537785 | FDR correction BH |
| CG5382~Q9VD26 | VMAT~Control | VMAT | Control | 17.18633369 | 0.051091842 | 16.99843263 | 0.262897419 | 1.139105261 | 0.187901068 | 0.167803125 | 0.1477577 | 0.830449879 | Unpaired t-test | 0.47883109 | permutation FDR (250 perm) | 1 | 0.172889751 | FDR correction BH |
| Hrb87F~E1JIK0 | VMAT~Control | VMAT | Control | 14.11955796 | 0.052608716 | 13.933592 | 0.289429774 | 1.137578387 | 0.185965961 | 0.164345084 | 0.187854108 | 0.726179303 | Unpaired t-test | 0.486136344 | permutation FDR (250 perm) | 1 | 0.215711498 | FDR correction BH |
| RpL34b~Q9VHE5 | VMAT~Control | VMAT | Control | 20.05859752 | 0.235425363 | 19.87400633 | 0.730997361 | 1.136494885 | 0.184591191 | 0.137401081 | 0.602700409 | 0.219898514 | Unpaired t-test | 0.543708965 | permutation FDR (250 perm) | 1 | 0.637930199 | FDR correction BH |
| Snrk~Q0E981 | VMAT~Control | VMAT | Control | 14.78318379 | 0.072223863 | 14.5997427 | 0.116562891 | 1.135589244 | 0.18344109 | 0.172841735 | 0.0135426 | 1.868297958 | Unpaired t-test | 0.468999131 | permutation FDR (250 perm) | 1 | 0.018350987 | FDR correction BH |
| Dmel\CG34325~X2JFG6 | VMAT~Control | VMAT | Control | 15.3050827 | 0.091891014 | 15.12529348 | 0.158436306 | 1.132718378 | 0.179789216 | 0.16617765 | 0.052889571 | 1.27662996 | Unpaired t-test | 0.482330873 | permutation FDR (250 perm) | 1 | 0.065839264 | FDR correction BH |
| Srp68~Q9VSS2 | VMAT~Control | VMAT | Control | 17.06438935 | 0.222377844 | 16.8863649 | 0.259802305 | 1.131333637 | 0.178024451 | 0.154409477 | 0.271429827 | 0.56634243 | Unpaired t-test | 0.506503036 | permutation FDR (250 perm) | 1 | 0.304986068 | FDR correction BH |
| RhoGAPp190~M9PHG0 | VMAT~Control | VMAT | Control | 17.44451864 | 0.080236441 | 17.26753406 | 0.145308032 | 1.130518482 | 0.176984578 | 0.164754446 | 0.038336936 | 1.416382598 | Unpaired t-test | 0.485259227 | permutation FDR (250 perm) | 1 | 0.048702184 | FDR correction BH |
| ND-B14.5A~O97418 | VMAT~Control | VMAT | Control | 17.7980065 | 0.302435907 | 17.62128916 | 0.478551717 | 1.13030909 | 0.17671734 | 0.141016078 | 0.501076339 | 0.300096104 | Unpaired t-test | 0.535547856 | permutation FDR (250 perm) | 1 | 0.537372985 | FDR correction BH |
| Cdk12~M9PG50 | VMAT~Control | VMAT | Control | 16.53020668 | 0.852787501 | 16.35396162 | 0.876251285 | 1.129939133 | 0.176245061 | 0.113940222 | 0.75385588 | 0.122711673 | Unpaired t-test | 0.598938282 | permutation FDR (250 perm) | 1 | 0.779691011 | FDR correction BH |
| 14-3-3zeta~A0A0B4KEH0 | VMAT~Control | VMAT | Control | 17.39894877 | 0.111864109 | 17.22458987 | 0.15630824 | 1.128462833 | 0.174358903 | 0.160557349 | 0.069995436 | 1.154930274 | Unpaired t-test | 0.494014763 | permutation FDR (250 perm) | 1 | 0.085549978 | FDR correction BH |
| ValRS~Q0E993 | VMAT~Control | VMAT | Control | 15.96403684 | 0.114183699 | 15.79044615 | 0.097562746 | 1.127862107 | 0.173590694 | 0.162665117 | 0.027205481 | 1.565343591 | Unpaired t-test | 0.489587495 | permutation FDR (250 perm) | 1 | 0.035168425 | FDR correction BH |
| Tpi~P29613 | VMAT~Control | VMAT | Control | 17.5882952 | 0.377776021 | 17.41546526 | 0.599025351 | 1.127267519 | 0.172829931 | 0.13125829 | 0.597238359 | 0.223852306 | Unpaired t-test | 0.557440069 | permutation FDR (250 perm) | 1 | 0.632717866 | FDR correction BH |
| Rpn9~Q7KMP8 | VMAT~Control | VMAT | Control | 13.85362505 | 0.76968751 | 13.68084724 | 0.190515769 | 1.127226798 | 0.172777814 | 0.127548707 | 0.636589758 | 0.196140353 | Unpaired t-test | 0.566093426 | permutation FDR (250 perm) | 1 | 0.668689107 | FDR correction BH |
| Rbp1~Q02427-1 | VMAT~Control | VMAT | Control | 20.97941337 | 0.108419464 | 20.81042706 | 0.480989365 | 1.12426825 | 0.168986304 | 0.138456398 | 0.461163548 | 0.336145029 | Unpaired t-test | 0.54134777 | permutation FDR (250 perm) | 1 | 0.499116777 | FDR correction BH |
| AnxB9~A0A0B4KH34 | VMAT~Control | VMAT | Control | 16.6849821 | 0.177857044 | 16.51615397 | 0.186845076 | 1.124144994 | 0.168828128 | 0.151365953 | 0.174058345 | 0.759305151 | Unpaired t-test | 0.513040312 | permutation FDR (250 perm) | 1 | 0.200849678 | FDR correction BH |
| Nap1~Q9W1G7 | VMAT~Control | VMAT | Control | 17.79576468 | 0.212994252 | 17.62954684 | 0.282883808 | 1.1221129 | 0.166217838 | 0.143494083 | 0.318597871 | 0.49675713 | Unpaired t-test | 0.530054569 | permutation FDR (250 perm) | 1 | 0.354432604 | FDR correction BH |
| Taf2~E8NHB1 | VMAT~Control | VMAT | Control | 18.65522045 | 0.341588327 | 18.4894974 | 0.660689767 | 1.121728123 | 0.165723048 | 0.124358447 | 0.629097099 | 0.201282318 | Unpaired t-test | 0.573392131 | permutation FDR (250 perm) | 1 | 0.662296319 | FDR correction BH |
| Ufl1~A0A0B4K6K7 | VMAT~Control | VMAT | Control | 16.18374275 | 0.305930587 | 16.0193468 | 0.231423601 | 1.120696745 | 0.164395945 | 0.140323219 | 0.360513277 | 0.443078736 | Unpaired t-test | 0.537124296 | permutation FDR (250 perm) | 1 | 0.397307235 | FDR correction BH |
| Dmel\CG15432~Q9VQZ5 | VMAT~Control | VMAT | Control | 17.28179188 | 0.212735093 | 17.11783101 | 0.641760738 | 1.120358824 | 0.163960866 | 0.125895028 | 0.599509378 | 0.222204019 | Unpaired t-test | 0.569774319 | permutation FDR (250 perm) | 1 | 0.634838091 | FDR correction BH |
| Klp10A~A4V4A1 | VMAT~Control | VMAT | Control | 13.4074122 | 0.432838084 | 13.24419769 | 0.281147097 | 1.11977937 | 0.163214507 | 0.13260617 | 0.495649673 | 0.304825176 | Unpaired t-test | 0.55452987 | permutation FDR (250 perm) | 1 | 0.531795501 | FDR correction BH |
| CG30386~Q7KVP6 | VMAT~Control | VMAT | Control | 17.64985999 | 0.234069608 | 17.48673587 | 0.133927756 | 1.119709215 | 0.163124118 | 0.145568182 | 0.205983139 | 0.686168327 | Unpaired t-test | 0.525411867 | permutation FDR (250 perm) | 1 | 0.235495051 | FDR correction BH |
| GstE13~Q7JVI6 | VMAT~Control | VMAT | Control | 16.26399407 | 0.105085769 | 16.10196617 | 0.180039265 | 1.118858736 | 0.162027897 | 0.148210547 | 0.112852642 | 0.947488269 | Unpaired t-test | 0.519740035 | permutation FDR (250 perm) | 1 | 0.134372848 | FDR correction BH |

| | | | | | | | | | | | | | | | | | | |
|---|---|---|---|---|---|---|---|---|---|---|---|---|---|---|---|---|---|---|
| Tudor-SN~Q9W0S7 | VMAT~Control | VMAT | Control | 18.1329729 | 0.132685009 | 17.97232048 | 0.047599895 | 1.117792521 | 0.160652427 | 0.151125284 | 0.028944959 | 1.538427063 | Unpaired t-test | 0.513541396 | permutation FDR (250 perm) | 1 | 0.037253381 | FDR correction BH |
| CycT~M9PI89 | VMAT~Control | VMAT | Control | 18.51318599 | 0.347311867 | 18.35266313 | 0.714755956 | 1.117692138 | 0.16052286 | 0.118433181 | 0.661134935 | 0.179709894 | Unpaired t-test | 0.587246978 | permutation FDR (250 perm) | 1 | 0.691386778 | FDR correction BH |
| Tm1~A0A0B4KG68 | VMAT~Control | VMAT | Control | 17.04462432 | 0.151989625 | 16.88540218 | 0.17616089 | 1.116684894 | 0.159222144 | 0.14421623 | 0.156958556 | 0.804215007 | Unpaired t-test | 0.528384582 | permutation FDR (250 perm) | 1 | 0.182639861 | FDR correction BH |
| Dmel\CG31705~Q9VKG4 | VMAT~Control | VMAT | Control | 13.8108782 | 0.44303732 | 13.65174413 | 0.277176719 | 1.116616729 | 0.159134075 | 0.128987913 | 0.511395699 | 0.291242929 | Unpaired t-test | 0.562701861 | permutation FDR (250 perm) | 1 | 0.547442235 | FDR correction BH |
| Nsun2~Q9W4M9 | VMAT~Control | VMAT | Control | 13.95895922 | 0.672003239 | 13.80071124 | 0.34549818 | 1.115931118 | 0.158247978 | 0.118278901 | 0.649612849 | 0.187345393 | Unpaired t-test | 0.587637306 | permutation FDR (250 perm) | 1 | 0.679941595 | FDR correction BH |
| pain~Q9W0Y6 | VMAT~Control | VMAT | Control | 15.4663492 | 0.216547556 | 15.3113677 | 0.128655246 | 1.113407343 | 0.154981503 | 0.139291002 | 0.198902546 | 0.701359658 | Unpaired t-test | 0.539407536 | permutation FDR (250 perm) | 1 | 0.227731806 | FDR correction BH |
| Cpsf6~M9NF33 | VMAT~Control | VMAT | Control | 19.08615304 | 0.211964947 | 18.85152327 | 0.524420605 | 1.113135921 | 0.154629766 | 0.123411486 | 0.566636886 | 0.246695158 | Unpaired t-test | 0.575512532 | permutation FDR (250 perm) | 1 | 0.591063011 | FDR correction BH |
| sgg~A4V3W2 | VMAT~Control | VMAT | Control | 17.6899214 | 0.061740906 | 17.53561267 | 0.200636062 | 1.112888253 | 0.154308736 | 0.141065575 | 0.131268424 | 0.881839729 | Unpaired t-test | 0.535431789 | permutation FDR (250 perm) | 1 | 0.15489016 | FDR correction BH |
| Dmel\CG8726~A1Z782 | VMAT~Control | VMAT | Control | 16.82674571 | 0.120444499 | 16.67266362 | 0.125537365 | 1.112713437 | 0.154082096 | 0.142959429 | 0.075820403 | 1.120213913 | Unpaired t-test | 0.531201386 | permutation FDR (250 perm) | 1 | 0.092190717 | FDR correction BH |
| Rtnl1~E1JHT6 | VMAT~Control | VMAT | Control | 20.21563153 | 0.069684065 | 20.06255265 | 0.121862025 | 1.111939949 | 0.153078877 | 0.14403637 | 0.034946627 | 1.456594741 | Unpaired t-test | 0.528796882 | permutation FDR (250 perm) | 1 | 0.044587728 | FDR correction BH |
| Pmm2~Q9VTZ6 | VMAT~Control | VMAT | Control | 15.15135399 | 0.224176979 | 14.99885455 | 0.075169224 | 1.111493445 | 0.15249944 | 0.137916066 | 0.179823976 | 0.745152403 | Unpaired t-test | 0.542546557 | permutation FDR (250 perm) | 1 | 0.207096693 | FDR correction BH |
| Lis-1~A0A0B4K812 | VMAT~Control | VMAT | Control | 15.69594342 | 0.1054339 | 15.54400988 | 0.071620049 | 1.111057539 | 0.151933533 | 0.143740158 | 0.023677157 | 1.62567044 | Unpaired t-test | 0.529472499 | permutation FDR (250 perm) | 1 | 0.030844509 | FDR correction BH |
| Got1~Q7K221 | VMAT~Control | VMAT | Control | 16.45760019 | 0.811955576 | 16.30669736 | 0.460617193 | 1.110264055 | 0.150902835 | 0.106458667 | 0.725273846 | 0.139497983 | Unpaired t-test | 0.618356188 | permutation FDR (250 perm) | 1 | 0.752443647 | FDR correction BH |
| CG14632~X2JDC1 | VMAT~Control | VMAT | Control | 13.5158002 | 0.507646821 | 13.3687234 | 0.115387224 | 1.107323526 | 0.147076795 | 0.119301371 | 0.541737081 | 0.266211436 | Unpaired t-test | 0.58516753 | permutation FDR (250 perm) | 1 | 0.57808209 | FDR correction BH |
| l(3)80Fj~Q7PLL6 | VMAT~Control | VMAT | Control | 14.24852019 | 0.466649274 | 14.10257439 | 0.478527569 | 1.106455782 | 0.145945797 | 0.112359796 | 0.635899953 | 0.196611207 | Unpaired t-test | 0.602836928 | permutation FDR (250 perm) | 1 | 0.668262718 | FDR correction BH |
| CNBP~Q8T8R1 | VMAT~Control | VMAT | Control | 18.92681563 | 0.122220987 | 18.78097816 | 0.266905987 | 1.106372701 | 0.145837465 | 0.128913279 | 0.292624374 | 0.533689503 | Unpaired t-test | 0.562608997 | permutation FDR (250 perm) | 1 | 0.32755006 | FDR correction BH |
| Syx4~Q7KVY7 | VMAT~Control | VMAT | Control | 17.0351176 | 0.26478097 | 16.88951026 | 0.473555382 | 1.106196236 | 0.145607338 | 0.117176081 | 0.561788871 | 0.250426868 | Unpaired t-test | 0.590410013 | permutation FDR (250 perm) | 1 | 0.597853075 | FDR correction BH |
| spen~M9PBZ3 | VMAT~Control | VMAT | Control | 20.50132528 | 0.042537433 | 20.35933733 | 0.151545023 | 1.103424527 | 0.141987954 | 0.132650397 | 0.071325075 | 1.14675776 | Unpaired t-test | 0.554429437 | permutation FDR (250 perm) | 1 | 0.087039517 | FDR correction BH |
| f-cup~Q7KSK8 | VMAT~Control | VMAT | Control | 17.41552315 | 0.063347578 | 17.27390753 | 0.215947752 | 1.103139788 | 0.141615619 | 0.128666112 | 0.189712845 | 0.721903262 | Unpaired t-test | 0.563269896 | permutation FDR (250 perm) | 1 | 0.217633547 | FDR correction BH |
| Khc~P17210 | VMAT~Control | VMAT | Control | 16.46334701 | 0.15741072 | 16.32200751 | 0.264773281 | 1.102928674 | 0.141339495 | 0.124226587 | 0.329062441 | 0.482721686 | Unpaired t-test | 0.573746649 | permutation FDR (250 perm) | 1 | 0.365211214 | FDR correction BH |
| blp~A0A0B4LH72 | VMAT~Control | VMAT | Control | 13.73431553 | 0.441766725 | 13.59299993 | 0.410839274 | 1.102910406 | 0.141315599 | 0.111290088 | 0.611835997 | 0.213364975 | Unpaired t-test | 0.605362656 | permutation FDR (250 perm) | 1 | 0.646437135 | FDR correction BH |
| eEF2~P13060 | VMAT~Control | VMAT | Control | 19.13403811 | 0.084779506 | 18.9935131 | 0.119954941 | 1.102306189 | 0.140525018 | 0.131862778 | 0.058116076 | 1.235703716 | Unpaired t-test | 0.556046733 | permutation FDR (250 perm) | 1 | 0.072078632 | FDR correction BH |
| jdp~Q9TVP3 | VMAT~Control | VMAT | Control | 20.94095305 | 0.162926998 | 20.80322955 | 0.153623884 | 1.100167739 | 0.137723504 | 0.125186638 | 0.199082203 | 0.700967563 | Unpaired t-test | 0.571412019 | permutation FDR (250 perm) | 1 | 0.227826692 | FDR correction BH |
| GstE9~Q7K8X7 | VMAT~Control | VMAT | Control | 14.83064545 | 0.235451437 | 14.69321577 | 0.356246048 | 1.099943698 | 0.13742968 | 0.115393019 | 0.48822519 | 0.311379817 | Unpaired t-test | 0.595169616 | permutation FDR (250 perm) | 1 | 0.525266041 | FDR correction BH |
| Alk~Q7KJ08 | VMAT~Control | VMAT | Control | 15.51555937 | 0.222021068 | 15.37846699 | 0.224555121 | 1.099686565 | 0.137092382 | 0.120094574 | 0.355588095 | 0.449052787 | Unpaired t-test | 0.583165803 | permutation FDR (250 perm) | 1 | 0.392614623 | FDR correction BH |
| Ns1~Q8MT06 | VMAT~Control | VMAT | Control | 17.50395087 | 0.190781342 | 17.3679074 | 0.476609487 | 1.098887327 | 0.136043469 | 0.110641473 | 0.566636886 | 0.246695158 | Unpaired t-test | 0.607108236 | permutation FDR (250 perm) | 1 | 0.602195589 | FDR correction BH |
| Syp~Q95TW4 | VMAT~Control | VMAT | Control | 17.26069804 | 0.102419986 | 17.12466255 | 0.148875609 | 1.098881253 | 0.136035495 | 0.12586403 | 0.123218601 | 0.909323725 | Unpaired t-test | 0.569843493 | permutation FDR (250 perm) | 1 | 0.145830361 | FDR correction BH |
| Fmo-2~Q7K3U4 | VMAT~Control | VMAT | Control | 16.59549844 | 0.184745579 | 16.46088724 | 0.145104487 | 1.097796921 | 0.134611199 | 0.121813661 | 0.229002086 | 0.640160562 | Unpaired t-test | 0.579585134 | permutation FDR (250 perm) | 1 | 0.260546597 | FDR correction BH |
| Ist1~M9PES8 | VMAT~Control | VMAT | Control | 18.33505125 | 0.076916881 | 18.2054531 | 0.148358644 | 1.09398894 | 0.129598153 | 0.120586164 | 0.113560202 | 0.944773842 | Unpaired t-test | 0.581994819 | permutation FDR (250 perm) | 1 | 0.135146975 | FDR correction BH |
| B4~Q9VJX3 | VMAT~Control | VMAT | Control | 13.96385549 | 0.675996507 | 13.83528552 | 0.774297102 | 1.093209553 | 0.128569972 | 0.088081209 | 0.785409703 | 0.104903738 | Unpaired t-test | 0.669494411 | permutation FDR (250 perm) | 1 | 0.808065752 | FDR correction BH |
| Dmel\CG11307~B5RJJ4 | VMAT~Control | VMAT | Control | 17.09102287 | 0.077948698 | 16.963111 | 0.24155523 | 1.092710984 | 0.127911867 | 0.114872458 | 0.286116506 | 0.543457087 | Unpaired t-test | 0.596509279 | permutation FDR (250 perm) | 1 | 0.320722979 | FDR correction BH |
| Stim~P83094 | VMAT~Control | VMAT | Control | 17.93942513 | 0.183083401 | 17.81321386 | 0.219751188 | 1.091423696 | 0.126211271 | 0.111897944 | 0.347067364 | 0.459586222 | Unpaired t-test | 0.604041415 | permutation FDR (250 perm) | 1 | 0.38407379 | FDR correction BH |
| Fkbp39~P54397 | VMAT~Control | VMAT | Control | 15.92609661 | 0.263670221 | 15.8014946 | 0.151798298 | 1.090206937 | 0.124602005 | 0.109678852 | 0.381339726 | 0.418687949 | Unpaired t-test | 0.609631738 | permutation FDR (250 perm) | 1 | 0.418886475 | FDR correction BH |
| uif~E1JHA6 | VMAT~Control | VMAT | Control | 16.00508337 | 0.446153441 | 15.88146082 | 1.073647769 | 1.08946704 | 0.123622551 | 0.081332967 | 0.816871111 | 0.087846463 | Unpaired t-test | 0.68964433 | permutation FDR (250 perm) | 1 | 0.838236528 | FDR correction BH |
| tweek~M9MSI8 | VMAT~Control | VMAT | Control | 17.94532897 | 0.179136753 | 17.82206031 | 0.201913372 | 1.089199824 | 0.123268654 | 0.109991209 | 0.331244777 | 0.479850961 | Unpaired t-test | 0.608810694 | permutation FDR (250 perm) | 1 | 0.36746004 | FDR correction BH |
| ytr~Q9W1I6 | VMAT~Control | VMAT | Control | 18.17872651 | 0.196701498 | 18.058602 | 0.596288483 | 1.086828657 | 0.120124512 | 0.093788442 | 0.67787064 | 0.168853176 | Unpaired t-test | 0.652869191 | permutation FDR (250 perm) | 1 | 0.70669065 | FDR correction BH |
| Hsp23~M9NE68 | VMAT~Control | VMAT | Control | 16.94965621 | 0.130965361 | 16.83096115 | 0.334793447 | 1.085752336 | 0.118695057 | 0.102255251 | 0.477312515 | 0.321197179 | Unpaired t-test | 0.629489013 | permutation FDR (250 perm) | 1 | 0.514937516 | FDR correction BH |
| Ddc~P05031 | VMAT~Control | VMAT | Control | 16.10187237 | 0.298804311 | 15.98360998 | 0.426117038 | 1.085426761 | 0.118262384 | 0.095933896 | 0.622394474 | 0.205934272 | Unpaired t-test | 0.647037452 | permutation FDR (250 perm) | 1 | 0.655533151 | FDR correction BH |
| Pde6~A0A0B4KFZ3 | VMAT~Control | VMAT | Control | 14.77551177 | 0.156360087 | 14.65734571 | 0.338018697 | 1.085354288 | 0.118166053 | 0.101294765 | 0.494244953 | 0.306057757 | Unpaired t-test | 0.632046512 | permutation FDR (250 perm) | 1 | 0.53077218 | FDR correction BH |
| eIF1A~Q9VEA1 | VMAT~Control | VMAT | Control | 16.28311306 | 0.1780636 | 16.16592694 | 0.474350686 | 1.08461733 | 0.117186128 | 0.095538138 | 0.616280965 | 0.210221246 | Unpaired t-test | 0.648167025 | permutation FDR (250 perm) | 1 | 0.650257908 | FDR correction BH |
| Prp4k~X2JG22 | VMAT~Control | VMAT | Control | 19.30175298 | 0.396726995 | 19.18609204 | 0.593739755 | 1.0834713 | 0.115660939 | 0.087665152 | 0.724756337 | 0.139807979 | Unpaired t-test | 0.670837489 | permutation FDR (250 perm) | 1 | 0.752238279 | FDR correction BH |
| Tm1~A0A0B4KG06 | VMAT~Control | VMAT | Control | 18.62671491 | 0.074591217 | 18.51193494 | 0.152053017 | 1.082809886 | 0.114779965 | 0.10669845 | 0.160618904 | 0.794203342 | Unpaired t-test | 0.617640362 | permutation FDR (250 perm) | 1 | 0.186346427 | FDR correction BH |
| Dmel\CG5068~Q95R98 | VMAT~Control | VMAT | Control | 15.72673896 | 0.120560094 | 15.61389489 | 0.21388431 | 1.081357883 | 0.112844072 | 0.10167956 | 0.328300809 | 0.483728047 | Unpaired t-test | 0.630832041 | permutation FDR (250 perm) | 1 | 0.364537785 | FDR correction BH |
| Vha26~A0A0B4LGS4 | VMAT~Control | VMAT | Control | 16.66281612 | 0.122661608 | 16.5506314 | 0.259045554 | 1.080953787 | 0.112184723 | 0.099438681 | 0.401990405 | 0.395784313 | Unpaired t-test | 0.637279931 | permutation FDR (250 perm) | 1 | 0.439088504 | FDR correction BH |
| Ulp1~Q9VWK5 | VMAT~Control | VMAT | Control | 14.97441208 | 2.197302553 | 14.86266024 | 1.6182412 | 1.080539519 | 0.111751838 | 0.050329667 | 0.928848 | 0.03205535 | Unpaired t-test | 0.793516888 | permutation FDR (250 perm) | 1 | 0.940185446 | FDR correction BH |
| mipIe1~Q9W0S5 | VMAT~Control | VMAT | Control | 16.40908448 | 0.132715942 | 16.29823448 | 0.511905239 | 1.079864276 | 0.110849997 | 0.089648221 | 0.649326919 | 0.187536592 | Unpaired t-test | 0.664797936 | permutation FDR (250 perm) | 1 | 0.679941595 | FDR correction BH |
| RpS25~P48588 | VMAT~Control | VMAT | Control | 16.67062964 | 0.214214729 | 16.56170142 | 0.328794236 | 1.078426773 | 0.108928218 | 0.092665778 | 0.548688616 | 0.260674051 | Unpaired t-test | 0.656313253 | permutation FDR (250 perm) | 1 | 0.584969656 | FDR correction BH |
| Argl~Q9VLG9 | VMAT~Control | VMAT | Control | 13.89727825 | 0.417941186 | 13.79013073 | 0.656747019 | 1.077096504 | 0.107147516 | 0.079478302 | 0.764568023 | 0.11658387 | Unpaired t-test | 0.695328179 | permutation FDR (250 perm) | 1 | 0.78938295 | FDR correction BH |
| Dmel\CG6195~Q9VDV2 | VMAT~Control | VMAT | Control | 16.64205089 | 0.195261938 | 16.53584221 | 0.559018951 | 1.076395808 | 0.106208678 | 0.083971848 | 0.696804078 | 0.156889316 | Unpaired t-test | 0.681577997 | permutation FDR (250 perm) | 1 | 0.724503887 | FDR correction BH |
| Lamtor3~M9NEQ5 | VMAT~Control | VMAT | Control | 20.31003359 | 0.229099047 | 20.20415044 | 0.435063749 | 1.076152955 | 0.105883144 | 0.086797004 | 0.640511115 | 0.193473329 | Unpaired t-test | 0.673017619 | permutation FDR (250 perm) | 1 | 0.671908719 | FDR correction BH |
| Dmel\CG12896~A1Z893 | VMAT~Control | VMAT | Control | 16.0333491 | 0.22808457 | 15.92854453 | 0.239715452 | 1.075348714 | 0.104804573 | 0.091295005 | 0.494968268 | 0.305422643 | Unpaired t-test | 0.65989166 | permutation FDR (250 perm) | 1 | 0.531306567 | FDR correction BH |
| Sbf~Q7KSP6 | VMAT~Control | VMAT | Control | 15.75452186 | 0.180040296 | 15.65016207 | 0.273256737 | 1.07501724 | 0.104359797 | 0.091037018 | 0.49208328 | 0.307961391 | Unpaired t-test | 0.660646604 | permutation FDR (250 perm) | 1 | 0.528691941 | FDR correction BH |
| Ehbp1~A1ZAN2 | VMAT~Control | VMAT | Control | 16.35029141 | 0.158037479 | 16.24745488 | 0.180547911 | 1.073882782 | 0.102836527 | 0.092870881 | 0.360485608 | 0.443112069 | Unpaired t-test | 0.655672978 | permutation FDR (250 perm) | 1 | 0.397307235 | FDR correction BH |
| dod~P54353 | VMAT~Control | VMAT | Control | 16.30037449 | 0.048314736 | 16.19753983 | 0.094146711 | 1.07388139 | 0.102834656 | 0.09818785 | 0.054892818 | 1.260484475 | Unpaired t-test | 0.640768303 | permutation FDR (250 perm) | 1 | 0.068260799 | FDR correction BH |
| Srpk79D~Q8T3S1 | VMAT~Control | VMAT | Control | 15.65494676 | 0.302071764 | 15.55245001 | 0.320116313 | 1.073629895 | 0.102496749 | 0.08563977 | 0.61389435 | 0.211906364 | Unpaired t-test | 0.676464117 | permutation FDR (250 perm) | 1 | 0.648320906 | FDR correction BH |
| eIF3m~A0A0B4LFB3 | VMAT~Control | VMAT | Control | 15.8986634 | 0.144027802 | 15.79705986 | 0.207585324 | 1.072965387 | 0.101603537 | 0.09128867 | 0.389678961 | 0.409293041 | Unpaired t-test | 0.659910576 | permutation FDR (250 perm) | 1 | 0.426653151 | FDR correction BH |
| Dmel\CG31522~Q9VN29 | VMAT~Control | VMAT | Control | 17.15756257 | 0.262806874 | 17.05688206 | 0.09039236 | 1.07227913 | 0.100680509 | 0.089550417 | 0.436760909 | 0.359756239 | Unpaired t-test | 0.665122958 | permutation FDR (250 perm) | 1 | 0.475328331 | FDR correction BH |
| Dmel\CG14894~Q9VEV3 | VMAT~Control | VMAT | Control | 19.37429664 | 0.259946804 | 19.2739133 | 0.247768758 | 1.072058286 | 0.100383345 | 0.086492634 | 0.545934476 | 0.262859479 | Unpaired t-test | 0.673882252 | permutation FDR (250 perm) | 1 | 0.582297126 | FDR correction BH |
| p115~Q9W3N6 | VMAT~Control | VMAT | Control | 16.71942176 | 0.222521119 | 16.62014197 | 0.243542807 | 1.071238561 | 0.099279799 | 0.086515904 | 0.516236511 | 0.287151283 | Unpaired t-test | 0.673823807 | permutation FDR (250 perm) | 1 | 0.552373067 | FDR correction BH |
| Dmel\CG12237~Q9VWF0 | VMAT~Control | VMAT | Control | 15.71852493 | 0.084549577 | 15.6203502 | 0.1550422 | 1.070418334 | 0.098174732 | 0.090988729 | 0.242193136 | 0.615838169 | Unpaired t-test | 0.660811694 | permutation FDR (250 perm) | 1 | 0.275022983 | FDR correction BH |
| Dmel\CG11808~Q7K1H5 | VMAT~Control | VMAT | Control | 18.15065519 | 0.430959014 | 18.05292883 | 0.680580733 | 1.070085712 | 0.097726358 | 0.071844183 | 0.791700486 | 0.101439088 | Unpaired t-test | 0.7196103 | permutation FDR (250 perm) | 1 | 0.814182152 | FDR correction BH |
| CHMP2B~Q9VRJ5 | VMAT~Control | VMAT | Control | 17.25368675 | 0.11491862 | 17.15652519 | 0.154945808 | 1.069666871 | 0.097161564 | 0.089444948 | 0.286373296 | 0.543067482 | Unpaired t-test | 0.66550129 | permutation FDR (250 perm) | 1 | 0.320858038 | FDR correction BH |
| Fak~B7YZL9 | VMAT~Control | VMAT | Control | 15.52256522 | 0.199972279 | 15.42786755 | 0.168607302 | 1.067841605 | 0.094697665 | 0.084780367 | 0.437040299 | 0.359478515 | Unpaired t-test | 0.679033949 | permutation FDR (250 perm) | 1 | 0.475412599 | FDR correction BH |
| CG9216~Q8IR21 | VMAT~Control | VMAT | Control | 19.57780381 | 0.174362194 | 19.48444838 | 0.576075896 | 1.066848582 | 0.093355428 | 0.073556215 | 0.735910071 | 0.133175253 | Unpaired t-test | 0.713764807 | permutation FDR (250 perm) | 1 | 0.76194845 | FDR correction BH |
| dj-1beta~Q9VA37 | VMAT~Control | VMAT | Control | 16.54130983 | 0.15569713 | 16.45581091 | 0.149769637 | 1.061054614 | 0.085498916 | 0.077966192 | 0.396967123 | 0.40124546 | Unpaired t-test | 0.699759553 | permutation FDR (250 perm) | 1 | 0.434027222 | FDR correction BH |
| Srp19~P49963 | VMAT~Control | VMAT | Control | 15.66429073 | 0.110058858 | 15.57907774 | 0.192442084 | 1.060844352 | 0.085212998 | 0.077526746 | 0.410199026 | 0.387005375 | Unpaired t-test | 0.701082009 | permutation FDR (250 perm) | 1 | 0.447455286 | FDR correction BH |
| anon-WO0118547.344~Q9VHN7 | VMAT~Control | VMAT | Control | 19.29269987 | 0.078997209 | 19.20913016 | 0.219397451 | 1.059636693 | 0.083569708 | 0.075677729 | 0.441544964 | 0.355025064 | Unpaired t-test | 0.707077716 | permutation FDR (250 perm) | 1 | 0.479647829 | FDR correction BH |
| vimar~A1Z6S7 | VMAT~Control | VMAT | Control | 15.51679017 | 0.126287197 | 15.43553434 | 0.395836307 | 1.057938549 | 0.081255831 | 0.068523232 | 0.67118966 | 0.173154742 | Unpaired t-test | 0.730395538 | permutation FDR (250 perm) | 1 | 0.700656523 | FDR correction BH |
| Spn77Bb~Q9VWB4 | VMAT~Control | VMAT | Control | 16.92468816 | 0.292504222 | 16.8440679 | 0.337987484 | 1.057472584 | 0.080620261 | 0.067189312 | 0.695209411 | 0.157884358 | Unpaired t-test | 0.735121407 | permutation FDR (250 perm) | 1 | 0.723165246 | FDR correction BH |
| CaMKI~Q7JMV3 | VMAT~Control | VMAT | Control | 17.43352547 | 0.148298574 | 17.35416857 | 0.254475686 | 1.05654697 | 0.079356905 | 0.070120639 | 0.56027517 | 0.251598624 | Unpaired t-test | 0.725327039 | permutation FDR (250 perm) | 1 | 0.596781787 | FDR correction BH |
| Pgi~P52029 | VMAT~Control | VMAT | Control | 18.57877675 | 0.544510566 | 18.49987418 | 0.577392785 | 1.056214288 | 0.078902563 | 0.058233711 | 0.828550992 | 0.081680758 | Unpaired t-test | 0.765236301 | permutation FDR (250 perm) | 1 | 0.849481287 | FDR correction BH |
| CCT7~Q9VHL2 | VMAT~Control | VMAT | Control | 17.83943307 | 0.090484318 | 17.7610578 | 0.11312722 | 1.055828318 | 0.078375265 | 0.07360669 | 0.254183862 | 0.594852026 | Unpaired t-test | 0.713608584 | permutation FDR (250 perm) | 1 | 0.287806066 | FDR correction BH |
| Rpn11~Q9V3H2 | VMAT~Control | VMAT | Control | 17.53859485 | 0.114103393 | 17.46144854 | 0.281358046 | 1.0549293 | 0.077146315 | 0.067923604 | 0.582452686 | 0.234739348 | Unpaired t-test | 0.732528529 | permutation FDR (250 perm) | 1 | 0.617331662 | FDR correction BH |
| pkm~Q7KV34 | VMAT~Control | VMAT | Control | 17.65743485 | 0.13705681 | 17.58158324 | 0.171317002 | 1.054012228 | 0.075891604 | 0.069110714 | 0.4571298 | 0.339960466 | Unpaired t-test | 0.728441012 | permutation FDR (250 perm) | 1 | 0.494978633 | FDR correction BH |
| rut~M9PEL4 | VMAT~Control | VMAT | Control | 13.60048067 | 0.276940734 | 13.52886993 | 0.354563085 | 1.050889324 | 0.071610738 | 0.059615901 | 0.729298335 | 0.137094778 | Unpaired t-test | 0.760303082 | permutation FDR (250 perm) | 1 | 0.755619842 | FDR correction BH |
| Paics~Q9I7S8 | VMAT~Control | VMAT | Control | 17.3229197 | 0.125240631 | 17.25170484 | 0.153525625 | 1.050600996 | 0.071214857 | 0.06541838 | 0.440243961 | 0.356306593 | Unpaired t-test | 0.741148005 | permutation FDR (250 perm) | 1 | 0.478455348 | FDR correction BH |
| beta-PheRS~Q9VCA5 | VMAT~Control | VMAT | Control | 16.29030599 | 0.12958332 | 16.22018049 | 0.119200674 | 1.049808007 | 0.070125507 | 0.065006813 | 0.394073468 | 0.404422804 | Unpaired t-test | 0.742625483 | permutation FDR (250 perm) | 1 | 0.431063635 | FDR correction BH |
| GluClalpha~A0A0B4LHB7 | VMAT~Control | VMAT | Control | 16.52982765 | 0.248529922 | 16.46365756 | 0.214058522 | 1.046933708 | 0.066170093 | 0.057705357 | 0.661547696 | 0.179438839 | Unpaired t-test | 0.766449957 | permutation FDR (250 perm) | 1 | 0.691511224 | FDR correction BH |
| Rpt4~Q9W414 | VMAT~Control | VMAT | Control | 16.00519188 | 0.096140951 | 15.93915386 | 0.075727006 | 1.046837871 | 0.066038022 | 0.062611218 | 0.255361284 | 0.592844947 | Unpaired t-test | 0.749844968 | permutation FDR (250 perm) | 1 | 0.288583995 | FDR correction BH |
| Dmel\CG6621~A0A0B4JDC3 | VMAT~Control | VMAT | Control | 20.693243 | 0.071572077 | 20.6274282 | 0.457020149 | 1.04667591 | 0.065814799 | 0.054533156 | 0.75692659 | 0.120946238 | Unpaired t-test | 0.77767593 | permutation FDR (250 perm) | 1 | 0.782179628 | FDR correction BH |
| Fer1HCH~H1UUD2 | VMAT~Control | VMAT | Control | 15.50321172 | 0.328595811 | 15.44394641 | 0.126085361 | 1.041935026 | 0.059265315 | 0.05120559 | 0.714390075 | 0.146064587 | Unpaired t-test | 0.790206071 | permutation FDR (250 perm) | 1 | 0.741806016 | FDR correction BH |
| DAAM~Q7YZA4 | VMAT~Control | VMAT | Control | 15.15835301 | 0.196153876 | 15.09953591 | 0.324786353 | 1.041611367 | 0.058817098 | 0.050284625 | 0.736053855 | 0.133090408 | Unpaired t-test | 0.79367251 | permutation FDR (250 perm) | 1 | 0.76194845 | FDR correction BH |
| RpL7A~X2JCS6 | VMAT~Control | VMAT | Control | 18.36531024 | 0.098218006 | 18.306759 | 0.108079572 | 1.041419442 | 0.058551245 | 0.054961618 | 0.39106094 | 0.40775556 | Unpaired t-test | 0.776509837 | permutation FDR (250 perm) | 1 | 0.427967202 | FDR correction BH |
| FOHSDR~Q9VDC0 | VMAT~Control | VMAT | Control | 15.45079473 | 0.290383595 | 15.39665154 | 0.424475578 | 1.038242319 | 0.054143199 | 0.044018827 | 0.818645712 | 0.086904009 | Unpaired t-test | 0.816766866 | permutation FDR (250 perm) | 1 | 0.839691506 | FDR correction BH |
| RpL13~M9PFF0 | VMAT~Control | VMAT | Control | 18.66790569 | 0.066359434 | 18.61605972 | 0.282018413 | 1.036590422 | 0.051845969 | 0.045898979 | 0.697461771 | 0.156479592 | Unpaired t-test | 0.809426132 | permutation FDR (250 perm) | 1 | 0.724867554 | FDR correction BH |
| Prp40~Q9VQK5 | VMAT~Control | VMAT | Control | 17.05866738 | 0.498050346 | 17.00791489 | 0.571589145 | 1.035805039 | 0.050752481 | 0.037901909 | 0.883984373 | 0.053555412 | Unpaired t-test | 0.840867264 | permutation FDR (250 perm) | 1 | 0.901602779 | FDR correction BH |
| CG1354-RA~E6PBV6 | VMAT~Control | VMAT | Control | 16.47434842 | 0.106183902 | 16.4252989 | 0.327966492 | 1.034583809 | 0.049049515 | 0.042497769 | 0.756909352 | 0.120956129 | Unpaired t-test | 0.822809564 | permutation FDR (250 perm) | 1 | 0.782179628 | FDR correction BH |
| anon-WO0172774.24~Q9Y114 | VMAT~Control | VMAT | Control | 17.21719629 | 0.264941254 | 17.17115191 | 0.335671449 | 1.032430289 | 0.046044372 | 0.03865238 | 0.814602459 | 0.089054284 | Unpaired t-test | 0.838006832 | permutation FDR (250 perm) | 1 | 0.836273087 | FDR correction BH |
| Gmppb~E2QCZ9 | VMAT~Control | VMAT | Control | 15.72685856 | 0.135542931 | 15.68092215 | 0.141275751 | 1.032353029 | 0.045936407 | 0.042238179 | 0.611259557 | 0.213774338 | Unpaired t-test | 0.823861657 | permutation FDR (250 perm) | 1 | 0.646118094 | FDR correction BH |
| Dmel\CG1307~E1JJ73 | VMAT~Control | VMAT | Control | 15.77691156 | 0.293712288 | 15.7330467 | 0.535779649 | 1.030871753 | 0.043864864 | 0.034451111 | 0.875659832 | 0.057664572 | Unpaired t-test | 0.854036674 | permutation FDR (250 perm) | 1 | 0.894662866 | FDR correction BH |
| Dmel\CG9281~Q9VXR5 | VMAT~Control | VMAT | Control | 16.37717746 | 0.176095602 | 16.3347952 | 0.057789946 | 1.029812911 | 0.042382263 | 0.039138298 | 0.62021273 | 0.207459324 | Unpaired t-test | 0.836175918 | permutation FDR (250 perm) | 1 | 0.653527648 | FDR correction BH |
| Ptp61F~Q9W0G1-2 | VMAT~Control | VMAT | Control | 17.36170606 | 0.309318346 | 17.31974564 | 0.291355023 | 1.029511839 | 0.041960421 | 0.035259835 | 0.829687892 | 0.081085248 | Unpaired t-test | 0.850768443 | permutation FDR (250 perm) | 1 | 0.850276577 | FDR correction BH |
| Dmel\CG7332~X2JL49 | VMAT~Control | VMAT | Control | 17.43353957 | 0.108010767 | 17.39260941 | 0.158310386 | 1.028776899 | 0.040930152 | 0.037699075 | 0.643230866 | 0.191633124 | Unpaired t-test | 0.841686726 | permutation FDR (250 perm) | 1 | 0.674461228 | FDR correction BH |
| CG3641~Q9VFF2 | VMAT~Control | VMAT | Control | 15.68888535 | 0.833846881 | 15.65326703 | 0.952110633 | 1.024996024 | 0.035618313 | 0.022744682 | 0.951062977 | 0.021790724 | Unpaired t-test | 0.90193356 | permutation FDR (250 perm) | 1 | 0.958716852 | FDR correction BH |
| eff~A0A0B4KHJ1 | VMAT~Control | VMAT | Control | 16.00952289 | 0.264757185 | 15.97559856 | 0.165318833 | 1.023793203 | 0.033924333 | 0.029768896 | 0.812895662 | 0.089965194 | Unpaired t-test | 0.872963342 | permutation FDR (250 perm) | 1 | 0.834884987 | FDR correction BH |
| CG8833~Q9VUA0 | VMAT~Control | VMAT | Control | 17.16887758 | 0.35804824 | 17.13553066 | 0.503454193 | 1.023338532 | 0.033346924 | 0.026128134 | 0.906321007 | 0.042717954 | Unpaired t-test | 0.88828474 | permutation FDR (250 perm) | 1 | 0.921191559 | FDR correction BH |
| fray~A0A126GUV7 | VMAT~Control | VMAT | Control | 17.28830406 | 0.192655426 | 17.2584495 | 0.244732475 | 1.020909199 | 0.029854556 | 0.026204502 | 0.834595474 | 0.078523975 | Unpaired t-test | 0.887936914 | permutation FDR (250 perm) | 1 | 0.854468209 | FDR correction BH |
| Klc~M9PFG7 | VMAT~Control | VMAT | Control | 18.06035782 | 0.179934085 | 18.03213753 | 0.131204013 | 1.019753381 | 0.028220291 | 0.025664377 | 0.782676803 | 0.106417537 | Unpaired t-test | 0.890219949 | permutation FDR (250 perm) | 1 | 0.805958528 | FDR correction BH |
| S-Lap2~Q9VSM7 | VMAT~Control | VMAT | Control | 14.44471299 | 0.372596642 | 14.41974442 | 0.333094169 | 1.017457524 | 0.024968567 | 0.020407352 | 0.913262274 | 0.039404482 | Unpaired t-test | 0.91116766 | permutation FDR (250 perm) | 1 | 0.927045879 | FDR correction BH |
| CanA-14F~M9NEL1 | VMAT~Control | VMAT | Control | 18.70612248 | 0.084537794 | 18.68576792 | 0.136552337 | 1.014208707 | 0.020354566 | 0.018990592 | 0.782653318 | 0.106430569 | Unpaired t-test | 0.916859209 | permutation FDR (250 perm) | 1 | 0.805958528 | FDR correction BH |
| DSCR3~Q9VPC3 | VMAT~Control | VMAT | Control | 15.62759479 | 0.509402822 | 15.60771506 | 0.78819395 | 1.013874956 | 0.019879732 | 0.014002768 | 0.963153739 | 0.016304385 | Unpaired t-test | 0.938545918 | permutation FDR (250 perm) | 1 | 0.968493184 | FDR correction BH |
| l(2)gl~M9NCX1 | VMAT~Control | VMAT | Control | 17.78554036 | 0.032173003 | 17.76712676 | 0.096587355 | 1.012845138 | 0.018413605 | 0.017611768 | 0.694403092 | 0.158388354 | Unpaired t-test | 0.922442177 | permutation FDR (250 perm) | 1 | 0.722645835 | FDR correction BH |
| dre4~Q8IRG6 | VMAT~Control | VMAT | Control | 17.10479855 | 0.235322273 | 17.08733372 | 0.574040592 | 1.012179264 | 0.017464824 | 0.013671604 | 0.951049069 | 0.021797075 | Unpaired t-test | 0.939916667 | permutation FDR (250 perm) | 1 | 0.958716852 | FDR correction BH |
| RpL23A~Q9W0A8 | VMAT~Control | VMAT | Control | 17.6522302 | 0.120179627 | 17.6348287 | 0.354472672 | 1.012134836 | 0.017401498 | 0.014906351 | 0.919257371 | 0.036562879 | Unpaired t-test | 0.93447619 | permutation FDR (250 perm) | 1 | 0.932729246 | FDR correction BH |
| cpa~Q9W2N0 | VMAT~Control | VMAT | Control | 16.00766158 | 0.184678586 | 15.99124973 | 0.224494925 | 1.011440775 | 0.016411846 | 0.014523716 | 0.90204262 | 0.044772942 | Unpaired t-test | 0.936183673 | permutation FDR (250 perm) | 1 | 0.918032134 | FDR correction BH |
| Snx1~Q9VQQ6 | VMAT~Control | VMAT | Control | 13.97150349 | 0.658222251 | 13.95625934 | 0.660444553 | 1.01062246 | 0.015244147 | 0.010758046 | 0.971557901 | 0.012531312 | Unpaired t-test | 0.952606885 | permutation FDR (250 perm) | 1 | 0.975335591 | FDR correction BH |
| Tspo~Q9VPR1 | VMAT~Control | VMAT | Control | 17.00855998 | 0.287527319 | 16.99474691 | 0.17760164 | 1.009620475 | 0.013813074 | 0.0119991 | 0.929002272 | 0.031983224 | Unpaired t-test | 0.947477891 | permutation FDR (250 perm) | 1 | 0.940185446 | FDR correction BH |
| CK02656~Q9VPJ9 | VMAT~Control | VMAT | Control | 17.66608043 | 0.081281792 | 17.65302543 | 0.179572566 | 1.009090153 | 0.013055001 | 0.011997416 | 0.885209055 | 0.052954152 | Unpaired t-test | 0.947482993 | permutation FDR (250 perm) | 1 | 0.902460856 | FDR correction BH |
| Dmel\CG9213~X2JKF2 | VMAT~Control | VMAT | Control | 16.98934535 | 0.409573798 | 16.97650293 | 0.671886635 | 1.008941427 | 0.012842423 | 0.009499508 | 0.97160661 | 0.012509539 | Unpaired t-test | 0.958191245 | permutation FDR (250 perm) | 1 | 0.975335591 | FDR correction BH |
| Fas3~A8DZ14 | VMAT~Control | VMAT | Control | 15.48559425 | 0.209186049 | 15.47293265 | 0.313921039 | 1.008814974 | 0.012661595 | 0.010833877 | 0.941653426 | 0.026108909 | Unpaired t-test | 0.952251594 | permutation FDR (250 perm) | 1 | 0.95135286 | FDR correction BH |
| Nak~Q9VJ30 | VMAT~Control | VMAT | Control | 16.25603485 | 0.252167024 | 16.24543204 | 0.225948181 | 1.007376382 | 0.010602812 | 0.009208462 | 0.945556395 | 0.024312564 | Unpaired t-test | 0.959415215 | permutation FDR (250 perm) | 1 | 0.954886209 | FDR correction BH |
| CG34039-RA~A8E6W0 | VMAT~Control | VMAT | Control | 13.93619902 | 0.74013627 | 13.9265342 | 0.928554276 | 1.006721635 | 0.009664824 | 0.00631259 | 0.985837454 | 0.006194686 | Unpaired t-test | 0.972452189 | permutation FDR (250 perm) | 1 | 0.986675751 | FDR correction BH |
| CG1809~Q9I7R7 | VMAT~Control | VMAT | Control | 16.84549746 | 0.181227336 | 16.83699311 | 0.533145383 | 1.00591217 | 0.008504343 | 0.006793538 | 0.973724795 | 0.011563771 | Unpaired t-test | 0.970347641 | permutation FDR (250 perm) | 1 | 0.977045255 | FDR correction BH |
| Ncc69~M9PCA7 | VMAT~Control | VMAT | Control | 17.51476763 | 0.106464785 | 17.50662615 | 0.264831073 | 1.005659198 | 0.008141481 | 0.007219877 | 0.95040182 | 0.022092741 | Unpaired t-test | 0.96839949 | permutation FDR (250 perm) | 1 | 0.958716852 | FDR correction BH |
| swm~Q9VIV2 | VMAT~Control | VMAT | Control | 16.20418438 | 0.371418775 | 16.19616441 | 0.736438407 | 1.005574497 | 0.008019967 | 0.00585886 | 0.98308109 | 0.007410658 | Unpaired t-test | 0.97438674 | permutation FDR (250 perm) | 1 | 0.984932855 | FDR correction BH |
| CG4968~Q9VL00 | VMAT~Control | VMAT | Control | 16.92351383 | 0.076011013 | 16.91641761 | 0.142846681 | 1.004930844 | 0.007096224 | 0.006617364 | 0.92382025 | 0.034412522 | Unpaired t-test | 0.971155121 | permutation FDR (250 perm) | 1 | 0.936148458 | FDR correction BH |
| Pak3~Q9VEV1 | VMAT~Control | VMAT | Control | 15.8732887 | 0.22064415 | 15.86770946 | 0.582293176 | 1.003874727 | 0.005579248 | 0.004363977 | 0.984409723 | 0.006824105 | Unpaired t-test | 0.981026774 | permutation FDR (250 perm) | 1 | 0.985665882 | FDR correction BH |
| gek~Q9W1B0 | VMAT~Control | VMAT | Control | 17.5097081 | 0.255749079 | 17.50592643 | 0.257738445 | 1.002624692 | 0.00378167 | 0.003253384 | 0.981877928 | 0.007942502 | Unpaired t-test | 0.985359388 | permutation FDR (250 perm) | 1 | 0.984386986 | FDR correction BH |
| RhoGAP93B~Q9VDE9 | VMAT~Control | VMAT | Control | 16.03303735 | 0.289798422 | 16.02968444 | 0.080680947 | 1.002326766 | 0.003352915 | 0.002955332 | 0.980606674 | 0.008505155 | Unpaired t-test | 0.986650807 | permutation FDR (250 perm) | 1 | 0.983531363 | FDR correction BH |
| Mlc1~A0A0B4LHS1 | VMAT~Control | VMAT | Control | 20.44872314 | 0.290321648 | 20.44643492 | 0.328738051 | 1.001587333 | 0.002288222 | 0.001913004 | 0.990921362 | 0.003960809 | Unpaired t-test | 0.991469839 | permutation FDR (250 perm) | 1 | 0.991342493 | FDR correction BH |
| RpL36A~Q9VLT7 | VMAT~Control | VMAT | Control | 20.11065506 | 0.221457941 | 20.11180418 | 0.557899804 | 0.999203807 | -0.001149122 | -0.000905934 | 0.996668648 | 0.001449203 | Unpaired t-test | 0.995937128 | permutation FDR (250 perm) | 1 | 0.996668648 | FDR correction BH |
| Arpc2~X2JAI2 | VMAT~Control | VMAT | Control | 15.00785721 | 0.206478211 | 15.01110145 | 0.26658911 | 0.997753791 | -0.003244239 | -0.002819117 | 0.983259222 | 0.007331972 | Unpaired t-test | 0.987250637 | permutation FDR (250 perm) | 1 | 0.984932855 | FDR correction BH |
| Dmel\CG5653~Q9VSR8 | VMAT~Control | VMAT | Control | 16.40970115 | 0.092281335 | 16.4154642 | 0.259125647 | 0.996013322 | -0.005763056 | -0.005131777 | 0.963556091 | 0.016122999 | Unpaired t-test | 0.977718657 | permutation FDR (250 perm) | 1 | 0.968493184 | FDR correction BH |
| ArfGAP3~M9PGG0 | VMAT~Control | VMAT | Control | 19.99718305 | 0.076940112 | 20.00328312 | 0.082487473 | 0.995780675 | -0.006100077 | -0.005807131 | 0.906147204 | 0.042801245 | Unpaired t-test | 0.974607735 | permutation FDR (250 perm) | 1 | 0.921191559 | FDR correction BH |
| fws~M9NCV1 | VMAT~Control | VMAT | Control | 16.02077611 | 0.207352218 | 16.0281595 | 0.260943602 | 0.994895302 | -0.007383384 | -0.006425616 | 0.961468574 | 0.017064906 | Unpaired t-test | 0.971945601 | permutation FDR (250 perm) | 1 | 0.96722095 | FDR correction BH |
| betaCOP~P45437 | VMAT~Control | VMAT | Control | 17.0385487 | 0.117860211 | 17.04595232 | 0.063027402 | 0.99488135 | -0.007403616 | -0.006986046 | 0.903877119 | 0.043890607 | Unpaired t-test | 0.969470463 | permutation FDR (250 perm) | 1 | 0.919501615 | FDR correction BH |
| Hsp68~O97125 | VMAT~Control | VMAT | Control | 17.33372357 | 0.165251779 | 17.34158801 | 0.153058251 | 0.994563619 | -0.007864436 | -0.007144731 | 0.939310381 | 0.027190878 | Unpaired t-test | 0.968698683 | permutation FDR (250 perm) | 1 | 0.949393146 | FDR correction BH |
| RpS3~G3M3A2 | VMAT~Control | VMAT | Control | 20.52459664 | 0.131023672 | 20.53343221 | 0.071615033 | 0.993894364 | -0.008835572 | -0.008282491 | 0.897360106 | 0.047033242 | Unpaired t-test | 0.963738207 | permutation FDR (250 perm) | 1 | 0.914056984 | FDR correction BH |
| CG1846~Q9VY47 | VMAT~Control | VMAT | Control | 13.50421209 | 0.251862183 | 13.5148964 | 0.357927334 | 0.992621555 | -0.010684311 | -0.008935405 | 0.957542171 | 0.01884209 | Unpaired t-test | 0.96078878 | permutation FDR (250 perm) | 1 | 0.963682886 | FDR correction BH |
| alphaCOP~Q9W0B8 | VMAT~Control | VMAT | Control | 17.9769025 | 0.16597391 | 17.98859217 | 0.220301545 | 0.991930077 | -0.011689669 | -0.010406049 | 0.926372873 | 0.033214171 | Unpaired t-test | 0.954235444 | permutation FDR (250 perm) | 1 | 0.938331215 | FDR correction BH |
| Sec24CD~Q9VQ94 | VMAT~Control | VMAT | Control | 16.78463277 | 0.161949367 | 16.79732908 | 0.462046603 | 0.991238198 | -0.012696311 | -0.010415702 | 0.95490294 | 0.02004077 | Unpaired t-test | 0.954185945 | permutation FDR (250 perm) | 1 | 0.961437776 | FDR correction BH |
| CG11572~H9XVM5 | VMAT~Control | VMAT | Control | 15.75602667 | 0.258363595 | 15.77131357 | 0.153814864 | 0.989459868 | -0.015286901 | -0.013474929 | 0.9117393 | 0.040129325 | Unpaired t-test | 0.940797619 | permutation FDR (250 perm) | 1 | 0.925899186 | FDR correction BH |
| Mob4~A0A0B4K6X9 | VMAT~Control | VMAT | Control | 17.15305144 | 0.100498999 | 17.16849452 | 0.248221162 | 0.98935276 | -0.01544308 | -0.013791404 | 0.899954744 | 0.045779329 | Unpaired t-test | 0.939455782 | permutation FDR (250 perm) | 1 | 0.916303403 | FDR correction BH |
| Esyt2~Q9VC62 | VMAT~Control | VMAT | Control | 17.58145077 | 0.070204389 | 17.59934845 | 0.087782514 | 0.98767091 | -0.017897676 | -0.017041054 | 0.729204328 | 0.137150763 | Unpaired t-test | 0.925081633 | permutation FDR (250 perm) | 1 | 0.755619842 | FDR correction BH |
| Ssrp~Q05344 | VMAT~Control | VMAT | Control | 16.946311 | 0.169409292 | 16.96641266 | 0.672649015 | 0.986163209 | -0.020101664 | -0.015342304 | 0.949610462 | 0.022454509 | Unpaired t-test | 0.932554422 | permutation FDR (250 perm) | 1 | 0.958569051 | FDR correction BH |
| jdp~Q9TVP3-2 | VMAT~Control | VMAT | Control | 17.89539102 | 0.257121917 | 17.91630002 | 0.231679446 | 0.985611506 | -0.020908997 | -0.018106448 | 0.89522356 | 0.048068497 | Unpaired t-test | 0.920232993 | permutation FDR (250 perm) | 1 | 0.912275438 | FDR correction BH |
| dsh~P51140 | VMAT~Control | VMAT | Control | 17.35766248 | 0.205035786 | 17.3791023 | 0.423628191 | 0.985248925 | -0.021439825 | -0.017711897 | 0.920878561 | 0.035797638 | Unpaired t-test | 0.921993197 | permutation FDR (250 perm) | 1 | 0.933569394 | FDR correction BH |
| loopin-1~Q7K2S9 | VMAT~Control | VMAT | Control | 15.83328444 | 0.362766155 | 15.85686945 | 0.265513179 | 0.983785022 | -0.023585005 | -0.019637061 | 0.908935746 | 0.041466817 | Unpaired t-test | 0.914028073 | permutation FDR (250 perm) | 1 | 0.923450473 | FDR correction BH |
| Gdi~Q9VLB7 | VMAT~Control | VMAT | Control | 15.01741992 | 0.223979905 | 15.04159974 | 0.268920422 | 0.983379496 | -0.024179821 | -0.02090748 | 0.88029804 | 0.055370265 | Unpaired t-test | 0.909041702 | permutation FDR (250 perm) | 1 | 0.898621676 | FDR correction BH |
| Nph~Q9VAC4 | VMAT~Control | VMAT | Control | 16.75004926 | 0.183625172 | 16.77440808 | 0.250353419 | 0.983257492 | -0.024358821 | -0.021388985 | 0.864240015 | 0.06336563 | Unpaired t-test | 0.906956596 | permutation FDR (250 perm) | 1 | 0.883762378 | FDR correction BH |
| l(1)G0196~X2JCY0 | VMAT~Control | VMAT | Control | 16.46810559 | 0.256616936 | 16.49263129 | 0.238132657 | 0.983143759 | -0.024525706 | -0.021205687 | 0.878637403 | 0.056190313 | Unpaired t-test | 0.907739574 | permutation FDR (250 perm) | 1 | 0.897315595 | FDR correction BH |
| Panx~Q9W2H9 | VMAT~Control | VMAT | Control | 14.34742931 | 1.02949484 | 14.37276384 | 1.057159642 | 0.982592726 | -0.025334536 | -0.01526254 | 0.970131906 | 0.013169212 | Unpaired t-test | 0.932916667 | permutation FDR (250 perm) | 1 | 0.974686516 | FDR correction BH |

| | | | | | | | | | | | | | | | | | | |
|---|---|---|---|---|---|---|---|---|---|---|---|---|---|---|---|---|---|---|
| Gss2~Q86B44 | VMAT~Control | VMAT | Control | 16.41887863 | 0.125601423 | 16.44450522 | 0.131576945 | 0.982393833 | -0.02562659 | -0.023698725 | 0.759220998 | 0.119631789 | Unpaired t-test | 0.898253089 | permutation FDR (250 perm) | 1 | 0.784206331 | FDR correction BH |
| Tbc1d22~Q95RC5 | VMAT~Control | VMAT | Control | 16.26557366 | 0.10731999 | 16.29337891 | 0.055797775 | 0.980911409 | -0.02780525 | -0.026378332 | 0.618408439 | 0.208724593 | Unpaired t-test | 0.887229327 | permutation FDR (250 perm) | 1 | 0.651918256 | FDR correction BH |
| Srrm1~Q9VU43 | VMAT~Control | VMAT | Control | 17.93041251 | 0.522223321 | 17.95863184 | 0.863802027 | 0.980629908 | -0.028219331 | -0.019442654 | 0.951385967 | 0.021643259 | Unpaired t-test | 0.914907699 | permutation FDR (250 perm) | 1 | 0.958716852 | FDR correction BH |
| Dmel\CG6071~Q9VTL4 | VMAT~Control | VMAT | Control | 21.6060212 | 0.452155024 | 21.63472199 | 0.646599462 | 0.980302703 | -0.028700794 | -0.021214974 | 0.936777569 | 0.028363517 | Unpaired t-test | 0.907714043 | permutation FDR (250 perm) | 1 | 0.947239861 | FDR correction BH |
| Drice~O01382 | VMAT~Control | VMAT | Control | 13.78616161 | 0.815586051 | 13.81534132 | 0.771192814 | 0.979977335 | -0.029179712 | -0.019427498 | 0.954790851 | 0.020091751 | Unpaired t-test | 0.914982561 | permutation FDR (250 perm) | 1 | 0.961437776 | FDR correction BH |
| Sec24AB~A0A0B4LEK8 | VMAT~Control | VMAT | Control | 15.53992744 | 0.25551878 | 15.57073682 | 0.282733509 | 0.978870977 | -0.030809381 | -0.026323181 | 0.860154017 | 0.065423778 | Unpaired t-test | 0.887459506 | permutation FDR (250 perm) | 1 | 0.879966343 | FDR correction BH |
| Dmel\CG6907~X2J4N6 | VMAT~Control | VMAT | Control | 15.54529757 | 0.158885391 | 15.58004983 | 0.08858676 | 0.976199378 | -0.034752262 | -0.032137736 | 0.678303741 | 0.168575788 | Unpaired t-test | 0.863038363 | permutation FDR (250 perm) | 1 | 0.706829131 | FDR correction BH |
| EG:BACR25B3.6~Q7K6T9 | VMAT~Control | VMAT | Control | 16.07719442 | 0.065511768 | 16.11245195 | 0.152026947 | 0.975857544 | -0.035257537 | -0.032827256 | 0.644134491 | 0.191023446 | Unpaired t-test | 0.860262575 | permutation FDR (250 perm) | 1 | 0.67510801 | FDR correction BH |
| kmr~A8JR01 | VMAT~Control | VMAT | Control | 17.07418271 | 0.154425196 | 17.10987059 | 0.221338861 | 0.9755665 | -0.035687877 | -0.031844378 | 0.773520744 | 0.111528035 | Unpaired t-test | 0.86427792 | permutation FDR (250 perm) | 1 | 0.797926307 | FDR correction BH |
| TER94~Q7KN62 | VMAT~Control | VMAT | Control | 18.59732614 | 0.081940972 | 18.63607261 | 0.088590093 | 0.973500438 | -0.038746467 | -0.036762482 | 0.489216703 | 0.310498723 | Unpaired t-test | 0.845558685 | permutation FDR (250 perm) | 1 | 0.525852109 | FDR correction BH |
| snama~Q9XZ21 | VMAT~Control | VMAT | Control | 17.33700278 | 0.355660187 | 17.37775273 | 0.481667612 | 0.972149474 | -0.040749941 | -0.032143064 | 0.882068539 | 0.054497668 | Unpaired t-test | 0.863017903 | permutation FDR (250 perm) | 1 | 0.900038726 | FDR correction BH |
| alpha-Cat~P35220 | VMAT~Control | VMAT | Control | 15.57559558 | 0.134612074 | 15.62109147 | 0.336887612 | 0.968956708 | -0.045495886 | -0.039144906 | 0.784872093 | 0.105201113 | Unpaired t-test | 0.836157131 | permutation FDR (250 perm) | 1 | 0.807865722 | FDR correction BH |
| dSpb1~Q9VXK5 | VMAT~Control | VMAT | Control | 18.11034722 | 0.39919341 | 18.15902945 | 0.506548035 | 0.966819029 | -0.048682226 | -0.037784277 | 0.869330043 | 0.060815311 | Unpaired t-test | 0.841379428 | permutation FDR (250 perm) | 1 | 0.888581382 | FDR correction BH |
| jp~Q7JV09 | VMAT~Control | VMAT | Control | 15.94491087 | 0.076908981 | 15.99643702 | 0.131219354 | 0.964915055 | -0.051526153 | -0.048244564 | 0.466215747 | 0.331413062 | Unpaired t-test | 0.80094359 | permutation FDR (250 perm) | 1 | 0.503889747 | FDR correction BH |
| Dmel\CG13842~Q9VCV8 | VMAT~Control | VMAT | Control | 22.72772636 | 0.045349654 | 22.78014436 | 0.474017132 | 0.964318751 | -0.052417993 | -0.043215123 | 0.810547902 | 0.091221314 | Unpaired t-test | 0.820006832 | permutation FDR (250 perm) | 1 | 0.832837085 | FDR correction BH |
| ringer~M9NFZ9 | VMAT~Control | VMAT | Control | 18.59090064 | 0.193803083 | 18.64730405 | 0.292859854 | 0.961658518 | -0.056403406 | -0.048747516 | 0.726958735 | 0.138490241 | Unpaired t-test | 0.79914188 | permutation FDR (250 perm) | 1 | 0.753859411 | FDR correction BH |
| Asator~L0MPN7 | VMAT~Control | VMAT | Control | 17.70790918 | 0.071822577 | 17.76593225 | 0.079231914 | 0.960579502 | -0.058023072 | -0.055374771 | 0.252918887 | 0.597018737 | Unpaired t-test | 0.77493071 | permutation FDR (250 perm) | 1 | 0.286787602 | FDR correction BH |
| Pp2B-14D~M9NE01 | VMAT~Control | VMAT | Control | 16.9498991 | 0.06888397 | 17.00799054 | 0.113756298 | 0.960533986 | -0.058091434 | -0.054830473 | 0.351724301 | 0.453797625 | Unpaired t-test | 0.776947819 | permutation FDR (250 perm) | 1 | 0.388713147 | FDR correction BH |
| Sarm~A8JNN3 | VMAT~Control | VMAT | Control | 17.86577222 | 0.173152373 | 17.92413666 | 0.102669051 | 0.960352239 | -0.058364439 | -0.053544119 | 0.531386878 | 0.274589174 | Unpaired t-test | 0.781448482 | permutation FDR (250 perm) | 1 | 0.567552047 | FDR correction BH |
| Men~Q2QBM1 | VMAT~Control | VMAT | Control | 15.62206112 | 0.17549087 | 15.68290061 | 0.26385653 | 0.958706092 | -0.060839496 | -0.05328776 | 0.676803249 | 0.169537565 | Unpaired t-test | 0.782438649 | permutation FDR (250 perm) | 1 | 0.705890496 | FDR correction BH |
| Mo25~M9PI82 | VMAT~Control | VMAT | Control | 17.17074834 | 0.128162475 | 17.23702551 | 0.076868128 | 0.955099422 | -0.066277175 | -0.06212507 | 0.344751874 | 0.462493365 | Unpaired t-test | 0.751299658 | permutation FDR (250 perm) | 1 | 0.381724323 | FDR correction BH |
| wdb~Q9VB23 | VMAT~Control | VMAT | Control | 17.49220621 | 0.050524683 | 17.55867526 | 0.115578729 | 0.954972409 | -0.066469044 | -0.062919662 | 0.265958188 | 0.575186635 | Unpaired t-test | 0.748820557 | permutation FDR (250 perm) | 1 | 0.299552906 | FDR correction BH |
| CG9545~Q9VMC7 | VMAT~Control | VMAT | Control | 16.68794153 | 0.074271905 | 16.75578275 | 0.117755784 | 0.954064546 | -0.067841222 | -0.063864871 | 0.301445105 | 0.520791763 | Unpaired t-test | 0.746478559 | permutation FDR (250 perm) | 1 | 0.336942915 | FDR correction BH |
| Dmel\CG15784~Q9W4C1 | VMAT~Control | VMAT | Control | 15.37643749 | 0.48686848 | 15.45278275 | 0.62245315 | 0.948457308 | -0.076345259 | -0.056409608 | 0.833310786 | 0.079192997 | Unpaired t-test | 0.771163388 | permutation FDR (250 perm) | 1 | 0.85361775 | FDR correction BH |
| anon-WO0118547.327~Q9VHI8 | VMAT~Control | VMAT | Control | 15.58951064 | 0.120360074 | 15.66587735 | 0.079814816 | 0.948443207 | -0.076366707 | -0.071733693 | 0.264399036 | 0.577740133 | Unpaired t-test | 0.719883262 | permutation FDR (250 perm) | 1 | 0.298224883 | FDR correction BH |
| APC10~Q9V831 | VMAT~Control | VMAT | Control | 13.75769794 | 0.306950169 | 13.8345404 | 0.406124632 | 0.948130496 | -0.076842457 | -0.062592392 | 0.742699008 | 0.129187156 | Unpaired t-test | 0.749898073 | permutation FDR (250 perm) | 1 | 0.768489435 | FDR correction BH |
| Dmel\CG13185~A8DYB0 | VMAT~Control | VMAT | Control | 18.95706254 | 0.180323215 | 19.03455535 | 0.227826338 | 0.947703186 | -0.077492807 | -0.068581381 | 0.56417939 | 0.248582783 | Unpaired t-test | 0.730199914 | permutation FDR (250 perm) | 1 | 0.60012575 | FDR correction BH |
| Pde8~Q6NNF2-4 | VMAT~Control | VMAT | Control | 13.54960656 | 0.341406186 | 13.62920036 | 0.482251242 | 0.946324056 | -0.079593795 | -0.062957622 | 0.769422007 | 0.113835396 | Unpaired t-test | 0.749344192 | permutation FDR (250 perm) | 1 | 0.794046209 | FDR correction BH |
| Usp2~X2JLM6 | VMAT~Control | VMAT | Control | 14.12496566 | 0.588770796 | 14.20846644 | 0.644030544 | 0.943764766 | -0.083500784 | -0.060062245 | 0.834866984 | 0.078382713 | Unpaired t-test | 0.758660959 | permutation FDR (250 perm) | 1 | 0.854468209 | FDR correction BH |
| rump~Q9VHC7 | VMAT~Control | VMAT | Control | 15.15989312 | 0.147838651 | 15.24543306 | 0.426664691 | 0.942431759 | -0.085539938 | -0.07116822 | 0.680830846 | 0.166960776 | Unpaired t-test | 0.721852361 | permutation FDR (250 perm) | 1 | 0.709148589 | FDR correction BH |
| ClC-b~Q7JZ25 | VMAT~Control | VMAT | Control | 16.85961062 | 0.29356255 | 16.94592254 | 0.207484682 | 0.941927603 | -0.086311917 | -0.07435771 | 0.603090686 | 0.219617379 | Unpaired t-test | 0.711150215 | permutation FDR (250 perm) | 1 | 0.638056393 | FDR correction BH |
| Dhrs4~Q9VRJ4 | VMAT~Control | VMAT | Control | 18.69098476 | 0.045983476 | 18.77999914 | 0.208195827 | 0.940164831 | -0.089014381 | -0.081265551 | 0.372536215 | 0.428831502 | Unpaired t-test | 0.689888316 | permutation FDR (250 perm) | 1 | 0.409598436 | FDR correction BH |
| Dmel\CG6937~Q9VCY1 | VMAT~Control | VMAT | Control | 14.78822956 | 0.168118443 | 14.8781354 | 0.295389029 | 0.939584068 | -0.089905844 | -0.078043335 | 0.56732207 | 0.246170321 | Unpaired t-test | 0.699471018 | permutation FDR (250 perm) | 1 | 0.602612356 | FDR correction BH |
| betaggt-I~M9PEL6 | VMAT~Control | VMAT | Control | 16.09934541 | 0.114333545 | 16.19035427 | 0.252348628 | 0.938865978 | -0.091008865 | -0.080976181 | 0.479481493 | 0.319228151 | Unpaired t-test | 0.690666667 | permutation FDR (250 perm) | 1 | 0.516567246 | FDR correction BH |
| Ca-alpha1T~M9NGR9 | VMAT~Control | VMAT | Control | 20.72563822 | 0.101890538 | 20.81821671 | 0.138706748 | 0.937845061 | -0.092578496 | -0.08596207 | 0.256758198 | 0.590475682 | Unpaired t-test | 0.675449936 | permutation FDR (250 perm) | 1 | 0.290023415 | FDR correction BH |
| Dbp80~O61305 | VMAT~Control | VMAT | Control | 15.87282576 | 0.108848159 | 15.96575341 | 0.060992358 | 0.937618119 | -0.092927645 | -0.088016365 | 0.126807369 | 0.896855507 | Unpaired t-test | 0.669676698 | permutation FDR (250 perm) | 1 | 0.14985168 | FDR correction BH |
| shi~X2JE15 | VMAT~Control | VMAT | Control | 14.57990545 | 1.811907931 | 14.67875025 | 1.76819499 | 0.933780392 | -0.0988448 | -0.04635786 | 0.932154638 | 0.030512035 | Unpaired t-test | 0.807596926 | permutation FDR (250 perm) | 1 | 0.942970356 | FDR correction BH |
| spas~A0A126GV13 | VMAT~Control | VMAT | Control | 16.00264214 | 0.286569362 | 16.10274361 | 0.350581839 | 0.932967375 | -0.100101463 | -0.083244515 | 0.631751478 | 0.199453733 | Unpaired t-test | 0.683903739 | permutation FDR (250 perm) | 1 | 0.664793464 | FDR correction BH |
| csw~P29349-2 | VMAT~Control | VMAT | Control | 16.70624078 | 0.106294493 | 16.81119754 | 0.183402876 | 0.92983281 | -0.104956761 | -0.09586844 | 0.294153772 | 0.531425578 | Unpaired t-test | 0.647221696 | permutation FDR (250 perm) | 1 | 0.328949159 | FDR correction BH |
| Tailor~Q86BP3 | VMAT~Control | VMAT | Control | 13.80031057 | 0.37621501 | 13.90548926 | 0.738296504 | 0.929689784 | -0.105178692 | -0.076740706 | 0.782332206 | 0.106608791 | Unpaired t-test | 0.703658222 | permutation FDR (250 perm) | 1 | 0.805958528 | FDR correction BH |
| Taf8~Q9VWY6 | VMAT~Control | VMAT | Control | 17.15389047 | 0.263224886 | 17.26103356 | 0.478580908 | 0.928424766 | -0.107143087 | -0.086109538 | 0.670251189 | 0.173762407 | Unpaired t-test | 0.675070047 | permutation FDR (250 perm) | 1 | 0.699987266 | FDR correction BH |
| RAF2~Q9VV52 | VMAT~Control | VMAT | Control | 16.77967935 | 0.316213549 | 16.8907942 | 0.358852339 | 0.925872313 | -0.11111485 | -0.091535424 | 0.614734868 | 0.211312153 | Unpaired t-test | 0.659415054 | permutation FDR (250 perm) | 1 | 0.648917435 | FDR correction BH |
| Rpn1~Q9VW54 | VMAT~Control | VMAT | Control | 16.4068828 | 0.107003592 | 16.51909273 | 0.088323424 | 0.925169797 | -0.112209926 | -0.105654124 | 0.100666382 | 0.99711554 | Unpaired t-test | 0.620626132 | permutation FDR (250 perm) | 1 | 0.120717607 | FDR correction BH |
| WASp~Q9VAT0 | VMAT~Control | VMAT | Control | 16.2544174 | 0.175431762 | 16.36883701 | 0.304162997 | 0.923753855 | -0.114419614 | -0.09889083 | 0.482930648 | 0.316115233 | Unpaired t-test | 0.638806202 | permutation FDR (250 perm) | 1 | 0.520045171 | FDR correction BH |
| anon-EST:Posey214~Q9VZ01 | VMAT~Control | VMAT | Control | 14.81226017 | 2.012378305 | 14.93041025 | 1.598710445 | 0.92136833 | -0.118150085 | -0.054969002 | 0.920159174 | 0.03613704 | Unpaired t-test | 0.776484175 | permutation FDR (250 perm) | 1 | 0.933242006 | FDR correction BH |
| Vps60~M9PCV9 | VMAT~Control | VMAT | Control | 18.71087602 | 0.088311258 | 18.82958631 | 0.096045499 | 0.921010632 | -0.118710285 | -0.112165431 | 0.069287136 | 1.159347391 | Unpaired t-test | 0.603345988 | permutation FDR (250 perm) | 1 | 0.084772306 | FDR correction BH |
| Pp2A-29B~M9NDI6 | VMAT~Control | VMAT | Control | 18.17185752 | 0.069975915 | 18.29284258 | 0.096621622 | 0.919559571 | -0.120985056 | -0.114857157 | 0.046756025 | 1.330162418 | Unpaired t-test | 0.596562797 | permutation FDR (250 perm) | 1 | 0.058575669 | FDR correction BH |
| Pp1-87B~P12982 | VMAT~Control | VMAT | Control | 16.40518856 | 0.049396199 | 16.52751812 | 0.337972206 | 0.918702999 | -0.122329557 | -0.10611962 | 0.44183586 | 0.354739039 | Unpaired t-test | 0.619329021 | permutation FDR (250 perm) | 1 | 0.479742442 | FDR correction BH |
| IMP~Q9VUW2 | VMAT~Control | VMAT | Control | 16.23124057 | 0.162317439 | 16.35379935 | 0.264675018 | 0.918557041 | -0.122558782 | -0.107616046 | 0.398143686 | 0.399960167 | Unpaired t-test | 0.615149633 | permutation FDR (250 perm) | 1 | 0.43511153 | FDR correction BH |
| Rpn8~A0A0B4KFH4 | VMAT~Control | VMAT | Control | 16.12967096 | 0.15890179 | 16.25294967 | 0.13777939 | 0.918098778 | -0.123278714 | -0.112680411 | 0.219264088 | 0.659032493 | Unpaired t-test | 0.601987921 | permutation FDR (250 perm) | 1 | 0.249950442 | FDR correction BH |
| Lmpt~Q9VVB7 | VMAT~Control | VMAT | Control | 17.48349245 | 0.136044433 | 17.60954614 | 0.07620208 | 0.916334544 | -0.126053688 | -0.11783637 | 0.100794338 | 0.996563865 | Unpaired t-test | 0.588841105 | permutation FDR (250 perm) | 1 | 0.120809506 | FDR correction BH |
| metro~A0A0B4LF49 | VMAT~Control | VMAT | Control | 15.98248061 | 0.172026967 | 16.10921602 | 0.225006062 | 0.915901642 | -0.126735417 | -0.112487153 | 0.340639639 | 0.467704816 | Unpaired t-test | 0.602488352 | permutation FDR (250 perm) | 1 | 0.377526229 | FDR correction BH |
| Rpt5~Q9V3V6 | VMAT~Control | VMAT | Control | 17.64266623 | 0.087131121 | 17.77162473 | 0.101693383 | 0.914491394 | -0.128958501 | -0.121671721 | 0.05674791 | 1.246050127 | Unpaired t-test | 0.579891098 | permutation FDR (250 perm) | 1 | 0.070418862 | FDR correction BH |
| Moca-cyp~Q9VAW1 | VMAT~Control | VMAT | Control | 16.82514512 | 0.272867543 | 16.95505895 | 0.612068374 | 0.913886033 | -0.129913832 | -0.099957196 | 0.673862636 | 0.171428624 | Unpaired t-test | 0.635848407 | permutation FDR (250 perm) | 1 | 0.703135038 | FDR correction BH |
| pix~Q9VSS1 | VMAT~Control | VMAT | Control | 16.02246896 | 0.136006653 | 16.15264226 | 0.1217731 | 0.913721687 | -0.130173297 | -0.120347937 | 0.141918567 | 0.847960782 | Unpaired t-test | 0.58252677 | permutation FDR (250 perm) | 1 | 0.166704744 | FDR correction BH |
| arg~O76895 | VMAT~Control | VMAT | Control | 18.24342128 | 0.189143815 | 18.3739201 | 0.396678884 | 0.913515541 | -0.130498822 | -0.109063964 | 0.521705514 | 0.282574573 | Unpaired t-test | 0.611242777 | permutation FDR (250 perm) | 1 | 0.557971277 | FDR correction BH |
| mGluR~L0MPZ9 | VMAT~Control | VMAT | Control | 16.13594029 | 0.147798761 | 16.26871144 | 0.184862729 | 0.912077834 | -0.132771149 | -0.120062778 | 0.238237696 | 0.622989521 | Unpaired t-test | 0.583253886 | permutation FDR (250 perm) | 1 | 0.270661938 | FDR correction BH |
| Slmap~Q9W5R5 | VMAT~Control | VMAT | Control | 18.0647891 | 0.202340671 | 18.20143437 | 0.150158811 | 0.909631886 | -0.136645269 | -0.122806799 | 0.253139885 | 0.596639421 | Unpaired t-test | 0.576954192 | permutation FDR (250 perm) | 1 | 0.286899995 | FDR correction BH |
| sxc~Q7KJA9 | VMAT~Control | VMAT | Control | 16.73246437 | 0.080643089 | 16.8704322 | 0.113872886 | 0.908798379 | -0.137967834 | -0.129863999 | 0.051479056 | 1.288369429 | Unpaired t-test | 0.56066984 | permutation FDR (250 perm) | 1 | 0.064151242 | FDR correction BH |
| Dlic~Q9VZ20 | VMAT~Control | VMAT | Control | 17.65988956 | 0.059623822 | 17.79969634 | 0.2238728 | 0.90764071 | -0.139806776 | -0.126681449 | 0.206974352 | 0.684083468 | Unpaired t-test | 0.568243945 | permutation FDR (250 perm) | 1 | 0.23651341 | FDR correction BH |
| Cdep~A0A0C4DHA1 | VMAT~Control | VMAT | Control | 16.91781851 | 0.09543147 | 17.05767767 | 0.248870524 | 0.907607749 | -0.139859168 | -0.124963469 | 0.267859968 | 0.572092187 | Unpaired t-test | 0.571941202 | permutation FDR (250 perm) | 1 | 0.301194784 | FDR correction BH |
| Tm1~Q6NN28 | VMAT~Control | VMAT | Control | 16.36682418 | 0.095386207 | 16.50695737 | 0.216248642 | 0.907435371 | -0.140133199 | -0.126737137 | 0.21440294 | 0.668769263 | Unpaired t-test | 0.568074394 | permutation FDR (250 perm) | 1 | 0.244645914 | FDR correction BH |
| Lnk~A0A0B4LIP5 | VMAT~Control | VMAT | Control | 13.37850398 | 0.314135436 | 13.51923228 | 0.326707583 | 0.90706114 | -0.140728297 | -0.117011149 | 0.503305259 | 0.298168532 | Unpaired t-test | 0.590843332 | permutation FDR (250 perm) | 1 | 0.539517568 | FDR correction BH |
| Grx4~Q9VJZ6 | VMAT~Control | VMAT | Control | 16.80505732 | 0.089136791 | 16.95173338 | 0.256734819 | 0.903329318 | -0.146676062 | -0.130781111 | 0.255271161 | 0.592998247 | Unpaired t-test | 0.558540891 | permutation FDR (250 perm) | 1 | 0.288583995 | FDR correction BH |
| Tps1~Q9Y119 | VMAT~Control | VMAT | Control | 18.79190033 | 0.088714358 | 18.94097932 | 0.058307243 | 0.901826005 | -0.149078984 | -0.142322065 | 0.010508833 | 1.978445491 | Unpaired t-test | 0.532623647 | permutation FDR (250 perm) | 1 | 0.014466546 | FDR correction BH |
| Dmel\CG9170~Q9VXK9 | VMAT~Control | VMAT | Control | 15.60952961 | 0.202680241 | 15.76552384 | 0.093566887 | 0.89751364 | -0.155994231 | -0.141834353 | 0.149225501 | 0.826156955 | Unpaired t-test | 0.533711563 | permutation FDR (250 perm) | 1 | 0.174244459 | FDR correction BH |
| Dmel\CG9784~Q9VXE7 | VMAT~Control | VMAT | Control | 16.36208493 | 0.091082241 | 16.52041891 | 0.113447371 | 0.896059239 | -0.158333982 | -0.148661545 | 0.035236755 | 1.453004095 | Unpaired t-test | 0.518679376 | permutation FDR (250 perm) | 1 | 0.044933543 | FDR correction BH |
| kek5~Q9VWI6 | VMAT~Control | VMAT | Control | 15.52356591 | 0.2948895 | 15.68341127 | 0.531354965 | 0.895121011 | -0.159845362 | -0.125687201 | 0.569468016 | 0.244530663 | Unpaired t-test | 0.570234328 | permutation FDR (250 perm) | 1 | 0.604385802 | FDR correction BH |
| MYPT-75D~M9PFV2 | VMAT~Control | VMAT | Control | 16.04420659 | 0.122547891 | 16.20520191 | 0.10087399 | 0.894407806 | -0.160995315 | -0.150324687 | 0.046724754 | 1.330452973 | Unpaired t-test | 0.51363676 | permutation FDR (250 perm) | 1 | 0.058573962 | FDR correction BH |
| CG32543~Q9VWP8 | VMAT~Control | VMAT | Control | 16.82179361 | 0.174241842 | 16.98626174 | 0.178360417 | 0.892257399 | -0.164468135 | -0.14796817 | 0.17101003 | 0.766978417 | Unpaired t-test | 0.520102209 | permutation FDR (250 perm) | 1 | 0.197622784 | FDR correction BH |
| Ndae1~M9PC92 | VMAT~Control | VMAT | Control | 15.36649648 | 0.162971942 | 15.532352 | 0.194812391 | 0.891399764 | -0.165855516 | -0.148937887 | 0.174942304 | 0.757105157 | Unpaired t-test | 0.518069324 | permutation FDR (250 perm) | 1 | 0.201770791 | FDR correction BH |
| Dmel\CG15894~X2JIM3 | VMAT~Control | VMAT | Control | 17.48509985 | 0.146670563 | 17.65170653 | 0.132446292 | 0.890935766 | -0.166606673 | -0.153077804 | 0.088769905 | 1.051734246 | Unpaired t-test | 0.509431049 | permutation FDR (250 perm) | 1 | 0.107051412 | FDR correction BH |
| Dmel\CG7878~Q7K4L8 | VMAT~Control | VMAT | Control | 15.00530369 | 0.122504403 | 15.17425402 | 0.364168439 | 0.889489618 | -0.168950328 | -0.144176609 | 0.348676343 | 0.457577518 | Unpaired t-test | 0.528476397 | permutation FDR (250 perm) | 1 | 0.385525651 | FDR correction BH |
| cg9346~Q9W2N6 | VMAT~Control | VMAT | Control | 17.83214933 | 0.215677472 | 18.00363312 | 0.488473116 | 0.887928992 | -0.171483787 | -0.138427546 | 0.489125562 | 0.31057964 | Unpaired t-test | 0.541448246 | permutation FDR (250 perm) | 1 | 0.525852109 | FDR correction BH |
| Hsc70-3~F3YDH0 | VMAT~Control | VMAT | Control | 17.95026894 | 0.087181283 | 18.12367092 | 0.108919046 | 0.886749192 | -0.173401984 | -0.163218427 | 0.019476532 | 1.710488363 | Unpaired t-test | 0.48847069 | permutation FDR (250 perm) | 1 | 0.025771645 | FDR correction BH |
| LeuRS~Q9VQR8 | VMAT~Control | VMAT | Control | 16.03852643 | 0.102111287 | 16.21256823 | 0.068779868 | 0.886356018 | -0.174041799 | -0.164959328 | 0.010141313 | 1.993905794 | Unpaired t-test | 0.484884064 | permutation FDR (250 perm) | 1 | 0.014017999 | FDR correction BH |
| His3.3A~C0HL66 | VMAT~Control | VMAT | Control | 18.00256757 | 0.36309716 | 18.17837752 | 0.440142481 | 0.885270375 | -0.175809952 | -0.140068362 | 0.506503284 | 0.295417734 | Unpaired t-test | 0.537668255 | permutation FDR (250 perm) | 1 | 0.542698558 | FDR correction BH |
| Surf6~Q9VDS6 | VMAT~Control | VMAT | Control | 16.88860543 | 0.310998205 | 17.06964319 | 0.454947718 | 0.882068277 | -0.181037762 | -0.145242259 | 0.479471809 | 0.319236922 | Unpaired t-test | 0.526182763 | permutation FDR (250 perm) | 1 | 0.516567246 | FDR correction BH |
| Hex-C~Q7JYW9 | VMAT~Control | VMAT | Control | 15.80810264 | 0.168359161 | 15.98970907 | 0.095161494 | 0.881720663 | -0.181606425 | -0.167150023 | 0.062098379 | 1.206919738 | Unpaired t-test | 0.480225792 | permutation FDR (250 perm) | 1 | 0.076493764 | FDR correction BH |
| FBgn 31395~Q9VQ84 | VMAT~Control | VMAT | Control | 15.52603338 | 0.110513845 | 15.70889367 | 0.223769665 | 0.880954686 | -0.182860283 | -0.1645001 | 0.132386496 | 0.87815314 | Unpaired t-test | 0.485775076 | permutation FDR (250 perm) | 1 | 0.156131168 | FDR correction BH |
| orb2~A4V1P2 | VMAT~Control | VMAT | Control | 19.00888247 | 0.105448863 | 19.19496217 | 0.126399449 | 0.878991002 | -0.186079698 | -0.173320598 | 0.029987829 | 1.52305498 | Unpaired t-test | 0.4681755 | permutation FDR (250 perm) | 1 | 0.038511374 | FDR correction BH |
| SmydA-1~Q7K561 | VMAT~Control | VMAT | Control | 14.90748758 | 0.155556909 | 15.0943593 | 0.206961608 | 0.87850858 | -0.186871718 | -0.167480012 | 0.137612454 | 0.861342261 | Unpaired t-test | 0.479531046 | permutation FDR (250 perm) | 1 | 0.16180805 | FDR correction BH |
| AGO2~Q9VUQ5-2 | VMAT~Control | VMAT | Control | 14.26968599 | 0.298329477 | 14.45750255 | 0.352759201 | 0.87793342 | -0.18781656 | -0.155656326 | 0.384716035 | 0.414859712 | Unpaired t-test | 0.504114534 | permutation FDR (250 perm) | 1 | 0.421807893 | FDR correction BH |
| Dmel\CG12822~Q8MR50 | VMAT~Control | VMAT | Control | 13.37880684 | 0.551987798 | 13.56888042 | 0.159515099 | 0.876561013 | -0.190073582 | -0.1512172 | 0.476483182 | 0.321952423 | Unpaired t-test | 0.513340269 | permutation FDR (250 perm) | 1 | 0.514278501 | FDR correction BH |
| Rpn6~A0A0B4K7Z5 | VMAT~Control | VMAT | Control | 15.38609805 | 0.086525331 | 15.58036104 | 0.187225674 | 0.874019278 | -0.194262993 | -0.177857601 | 0.061445425 | 1.211510447 | Unpaired t-test | 0.459365493 | permutation FDR (250 perm) | 1 | 0.075768743 | FDR correction BH |
| eEF1gamma~A0A0B4JD11 | VMAT~Control | VMAT | Control | 18.030234 | 0.243066885 | 18.22451108 | 0.099311496 | 0.874010746 | -0.194277078 | -0.173861253 | 0.129035258 | 0.889291604 | Unpaired t-test | 0.467077324 | permutation FDR (250 perm) | 1 | 0.152331494 | FDR correction BH |
| eEF1alpha2~A4V3Q6 | VMAT~Control | VMAT | Control | 20.70640486 | 0.162384003 | 20.90346625 | 0.184442399 | 0.872325587 | -0.197061387 | -0.177549138 | 0.103199446 | 0.986322635 | Unpaired t-test | 0.45997914 | permutation FDR (250 perm) | 1 | 0.123503556 | FDR correction BH |
| SmB~Q05856 | VMAT~Control | VMAT | Control | 14.34497835 | 0.238124355 | 14.54340434 | 0.174569752 | 0.871500871 | -0.198425988 | -0.175281196 | 0.163813083 | 0.785651417 | Unpaired t-test | 0.464258905 | permutation FDR (250 perm) | 1 | 0.189865089 | FDR correction BH |
| Ttc7~A0A0B4K7H0 | VMAT~Control | VMAT | Control | 15.27891466 | 0.182262328 | 15.48000515 | 0.336097937 | 0.869892786 | -0.201090495 | -0.171727469 | 0.266808107 | 0.573800979 | Unpaired t-test | 0.471143354 | permutation FDR (250 perm) | 1 | 0.300222889 | FDR correction BH |
| Tsc1~Q9VCC9 | VMAT~Control | VMAT | Control | 16.55497712 | 0.219116842 | 16.76218311 | 0.545379238 | 0.866213168 | -0.20720599 | -0.164078065 | 0.448801952 | 0.347945263 | Unpaired t-test | 0.486695614 | permutation FDR (250 perm) | 1 | 0.486632794 | FDR correction BH |
| RpL19~P36241 | VMAT~Control | VMAT | Control | 18.42583316 | 0.065358812 | 18.63878495 | 0.224451149 | 0.86277018 | -0.212951781 | -0.192795629 | 0.069003538 | 1.161128644 | Unpaired t-test | 0.431088696 | permutation FDR (250 perm) | 1 | 0.084513178 | FDR correction BH |
| Dmel\CG8289~X2JFW6 | VMAT~Control | VMAT | Control | 15.65988461 | 0.24810656 | 15.8757899 | 0.474291213 | 0.86100571 | -0.215905289 | -0.174204559 | 0.388294877 | 0.410838339 | Unpaired t-test | 0.466350999 | permutation FDR (250 perm) | 1 | 0.425335571 | FDR correction BH |
| RyR~Q24498-5 | VMAT~Control | VMAT | Control | 21.35122375 | 0.082896379 | 21.56722288 | 0.110190406 | 0.860949706 | -0.215999132 | -0.20345292 | 0.005700708 | 2.24407123 | Unpaired t-test | 0.411829565 | permutation FDR (250 perm) | 1 | 0.00816766 | FDR correction BH |
| eIF3i~O02195 | VMAT~Control | VMAT | Control | 14.21371914 | 0.213674308 | 14.43177971 | 0.125554866 | 0.859720396 | -0.218060562 | -0.196303487 | 0.077471398 | 1.110858606 | Unpaired t-test | 0.424566957 | permutation FDR (250 perm) | 1 | 0.094052435 | FDR correction BH |
| CG17788~Q8SWS2 | VMAT~Control | VMAT | Control | 17.37039406 | 0.161774078 | 17.5885131 | 0.070497966 | 0.859685554 | -0.218119031 | -0.202164472 | 0.019997432 | 1.699025773 | Unpaired t-test | 0.414086957 | permutation FDR (250 perm) | 1 | 0.026401545 | FDR correction BH |
| mmd~M9NDY9 | VMAT~Control | VMAT | Control | 16.61059028 | 0.058434981 | 16.82882248 | 0.158260244 | 0.859618119 | -0.218232203 | -0.202922397 | 0.016037279 | 1.794869307 | Unpaired t-test | 0.412822609 | permutation FDR (250 perm) | 1 | 0.021413361 | FDR correction BH |
| Myo61F~Q23979-2 | VMAT~Control | VMAT | Control | 17.38158887 | 0.080073902 | 17.60023238 | 0.157458827 | 0.859373079 | -0.218643511 | -0.202635295 | 0.019867788 | 1.701850492 | Unpaired t-test | 0.413241739 | permutation FDR (250 perm) | 1 | 0.026245484 | FDR correction BH |
| KrT95D~Q9VCE7 | VMAT~Control | VMAT | Control | 18.05492613 | 0.139804088 | 18.27448139 | 0.153002601 | 0.858830149 | -0.219555257 | -0.200931403 | 0.039360075 | 1.404944084 | Unpaired t-test | 0.41628 | permutation FDR (250 perm) | 1 | 0.049840568 | FDR correction BH |
| Nup93-1~Q9XZ06 | VMAT~Control | VMAT | Control | 15.44723959 | 0.147619088 | 15.66769722 | 0.257388713 | 0.858293138 | -0.22045763 | -0.194630993 | 0.127619672 | 0.894082377 | Unpaired t-test | 0.42774087 | permutation FDR (250 perm) | 1 | 0.150735929 | FDR correction BH |
| mts~X2JDI1 | VMAT~Control | VMAT | Control | 16.75487585 | 0.096036518 | 16.9765709 | 0.247389562 | 0.857557286 | -0.221695047 | -0.198175586 | 0.091306306 | 1.039499226 | Unpaired t-test | 0.421191304 | permutation FDR (250 perm) | 1 | 0.109997464 | FDR correction BH |
| Dmel\CG8353~Q9VLR2 | VMAT~Control | VMAT | Control | 13.1658815 | 0.519077604 | 13.38888245 | 0.400042024 | 0.856781393 | -0.223000946 | -0.172457386 | 0.464285834 | 0.333214567 | Unpaired t-test | 0.469767159 | permutation FDR (250 perm) | 1 | 0.502034384 | FDR correction BH |
| norpA~X2JCI4 | VMAT~Control | VMAT | Control | 15.86808156 | 0.154054559 | 16.09187606 | 0.185186912 | 0.856310251 | -0.223794499 | -0.202030125 | 0.064478892 | 1.190582435 | Unpaired t-test | 0.414346087 | permutation FDR (250 perm) | 1 | 0.079301626 | FDR correction BH |
| Dmel\CG3226~Q9W3Y3 | VMAT~Control | VMAT | Control | 18.59708947 | 0.220672639 | 18.82132712 | 0.364106913 | 0.856047265 | -0.224237641 | -0.188370888 | 0.266188437 | 0.574810814 | Unpaired t-test | 0.439062609 | permutation FDR (250 perm) | 1 | 0.299668857 | FDR correction BH |
| Gat~Q8IMC1 | VMAT~Control | VMAT | Control | 13.7794624 | 1.011597702 | 14.00419674 | 0.869379644 | 0.855752588 | -0.224734344 | -0.140765568 | 0.714233737 | 0.146159639 | Unpaired t-test | 0.536098744 | permutation FDR (250 perm) | 1 | 0.741806016 | FDR correction BH |
| Usp16-45~Q9W4C3 | VMAT~Control | VMAT | Control | 13.35681819 | 0.517089831 | 13.58453931 | 0.296821391 | 0.853982776 | -0.227721122 | -0.179783597 | 0.413074226 | 0.383971903 | Unpaired t-test | 0.455450674 | permutation FDR (250 perm) | 1 | 0.450382921 | FDR correction BH |
| Vps24~Q9VN02 | VMAT~Control | VMAT | Control | 16.43614433 | 0.081546256 | 16.66439764 | 0.354059308 | 0.853667817 | -0.228253303 | -0.196349366 | 0.190385272 | 0.720366652 | Unpaired t-test | 0.424495652 | permutation FDR (250 perm) | 1 | 0.218298553 | FDR correction BH |
| eIF3c~A0A0B4LFL2 | VMAT~Control | VMAT | Control | 17.43033713 | 0.150345696 | 17.66240182 | 0.09501671 | 0.851415529 | -0.232064691 | -0.214966538 | 0.015362599 | 1.813535319 | Unpaired t-test | 0.392856397 | permutation FDR (250 perm) | 1 | 0.020606015 | FDR correction BH |
| hlk~Q9V9A9 | VMAT~Control | VMAT | Control | 15.23496505 | 0.197789958 | 15.46952661 | 0.23089761 | 0.849943264 | -0.234561554 | -0.206486313 | 0.115214069 | 0.938494487 | Unpaired t-test | 0.406594783 | permutation FDR (250 perm) | 1 | 0.136976726 | FDR correction BH |
| His1:CG33807~Q4ABD8 | VMAT~Control | VMAT | Control | 18.89780753 | 0.550774232 | 19.13383475 | 0.715939077 | 0.849080229 | -0.236027215 | -0.168115265 | 0.571974665 | 0.242623207 | Unpaired t-test | 0.478167607 | permutation FDR (250 perm) | 1 | 0.606772583 | FDR correction BH |
| Plc21C~P25455-2 | VMAT~Control | VMAT | Control | 16.92744268 | 0.049641065 | 17.16362137 | 0.160387989 | 0.848991086 | -0.236178689 | -0.219683802 | 0.010412362 | 1.982450751 | Unpaired t-test | 0.385302569 | permutation FDR (250 perm) | 1 | 0.014358934 | FDR correction BH |
| Atox1~Q95RR1 | VMAT~Control | VMAT | Control | 14.52091225 | 0.103745671 | 14.75867663 | 0.468558913 | 0.848058461 | -0.237764374 | -0.195751923 | 0.293867674 | 0.531848184 | Unpaired t-test | 0.425563478 | permutation FDR (250 perm) | 1 | 0.328785411 | FDR correction BH |
| CysRS~Q7KN90 | VMAT~Control | VMAT | Control | 17.36333418 | 0.075424837 | 17.60263938 | 0.176464949 | 0.847153202 | -0.239305201 | -0.220390416 | 0.019174787 | 1.717269462 | Unpaired t-test | 0.384114933 | permutation FDR (250 perm) | 1 | 0.025415229 | FDR correction BH |
| mge~Q9VZL1 | VMAT~Control | VMAT | Control | 15.33996849 | 0.147419864 | 15.57929077 | 0.304907301 | 0.847143173 | -0.239322279 | -0.20784241 | 0.145164878 | 0.838138448 | Unpaired t-test | 0.404354783 | permutation FDR (250 perm) | 1 | 0.170178348 | FDR correction BH |
| sra~A0A0B4LHI3 | VMAT~Control | VMAT | Control | 16.78138435 | 0.155787537 | 17.02186768 | 0.093721748 | 0.846461687 | -0.240483326 | -0.222400757 | 0.014343279 | 1.843351542 | Unpaired t-test | 0.381178214 | permutation FDR (250 perm) | 1 | 0.019337961 | FDR correction BH |
| Dmel\CG7255~Q2PDY3 | VMAT~Control | VMAT | Control | 15.43037202 | 0.047135787 | 15.67101641 | 0.104797593 | 0.846367192 | -0.240644391 | -0.228882283 | 0.000863872 | 3.063550673 | Unpaired t-test | 0.370966449 | permutation FDR (250 perm) | 1 | 0.0013938 | FDR correction BH |
| Dmel\CG1440~Q9W3F6 | VMAT~Control | VMAT | Control | 15.63817634 | 0.028181637 | 15.88096561 | 0.140882443 | 0.845109819 | -0.242789268 | -0.228131214 | 0.00360988 | 2.442507294 | Unpaired t-test | 0.372127233 | permutation FDR (250 perm) | 1 | 0.005297791 | FDR correction BH |
| Fas3~X2JAT7 | VMAT~Control | VMAT | Control | 21.15741594 | 0.115479845 | 21.40100508 | 0.140885422 | 0.844641397 | -0.243589137 | -0.225239563 | 0.013572791 | 1.867330822 | Unpaired t-test | 0.37664488 | permutation FDR (250 perm) | 1 | 0.01837283 | FDR correction BH |
| Atat~Q9VSY4 | VMAT~Control | VMAT | Control | 17.4970885 | 0.163998892 | 17.74243603 | 0.148994626 | 0.84361255 | -0.245347537 | -0.223227702 | 0.032767372 | 1.484558383 | Unpaired t-test | 0.379823965 | permutation FDR (250 perm) | 1 | 0.041989328 | FDR correction BH |
| beta'COP~O62621 | VMAT~Control | VMAT | Control | 20.61437138 | 0.127534353 | 20.86050259 | 0.338255637 | 0.843154427 | -0.246131204 | -0.2118775 | 0.158870157 | 0.798957675 | Unpaired t-test | 0.397561739 | permutation FDR (250 perm) | 1 | 0.184590498 | FDR correction BH |
| Nrg~E1JJF9 | VMAT~Control | VMAT | Control | 18.25421122 | 0.080446482 | 18.50149472 | 0.05649926 | 0.842481258 | -0.247283502 | -0.236869948 | 0.000219994 | 3.657588284 | Unpaired t-test | 0.358920663 | permutation FDR (250 perm) | 1 | 0.000402071 | FDR correction BH |
| Mnn1~Q9VM47 | VMAT~Control | VMAT | Control | 13.25700519 | 0.325738959 | 13.50993118 | 0.181928655 | 0.839192689 | -0.252925985 | -0.216758604 | 0.160513276 | 0.794489042 | Unpaired t-test | 0.390157597 | permutation FDR (250 perm) | 1 | 0.186315706 | FDR correction BH |
| Hsp27~X2JC82 | VMAT~Control | VMAT | Control | 16.67949276 | 0.081621886 | 16.93763092 | 0.145647927 | 0.836166323 | -0.258138156 | -0.24020303 | 0.006151389 | 2.211026804 | Unpaired t-test | 0.353815962 | permutation FDR (250 perm) | 1 | 0.008775982 | FDR correction BH |
| Naus~Q8SX68 | VMAT~Control | VMAT | Control | 17.74051965 | 0.21442456 | 18.00110175 | 0.219656722 | 0.834751046 | -0.260582098 | -0.229127775 | 0.086879942 | 1.061080479 | Unpaired t-test | 0.370612636 | permutation FDR (250 perm) | 1 | 0.104933496 | FDR correction BH |
| Atg18a~M9PEY7 | VMAT~Control | VMAT | Control | 19.95817546 | 0.102180429 | 20.21877339 | 0.241344369 | 0.834741887 | -0.260597929 | -0.233258314 | 0.050404106 | 1.297534084 | Unpaired t-test | 0.364162092 | permutation FDR (250 perm) | 1 | 0.06284495 | FDR correction BH |
| Pkc53E~A0A0B4KFT3 | VMAT~Control | VMAT | Control | 14.07397475 | 0.86362757 | 14.33604401 | 0.958907281 | 0.833891007 | -0.262069265 | -0.166169181 | 0.659456241 | 0.180814017 | Unpaired t-test | 0.482348241 | permutation FDR (250 perm) | 1 | 0.689937774 | FDR correction BH |
| Nurf-38~O77460 | VMAT~Control | VMAT | Control | 17.79732307 | 0.112840351 | 18.06006642 | 0.120112163 | 0.833501472 | -0.262743348 | -0.244707902 | 0.005138782 | 2.289139797 | Unpaired t-test | 0.347689219 | permutation FDR (250 perm) | 1 | 0.007416734 | FDR correction BH |
| milt~D0Z747 | VMAT~Control | VMAT | Control | 15.93414255 | 0.130928264 | 16.19767577 | 0.148109037 | 0.833045257 | -0.26353322 | -0.242127581 | 0.013785806 | 1.860567836 | Unpaired t-test | 0.351195462 | permutation FDR (250 perm) | 1 | 0.01862904 | FDR correction BH |
| faf~A0A0B4K6W2 | VMAT~Control | VMAT | Control | 16.71305966 | 0.162071883 | 16.97689058 | 0.077441364 | 0.832873377 | -0.263830918 | -0.24421329 | 0.008228317 | 2.084688972 | Unpaired t-test | 0.348424269 | permutation FDR (250 perm) | 1 | 0.011591537 | FDR correction BH |
| Hsp110~Q9VUC1 | VMAT~Control | VMAT | Control | 17.9204621 | 0.123634488 | 18.18705526 | 0.078549348 | 0.831280251 | -0.266593158 | -0.250203228 | 0.002251164 | 2.647592837 | Unpaired t-test | 0.340120524 | permutation FDR (250 perm) | 1 | 0.003396949 | FDR correction BH |
| RpL36~X2JC35 | VMAT~Control | VMAT | Control | 15.81812464 | 0.104532185 | 16.08770233 | 0.466897958 | 0.829562339 | -0.269577696 | -0.222062481 | 0.236323668 | 0.626492781 | Unpaired t-test | 0.38160453 | permutation FDR (250 perm) | 1 | 0.268617052 | FDR correction BH |
| Cf2~M9PB35 | VMAT~Control | VMAT | Control | 13.54293025 | 0.311351668 | 13.81302322 | 0.539134382 | 0.829266101 | -0.270092976 | -0.211269901 | 0.35488918 | 0.449907241 | Unpaired t-test | 0.398485217 | permutation FDR (250 perm) | 1 | 0.392026809 | FDR correction BH |
| qkr54B~Q7KNS5 | VMAT~Control | VMAT | Control | 13.68734071 | 0.37005873 | 13.95930884 | 0.74045903 | 0.82818896 | -0.271968123 | -0.198488574 | 0.479419234 | 0.319284546 | Unpaired t-test | 0.420648696 | permutation FDR (250 perm) | 1 | 0.516567246 | FDR correction BH |
| hppy~A0A0B4LFQ3 | VMAT~Control | VMAT | Control | 14.52575476 | 0.045423389 | 14.80134258 | 0.208542475 | 0.826113656 | -0.275587815 | -0.251575046 | 0.016183064 | 1.790939254 | Unpaired t-test | 0.338214847 | permutation FDR (250 perm) | 1 | 0.021583531 | FDR correction BH |
| AlaRS~Q9VLM8 | VMAT~Control | VMAT | Control | 16.05514039 | 0.199694542 | 16.33831567 | 0.120971693 | 0.821780341 | -0.283175277 | -0.256403051 | 0.02185742 | 1.660401095 | Unpaired t-test | 0.33166623 | permutation FDR (250 perm) | 1 | 0.028664272 | FDR correction BH |
| Dmel\CG34133~Q0KHZ4 | VMAT~Control | VMAT | Control | 16.4696278 | 0.122668085 | 16.7529906 | 0.131796863 | 0.821673532 | -0.283362799 | -0.262246539 | 0.00554646 | 2.255984149 | Unpaired t-test | 0.323917868 | permutation FDR (250 perm) | 1 | 0.007961199 | FDR correction BH |
| lic~O62602 | VMAT~Control | VMAT | Control | 16.1475557 | 0.090898304 | 16.43138004 | 0.060427161 | 0.821410704 | -0.283824348 | -0.270614615 | 0.000169598 | 3.770578648 | Unpaired t-test | 0.312718218 | permutation FDR (250 perm) | 1 | 0.000316853 | FDR correction BH |
| MAPk-Ak2~A4V402 | VMAT~Control | VMAT | Control | 18.03514615 | 0.089683899 | 18.31920103 | 0.260695219 | 0.821279461 | -0.284054877 | -0.252877036 | 0.043960761 | 1.356934801 | Unpaired t-test | 0.336403493 | permutation FDR (250 perm) | 1 | 0.055220721 | FDR correction BH |
| Usp7~X2JEX7 | VMAT~Control | VMAT | Control | 18.42469663 | 0.075214489 | 18.71024062 | 0.191086013 | 0.820432194 | -0.28554399 | -0.26152597 | 0.011075488 | 1.955637126 | Unpaired t-test | 0.324889471 | permutation FDR (250 perm) | 1 | 0.015193298 | FDR correction BH |
| RpS21~M9PEA6 | VMAT~Control | VMAT | Control | 18.57374013 | 0.161117727 | 18.85967291 | 0.37519923 | 0.820211123 | -0.285932786 | -0.241780985 | 0.148462514 | 0.828383188 | Unpaired t-test | 0.351689354 | permutation FDR (250 perm) | 1 | 0.1735257 | FDR correction BH |
| cib~O97428 | VMAT~Control | VMAT | Control | 21.70953972 | 0.057314708 | 22.00163921 | 0.154191636 | 0.816712671 | -0.292099485 | -0.27208331 | 0.002640957 | 2.578238684 | Unpaired t-test | 0.311001748 | permutation FDR (250 perm) | 1 | 0.003944678 | FDR correction BH |
| Uba5~Q9VYY3 | VMAT~Control | VMAT | Control | 16.18366328 | 0.102873198 | 16.47861469 | 0.136560913 | 0.815099782 | -0.294951415 | -0.274000905 | 0.003173073 | 2.498519991 | Unpaired t-test | 0.308550699 | permutation FDR (250 perm) | 1 | 0.004685955 | FDR correction BH |
| CG10860~M9MRV3 | VMAT~Control | VMAT | Control | 13.52707291 | 0.302979645 | 13.82206045 | 0.723823639 | 0.815079369 | -0.294987546 | -0.218360803 | 0.420207933 | 0.376535752 | Unpaired t-test | 0.387392251 | permutation FDR (250 perm) | 1 | 0.457948831 | FDR correction BH |

| | | | | | | | | | | | | | | | | | | |
|---|---|---|---|---|---|---|---|---|---|---|---|---|---|---|---|---|---|---|
| Dmel\CG6432~Q9VC92 | VMAT~Control | VMAT | Control | 13.65677777 | 0.553536223 | 13.95208159 | 0.514732894 | 0.814900699 | -0.295303827 | -0.220698945 | 0.402838416 | 0.394869121 | Unpaired t-test | 0.383569874 | permutation FDR (250 perm) | 1 | 0.439629871 | FDR correction BH |
| Ak1~Q9VTV3 | VMAT~Control | VMAT | Control | 17.35793617 | 0.222632949 | 17.65558885 | 0.252126694 | 0.813575041 | -0.297652674 | -0.258733557 | 0.076025563 | 1.119040353 | Unpaired t-test | 0.328515509 | permutation FDR (250 perm) | 1 | 0.09239245 | FDR correction BH |
| Dsor1~X2JAZ3 | VMAT~Control | VMAT | Control | 16.76047883 | 0.080078727 | 17.06155943 | 0.093390681 | 0.811644231 | -0.301080607 | -0.285379829 | 0.000272102 | 3.565268012 | Unpaired t-test | 0.295293912 | permutation FDR (250 perm) | 1 | 0.000490826 | FDR correction BH |
| pst~Q8IQ82 | VMAT~Control | VMAT | Control | 15.44049106 | 0.312891712 | 15.74338538 | 0.50695129 | 0.810624496 | -0.302894323 | -0.239173478 | 0.282088821 | 0.549614124 | Unpaired t-test | 0.355368513 | permutation FDR (250 perm) | 1 | 0.316358783 | FDR correction BH |
| ttm50~A0A0S0X8K7 | VMAT~Control | VMAT | Control | 14.07378474 | 1.440773429 | 14.37877681 | 0.826129702 | 0.809446668 | -0.304992066 | -0.175007146 | 0.690003669 | 0.1611486 | Unpaired t-test | 0.464813206 | permutation FDR (250 perm) | 1 | 0.718385067 | FDR correction BH |
| Plap~Q9VPY2 | VMAT~Control | VMAT | Control | 15.01356115 | 0.102498921 | 15.32056365 | 0.114139205 | 0.808319471 | -0.307002495 | -0.287292556 | 0.001188412 | 2.925032966 | Unpaired t-test | 0.293082786 | permutation FDR (250 perm) | 1 | 0.001879838 | FDR correction BH |
| anon-WO0118547.200~Q9VJ06 | VMAT~Control | VMAT | Control | 19.62120665 | 0.097575686 | 19.92822985 | 0.120752188 | 0.808307874 | -0.307023194 | -0.287090701 | 0.001289886 | 2.889448627 | Unpaired t-test | 0.293335085 | permutation FDR (250 perm) | 1 | 0.002021566 | FDR correction BH |
| fabp~Q8INK2 | VMAT~Control | VMAT | Control | 16.16035629 | 0.296196986 | 16.46761386 | 0.557639131 | 0.808176569 | -0.30725757 | -0.239599357 | 0.302080776 | 0.519876911 | Unpaired t-test | 0.354740515 | permutation FDR (250 perm) | 1 | 0.337493188 | FDR correction BH |
| SerRS~Q9VQL1 | VMAT~Control | VMAT | Control | 17.21216425 | 0.097039096 | 17.52100266 | 0.121799795 | 0.807291491 | -0.308838409 | -0.288729957 | 0.001265252 | 2.897822965 | Unpaired t-test | 0.291290407 | permutation FDR (250 perm) | 1 | 0.001988623 | FDR correction BH |
| Ufd1~Q9VTF9 | VMAT~Control | VMAT | Control | 15.6682995 | 0.077109615 | 15.97720188 | 0.111435047 | 0.807255695 | -0.30890238 | -0.291251636 | 0.000465968 | 3.331644014 | Unpaired t-test | 0.288420499 | permutation FDR (250 perm) | 1 | 0.000789696 | FDR correction BH |
| anon-WO02059370.23~Q9VR30 | VMAT~Control | VMAT | Control | 15.31593697 | 0.225218922 | 15.62569044 | 0.262079333 | 0.806779609 | -0.309753474 | -0.268292267 | 0.072852924 | 1.137553012 | Unpaired t-test | 0.315781564 | permutation FDR (250 perm) | 1 | 0.088811902 | FDR correction BH |
| Septin2~P54359 | VMAT~Control | VMAT | Control | 18.24579391 | 0.139187103 | 18.55885805 | 0.127899601 | 0.804930352 | -0.313064138 | -0.288661923 | 0.00408593 | 2.38870908 | Unpaired t-test | 0.291381516 | permutation FDR (250 perm) | 1 | 0.005951905 | FDR correction BH |
| LRR~E1JGZ8 | VMAT~Control | VMAT | Control | 16.92547781 | 0.630118491 | 17.23856125 | 0.592955013 | 0.804919583 | -0.313083439 | -0.225735478 | 0.437279745 | 0.359240638 | Unpaired t-test | 0.375837908 | permutation FDR (250 perm) | 1 | 0.475453358 | FDR correction BH |
| CG17333~Q9VZ64 | VMAT~Control | VMAT | Control | 14.87720124 | 0.281706102 | 15.1904255 | 0.279701067 | 0.804841015 | -0.313224267 | -0.266000264 | 0.108145868 | 0.965990071 | Unpaired t-test | 0.318792486 | permutation FDR (250 perm) | 1 | 0.129226077 | FDR correction BH |
| VhaSFD~Q9V3J1-2 | VMAT~Control | VMAT | Control | 19.84664769 | 0.162827181 | 20.16041674 | 0.252374329 | 0.804537152 | -0.313769051 | -0.276614922 | 0.0416202 | 1.380695836 | Unpaired t-test | 0.305361888 | permutation FDR (250 perm) | 1 | 0.052476674 | FDR correction BH |
| hpo~A0A0B4KFW0 | VMAT~Control | VMAT | Control | 14.46399812 | 0.137231153 | 14.77851158 | 0.195847693 | 0.804122131 | -0.314513459 | -0.284126839 | 0.014764671 | 1.830776233 | Unpaired t-test | 0.296904074 | permutation FDR (250 perm) | 1 | 0.019860591 | FDR correction BH |
| ppan~Q9VDE5 | VMAT~Control | VMAT | Control | 15.72399033 | 0.466545859 | 16.04076385 | 0.502043099 | 0.802863411 | -0.316773528 | -0.242459582 | 0.325708956 | 0.487170299 | Unpaired t-test | 0.350685864 | permutation FDR (250 perm) | 1 | 0.362172359 | FDR correction BH |
| aPKC~A0A0B4JD78 | VMAT~Control | VMAT | Control | 16.37586544 | 0.082756016 | 16.6947953 | 0.246654973 | 0.801664304 | -0.318929859 | -0.285689709 | 0.020793022 | 1.682082377 | Unpaired t-test | 0.294938239 | permutation FDR (250 perm) | 1 | 0.027375154 | FDR correction BH |
| tmod~A8JRH3 | VMAT~Control | VMAT | Control | 18.36695214 | 0.202258679 | 18.68766137 | 0.187202414 | 0.800676167 | -0.320709231 | -0.285518909 | 0.026396452 | 1.578454439 | Unpaired t-test | 0.295139728 | permutation FDR (250 perm) | 1 | 0.034216547 | FDR correction BH |
| raptor~Q9W437 | VMAT~Control | VMAT | Control | 15.21737076 | 0.265826856 | 15.54024845 | 0.722831954 | 0.799473604 | -0.322877693 | -0.24016007 | 0.37062328 | 0.431067304 | Unpaired t-test | 0.353864806 | permutation FDR (250 perm) | 1 | 0.407876205 | FDR correction BH |
| Dmel\CG6664~Q9VVC8 | VMAT~Control | VMAT | Control | 15.68594463 | 0.386853877 | 16.00995192 | 0.255083353 | 0.798847881 | -0.324007289 | -0.268388807 | 0.148998585 | 0.826817857 | Unpaired t-test | 0.31566623 | permutation FDR (250 perm) | 1 | 0.17406584 | FDR correction BH |
| Rpn2~A0A0B4KHE9 | VMAT~Control | VMAT | Control | 16.16556812 | 0.138657396 | 16.49088079 | 0.169830699 | 0.798125395 | -0.325312665 | -0.296264167 | 0.00777565 | 2.109263271 | Unpaired t-test | 0.283016221 | permutation FDR (250 perm) | 1 | 0.010980133 | FDR correction BH |
| Dmel\CG7888~Q7YTZ0 | VMAT~Control | VMAT | Control | 14.97422448 | 0.115516079 | 15.30359738 | 0.317935408 | 0.79588236 | -0.329372894 | -0.286092993 | 0.054497983 | 1.263619568 | Unpaired t-test | 0.294486202 | permutation FDR (250 perm) | 1 | 0.06780563 | FDR correction BH |
| Nmt~O61613 | VMAT~Control | VMAT | Control | 18.48006397 | 0.180352242 | 18.81013215 | 0.598472488 | 0.795498887 | -0.330068182 | -0.257959683 | 0.265012569 | 0.576733528 | Unpaired t-test | 0.329567497 | permutation FDR (250 perm) | 1 | 0.298773749 | FDR correction BH |
| mEFG1~Q9VM33 | VMAT~Control | VMAT | Control | 14.50081669 | 1.118343636 | 14.83305345 | 0.915014484 | 0.794304035 | -0.332236763 | -0.201819051 | 0.61833019 | 0.208779548 | Unpaired t-test | 0.414728696 | permutation FDR (250 perm) | 1 | 0.651918256 | FDR correction BH |
| Dys~Q9VDW6-4 | VMAT~Control | VMAT | Control | 17.8730017 | 0.080915868 | 18.20798162 | 0.092073961 | 0.792795174 | -0.334979915 | -0.317571333 | 0.00011406 | 3.942866909 | Unpaired t-test | 0.259903424 | permutation FDR (250 perm) | 1 | 0.000220441 | FDR correction BH |
| Dmel\CG7766~X2JJ32 | VMAT~Control | VMAT | Control | 16.64126874 | 0.034845266 | 16.97630991 | 0.063466367 | 0.79276151 | -0.335041177 | -0.32453295 | 1.16E-06 | 5.935007099 | Unpaired t-test | 0.253185055 | permutation FDR (250 perm) | 1 | 3.85E-06 | FDR correction BH |
| cg4266~A0A0B4LH25 | VMAT~Control | VMAT | Control | 13.36782942 | 0.445290923 | 13.70452731 | 0.371403174 | 0.791851666 | -0.336697894 | -0.26736566 | 0.223291554 | 0.651127703 | Unpaired t-test | 0.317025775 | permutation FDR (250 perm) | 1 | 0.254295268 | FDR correction BH |
| Zyx~Q8I0G8 | VMAT~Control | VMAT | Control | 17.19331995 | 0.182657039 | 17.53489677 | 0.366099405 | 0.789178294 | -0.341576819 | -0.288744821 | 0.091482695 | 1.038661051 | Unpaired t-test | 0.291271134 | permutation FDR (250 perm) | 1 | 0.110153587 | FDR correction BH |
| EP3527~E1JIZ3 | VMAT~Control | VMAT | Control | 13.98682492 | 1.0397021 | 14.32862028 | 0.754304607 | 0.789058758 | -0.341795358 | -0.217088929 | 0.565065269 | 0.247901385 | Unpaired t-test | 0.389584676 | permutation FDR (250 perm) | 1 | 0.600796587 | FDR correction BH |
| speck~Q94521-2 | VMAT~Control | VMAT | Control | 16.41299249 | 0.163939711 | 16.75535214 | 0.170999374 | 0.788750189 | -0.342359649 | -0.309564239 | 0.008996223 | 2.045939769 | Unpaired t-test | 0.268412462 | permutation FDR (250 perm) | 1 | 0.012538253 | FDR correction BH |
| smash~E2QCZ8 | VMAT~Control | VMAT | Control | 13.23080832 | 0.304016498 | 13.57647658 | 0.493352682 | 0.786943378 | -0.34566826 | -0.27452262 | 0.211851837 | 0.673967765 | Unpaired t-test | 0.307912587 | permutation FDR (250 perm) | 1 | 0.241969541 | FDR correction BH |
| Brd7-9~Q9VLX2 | VMAT~Control | VMAT | Control | 15.2734276 | 0.419229818 | 15.61938407 | 0.685808249 | 0.786786184 | -0.345956471 | -0.254479284 | 0.358536891 | 0.445466152 | Unpaired t-test | 0.334227074 | permutation FDR (250 perm) | 1 | 0.395499457 | FDR correction BH |
| Cwc25~Q9VQF5 | VMAT~Control | VMAT | Control | 15.36740697 | 0.238534398 | 15.71530499 | 0.335979195 | 0.785728053 | -0.347898025 | -0.293765328 | 0.088358328 | 1.05375251 | Unpaired t-test | 0.285706527 | permutation FDR (250 perm) | 1 | 0.10660969 | FDR correction BH |
| R5_2R:11947059..11948303~Q7K4L9 | VMAT~Control | VMAT | Control | 15.36510177 | 0.234102028 | 15.71325378 | 0.187482936 | 0.785589741 | -0.348152005 | -0.306977295 | 0.026690386 | 1.573645145 | Unpaired t-test | 0.27102807 | permutation FDR (250 perm) | 1 | 0.034559499 | FDR correction BH |
| Gs2~X2JJG8 | VMAT~Control | VMAT | Control | 20.01875373 | 0.06487614 | 20.36714089 | 0.072445836 | 0.7854617 | -0.348387165 | -0.333866974 | 1.16E-05 | 4.934063453 | Unpaired t-test | 0.244094945 | permutation FDR (250 perm) | 1 | 2.87E-05 | FDR correction BH |
| Hyccin~Q7K1C5 | VMAT~Control | VMAT | Control | 15.38371774 | 0.126763514 | 15.73496939 | 0.176735254 | 0.783903703 | -0.351251655 | -0.320115029 | 0.004758532 | 2.322526973 | Unpaired t-test | 0.257592267 | permutation FDR (250 perm) | 1 | 0.006876357 | FDR correction BH |
| SH3PX1~Q9NCC3 | VMAT~Control | VMAT | Control | 16.54661375 | 0.186156684 | 16.9002802 | 0.566499142 | 0.782592699 | -0.353666443 | -0.279208694 | 0.214261474 | 0.669055911 | Unpaired t-test | 0.302243007 | permutation FDR (250 perm) | 1 | 0.24460306 | FDR correction BH |
| shrb~Q8T0Q4 | VMAT~Control | VMAT | Control | 18.69328637 | 0.06556126 | 19.04830049 | 0.243088264 | 0.781861991 | -0.355014121 | -0.319086056 | 0.010280947 | 1.987966893 | Unpaired t-test | 0.258543058 | permutation FDR (250 perm) | 1 | 0.014202669 | FDR correction BH |
| arm~M9PGG4 | VMAT~Control | VMAT | Control | 16.12352459 | 0.18627183 | 16.4830762 | 0.304183724 | 0.779406782 | -0.35955161 | -0.310087974 | 0.047850968 | 1.320109271 | Unpaired t-test | 0.267831505 | permutation FDR (250 perm) | 1 | 0.059883668 | FDR correction BH |
| clu~A1ZAB5 | VMAT~Control | VMAT | Control | 16.0342619 | 0.26650935 | 16.39416853 | 0.193969876 | 0.779215006 | -0.359906633 | -0.313668169 | 0.034759602 | 1.458925208 | Unpaired t-test | 0.263894691 | permutation FDR (250 perm) | 1 | 0.044373158 | FDR correction BH |
| BicD~M9NDR2 | VMAT~Control | VMAT | Control | 16.84803723 | 0.085861627 | 17.20804745 | 0.158118372 | 0.779159061 | -0.360010219 | -0.333199121 | 0.001189872 | 2.924499822 | Unpaired t-test | 0.244664615 | permutation FDR (250 perm) | 1 | 0.001879838 | FDR correction BH |
| Reps~M9NDL7 | VMAT~Control | VMAT | Control | 17.16587961 | 0.152735068 | 17.52626037 | 0.098467463 | 0.778958965 | -0.360380764 | -0.333294048 | 0.001265488 | 2.897742106 | Unpaired t-test | 0.244574945 | permutation FDR (250 perm) | 1 | 0.001988623 | FDR correction BH |
| RSG7~Q9VXA3 | VMAT~Control | VMAT | Control | 15.63163073 | 0.186366005 | 15.9955807 | 0.147841011 | 0.777034217 | -0.363949966 | -0.328954061 | 0.006535878 | 2.184696033 | Unpaired t-test | 0.248734945 | permutation FDR (250 perm) | 1 | 0.00930197 | FDR correction BH |
| Arr2~P19107 | VMAT~Control | VMAT | Control | 16.92415644 | 0.201796725 | 17.29562929 | 0.117996882 | 0.772992946 | -0.371472846 | -0.336313907 | 0.005239065 | 2.280746176 | Unpaired t-test | 0.241749451 | permutation FDR (250 perm) | 1 | 0.007552211 | FDR correction BH |
| SNF4Agamma~Q9VDD2 | VMAT~Control | VMAT | Control | 15.52113723 | 0.167211107 | 15.89329895 | 0.298937557 | 0.772623932 | -0.372161729 | -0.322726 | 0.035477865 | 1.450042523 | Unpaired t-test | 0.254996044 | permutation FDR (250 perm) | 1 | 0.04521651 | FDR correction BH |
| CG14318~A0A0B4LIC0 | VMAT~Control | VMAT | Control | 15.21671615 | 0.327856529 | 15.58972192 | 0.239718597 | 0.772172045 | -0.37300577 | -0.31566941 | 0.067101545 | 1.173267483 | Unpaired t-test | 0.261947322 | permutation FDR (250 perm) | 1 | 0.082312161 | FDR correction BH |
| HIP-R~E2QD63 | VMAT~Control | VMAT | Control | 18.44980785 | 0.15430737 | 18.82392747 | 0.207244326 | 0.771576109 | -0.374119621 | -0.335367354 | 0.008904312 | 2.050399607 | Unpaired t-test | 0.242607473 | permutation FDR (250 perm) | 1 | 0.012424868 | FDR correction BH |
| eIF2beta~P41375 | VMAT~Control | VMAT | Control | 15.91058946 | 0.096136844 | 16.29016635 | 0.219215858 | 0.768662987 | -0.379576895 | -0.342872585 | 0.005305143 | 2.275302892 | Unpaired t-test | 0.235790677 | permutation FDR (250 perm) | 1 | 0.007633439 | FDR correction BH |
| Cyp313a1~Q9VFJ0 | VMAT~Control | VMAT | Control | 15.70121796 | 0.411194102 | 16.08172091 | 0.317946495 | 0.768169747 | -0.380502948 | -0.308736428 | 0.132691976 | 0.877155337 | Unpaired t-test | 0.269286529 | permutation FDR (250 perm) | 1 | 0.156413076 | FDR correction BH |
| LysRS~Q8SXM8 | VMAT~Control | VMAT | Control | 15.97192936 | 0.185571785 | 16.35690035 | 0.242629011 | 0.765794396 | -0.384970992 | -0.338702293 | 0.018219486 | 1.739463888 | Unpaired t-test | 0.239597186 | permutation FDR (250 perm) | 1 | 0.024189887 | FDR correction BH |
| Fkbp59~X2J5E8 | VMAT~Control | VMAT | Control | 17.69330102 | 0.11200913 | 18.07932858 | 0.249849042 | 0.76523377 | -0.386027554 | -0.343915 | 0.010288788 | 1.987635779 | Unpaired t-test | 0.234671944 | permutation FDR (250 perm) | 1 | 0.014205165 | FDR correction BH |
| flw~H5V895 | VMAT~Control | VMAT | Control | 15.5390795 | 0.181837655 | 15.92600073 | 0.234771404 | 0.76475989 | -0.386921236 | -0.341561066 | 0.015471497 | 1.810467652 | Unpaired t-test | 0.237004398 | permutation FDR (250 perm) | 1 | 0.02072846 | FDR correction BH |
| Pdcd4~Q9VY91 | VMAT~Control | VMAT | Control | 15.84368821 | 0.072145365 | 16.23083685 | 0.074671736 | 0.764639355 | -0.387148639 | -0.369969293 | 8.18E-06 | 5.087019931 | Unpaired t-test | 0.2122382 | permutation FDR (250 perm) | 1 | 2.11E-05 | FDR correction BH |
| DIP2~M9PDH5 | VMAT~Control | VMAT | Control | 17.6556967 | 0.097009883 | 18.04289603 | 0.201684516 | 0.764612492 | -0.387199325 | -0.351971381 | 0.003116224 | 2.506371293 | Unpaired t-test | 0.227478873 | permutation FDR (250 perm) | 1 | 0.00461358 | FDR correction BH |
| SF2~Q9V3W7 | VMAT~Control | VMAT | Control | 16.91871338 | 0.428654759 | 17.30649492 | 0.375346106 | 0.764303985 | -0.387781543 | -0.309037134 | 0.159013989 | 0.798564667 | Unpaired t-test | 0.268942519 | permutation FDR (250 perm) | 1 | 0.184666468 | FDR correction BH |
| Dscam2~M9PEU0 | VMAT~Control | VMAT | Control | 14.20479591 | 0.18040127 | 14.59556642 | 0.381494648 | 0.762722143 | -0.39077051 | -0.328731208 | 0.06521833 | 1.185630324 | Unpaired t-test | 0.248931868 | permutation FDR (250 perm) | 1 | 0.080085524 | FDR correction BH |
| Dmel\CG10550~Q9VBS7 | VMAT~Control | VMAT | Control | 13.51576452 | 0.547111681 | 13.90777254 | 0.793921143 | 0.762068182 | -0.392008014 | -0.273902751 | 0.384673269 | 0.414907992 | Unpaired t-test | 0.308662587 | permutation FDR (250 perm) | 1 | 0.421807893 | FDR correction BH |
| HBS1~Q9W074 | VMAT~Control | VMAT | Control | 16.58339489 | 0.094769049 | 16.97795257 | 0.128125462 | 0.760722572 | -0.394557682 | -0.36830823 | 0.000248896 | 3.603982578 | Unpaired t-test | 0.213790913 | permutation FDR (250 perm) | 1 | 0.000452782 | FDR correction BH |
| HisRS~Q9VWT1 | VMAT~Control | VMAT | Control | 15.43081996 | 0.101149681 | 15.82697665 | 0.191223148 | 0.759879894 | -0.396156689 | -0.361211446 | 0.002162247 | 2.665094668 | Unpaired t-test | 0.21968268 | permutation FDR (250 perm) | 1 | 0.003279594 | FDR correction BH |
| Rrp6~Q9VFF3 | VMAT~Control | VMAT | Control | 13.34835876 | 0.177438318 | 13.74568092 | 0.266621204 | 0.759266278 | -0.397322162 | -0.34754413 | 0.0196501 | 1.706635225 | Unpaired t-test | 0.231596831 | permutation FDR (250 perm) | 1 | 0.025986706 | FDR correction BH |
| Atg3~Q9VVS6 | VMAT~Control | VMAT | Control | 18.21213415 | 0.187115042 | 18.61651873 | 0.507587996 | 0.755558525 | -0.404384585 | -0.325609048 | 0.125579001 | 0.901082976 | Unpaired t-test | 0.25213011 | permutation FDR (250 perm) | 1 | 0.14847462 | FDR correction BH |
| Oga~Q9VDC9 | VMAT~Control | VMAT | Control | 15.34043781 | 0.173183525 | 15.74547646 | 0.254537272 | 0.755216064 | -0.405038643 | -0.356020944 | 0.014739025 | 1.831531232 | Unpaired t-test | 0.224103919 | permutation FDR (250 perm) | 1 | 0.01983743 | FDR correction BH |
| Dmel\CG6686~Q9VKD8 | VMAT~Control | VMAT | Control | 16.65379462 | 0.368991995 | 17.06000274 | 0.498727677 | 0.75460412 | -0.406208118 | -0.317984304 | 0.173883606 | 0.759741363 | Unpaired t-test | 0.259452151 | permutation FDR (250 perm) | 1 | 0.200746448 | FDR correction BH |
| p8~Q9VKC5 | VMAT~Control | VMAT | Control | 14.75226263 | 1.349298439 | 15.15982615 | 1.269275545 | 0.753895508 | -0.407563519 | -0.222900812 | 0.633369217 | 0.198343048 | Unpaired t-test | 0.380369499 | permutation FDR (250 perm) | 1 | 0.665900463 | FDR correction BH |
| Shark~Q24145 | VMAT~Control | VMAT | Control | 13.27699657 | 0.33588349 | 13.68494319 | 0.482757851 | 0.753695343 | -0.407946616 | -0.322995403 | 0.151931865 | 0.818351132 | Unpaired t-test | 0.254693626 | permutation FDR (250 perm) | 1 | 0.177053272 | FDR correction BH |
| cpb~M9PBN7 | VMAT~Control | VMAT | Control | 15.19725706 | 0.302503013 | 15.60576722 | 0.486941416 | 0.75340099 | -0.408510165 | -0.325151956 | 0.142141577 | 0.847278871 | Unpaired t-test | 0.252603077 | permutation FDR (250 perm) | 1 | 0.166883427 | FDR correction BH |
| Gint3~Q7K0S5 | VMAT~Control | VMAT | Control | 16.74507244 | 0.127700383 | 17.15466855 | 0.049987065 | 0.752834108 | -0.409596103 | -0.385927633 | 5.51E-05 | 4.258953906 | Unpaired t-test | 0.199639735 | permutation FDR (250 perm) | 1 | 0.000115163 | FDR correction BH |
| mop~Q9VUH6 | VMAT~Control | VMAT | Control | 16.90161743 | 0.264314107 | 17.31213089 | 0.221218806 | 0.752355561 | -0.410513458 | -0.355686946 | 0.023768137 | 1.62400485 | Unpaired t-test | 0.224316901 | permutation FDR (250 perm) | 1 | 0.030945905 | FDR correction BH |
| BcDNA.LD23634~Q9V3Y5 | VMAT~Control | VMAT | Control | 15.63744356 | 0.527393483 | 16.04799382 | 0.484015049 | 0.752336371 | -0.410550257 | -0.310992414 | 0.228614229 | 0.640896742 | Unpaired t-test | 0.266767881 | permutation FDR (250 perm) | 1 | 0.260231091 | FDR correction BH |
| UGP~E1JI91 | VMAT~Control | VMAT | Control | 20.41602021 | 0.074884137 | 20.82821777 | 0.086198359 | 0.751477825 | -0.412197562 | -0.392171611 | 1.09E-05 | 4.963225192 | Unpaired t-test | 0.194597792 | permutation FDR (250 perm) | 1 | 2.71E-05 | FDR correction BH |
| parvin~Q9VWD0 | VMAT~Control | VMAT | Control | 16.99698201 | 0.181106446 | 17.41215676 | 0.234765886 | 0.74992865 | -0.415174755 | -0.36656767 | 0.010671998 | 1.971754274 | Unpaired t-test | 0.215283635 | permutation FDR (250 perm) | 1 | 0.014673997 | FDR correction BH |
| Srp54k~Q9V3D9 | VMAT~Control | VMAT | Control | 17.84721997 | 0.210584512 | 18.26434713 | 0.108038201 | 0.748914455 | -0.417127159 | -0.377201461 | 0.002771079 | 2.557351038 | Unpaired t-test | 0.206617219 | permutation FDR (250 perm) | 1 | 0.004120733 | FDR correction BH |
| Dif~E1JHK2 | VMAT~Control | VMAT | Control | 15.28542633 | 0.258391463 | 15.70284031 | 0.368076073 | 0.748765577 | -0.417413983 | -0.347520652 | 0.064689981 | 1.189162976 | Unpaired t-test | 0.231616197 | permutation FDR (250 perm) | 1 | 0.079519695 | FDR correction BH |
| Dmel\CG11334~A0A0B4KI38 | VMAT~Control | VMAT | Control | 17.05794984 | 0.120035271 | 17.47662026 | 0.177717048 | 0.748113762 | -0.418670424 | -0.382030596 | 0.001409352 | 2.850980555 | Unpaired t-test | 0.202742605 | permutation FDR (250 perm) | 1 | 0.002194217 | FDR correction BH |
| LamC~Q03427 | VMAT~Control | VMAT | Control | 16.22091146 | 0.198159241 | 16.63998763 | 0.098499556 | 0.74790339 | -0.419076171 | -0.38133753 | 0.001731043 | 2.761692155 | Unpaired t-test | 0.203242384 | permutation FDR (250 perm) | 1 | 0.002661577 | FDR correction BH |
| Dcr-2~A0A0B4LFK8 | VMAT~Control | VMAT | Control | 16.38774073 | 0.230028796 | 16.80882428 | 0.253000173 | 0.746863474 | -0.421083551 | -0.365232299 | 0.020351296 | 1.691407937 | Unpaired t-test | 0.216421703 | permutation FDR (250 perm) | 1 | 0.0268236 | FDR correction BH |
| cg11777~Q8MKJ6 | VMAT~Control | VMAT | Control | 16.39858683 | 0.109665496 | 16.82206439 | 0.045483122 | 0.745625154 | -0.423477564 | -0.402126764 | 1.21E-05 | 4.917564545 | Unpaired t-test | 0.187043747 | permutation FDR (250 perm) | 1 | 2.97E-05 | FDR correction BH |
| Sec13~Q9V3J4 | VMAT~Control | VMAT | Control | 15.31253831 | 0.252817089 | 15.73609622 | 0.307389301 | 0.745583632 | -0.423557907 | -0.359559427 | 0.038635229 | 1.413016511 | Unpaired t-test | 0.22112825 | permutation FDR (250 perm) | 1 | 0.049054654 | FDR correction BH |
| HINT1~Q86BQ4 | VMAT~Control | VMAT | Control | 13.99199018 | 0.655137979 | 14.41674431 | 1.00184161 | 0.744965677 | -0.424754137 | -0.276653272 | 0.44593781 | 0.350725703 | Unpaired t-test | 0.305305944 | permutation FDR (250 perm) | 1 | 0.483973078 | FDR correction BH |
| rudhira~Q9VYU0 | VMAT~Control | VMAT | Control | 15.94139319 | 0.137194418 | 16.36853973 | 0.286599301 | 0.743731331 | -0.427146545 | -0.374001136 | 0.013208325 | 1.879152259 | Unpaired t-test | 0.208935157 | permutation FDR (250 perm) | 1 | 0.017941371 | FDR correction BH |
| Hsc70-1~P29843 | VMAT~Control | VMAT | Control | 15.63488556 | 0.148390513 | 16.0660741 | 0.242338505 | 0.741650538 | -0.431188539 | -0.382571085 | 0.006850902 | 2.164252214 | Unpaired t-test | 0.20227638 | permutation FDR (250 perm) | 1 | 0.009732664 | FDR correction BH |
| BcDNA:GH02439~Q95T61 | VMAT~Control | VMAT | Control | 16.68720765 | 0.067569978 | 17.1191485 | 0.149155168 | 0.741263897 | -0.431940848 | -0.40246821 | 0.000151375 | 3.819944425 | Unpaired t-test | 0.186801591 | permutation FDR (250 perm) | 1 | 0.000285802 | FDR correction BH |
| MnM~Q9VZT8 | VMAT~Control | VMAT | Control | 17.51395894 | 0.252833499 | 17.9459035 | 0.253777568 | 0.74126199 | -0.431944559 | -0.37230028 | 0.022459277 | 1.648604231 | Unpaired t-test | 0.210292016 | permutation FDR (250 perm) | 1 | 0.029371743 | FDR correction BH |
| Dmel\CG6891~Q9VWQ7 | VMAT~Control | VMAT | Control | 15.53593971 | 0.091134825 | 15.96862308 | 0.288615282 | 0.740882483 | -0.43268337 | -0.381099765 | 0.009545552 | 2.020198931 | Unpaired t-test | 0.203436645 | permutation FDR (250 perm) | 1 | 0.013241149 | FDR correction BH |
| Dmel\CG9727~Q9VI23 | VMAT~Control | VMAT | Control | 13.69863266 | 0.485404122 | 14.13361144 | 0.905576301 | 0.739704631 | -0.434978786 | -0.298033437 | 0.366134102 | 0.436359819 | Unpaired t-test | 0.280971504 | permutation FDR (250 perm) | 1 | 0.403124264 | FDR correction BH |
| CG17276~Q9VDH1 | VMAT~Control | VMAT | Control | 15.75840604 | 0.097360965 | 16.1952313 | 0.478306572 | 0.738758507 | -0.436825257 | -0.358555555 | 0.073252661 | 1.135176595 | Unpaired t-test | 0.221888987 | permutation FDR (250 perm) | 1 | 0.08920681 | FDR correction BH |
| BEST:GH22856~Q7KVP4 | VMAT~Control | VMAT | Control | 19.29131013 | 0.355878337 | 19.73189604 | 0.634608017 | 0.7368353 | -0.440585917 | -0.332421103 | 0.205528723 | 0.687127476 | Unpaired t-test | 0.245415385 | permutation FDR (250 perm) | 1 | 0.235089705 | FDR correction BH |
| trus~Q9VG62 | VMAT~Control | VMAT | Control | 15.70405935 | 0.116025538 | 16.14657696 | 0.161267165 | 0.735849376 | -0.44251761 | -0.406409326 | 0.000552882 | 3.257367728 | Unpaired t-test | 0.18410084 | permutation FDR (250 perm) | 1 | 0.000921731 | FDR correction BH |
| deltaCOP~Q7K7G0 | VMAT~Control | VMAT | Control | 20.70552247 | 0.027632853 | 21.14881885 | 0.059490295 | 0.735452272 | -0.443296375 | -0.430662938 | 3.26E-08 | 7.487334117 | Unpaired t-test | 0.167182624 | permutation FDR (250 perm) | 1 | 1.92E-07 | FDR correction BH |
| His2B:CG33910~P02283 | VMAT~Control | VMAT | Control | 17.64241885 | 0.244636893 | 18.08693687 | 0.443857431 | 0.73482977 | -0.444518019 | -0.362383025 | 0.078275993 | 1.106371415 | Unpaired t-test | 0.218874283 | permutation FDR (250 perm) | 1 | 0.094931317 | FDR correction BH |
| AnxB10~P22465-2 | VMAT~Control | VMAT | Control | 14.61625756 | 0.127305642 | 15.06335573 | 0.383487511 | 0.733516755 | -0.447098175 | -0.37867089 | 0.032867424 | 1.483234327 | Unpaired t-test | 0.205416336 | permutation FDR (250 perm) | 1 | 0.042094623 | FDR correction BH |
| Alsin2~Q9VVI1 | VMAT~Control | VMAT | Control | 14.99620015 | 0.331187883 | 15.4452269 | 0.456214611 | 0.732536857 | -0.449026746 | -0.35861379 | 0.105252653 | 0.977766947 | Unpaired t-test | 0.221827313 | permutation FDR (250 perm) | 1 | 0.125896721 | FDR correction BH |
| Hsp60A~O02649 | VMAT~Control | VMAT | Control | 14.27054092 | 0.21739067 | 14.72243466 | 0.924544977 | 0.731082569 | -0.451893741 | -0.317175144 | 0.312381723 | 0.505314384 | Unpaired t-test | 0.260314311 | permutation FDR (250 perm) | 1 | 0.348010684 | FDR correction BH |
| Dmel\CG31751~Q9VJ68 | VMAT~Control | VMAT | Control | 20.19644366 | 0.045920413 | 20.65286449 | 0.096331834 | 0.728792073 | -0.456420827 | -0.435630256 | 2.39E-06 | 5.621739936 | Unpaired t-test | 0.163772949 | permutation FDR (250 perm) | 1 | 7.17E-06 | FDR correction BH |
| Ns2~Q7JXU4 | VMAT~Control | VMAT | Control | 13.59854925 | 0.314414691 | 14.05571818 | 0.493246231 | 0.728414258 | -0.457168933 | -0.362375006 | 0.111102447 | 0.954276378 | Unpaired t-test | 0.218876048 | permutation FDR (250 perm) | 1 | 0.1325571 | FDR correction BH |
| koi~A1Z6Q0 | VMAT~Control | VMAT | Control | 16.62296362 | 0.206658214 | 17.08229798 | 0.337761623 | 0.727321758 | -0.459334359 | -0.390231335 | 0.026770012 | 1.572351435 | Unpaired t-test | 0.196241943 | permutation FDR (250 perm) | 1 | 0.034643545 | FDR correction BH |
| PIP4K~Q8SXX1 | VMAT~Control | VMAT | Control | 17.30350666 | 0.090095512 | 17.76328931 | 0.106720847 | 0.727095789 | -0.459782654 | -0.432752723 | 2.42E-05 | 4.615835102 | Unpaired t-test | 0.165679078 | permutation FDR (250 perm) | 1 | 5.52E-05 | FDR correction BH |
| Dmel\CG1416~Q9V9Q4 | VMAT~Control | VMAT | Control | 17.26621215 | 0.043901656 | 17.72680164 | 0.119950736 | 0.726689269 | -0.460589494 | -0.435700697 | 1.10E-05 | 4.958924118 | Unpaired t-test | 0.163749889 | permutation FDR (250 perm) | 1 | 2.73E-05 | FDR correction BH |
| jub~M9MSC9 | VMAT~Control | VMAT | Control | 14.85831277 | 0.332027819 | 15.32263053 | 0.36599614 | 0.724813756 | -0.464317759 | -0.380277943 | 0.061970325 | 1.207816228 | Unpaired t-test | 0.204121854 | permutation FDR (250 perm) | 1 | 0.076375992 | FDR correction BH |
| ReepA~A0A0B4JCU3 | VMAT~Control | VMAT | Control | 16.25234075 | 0.173827011 | 16.71885643 | 0.335876683 | 0.723710354 | -0.466515683 | -0.39902722 | 0.020189108 | 1.694882862 | Unpaired t-test | 0.189248565 | permutation FDR (250 perm) | 1 | 0.02662474 | FDR correction BH |
| RhoGDI~Q9VW59 | VMAT~Control | VMAT | Control | 15.24881167 | 0.076806272 | 15.71553692 | 0.327770739 | 0.723605232 | -0.466725256 | -0.405652554 | 0.011249743 | 1.94885741 | Unpaired t-test | 0.184643963 | permutation FDR (250 perm) | 1 | 0.015405407 | FDR correction BH |
| gfzf~Q6NP69 | VMAT~Control | VMAT | Control | 16.01574159 | 0.245668946 | 16.48312142 | 0.709435083 | 0.723276994 | -0.467379831 | -0.349899816 | 0.194093836 | 0.711988257 | Unpaired t-test | 0.229367958 | permutation FDR (250 perm) | 1 | 0.222442497 | FDR correction BH |
| CdGAPr~E1JHM1 | VMAT~Control | VMAT | Control | 16.73220891 | 0.123320494 | 17.20477991 | 0.13287155 | 0.720679148 | -0.472570992 | -0.437132116 | 0.000166258 | 3.779218154 | Unpaired t-test | 0.162854102 | permutation FDR (250 perm) | 1 | 0.000311353 | FDR correction BH |
| Hcs~Q9VNC3 | VMAT~Control | VMAT | Control | 17.75125818 | 0.089945048 | 18.22648128 | 0.097976164 | 0.719355538 | -0.475223104 | -0.44854366 | 1.19E-05 | 4.92409762 | Unpaired t-test | 0.15542984 | permutation FDR (250 perm) | 1 | 2.93E-05 | FDR correction BH |
| eEF5~Q9GU68 | VMAT~Control | VMAT | Control | 18.07575714 | 0.128590658 | 18.55131933 | 0.169635876 | 0.719186481 | -0.475562193 | -0.434225442 | 0.000540843 | 3.266928432 | Unpaired t-test | 0.164721951 | permutation FDR (250 perm) | 1 | 0.000903581 | FDR correction BH |
| tho2~E2QCS8 | VMAT~Control | VMAT | Control | 14.12686366 | 0.18109415 | 14.60312025 | 0.748379211 | 0.718840407 | -0.476256587 | -0.354266707 | 0.196739139 | 0.706109234 | Unpaired t-test | 0.225540493 | permutation FDR (250 perm) | 1 | 0.225364444 | FDR correction BH |
| Dmel\CG8003~Q9VTB0 | VMAT~Control | VMAT | Control | 15.88410508 | 0.134642257 | 16.36108898 | 0.153415423 | 0.718478108 | -0.476983896 | -0.437084594 | 0.000386955 | 3.412339599 | Unpaired t-test | 0.162878936 | permutation FDR (250 perm) | 1 | 0.000666832 | FDR correction BH |
| puml~Q7JQU9 | VMAT~Control | VMAT | Control | 16.35881715 | 0.086612986 | 16.84079854 | 0.21331174 | 0.715993605 | -0.481981393 | -0.436989043 | 0.000865864 | 3.062550305 | Unpaired t-test | 0.162937472 | permutation FDR (250 perm) | 1 | 0.001396057 | FDR correction BH |
| cg9143~Q6NQY9 | VMAT~Control | VMAT | Control | 13.5814592 | 0.484841182 | 14.06537796 | 0.414839947 | 0.715032755 | -0.483918764 | -0.376483822 | 0.120781894 | 0.917998163 | Unpaired t-test | 0.206990737 | permutation FDR (250 perm) | 1 | 0.143234549 | FDR correction BH |
| CYLD~Q8IPC5 | VMAT~Control | VMAT | Control | 16.34808287 | 0.050047218 | 16.8366347 | 0.147220431 | 0.712740182 | -0.488551834 | -0.456787169 | 3.60E-05 | 4.443337729 | Unpaired t-test | 0.150162739 | permutation FDR (250 perm) | 1 | 7.90E-05 | FDR correction BH |
| Unc-115a~A0A0B4LGZ0 | VMAT~Control | VMAT | Control | 18.92158446 | 0.081386975 | 19.41266225 | 0.08975059 | 0.71149337 | -0.491077782 | -0.465837302 | 3.89E-06 | 5.410596638 | Unpaired t-test | 0.144117386 | permutation FDR (250 perm) | 1 | 1.11E-05 | FDR correction BH |
| tzn~Q8IPU3 | VMAT~Control | VMAT | Control | 18.24307438 | 0.06878228 | 18.73502357 | 0.066159688 | 0.711063747 | -0.491949192 | -0.471812039 | 4.26E-07 | 6.370328485 | Unpaired t-test | 0.140222519 | permutation FDR (250 perm) | 1 | 1.62E-06 | FDR correction BH |
| Ndf~X2JDP6 | VMAT~Control | VMAT | Control | 16.16494233 | 0.121284872 | 16.65696663 | 0.186182733 | 0.711026731 | -0.492024296 | -0.447550301 | 0.000577391 | 3.238529922 | Unpaired t-test | 0.156037284 | permutation FDR (250 perm) | 1 | 0.000959872 | FDR correction BH |
| Unc-89~A8DYP0-3 | VMAT~Control | VMAT | Control | 18.50681427 | 0.07316463 | 19.00077211 | 0.053024107 | 0.710074428 | -0.493957843 | -0.474772545 | 2.45E-07 | 6.609965194 | Unpaired t-test | 0.138294615 | permutation FDR (250 perm) | 1 | 1.02E-06 | FDR correction BH |
| eas~P54352-2 | VMAT~Control | VMAT | Control | 16.0092444 | 0.211583915 | 16.50589305 | 0.097321944 | 0.708751283 | -0.496648653 | -0.449800544 | 0.000758819 | 3.119861731 | Unpaired t-test | 0.15462167 | permutation FDR (250 perm) | 1 | 0.001236166 | FDR correction BH |
| alpha-Est5~Q95U30 | VMAT~Control | VMAT | Control | 15.04786593 | 0.180686058 | 15.54618234 | 0.094015547 | 0.707932441 | -0.498316406 | -0.45671456 | 0.000272814 | 3.564133367 | Unpaired t-test | 0.150200089 | permutation FDR (250 perm) | 1 | 0.000491357 | FDR correction BH |
| Rip11~Q494I2 | VMAT~Control | VMAT | Control | 13.56544874 | 0.370892443 | 14.06400349 | 0.579842226 | 0.707815495 | -0.498554749 | -0.381209347 | 0.136384842 | 0.865233896 | Unpaired t-test | 0.203343046 | permutation FDR (250 perm) | 1 | 0.160524959 | FDR correction BH |
| Lsd-1~A0A0B4KHZ1 | VMAT~Control | VMAT | Control | 16.95116652 | 0.180625245 | 17.45028507 | 0.510332254 | 0.707538934 | -0.499118557 | -0.401834113 | 0.06620803 | 1.179089334 | Unpaired t-test | 0.187243482 | permutation FDR (250 perm) | 1 | 0.081258448 | FDR correction BH |
| ALiX~Q9VB05 | VMAT~Control | VMAT | Control | 17.09495387 | 0.230831244 | 17.59457943 | 0.287759164 | 0.707290328 | -0.499625562 | -0.42887131 | 0.01271093 | 1.895822656 | Unpaired t-test | 0.168334958 | permutation FDR (250 perm) | 1 | 0.017305686 | FDR correction BH |
| Shaw~M9PEE2 | VMAT~Control | VMAT | Control | 22.30919015 | 0.385791814 | 22.81149217 | 0.541296947 | 0.705979393 | -0.502302022 | -0.387200248 | 0.121961453 | 0.913777411 | Unpaired t-test | 0.19855011 | permutation FDR (250 perm) | 1 | 0.144487801 | FDR correction BH |
| Yp1~X2JD55 | VMAT~Control | VMAT | Control | 18.49510038 | 0.32652536 | 19.00123282 | 0.395347888 | 0.704107472 | -0.506132443 | -0.411720262 | 0.05180721 | 1.285609792 | Unpaired t-test | 0.180329204 | permutation FDR (250 perm) | 1 | 0.064526018 | FDR correction BH |
| Cndp2~Q8MT58 | VMAT~Control | VMAT | Control | 15.61411169 | 0.216848462 | 16.12547181 | 0.166089465 | 0.701560721 | -0.511360118 | -0.455694813 | 0.001869308 | 2.728319072 | Unpaired t-test | 0.150854602 | permutation FDR (250 perm) | 1 | 0.002851816 | FDR correction BH |
| Dmel\CG1578~Q8MRI4 | VMAT~Control | VMAT | Control | 16.54840034 | 0.103890864 | 17.06331362 | 0.194538351 | 0.699835 | -0.514913277 | -0.468687098 | 0.000389478 | 3.409516652 | Unpaired t-test | 0.142288256 | permutation FDR (250 perm) | 1 | 0.000670689 | FDR correction BH |
| sds22~Q9VEK8 | VMAT~Control | VMAT | Control | 15.33609679 | 0.19658439 | 15.85175225 | 0.085784745 | 0.699475071 | -0.515655454 | -0.470522355 | 0.000311937 | 3.505932558 | Unpaired t-test | 0.141151758 | permutation FDR (250 perm) | 1 | 0.000551712 | FDR correction BH |
| twf~Q9VFM9 | VMAT~Control | VMAT | Control | 17.0306896 | 0.139731412 | 17.54709011 | 0.140202182 | 0.69911393 | -0.516400515 | -0.474404824 | 0.000165258 | 3.781838031 | Unpaired t-test | 0.138518914 | permutation FDR (250 perm) | 1 | 0.000309974 | FDR correction BH |
| Npl4~Q9VBP9-2 | VMAT~Control | VMAT | Control | 16.53493291 | 0.055954647 | 17.05528618 | 0.152534362 | 0.697201088 | -0.520353274 | -0.485105336 | 3.06E-05 | 4.513866943 | Unpaired t-test | 0.13218198 | permutation FDR (250 perm) | 1 | 6.82E-05 | FDR correction BH |
| subdued~Q9VDV4 | VMAT~Control | VMAT | Control | 14.77672297 | 0.077808927 | 15.30062658 | 0.207458078 | 0.695487451 | -0.523903612 | -0.47667081 | 0.000353969 | 3.451034905 | Unpaired t-test | 0.137329176 | permutation FDR (250 perm) | 1 | 0.000617217 | FDR correction BH |
| Dph2~Q9VFE9 | VMAT~Control | VMAT | Control | 16.91310914 | 0.098053805 | 17.43760996 | 0.218836619 | 0.695199611 | -0.52450082 | -0.473700344 | 0.000631687 | 3.199497929 | Unpaired t-test | 0.139006676 | permutation FDR (250 perm) | 1 | 0.001044236 | FDR correction BH |
| NO66~E2QD64 | VMAT~Control | VMAT | Control | 15.57643511 | 0.159500876 | 16.10389328 | 0.161044943 | 0.693775994 | -0.527458174 | -0.478912419 | 0.000399291 | 3.398710864 | Unpaired t-test | 0.135887701 | permutation FDR (250 perm) | 1 | 0.000686582 | FDR correction BH |
| Dmel\CG1943~Q9VI56 | VMAT~Control | VMAT | Control | 16.97807727 | 0.1862706 | 17.50838319 | 0.191638338 | 0.692407895 | -0.530305919 | -0.473691558 | 0.001260221 | 2.899553363 | Unpaired t-test | 0.139015576 | permutation FDR (250 perm) | 1 | 0.001984321 | FDR correction BH |
| pps~Q9VG78 | VMAT~Control | VMAT | Control | 15.95221199 | 0.430410064 | 16.48627754 | 0.671620792 | 0.690605849 | -0.534065543 | -0.393638 | 0.165258123 | 0.781837184 | Unpaired t-test | 0.1934234 | permutation FDR (250 perm) | 1 | 0.191257435 | FDR correction BH |
| Pkn~A0A0B4K6U3 | VMAT~Control | VMAT | Control | 15.23958011 | 0.29938749 | 15.77428483 | 0.345506716 | 0.690299951 | -0.534704714 | -0.443939403 | 0.025806009 | 1.58827916 | Unpaired t-test | 0.158347093 | permutation FDR (250 perm) | 1 | 0.033488062 | FDR correction BH |
| I-2~Q9V3C7 | VMAT~Control | VMAT | Control | 16.70354092 | 0.137294116 | 17.24233685 | 0.146881965 | 0.688345159 | -0.538795934 | -0.494346519 | 0.000133482 | 3.874576512 | Unpaired t-test | 0.126876565 | permutation FDR (250 perm) | 1 | 0.000254633 | FDR correction BH |
| B52~A0A0C4DHG5 | VMAT~Control | VMAT | Control | 16.76352514 | 0.282883813 | 17.30454777 | 0.441714862 | 0.687283567 | -0.54102263 | -0.438224582 | 0.043777766 | 1.3587464 | Unpaired t-test | 0.162225377 | permutation FDR (250 perm) | 1 | 0.055049606 | FDR correction BH |
| Dmel\CG6144~Q9VKU5 | VMAT~Control | VMAT | Control | 14.72594812 | 0.27226753 | 15.26961069 | 0.173975586 | 0.686027079 | -0.543662571 | -0.475023094 | 0.003707238 | 2.43094956 | Unpaired t-test | 0.138143302 | permutation FDR (250 perm) | 1 | 0.005435245 | FDR correction BH |
| Rs1~Q7JQN4 | VMAT~Control | VMAT | Control | 14.97004615 | 0.548906953 | 15.51401089 | 0.429850959 | 0.685883407 | -0.54396474 | -0.41467307 | 0.111644258 | 0.952163608 | Unpaired t-test | 0.17820885 | permutation FDR (250 perm) | 1 | 0.133136061 | FDR correction BH |
| Dmel\CG8475~M9PCI5 | VMAT~Control | VMAT | Control | 15.78737116 | 0.084634684 | 16.3318674 | 0.112885637 | 0.685630771 | -0.544496235 | -0.51217923 | 6.03E-06 | 5.219758762 | Unpaired t-test | 0.116277653 | permutation FDR (250 perm) | 1 | 1.64E-05 | FDR correction BH |
| GS~M9NFH8 | VMAT~Control | VMAT | Control | 18.55254069 | 0.07350909 | 19.10121532 | 0.120930682 | 0.683647891 | -0.54867463 | -0.516016212 | 5.79E-06 | 5.237570131 | Unpaired t-test | 0.114244733 | permutation FDR (250 perm) | 1 | 1.58E-05 | FDR correction BH |
| eag~M9PHM3 | VMAT~Control | VMAT | Control | 14.12411051 | 0.325524623 | 14.67474974 | 0.134882059 | 0.682717562 | -0.550639231 | -0.475680797 | 0.005781147 | 2.237985977 | Unpaired t-test | 0.137834372 | permutation FDR (250 perm) | 1 | 0.008277871 | FDR correction BH |
| mtm~Q9VMI9 | VMAT~Control | VMAT | Control | 21.15227564 | 0.143049284 | 21.70880052 | 0.238513682 | 0.679938008 | -0.556524878 | -0.494961576 | 0.001189301 | 2.924708371 | Unpaired t-test | 0.126470483 | permutation FDR (250 perm) | 1 | 0.001879838 | FDR correction BH |
| endos~Q9VUB8 | VMAT~Control | VMAT | Control | 17.09643654 | 0.113845257 | 17.66017159 | 0.105795103 | 0.676548349 | -0.563735056 | -0.527100015 | 1.04E-05 | 4.981520635 | Unpaired t-test | 0.108256963 | permutation FDR (250 perm) | 1 | 2.60E-05 | FDR correction BH |

| | | | | | | | | | | | | | | | | | | |
|---|---|---|---|---|---|---|---|---|---|---|---|---|---|---|---|---|---|---|
| Hrb27C~E1JHA4 | VMAT~Control | VMAT | Control | 18.26999147 | 0.094755125 | 18.8402694 | 0.134097466 | 0.67348703 | -0.570277933 | -0.531266431 | 1.53E-05 | 4.816609051 | Unpaired t-test | 0.105919137 | permutation FDR (250 perm) | 1 | 3.65E-05 | FDR correction BH |
| Septin1~P42207 | VMAT~Control | VMAT | Control | 15.05046793 | 0.131059517 | 15.62224034 | 0.356634246 | 0.672789731 | -0.57177241 | -0.488727649 | 0.007182286 | 2.143737316 | Unpaired t-test | 0.130125949 | permutation FDR (250 perm) | 1 | 0.010178869 | FDR correction BH |
| Pfdn5~E1JIT4 | VMAT~Control | VMAT | Control | 15.90045771 | 0.200072841 | 16.47303167 | 0.234752964 | 0.672416038 | -0.57257396 | -0.50316681 | 0.00197727 | 2.703934046 | Unpaired t-test | 0.121565996 | permutation FDR (250 perm) | 1 | 0.003004838 | FDR correction BH |
| smash~B7Z0T2 | VMAT~Control | VMAT | Control | 16.30034417 | 0.113784328 | 16.87522719 | 0.197843911 | 0.671340685 | -0.574883019 | -0.521640381 | 0.000217697 | 3.662147285 | Unpaired t-test | 0.111244165 | permutation FDR (250 perm) | 1 | 0.000398491 | FDR correction BH |
| eIF3e~O77410 | VMAT~Control | VMAT | Control | 16.9459215 | 0.146656625 | 17.52342744 | 0.049447951 | 0.670121251 | -0.577505936 | -0.540121662 | 8.13E-06 | 5.089870877 | Unpaired t-test | 0.101259559 | permutation FDR (250 perm) | 1 | 2.10E-05 | FDR correction BH |
| Dmel\CG12608~Q9VXW3 | VMAT~Control | VMAT | Control | 13.54252203 | 0.317812845 | 14.12425958 | 0.526378994 | 0.668158577 | -0.58173755 | -0.456270621 | 0.060476307 | 1.218414739 | Unpaired t-test | 0.150498888 | permutation FDR (250 perm) | 1 | 0.074651928 | FDR correction BH |
| Gfat2~Q9VAW3 | VMAT~Control | VMAT | Control | 16.09833577 | 0.23157393 | 16.68396655 | 0.272655709 | 0.666357928 | -0.585630778 | -0.504862928 | 0.00438433 | 2.35809675 | Unpaired t-test | 0.120606711 | permutation FDR (250 perm) | 1 | 0.006362955 | FDR correction BH |
| Crk~Q9XYM0 | VMAT~Control | VMAT | Control | 16.61636057 | 0.117528018 | 17.20404629 | 0.178775628 | 0.66540946 | -0.587685718 | -0.536366073 | 0.00010944 | 3.960822652 | Unpaired t-test | 0.103124494 | permutation FDR (250 perm) | 1 | 0.000212911 | FDR correction BH |
| Ate1~A0A0B4KFD5 | VMAT~Control | VMAT | Control | 14.74286083 | 0.111297673 | 15.33306681 | 0.266169393 | 0.664248063 | -0.59020598 | -0.52275864 | 0.001019253 | 2.991717916 | Unpaired t-test | 0.110639138 | permutation FDR (250 perm) | 1 | 0.001629974 | FDR correction BH |
| RhoGAP92B~A0A0B4LHN0 | VMAT~Control | VMAT | Control | 15.34190547 | 0.230331389 | 15.93226538 | 0.484202522 | 0.664177196 | -0.590359906 | -0.476175994 | 0.033564175 | 1.474124027 | Unpaired t-test | 0.137540516 | permutation FDR (250 perm) | 1 | 0.042940254 | FDR correction BH |
| Ank~Q0KIE7 | VMAT~Control | VMAT | Control | 17.4940741 | 0.105826596 | 18.08732765 | 0.147419171 | 0.662846377 | -0.59325355 | -0.548721383 | 2.57E-05 | 4.58990508 | Unpaired t-test | 0.096920306 | permutation FDR (250 perm) | 1 | 5.82E-05 | FDR correction BH |
| alph~Q9VAK1 | VMAT~Control | VMAT | Control | 17.6321325 | 0.153362919 | 18.22565668 | 0.317998181 | 0.662722051 | -0.593524172 | -0.512592061 | 0.003727368 | 2.428597691 | Unpaired t-test | 0.116046574 | permutation FDR (250 perm) | 1 | 0.005460003 | FDR correction BH |
| CG12167~Q9VNC4 | VMAT~Control | VMAT | Control | 13.24489784 | 0.313365546 | 13.83857206 | 0.456296627 | 0.662653128 | -0.59367422 | -0.475872152 | 0.037430526 | 1.426774075 | Unpaired t-test | 0.137707925 | permutation FDR (250 perm) | 1 | 0.047576381 | FDR correction BH |
| CRMP~Q9V3N7 | VMAT~Control | VMAT | Control | 22.07470022 | 0.149761358 | 22.66873702 | 0.081584325 | 0.662486609 | -0.594036801 | -0.551941052 | 1.49E-05 | 4.827580686 | Unpaired t-test | 0.095160738 | permutation FDR (250 perm) | 1 | 3.58E-05 | FDR correction BH |
| Dmel\CG9330~Q9VW13 | VMAT~Control | VMAT | Control | 15.74241515 | 0.120083708 | 16.3373833 | 0.055307229 | 0.662059071 | -0.594968151 | -0.561754278 | 1.50E-06 | 5.823674016 | Unpaired t-test | 0.09023683 | permutation FDR (250 perm) | 1 | 4.81E-06 | FDR correction BH |
| epsilonCOP~Q9Y0Y5 | VMAT~Control | VMAT | Control | 16.46776351 | 0.136483723 | 17.06307483 | 0.221805952 | 0.661901611 | -0.595311312 | -0.53320877 | 0.000456435 | 3.340620812 | Unpaired t-test | 0.104819407 | permutation FDR (250 perm) | 1 | 0.000777459 | FDR correction BH |
| PpD3~Q9VH81 | VMAT~Control | VMAT | Control | 16.85973909 | 0.056216641 | 17.45787049 | 0.149148838 | 0.660609033 | -0.598131398 | -0.558332301 | 7.73E-06 | 5.11162178 | Unpaired t-test | 0.091857722 | permutation FDR (250 perm) | 1 | 2.00E-05 | FDR correction BH |
| Vha14-1~Q24583 | VMAT~Control | VMAT | Control | 17.07630169 | 0.061525065 | 17.67713586 | 0.185995514 | 0.659372598 | -0.600834163 | -0.552434086 | 4.42E-05 | 4.354994451 | Unpaired t-test | 0.094910401 | permutation FDR (250 perm) | 1 | 9.43E-05 | FDR correction BH |
| spg~B7Z0R2 | VMAT~Control | VMAT | Control | 13.09473492 | 0.334932509 | 13.69742045 | 0.247341489 | 0.658526988 | -0.602685528 | -0.508079628 | 0.008918921 | 2.049687689 | Unpaired t-test | 0.11870962 | permutation FDR (250 perm) | 1 | 0.012437879 | FDR correction BH |
| mod(mdg4)~Q86B87-26 | VMAT~Control | VMAT | Control | 13.21507332 | 0.330418234 | 13.82276448 | 0.47088442 | 0.656246095 | -0.607691163 | -0.483346435 | 0.039804522 | 1.400067587 | Unpaired t-test | 0.133184128 | permutation FDR (250 perm) | 1 | 0.050322151 | FDR correction BH |
| CTPsyn~X2JCY2 | VMAT~Control | VMAT | Control | 15.67942344 | 0.055277141 | 16.28838375 | 0.13054942 | 0.655669044 | -0.608960312 | -0.57265327 | 2.30E-06 | 5.638780322 | Unpaired t-test | 0.085104505 | permutation FDR (250 perm) | 1 | 6.95E-06 | FDR correction BH |
| chas~X2JE77 | VMAT~Control | VMAT | Control | 13.43726399 | 0.448912875 | 14.04669283 | 0.584903492 | 0.655456144 | -0.609428842 | -0.458307549 | 0.094319467 | 1.025398664 | Unpaired t-test | 0.149116941 | permutation FDR (250 perm) | 1 | 0.113453257 | FDR correction BH |
| CT28495~Q9VG23 | VMAT~Control | VMAT | Control | 15.29428704 | 0.199169039 | 15.90382896 | 0.232713255 | 0.655404769 | -0.609541925 | -0.536103989 | 0.001235696 | 2.908088508 | Unpaired t-test | 0.103264719 | permutation FDR (250 perm) | 1 | 0.001947006 | FDR correction BH |
| Sod1~P61851 | VMAT~Control | VMAT | Control | 16.6349146 | 0.080145243 | 17.24654759 | 0.250707145 | 0.654455505 | -0.611632986 | -0.547220119 | 0.000403546 | 3.394107447 | Unpaired t-test | 0.097730752 | permutation FDR (250 perm) | 1 | 0.000692381 | FDR correction BH |
| Madm~A0A126GUQ5 | VMAT~Control | VMAT | Control | 14.55544701 | 0.247504106 | 15.17381261 | 0.199807975 | 0.651408478 | -0.618365599 | -0.541355431 | 0.001450601 | 2.838452049 | Unpaired t-test | 0.100653171 | permutation FDR (250 perm) | 1 | 0.002251766 | FDR correction BH |
| Zasp66~Q9VSN2 | VMAT~Control | VMAT | Control | 17.62600678 | 0.268919036 | 18.24525637 | 0.285454418 | 0.651009461 | -0.619249584 | -0.526847738 | 0.005439763 | 2.264420035 | Unpaired t-test | 0.108438455 | permutation FDR (250 perm) | 1 | 0.007812814 | FDR correction BH |
| Csk~Q7KSQ2 | VMAT~Control | VMAT | Control | 14.20438098 | 0.06878098 | 14.82492754 | 0.037370383 | 0.650424468 | -0.620546565 | -0.599557979 | 6.96E-09 | 8.157493993 | Unpaired t-test | 0.073761302 | permutation FDR (250 perm) | 1 | 5.67E-08 | FDR correction BH |
| TBCB~A1ZBM2 | VMAT~Control | VMAT | Control | 16.86918718 | 0.09772497 | 17.49129492 | 0.128886754 | 0.649721009 | -0.62210774 | -0.580142937 | 6.21E-06 | 5.206658494 | Unpaired t-test | 0.081679279 | permutation FDR (250 perm) | 1 | 1.68E-05 | FDR correction BH |
| RpL22~P50887 | VMAT~Control | VMAT | Control | 17.46560387 | 0.193357558 | 18.08793268 | 0.127056582 | 0.649621456 | -0.622328812 | -0.563974227 | 0.00012957 | 3.887496682 | Unpaired t-test | 0.089100405 | permutation FDR (250 perm) | 1 | 0.000247973 | FDR correction BH |
| Tap42~Q9VJZ5 | VMAT~Control | VMAT | Control | 15.87970225 | 0.182717205 | 16.50617716 | 0.210243084 | 0.647757217 | -0.62647491 | -0.55707979 | 0.000514788 | 3.288371241 | Unpaired t-test | 0.092488068 | permutation FDR (250 perm) | 1 | 0.000866199 | FDR correction BH |
| Ankrd49~Q9V3Y0 | VMAT~Control | VMAT | Control | 14.65249633 | 0.213124249 | 15.2797284 | 0.380530955 | 0.647417348 | -0.62723207 | -0.524857703 | 0.009241921 | 2.034237733 | Unpaired t-test | 0.109561545 | permutation FDR (250 perm) | 1 | 0.012850256 | FDR correction BH |
| Dmel\CG7139~Q9XZ11 | VMAT~Control | VMAT | Control | 12.90033147 | 0.33102638 | 13.5318972 | 0.423816142 | 0.645475513 | -0.631565729 | -0.509122364 | 0.025332659 | 1.596319222 | Unpaired t-test | 0.11810385 | permutation FDR (250 perm) | 1 | 0.032910088 | FDR correction BH |
| sif~Q8IQ62 | VMAT~Control | VMAT | Control | 21.14656683 | 0.11820594 | 21.78567695 | 0.214004225 | 0.642108892 | -0.639110118 | -0.576120157 | 0.00016261 | 3.788853521 | Unpaired t-test | 0.083513514 | permutation FDR (250 perm) | 1 | 0.000305493 | FDR correction BH |
| Asap~A1Z7A6-2 | VMAT~Control | VMAT | Control | 15.10895727 | 0.220590047 | 15.74866907 | 0.412644066 | 0.641841155 | -0.639711797 | -0.529013816 | 0.012103913 | 1.917074212 | Unpaired t-test | 0.107203953 | permutation FDR (250 perm) | 1 | 0.016517456 | FDR correction BH |
| Dmel\CG9705~M9PFS8 | VMAT~Control | VMAT | Control | 18.59674096 | 0.126259486 | 19.23903722 | 0.069921163 | 0.640692384 | -0.642296254 | -0.60335272 | 1.66E-06 | 5.779221898 | Unpaired t-test | 0.072409235 | permutation FDR (250 perm) | 1 | 5.25E-06 | FDR correction BH |
| yrt~A0T1Z5 | VMAT~Control | VMAT | Control | 15.89991775 | 0.09106448 | 16.54894134 | 0.146048219 | 0.637711766 | -0.649023595 | -0.602637842 | 7.41E-06 | 5.130377172 | Unpaired t-test | 0.072662743 | permutation FDR (250 perm) | 1 | 1.94E-05 | FDR correction BH |
| p47~Q7K3Z3 | VMAT~Control | VMAT | Control | 18.17102474 | 0.087019673 | 18.82028375 | 0.1109691 | 0.637607717 | -0.649259005 | -0.610741987 | 1.22E-06 | 5.914698767 | Unpaired t-test | 0.069502943 | permutation FDR (250 perm) | 1 | 4.00E-06 | FDR correction BH |
| CG7687~Q8MQJ5 | VMAT~Control | VMAT | Control | 14.78707904 | 0.179800472 | 15.43963901 | 0.155492832 | 0.636150503 | -0.65255997 | -0.589854156 | 0.000109982 | 3.958679096 | Unpaired t-test | 0.077631033 | permutation FDR (250 perm) | 1 | 0.000213435 | FDR correction BH |
| nrm~Q0E8B8 | VMAT~Control | VMAT | Control | 17.38993163 | 0.087444276 | 18.04918863 | 0.132146282 | 0.633204319 | -0.659256999 | -0.615630435 | 3.07E-06 | 5.512931177 | Unpaired t-test | 0.067692029 | permutation FDR (250 perm) | 1 | 8.99E-06 | FDR correction BH |
| gammaCOP~A0A0B4KHL2 | VMAT~Control | VMAT | Control | 18.17621482 | 0.062980345 | 18.83738155 | 0.088448607 | 0.632366685 | -0.66116673 | -0.630548203 | 8.83E-08 | 7.053954833 | Unpaired t-test | 0.062407978 | permutation FDR (250 perm) | 1 | 4.30E-07 | FDR correction BH |
| mora~Q9VCC0 | VMAT~Control | VMAT | Control | 17.40625104 | 0.057814595 | 18.06790528 | 0.223429899 | 0.632153034 | -0.66165424 | -0.599752649 | 7.73E-05 | 4.112015253 | Unpaired t-test | 0.073674503 | permutation FDR (250 perm) | 1 | 0.000155721 | FDR correction BH |
| Txl~Q9VRP3 | VMAT~Control | VMAT | Control | 18.90181495 | 0.099461228 | 19.56422988 | 0.192930213 | 0.631819808 | -0.662414929 | -0.603802751 | 4.60E-05 | 4.336958561 | Unpaired t-test | 0.072220914 | permutation FDR (250 perm) | 1 | 9.78E-05 | FDR correction BH |
| p23~A0A0B4K6D2 | VMAT~Control | VMAT | Control | 16.98697755 | 0.079474561 | 17.65299595 | 0.189371432 | 0.630243656 | -0.666018404 | -0.60999341 | 2.75E-05 | 4.56010311 | Unpaired t-test | 0.069828882 | permutation FDR (250 perm) | 1 | 6.16E-05 | FDR correction BH |
| Mi-2~M9PIA6 | VMAT~Control | VMAT | Control | 15.73798784 | 0.153641998 | 16.40551485 | 0.290139757 | 0.629584963 | -0.667527012 | -0.582065609 | 0.001064117 | 2.973010474 | Unpaired t-test | 0.080868468 | permutation FDR (250 perm) | 1 | 0.00169596 | FDR correction BH |
| dnc~E2QD73 | VMAT~Control | VMAT | Control | 16.39310149 | 0.110662133 | 17.06103749 | 0.130514772 | 0.629406505 | -0.667936005 | -0.620455761 | 5.45E-06 | 5.263905697 | Unpaired t-test | 0.065951087 | permutation FDR (250 perm) | 1 | 1.50E-05 | FDR correction BH |
| Dmel\CG8578~Q9VXN3 | VMAT~Control | VMAT | Control | 17.97462386 | 0.059572764 | 18.64350088 | 0.060280608 | 0.628996101 | -0.668877022 | -0.644451271 | 7.27E-09 | 8.138704097 | Unpaired t-test | 0.057754991 | permutation FDR (250 perm) | 1 | 5.84E-08 | FDR correction BH |
| dos~Q9VZZ9-2 | VMAT~Control | VMAT | Control | 13.39708591 | 0.253174754 | 14.06614093 | 0.918516684 | 0.628918502 | -0.669055017 | -0.469152898 | 0.147438404 | 0.831389378 | Unpaired t-test | 0.142089898 | permutation FDR (250 perm) | 1 | 0.172722688 | FDR correction BH |
| Gart~X2J516 | VMAT~Control | VMAT | Control | 14.14470781 | 0.172480233 | 14.81634034 | 0.397603793 | 0.62779588 | -0.671632533 | -0.56258936 | 0.006069964 | 2.216813901 | Unpaired t-test | 0.089746961 | permutation FDR (250 perm) | 1 | 0.008665067 | FDR correction BH |
| kermit~Q7JX82 | VMAT~Control | VMAT | Control | 16.69552384 | 0.35045392 | 17.36760989 | 0.53155715 | 0.627598561 | -0.67208605 | -0.523131925 | 0.039926749 | 1.398736049 | Unpaired t-test | 0.110398564 | permutation FDR (250 perm) | 1 | 0.05044958 | FDR correction BH |
| Sec16~X2JJP5 | VMAT~Control | VMAT | Control | 16.84521799 | 0.172828184 | 17.51948704 | 0.113038252 | 0.626649634 | -0.674269051 | -0.617261835 | 2.60E-05 | 4.585240197 | Unpaired t-test | 0.06717029 | permutation FDR (250 perm) | 1 | 5.87E-05 | FDR correction BH |
| Ldsdh1~Q9W3K9 | VMAT~Control | VMAT | Control | 13.37996724 | 0.37750921 | 14.05476295 | 0.989613653 | 0.626420918 | -0.674795706 | -0.4578994 | 0.184726971 | 0.733469692 | Unpaired t-test | 0.149417519 | permutation FDR (250 perm) | 1 | 0.212535332 | FDR correction BH |
| ArgRS~Q9VXN4 | VMAT~Control | VMAT | Control | 14.95694404 | 0.181746676 | 15.6333746 | 0.099664837 | 0.625711463 | -0.67643056 | -0.619046008 | 2.61E-05 | 4.583345257 | Unpaired t-test | 0.066492754 | permutation FDR (250 perm) | 1 | 5.89E-05 | FDR correction BH |
| RhoGAP1A~X2J9V9 | VMAT~Control | VMAT | Control | 16.82685381 | 0.122308648 | 17.50351235 | 0.134834811 | 0.625612593 | -0.676658541 | -0.625717431 | 8.41E-06 | 5.075017803 | Unpaired t-test | 0.064099728 | permutation FDR (250 perm) | 1 | 2.16E-05 | FDR correction BH |
| Gclc~A4V448 | VMAT~Control | VMAT | Control | 16.3421107 | 0.078371382 | 17.02384483 | 0.115256119 | 0.623415474 | -0.68173413 | -0.641733925 | 6.95E-07 | 6.157863369 | Unpaired t-test | 0.058604356 | permutation FDR (250 perm) | 1 | 2.46E-06 | FDR correction BH |
| Dmel\CG12279~Q9VG33 | VMAT~Control | VMAT | Control | 16.00373082 | 0.133296721 | 16.68818349 | 0.259320821 | 0.622241848 | -0.68445267 | -0.60549825 | 0.000373903 | 3.427240595 | Unpaired t-test | 0.071536442 | permutation FDR (250 perm) | 1 | 0.000646707 | FDR correction BH |
| bond~Q9VCY7 | VMAT~Control | VMAT | Control | 15.56605668 | 0.418394357 | 16.25540174 | 0.329285818 | 0.620135307 | -0.689345064 | -0.556771854 | 0.015968042 | 1.796748335 | Unpaired t-test | 0.092655561 | permutation FDR (250 perm) | 1 | 0.021333014 | FDR correction BH |
| nsl1~E1JIM1 | VMAT~Control | VMAT | Control | 13.1537693 | 0.300339779 | 13.84508917 | 0.333817299 | 0.619287028 | -0.691319869 | -0.575707778 | 0.006304604 | 2.200342195 | Unpaired t-test | 0.08369009 | permutation FDR (250 perm) | 1 | 0.008983679 | FDR correction BH |
| APP-BP1~M9PEU5 | VMAT~Control | VMAT | Control | 14.55385734 | 0.268467151 | 15.24634515 | 0.390844139 | 0.618785884 | -0.692487809 | -0.571334288 | 0.008492914 | 2.070943278 | Unpaired t-test | 0.085704505 | permutation FDR (250 perm) | 1 | 0.011928592 | FDR correction BH |
| Acbp1~Q9VLS4 | VMAT~Control | VMAT | Control | 13.71760785 | 0.62918774 | 14.41258276 | 0.765655999 | 0.61772006 | -0.694974912 | -0.481553226 | 0.147927714 | 0.829950454 | Unpaired t-test | 0.134298707 | permutation FDR (250 perm) | 1 | 0.172986507 | FDR correction BH |
| Vha55~E1JIJ5 | VMAT~Control | VMAT | Control | 22.29408753 | 0.071022837 | 22.98982874 | 0.102089413 | 0.61739204 | -0.695741212 | -0.65908462 | 1.97E-07 | 6.704615937 | Unpaired t-test | 0.053137665 | permutation FDR (250 perm) | 1 | 8.59E-07 | FDR correction BH |
| upSET~Q9VUB5 | VMAT~Control | VMAT | Control | 14.55836374 | 0.207389224 | 15.26171309 | 0.389457019 | 0.614144757 | -0.70334935 | -0.587433627 | 0.005143367 | 2.288752519 | Unpaired t-test | 0.078669071 | permutation FDR (250 perm) | 1 | 0.007418802 | FDR correction BH |
| Trx-2~Q9V429 | VMAT~Control | VMAT | Control | 20.88032 | 0.248512787 | 21.58417822 | 0.420586364 | 0.613928173 | -0.703858219 | -0.57765615 | 0.009148397 | 2.038654994 | Unpaired t-test | 0.082765766 | permutation FDR (250 perm) | 1 | 0.012727734 | FDR correction BH |
| CG13621~Q9VC53 | VMAT~Control | VMAT | Control | 13.45876061 | 0.323531127 | 14.16601391 | 0.598006584 | 0.61248512 | -0.707253299 | -0.542344219 | 0.042339446 | 1.373254827 | Unpaired t-test | 0.100122412 | permutation FDR (250 perm) | 1 | 0.053297891 | FDR correction BH |
| eEF1alpha1~P08736 | VMAT~Control | VMAT | Control | 22.35604072 | 0.093264869 | 23.06539338 | 0.106567721 | 0.611594501 | -0.709352661 | -0.667103351 | 5.57E-07 | 6.253989926 | Unpaired t-test | 0.050941818 | permutation FDR (250 perm) | 1 | 2.04E-06 | FDR correction BH |
| CIAPIN1~Q9V3Y2 | VMAT~Control | VMAT | Control | 19.40476289 | 0.041705736 | 20.11978812 | 0.146464343 | 0.609194478 | -0.715025229 | -0.669433724 | 1.02E-06 | 5.993295167 | Unpaired t-test | 0.050234545 | permutation FDR (250 perm) | 1 | 3.42E-06 | FDR correction BH |
| dlg1~P31007-6 | VMAT~Control | VMAT | Control | 17.73653141 | 0.061386307 | 18.45634751 | 0.115383454 | 0.607174834 | -0.7198161 | -0.68006666 | 2.29E-07 | 6.640467555 | Unpaired t-test | 0.047379718 | permutation FDR (250 perm) | 1 | 9.65E-07 | FDR correction BH |
| pnut~P40797 | VMAT~Control | VMAT | Control | 16.22350591 | 0.157319651 | 16.94764627 | 0.146797301 | 0.605357646 | -0.724140355 | -0.660574722 | 2.00E-05 | 4.698134198 | Unpaired t-test | 0.052757837 | permutation FDR (250 perm) | 1 | 4.64E-05 | FDR correction BH |
| vari~Q2PDR9 | VMAT~Control | VMAT | Control | 14.17470494 | 0.679019338 | 14.90232475 | 0.192483148 | 0.603899418 | -0.727619812 | -0.55305735 | 0.043858346 | 1.35794775 | Unpaired t-test | 0.094609635 | permutation FDR (250 perm) | 1 | 0.055121488 | FDR correction BH |
| ade3~Q8T9H8 | VMAT~Control | VMAT | Control | 13.54740647 | 0.387441455 | 14.28755678 | 0.815573825 | 0.598676975 | -0.74015031 | -0.527247753 | 0.096705921 | 1.014546935 | Unpaired t-test | 0.108168913 | permutation FDR (250 perm) | 1 | 0.116148719 | FDR correction BH |
| PTP-ER~Q9W2F3 | VMAT~Control | VMAT | Control | 14.73312491 | 1.421586126 | 15.47981911 | 0.336892712 | 0.595967596 | -0.746694203 | -0.451621966 | 0.27972393 | 0.553270379 | Unpaired t-test | 0.15343936 | permutation FDR (250 perm) | 1 | 0.314005785 | FDR correction BH |
| nudC~Q9VVA6 | VMAT~Control | VMAT | Control | 19.21832206 | 0.095053605 | 19.96534395 | 0.218537875 | 0.595832244 | -0.747021897 | -0.67507408 | 3.67E-05 | 4.434777706 | Unpaired t-test | 0.048707594 | permutation FDR (250 perm) | 1 | 8.04E-05 | FDR correction BH |
| OtopLa~Q9W4A6 | VMAT~Control | VMAT | Control | 17.87012953 | 0.081903698 | 18.61865635 | 0.275463676 | 0.595211036 | -0.748526818 | -0.66328115 | 0.000167373 | 3.776315106 | Unpaired t-test | 0.051907315 | permutation FDR (250 perm) | 1 | 0.000312943 | FDR correction BH |
| Cortactin~Q9VDF4 | VMAT~Control | VMAT | Control | 18.47122356 | 0.078403748 | 19.22439507 | 0.116716473 | 0.593297863 | -0.753171509 | -0.708613433 | 2.97E-07 | 6.52684362 | Unpaired t-test | 0.040925182 | permutation FDR (250 perm) | 1 | 1.19E-06 | FDR correction BH |
| CG3817~Q9VFE6 | VMAT~Control | VMAT | Control | 14.02951197 | 1.315702832 | 14.79067718 | 1.795765062 | 0.590019602 | -0.761165209 | -0.381426545 | 0.462200524 | 0.335169566 | Unpaired t-test | 0.203185872 | permutation FDR (250 perm) | 1 | 0.500009207 | FDR correction BH |
| Nup98-96~Q9VCH5 | VMAT~Control | VMAT | Control | 17.36873126 | 0.141670841 | 18.13143476 | 0.298902795 | 0.589390823 | -0.762703499 | -0.664417524 | 0.00042776 | 3.368799915 | Unpaired t-test | 0.051610909 | permutation FDR (250 perm) | 1 | 0.000731652 | FDR correction BH |
| Dmel\CG8066~Q9VFI3 | VMAT~Control | VMAT | Control | 17.95397623 | 0.146316668 | 18.71697361 | 0.147986561 | 0.589270773 | -0.762997382 | -0.698032636 | 9.50E-06 | 5.022435616 | Unpaired t-test | 0.043123063 | permutation FDR (250 perm) | 1 | 2.41E-05 | FDR correction BH |
| Mcm3~X2JAI4 | VMAT~Control | VMAT | Control | 13.51766004 | 0.369685613 | 14.28092354 | 0.203378607 | 0.589162088 | -0.763263497 | -0.64210168 | 0.002342531 | 2.630314614 | Unpaired t-test | 0.058479129 | permutation FDR (250 perm) | 1 | 0.003523526 | FDR correction BH |
| DCTN2-p50~Q7K2D2 | VMAT~Control | VMAT | Control | 17.19951539 | 0.188416061 | 17.96405403 | 0.214432957 | 0.588641579 | -0.764538644 | -0.677988385 | 0.000134065 | 3.872686032 | Unpaired t-test | 0.047972715 | permutation FDR (250 perm) | 1 | 0.000255537 | FDR correction BH |
| GlyT~Q9W1J0 | VMAT~Control | VMAT | Control | 17.13140772 | 0.130322967 | 17.89688852 | 0.246021641 | 0.58825729 | -0.765480801 | -0.680725319 | 0.000108598 | 3.964178821 | Unpaired t-test | 0.047203274 | permutation FDR (250 perm) | 1 | 0.000211973 | FDR correction BH |
| nahoda~Q9W1X5 | VMAT~Control | VMAT | Control | 15.11936158 | 0.13897401 | 15.88676372 | 0.418797178 | 0.587474389 | -0.767402137 | -0.640925348 | 0.003015259 | 2.520675386 | Unpaired t-test | 0.058892922 | permutation FDR (250 perm) | 1 | 0.004472539 | FDR correction BH |
| CAP~A0A0B4K7L3 | VMAT~Control | VMAT | Control | 19.20185857 | 0.210376577 | 19.97338375 | 0.104429282 | 0.585797857 | -0.77152518 | -0.698189501 | 2.47E-05 | 4.607788309 | Unpaired t-test | 0.043097539 | permutation FDR (250 perm) | 1 | 5.61E-05 | FDR correction BH |
| Nfl~L0MPW8 | VMAT~Control | VMAT | Control | 13.63403666 | 0.230882473 | 14.40769538 | 1.065868483 | 0.584932186 | -0.77365872 | -0.520027759 | 0.143764271 | 0.842349034 | Unpaired t-test | 0.112053836 | permutation FDR (250 perm) | 1 | 0.168620375 | FDR correction BH |
| cin~P39205 | VMAT~Control | VMAT | Control | 15.68484055 | 0.232744423 | 16.4606687 | 0.117473423 | 0.584053267 | -0.775828142 | -0.69481713 | 5.68E-05 | 4.2456587 | Unpaired t-test | 0.043830524 | permutation FDR (250 perm) | 1 | 0.000118323 | FDR correction BH |
| tamo~B7YZS3 | VMAT~Control | VMAT | Control | 13.13597711 | 0.282352615 | 13.91303731 | 0.476514021 | 0.583554699 | -0.777060202 | -0.622791652 | 0.010563221 | 1.976203626 | Unpaired t-test | 0.065149071 | permutation FDR (250 perm) | 1 | 0.014532918 | FDR correction BH |
| Usp14~Q9VKZ8 | VMAT~Control | VMAT | Control | 18.00490346 | 0.057629095 | 18.78849544 | 0.09541191 | 0.580918636 | -0.783591983 | -0.746385515 | 2.23E-08 | 7.652233897 | Unpaired t-test | 0.033966039 | permutation FDR (250 perm) | 1 | 1.41E-07 | FDR correction BH |
| CG16790~Q9VHB3-2 | VMAT~Control | VMAT | Control | 14.39791334 | 0.084803197 | 15.1840306 | 0.533863001 | 0.579902693 | -0.786117257 | -0.63307504 | 0.00869314 | 2.060823322 | Unpaired t-test | 0.061563719 | permutation FDR (250 perm) | 1 | 0.012159033 | FDR correction BH |
| Dmel\CG46466~A0A6H2EEL7 | VMAT~Control | VMAT | Control | 15.85170958 | 0.126933556 | 16.63790668 | 0.199441389 | 0.579870599 | -0.786197103 | -0.71102401 | 2.22E-05 | 4.653642121 | Unpaired t-test | 0.040427202 | permutation FDR (250 perm) | 1 | 5.09E-05 | FDR correction BH |
| Dmel\CG13994~Q9VMI2 | VMAT~Control | VMAT | Control | 17.43806194 | 0.151167908 | 18.22512073 | 0.379257338 | 0.579524362 | -0.787058785 | -0.665540536 | 0.001535572 | 2.813729946 | Unpaired t-test | 0.051367273 | permutation FDR (250 perm) | 1 | 0.00237499 | FDR correction BH |
| U2af50~M9NH51 | VMAT~Control | VMAT | Control | 14.9352606 | 0.690048651 | 15.7225659 | 0.624203793 | 0.579425347 | -0.787305298 | -0.55595773 | 0.087773011 | 1.056639002 | Unpaired t-test | 0.093037371 | permutation FDR (250 perm) | 1 | 0.105957779 | FDR correction BH |
| Cog1~A0A0B4KH76 | VMAT~Control | VMAT | Control | 14.58809951 | 0.079291416 | 15.37636422 | 0.70843124 | 0.57904015 | -0.788264708 | -0.597714356 | 0.032957127 | 1.482050651 | Unpaired t-test | 0.074603978 | permutation FDR (250 perm) | 1 | 0.042186557 | FDR correction BH |
| Fcp1~Q9W147 | VMAT~Control | VMAT | Control | 13.44493094 | 0.154793323 | 14.23478905 | 0.055562089 | 0.578400975 | -0.789858109 | -0.73574403 | 8.24E-07 | 6.084158667 | Unpaired t-test | 0.035818515 | permutation FDR (250 perm) | 1 | 2.84E-06 | FDR correction BH |
| Lmpt~M9MRX0 | VMAT~Control | VMAT | Control | 17.91898629 | 0.065964277 | 18.71007179 | 0.117880659 | 0.577909101 | -0.791085505 | -0.746018182 | 1.28E-07 | 6.892866489 | Unpaired t-test | 0.034013768 | permutation FDR (250 perm) | 1 | 5.97E-07 | FDR correction BH |
| anon-WO0172774.138~Q9VSC5 | VMAT~Control | VMAT | Control | 16.63396478 | 0.095328452 | 17.42506154 | 0.209975387 | 0.577904594 | -0.791096758 | -0.717139425 | 1.70E-05 | 4.770134033 | Unpaired t-test | 0.039157991 | permutation FDR (250 perm) | 1 | 3.99E-05 | FDR correction BH |
| Eb1~Q7JZD3 | VMAT~Control | VMAT | Control | 17.30508106 | 0.083347006 | 18.09801614 | 0.162089464 | 0.57716868 | -0.79293508 | -0.733173798 | 2.05E-06 | 5.689217161 | Unpaired t-test | 0.036245309 | permutation FDR (250 perm) | 1 | 6.33E-06 | FDR correction BH |
| eIF5~Q9VXK6 | VMAT~Control | VMAT | Control | 14.75400418 | 0.393741545 | 15.5471367 | 0.384200475 | 0.577089694 | -0.793132528 | -0.63653 | 0.009116491 | 2.040172294 | Unpaired t-test | 0.060342857 | permutation FDR (250 perm) | 1 | 0.012698355 | FDR correction BH |
| zetaCOP~Q9VV89 | VMAT~Control | VMAT | Control | 19.25018759 | 0.132809263 | 20.04638819 | 0.064714458 | 0.575863746 | -0.796200596 | -0.746855792 | 2.81E-07 | 6.551683669 | Unpaired t-test | 0.033907297 | permutation FDR (250 perm) | 1 | 1.14E-06 | FDR correction BH |
| Rok~X2JE40 | VMAT~Control | VMAT | Control | 14.8816159 | 0.133553317 | 15.68394635 | 0.239725245 | 0.573422155 | -0.802330448 | -0.714629025 | 6.57E-05 | 4.182177698 | Unpaired t-test | 0.039649315 | permutation FDR (250 perm) | 1 | 0.000134331 | FDR correction BH |
| bun~X2J847 | VMAT~Control | VMAT | Control | 15.27839565 | 0.177487641 | 16.08081467 | 0.117374991 | 0.573386951 | -0.802419022 | -0.732694558 | 7.41E-06 | 5.130436788 | Unpaired t-test | 0.036311213 | permutation FDR (250 perm) | 1 | 1.94E-05 | FDR correction BH |
| wrd~Q8IN90 | VMAT~Control | VMAT | Control | 17.32226993 | 0.105573041 | 18.12737982 | 0.223256525 | 0.572318485 | -0.805109891 | -0.725034356 | 2.63E-05 | 4.579608363 | Unpaired t-test | 0.037608417 | permutation FDR (250 perm) | 1 | 5.93E-05 | FDR correction BH |
| Dmel\CG1703~Q9VYW4 | VMAT~Control | VMAT | Control | 16.98633594 | 0.222280155 | 17.79579126 | 0.152949776 | 0.570597241 | -0.809455324 | -0.722298093 | 5.31E-05 | 4.274887848 | Unpaired t-test | 0.038177493 | permutation FDR (250 perm) | 1 | 0.00011141 | FDR correction BH |
| Dmel\CG5466~Q9VDK2 | VMAT~Control | VMAT | Control | 13.60615409 | 0.064168877 | 14.41588685 | 1.063826424 | 0.570487521 | -0.809732766 | -0.548368219 | 0.120181781 | 0.920161365 | Unpaired t-test | 0.097134624 | permutation FDR (250 perm) | 1 | 0.142666622 | FDR correction BH |
| IleRS~Q8MSW0 | VMAT~Control | VMAT | Control | 14.94676442 | 0.216582638 | 15.75817084 | 0.302104324 | 0.569826091 | -0.811406414 | -0.695746957 | 0.000641007 | 3.193137498 | Unpaired t-test | 0.043597267 | permutation FDR (250 perm) | 1 | 0.001058898 | FDR correction BH |
| Cen~Q9VIK6 | VMAT~Control | VMAT | Control | 16.15671799 | 0.130175871 | 16.969495 | 0.122398372 | 0.569284999 | -0.812777009 | -0.752634828 | 1.36E-06 | 5.866430446 | Unpaired t-test | 0.032976595 | permutation FDR (250 perm) | 1 | 4.40E-06 | FDR correction BH |
| Pglym78~Q9VAN7 | VMAT~Control | VMAT | Control | 19.93271147 | 0.198221914 | 20.74901999 | 0.227467365 | 0.567893176 | -0.81630852 | -0.719257597 | 0.000123646 | 3.907821471 | Unpaired t-test | 0.038764168 | permutation FDR (250 perm) | 1 | 0.000237601 | FDR correction BH |
| DCTN3-p24~Q9W1V8 | VMAT~Control | VMAT | Control | 16.50928558 | 0.226220645 | 17.32711047 | 0.283236123 | 0.567296595 | -0.81782489 | -0.703741293 | 0.000503017 | 3.298417006 | Unpaired t-test | 0.041870556 | permutation FDR (250 perm) | 1 | 0.000847604 | FDR correction BH |
| Dmel\CG12567~Q7PLS1 | VMAT~Control | VMAT | Control | 15.808539 | 0.169017738 | 16.629084 | 0.295490437 | 0.566228001 | -0.820545001 | -0.712131704 | 0.000305769 | 3.514606596 | Unpaired t-test | 0.040202647 | permutation FDR (250 perm) | 1 | 0.000543641 | FDR correction BH |
| Snx6~Q9VLQ9 | VMAT~Control | VMAT | Control | 16.39889348 | 0.265353683 | 17.2220718 | 0.21038506 | 0.565195418 | -0.823178325 | -0.714910371 | 0.00028663 | 3.542678909 | Unpaired t-test | 0.039594521 | permutation FDR (250 perm) | 1 | 0.000514273 | FDR correction BH |
| Septin4~Q7KUX3 | VMAT~Control | VMAT | Control | 16.59816334 | 0.077160562 | 17.42488265 | 0.109318533 | 0.563809889 | -0.826719313 | -0.780041418 | 7.69E-08 | 7.114223805 | Unpaired t-test | 0.028987109 | permutation FDR (250 perm) | 1 | 3.79E-07 | FDR correction BH |
| eIF4H1~X2JE34 | VMAT~Control | VMAT | Control | 16.45434375 | 0.165602923 | 17.28197545 | 0.224017702 | 0.563453437 | -0.827631702 | -0.735943503 | 5.76E-05 | 4.239647539 | Unpaired t-test | 0.035785518 | permutation FDR (250 perm) | 1 | 0.000119866 | FDR correction BH |
| RpLP1~M9PBK5 | VMAT~Control | VMAT | Control | 19.95669601 | 0.106468826 | 20.79218738 | 0.265438718 | 0.560392143 | -0.835491367 | -0.740748851 | 6.62E-05 | 4.179211657 | Unpaired t-test | 0.034946306 | permutation FDR (250 perm) | 1 | 0.000135134 | FDR correction BH |
| TM4SF~O76137 | VMAT~Control | VMAT | Control | 16.01410845 | 0.31425134 | 16.85323381 | 0.293352176 | 0.558982353 | -0.839125356 | -0.70381382 | 0.001410842 | 2.850521765 | Unpaired t-test | 0.04185597 | permutation FDR (250 perm) | 1 | 0.002195057 | FDR correction BH |
| Dmel\CG9265~Q9VIG6 | VMAT~Control | VMAT | Control | 13.57828627 | 0.267614537 | 14.42030847 | 0.803265129 | 0.557861078 | -0.842022197 | -0.610761618 | 0.050370013 | 1.297827933 | Unpaired t-test | 0.069492078 | permutation FDR (250 perm) | 1 | 0.062835724 | FDR correction BH |
| Dmel\CG33722~Q6NL44 | VMAT~Control | VMAT | Control | 15.48106009 | 0.088353518 | 16.32484781 | 0.193317427 | 0.557178801 | -0.843787727 | -0.770543173 | 4.68E-06 | 5.329442546 | Unpaired t-test | 0.030367985 | permutation FDR (250 perm) | 1 | 1.31E-05 | FDR correction BH |
| yki~A0A0C4DHD4 | VMAT~Control | VMAT | Control | 15.4802219 | 0.242840327 | 16.32468315 | 0.260018578 | 0.556918743 | -0.844461247 | -0.728542357 | 0.000343893 | 3.463576656 | Unpaired t-test | 0.037024703 | permutation FDR (250 perm) | 1 | 0.00060143 | FDR correction BH |
| psq~A4UZC9 | VMAT~Control | VMAT | Control | 13.39718327 | 0.326418252 | 14.24227861 | 0.999999561 | 0.556674021 | -0.845095338 | -0.574724504 | 0.102650543 | 0.988638749 | Unpaired t-test | 0.084147748 | permutation FDR (250 perm) | 1 | 0.122909145 | FDR correction BH |
| CG10517~Q961L8 | VMAT~Control | VMAT | Control | 15.90339412 | 0.109326251 | 16.75034466 | 0.102641995 | 0.55595864 | -0.846950535 | -0.79372079 | 1.80E-07 | 6.743703897 | Unpaired t-test | 0.027243542 | permutation FDR (250 perm) | 1 | 7.97E-07 | FDR correction BH |
| pins~Q9VB22 | VMAT~Control | VMAT | Control | 15.05985536 | 0.228378292 | 15.90740023 | 0.263844493 | 0.555729654 | -0.847544869 | -0.733133493 | 0.000288504 | 3.539847596 | Unpaired t-test | 0.036252632 | permutation FDR (250 perm) | 1 | 0.000516856 | FDR correction BH |
| Dmel\CG7789~Q9VAG9 | VMAT~Control | VMAT | Control | 18.85831191 | 0.07775034 | 19.70786933 | 0.088306394 | 0.554954957 | -0.849557416 | -0.807090154 | 1.72E-08 | 7.764449625 | Unpaired t-test | 0.025838337 | permutation FDR (250 perm) | 1 | 1.16E-07 | FDR correction BH |
| Pax~Q9VIX2 | VMAT~Control | VMAT | Control | 17.94179328 | 0.107455761 | 18.79173914 | 0.123274528 | 0.554805557 | -0.849945857 | -0.792021668 | 3.95E-07 | 6.403820161 | Unpaired t-test | 0.027457815 | permutation FDR (250 perm) | 1 | 1.52E-06 | FDR correction BH |
| Dmel\CG13917~Q9W0D3 | VMAT~Control | VMAT | Control | 15.0284835 | 0.221518667 | 15.87984009 | 0.073569138 | 0.554263309 | -0.851356588 | -0.770886296 | 9.94E-06 | 5.002581404 | Unpaired t-test | 0.030298068 | permutation FDR (250 perm) | 1 | 2.49E-05 | FDR correction BH |
| AMPdeam~X2JEY5 | VMAT~Control | VMAT | Control | 18.16972318 | 0.165095403 | 19.02124394 | 0.209575274 | 0.55420024 | -0.85152076 | -0.760752847 | 3.15E-05 | 4.501171637 | Unpaired t-test | 0.031761139 | permutation FDR (250 perm) | 1 | 7.00E-05 | FDR correction BH |
| eIF1~Q9VZS3 | VMAT~Control | VMAT | Control | 13.55655369 | 0.659108234 | 14.41362215 | 0.387072819 | 0.552073224 | -0.857068465 | -0.638729603 | 0.031057428 | 1.507834509 | Unpaired t-test | 0.059629764 | permutation FDR (250 perm) | 1 | 0.039841518 | FDR correction BH |
| msn~Q9W002 | VMAT~Control | VMAT | Control | 14.8098374 | 0.355226526 | 15.67759472 | 0.571240898 | 0.547998056 | -0.86775732 | -0.667078259 | 0.016259212 | 1.788900505 | Unpaired t-test | 0.050943636 | permutation FDR (250 perm) | 1 | 0.021653184 | FDR correction BH |
| Dmel\CG9391~Q7KTW5 | VMAT~Control | VMAT | Control | 19.50448251 | 0.13533939 | 20.37670331 | 0.101147648 | 0.546305255 | -0.872220794 | -0.810944708 | 4.20E-07 | 6.376275432 | Unpaired t-test | 0.025433858 | permutation FDR (250 perm) | 1 | 1.60E-06 | FDR correction BH |
| sif~A8JNL5 | VMAT~Control | VMAT | Control | 17.3367083 | 0.173562094 | 18.20918908 | 0.27298662 | 0.546206815 | -0.872480779 | -0.762212337 | 0.000126788 | 3.896923339 | Unpaired t-test | 0.031584559 | permutation FDR (250 perm) | 1 | 0.000243044 | FDR correction BH |
| CG1785~Q9W3C2 | VMAT~Control | VMAT | Control | 13.47176083 | 0.407171057 | 14.34564892 | 1.078350815 | 0.545674263 | -0.873888096 | -0.576638869 | 0.120883492 | 0.917633004 | Unpaired t-test | 0.083243243 | permutation FDR (250 perm) | 1 | 0.14328285 | FDR correction BH |
| capt~Q9VPX7 | VMAT~Control | VMAT | Control | 19.24889836 | 0.051076807 | 20.12322185 | 0.156262428 | 0.545509608 | -0.874323489 | -0.814444598 | 3.18E-07 | 6.497550465 | Unpaired t-test | 0.024980574 | permutation FDR (250 perm) | 1 | 1.26E-06 | FDR correction BH |
| tws~A0A0B4LGZ3 | VMAT~Control | VMAT | Control | 13.80139395 | 0.541402818 | 14.67662071 | 0.144747859 | 0.545168172 | -0.875226759 | -0.699830487 | 0.005802105 | 2.236414399 | Unpaired t-test | 0.04274567 | permutation FDR (250 perm) | 1 | 0.00830283 | FDR correction BH |
| Strn-Mlck~A1ZA73 | VMAT~Control | VMAT | Control | 19.30074607 | 0.135172986 | 20.1763286 | 0.075004722 | 0.545033748 | -0.875582533 | -0.818964374 | 1.76E-07 | 6.755386772 | Unpaired t-test | 0.024546043 | permutation FDR (250 perm) | 1 | 7.80E-07 | FDR correction BH |
| fmt~M9PEB1 | VMAT~Control | VMAT | Control | 16.44174014 | 0.192103788 | 17.31733489 | 0.125197865 | 0.545029133 | -0.875594748 | -0.794157141 | 6.63E-06 | 5.178795248 | Unpaired t-test | 0.027202952 | permutation FDR (250 perm) | 1 | 1.77E-05 | FDR correction BH |
| Abi~A0A0B4KH51 | VMAT~Control | VMAT | Control | 15.13436806 | 0.261849829 | 16.01136237 | 0.223825585 | 0.544500656 | -0.876994308 | -0.759924824 | 0.000200289 | 3.698343181 | Unpaired t-test | 0.03187322 | permutation FDR (250 perm) | 1 | 0.000368799 | FDR correction BH |
| Nadk1a~E1JH60 | VMAT~Control | VMAT | Control | 13.34567051 | 0.304161658 | 14.22451034 | 0.350926879 | 0.543804564 | -0.878839834 | -0.727706556 | 0.001739348 | 2.759613573 | Unpaired t-test | 0.037165599 | permutation FDR (250 perm) | 1 | 0.002670858 | FDR correction BH |
| CG34108~Q9VH10 | VMAT~Control | VMAT | Control | 17.07011532 | 0.183113573 | 17.94973081 | 0.219474851 | 0.543512272 | -0.879615483 | -0.779919991 | 4.29E-05 | 4.367402898 | Unpaired t-test | 0.029001842 | permutation FDR (250 perm) | 1 | 9.19E-05 | FDR correction BH |
| hts~Q02645-2 | VMAT~Control | VMAT | Control | 20.58146717 | 0.032251773 | 21.46187036 | 0.101468265 | 0.543215597 | -0.880403192 | -0.840388037 | 4.62E-09 | 8.335666326 | Unpaired t-test | 0.022287037 | permutation FDR (250 perm) | 1 | 4.12E-08 | FDR correction BH |
| Dmel\CG8441~Q7K1V1 | VMAT~Control | VMAT | Control | 13.38044455 | 0.499185584 | 14.26442553 | 0.851088638 | 0.541870127 | -0.883980981 | -0.613340343 | 0.072982458 | 1.136781516 | Unpaired t-test | 0.068554348 | permutation FDR (250 perm) | 1 | 0.088923761 | FDR correction BH |
| Non2~Q9VZY0 | VMAT~Control | VMAT | Control | 16.72215912 | 0.161479525 | 17.61339825 | 0.310845714 | 0.539150841 | -0.891239135 | -0.770532902 | 0.000201259 | 3.696245009 | Unpaired t-test | 0.030367985 | permutation FDR (250 perm) | 1 | 0.000370128 | FDR correction BH |
| AIMP2~Q8T060 | VMAT~Control | VMAT | Control | 15.83074631 | 0.151132737 | 16.72239116 | 0.182016857 | 0.538999241 | -0.891644853 | -0.806332551 | 7.44E-06 | 5.128268246 | Unpaired t-test | 0.025884527 | permutation FDR (250 perm) | 1 | 1.95E-05 | FDR correction BH |
| ATPCL~E2QCF1 | VMAT~Control | VMAT | Control | 19.834577 | 0.048160892 | 20.73227714 | 0.070041321 | 0.536741691 | -0.897700143 | -0.864824834 | 4.21E-10 | 9.376176292 | Unpaired t-test | 0.020474733 | permutation FDR (250 perm) | 1 | 6.34E-09 | FDR correction BH |
| CG17683~Q8SYS7-2 | VMAT~Control | VMAT | Control | 16.98485758 | 0.236502331 | 17.88640243 | 0.449645065 | 0.535313206 | -0.901544852 | -0.734631726 | 0.002652106 | 2.576409076 | Unpaired t-test | 0.036025653 | permutation FDR (250 perm) | 1 | 0.003956311 | FDR correction BH |
| Cdc37~Q24276 | VMAT~Control | VMAT | Control | 16.01289736 | 0.126783698 | 16.91458569 | 0.260116265 | 0.535259972 | -0.901688326 | -0.798371209 | 3.86E-05 | 4.41302246 | Unpaired t-test | 0.026730627 | permutation FDR (250 perm) | 1 | 8.39E-05 | FDR correction BH |
| cnk~Q7KNQ9 | VMAT~Control | VMAT | Control | 13.22394704 | 0.18462208 | 14.12676915 | 0.558250181 | 0.534839488 | -0.902822108 | -0.714848579 | 0.006402285 | 2.193665016 | Unpaired t-test | 0.039610959 | permutation FDR (250 perm) | 1 | 0.009117349 | FDR correction BH |
| CSN5~A0A0B4KHM2 | VMAT~Control | VMAT | Control | 13.76792242 | 0.320692222 | 14.67164423 | 0.297479032 | 0.534506052 | -0.903721811 | -0.755859962 | 0.000950926 | 3.02185309 | Unpaired t-test | 0.0324573 | permutation FDR (250 perm) | 1 | 0.001522776 | FDR correction BH |

| | | | | | | | | | | | | | | | | | | |
|---|---|---|---|---|---|---|---|---|---|---|---|---|---|---|---|---|---|---|
| pr~P48611 | VMAT~Control | VMAT | Control | 15.35065618 | 0.303474237 | 16.25548275 | 0.754816488 | 0.534096905 | -0.904826571 | -0.663447523 | 0.032156092 | 1.492736734 | Unpaired t-test | 0.051856429 | permutation FDR (250 perm) | 1 | 0.041228454 | FDR correction BH |
| eIF2Bgamma~A1ZA22 | VMAT~Control | VMAT | Control | 13.55884558 | 0.40410437 | 14.46425013 | 0.652336513 | 0.533882977 | -0.905404546 | -0.674078184 | 0.024805306 | 1.605455417 | Unpaired t-test | 0.048940427 | permutation FDR (250 perm) | 1 | 0.032242788 | FDR correction BH |
| CG3791~M9NFQ4 | VMAT~Control | VMAT | Control | 16.64793158 | 0.142604579 | 17.55796191 | 0.162785662 | 0.532173903 | -0.910030332 | -0.829726491 | 2.79E-06 | 5.554877424 | Unpaired t-test | 0.023357705 | permutation FDR (250 perm) | 1 | 8.23E-06 | FDR correction BH |
| Sf3b3~Q9W0M7 | VMAT~Control | VMAT | Control | 17.69310661 | 0.124068471 | 18.60379253 | 0.081828689 | 0.531932129 | -0.910685915 | -0.853928361 | 8.32E-08 | 7.079898823 | Unpaired t-test | 0.021241315 | permutation FDR (250 perm) | 1 | 4.07E-07 | FDR correction BH |
| CG31367~A0A0B4KFB8 | VMAT~Control | VMAT | Control | 19.14988851 | 0.202612785 | 20.0617888 | 0.235325696 | 0.531484572 | -0.911900282 | -0.800703305 | 6.34E-05 | 4.197867858 | Unpaired t-test | 0.02647786 | permutation FDR (250 perm) | 1 | 0.000130243 | FDR correction BH |
| Gfat1~A8Y5A0 | VMAT~Control | VMAT | Control | 16.52211634 | 0.137370604 | 17.43761564 | 0.081062655 | 0.530160358 | -0.915499296 | -0.8545424 | 1.55E-07 | 6.809947391 | Unpaired t-test | 0.021197405 | permutation FDR (250 perm) | 1 | 7.00E-07 | FDR correction BH |
| BcDNA:RE63412~M9PG40 | VMAT~Control | VMAT | Control | 17.77542311 | 0.143770755 | 18.69308547 | 0.117326233 | 0.529366072 | -0.917662363 | -0.847342648 | 6.28E-07 | 6.202100448 | Unpaired t-test | 0.02176378 | permutation FDR (250 perm) | 1 | 2.26E-06 | FDR correction BH |
| PAPLA1~Q9VLS7 | VMAT~Control | VMAT | Control | 16.7414846 | 0.064399449 | 17.66009197 | 0.157659057 | 0.529019437 | -0.918607366 | -0.853595369 | 2.79E-07 | 6.555171247 | Unpaired t-test | 0.021263548 | permutation FDR (250 perm) | 1 | 1.13E-06 | FDR correction BH |
| Synd~Q9VDI1 | VMAT~Control | VMAT | Control | 16.09454665 | 0.166365866 | 17.01345181 | 0.151915274 | 0.528910251 | -0.918905158 | -0.834796857 | 3.67E-06 | 5.4351534 | Unpaired t-test | 0.022876446 | permutation FDR (250 perm) | 1 | 1.05E-05 | FDR correction BH |
| anon-WO0172774.58~Q9VTZ7 | VMAT~Control | VMAT | Control | 17.40693688 | 0.099978635 | 18.3266872 | 0.105788609 | 0.528600494 | -0.919750322 | -0.863538094 | 6.20E-08 | 7.207424382 | Unpaired t-test | 0.020573018 | permutation FDR (250 perm) | 1 | 3.20E-07 | FDR correction BH |
| eRF3~Q9VK85 | VMAT~Control | VMAT | Control | 20.03775464 | 0.243132977 | 20.95947536 | 0.16484616 | 0.527879036 | -0.921720723 | -0.81469569 | 3.64E-05 | 4.438513684 | Unpaired t-test | 0.024950971 | permutation FDR (250 perm) | 1 | 7.98E-05 | FDR correction BH |
| Pu~P48596-3 | VMAT~Control | VMAT | Control | 18.72108994 | 0.324011876 | 19.64797038 | 0.460458497 | 0.52599448 | -0.926880436 | -0.740440579 | 0.004238817 | 2.372755352 | Unpaired t-test | 0.034990826 | permutation FDR (250 perm) | 1 | 0.006163172 | FDR correction BH |
| GH26~Q9VKI8 | VMAT~Control | VMAT | Control | 13.94169853 | 0.699422812 | 14.86858089 | 0.302281589 | 0.525993776 | -0.926882365 | -0.691314266 | 0.021556907 | 1.666413545 | Unpaired t-test | 0.044526412 | permutation FDR (250 perm) | 1 | 0.028301707 | FDR correction BH |
| Dmel\CG31251~Q8IN95 | VMAT~Control | VMAT | Control | 14.24475835 | 0.466665377 | 15.17181478 | 0.227542342 | 0.525930318 | -0.927056429 | -0.752367175 | 0.002548051 | 2.593791887 | Unpaired t-test | 0.033005966 | permutation FDR (250 perm) | 1 | 0.003815593 | FDR correction BH |
| Cypl~Q9W0Q2 | VMAT~Control | VMAT | Control | 15.60804608 | 0.176136234 | 16.54778198 | 0.223426279 | 0.521328308 | -0.939735894 | -0.833664843 | 2.35E-05 | 4.62829137 | Unpaired t-test | 0.023007867 | permutation FDR (250 perm) | 1 | 5.38E-05 | FDR correction BH |
| beta-Spec~Q00963 | VMAT~Control | VMAT | Control | 16.6317351 | 0.074904432 | 17.57294391 | 0.148123013 | 0.520796334 | -0.941208803 | -0.876169956 | 1.74E-07 | 6.76007193 | Unpaired t-test | 0.019721448 | permutation FDR (250 perm) | 1 | 7.73E-07 | FDR correction BH |
| Dll~A2VEF7 | VMAT~Control | VMAT | Control | 13.26373696 | 0.521686462 | 14.20845846 | 0.502491872 | 0.519529831 | -0.944721506 | -0.713573337 | 0.015394215 | 1.812642449 | Unpaired t-test | 0.03986484 | permutation FDR (250 perm) | 1 | 0.020636664 | FDR correction BH |
| Sbds~Q9VRY5 | VMAT~Control | VMAT | Control | 13.85532076 | 0.084483775 | 14.80405404 | 0.39638228 | 0.518087154 | -0.948733284 | -0.803160974 | 0.000381858 | 3.418097549 | Unpaired t-test | 0.026234534 | permutation FDR (250 perm) | 1 | 0.000658531 | FDR correction BH |
| cora~Q9V8R9 | VMAT~Control | VMAT | Control | 21.78722323 | 0.047083954 | 22.74325829 | 0.050407239 | 0.515471631 | -0.956035064 | -0.927426436 | 2.87E-11 | 10.54207919 | Unpaired t-test | 0.016855275 | permutation FDR (250 perm) | 1 | 8.14E-10 | FDR correction BH |
| GlyS~Q9VFC8 | VMAT~Control | VMAT | Control | 19.1354285 | 0.076704145 | 20.09277445 | 0.064910108 | 0.515003466 | -0.957345953 | -0.916175415 | 1.16E-09 | 8.936896378 | Unpaired t-test | 0.017421102 | permutation FDR (250 perm) | 1 | 1.35E-08 | FDR correction BH |
| GckIII~Q9VEN3 | VMAT~Control | VMAT | Control | 16.91388334 | 0.158300744 | 17.87191881 | 0.118265758 | 0.514757385 | -0.958035471 | -0.880248297 | 7.55E-07 | 6.122305629 | Unpaired t-test | 0.01945961 | permutation FDR (250 perm) | 1 | 2.63E-06 | FDR correction BH |
| Dmel\CG14411~Q9VY15 | VMAT~Control | VMAT | Control | 17.71904935 | 0.091615725 | 18.68019721 | 0.104581282 | 0.513648075 | -0.961147857 | -0.904883752 | 2.62E-08 | 7.5820426 | Unpaired t-test | 0.017974849 | permutation FDR (250 perm) | 1 | 1.61E-07 | FDR correction BH |
| Clamp~Q9V9N4 | VMAT~Control | VMAT | Control | 13.29654957 | 0.423701757 | 14.25803728 | 0.215312323 | 0.513527089 | -0.961487713 | -0.792948378 | 0.001102154 | 2.957757765 | Unpaired t-test | 0.027317658 | permutation FDR (250 perm) | 1 | 0.00175302 | FDR correction BH |
| Dmel\CG17002~A0A0B4JD24 | VMAT~Control | VMAT | Control | 13.29755878 | 0.142254155 | 14.25924178 | 0.221655378 | 0.513457579 | -0.961683006 | -0.860346495 | 9.85E-06 | 5.006745584 | Unpaired t-test | 0.020765879 | permutation FDR (250 perm) | 1 | 2.47E-05 | FDR correction BH |
| r2d2~Q9VLW8 | VMAT~Control | VMAT | Control | 15.41069111 | 0.396639198 | 16.37522709 | 0.325429715 | 0.512443205 | -0.964535978 | -0.784529 | 0.001817816 | 2.740450169 | Unpaired t-test | 0.028338858 | permutation FDR (250 perm) | 1 | 0.002782274 | FDR correction BH |
| Prx6a~Q9VQI7 | VMAT~Control | VMAT | Control | 16.20577712 | 0.144314863 | 17.177953 | 0.211131267 | 0.509736693 | -0.972175888 | -0.872399067 | 6.90E-06 | 5.161425816 | Unpaired t-test | 0.019992582 | permutation FDR (250 perm) | 1 | 1.83E-05 | FDR correction BH |
| Dmel\CG8547~Q0E980 | VMAT~Control | VMAT | Control | 17.18483918 | 0.152503163 | 18.15762579 | 0.217289486 | 0.509520954 | -0.972786616 | -0.869553422 | 9.54E-06 | 5.020439731 | Unpaired t-test | 0.020157626 | permutation FDR (250 perm) | 1 | 2.41E-05 | FDR correction BH |
| Mec2~Q9VWL0 | VMAT~Control | VMAT | Control | 15.63291234 | 0.077174134 | 16.6096465 | 0.596443642 | 0.508128695 | -0.976734158 | -0.76971155 | 0.004601083 | 2.337139931 | Unpaired t-test | 0.030456302 | permutation FDR (250 perm) | 1 | 0.006652917 | FDR correction BH |
| AdSS~Q9Y0Y2 | VMAT~Control | VMAT | Control | 15.06732628 | 0.421605475 | 16.05133745 | 0.302811899 | 0.505572128 | -0.984011163 | -0.798619237 | 0.001719523 | 2.764592043 | Unpaired t-test | 0.026697417 | permutation FDR (250 perm) | 1 | 0.002645593 | FDR correction BH |
| alpha-Spec~M9PBI5 | VMAT~Control | VMAT | Control | 18.45770482 | 0.064894979 | 19.44392407 | 0.115797806 | 0.504798926 | -0.986219254 | -0.930953995 | 1.31E-08 | 7.884255364 | Unpaired t-test | 0.016700607 | permutation FDR (250 perm) | 1 | 9.15E-08 | FDR correction BH |
| bel~A0A0B4KGU4 | VMAT~Control | VMAT | Control | 17.18245791 | 0.112034452 | 18.16947474 | 0.197662482 | 0.504519933 | -0.987016826 | -0.895977268 | 2.07E-06 | 5.683440775 | Unpaired t-test | 0.018449093 | permutation FDR (250 perm) | 1 | 6.40E-06 | FDR correction BH |
| BcDNA:RH17411~Q7JRC0 | VMAT~Control | VMAT | Control | 21.56264675 | 0.195303832 | 22.55036839 | 0.135434081 | 0.504273517 | -0.987721634 | -0.892824831 | 3.10E-06 | 5.508637051 | Unpaired t-test | 0.018664807 | permutation FDR (250 perm) | 1 | 9.07E-06 | FDR correction BH |
| Cka~M9PF42 | VMAT~Control | VMAT | Control | 16.48324337 | 0.158386492 | 17.47596074 | 0.069860122 | 0.502530346 | -0.992717378 | -0.921386673 | 1.56E-07 | 6.806368413 | Unpaired t-test | 0.017187675 | permutation FDR (250 perm) | 1 | 7.02E-07 | FDR correction BH |
| Trxr1~A0A6H2EFX8 | VMAT~Control | VMAT | Control | 14.14956993 | 0.442148211 | 15.14917286 | 0.252928986 | 0.500137631 | -0.999602937 | -0.814140363 | 0.001360216 | 2.866392044 | Unpaired t-test | 0.025012026 | permutation FDR (250 perm) | 1 | 0.002123308 | FDR correction BH |
| Rpn13~Q7K2G1 | VMAT~Control | VMAT | Control | 17.04828006 | 0.074427613 | 18.04907192 | 0.149264289 | 0.499725637 | -1.000791862 | -0.931323265 | 1.02E-07 | 6.993103489 | Unpaired t-test | 0.016693134 | permutation FDR (250 perm) | 1 | 4.86E-07 | FDR correction BH |
| Elp2~Q7K4B3 | VMAT~Control | VMAT | Control | 14.19516277 | 0.131567992 | 15.19596295 | 0.229285477 | 0.499722754 | -1.000800185 | -0.894992557 | 7.15E-06 | 5.145614901 | Unpaired t-test | 0.018519758 | permutation FDR (250 perm) | 1 | 1.89E-05 | FDR correction BH |
| Prps~M9PEP5 | VMAT~Control | VMAT | Control | 17.11609465 | 0.070023087 | 18.11788041 | 0.153064796 | 0.499381487 | -1.001785755 | -0.931654893 | 1.10E-07 | 6.959473716 | Unpaired t-test | 0.016689397 | permutation FDR (250 perm) | 1 | 5.20E-07 | FDR correction BH |
| Nadsyn~C5WLN1 | VMAT~Control | VMAT | Control | 17.15704975 | 0.092396676 | 18.15914545 | 0.044288655 | 0.499274215 | -1.002095694 | -0.958188829 | 8.95E-10 | 9.048374805 | Unpaired t-test | 0.015468165 | permutation FDR (250 perm) | 1 | 1.13E-08 | FDR correction BH |
| bark~M9NDE3-2 | VMAT~Control | VMAT | Control | 14.8348779 | 0.144285074 | 15.83967398 | 0.753440324 | 0.498340567 | -1.004796076 | -0.748132936 | 0.015071084 | 1.821855513 | Unpaired t-test | 0.033714548 | permutation FDR (250 perm) | 1 | 0.020238067 | FDR correction BH |
| Nt5b~Q9VWV2 | VMAT~Control | VMAT | Control | 15.35135003 | 0.144986983 | 16.3569591 | 0.157476834 | 0.498059819 | -1.005609069 | -0.917753474 | 1.01E-06 | 5.995519266 | Unpaired t-test | 0.017352007 | permutation FDR (250 perm) | 1 | 3.41E-06 | FDR correction BH |
| Sf3b5~Q9VHI4 | VMAT~Control | VMAT | Control | 15.72445119 | 0.248461963 | 16.73089905 | 0.278588282 | 0.497770329 | -1.006447858 | -0.862467546 | 0.000127138 | 3.895723489 | Unpaired t-test | 0.020623088 | permutation FDR (250 perm) | 1 | 0.000243518 | FDR correction BH |
| mbo~Q9GYU8 | VMAT~Control | VMAT | Control | 15.52060166 | 0.094291507 | 16.53417219 | 0.122012204 | 0.495318865 | -1.013570527 | -0.948183184 | 4.25E-08 | 7.371522768 | Unpaired t-test | 0.015872719 | permutation FDR (250 perm) | 1 | 2.39E-07 | FDR correction BH |
| dl~C7LA93 | VMAT~Control | VMAT | Control | 16.66924676 | 0.340497335 | 17.68294589 | 0.281668694 | 0.495274712 | -1.013699133 | -0.846425392 | 0.000444537 | 3.352091873 | Unpaired t-test | 0.021830477 | permutation FDR (250 perm) | 1 | 0.00075884 | FDR correction BH |
| pAbp~P21187 | VMAT~Control | VMAT | Control | 17.74186189 | 0.069630371 | 18.75910398 | 0.170347671 | 0.494059914 | -1.017242089 | -0.939888925 | 2.21E-07 | 6.655381034 | Unpaired t-test | 0.016304958 | permutation FDR (250 perm) | 1 | 9.43E-07 | FDR correction BH |
| ATPsynCF6~D1Z3A2 | VMAT~Control | VMAT | Control | 13.32012787 | 0.610193739 | 14.34082679 | 0.930101196 | 0.492877518 | -1.020698919 | -0.681611704 | 0.067311437 | 1.171911137 | Unpaired t-test | 0.047020928 | permutation FDR (250 perm) | 1 | 0.082526627 | FDR correction BH |
| HspB8~Q8IQW5 | VMAT~Control | VMAT | Control | 19.43103171 | 0.137749568 | 20.45188884 | 0.254085383 | 0.492823468 | -1.020857137 | -0.904009371 | 1.32E-05 | 4.880308732 | Unpaired t-test | 0.018028878 | permutation FDR (250 perm) | 1 | 3.20E-05 | FDR correction BH |
| Doa~A0A0B4K6R9 | VMAT~Control | VMAT | Control | 15.23419619 | 0.217564857 | 16.26186612 | 0.504900286 | 0.49050171 | -1.02766993 | -0.824861695 | 0.001888493 | 2.723884723 | Unpaired t-test | 0.023864877 | permutation FDR (250 perm) | 1 | 0.002877354 | FDR correction BH |
| Bruce~A0A0B4KH36 | VMAT~Control | VMAT | Control | 20.64804021 | 0.17278411 | 21.68037599 | 0.168266106 | 0.488917929 | -1.032335784 | -0.931829568 | 2.37E-06 | 5.624900642 | Unpaired t-test | 0.016676319 | permutation FDR (250 perm) | 1 | 7.13E-06 | FDR correction BH |
| e(y)3~M9PHL0 | VMAT~Control | VMAT | Control | 13.23254307 | 0.383138633 | 14.26626216 | 1.070711688 | 0.488449362 | -1.033719091 | -0.685230973 | 0.069502156 | 1.158001722 | Unpaired t-test | 0.046072793 | permutation FDR (250 perm) | 1 | 0.084991208 | FDR correction BH |
| mud~M9PJN0 | VMAT~Control | VMAT | Control | 14.17534138 | 1.721924957 | 15.20961541 | 1.869100541 | 0.488261512 | -1.034274034 | -0.484089374 | 0.384219986 | 0.415420049 | Unpaired t-test | 0.132806423 | permutation FDR (250 perm) | 1 | 0.421656805 | FDR correction BH |
| Zpr1~Q9W379 | VMAT~Control | VMAT | Control | 14.58429516 | 0.444454831 | 15.62198559 | 0.550264642 | 0.487106647 | -1.037690424 | -0.788319379 | 0.00828364 | 2.081778791 | Unpaired t-test | 0.027858065 | permutation FDR (250 perm) | 1 | 0.011652796 | FDR correction BH |
| Sms~Q9VTW1 | VMAT~Control | VMAT | Control | 13.29030635 | 0.407108227 | 14.33074451 | 1.012433706 | 0.486179795 | -1.040438157 | -0.699215425 | 0.058821704 | 1.230462396 | Unpaired t-test | 0.042891522 | permutation FDR (250 perm) | 1 | 0.072762108 | FDR correction BH |
| BEST:LD19244~Q9VID9 | VMAT~Control | VMAT | Control | 16.98296723 | 0.42038233 | 18.02934807 | 0.181671162 | 0.484181264 | -1.046380841 | -0.868506293 | 0.000457871 | 3.339256496 | Unpaired t-test | 0.020203987 | permutation FDR (250 perm) | 1 | 0.000778778 | FDR correction BH |
| tacc~A0A0B4JCV4 | VMAT~Control | VMAT | Control | 16.40787828 | 0.170031736 | 17.45475918 | 0.298997359 | 0.484013469 | -1.0468809 | -0.907313639 | 4.71E-05 | 4.32719529 | Unpaired t-test | 0.01784816 | permutation FDR (250 perm) | 1 | 9.97E-05 | FDR correction BH |
| goe~X2JFT2 | VMAT~Control | VMAT | Control | 14.66008307 | 0.263525286 | 15.70805847 | 0.338396672 | 0.48364641 | -1.047975404 | -0.879313193 | 0.000275537 | 3.559820669 | Unpaired t-test | 0.019511606 | permutation FDR (250 perm) | 1 | 0.000495882 | FDR correction BH |
| shot~A1Z9J1 | VMAT~Control | VMAT | Control | 13.30003052 | 0.469200367 | 14.35067131 | 0.549687633 | 0.482753694 | -1.050640797 | -0.794012462 | 0.008710156 | 2.059974079 | Unpaired t-test | 0.027217712 | permutation FDR (250 perm) | 1 | 0.012168372 | FDR correction BH |
| Fim~O61604 | VMAT~Control | VMAT | Control | 17.73305165 | 0.105458147 | 18.78662114 | 0.256772359 | 0.48177469 | -1.053569492 | -0.937222826 | 6.99E-06 | 5.155401197 | Unpaired t-test | 0.016431978 | permutation FDR (250 perm) | 1 | 1.85E-05 | FDR correction BH |
| Iswi~Q24368 | VMAT~Control | VMAT | Control | 13.85414989 | 0.419422936 | 14.90885581 | 0.35720655 | 0.481395337 | -1.054705926 | -0.846216319 | 0.001609276 | 2.793369333 | Unpaired t-test | 0.021839741 | permutation FDR (250 perm) | 1 | 0.00248409 | FDR correction BH |
| Dmel\CG2767~Q9VHX4 | VMAT~Control | VMAT | Control | 16.48979533 | 0.125407169 | 17.54737574 | 0.1765341 | 0.480437139 | -1.057580416 | -0.964205447 | 7.05E-07 | 6.151774773 | Unpaired t-test | 0.015208255 | permutation FDR (250 perm) | 1 | 2.48E-06 | FDR correction BH |
| vito~Q8SWS7 | VMAT~Control | VMAT | Control | 14.89414035 | 1.264731145 | 15.95326027 | 0.855338373 | 0.479924735 | -1.059119926 | -0.62937576 | 0.151919069 | 0.818387709 | Unpaired t-test | 0.062852221 | permutation FDR (250 perm) | 1 | 0.177053272 | FDR correction BH |
| Eip93F~Q9VD60 | VMAT~Control | VMAT | Control | 13.26053502 | 0.409510917 | 14.32052245 | 0.819154375 | 0.479636239 | -1.059987428 | -0.751996701 | 0.02703852 | 1.568017085 | Unpaired t-test | 0.033044516 | permutation FDR (250 perm) | 1 | 0.0349718 | FDR correction BH |
| Gmap~A0A4D6K5M0 | VMAT~Control | VMAT | Control | 15.91985273 | 0.160223219 | 16.98329932 | 0.104101023 | 0.478487589 | -1.063446592 | -0.979728779 | 2.07E-07 | 6.683520863 | Unpaired t-test | 0.014463128 | permutation FDR (250 perm) | 1 | 8.94E-07 | FDR correction BH |
| cact~M9PD95 | VMAT~Control | VMAT | Control | 14.96123425 | 0.214647679 | 16.02514004 | 0.197832795 | 0.478335311 | -1.063905798 | -0.941054755 | 1.00E-05 | 4.999703881 | Unpaired t-test | 0.016256314 | permutation FDR (250 perm) | 1 | 2.51E-05 | FDR correction BH |
| Msp300~M9MRD1 | VMAT~Control | VMAT | Control | 21.36650246 | 0.235164113 | 22.43144427 | 0.185434482 | 0.47799194 | -1.064941805 | -0.939158929 | 1.24E-05 | 4.905877407 | Unpaired t-test | 0.01632928 | permutation FDR (250 perm) | 1 | 3.04E-05 | FDR correction BH |
| Pak~Q9VI13 | VMAT~Control | VMAT | Control | 13.92969455 | 0.334185312 | 14.99639144 | 0.286902333 | 0.477410799 | -1.066696893 | -0.891161672 | 0.000294905 | 3.530317166 | Unpaired t-test | 0.018757787 | permutation FDR (250 perm) | 1 | 0.000527113 | FDR correction BH |
| homer~Q0E8S7 | VMAT~Control | VMAT | Control | 17.38741541 | 0.022929139 | 18.45415876 | 0.108649549 | 0.477395426 | -1.066743351 | -1.016275339 | 1.07E-09 | 8.972132963 | Unpaired t-test | 0.012966557 | permutation FDR (250 perm) | 1 | 1.27E-08 | FDR correction BH |
| Acn~Q9VJ12 | VMAT~Control | VMAT | Control | 15.21833834 | 0.281879632 | 16.28633303 | 0.41016792 | 0.476981528 | -1.067994698 | -0.873563084 | 0.000725373 | 3.139438741 | Unpaired t-test | 0.01990357 | permutation FDR (250 perm) | 1 | 0.001182498 | FDR correction BH |
| GluProRS~P28668 | VMAT~Control | VMAT | Control | 16.24587656 | 0.264788834 | 17.31517994 | 0.155477168 | 0.47654905 | -1.06930338 | -0.940194283 | 1.49E-05 | 4.826657584 | Unpaired t-test | 0.016299345 | permutation FDR (250 perm) | 1 | 3.58E-05 | FDR correction BH |
| Capr~E4NKG1 | VMAT~Control | VMAT | Control | 16.54570632 | 0.109206843 | 17.61521448 | 0.490712388 | 0.476481412 | -1.069508159 | -0.873194726 | 0.000771881 | 3.112449753 | Unpaired t-test | 0.019925823 | permutation FDR (250 perm) | 1 | 0.001256575 | FDR correction BH |
| Dmel\CG7611~E1JHX5 | VMAT~Control | VMAT | Control | 16.24240791 | 0.081410593 | 17.31254142 | 0.272198659 | 0.476274924 | -1.070133503 | -0.949492101 | 7.48E-06 | 5.125949707 | Unpaired t-test | 0.015827796 | permutation FDR (250 perm) | 1 | 1.95E-05 | FDR correction BH |
| Pex1~Q9VUC7 | VMAT~Control | VMAT | Control | 13.45022919 | 0.167691105 | 14.52232431 | 0.576243374 | 0.475627778 | -1.072095119 | -0.845238236 | 0.002540851 | 2.595020775 | Unpaired t-test | 0.021908291 | permutation FDR (250 perm) | 1 | 0.003807233 | FDR correction BH |
| Clic~Q9VY78 | VMAT~Control | VMAT | Control | 16.05891984 | 0.043432802 | 17.13123406 | 0.37038939 | 0.475555552 | -1.072314215 | -0.919038672 | 7.54E-05 | 4.122530658 | Unpaired t-test | 0.017284781 | permutation FDR (250 perm) | 1 | 0.000152257 | FDR correction BH |
| CG11016~B7YZX9 | VMAT~Control | VMAT | Control | 13.45544472 | 0.334236448 | 14.54523499 | 0.599904179 | 0.46982967 | -1.089790271 | -0.833736912 | 0.005281663 | 2.277229317 | Unpaired t-test | 0.023002314 | permutation FDR (250 perm) | 1 | 0.007604302 | FDR correction BH |
| CG17528~Q7PLI7 | VMAT~Control | VMAT | Control | 14.27132131 | 0.347248006 | 15.36415492 | 0.367642508 | 0.468839619 | -1.092833606 | -0.891264943 | 0.000689584 | 3.161413131 | Unpaired t-test | 0.018737331 | permutation FDR (250 perm) | 1 | 0.001130418 | FDR correction BH |
| CSN3~Q8SYG2 | VMAT~Control | VMAT | Control | 13.94477223 | 0.249771041 | 15.03803709 | 0.621389245 | 0.468699493 | -1.093264862 | -0.841294619 | 0.004460353 | 2.350630729 | Unpaired t-test | 0.022201019 | permutation FDR (250 perm) | 1 | 0.006464171 | FDR correction BH |
| Dmel\CG11444~Q9W4J4 | VMAT~Control | VMAT | Control | 16.14857949 | 0.122400337 | 17.24275893 | 0.116654276 | 0.468402461 | -1.094179439 | -1.017256968 | 4.94E-08 | 7.306452768 | Unpaired t-test | 0.012936411 | permutation FDR (250 perm) | 1 | 2.68E-07 | FDR correction BH |
| Gclm~Q9VCW6 | VMAT~Control | VMAT | Control | 16.03014268 | 0.12254329 | 17.12678162 | 0.096028828 | 0.467604611 | -1.09663894 | -1.025255262 | 2.19E-08 | 7.660547088 | Unpaired t-test | 0.012650636 | permutation FDR (250 perm) | 1 | 1.39E-07 | FDR correction BH |
| PSMG1~Q9VNI4 | VMAT~Control | VMAT | Control | 13.262661 | 0.142905162 | 14.36065945 | 1.148071338 | 0.467164177 | -1.097998446 | -0.723607367 | 0.059805186 | 1.223261157 | Unpaired t-test | 0.037912168 | permutation FDR (250 perm) | 1 | 0.073862228 | FDR correction BH |
| CG5009~Q7KML2 | VMAT~Control | VMAT | Control | 13.8316371 | 0.449534435 | 14.93166904 | 0.909235695 | 0.466506169 | -1.100031936 | -0.756760873 | 0.035749487 | 1.446730186 | Unpaired t-test | 0.032325069 | permutation FDR (250 perm) | 1 | 0.045538037 | FDR correction BH |
| me31B~P23128 | VMAT~Control | VMAT | Control | 19.07742948 | 0.124392073 | 20.17904937 | 0.074721935 | 0.465992976 | -1.101619885 | -1.034486935 | 1.06E-08 | 7.97502763 | Unpaired t-test | 0.012364839 | permutation FDR (250 perm) | 1 | 7.77E-08 | FDR correction BH |
| Dmel\CG42748~Q9V9N7 | VMAT~Control | VMAT | Control | 17.19487729 | 0.254209984 | 18.30016104 | 0.256357276 | 0.464811042 | -1.105283753 | -0.951635557 | 4.48E-05 | 4.348519699 | Unpaired t-test | 0.015741573 | permutation FDR (250 perm) | 1 | 9.56E-05 | FDR correction BH |
| sta~P38979 | VMAT~Control | VMAT | Control | 19.47177208 | 0.118803035 | 20.57811233 | 0.074326015 | 0.464470784 | -1.106340243 | -1.041093465 | 7.24E-09 | 8.139975887 | Unpaired t-test | 0.012128544 | permutation FDR (250 perm) | 1 | 5.84E-08 | FDR correction BH |
| coro~A0A0B4KEJ7 | VMAT~Control | VMAT | Control | 18.60454048 | 0.184926841 | 19.71586014 | 0.323153693 | 0.46287044 | -1.111319662 | -0.952688454 | 5.54E-05 | 4.25654021 | Unpaired t-test | 0.015687266 | permutation FDR (250 perm) | 1 | 0.000115506 | FDR correction BH |
| hts~A0A0B4K7A5 | VMAT~Control | VMAT | Control | 17.63872105 | 0.091439014 | 18.7507104 | 0.112970602 | 0.46265563 | -1.111989349 | -1.044123792 | 9.82E-09 | 8.007816791 | Unpaired t-test | 0.012005671 | permutation FDR (250 perm) | 1 | 7.39E-08 | FDR correction BH |
| Lpin~A0A0B4JD00 | VMAT~Control | VMAT | Control | 15.71481622 | 0.231175623 | 16.82756923 | 0.281423355 | 0.462410797 | -1.11275301 | -0.956898241 | 4.56E-05 | 4.341206547 | Unpaired t-test | 0.015520599 | permutation FDR (250 perm) | 1 | 9.71E-05 | FDR correction BH |
| CLIP-190~E1JHJ9 | VMAT~Control | VMAT | Control | 16.30939768 | 0.100716196 | 17.42859137 | 0.054014545 | 0.460351039 | -1.119193691 | -1.064772814 | 8.83E-10 | 9.053956088 | Unpaired t-test | 0.011257224 | permutation FDR (250 perm) | 1 | 1.12E-08 | FDR correction BH |
| Hex-A~Q9W330 | VMAT~Control | VMAT | Control | 19.38295828 | 0.193910034 | 20.50586124 | 0.266431843 | 0.459168963 | -1.122902966 | -0.97867703 | 1.80E-05 | 4.744906462 | Unpaired t-test | 0.014525129 | permutation FDR (250 perm) | 1 | 4.20E-05 | FDR correction BH |
| Jarid2~Q9VT00 | VMAT~Control | VMAT | Control | 13.6592067 | 0.339516535 | 14.78220576 | 0.654937951 | 0.459138381 | -1.122999058 | -0.844414995 | 0.00672665 | 2.172201138 | Unpaired t-test | 0.021978694 | permutation FDR (250 perm) | 1 | 0.009567695 | FDR correction BH |
| CG5543~Q9W1J3 | VMAT~Control | VMAT | Control | 13.45322225 | 0.324796974 | 14.57634267 | 1.12892228 | 0.459099757 | -1.123120425 | -0.736303838 | 0.058245165 | 1.234740117 | Unpaired t-test | 0.035710357 | permutation FDR (250 perm) | 1 | 0.072200695 | FDR correction BH |
| Prx2~X2JF59 | VMAT~Control | VMAT | Control | 19.26211919 | 0.132376123 | 20.38662435 | 0.176276236 | 0.458659315 | -1.124505156 | -1.023592486 | 4.70E-07 | 6.327794586 | Unpaired t-test | 0.012710985 | permutation FDR (250 perm) | 1 | 1.76E-06 | FDR correction BH |
| pod1~D6W4T8 | VMAT~Control | VMAT | Control | 16.53740628 | 0.072045974 | 17.66589177 | 0.140419614 | 0.457395637 | -1.128485492 | -1.054087114 | 1.89E-08 | 7.723388726 | Unpaired t-test | 0.011627423 | permutation FDR (250 perm) | 1 | 1.24E-07 | FDR correction BH |
| mib1~M9PI73 | VMAT~Control | VMAT | Control | 14.11369139 | 0.140485691 | 15.24425838 | 0.20028942 | 0.456736188 | -1.130566993 | -1.019071045 | 1.18E-06 | 5.929608262 | Unpaired t-test | 0.012870877 | permutation FDR (250 perm) | 1 | 3.89E-06 | FDR correction BH |
| Sec31~A1Z7J7 | VMAT~Control | VMAT | Control | 17.4218674 | 0.13142192 | 18.55401725 | 0.192087219 | 0.456235356 | -1.132149842 | -1.025418396 | 7.32E-07 | 6.135615451 | Unpaired t-test | 0.012644979 | permutation FDR (250 perm) | 1 | 2.56E-06 | FDR correction BH |
| GlyRS~Q9VUK8-2 | VMAT~Control | VMAT | Control | 15.51022759 | 0.083316176 | 16.6468021 | 0.061693546 | 0.45483825 | -1.136574511 | -1.086214265 | 2.91E-10 | 9.536007675 | Unpaired t-test | 0.010458472 | permutation FDR (250 perm) | 1 | 4.86E-09 | FDR correction BH |
| elB~Q9VJS8-2 | VMAT~Control | VMAT | Control | 13.52638009 | 0.465818819 | 14.66427232 | 0.224341928 | 0.454423002 | -1.13789223 | -0.924197965 | 0.000603759 | 3.219136692 | Unpaired t-test | 0.017043884 | permutation FDR (250 perm) | 1 | 0.000999471 | FDR correction BH |
| CG3737~M9NGH0 | VMAT~Control | VMAT | Control | 13.06594125 | 0.32261122 | 14.2041725 | 0.335183632 | 0.454316231 | -1.138231245 | -0.94220464 | 0.000272701 | 3.564312506 | Unpaired t-test | 0.016200187 | permutation FDR (250 perm) | 1 | 0.000491357 | FDR correction BH |
| Dp~A0A0B4KEQ0 | VMAT~Control | VMAT | Control | 13.22956524 | 0.345809777 | 14.36887513 | 0.795334148 | 0.453976686 | -1.139309886 | -0.820916753 | 0.014849014 | 1.828302384 | Unpaired t-test | 0.024331328 | permutation FDR (250 perm) | 1 | 0.019962638 | FDR correction BH |
| Cyt-c-p~X2J6D4 | VMAT~Control | VMAT | Control | 16.6762066 | 0.137650182 | 17.81660435 | 0.262995269 | 0.453634494 | -1.140397748 | -1.006750637 | 6.28E-06 | 5.202234804 | Unpaired t-test | 0.013333961 | permutation FDR (250 perm) | 1 | 1.69E-05 | FDR correction BH |
| cyst~Q9VIV0 | VMAT~Control | VMAT | Control | 14.91072344 | 0.264824873 | 16.05124648 | 0.25049923 | 0.4535951 | -1.140523041 | -0.980654115 | 3.73E-05 | 4.42792746 | Unpaired t-test | 0.014433067 | permutation FDR (250 perm) | 1 | 8.14E-05 | FDR correction BH |
| Dmel\CG10565~Q9VP77 | VMAT~Control | VMAT | Control | 13.48260565 | 0.377488586 | 14.62395383 | 0.306983467 | 0.453335744 | -1.141348179 | -0.93737967 | 0.000375919 | 3.424905893 | Unpaired t-test | 0.016428237 | permutation FDR (250 perm) | 1 | 0.000649716 | FDR correction BH |
| CalpB~M9NE73 | VMAT~Control | VMAT | Control | 14.93983825 | 0.419733511 | 16.09229632 | 0.108684638 | 0.449858109 | -1.152458066 | -0.965287555 | 0.000142472 | 3.846269854 | Unpaired t-test | 0.015153846 | permutation FDR (250 perm) | 1 | 0.000270249 | FDR correction BH |
| LUBEL~Q8IPJ3-4 | VMAT~Control | VMAT | Control | 13.55131978 | 0.414501984 | 14.7049245 | 0.183124311 | 0.449500704 | -1.153604719 | -0.959214545 | 0.00020038 | 3.698145598 | Unpaired t-test | 0.015428304 | permutation FDR (250 perm) | 1 | 0.000368799 | FDR correction BH |
| Dmel\CG8004~Q9VW58 | VMAT~Control | VMAT | Control | 16.77668982 | 0.116848214 | 17.93352038 | 0.184795935 | 0.44849675 | -1.156830564 | -1.053792461 | 3.34E-07 | 6.476580312 | Unpaired t-test | 0.011633097 | permutation FDR (250 perm) | 1 | 1.31E-06 | FDR correction BH |
| Chd64~Q9VZI1 | VMAT~Control | VMAT | Control | 19.93090063 | 0.091526213 | 21.09442529 | 0.175908036 | 0.446420547 | -1.163524664 | -1.068748114 | 1.26E-07 | 6.900833 | Unpaired t-test | 0.011080569 | permutation FDR (250 perm) | 1 | 5.87E-07 | FDR correction BH |
| Cat~P17336 | VMAT~Control | VMAT | Control | 18.52872224 | 0.134194621 | 19.70529313 | 0.126461983 | 0.442401784 | -1.176570893 | -1.086938551 | 5.65E-08 | 7.24796434 | Unpaired t-test | 0.010428097 | permutation FDR (250 perm) | 1 | 3.01E-07 | FDR correction BH |
| Stip1~Q9VPN5 | VMAT~Control | VMAT | Control | 15.94354328 | 0.245583108 | 17.12097979 | 0.416002507 | 0.442136424 | -1.177436505 | -0.968253739 | 0.000280835 | 3.551548537 | Unpaired t-test | 0.014988743 | permutation FDR (250 perm) | 1 | 0.000505031 | FDR correction BH |
| Spn~M9NDX8 | VMAT~Control | VMAT | Control | 16.24561493 | 0.124360199 | 17.43386137 | 0.142231381 | 0.43883593 | -1.188246441 | -1.095670128 | 6.49E-08 | 7.187976429 | Unpaired t-test | 0.010208076 | permutation FDR (250 perm) | 1 | 3.31E-07 | FDR correction BH |
| Tdrd3~Q9VUH8 | VMAT~Control | VMAT | Control | 13.79038617 | 0.338226118 | 14.98525905 | 0.245854017 | 0.436824935 | -1.194872884 | -1.006634356 | 7.94E-05 | 4.1004048 | Unpaired t-test | 0.01333773 | permutation FDR (250 perm) | 1 | 0.000159259 | FDR correction BH |
| Dhx15~Q7K3M5 | VMAT~Control | VMAT | Control | 15.48787934 | 0.08156557 | 16.68422749 | 0.133734106 | 0.436378477 | -1.196348147 | -1.118026048 | 9.99E-09 | 8.000272755 | Unpaired t-test | 0.009383444 | permutation FDR (250 perm) | 1 | 7.47E-08 | FDR correction BH |
| DCTN1-p150~P13496 | VMAT~Control | VMAT | Control | 15.2833551 | 0.113554947 | 16.48959277 | 0.082223394 | 0.43339738 | -1.206237662 | -1.135070613 | 3.14E-09 | 8.503741185 | Unpaired t-test | 0.008865174 | permutation FDR (250 perm) | 1 | 3.04E-08 | FDR correction BH |
| Abp1~Q9VU84 | VMAT~Control | VMAT | Control | 17.30949684 | 0.121181474 | 18.51817131 | 0.193057531 | 0.43266596 | -1.208674473 | -1.096863108 | 3.27E-07 | 6.485489503 | Unpaired t-test | 0.010166271 | permutation FDR (250 perm) | 1 | 1.29E-06 | FDR correction BH |
| Dmel\CG4593~Q9W3T2 | VMAT~Control | VMAT | Control | 13.94598394 | 0.418227974 | 15.1561057 | 0.285081311 | 0.432232134 | -1.210121764 | -0.986761932 | 0.00032541 | 3.48756936 | Unpaired t-test | 0.014193791 | permutation FDR (250 perm) | 1 | 0.000572507 | FDR correction BH |
| sw~Q24246 | VMAT~Control | VMAT | Control | 16.32884333 | 0.352905676 | 17.5417432 | 0.5145646 | 0.431400613 | -1.21289987 | -0.948288566 | 0.001451117 | 2.83829763 | Unpaired t-test | 0.015868975 | permutation FDR (250 perm) | 1 | 0.002251766 | FDR correction BH |
| mlt~Q7K549 | VMAT~Control | VMAT | Control | 17.68232372 | 0.085958452 | 18.8959983 | 0.236148424 | 0.431169017 | -1.213674584 | -1.091053618 | 7.82E-07 | 6.106577965 | Unpaired t-test | 0.010331591 | permutation FDR (250 perm) | 1 | 2.72E-06 | FDR correction BH |
| kraken~M9PBM0 | VMAT~Control | VMAT | Control | 14.07999345 | 1.152943545 | 15.29970861 | 1.530618225 | 0.429367481 | -1.219715164 | -0.656827411 | 0.18509886 | 0.732596256 | Unpaired t-test | 0.053840145 | permutation FDR (250 perm) | 1 | 0.212755233 | FDR correction BH |
| Pex19~Q8IP97 | VMAT~Control | VMAT | Control | 13.36317691 | 0.284072334 | 14.58472492 | 1.275675139 | 0.428822345 | -1.221548011 | -0.770949027 | 0.063129189 | 1.199769788 | Unpaired t-test | 0.030296228 | permutation FDR (250 perm) | 1 | 0.077682233 | FDR correction BH |
| NiPp1~Q9V7W9 | VMAT~Control | VMAT | Control | 13.34296769 | 0.444082012 | 14.56580533 | 0.319514191 | 0.428439191 | -1.222837639 | -0.982465413 | 0.00053887 | 3.268516372 | Unpaired t-test | 0.014363551 | permutation FDR (250 perm) | 1 | 0.000902204 | FDR correction BH |
| Uba1~Q8T0L3 | VMAT~Control | VMAT | Control | 16.32722499 | 0.073128748 | 17.55164222 | 0.076603763 | 0.427970353 | -1.224417234 | -1.169048382 | 1.72E-10 | 9.763428721 | Unpaired t-test | 0.007732569 | permutation FDR (250 perm) | 1 | 3.27E-09 | FDR correction BH |
| Uck~B7Z0P8 | VMAT~Control | VMAT | Control | 13.48312892 | 0.560556618 | 14.70865257 | 0.217831093 | 0.427642265 | -1.225523649 | -0.965776712 | 0.001047486 | 2.979851617 | Unpaired t-test | 0.015116323 | permutation FDR (250 perm) | 1 | 0.001672851 | FDR correction BH |
| mbf1~Q9XZ03 | VMAT~Control | VMAT | Control | 15.79185951 | 0.29666208 | 17.02049477 | 0.417556406 | 0.426720917 | -1.228635265 | -0.999647561 | 0.000317394 | 3.49840115 | Unpaired t-test | 0.013623352 | permutation FDR (250 perm) | 1 | 0.00056092 | FDR correction BH |
| Dmel\CG10435~Q9VHX1 | VMAT~Control | VMAT | Control | 15.00106284 | 1.464275047 | 16.231532 | 0.043699001 | 0.426178831 | -1.230469162 | -0.743425142 | 0.089790532 | 1.046769453 | Unpaired t-test | 0.034469022 | permutation FDR (250 perm) | 1 | 0.108226786 | FDR correction BH |
| Dmel\CG8545~A1Z909 | VMAT~Control | VMAT | Control | 13.8121244 | 0.303205173 | 15.04432326 | 0.292323142 | 0.425668175 | -1.232198864 | -1.036895434 | 6.54E-05 | 4.184478541 | Unpaired t-test | 0.012277883 | permutation FDR (250 perm) | 1 | 0.000133854 | FDR correction BH |
| Dmel\CG10863~Q9Y112 | VMAT~Control | VMAT | Control | 15.82412428 | 0.360187506 | 17.05665535 | 0.473592394 | 0.425570167 | -1.232531075 | -0.973492498 | 0.000933359 | 3.029951417 | Unpaired t-test | 0.014771831 | permutation FDR (250 perm) | 1 | 0.0014977 | FDR correction BH |
| cass~Q9V431 | VMAT~Control | VMAT | Control | 13.29569474 | 0.298007101 | 14.53061841 | 0.853252305 | 0.424864976 | -1.234923677 | -0.879456262 | 0.012141328 | 1.915733809 | Unpaired t-test | 0.019506035 | permutation FDR (250 perm) | 1 | 0.016558914 | FDR correction BH |
| Vrp1~Q8SX98 | VMAT~Control | VMAT | Control | 14.69242447 | 0.294918823 | 15.93817049 | 0.256765012 | 0.421689783 | -1.245746027 | -1.060322673 | 3.20E-05 | 4.494351213 | Unpaired t-test | 0.011422895 | permutation FDR (250 perm) | 1 | 7.09E-05 | FDR correction BH |
| Dmel\CG16812~Q9VK18 | VMAT~Control | VMAT | Control | 13.42010862 | 0.189653321 | 14.66844231 | 0.348945325 | 0.420934104 | -1.248333692 | -1.06005461 | 3.59E-05 | 4.444830468 | Unpaired t-test | 0.011443709 | permutation FDR (250 perm) | 1 | 7.89E-05 | FDR correction BH |
| viaf~Q8MR62 | VMAT~Control | VMAT | Control | 17.54635692 | 0.084955287 | 18.79855457 | 0.209412983 | 0.419808229 | -1.252197648 | -1.137259885 | 2.16E-07 | 6.665263169 | Unpaired t-test | 0.008817922 | permutation FDR (250 perm) | 1 | 9.28E-07 | FDR correction BH |
| Dmel\CG17816~A8JQV2 | VMAT~Control | VMAT | Control | 17.76766078 | 0.095102733 | 19.02069142 | 0.17344958 | 0.419565909 | -1.253030635 | -1.151191751 | 6.06E-08 | 7.217705814 | Unpaired t-test | 0.008339695 | permutation FDR (250 perm) | 1 | 3.15E-07 | FDR correction BH |
| Slu7~Q9VAQ7 | VMAT~Control | VMAT | Control | 16.38067321 | 0.241521134 | 17.63656318 | 0.267771754 | 0.418735177 | -1.255889974 | -1.08148314 | 1.49E-05 | 4.827059085 | Unpaired t-test | 0.010612903 | permutation FDR (250 perm) | 1 | 3.58E-05 | FDR correction BH |
| Gli~Q7KT70 | VMAT~Control | VMAT | Control | 14.87263884 | 0.272203774 | 16.13661584 | 0.449801298 | 0.416394522 | -1.263977006 | -1.023360544 | 0.00031195 | 3.505915644 | Unpaired t-test | 0.012716643 | permutation FDR (250 perm) | 1 | 0.000551712 | FDR correction BH |
| dnc~H8F4P8 | VMAT~Control | VMAT | Control | 14.7403161 | 0.304514078 | 16.00594482 | 0.128639678 | 0.415918074 | -1.265628716 | -1.102621831 | 6.47E-06 | 5.188956842 | Unpaired t-test | 0.009943943 | permutation FDR (250 perm) | 1 | 1.74E-05 | FDR correction BH |
| Ald1~C8VV14 | VMAT~Control | VMAT | Control | 17.29119407 | 0.148628844 | 18.55887504 | 0.203473284 | 0.415326845 | -1.26768097 | -1.139296907 | 5.35E-07 | 6.27170872 | Unpaired t-test | 0.008747735 | permutation FDR (250 perm) | 1 | 1.97E-06 | FDR correction BH |
| Clbn~Q9VBX1 | VMAT~Control | VMAT | Control | 17.69270426 | 0.425092937 | 18.96536553 | 0.467810425 | 0.413895576 | -1.272661266 | -0.992186294 | 0.00113946 | 2.94330082 | Unpaired t-test | 0.013914353 | permutation FDR (250 perm) | 1 | 0.001809912 | FDR correction BH |
| pont~Q9VH07 | VMAT~Control | VMAT | Control | 15.95797408 | 0.668391039 | 17.2310912 | 0.458725588 | 0.413764818 | -1.273117117 | -0.934370549 | 0.005616059 | 2.250568325 | Unpaired t-test | 0.01656662 | permutation FDR (250 perm) | 1 | 0.008054709 | FDR correction BH |
| Got2~Q9VQ61 | VMAT~Control | VMAT | Control | 16.13373443 | 0.06995762 | 17.41167985 | 0.189638633 | 0.412382376 | -1.277945418 | -1.17200156 | 6.17E-08 | 7.209758186 | Unpaired t-test | 0.007642789 | permutation FDR (250 perm) | 1 | 3.19E-07 | FDR correction BH |
| msps~Q9VEZ3 | VMAT~Control | VMAT | Control | 13.68815693 | 0.664490484 | 14.97607776 | 0.254785172 | 0.409540821 | -1.287920837 | -0.976981734 | 0.002335956 | 2.631535376 | Unpaired t-test | 0.01461907 | permutation FDR (250 perm) | 1 | 0.003515882 | FDR correction BH |
| shep~Q8MSV2-1 | VMAT~Control | VMAT | Control | 15.82831606 | 0.075567628 | 17.1175797 | 0.150098305 | 0.409159812 | -1.289263644 | -1.1991442 | 9.56E-09 | 8.019371187 | Unpaired t-test | 0.006961612 | permutation FDR (250 perm) | 1 | 7.28E-08 | FDR correction BH |
| eIF3d1~Q9VCK0 | VMAT~Control | VMAT | Control | 16.11079603 | 0.131902683 | 17.40315966 | 0.24786561 | 0.408281574 | -1.292363636 | -1.148188728 | 1.22E-06 | 5.913641849 | Unpaired t-test | 0.008444656 | permutation FDR (250 perm) | 1 | 4.00E-06 | FDR correction BH |
| eIF4G1~O61380 | VMAT~Control | VMAT | Control | 17.62527458 | 0.148701409 | 18.92305165 | 0.109865628 | 0.406752448 | -1.297777068 | -1.198667401 | 2.26E-08 | 7.64594714 | Unpaired t-test | 0.006971209 | permutation FDR (250 perm) | 1 | 1.43E-07 | FDR correction BH |
| cer~A1ZBK7 | VMAT~Control | VMAT | Control | 15.57233753 | 0.237624408 | 16.87256451 | 0.237807862 | 0.406062308 | -1.300226976 | -1.130293167 | 5.91E-06 | 5.22818136 | Unpaired t-test | 0.008998571 | permutation FDR (250 perm) | 1 | 1.61E-05 | FDR correction BH |
| Argk1~P48610-2 | VMAT~Control | VMAT | Control | 19.65387049 | 0.024889961 | 20.95447921 | 0.096336997 | 0.405954876 | -1.300608721 | -1.245199908 | 5.13E-11 | 10.29013443 | Unpaired t-test | 0.006005792 | permutation FDR (250 perm) | 1 | 1.33E-09 | FDR correction BH |
| Vha13~A0A0B4LHE7 | VMAT~Control | VMAT | Control | 15.68567754 | 0.048558552 | 16.98858864 | 0.296533957 | 0.405307535 | -1.302911098 | -1.148566404 | 2.11E-06 | 5.676034092 | Unpaired t-test | 0.008431298 | permutation FDR (250 perm) | 1 | 6.49E-06 | FDR correction BH |

| | | | | | | | | | | | | | | | | | | |
|---|---|---|---|---|---|---|---|---|---|---|---|---|---|---|---|---|---|---|
| Graf~X2JDY8-2 | VMAT~Control | VMAT | Control | 14.96054414 | 0.367336042 | 16.26511476 | 0.227495804 | 0.40484158 | -1.304570621 | -1.093309829 | 5.03E-05 | 4.298117509 | Unpaired t-test | 0.010265083 | permutation FDR (250 perm) | 1 | 0.000106366 | FDR correction BH |
| Dmel\CG2698~A0A0B4KFB0 | VMAT~Control | VMAT | Control | 13.19701224 | 0.321495401 | 14.50650437 | 0.410684597 | 0.403462885 | -1.309492129 | -1.061824586 | 0.000223251 | 3.651205946 | Unpaired t-test | 0.011369617 | permutation FDR (250 perm) | 1 | 0.000407706 | FDR correction BH |
| GstE11~Q7JVZ8 | VMAT~Control | VMAT | Control | 15.55995545 | 0.330712414 | 16.8708753 | 0.433021684 | 0.403063808 | -1.310919847 | -1.054072617 | 0.000310149 | 3.50842999 | Unpaired t-test | 0.011627423 | permutation FDR (250 perm) | 1 | 0.000549767 | FDR correction BH |
| mal~X2JFY7 | VMAT~Control | VMAT | Control | 15.97432035 | 0.068615807 | 17.28867131 | 0.133333469 | 0.402106355 | -1.314350958 | -1.231748567 | 2.62E-09 | 8.582470647 | Unpaired t-test | 0.006267246 | permutation FDR (250 perm) | 1 | 2.64E-08 | FDR correction BH |
| Sap130~M9PI47 | VMAT~Control | VMAT | Control | 13.52933863 | 0.276485131 | 14.84763074 | 0.520876164 | 0.401009382 | -1.318292105 | -1.043179063 | 0.000538344 | 3.26894044 | Unpaired t-test | 0.012041588 | permutation FDR (250 perm) | 1 | 0.000901965 | FDR correction BH |
| cN-IIIB~Q9W197 | VMAT~Control | VMAT | Control | 13.48578609 | 0.249841724 | 14.80466445 | 0.368549188 | 0.400846461 | -1.318878358 | -1.099869329 | 5.90E-05 | 4.228989908 | Unpaired t-test | 0.010044656 | permutation FDR (250 perm) | 1 | 0.000122509 | FDR correction BH |
| Dmel\CG3548~Q9W140 | VMAT~Control | VMAT | Control | 13.69944768 | 0.455844125 | 15.02002421 | 0.167558002 | 0.400374908 | -1.320576534 | -1.084933744 | 0.000118772 | 3.925285107 | Unpaired t-test | 0.010507831 | permutation FDR (250 perm) | 1 | 0.000228984 | FDR correction BH |
| l(2)37Cb~Q9VIZ3 | VMAT~Control | VMAT | Control | 13.50619261 | 0.195854008 | 14.82749209 | 0.311650801 | 0.400174327 | -1.321299479 | -1.134540817 | 1.14E-05 | 4.941752708 | Unpaired t-test | 0.008880419 | permutation FDR (250 perm) | 1 | 2.83E-05 | FDR correction BH |
| Trl~Q2PDY2 | VMAT~Control | VMAT | Control | 13.2598571 | 0.283836508 | 14.58137149 | 1.369789455 | 0.400114721 | -1.321514386 | -0.8129387 | 0.060797243 | 1.216116114 | Unpaired t-test | 0.025143386 | permutation FDR (250 perm) | 1 | 0.075008758 | FDR correction BH |
| Dmel\CG8516~Q9VH63 | VMAT~Control | VMAT | Control | 14.21588522 | 0.23785977 | 15.53749778 | 0.293758174 | 0.400087495 | -1.321612559 | -1.130511825 | 1.44E-05 | 4.841935528 | Unpaired t-test | 0.008990948 | permutation FDR (250 perm) | 1 | 3.46E-05 | FDR correction BH |
| Dp1~Q7KN75 | VMAT~Control | VMAT | Control | 18.74763907 | 0.130158873 | 20.07495031 | 0.199334515 | 0.398510255 | -1.327311244 | -1.199594602 | 2.04E-07 | 6.690661972 | Unpaired t-test | 0.006957774 | permutation FDR (250 perm) | 1 | 8.84E-07 | FDR correction BH |
| BcDNA:RE20991~Q8SXU5 | VMAT~Control | VMAT | Control | 14.62507901 | 0.222569218 | 15.9621407 | 0.248871949 | 0.395826005 | -1.337061698 | -1.163355669 | 4.33E-06 | 5.36349257 | Unpaired t-test | 0.007893079 | permutation FDR (250 perm) | 1 | 1.22E-05 | FDR correction BH |
| Pcf11~A0A0B4LFF9 | VMAT~Control | VMAT | Control | 13.89607396 | 0.086265492 | 15.23774824 | 0.438474279 | 0.39456249 | -1.341674285 | -1.118201066 | 5.28E-05 | 4.277683343 | Unpaired t-test | 0.009370776 | permutation FDR (250 perm) | 1 | 0.000110893 | FDR correction BH |
| CCT2~Q9W392 | VMAT~Control | VMAT | Control | 17.95225293 | 0.137954354 | 19.30063899 | 0.138185488 | 0.392731153 | -1.348386052 | -1.240096925 | 2.65E-08 | 7.577549176 | Unpaired t-test | 0.00611288 | permutation FDR (250 perm) | 1 | 1.62E-07 | FDR correction BH |
| Drp1~Q9VQE0 | VMAT~Control | VMAT | Control | 17.92475904 | 0.067845473 | 19.27949719 | 0.180269301 | 0.391005781 | -1.354738156 | -1.247296697 | 2.22E-08 | 7.654341013 | Unpaired t-test | 0.00596139 | permutation FDR (250 perm) | 1 | 1.41E-07 | FDR correction BH |
| CG-6428~Q9W4N6 | VMAT~Control | VMAT | Control | 13.54678654 | 0.111888637 | 14.90381699 | 0.320158559 | 0.390385007 | -1.357030447 | -1.17831426 | 4.36E-06 | 5.360239247 | Unpaired t-test | 0.007488751 | permutation FDR (250 perm) | 1 | 1.23E-05 | FDR correction BH |
| Zw~P12646 | VMAT~Control | VMAT | Control | 17.52424696 | 0.252838312 | 18.8842755 | 0.418913157 | 0.389574584 | -1.360028538 | -1.115854886 | 9.95E-05 | 4.002366512 | Unpaired t-test | 0.009440533 | permutation FDR (250 perm) | 1 | 0.000195753 | FDR correction BH |
| Wdr62~Q0E8U7 | VMAT~Control | VMAT | Control | 14.94652555 | 0.092217752 | 16.30778386 | 0.239454875 | 0.389242647 | -1.361258309 | -1.221128553 | 3.26E-07 | 6.487327628 | Unpaired t-test | 0.006484089 | permutation FDR (250 perm) | 1 | 1.29E-06 | FDR correction BH |
| bbg~Q15K24 | VMAT~Control | VMAT | Control | 13.43442304 | 0.264302834 | 14.79880578 | 0.47653862 | 0.38840058 | -1.364382738 | -1.097036481 | 0.000228116 | 3.641843709 | Unpaired t-test | 0.01015867 | permutation FDR (250 perm) | 1 | 0.000415946 | FDR correction BH |
| eIF2Bbeta~O76863 | VMAT~Control | VMAT | Control | 13.57470152 | 0.558612363 | 14.94724037 | 0.269575616 | 0.386210999 | -1.372538844 | -1.074489125 | 0.000580169 | 3.236445531 | Unpaired t-test | 0.01085389 | permutation FDR (250 perm) | 1 | 0.00096313 | FDR correction BH |
| Idh~Q7KUB0 | VMAT~Control | VMAT | Control | 19.13579513 | 0.050804208 | 20.50863132 | 0.067836908 | 0.386131409 | -1.372836184 | -1.322702736 | 6.12E-12 | 11.2135491 | Unpaired t-test | 0.004312167 | permutation FDR (250 perm) | 1 | 2.88E-10 | FDR correction BH |
| holn1~Q9VKV5 | VMAT~Control | VMAT | Control | 13.64345319 | 0.415997125 | 15.02498248 | 0.361658942 | 0.38381173 | -1.381529291 | -1.108312483 | 0.00022636 | 3.645201216 | Unpaired t-test | 0.009719448 | permutation FDR (250 perm) | 1 | 0.000413062 | FDR correction BH |
| hts~Q02645-3 | VMAT~Control | VMAT | Control | 16.84459285 | 0.103933293 | 18.2277073 | 0.612889438 | 0.383390248 | -1.383114453 | -1.082244431 | 0.00055742 | 3.253817666 | Unpaired t-test | 0.010565465 | permutation FDR (250 perm) | 1 | 0.000927982 | FDR correction BH |
| CG3760~Q8MMC4-2 | VMAT~Control | VMAT | Control | 17.62941407 | 0.04887062 | 19.01407111 | 0.131520857 | 0.382980532 | -1.384657036 | -1.302903455 | 8.19E-10 | 9.086843179 | Unpaired t-test | 0.004689255 | permutation FDR (250 perm) | 1 | 1.05E-08 | FDR correction BH |
| Dmel\CG11586~Q9VZF0 | VMAT~Control | VMAT | Control | 13.49240692 | 0.315126501 | 14.87989738 | 0.254810543 | 0.382229105 | -1.387490458 | -1.174608641 | 1.73E-05 | 4.762619728 | Unpaired t-test | 0.00757935 | permutation FDR (250 perm) | 1 | 4.05E-05 | FDR correction BH |
| SCAR~Q9VKM2 | VMAT~Control | VMAT | Control | 15.22446722 | 0.211271365 | 16.62289208 | 0.679855327 | 0.379343083 | -1.398424865 | -1.060712102 | 0.001351262 | 2.869260579 | Unpaired t-test | 0.011403974 | permutation FDR (250 perm) | 1 | 0.002112131 | FDR correction BH |
| CSN7~Q9V4S8 | VMAT~Control | VMAT | Control | 15.56545711 | 0.174207466 | 16.96820557 | 0.133699082 | 0.378207937 | -1.402748456 | -1.277307157 | 5.59E-08 | 7.252522571 | Unpaired t-test | 0.005233446 | permutation FDR (250 perm) | 1 | 3.00E-07 | FDR correction BH |
| Atx2~Q8SWR8 | VMAT~Control | VMAT | Control | 16.24957377 | 0.081401371 | 17.65838418 | 0.145789032 | 0.376622107 | -1.408810409 | -1.310919519 | 3.79E-09 | 8.420923739 | Unpaired t-test | 0.004532946 | permutation FDR (250 perm) | 1 | 3.49E-08 | FDR correction BH |
| Dmel\CG34120~E2QD66 | VMAT~Control | VMAT | Control | 13.8252389 | 1.388591747 | 15.23590733 | 2.200521781 | 0.376137375 | -1.410668429 | -0.651983519 | 0.253271514 | 0.596413654 | Unpaired t-test | 0.055355132 | permutation FDR (250 perm) | 1 | 0.286911041 | FDR correction BH |
| Dmel\CG9643~Q9VQJ8 | VMAT~Control | VMAT | Control | 14.67088707 | 0.192843905 | 16.08353309 | 0.650237149 | 0.375622131 | -1.412646029 | -1.083887654 | 0.000897903 | 3.046770577 | Unpaired t-test | 0.010542003 | permutation FDR (250 perm) | 1 | 0.001444746 | FDR correction BH |
| eIF3f1~Q9VN50 | VMAT~Control | VMAT | Control | 13.30647369 | 0.347615511 | 14.73500557 | 1.208075034 | 0.371508758 | -1.428531875 | -0.914442465 | 0.029311969 | 1.532955006 | Unpaired t-test | 0.017508166 | permutation FDR (250 perm) | 1 | 0.037705123 | FDR correction BH |
| Pi4KIIIalpha~Q9W4X4 | VMAT~Control | VMAT | Control | 13.42574454 | 0.616975898 | 14.86505819 | 0.365833376 | 0.36874269 | -1.439313644 | -1.089746656 | 0.001166418 | 2.9331458 | Unpaired t-test | 0.010363895 | permutation FDR (250 perm) | 1 | 0.001848988 | FDR correction BH |
| axo~M9PE65 | VMAT~Control | VMAT | Control | 18.10025684 | 0.17118423 | 19.54104186 | 0.168661329 | 0.36836681 | -1.440785017 | -1.300967957 | 1.02E-07 | 6.989771911 | Unpaired t-test | 0.004739593 | permutation FDR (250 perm) | 1 | 4.88E-07 | FDR correction BH |
| Dmel\CG11550~Q9V9V4 | VMAT~Control | VMAT | Control | 13.4737215 | 0.177746869 | 14.91825409 | 0.184210512 | 0.367411174 | -1.444532591 | -1.296150311 | 1.82E-07 | 6.7401973 | Unpaired t-test | 0.004839865 | permutation FDR (250 perm) | 1 | 8.02E-07 | FDR correction BH |
| Mad1~Q7KND8 | VMAT~Control | VMAT | Control | 13.3146918 | 0.297106339 | 14.76279517 | 0.356459046 | 0.366502928 | -1.44810337 | -1.199231615 | 3.82E-05 | 4.418402406 | Unpaired t-test | 0.006961612 | permutation FDR (250 perm) | 1 | 8.30E-05 | FDR correction BH |
| stmA~A0A0B4LFU5 | VMAT~Control | VMAT | Control | 15.1162038 | 0.282694198 | 16.56446429 | 0.112229323 | 0.366463016 | -1.448260487 | -1.274851256 | 8.92E-07 | 6.049596812 | Unpaired t-test | 0.005297245 | permutation FDR (250 perm) | 1 | 3.04E-06 | FDR correction BH |
| MetRS~A1ZBE9 | VMAT~Control | VMAT | Control | 16.0211715 | 0.205837854 | 17.4736525 | 0.166467785 | 0.365392518 | -1.452481001 | -1.298725164 | 2.37E-07 | 6.624945647 | Unpaired t-test | 0.004787996 | permutation FDR (250 perm) | 1 | 9.93E-07 | FDR correction BH |
| ntc~Q8SX86 | VMAT~Control | VMAT | Control | 13.58412633 | 0.472582926 | 15.03864853 | 0.144145421 | 0.364875909 | -1.454522196 | -1.1912957 | 6.21E-05 | 4.206838886 | Unpaired t-test | 0.007165868 | permutation FDR (250 perm) | 1 | 0.000127803 | FDR correction BH |
| Pex3~Q9VUL8 | VMAT~Control | VMAT | Control | 13.8492867 | 0.558411737 | 15.31721961 | 0.328234077 | 0.361499884 | -1.467932911 | -1.13821826 | 0.000486608 | 3.31282066 | Unpaired t-test | 0.008800763 | permutation FDR (250 perm) | 1 | 0.000821718 | FDR correction BH |
| DENR~Q9VX98 | VMAT~Control | VMAT | Control | 13.57828067 | 0.725788175 | 15.04720443 | 0.156974906 | 0.361251688 | -1.468923764 | -1.102723415 | 0.001287468 | 2.890263521 | Unpaired t-test | 0.009938242 | permutation FDR (250 perm) | 1 | 0.002019121 | FDR correction BH |
| GstE12~Q9XYZ9 | VMAT~Control | VMAT | Control | 17.6919258 | 0.118459369 | 19.1643218 | 0.141143673 | 0.360383284 | -1.472396001 | -1.360298554 | 6.44E-09 | 8.190936021 | Unpaired t-test | 0.003688867 | permutation FDR (250 perm) | 1 | 5.36E-08 | FDR correction BH |
| sle~Q8INM3-2 | VMAT~Control | VMAT | Control | 13.50154466 | 0.268706593 | 14.9869666 | 0.17168431 | 0.357144066 | -1.485421945 | -1.300032772 | 1.09E-06 | 5.961676002 | Unpaired t-test | 0.004757018 | permutation FDR (250 perm) | 1 | 3.65E-06 | FDR correction BH |
| aPKC~A1Z9X0 | VMAT~Control | VMAT | Control | 13.03162906 | 0.271141669 | 14.51936538 | 1.036157429 | 0.356571595 | -1.487736316 | -1.005916133 | 0.011136098 | 1.953266954 | Unpaired t-test | 0.013367876 | permutation FDR (250 perm) | 1 | 0.015263225 | FDR correction BH |
| lolal~E1JGK7 | VMAT~Control | VMAT | Control | 13.26460626 | 0.081968266 | 14.76184174 | 1.232865333 | 0.354231525 | -1.497235483 | -0.96435854 | 0.021949492 | 1.658575527 | Unpaired t-test | 0.01520075 | permutation FDR (250 perm) | 1 | 0.028768989 | FDR correction BH |
| Nacalpha~A0A0B4LEY6 | VMAT~Control | VMAT | Control | 14.86713964 | 0.51007905 | 16.36473063 | 0.187987137 | 0.354144248 | -1.497590984 | -1.2047102 | 0.000106901 | 3.97101833 | Unpaired t-test | 0.006847816 | permutation FDR (250 perm) | 1 | 0.000209181 | FDR correction BH |
| DCP1~Q9W1H5 | VMAT~Control | VMAT | Control | 15.95363041 | 0.148769055 | 17.4512421 | 0.064951435 | 0.354139166 | -1.497611686 | -1.396249509 | 1.58E-09 | 8.800000787 | Unpaired t-test | 0.003121286 | permutation FDR (250 perm) | 1 | 1.77E-08 | FDR correction BH |
| Mfe2~X2JFD6 | VMAT~Control | VMAT | Control | 13.2739822 | 0.303984848 | 14.77390628 | 0.159078987 | 0.353571996 | -1.499924081 | -1.300396539 | 1.96E-06 | 5.70854358 | Unpaired t-test | 0.00475121 | permutation FDR (250 perm) | 1 | 6.08E-06 | FDR correction BH |
| Drep2~Q7K304 | VMAT~Control | VMAT | Control | 16.53530461 | 0.362103771 | 18.04119022 | 0.395919532 | 0.352113974 | -1.505885611 | -1.214476359 | 9.19E-05 | 4.03665673 | Unpaired t-test | 0.00662747 | permutation FDR (250 perm) | 1 | 0.00018211 | FDR correction BH |
| MEP-1~Q0E8J0 | VMAT~Control | VMAT | Control | 13.53718433 | 0.387224769 | 15.04592584 | 0.225722733 | 0.351417634 | -1.508741508 | -1.256817066 | 2.00E-05 | 4.698947062 | Unpaired t-test | 0.005732496 | permutation FDR (250 perm) | 1 | 4.63E-05 | FDR correction BH |
| eIF2D~Q9NG60 | VMAT~Control | VMAT | Control | 13.79913171 | 0.437835289 | 15.31606576 | 0.482286026 | 0.349427714 | -1.516934058 | -1.174727444 | 0.000397067 | 3.401136133 | Unpaired t-test | 0.007577438 | permutation FDR (250 perm) | 1 | 0.000683257 | FDR correction BH |
| hyd~A0A0B4LGZ6 | VMAT~Control | VMAT | Control | 13.32209374 | 0.262702782 | 14.83965173 | 1.489392285 | 0.349276627 | -1.51755799 | -0.905270713 | 0.048695965 | 1.312507023 | Unpaired t-test | 0.017959944 | permutation FDR (250 perm) | 1 | 0.060876422 | FDR correction BH |
| Moe~M9NG50 | VMAT~Control | VMAT | Control | 14.88183687 | 0.283815254 | 16.41421016 | 0.450724857 | 0.345708197 | -1.532373282 | -1.237578123 | 7.51E-05 | 4.124450773 | Unpaired t-test | 0.006174626 | permutation FDR (250 perm) | 1 | 0.000151715 | FDR correction BH |
| Mcm5~Q9VGW6 | VMAT~Control | VMAT | Control | 12.85634001 | 0.394358082 | 14.38977006 | 0.957232059 | 0.34545506 | -1.53343005 | -1.048146023 | 0.007852824 | 2.104974144 | Unpaired t-test | 0.011844991 | permutation FDR (250 perm) | 1 | 0.011082462 | FDR correction BH |
| Mtap~A0A0B4LFH0 | VMAT~Control | VMAT | Control | 17.59926909 | 0.285271085 | 19.13515109 | 0.375946932 | 0.344868437 | -1.535881999 | -1.268220873 | 2.67E-05 | 4.573231414 | Unpaired t-test | 0.005453842 | permutation FDR (250 perm) | 1 | 6.00E-05 | FDR correction BH |
| SmD2~Q9VI10 | VMAT~Control | VMAT | Control | 13.19226022 | 0.294128727 | 14.73219503 | 1.019257171 | 0.343900995 | -1.539934806 | -1.044430526 | 0.008781311 | 2.056440637 | Unpaired t-test | 0.011990548 | permutation FDR (250 perm) | 1 | 0.012260502 | FDR correction BH |
| Tlk~B7Z120 | VMAT~Control | VMAT | Control | 13.41826457 | 0.203943434 | 14.95825481 | 0.395456037 | 0.343887782 | -1.539990236 | -1.284409845 | 1.57E-05 | 4.803468016 | Unpaired t-test | 0.005079826 | permutation FDR (250 perm) | 1 | 3.75E-05 | FDR correction BH |
| cher~B7Z0L2 | VMAT~Control | VMAT | Control | 13.54116762 | 0.30041572 | 15.08230139 | 0.246889426 | 0.343615311 | -1.541133771 | -1.312833566 | 4.75E-06 | 5.323067481 | Unpaired t-test | 0.004507752 | permutation FDR (250 perm) | 1 | 1.32E-05 | FDR correction BH |
| Mlc2~P18432 | VMAT~Control | VMAT | Control | 17.98699966 | 0.231262299 | 19.5323569 | 0.2260961 | 0.342610854 | -1.54535724 | -1.350083048 | 8.64E-07 | 6.063582232 | Unpaired t-test | 0.00381933 | permutation FDR (250 perm) | 1 | 2.96E-06 | FDR correction BH |
| Map60~Q7K180 | VMAT~Control | VMAT | Control | 13.49159905 | 0.423076836 | 15.03822059 | 0.149453046 | 0.342310741 | -1.546621533 | -1.288138498 | 1.63E-05 | 4.788019547 | Unpaired t-test | 0.005006289 | permutation FDR (250 perm) | 1 | 3.86E-05 | FDR correction BH |
| Dmel\CG8635~A0A0B4LEV3 | VMAT~Control | VMAT | Control | 17.18656326 | 0.176198039 | 18.73497174 | 0.301056166 | 0.341887013 | -1.548408475 | -1.339453279 | 1.70E-06 | 5.769045897 | Unpaired t-test | 0.003990291 | permutation FDR (250 perm) | 1 | 5.36E-06 | FDR correction BH |
| Pfrx~Q9VWH7 | VMAT~Control | VMAT | Control | 17.63286315 | 0.071455929 | 19.18146538 | 0.181122288 | 0.341841101 | -1.548602225 | -1.424557333 | 6.74E-09 | 8.171279596 | Unpaired t-test | 0.002711632 | permutation FDR (250 perm) | 1 | 5.54E-08 | FDR correction BH |
| Taf5~P49846 | VMAT~Control | VMAT | Control | 13.73431766 | 0.510726929 | 15.28356781 | 0.389742043 | 0.341687612 | -1.54925015 | -1.203476841 | 0.000304761 | 3.516041058 | Unpaired t-test | 0.00687278 | permutation FDR (250 perm) | 1 | 0.000542668 | FDR correction BH |
| Ythdf~Q9VBZ5-3 | VMAT~Control | VMAT | Control | 13.11739239 | 0.403680586 | 14.68193861 | 0.620671 | 0.338084032 | -1.564546218 | -1.1753639 | 0.000810206 | 3.091404419 | Unpaired t-test | 0.007564054 | permutation FDR (250 perm) | 1 | 0.001315328 | FDR correction BH |
| Phf5a~Q9VMC8 | VMAT~Control | VMAT | Control | 13.43933743 | 0.528954061 | 15.00922668 | 0.188275448 | 0.336834252 | -1.569889246 | -1.254813561 | 9.48E-05 | 4.023282873 | Unpaired t-test | 0.005778851 | permutation FDR (250 perm) | 1 | 0.000187332 | FDR correction BH |
| rhea~Q9VSL8 | VMAT~Control | VMAT | Control | 17.48745115 | 0.126649311 | 19.05794067 | 0.116402667 | 0.336694131 | -1.57048952 | -1.458304955 | 1.76E-09 | 8.755605088 | Unpaired t-test | 0.002371387 | permutation FDR (250 perm) | 1 | 1.92E-08 | FDR correction BH |
| Sip1~Q7K5M6 | VMAT~Control | VMAT | Control | 17.46390421 | 0.109171088 | 19.03719511 | 0.079719435 | 0.336040983 | -1.573290901 | -1.483601075 | 1.61E-10 | 9.791906998 | Unpaired t-test | 0.002081572 | permutation FDR (250 perm) | 1 | 3.14E-09 | FDR correction BH |
| yps~M9NFR5 | VMAT~Control | VMAT | Control | 13.41506513 | 0.349187971 | 14.99678835 | 0.186159509 | 0.334082608 | -1.581723217 | -1.343897028 | 4.40E-06 | 5.356198115 | Unpaired t-test | 0.003914563 | permutation FDR (250 perm) | 1 | 1.23E-05 | FDR correction BH |
| Nost~D5SHR1 | VMAT~Control | VMAT | Control | 13.57708265 | 0.534270079 | 15.16497402 | 0.238162064 | 0.332657308 | -1.587891367 | -1.258635693 | 0.000120377 | 3.919455578 | Unpaired t-test | 0.005682279 | permutation FDR (250 perm) | 1 | 0.000231889 | FDR correction BH |
| twin~Q9VCB6 | VMAT~Control | VMAT | Control | 13.91804034 | 0.540217008 | 15.51160254 | 0.313285101 | 0.33135229 | -1.593562207 | -1.245672775 | 0.000196679 | 3.706241304 | Unpaired t-test | 0.006 | permutation FDR (250 perm) | 1 | 0.000362839 | FDR correction BH |
| CG18490~Q9VTH2 | VMAT~Control | VMAT | Control | 13.49759266 | 0.26291726 | 15.09875334 | 0.484733697 | 0.329611691 | -1.601160678 | -1.284407781 | 6.96E-05 | 4.157352107 | Unpaired t-test | 0.005079826 | permutation FDR (250 perm) | 1 | 0.000141619 | FDR correction BH |
| Nopp140~M9MSM4 | VMAT~Control | VMAT | Control | 13.41655265 | 0.283327963 | 15.02424492 | 0.47954338 | 0.328122796 | -1.607692269 | -1.287087736 | 7.31E-05 | 4.136155018 | Unpaired t-test | 0.005023706 | permutation FDR (250 perm) | 1 | 0.000147808 | FDR correction BH |
| Taf3~Q9V4D4 | VMAT~Control | VMAT | Control | 14.98252133 | 0.348712265 | 16.59728481 | 0.307044268 | 0.326518473 | -1.614763481 | -1.33696088 | 1.52E-05 | 4.819087318 | Unpaired t-test | 0.004036893 | permutation FDR (250 perm) | 1 | 3.63E-05 | FDR correction BH |
| Dmel\CG17816~A8JQV3 | VMAT~Control | VMAT | Control | 18.72372402 | 0.100420458 | 20.34513958 | 0.102803358 | 0.325016404 | -1.621415559 | -1.523501015 | 2.19E-10 | 9.65883214 | Unpaired t-test | 0.001751854 | permutation FDR (250 perm) | 1 | 3.83E-09 | FDR correction BH |
| tau~A0A0B4LIT4 | VMAT~Control | VMAT | Control | 16.46980433 | 0.097595059 | 18.09248363 | 0.158892606 | 0.324731829 | -1.622679294 | -1.497775825 | 2.81E-09 | 8.55184749 | Unpaired t-test | 0.001975406 | permutation FDR (250 perm) | 1 | 2.79E-08 | FDR correction BH |
| CG5995~Q9I7H8 | VMAT~Control | VMAT | Control | 13.28843442 | 0.443944832 | 14.9221562 | 0.066439231 | 0.322255798 | -1.633721777 | -1.360585368 | 1.01E-05 | 4.994294451 | Unpaired t-test | 0.003688867 | permutation FDR (250 perm) | 1 | 2.54E-05 | FDR correction BH |
| BEST:CK00496~Q9VRP2 | VMAT~Control | VMAT | Control | 15.97670787 | 0.096010721 | 17.62158773 | 0.082197028 | 0.319773022 | -1.644879865 | -1.556879896 | 5.35E-11 | 10.27133564 | Unpaired t-test | 0.001459593 | permutation FDR (250 perm) | 1 | 1.37E-09 | FDR correction BH |
| Hip1~Q86BS5 | VMAT~Control | VMAT | Control | 13.75712778 | 1.107116981 | 15.40323866 | 0.522940697 | 0.319500285 | -1.646110878 | -1.063673273 | 0.013203038 | 1.879326117 | Unpaired t-test | 0.011301467 | permutation FDR (250 perm) | 1 | 0.017941371 | FDR correction BH |
| Cisd2~Q9VAM6 | VMAT~Control | VMAT | Control | 18.12651062 | 0.11021175 | 19.77466434 | 0.112378773 | 0.319048196 | -1.648153717 | -1.539765555 | 4.58E-10 | 9.339590624 | Unpaired t-test | 0.001592483 | permutation FDR (250 perm) | 1 | 6.82E-09 | FDR correction BH |
| eIF3j~A0A0B4LF26 | VMAT~Control | VMAT | Control | 16.07990716 | 0.167724466 | 17.72976927 | 0.120689389 | 0.318670614 | -1.649862108 | -1.510296605 | 6.49E-09 | 8.187715001 | Unpaired t-test | 0.001875862 | permutation FDR (250 perm) | 1 | 5.38E-08 | FDR correction BH |
| CYC4~Q7K231 | VMAT~Control | VMAT | Control | 13.67679468 | 0.37809823 | 15.32718803 | 0.506046728 | 0.318553292 | -1.650393348 | -1.286852749 | 0.000163364 | 3.786842417 | Unpaired t-test | 0.005025641 | permutation FDR (250 perm) | 1 | 0.000306667 | FDR correction BH |
| Tango4~Q9VYQ9 | VMAT~Control | VMAT | Control | 13.29954359 | 0.30949484 | 14.9611605 | 0.41068858 | 0.316084694 | -1.661616917 | -1.350930906 | 2.84E-05 | 4.546567943 | Unpaired t-test | 0.003807674 | permutation FDR (250 perm) | 1 | 6.35E-05 | FDR correction BH |
| eIF5B~Q9VZP5 | VMAT~Control | VMAT | Control | 15.52387032 | 0.242383223 | 17.18968984 | 0.265318575 | 0.315165273 | -1.665819519 | -1.435168997 | 1.14E-06 | 5.94191098 | Unpaired t-test | 0.002604697 | permutation FDR (250 perm) | 1 | 3.81E-06 | FDR correction BH |
| Golgin84~Q8SZ63 | VMAT~Control | VMAT | Control | 14.54742929 | 1.125942722 | 16.21609263 | 0.140201677 | 0.314544635 | -1.668663341 | -1.106962346 | 0.008170863 | 2.087732088 | Unpaired t-test | 0.009753919 | permutation FDR (250 perm) | 1 | 0.011517491 | FDR correction BH |
| eRF1~Q9VPH7 | VMAT~Control | VMAT | Control | 16.79222696 | 0.177498308 | 18.46933224 | 0.161291914 | 0.312709449 | -1.677105281 | -1.514648091 | 2.34E-08 | 7.630043036 | Unpaired t-test | 0.001843273 | permutation FDR (250 perm) | 1 | 1.48E-07 | FDR correction BH |
| RanBPM~A0A0B4KFL0 | VMAT~Control | VMAT | Control | 14.30680491 | 0.336264307 | 15.98657609 | 0.619707916 | 0.312132138 | -1.679771186 | -1.277088516 | 0.00033424 | 3.475941659 | Unpaired t-test | 0.005237313 | permutation FDR (250 perm) | 1 | 0.000585505 | FDR correction BH |
| tud~A0A0B4LG05 | VMAT~Control | VMAT | Control | 14.36741417 | 0.376073715 | 16.04756985 | 0.463696536 | 0.312048963 | -1.680155676 | -1.326089361 | 8.99E-05 | 4.046102137 | Unpaired t-test | 0.004265761 | permutation FDR (250 perm) | 1 | 0.000178372 | FDR correction BH |
| Trmt6~Q8MS69 | VMAT~Control | VMAT | Control | 13.533512 | 0.329554347 | 15.22197335 | 0.418981166 | 0.310257643 | -1.688461345 | -1.363431565 | 3.36E-05 | 4.473135009 | Unpaired t-test | 0.003642197 | permutation FDR (250 perm) | 1 | 7.42E-05 | FDR correction BH |
| vlc~A1Z6H4 | VMAT~Control | VMAT | Control | 15.72010599 | 0.289439437 | 17.41057266 | 0.244019546 | 0.309826689 | -1.690466667 | -1.445702215 | 1.61E-06 | 5.792649888 | Unpaired t-test | 0.002484581 | permutation FDR (250 perm) | 1 | 5.11E-06 | FDR correction BH |
| mub~Q95SZ9 | VMAT~Control | VMAT | Control | 17.10552784 | 0.085583905 | 18.79662556 | 0.241673823 | 0.309691198 | -1.691097715 | -1.517146627 | 4.11E-08 | 7.386075161 | Unpaired t-test | 0.001821411 | permutation FDR (250 perm) | 1 | 2.33E-07 | FDR correction BH |
| Pym~P82804 | VMAT~Control | VMAT | Control | 13.39925152 | 0.353200515 | 15.0927989 | 0.147755324 | 0.309165795 | -1.69354738 | -1.445968056 | 1.76E-06 | 5.7551514 | Unpaired t-test | 0.002482624 | permutation FDR (250 perm) | 1 | 5.52E-06 | FDR correction BH |
| St2~Q9VHH0 | VMAT~Control | VMAT | Control | 14.10479476 | 0.924563527 | 15.80465655 | 0.781137752 | 0.30781559 | -1.699861792 | -1.102879927 | 0.010503575 | 1.978662861 | Unpaired t-test | 0.009932542 | permutation FDR (250 perm) | 1 | 0.014466546 | FDR correction BH |
| gw~Q8SY33-2 | VMAT~Control | VMAT | Control | 16.52521056 | 0.130570821 | 18.23174583 | 0.115541816 | 0.306395016 | -1.706535266 | -1.583097036 | 8.88E-10 | 9.051762496 | Unpaired t-test | 0.001275536 | permutation FDR (250 perm) | 1 | 1.12E-08 | FDR correction BH |
| grsm~Q9V3D8 | VMAT~Control | VMAT | Control | 18.69569395 | 0.090465616 | 20.40512291 | 0.146253221 | 0.305781077 | -1.709428967 | -1.587349446 | 7.63E-10 | 9.11760037 | Unpaired t-test | 0.001266833 | permutation FDR (250 perm) | 1 | 1.00E-08 | FDR correction BH |
| IMP~Q9VUW4 | VMAT~Control | VMAT | Control | 14.56497301 | 0.266019773 | 16.27720005 | 0.342148659 | 0.305188597 | -1.712227037 | -1.434241428 | 4.89E-06 | 5.310680795 | Unpaired t-test | 0.002624266 | permutation FDR (250 perm) | 1 | 1.35E-05 | FDR correction BH |
| RhoGAP19D~E1JJS2 | VMAT~Control | VMAT | Control | 17.51790849 | 0.321605241 | 19.23243336 | 0.321093402 | 0.3047029 | -1.714524862 | -1.424924348 | 7.38E-06 | 5.132186248 | Unpaired t-test | 0.002711002 | permutation FDR (250 perm) | 1 | 1.94E-05 | FDR correction BH |
| fliI~Q24020 | VMAT~Control | VMAT | Control | 13.55393324 | 0.35452853 | 15.28212184 | 0.423403932 | 0.301830689 | -1.728188595 | -1.385914614 | 3.73E-05 | 4.428753577 | Unpaired t-test | 0.00328335 | permutation FDR (250 perm) | 1 | 8.13E-05 | FDR correction BH |
| chico~M9PCQ8 | VMAT~Control | VMAT | Control | 15.8012484 | 0.288372303 | 17.53077571 | 0.169714008 | 0.301550742 | -1.729527311 | -1.504407054 | 4.15E-07 | 6.381438087 | Unpaired t-test | 0.001921182 | permutation FDR (250 perm) | 1 | 1.59E-06 | FDR correction BH |
| l(2)k01209~A1ZAS3 | VMAT~Control | VMAT | Control | 13.8402503 | 0.385259517 | 15.57472808 | 0.345596742 | 0.300517772 | -1.734477785 | -1.408475983 | 2.08E-05 | 4.682416469 | Unpaired t-test | 0.00294439 | permutation FDR (250 perm) | 1 | 4.79E-05 | FDR correction BH |
| Amph~Q7KLE5 | VMAT~Control | VMAT | Control | 19.19068954 | 0.069038375 | 20.92517986 | 0.09317824 | 0.30051516 | -1.734490322 | -1.648970926 | 1.35E-11 | 10.86981993 | Unpaired t-test | 0.00092678 | permutation FDR (250 perm) | 1 | 4.60E-10 | FDR correction BH |
| eIF3a~Q9VN25 | VMAT~Control | VMAT | Control | 16.63704419 | 0.162181408 | 18.37339606 | 0.225186226 | 0.300127648 | -1.736351866 | -1.544651199 | 6.81E-08 | 7.166551753 | Unpaired t-test | 0.001547749 | permutation FDR (250 perm) | 1 | 3.44E-07 | FDR correction BH |
| eIF4B~Q7PLL3 | VMAT~Control | VMAT | Control | 17.30123175 | 0.109754827 | 19.04062529 | 0.155652985 | 0.299495547 | -1.73939354 | -1.602869037 | 1.75E-09 | 8.756926099 | Unpaired t-test | 0.001144855 | permutation FDR (250 perm) | 1 | 1.92E-08 | FDR correction BH |
| Dmel\CG7706~E2QCZ5 | VMAT~Control | VMAT | Control | 13.02011524 | 0.288839345 | 14.76725501 | 0.235574521 | 0.297891783 | -1.747139764 | -1.497521698 | 1.03E-06 | 5.986643142 | Unpaired t-test | 0.001975406 | permutation FDR (250 perm) | 1 | 3.47E-06 | FDR correction BH |
| wupA~M9NFC0 | VMAT~Control | VMAT | Control | 20.14358494 | 0.355199854 | 21.89686728 | 0.212200492 | 0.296626142 | -1.753282344 | -1.479515258 | 2.60E-06 | 5.585143505 | Unpaired t-test | 0.002129666 | permutation FDR (250 perm) | 1 | 7.75E-06 | FDR correction BH |
| CG31291~Q6NR11 | VMAT~Control | VMAT | Control | 13.62420588 | 0.346587319 | 15.38542914 | 0.303126313 | 0.294997931 | -1.761223261 | -1.46048507 | 6.53E-06 | 5.185362449 | Unpaired t-test | 0.002345098 | permutation FDR (250 perm) | 1 | 1.75E-05 | FDR correction BH |
| Dmel\CG3638~A0A0S0X7W8 | VMAT~Control | VMAT | Control | 15.41559866 | 0.20700165 | 17.17835612 | 0.185951905 | 0.294684388 | -1.762757464 | -1.567674505 | 6.05E-08 | 7.218025234 | Unpaired t-test | 0.001373075 | permutation FDR (250 perm) | 1 | 3.15E-07 | FDR correction BH |
| bif~M9PGW8 | VMAT~Control | VMAT | Control | 15.2927782 | 0.407618142 | 17.0561806 | 0.619863985 | 0.294552682 | -1.763402405 | -1.324096152 | 0.000340167 | 3.468308166 | Unpaired t-test | 0.004290839 | permutation FDR (250 perm) | 1 | 0.000595355 | FDR correction BH |
| sgll~Q7KSW3 | VMAT~Control | VMAT | Control | 14.50277258 | 0.430243728 | 16.27487012 | 0.836885213 | 0.292782751 | -1.772097535 | -1.247227606 | 0.00179717 | 2.745410959 | Unpaired t-test | 0.00596139 | permutation FDR (250 perm) | 1 | 0.002754256 | FDR correction BH |
| Elp1~A0A0B4KFR6 | VMAT~Control | VMAT | Control | 13.8568622 | 0.550896792 | 15.6393017 | 0.091112239 | 0.290691441 | -1.782439499 | -1.42627642 | 3.15E-05 | 4.501707762 | Unpaired t-test | 0.002699267 | permutation FDR (250 perm) | 1 | 7.00E-05 | FDR correction BH |
| Nup54~Q9V6B9 | VMAT~Control | VMAT | Control | 13.68983594 | 0.523162662 | 15.47531121 | 0.356874962 | 0.2900804 | -1.785475274 | -1.391405731 | 8.86E-05 | 4.052597616 | Unpaired t-test | 0.003187926 | permutation FDR (250 perm) | 1 | 0.000176289 | FDR correction BH |
| CG32355~M9NF47 | VMAT~Control | VMAT | Control | 17.02561455 | 0.18515884 | 18.81420297 | 0.122579363 | 0.28945512 | -1.78858842 | -1.62701435 | 5.96E-09 | 8.224647727 | Unpaired t-test | 0.001026513 | permutation FDR (250 perm) | 1 | 4.99E-08 | FDR correction BH |
| bic~Q7KM15 | VMAT~Control | VMAT | Control | 15.24577053 | 0.513101619 | 17.0346087 | 0.518578492 | 0.289405016 | -1.788838168 | -1.348793592 | 0.000268103 | 3.571697845 | Unpaired t-test | 0.003838835 | permutation FDR (250 perm) | 1 | 0.000484727 | FDR correction BH |
| Taldo~Q9W1G0 | VMAT~Control | VMAT | Control | 14.34021713 | 0.835558831 | 16.12933331 | 0.270598116 | 0.289349252 | -1.78911618 | -1.284564517 | 0.001050179 | 2.978736741 | Unpaired t-test | 0.005077891 | permutation FDR (250 perm) | 1 | 0.001676014 | FDR correction BH |
| Pur-alpha~F3YDK4 | VMAT~Control | VMAT | Control | 15.4483306 | 0.260767223 | 17.24245381 | 0.230347242 | 0.288346776 | -1.794123204 | -1.552544663 | 4.25E-07 | 6.371702265 | Unpaired t-test | 0.00148662 | permutation FDR (250 perm) | 1 | 1.62E-06 | FDR correction BH |
| RanBP3~Q9VCI7 | VMAT~Control | VMAT | Control | 14.56317662 | 1.220854577 | 16.36236139 | 0.14972853 | 0.287336909 | -1.799184774 | -1.160709296 | 0.008418793 | 2.074750176 | Unpaired t-test | 0.007986635 | permutation FDR (250 perm) | 1 | 0.011831545 | FDR correction BH |
| HDAC6~Q9XYX1 | VMAT~Control | VMAT | Control | 13.48945407 | 0.299045188 | 15.28888035 | 0.267393139 | 0.287288814 | -1.799426273 | -1.525709474 | 1.55E-06 | 5.810684077 | Unpaired t-test | 0.001716263 | permutation FDR (250 perm) | 1 | 4.93E-06 | FDR correction BH |
| SF1~Q9VEJ1 | VMAT~Control | VMAT | Control | 14.1631786 | 0.905057152 | 15.96504051 | 0.734921744 | 0.286804206 | -1.801861912 | -1.184352296 | 0.006165146 | 2.210056669 | Unpaired t-test | 0.007363375 | permutation FDR (250 perm) | 1 | 0.00879028 | FDR correction BH |
| Dmel\CG13901~Q9W0M5 | VMAT~Control | VMAT | Control | 15.10203112 | 0.183431373 | 16.90878684 | 0.050319741 | 0.285832977 | -1.806755724 | -1.665114927 | 1.19E-09 | 8.924098142 | Unpaired t-test | 0.000849398 | permutation FDR (250 perm) | 1 | 1.38E-08 | FDR correction BH |
| RtGEF~Q9VIN1 | VMAT~Control | VMAT | Control | 13.64366975 | 0.618955844 | 15.45672419 | 0.454901762 | 0.284587767 | -1.813054445 | -1.349477292 | 0.000358765 | 3.445190334 | Unpaired t-test | 0.003825158 | permutation FDR (250 perm) | 1 | 0.000624192 | FDR correction BH |
| E(bx)~E1JHV7 | VMAT~Control | VMAT | Control | 13.69415073 | 0.478823818 | 15.5076811 | 0.195952089 | 0.284493901 | -1.813530371 | -1.472769831 | 1.41E-05 | 4.851461725 | Unpaired t-test | 0.002197349 | permutation FDR (250 perm) | 1 | 3.40E-05 | FDR correction BH |
| Dmel\CG14442~Q7YU88 | VMAT~Control | VMAT | Control | 13.02226577 | 0.281011021 | 14.83651882 | 0.294884021 | 0.284351427 | -1.814253053 | -1.534684444 | 1.65E-06 | 5.782493845 | Unpaired t-test | 0.001645895 | permutation FDR (250 perm) | 1 | 5.22E-06 | FDR correction BH |
| Tm1~P06754 | VMAT~Control | VMAT | Control | 19.19813605 | 0.161422246 | 21.01495731 | 0.235187532 | 0.28384569 | -1.816821259 | -1.611271603 | 5.75E-08 | 7.240364544 | Unpaired t-test | 0.001090909 | permutation FDR (250 perm) | 1 | 3.05E-07 | FDR correction BH |
| Agxt~Q9W3Z3 | VMAT~Control | VMAT | Control | 15.29404962 | 0.453329129 | 17.11835012 | 0.381718113 | 0.282377981 | -1.824300498 | -1.44209563 | 4.28E-05 | 4.368468026 | Unpaired t-test | 0.002510763 | permutation FDR (250 perm) | 1 | 9.18E-05 | FDR correction BH |
| Mtr4~Q9Y134 | VMAT~Control | VMAT | Control | 14.61215903 | 0.186782052 | 16.44764621 | 0.210398807 | 0.280196885 | -1.83548718 | -1.630353704 | 4.57E-08 | 7.340294611 | Unpaired t-test | 0.001015015 | permutation FDR (250 perm) | 1 | 2.53E-07 | FDR correction BH |
| l(1)G0020~X2JD07 | VMAT~Control | VMAT | Control | 13.6496233 | 0.22623591 | 15.48690547 | 0.168892099 | 0.279848483 | -1.837282169 | -1.63131345 | 4.68E-08 | 7.329895737 | Unpaired t-test | 0.001007007 | permutation FDR (250 perm) | 1 | 2.57E-07 | FDR correction BH |
| Acbp2~P42281 | VMAT~Control | VMAT | Control | 13.40540743 | 0.431812302 | 15.24526302 | 1.761812303 | 0.279349744 | -1.839855599 | -1.015806313 | 0.046729779 | 1.330406273 | Unpaired t-test | 0.01298163 | permutation FDR (250 perm) | 1 | 0.058573962 | FDR correction BH |
| Gs2~P20478-3 | VMAT~Control | VMAT | Control | 16.52545079 | 0.046011031 | 18.39362395 | 0.339197815 | 0.273920063 | -1.868173157 | -1.620154836 | 2.49E-07 | 6.603208075 | Unpaired t-test | 0.001064532 | permutation FDR (250 perm) | 1 | 1.03E-06 | FDR correction BH |
| Liprin-beta~Q9VU88 | VMAT~Control | VMAT | Control | 13.57100275 | 0.184473957 | 15.44673204 | 0.36644871 | 0.272489153 | -1.875729294 | -1.58493366 | 1.30E-06 | 5.88681389 | Unpaired t-test | 0.001273543 | permutation FDR (250 perm) | 1 | 4.21E-06 | FDR correction BH |
| Ank2~M9MRX4 | VMAT~Control | VMAT | Control | 20.05567087 | 0.084330164 | 21.93551864 | 0.263127669 | 0.271712385 | -1.879847768 | -1.673102466 | 3.05E-08 | 7.515219626 | Unpaired t-test | 0.00081206 | permutation FDR (250 perm) | 1 | 1.82E-07 | FDR correction BH |
| Zasp52~A0A126GUN6 | VMAT~Control | VMAT | Control | 19.14467541 | 0.079912985 | 21.02456779 | 0.111641413 | 0.271703983 | -1.879892379 | -1.771143884 | 3.24E-11 | 10.48965837 | Unpaired t-test | 0.000495683 | permutation FDR (250 perm) | 1 | 8.97E-10 | FDR correction BH |
| OtopLc~B7Z0W9 | VMAT~Control | VMAT | Control | 13.14307319 | 0.275470105 | 15.02846667 | 0.244429545 | 0.270669929 | -1.88539348 | -1.618781088 | 4.55E-07 | 6.342441367 | Unpaired t-test | 0.00107 | permutation FDR (250 perm) | 1 | 1.71E-06 | FDR correction BH |
| kay~A0A0B4LHU8 | VMAT~Control | VMAT | Control | 13.57786173 | 0.355423125 | 15.46433999 | 0.529815385 | 0.270466485 | -1.88647826 | -1.467713998 | 5.99E-05 | 4.222686771 | Unpaired t-test | 0.002278705 | permutation FDR (250 perm) | 1 | 0.000123982 | FDR correction BH |
| prom~A0AVW1 | VMAT~Control | VMAT | Control | 14.09913 | 0.977212341 | 15.98889193 | 0.273974977 | 0.269851585 | -1.889761934 | -1.299811708 | 0.001937504 | 2.712757428 | Unpaired t-test | 0.004758955 | permutation FDR (250 perm) | 1 | 0.002946308 | FDR correction BH |
| tth~Q9VYA9 | VMAT~Control | VMAT | Control | 13.28006997 | 0.230000505 | 15.17376736 | 0.339279029 | 0.269116473 | -1.89369739 | -1.600340898 | 1.18E-06 | 5.928555149 | Unpaired t-test | 0.00116026 | permutation FDR (250 perm) | 1 | 3.89E-06 | FDR correction BH |
| 65K~A0A0B4LGA2 | VMAT~Control | VMAT | Control | 13.38726141 | 0.221844585 | 15.28569997 | 1.442400915 | 0.26823352 | -1.898438559 | -1.148726533 | 0.015595876 | 1.806990223 | Unpaired t-test | 0.008421756 | permutation FDR (250 perm) | 1 | 0.020871343 | FDR correction BH |
| CG9776~Q9V468-2 | VMAT~Control | VMAT | Control | 13.32886596 | 0.310631707 | 15.22734139 | 0.343154992 | 0.268226665 | -1.89847543 | -1.572886165 | 3.49E-06 | 5.456977364 | Unpaired t-test | 0.001344622 | permutation FDR (250 perm) | 1 | 1.01E-05 | FDR correction BH |
| eIF3b~E2QCG7 | VMAT~Control | VMAT | Control | 13.51295063 | 0.278919623 | 15.41338279 | 0.084394746 | 0.267863115 | -1.900432161 | -1.68131983 | 4.58E-08 | 7.338753158 | Unpaired t-test | 0.000798391 | permutation FDR (250 perm) | 1 | 2.53E-07 | FDR correction BH |
| papi~Q9VQ91 | VMAT~Control | VMAT | Control | 13.30081787 | 0.247930548 | 15.20798483 | 0.262993311 | 0.266615587 | -1.907166967 | -1.641790412 | 3.42E-07 | 6.465474812 | Unpaired t-test | 0.000964411 | permutation FDR (250 perm) | 1 | 1.35E-06 | FDR correction BH |
| CG13335~Q8IH23 | VMAT~Control | VMAT | Control | 14.49350899 | 1.745311959 | 16.40183425 | 0.066165946 | 0.266401616 | -1.908325262 | -1.071437985 | 0.034660891 | 1.460160272 | Unpaired t-test | 0.010977715 | permutation FDR (250 perm) | 1 | 0.044271155 | FDR correction BH |
| Mf~Q70VI1 | VMAT~Control | VMAT | Control | 17.11173437 | 0.300969487 | 19.0276147 | 0.383134803 | 0.265010177 | -1.915880332 | -1.573117601 | 5.10E-06 | 5.292548134 | Unpaired t-test | 0.001344622 | permutation FDR (250 perm) | 1 | 1.41E-05 | FDR correction BH |
| Dmel\CG14615~Q9VR51 | VMAT~Control | VMAT | Control | 13.34894354 | 0.233816446 | 15.26503546 | 0.162695952 | 0.264971312 | -1.916091923 | -1.699583285 | 3.41E-08 | 7.467851315 | Unpaired t-test | 0.000724521 | permutation FDR (250 perm) | 1 | 1.99E-07 | FDR correction BH |
| spag~Q9V3E9 | VMAT~Control | VMAT | Control | 13.57365958 | 0.41997989 | 15.49254403 | 0.557747482 | 0.264458921 | -1.918884454 | -1.462298207 | 0.000108955 | 3.962753701 | Unpaired t-test | 0.002323529 | permutation FDR (250 perm) | 1 | 0.000212493 | FDR correction BH |
| Dmel\CG1965~Q9VI54 | VMAT~Control | VMAT | Control | 13.64159578 | 0.697089384 | 15.57843216 | 0.155016253 | 0.261188561 | -1.936836382 | -1.468008708 | 0.000121225 | 3.916408056 | Unpaired t-test | 0.002278705 | permutation FDR (250 perm) | 1 | 0.000233331 | FDR correction BH |

| | | | | | | | | | | | | | | | | | | |
|---|---|---|---|---|---|---|---|---|---|---|---|---|---|---|---|---|---|---|
| CCT8~Q7K3J0 | VMAT~Control | VMAT | Control | 20.59996473 | 0.145847723 | 22.53813921 | 0.129254068 | 0.260946421 | -1.938174477 | -1.782798363 | 7.59E-10 | 9.119859062 | Unpaired t-test | 0.000453482 | permutation FDR (250 perm) | 1 | 1.00E-08 | FDR correction BH |
| ras~A0A4D6K3W6 | VMAT~Control | VMAT | Control | 15.1780705 | 0.530932871 | 17.11976315 | 0.184690004 | 0.26031085 | -1.941692649 | -1.551621087 | 1.60E-05 | 4.795894121 | Unpaired t-test | 0.001488603 | permutation FDR (250 perm) | 1 | 3.80E-05 | FDR correction BH |
| bol~A4V1Q1 | VMAT~Control | VMAT | Control | 13.86598814 | 0.971169857 | 15.80925674 | 0.510783423 | 0.260026651 | -1.943268597 | -1.303570067 | 0.002686577 | 2.570800668 | Unpaired t-test | 0.004679574 | permutation FDR (250 perm) | 1 | 0.004000128 | FDR correction BH |
| Akr1B~Q9VTK9 | VMAT~Control | VMAT | Control | 15.28943909 | 0.289604989 | 17.23365971 | 0.113035546 | 0.259855117 | -1.944220626 | -1.706907572 | 6.85E-08 | 7.164539529 | Unpaired t-test | 0.000688541 | permutation FDR (250 perm) | 1 | 3.44E-07 | FDR correction BH |
| enc~M9PBM2 | VMAT~Control | VMAT | Control | 13.21099456 | 0.292409403 | 15.16321931 | 0.125679738 | 0.258417423 | -1.952224751 | -1.708974929 | 8.24E-08 | 7.084157794 | Unpaired t-test | 0.000686522 | permutation FDR (250 perm) | 1 | 4.04E-07 | FDR correction BH |
| Dmel\CG11710~Q8T921 | VMAT~Control | VMAT | Control | 13.42103577 | 0.306449237 | 15.37483061 | 0.433137818 | 0.25813634 | -1.953794839 | -1.579099187 | 9.14E-06 | 5.039084874 | Unpaired t-test | 0.001308765 | permutation FDR (250 perm) | 1 | 2.33E-05 | FDR correction BH |
| Patr-1~Q9VEN9 | VMAT~Control | VMAT | Control | 13.2623765 | 0.282440533 | 15.21713769 | 0.421433289 | 0.257963493 | -1.954761184 | -1.593274787 | 6.12E-06 | 5.213580311 | Unpaired t-test | 0.001212968 | permutation FDR (250 perm) | 1 | 1.66E-05 | FDR correction BH |
| Dmel\CG46280~Q9VG01 | VMAT~Control | VMAT | Control | 16.25882407 | 0.115732083 | 18.21432788 | 0.089336064 | 0.25783074 | -1.955503813 | -1.835493295 | 4.08E-11 | 10.38895163 | Unpaired t-test | 0.000326864 | permutation FDR (250 perm) | 1 | 1.08E-09 | FDR correction BH |
| Dmel\CG7974~Q9W0A9 | VMAT~Control | VMAT | Control | 13.60692483 | 0.409055381 | 15.56328905 | 0.725655272 | 0.257677019 | -1.956364218 | -1.42536835 | 0.000372568 | 3.428794542 | Unpaired t-test | 0.00270709 | permutation FDR (250 perm) | 1 | 0.000644871 | FDR correction BH |
| Hil~Q0E908 | VMAT~Control | VMAT | Control | 14.10246967 | 0.17836801 | 16.06259382 | 0.434312907 | 0.257006339 | -1.960124149 | -1.619973752 | 2.98E-06 | 5.526480108 | Unpaired t-test | 0.001066533 | permutation FDR (250 perm) | 1 | 8.73E-06 | FDR correction BH |
| Dmel\CG31974~Q9VPK3 | VMAT~Control | VMAT | Control | 13.22784009 | 0.498534838 | 15.18806395 | 0.445095952 | 0.256988577 | -1.96022386 | -1.509164276 | 6.40E-05 | 4.193587309 | Unpaired t-test | 0.001891626 | permutation FDR (250 perm) | 1 | 0.000131304 | FDR correction BH |
| Urod~Q9V595 | VMAT~Control | VMAT | Control | 14.10697651 | 1.119956003 | 16.07173062 | 0.340035719 | 0.256182864 | -1.964754119 | -1.289686104 | 0.003761673 | 2.424619009 | Unpaired t-test | 0.004971456 | permutation FDR (250 perm) | 1 | 0.005503404 | FDR correction BH |
| CG5822~Q7KTP7 | VMAT~Control | VMAT | Control | 13.54653779 | 0.33688676 | 15.51385931 | 0.311142534 | 0.25572737 | -1.967321519 | -1.632514775 | 2.32E-06 | 5.633760027 | Unpaired t-test | 0.000995493 | permutation FDR (250 perm) | 1 | 7.00E-06 | FDR correction BH |
| eIF4G2~Q9VCH1 | VMAT~Control | VMAT | Control | 13.37360583 | 0.5121103 | 15.34334324 | 0.15055651 | 0.255299495 | -1.969737411 | -1.590145783 | 8.97E-06 | 5.047191656 | Unpaired t-test | 0.001226933 | permutation FDR (250 perm) | 1 | 2.29E-05 | FDR correction BH |
| Tm2~A0A0B4KHJ9 | VMAT~Control | VMAT | Control | 18.63478304 | 0.211932842 | 20.61905114 | 0.251639338 | 0.252741048 | -1.984268102 | -1.729765678 | 9.67E-08 | 7.014427593 | Unpaired t-test | 0.000618807 | permutation FDR (250 perm) | 1 | 4.66E-07 | FDR correction BH |
| CG7701~A8DY97 | VMAT~Control | VMAT | Control | 15.95227254 | 0.476822664 | 17.93930174 | 0.348695676 | 0.252257802 | -1.987029206 | -1.571796162 | 2.01E-05 | 4.696291446 | Unpaired t-test | 0.001350598 | permutation FDR (250 perm) | 1 | 4.65E-05 | FDR correction BH |
| D19A~Q9VRV5 | VMAT~Control | VMAT | Control | 13.40691478 | 0.377804225 | 15.3955694 | 0.203601578 | 0.251973756 | -1.988654616 | -1.668429463 | 1.15E-06 | 5.940391997 | Unpaired t-test | 0.000831743 | permutation FDR (250 perm) | 1 | 3.81E-06 | FDR correction BH |
| pic~Q9XYZ5 | VMAT~Control | VMAT | Control | 13.71350933 | 0.230687271 | 15.70397437 | 0.25110831 | 0.251657754 | -1.990465047 | -1.727093728 | 1.32E-07 | 6.879489944 | Unpaired t-test | 0.000628918 | permutation FDR (250 perm) | 1 | 6.13E-07 | FDR correction BH |
| mnb~X2JFM3 | VMAT~Control | VMAT | Control | 13.29687265 | 0.29661581 | 15.29243312 | 0.062529848 | 0.250770498 | -1.995560466 | -1.757326463 | 4.19E-08 | 7.377865479 | Unpaired t-test | 0.000529949 | permutation FDR (250 perm) | 1 | 2.36E-07 | FDR correction BH |
| CG7065~Q7YZA2 | VMAT~Control | VMAT | Control | 14.27050871 | 0.371722353 | 16.27378515 | 0.392109396 | 0.249432881 | -2.003276439 | -1.613423567 | 8.60E-06 | 5.065378369 | Unpaired t-test | 0.001086913 | permutation FDR (250 perm) | 1 | 2.21E-05 | FDR correction BH |
| mask~A0A0B4KHR8 | VMAT~Control | VMAT | Control | 15.06789754 | 0.40329141 | 17.07316088 | 0.384031478 | 0.249089594 | -2.005263341 | -1.605433575 | 1.11E-05 | 4.953574015 | Unpaired t-test | 0.001128871 | permutation FDR (250 perm) | 1 | 2.75E-05 | FDR correction BH |
| Tm1~P06754-1 | VMAT~Control | VMAT | Control | 18.64525963 | 0.219132592 | 20.65053682 | 0.2156596 | 0.249087203 | -2.005277189 | -1.762884533 | 4.58E-08 | 7.339164839 | Unpaired t-test | 0.000511675 | permutation FDR (250 perm) | 1 | 2.53E-07 | FDR correction BH |
| CG5340~Q86B83 | VMAT~Control | VMAT | Control | 14.13988947 | 0.252515026 | 16.14687387 | 0.377198408 | 0.24879262 | -2.006984405 | -1.668317937 | 1.76E-06 | 5.753403481 | Unpaired t-test | 0.000831743 | permutation FDR (250 perm) | 1 | 5.53E-06 | FDR correction BH |
| BuGZ~Q9VJI6 | VMAT~Control | VMAT | Control | 13.66493934 | 0.169141055 | 15.6740347 | 0.149857886 | 0.248428851 | -2.009095363 | -1.824691229 | 2.28E-09 | 8.642837841 | Unpaired t-test | 0.00034898 | permutation FDR (250 perm) | 1 | 2.38E-08 | FDR correction BH |
| Dmel\CG15717~Q9VYG8 | VMAT~Control | VMAT | Control | 15.927712 | 0.443481906 | 17.93692338 | 0.275699156 | 0.248408873 | -2.009211386 | -1.628827798 | 6.19E-06 | 5.208115739 | Unpaired t-test | 0.001023023 | permutation FDR (250 perm) | 1 | 1.68E-05 | FDR correction BH |
| Dmel\CG8963~A0A0B4K891 | VMAT~Control | VMAT | Control | 14.11858601 | 0.274011052 | 16.12817231 | 0.482044895 | 0.248344328 | -2.0095863 | -1.610282199 | 1.05E-05 | 4.978299953 | Unpaired t-test | 0.001100899 | permutation FDR (250 perm) | 1 | 2.62E-05 | FDR correction BH |
| Dmel\CG10984~Q8T0E7 | VMAT~Control | VMAT | Control | 13.32069861 | 0.117625257 | 15.33453257 | 0.203507206 | 0.247614215 | -2.013833958 | -1.822276571 | 3.27E-09 | 8.485808746 | Unpaired t-test | 0.00035102 | permutation FDR (250 perm) | 1 | 3.13E-08 | FDR correction BH |
| Taf4~A4V201 | VMAT~Control | VMAT | Control | 14.4770722 | 0.114379719 | 16.49123292 | 0.458288563 | 0.247558139 | -2.014160713 | -1.662891875 | 2.46E-06 | 5.60993221 | Unpaired t-test | 0.000859438 | permutation FDR (250 perm) | 1 | 7.35E-06 | FDR correction BH |
| Dis3l2~Q9W0T7 | VMAT~Control | VMAT | Control | 13.17575327 | 0.433368848 | 15.20086503 | 0.254199622 | 0.245686116 | -2.025111766 | -1.653572103 | 4.08E-06 | 5.388819684 | Unpaired t-test | 0.00091675 | permutation FDR (250 perm) | 1 | 1.16E-05 | FDR correction BH |
| Top1~Q8IR39 | VMAT~Control | VMAT | Control | 15.17491015 | 0.573735375 | 17.20395701 | 0.272224068 | 0.245016896 | -2.029046859 | -1.580255333 | 3.13E-05 | 4.505148541 | Unpaired t-test | 0.001301445 | permutation FDR (250 perm) | 1 | 6.95E-05 | FDR correction BH |
| Cpsf160~Q9V726-2 | VMAT~Control | VMAT | Control | 13.3944404 | 0.254066454 | 15.42746312 | 0.196079346 | 0.244342594 | -2.033022718 | -1.777856221 | 6.05E-08 | 7.217892216 | Unpaired t-test | 0.000469512 | permutation FDR (250 perm) | 1 | 3.15E-07 | FDR correction BH |
| scaf6~Q59E33 | VMAT~Control | VMAT | Control | 13.44131359 | 0.341624419 | 15.47476559 | 0.284170283 | 0.244269901 | -2.033451992 | -1.696344365 | 1.29E-06 | 5.890364804 | Unpaired t-test | 0.000731855 | permutation FDR (250 perm) | 1 | 4.18E-06 | FDR correction BH |
| eIF4E1~M9PBZ9 | VMAT~Control | VMAT | Control | 16.30234532 | 0.101816857 | 18.33851417 | 0.151945383 | 0.243810329 | -2.036168849 | -1.882209046 | 2.50E-10 | 9.601512075 | Unpaired t-test | 0.000256148 | permutation FDR (250 perm) | 1 | 4.30E-09 | FDR correction BH |
| Art8~Q8SX32 | VMAT~Control | VMAT | Control | 14.84557222 | 0.098745718 | 16.8914475 | 0.694561048 | 0.242175482 | -2.045875281 | -1.557290159 | 6.72E-05 | 4.172943102 | Unpaired t-test | 0.001459593 | permutation FDR (250 perm) | 1 | 0.000136862 | FDR correction BH |
| Cpsf100~Q9V3D6 | VMAT~Control | VMAT | Control | 13.39371458 | 0.449352577 | 15.44635854 | 0.259315945 | 0.241041932 | -2.052643954 | -1.666082335 | 4.83E-06 | 5.316325183 | Unpaired t-test | 0.000841788 | permutation FDR (250 perm) | 1 | 1.34E-05 | FDR correction BH |
| Dmel\CG10907~Q9VTN7 | VMAT~Control | VMAT | Control | 13.42133161 | 0.280384367 | 15.47740562 | 0.060943368 | 0.240469529 | -2.056074001 | -1.822244476 | 1.85E-08 | 7.732436062 | Unpaired t-test | 0.00035102 | permutation FDR (250 perm) | 1 | 1.23E-07 | FDR correction BH |
| Jabba~B7YZK4 | VMAT~Control | VMAT | Control | 15.16014936 | 0.320824252 | 17.2323071 | 0.638006397 | 0.237803567 | -2.072157739 | -1.570568153 | 7.00E-05 | 4.154995067 | Unpaired t-test | 0.001356574 | permutation FDR (250 perm) | 1 | 0.000142231 | FDR correction BH |
| Spt5~Q9V460 | VMAT~Control | VMAT | Control | 14.04448169 | 0.238257446 | 16.12018532 | 0.629852531 | 0.237219806 | -2.075703628 | -1.59527403 | 4.23E-05 | 4.373312226 | Unpaired t-test | 0.001206191 | permutation FDR (250 perm) | 1 | 9.10E-05 | FDR correction BH |
| AGO1~Q7KY08 | VMAT~Control | VMAT | Control | 16.20118041 | 0.182968736 | 18.30429975 | 0.229772809 | 0.23275445 | -2.103119343 | -1.858935449 | 1.87E-08 | 7.729203017 | Unpaired t-test | 0.000278261 | permutation FDR (250 perm) | 1 | 1.23E-07 | FDR correction BH |
| GABA-B-R2~Q8IN24 | VMAT~Control | VMAT | Control | 13.21449779 | 0.253507334 | 15.33189708 | 0.28550008 | 0.230461987 | -2.117399287 | -1.808585033 | 2.14E-07 | 6.668798173 | Unpaired t-test | 0.000385321 | permutation FDR (250 perm) | 1 | 9.23E-07 | FDR correction BH |
| Dmel\CG6340~Q9VXU3 | VMAT~Control | VMAT | Control | 13.829211 | 0.221046756 | 15.94758835 | 0.573275779 | 0.2303058 | -2.118377353 | -1.661765552 | 1.63E-05 | 4.788985963 | Unpaired t-test | 0.000861446 | permutation FDR (250 perm) | 1 | 3.85E-05 | FDR correction BH |
| Dmel\CG12576~Q8IQ25 | VMAT~Control | VMAT | Control | 14.04413024 | 0.947672103 | 16.166575 | 0.282214416 | 0.22965741 | -2.12244476 | -1.471666233 | 0.000723977 | 3.140275477 | Unpaired t-test | 0.002215022 | permutation FDR (250 perm) | 1 | 0.00118104 | FDR correction BH |
| DCTN4-p62~Q7K130 | VMAT~Control | VMAT | Control | 15.45608022 | 0.43443152 | 17.58097977 | 0.743343233 | 0.229266974 | -2.124899548 | -1.534176056 | 0.000255051 | 3.593372641 | Unpaired t-test | 0.001647873 | permutation FDR (250 perm) | 1 | 0.000463264 | FDR correction BH |
| CG31145~A4VCL2-3 | VMAT~Control | VMAT | Control | 13.45166903 | 0.450067391 | 15.57813728 | 0.340370223 | 0.229017817 | -2.126468253 | -1.697977059 | 7.45E-06 | 5.127863723 | Unpaired t-test | 0.000726172 | permutation FDR (250 perm) | 1 | 1.95E-05 | FDR correction BH |
| fand~A1Z9G2 | VMAT~Control | VMAT | Control | 13.45378932 | 0.352845022 | 15.58576266 | 0.155701813 | 0.228145588 | -2.131973344 | -1.818348845 | 2.21E-07 | 6.655628506 | Unpaired t-test | 0.000367347 | permutation FDR (250 perm) | 1 | 9.43E-07 | FDR correction BH |
| Lmpt~Q8IQQ3 | VMAT~Control | VMAT | Control | 13.81615942 | 0.271271638 | 15.9617977 | 0.633449273 | 0.225994837 | -2.145638283 | -1.640182034 | 3.89E-05 | 4.410290855 | Unpaired t-test | 0.000970426 | permutation FDR (250 perm) | 1 | 8.44E-05 | FDR correction BH |
| Rel~Q94527-2 | VMAT~Control | VMAT | Control | 15.1142823 | 0.484919461 | 17.26082933 | 0.329429809 | 0.225852527 | -2.146547036 | -1.700677066 | 9.61E-06 | 5.017440782 | Unpaired t-test | 0.000722503 | permutation FDR (250 perm) | 1 | 2.43E-05 | FDR correction BH |
| vig2~Q960D3 | VMAT~Control | VMAT | Control | 15.30983323 | 0.378793089 | 17.4597497 | 0.347816943 | 0.22532566 | -2.149916476 | -1.747923776 | 2.94E-06 | 5.531962241 | Unpaired t-test | 0.000551724 | permutation FDR (250 perm) | 1 | 8.64E-06 | FDR correction BH |
| BcDNA:RH62365~Q9VMR9 | VMAT~Control | VMAT | Control | 17.29662369 | 0.45321097 | 19.44767352 | 0.403078035 | 0.225148718 | -2.151049829 | -1.692079859 | 1.27E-05 | 4.895767073 | Unpaired t-test | 0.00749245 | permutation FDR (250 perm) | 1 | 3.10E-05 | FDR correction BH |
| Pfdn6~M9PD09 | VMAT~Control | VMAT | Control | 13.39152391 | 0.448136492 | 15.54512233 | 0.223812402 | 0.224751334 | -2.153598419 | -1.759451251 | 2.28E-06 | 5.642364128 | Unpaired t-test | 0.000517766 | permutation FDR (250 perm) | 1 | 6.92E-06 | FDR correction BH |
| Dmel\CG5355~Q9VKW5 | VMAT~Control | VMAT | Control | 13.58930589 | 0.30490954 | 15.74307616 | 0.146534239 | 0.224724564 | -2.153770268 | -1.870746522 | 5.77E-08 | 7.238711497 | Unpaired t-test | 0.000264209 | permutation FDR (250 perm) | 1 | 3.05E-07 | FDR correction BH |
| Girdin~Q8SX64 | VMAT~Control | VMAT | Control | 13.82380552 | 0.832244725 | 15.99198872 | 0.147241114 | 0.222490679 | -2.168183199 | -1.573460434 | 0.000188654 | 3.724333901 | Unpaired t-test | 0.001344622 | permutation FDR (250 perm) | 1 | 0.000348854 | FDR correction BH |
| pcm~Q9VWI1 | VMAT~Control | VMAT | Control | 14.91954643 | 0.133980162 | 17.08802618 | 0.449201349 | 0.222444949 | -2.168479755 | -1.792673947 | 1.16E-06 | 5.933768934 | Unpaired t-test | 0.00040061 | permutation FDR (250 perm) | 1 | 3.86E-06 | FDR correction BH |
| lva~X2JAF1 | VMAT~Control | VMAT | Control | 16.68579406 | 0.182986163 | 18.85856569 | 0.176062294 | 0.221784179 | -2.172771637 | -1.951190186 | 3.31E-09 | 8.480357186 | Unpaired t-test | 0.00015448 | permutation FDR (250 perm) | 1 | 3.13E-08 | FDR correction BH |
| Ptx1~B7Z0S4 | VMAT~Control | VMAT | Control | 13.38336424 | 0.391801661 | 15.55779264 | 0.410228328 | 0.221529633 | -2.174428404 | -1.734421212 | 6.40E-06 | 5.193521556 | Unpaired t-test | 0.000602934 | permutation FDR (250 perm) | 1 | 1.72E-05 | FDR correction BH |
| tn~A0A0B4LFW8 | VMAT~Control | VMAT | Control | 13.36389715 | 0.391114694 | 15.53867639 | 0.621440481 | 0.221475767 | -2.174779242 | -1.637169563 | 5.91E-05 | 4.228641141 | Unpaired t-test | 0.000983968 | permutation FDR (250 perm) | 1 | 0.000122509 | FDR correction BH |
| fus~A0A0B4K880 | VMAT~Control | VMAT | Control | 13.2387004 | 0.489054564 | 15.41699439 | 0.552845009 | 0.220936856 | -2.17829399 | -1.637698639 | 6.09E-05 | 4.215680972 | Unpaired t-test | 0.000983968 | permutation FDR (250 perm) | 1 | 0.000125557 | FDR correction BH |
| unk~A0A0B4K6Y4 | VMAT~Control | VMAT | Control | 13.38107599 | 0.400565806 | 15.56326349 | 2.02471139 | 0.220341401 | -2.182187497 | -1.134766 | 0.039670648 | 1.401530701 | Unpaired t-test | 0.008870891 | permutation FDR (250 perm) | 1 | 0.050206831 | FDR correction BH |
| Prm~A4V1N8 | VMAT~Control | VMAT | Control | 15.30449346 | 0.52776217 | 17.48820447 | 0.524758282 | 0.22010884 | -2.183711006 | -1.638392474 | 6.38E-05 | 4.194865198 | Unpaired t-test | 0.000980451 | permutation FDR (250 perm) | 1 | 0.000131032 | FDR correction BH |
| e(y)1~Q27272 | VMAT~Control | VMAT | Control | 13.2352917 | 0.60971988 | 15.42397836 | 0.18503718 | 0.219351023 | -2.188686657 | -1.703317595 | 1.68E-05 | 4.774945861 | Unpaired t-test | 0.000712412 | permutation FDR (250 perm) | 1 | 3.95E-05 | FDR correction BH |
| wuho~Q9W415 | VMAT~Control | VMAT | Control | 13.32984199 | 0.345940057 | 15.52618431 | 0.343658357 | 0.21819012 | -2.196342318 | -1.803131073 | 1.49E-06 | 5.827168182 | Unpaired t-test | 0.000389002 | permutation FDR (250 perm) | 1 | 4.78E-06 | FDR correction BH |
| CG7083~Q9VSH9 | VMAT~Control | VMAT | Control | 13.52042254 | 0.26823938 | 15.72198376 | 0.264279641 | 0.217402251 | -2.20156122 | -1.884249858 | 1.30E-07 | 6.885960304 | Unpaired t-test | 0.000252049 | permutation FDR (250 perm) | 1 | 6.05E-07 | FDR correction BH |
| MED1~Q9VP05 | VMAT~Control | VMAT | Control | 13.39839169 | 0.203275318 | 15.60479687 | 0.209146807 | 0.216673529 | -2.206405184 | -1.951823836 | 1.10E-08 | 7.960279876 | Unpaired t-test | 0.00015448 | permutation FDR (250 perm) | 1 | 7.96E-08 | FDR correction BH |
| pzg~Q9VP57 | VMAT~Control | VMAT | Control | 15.21660006 | 0.113506972 | 17.42337357 | 0.076935987 | 0.21661822 | -2.206773502 | -2.07926527 | 6.52E-12 | 11.18550659 | Unpaired t-test | 6.27E-05 | permutation FDR (250 perm) | 1 | 3.01E-10 | FDR correction BH |
| Lasp~M9NDA6 | VMAT~Control | VMAT | Control | 15.19681973 | 0.307780204 | 17.408034 | 0.18016905 | 0.215952471 | -2.211214268 | -1.907053271 | 7.43E-08 | 7.128785922 | Unpaired t-test | 0.000215606 | permutation FDR (250 perm) | 1 | 3.68E-07 | FDR correction BH |
| sdt~X2JIZ6 | VMAT~Control | VMAT | Control | 13.45096496 | 0.60540061 | 15.66836061 | 0.306211131 | 0.21502918 | -2.217395643 | -1.701232094 | 2.58E-05 | 4.589058677 | Unpaired t-test | 0.000722503 | permutation FDR (250 perm) | 1 | 5.83E-05 | FDR correction BH |
| His2A:CG33859~Q4AB57 | VMAT~Control | VMAT | Control | 14.42785802 | 1.172843335 | 16.64954271 | 0.690580168 | 0.21439086 | -2.221684692 | -1.381059868 | 0.00446231 | 2.350440275 | Unpaired t-test | 0.00337129 | permutation FDR (250 perm) | 1 | 0.006464171 | FDR correction BH |
| par-6~O97111 | VMAT~Control | VMAT | Control | 14.31122092 | 0.528464928 | 16.53347075 | 0.231924394 | 0.214306895 | -2.222249826 | -1.766361473 | 6.15E-06 | 5.211099363 | Unpaired t-test | 0.000497715 | permutation FDR (250 perm) | 1 | 1.67E-05 | FDR correction BH |
| SRPK~Q0E965 | VMAT~Control | VMAT | Control | 13.75636043 | 0.579910215 | 15.97983233 | 0.75997136 | 0.214125439 | -2.223471892 | -1.557580383 | 0.000400774 | 3.397100484 | Unpaired t-test | 0.001459593 | permutation FDR (250 perm) | 1 | 0.000688629 | FDR correction BH |
| elav~M9NFP7 | VMAT~Control | VMAT | Control | 14.60473644 | 1.460854118 | 16.83167956 | 0.303936519 | 0.213610856 | -2.226943125 | -1.335655238 | 0.007525626 | 2.123457381 | Unpaired t-test | 0.004066019 | permutation FDR (250 perm) | 1 | 0.010646228 | FDR correction BH |
| Inos~O97477 | VMAT~Control | VMAT | Control | 13.75613413 | 0.411526169 | 15.99994162 | 0.128413657 | 0.211128392 | -2.243807493 | -1.881138888 | 3.89E-07 | 6.409704768 | Unpaired t-test | 0.000256148 | permutation FDR (250 perm) | 1 | 1.51E-06 | FDR correction BH |
| Dmel\CG12065~Q9W3D6 | VMAT~Control | VMAT | Control | 15.22690012 | 0.263856882 | 17.47424268 | 0.097274589 | 0.210611693 | -2.247342558 | -1.996282299 | 6.44E-09 | 8.191423208 | Unpaired t-test | 0.000113813 | permutation FDR (250 perm) | 1 | 5.36E-08 | FDR correction BH |
| eEF1beta~A0A0B4LFL3 | VMAT~Control | VMAT | Control | 16.30234941 | 0.082336107 | 18.55741513 | 0.209414189 | 0.209487241 | -2.25506572 | -2.04888342 | 7.04E-10 | 9.152403594 | Unpaired t-test | 8.13E-05 | permutation FDR (250 perm) | 1 | 9.37E-09 | FDR correction BH |
| cyp33~Q9V3G3 | VMAT~Control | VMAT | Control | 14.83715016 | 0.148085388 | 17.09241919 | 0.097679896 | 0.209457721 | -2.255269026 | -2.089497594 | 6.75E-11 | 10.17068773 | Unpaired t-test | 5.65E-05 | permutation FDR (250 perm) | 1 | 1.58E-09 | FDR correction BH |
| Cpsf5~Q9VSY2 | VMAT~Control | VMAT | Control | 13.40728016 | 0.319270704 | 15.66495593 | 0.323819465 | 0.209108589 | -2.257675773 | -1.876130718 | 6.05E-07 | 6.218055417 | Unpaired t-test | 0.000260113 | permutation FDR (250 perm) | 1 | 2.19E-06 | FDR correction BH |
| Tango11~Q961C9 | VMAT~Control | VMAT | Control | 15.57246885 | 0.167815587 | 17.83641891 | 0.090546979 | 0.208201151 | -2.263950053 | -2.086057358 | 1.33E-10 | 9.877295024 | Unpaired t-test | 6.06E-05 | permutation FDR (250 perm) | 1 | 2.65E-09 | FDR correction BH |
| XRCC1~Q9Y095 | VMAT~Control | VMAT | Control | 13.48001485 | 0.287152362 | 15.74921453 | 0.099393702 | 0.207444933 | -2.269199675 | -1.997721733 | 1.24E-08 | 7.905782532 | Unpaired t-test | 0.000113813 | permutation FDR (250 perm) | 1 | 8.78E-08 | FDR correction BH |
| CG7414~Q9VNX8 | VMAT~Control | VMAT | Control | 17.1810067 | 0.135089249 | 19.4505007 | 0.141372203 | 0.207402616 | -2.269494002 | -2.086992054 | 1.66E-10 | 9.780061029 | Unpaired t-test | 6.06E-05 | permutation FDR (250 perm) | 1 | 3.20E-09 | FDR correction BH |
| M7BP~A1Z720 | VMAT~Control | VMAT | Control | 13.62675144 | 0.137913137 | 15.90051026 | 0.556940526 | 0.206790408 | -2.273758827 | -1.809461577 | 4.76E-06 | 5.322654669 | Unpaired t-test | 0.000379205 | permutation FDR (250 perm) | 1 | 1.32E-05 | FDR correction BH |
| Amun~Q9VYV4 | VMAT~Control | VMAT | Control | 13.03562802 | 0.516438804 | 15.31586419 | 0.213945604 | 0.205864052 | -2.280236164 | -1.824199578 | 3.67E-06 | 5.435512779 | Unpaired t-test | 0.00034898 | permutation FDR (250 perm) | 1 | 1.05E-05 | FDR correction BH |
| kst~Q9VZQ3 | VMAT~Control | VMAT | Control | 15.75974809 | 0.248961308 | 18.06311721 | 0.412586058 | 0.202589441 | -2.303369115 | -1.894991618 | 8.61E-07 | 6.065105708 | Unpaired t-test | 0.000231914 | permutation FDR (250 perm) | 1 | 2.95E-06 | FDR correction BH |
| shot~A1Z9J3 | VMAT~Control | VMAT | Control | 17.21166846 | 0.140573912 | 19.51841687 | 0.124663572 | 0.202115461 | -2.30674841 | -2.127945211 | 9.53E-11 | 10.02099328 | Unpaired t-test | 4.41E-05 | permutation FDR (250 perm) | 1 | 2.02E-09 | FDR correction BH |
| AspRS-m~Q9VJH2 | VMAT~Control | VMAT | Control | 13.71100636 | 0.200569423 | 16.01803049 | 0.30302515 | 0.202076837 | -2.30702413 | -1.984514737 | 5.93E-08 | 7.226968732 | Unpaired t-test | 0.000119959 | permutation FDR (250 perm) | 1 | 3.11E-07 | FDR correction BH |
| beag~Q9VHH8 | VMAT~Control | VMAT | Control | 13.20087092 | 0.328743598 | 15.51760467 | 0.419099359 | 0.200721387 | -2.316733752 | -1.871036753 | 2.05E-06 | 5.68825079 | Unpaired t-test | 0.000264209 | permutation FDR (250 perm) | 1 | 6.33E-06 | FDR correction BH |
| Parg~O46043 | VMAT~Control | VMAT | Control | 13.36744178 | 0.338253949 | 15.68698446 | 0.092291966 | 0.200330963 | -2.319542671 | -2.005134576 | 4.00E-08 | 7.398390081 | Unpaired t-test | 0.000111917 | permutation FDR (250 perm) | 1 | 2.27E-07 | FDR correction BH |
| DIP1~Q9TW12 | VMAT~Control | VMAT | Control | 13.23477158 | 0.375406156 | 15.55645271 | 0.097832393 | 0.20003424 | -2.321681127 | -1.97843448 | 1.04E-07 | 6.982276607 | Unpaired t-test | 0.000128033 | permutation FDR (250 perm) | 1 | 4.95E-07 | FDR correction BH |
| Strn-Mlck~A0A0B4KF84 | VMAT~Control | VMAT | Control | 14.43624487 | 1.422321352 | 16.76109735 | 0.345036772 | 0.199595005 | -2.324852478 | -1.40514364 | 0.005251239 | 2.27973819 | Unpaired t-test | 0.002986355 | permutation FDR (250 perm) | 1 | 0.007565127 | FDR correction BH |
| tou~A1Z8M2 | VMAT~Control | VMAT | Control | 13.42154593 | 0.265185812 | 15.7502958 | 0.162417878 | 0.199056533 | -2.328749877 | -2.044429897 | 1.21E-08 | 7.917543266 | Unpaired t-test | 8.75E-05 | permutation FDR (250 perm) | 1 | 8.60E-08 | FDR correction BH |
| Tm1~Q95TA3 | VMAT~Control | VMAT | Control | 13.65376856 | 0.234258367 | 15.98549542 | 0.480333456 | 0.198646206 | -2.331726858 | -1.881947531 | 1.99E-06 | 5.700724273 | Unpaired t-test | 0.000256148 | permutation FDR (250 perm) | 1 | 6.17E-06 | FDR correction BH |
| Gk2~Q9W095 | VMAT~Control | VMAT | Control | 14.2284771 | 0.188257195 | 16.56070124 | 0.095354193 | 0.198577747 | -2.332224134 | -2.131101456 | 2.69E-10 | 9.570366337 | Unpaired t-test | 3.57E-05 | permutation FDR (250 perm) | 1 | 4.59E-09 | FDR correction BH |
| Dek~A1ZAK4 | VMAT~Control | VMAT | Control | 13.35834139 | 0.326098104 | 15.69977585 | 0.09595067 | 0.197314042 | -2.341434465 | -2.032464433 | 2.71E-08 | 7.566990673 | Unpaired t-test | 8.73E-05 | permutation FDR (250 perm) | 1 | 1.65E-07 | FDR correction BH |
| egl~Q9W1K4 | VMAT~Control | VMAT | Control | 14.27331135 | 1.320645637 | 16.6173333 | 0.44353878 | 0.196960474 | -2.344021957 | -1.444225947 | 0.003708158 | 2.430841827 | Unpaired t-test | 0.002498287 | permutation FDR (250 perm) | 1 | 0.005435245 | FDR correction BH |
| eIF2Bdelta~Q0E8X6 | VMAT~Control | VMAT | Control | 14.56360009 | 0.348035164 | 16.92441141 | 0.276632989 | 0.194681633 | -2.360811317 | -1.969273171 | 3.23E-07 | 6.491287232 | Unpaired t-test | 0.000142488 | permutation FDR (250 perm) | 1 | 1.28E-06 | FDR correction BH |
| RpS19a~E2QD65 | VMAT~Control | VMAT | Control | 18.13271221 | 0.05278079 | 20.50242051 | 0.100788282 | 0.19348474 | -2.369708305 | -2.25497425 | 5.02E-13 | 12.29949645 | Unpaired t-test | 2.77E-05 | permutation FDR (250 perm) | 1 | 6.76E-11 | FDR correction BH |
| Dmel\CG3436~Q9VPL0 | VMAT~Control | VMAT | Control | 13.53865739 | 0.558618282 | 15.90948808 | 0.139489456 | 0.193334272 | -2.370830689 | -1.885363824 | 3.37E-06 | 5.472360495 | Unpaired t-test | 0.000250128 | permutation FDR (250 perm) | 1 | 9.77E-06 | FDR correction BH |
| bru3~M9MRZ0 | VMAT~Control | VMAT | Control | 13.37059803 | 0.311474265 | 15.75352112 | 0.113573989 | 0.191720553 | -2.382923088 | -2.075234623 | 1.80E-08 | 7.745183326 | Unpaired t-test | 6.68E-05 | permutation FDR (250 perm) | 1 | 1.20E-07 | FDR correction BH |
| TpnC41C~A0A0B4K6P6 | VMAT~Control | VMAT | Control | 13.41927945 | 0.221996409 | 15.81563538 | 0.128500654 | 0.18994374 | -2.396355932 | -2.149752394 | 1.40E-09 | 8.85340799 | Unpaired t-test | 3.37E-05 | permutation FDR (250 perm) | 1 | 1.59E-08 | FDR correction BH |
| Dmel\CG1674~L0MLJ3 | VMAT~Control | VMAT | Control | 13.33603019 | 0.33508702 | 15.73280667 | 0.135556658 | 0.189888379 | -2.39677648 | -2.063245993 | 3.91E-08 | 7.407902156 | Unpaired t-test | 7.51E-05 | permutation FDR (250 perm) | 1 | 2.23E-07 | FDR correction BH |
| CAH1~Q9V396 | VMAT~Control | VMAT | Control | 13.65940955 | 0.637280527 | 16.06058371 | 0.32348353 | 0.189310435 | -2.401174156 | -1.819602339 | 2.03E-05 | 4.691880505 | Unpaired t-test | 0.000359184 | permutation FDR (250 perm) | 1 | 4.70E-05 | FDR correction BH |
| Dmel\CG12224~Q9VGF2 | VMAT~Control | VMAT | Control | 13.33705518 | 0.320801823 | 15.74087073 | 1.317831111 | 0.188964149 | -2.403815551 | -1.49624741 | 0.002673389 | 2.572937848 | Unpaired t-test | 0.001978368 | permutation FDR (250 perm) | 1 | 0.003983011 | FDR correction BH |
| Msp300~M9PCM8 | VMAT~Control | VMAT | Control | 13.43680843 | 0.33451902 | 15.84336921 | 0.231677702 | 0.188604921 | -2.406560778 | -2.036047622 | 1.17E-07 | 6.933428705 | Unpaired t-test | 8.74E-05 | permutation FDR (250 perm) | 1 | 5.49E-07 | FDR correction BH |
| scrib~E1JIZ2 | VMAT~Control | VMAT | Control | 16.79539927 | 0.157055949 | 19.20856179 | 0.077580887 | 0.18774384 | -2.413162522 | -2.237850494 | 3.05E-11 | 10.51582772 | Unpaired t-test | 2.76E-05 | permutation FDR (250 perm) | 1 | 8.54E-10 | FDR correction BH |
| up~E1JJP0 | VMAT~Control | VMAT | Control | 15.5972098 | 0.31952545 | 18.01188513 | 0.194732332 | 0.187547076 | -2.414675324 | -2.068523829 | 5.07E-08 | 7.294831859 | Unpaired t-test | 6.68E-05 | permutation FDR (250 perm) | 1 | 2.74E-07 | FDR correction BH |
| eIF3g1~Q9W4X7 | VMAT~Control | VMAT | Control | 16.98238301 | 0.096324367 | 19.40033783 | 0.118975675 | 0.187121233 | -2.417954825 | -2.263028747 | 7.85E-12 | 11.10493035 | Unpaired t-test | 2.77E-05 | permutation FDR (250 perm) | 1 | 3.24E-10 | FDR correction BH |
| Fbl6~Q494G8 | VMAT~Control | VMAT | Control | 13.55979816 | 0.295608079 | 15.98629325 | 0.108652952 | 0.18601681 | -2.426495093 | -2.126923918 | 9.18E-09 | 8.037187014 | Unpaired t-test | 4.41E-05 | permutation FDR (250 perm) | 1 | 7.04E-08 | FDR correction BH |
| sqd~Q08473-2 | VMAT~Control | VMAT | Control | 16.85197334 | 0.06671458 | 19.28627425 | 0.094956369 | 0.18501307 | -2.434300906 | -2.314196357 | 4.68E-13 | 12.3300769 | Unpaired t-test | 1.92E-05 | permutation FDR (250 perm) | 1 | 6.76E-11 | FDR correction BH |
| Bx42~M9PGQ6 | VMAT~Control | VMAT | Control | 13.56012028 | 0.232382691 | 16.01941154 | 0.08529325 | 0.181835871 | -2.459291265 | -2.214173815 | 7.67E-10 | 9.115309437 | Unpaired t-test | 2.75E-05 | permutation FDR (250 perm) | 1 | 1.00E-08 | FDR correction BH |
| Cbp20~Q9V3L6 | VMAT~Control | VMAT | Control | 13.41291376 | 0.190469281 | 15.8783377 | 0.582817625 | 0.181064555 | -2.465423942 | -1.934865076 | 4.18E-06 | 5.379304438 | Unpaired t-test | 0.000168551 | permutation FDR (250 perm) | 1 | 1.18E-05 | FDR correction BH |
| TfIIFbeta~P41900 | VMAT~Control | VMAT | Control | 13.50081013 | 0.398567647 | 15.96669835 | 0.316507106 | 0.181006296 | -2.465888218 | -2.008689259 | 7.59E-07 | 6.119541291 | Unpaired t-test | 0.000109845 | permutation FDR (250 perm) | 1 | 2.64E-06 | FDR correction BH |
| TBPH~Q8SXP8 | VMAT~Control | VMAT | Control | 13.36464263 | 0.253159928 | 15.83340078 | 0.29969908 | 0.18064658 | -2.468758153 | -2.10027051 | 6.45E-08 | 7.190312502 | Unpaired t-test | 5.03E-05 | permutation FDR (250 perm) | 1 | 3.29E-07 | FDR correction BH |
| Dmel\CG5787~Q9VK59 | VMAT~Control | VMAT | Control | 13.80213057 | 0.244721587 | 16.27949786 | 0.294036154 | 0.1795718 | -2.477367285 | -2.115450897 | 4.90E-08 | 7.309510461 | Unpaired t-test | 5.03E-05 | permutation FDR (250 perm) | 1 | 2.68E-07 | FDR correction BH |
| Dmel\CG33217~Q8MS38 | VMAT~Control | VMAT | Control | 13.40948076 | 0.546665825 | 15.89127474 | 0.074607712 | 0.179021656 | -2.481793973 | -1.990622399 | 1.51E-06 | 5.821846311 | Unpaired t-test | 0.000115822 | permutation FDR (250 perm) | 1 | 4.82E-06 | FDR correction BH |
| CG12259~Q9VAY7 | VMAT~Control | VMAT | Control | 13.42566794 | 0.508872447 | 15.90974188 | 0.324901705 | 0.178738963 | -2.484073933 | -1.955956719 | 3.39E-06 | 5.469265799 | Unpaired t-test | 0.00015242 | permutation FDR (250 perm) | 1 | 9.83E-06 | FDR correction BH |
| Dmel\CG5381~Q9VKZ6 | VMAT~Control | VMAT | Control | 14.82130648 | 0.254073764 | 17.30602625 | 0.210956933 | 0.178658966 | -2.484719774 | -2.164981531 | 1.16E-08 | 7.937392442 | Unpaired t-test | 3.17E-05 | permutation FDR (250 perm) | 1 | 8.29E-08 | FDR correction BH |
| Unr~M9ND61 | VMAT~Control | VMAT | Control | 13.40596122 | 0.551874033 | 15.89274977 | 0.340091547 | 0.178402959 | -2.486788552 | -1.927883666 | 6.36E-06 | 5.196564411 | Unpaired t-test | 0.000176773 | permutation FDR (250 perm) | 1 | 1.71E-05 | FDR correction BH |
| Dmel\CG11414~Q9W149 | VMAT~Control | VMAT | Control | 13.74232039 | 0.258623281 | 16.23128546 | 0.651532728 | 0.178134014 | -2.488965077 | -1.894924571 | 1.26E-05 | 4.900204324 | Unpaired t-test | 0.000231914 | permutation FDR (250 perm) | 1 | 3.08E-05 | FDR correction BH |
| BcDNA:RH31685~Q8INF8 | VMAT~Control | VMAT | Control | 13.19898595 | 0.261554347 | 15.69036986 | 0.134930423 | 0.177835603 | -2.491383911 | -2.201611526 | 3.67E-09 | 8.434945042 | Unpaired t-test | 2.75E-05 | permutation FDR (250 perm) | 1 | 3.42E-08 | FDR correction BH |
| Usp47~Q24574-2 | VMAT~Control | VMAT | Control | 13.25623473 | 0.397387275 | 15.75382043 | 0.668063766 | 0.177072772 | -2.497585702 | -1.853319949 | 2.98E-05 | 4.525817642 | Unpaired t-test | 0.000290389 | permutation FDR (250 perm) | 1 | 6.64E-05 | FDR correction BH |
| Df31~O16043 | VMAT~Control | VMAT | Control | 13.18291314 | 0.495157719 | 15.69346694 | 0.332686445 | 0.175488233 | -2.510553796 | -1.981836724 | 2.77E-06 | 5.558137958 | Unpaired t-test | 0.000119959 | permutation FDR (250 perm) | 1 | 8.19E-06 | FDR correction BH |
| Saf-B~Q8IMV6 | VMAT~Control | VMAT | Control | 13.39685836 | 0.224199484 | 15.92825713 | 0.082214544 | 0.172970897 | -2.531398774 | -2.287145665 | 4.05E-10 | 9.392161676 | Unpaired t-test | 2.13E-05 | permutation FDR (250 perm) | 1 | 6.28E-09 | FDR correction BH |
| Jasper~Q9VAA9 | VMAT~Control | VMAT | Control | 13.15644618 | 0.352604476 | 15.70971842 | 0.479291633 | 0.170368174 | -2.553272241 | -2.016640831 | 2.32E-06 | 5.634773142 | Unpaired t-test | 0.000103681 | permutation FDR (250 perm) | 1 | 7.00E-06 | FDR correction BH |
| Kank~Q7KMM8 | VMAT~Control | VMAT | Control | 15.41821899 | 0.699289233 | 17.97709233 | 0.210004147 | 0.16970802 | -2.558873347 | -1.928998625 | 1.41E-05 | 4.850730724 | Unpaired t-test | 0.000172662 | permutation FDR (250 perm) | 1 | 3.40E-05 | FDR correction BH |
| cher~A0A0B4JCY6 | VMAT~Control | VMAT | Control | 15.48252793 | 0.104088784 | 18.04862872 | 0.114016819 | 0.168859964 | -2.566100787 | -2.400373109 | 4.73E-12 | 11.32483189 | Unpaired t-test | 1.73E-05 | permutation FDR (250 perm) | 1 | 2.37E-10 | FDR correction BH |
| NC2alpha~Q9W2E3 | VMAT~Control | VMAT | Control | 13.49018365 | 0.265851267 | 16.0617651 | 0.396885836 | 0.168219698 | -2.571581446 | -2.118911958 | 2.84E-07 | 6.547112614 | Unpaired t-test | 5.04E-05 | permutation FDR (250 perm) | 1 | 1.14E-06 | FDR correction BH |
| up~P19351-6 | VMAT~Control | VMAT | Control | 13.36277831 | 0.245183146 | 15.93666804 | 0.307172618 | 0.167950764 | -2.573889734 | -2.189115897 | 4.40E-08 | 7.356180278 | Unpaired t-test | 3.17E-05 | permutation FDR (250 perm) | 1 | 2.46E-07 | FDR correction BH |
| Actn~M9MS06 | VMAT~Control | VMAT | Control | 14.06954873 | 0.273018057 | 16.64515686 | 0.180566367 | 0.167750837 | -2.575608132 | -2.246721457 | 7.48E-09 | 8.126020618 | Unpaired t-test | 2.76E-05 | permutation FDR (250 perm) | 1 | 5.99E-08 | FDR correction BH |
| RIC-3~E1JGP2 | VMAT~Control | VMAT | Control | 15.20034261 | 0.28258175 | 17.77710622 | 0.065712905 | 0.167616535 | -2.576763619 | -2.280833679 | 2.30E-09 | 8.638537589 | Unpaired t-test | 2.35E-05 | permutation FDR (250 perm) | 1 | 2.39E-08 | FDR correction BH |
| anon-WO0107627.3~Q9VZR5 | VMAT~Control | VMAT | Control | 13.95509 | 0.256186704 | 16.54362689 | 0.794930228 | 0.166254248 | -2.588536894 | -1.884615591 | 4.04E-05 | 4.393344686 | Unpaired t-test | 0.000252049 | permutation FDR (250 perm) | 1 | 8.72E-05 | FDR correction BH |
| Nelf-A~A0A0B4LIF3 | VMAT~Control | VMAT | Control | 13.24309759 | 0.352811506 | 15.83880439 | 0.407610029 | 0.165430047 | -2.595706797 | -2.091473493 | 8.05E-07 | 6.094456403 | Unpaired t-test | 5.65E-05 | permutation FDR (250 perm) | 1 | 2.79E-06 | FDR correction BH |
| Prosalpha3~P18053 | VMAT~Control | VMAT | Control | 13.31065921 | 0.182706933 | 15.90693958 | 0.476511713 | 0.165364291 | -2.596280365 | -2.11383848 | 4.82E-07 | 6.316914449 | Unpaired t-test | 5.03E-05 | permutation FDR (250 perm) | 1 | 1.80E-06 | FDR correction BH |
| Dmel\CG14109~Q9VU75 | VMAT~Control | VMAT | Control | 13.4147517 | 0.432270702 | 16.01392216 | 0.209867036 | 0.165033354 | -2.599170469 | -2.139417525 | 2.71E-07 | 6.566628055 | Unpaired t-test | 3.58E-05 | permutation FDR (250 perm) | 1 | 1.11E-06 | FDR correction BH |
| AOX1~Q9VF53 | VMAT~Control | VMAT | Control | 13.32603228 | 0.278018141 | 15.92888225 | 0.148203979 | 0.164612983 | -2.602849972 | -2.281408491 | 4.66E-09 | 8.331900236 | Unpaired t-test | 2.35E-05 | permutation FDR (250 perm) | 1 | 4.12E-08 | FDR correction BH |
| row~A0A0B4KER0 | VMAT~Control | VMAT | Control | 13.43879003 | 0.126474806 | 16.04171793 | 0.454570117 | 0.164604091 | -2.6029279 | -2.149382799 | 2.25E-07 | 6.647197343 | Unpaired t-test | 3.37E-05 | permutation FDR (250 perm) | 1 | 9.54E-07 | FDR correction BH |
| Gyf~Q7KQM6 | VMAT~Control | VMAT | Control | 16.09282302 | 0.317001705 | 18.69685138 | 0.361258975 | 0.164478582 | -2.604028359 | -2.14333733 | 2.67E-07 | 6.573412556 | Unpaired t-test | 3.58E-05 | permutation FDR (250 perm) | 1 | 1.10E-06 | FDR correction BH |
| mxt~M9PC59 | VMAT~Control | VMAT | Control | 14.47965569 | 0.323317286 | 17.08744536 | 0.509148307 | 0.164050322 | -2.607789672 | -2.053817509 | 2.16E-06 | 5.66479752 | Unpaired t-test | 7.92E-05 | permutation FDR (250 perm) | 1 | 6.62E-06 | FDR correction BH |
| Veneno~Q9VHR5 | VMAT~Control | VMAT | Control | 16.08720783 | 0.109118518 | 18.6968872 | 0.115439379 | 0.163835583 | -2.609679372 | -2.436585398 | 5.32E-12 | 11.27446409 | Unpaired t-test | 1.30E-05 | permutation FDR (250 perm) | 1 | 2.55E-10 | FDR correction BH |
| Taf6~M9PG32 | VMAT~Control | VMAT | Control | 14.11126988 | 0.291466808 | 16.72276262 | 0.236685269 | 0.163629782 | -2.611492742 | -2.236034838 | 2.47E-08 | 7.607616006 | Unpaired t-test | 2.76E-05 | permutation FDR (250 perm) | 1 | 1.53E-07 | FDR correction BH |
| ebi~Q95RJ9 | VMAT~Control | VMAT | Control | 13.20247674 | 0.331673991 | 15.81789719 | 0.136527025 | 0.163184909 | -2.615420449 | -2.253887767 | 1.56E-08 | 7.805675228 | Unpaired t-test | 2.76E-05 | permutation FDR (250 perm) | 1 | 1.06E-07 | FDR correction BH |
| TfIIA-L~P52654 | VMAT~Control | VMAT | Control | 13.59449274 | 0.306326894 | 16.20991492 | 0.335148804 | 0.163184713 | -2.615422183 | -2.173980129 | 1.50E-07 | 6.824682599 | Unpaired t-test | 3.17E-05 | permutation FDR (250 perm) | 1 | 6.80E-07 | FDR correction BH |
| Mbs~M9NEB2 | VMAT~Control | VMAT | Control | 13.2239881 | 0.337641001 | 15.83959032 | 0.42660792 | 0.16316435 | -2.615602218 | -2.103743095 | 8.16E-07 | 6.088314674 | Unpaired t-test | 5.03E-05 | permutation FDR (250 perm) | 1 | 2.82E-06 | FDR correction BH |
| Nup58~Q9VDV3 | VMAT~Control | VMAT | Control | 13.55665564 | 0.329220301 | 16.17343871 | 0.569086635 | 0.163030854 | -2.616783068 | -2.022208613 | 4.58E-06 | 5.339600255 | Unpaired t-test | 9.34E-05 | permutation FDR (250 perm) | 1 | 1.28E-05 | FDR correction BH |
| Nup358~A0A0B4K7J2-2 | VMAT~Control | VMAT | Control | 18.19153857 | 0.130453645 | 20.82342238 | 0.080968392 | 0.161333303 | -2.631883817 | -2.462778485 | 3.49E-12 | 11.45775035 | Unpaired t-test | 1.31E-05 | permutation FDR (250 perm) | 1 | 1.91E-10 | FDR correction BH |

| | | | | | | | | | | | | | | | | | | |
|---|---|---|---|---|---|---|---|---|---|---|---|---|---|---|---|---|---|---|
| Lsd-2~Q9VXY7 | VMAT~Control | VMAT | Control | 13.94466614 | 0.221698102 | 16.59258839 | 0.191346609 | 0.159549695 | -2.647922243 | -2.341287614 | 1.93E-09 | 8.714403653 | Unpaired t-test | 1.93E-05 | permutation FDR (250 perm) | 1 | 2.07E-08 | FDR correction BH |
| ninaC~M9PCU0 | VMAT~Control | VMAT | Control | 13.37330248 | 0.401056341 | 16.02513623 | 0.032341945 | 0.159117703 | -2.651833743 | -2.247430825 | 4.14E-08 | 7.382709018 | Unpaired t-test | 2.76E-05 | permutation FDR (250 perm) | 1 | 2.34E-07 | FDR correction BH |
| TpnC47D~A0A0B4LF64 | VMAT~Control | VMAT | Control | 13.26460272 | 0.347205985 | 15.93878352 | 0.573187017 | 0.156671992 | -2.6741808 | -2.057539663 | 4.47E-06 | 5.349641919 | Unpaired t-test | 7.92E-05 | permutation FDR (250 perm) | 1 | 1.25E-05 | FDR correction BH |
| rin~Q9VFT4 | VMAT~Control | VMAT | Control | 13.48473262 | 0.278138253 | 16.15960093 | 0.851447471 | 0.156597349 | -2.674868304 | -1.909828321 | 5.52E-05 | 4.258297617 | Unpaired t-test | 0.000213552 | permutation FDR (250 perm) | 1 | 0.000115235 | FDR correction BH |
| aru~Q9VPU7 | VMAT~Control | VMAT | Control | 14.74271935 | 0.520126711 | 17.41839704 | 0.340822217 | 0.156509519 | -2.675677692 | -2.093484518 | 2.26E-06 | 5.645577036 | Unpaired t-test | 5.65E-05 | permutation FDR (250 perm) | 1 | 6.89E-06 | FDR correction BH |
| lgs~Q961D9 | VMAT~Control | VMAT | Control | 13.3041234 | 0.437631451 | 15.9917171 | 0.180652861 | 0.155222146 | -2.687593691 | -2.217973059 | 1.72E-07 | 6.764531137 | Unpaired t-test | 2.75E-05 | permutation FDR (250 perm) | 1 | 7.68E-07 | FDR correction BH |
| GstD2~Q9VG98 | VMAT~Control | VMAT | Control | 13.98099805 | 1.045443316 | 16.67420919 | 0.412762711 | 0.154618929 | -2.693211142 | -1.792298271 | 0.000319967 | 3.49489507 | Unpaired t-test | 0.000410778 | permutation FDR (250 perm) | 1 | 0.000564196 | FDR correction BH |
| Dmel\CG15160~Q9VJ77 | VMAT~Control | VMAT | Control | 13.23365167 | 0.453634522 | 15.92934714 | 0.275002239 | 0.154352904 | -2.695695471 | -2.178800198 | 4.87E-07 | 6.312311978 | Unpaired t-test | 3.17E-05 | permutation FDR (250 perm) | 1 | 1.81E-06 | FDR correction BH |
| mod~A0A0B4K7G4 | VMAT~Control | VMAT | Control | 14.86815332 | 0.183778881 | 17.56825926 | 0.1812278 | 0.153881752 | -2.700105938 | -2.420690334 | 4.62E-10 | 9.335645754 | Unpaired t-test | 1.30E-05 | permutation FDR (250 perm) | 1 | 6.84E-09 | FDR correction BH |
| CG10057~Q9VBX4 | VMAT~Control | VMAT | Control | 13.19515651 | 0.509986988 | 15.8962454 | 0.295243127 | 0.153776943 | -2.70108889 | -2.137722714 | 1.27E-06 | 5.896990684 | Unpaired t-test | 3.58E-05 | permutation FDR (250 perm) | 1 | 4.13E-06 | FDR correction BH |
| su(r)~Q9W374 | VMAT~Control | VMAT | Control | 13.41877139 | 0.400927199 | 16.13066455 | 0.407469616 | 0.152629617 | -2.711893159 | -2.159759515 | 9.23E-07 | 6.034788891 | Unpaired t-test | 3.38E-05 | permutation FDR (250 perm) | 1 | 3.14E-06 | FDR correction BH |
| dop~Q9VUQ9 | VMAT~Control | VMAT | Control | 13.29925652 | 0.345986396 | 16.01267154 | 2.664478583 | 0.152468696 | -2.713415024 | -1.232476921 | 0.047515323 | 1.323166312 | Unpaired t-test | 0.006253739 | permutation FDR (250 perm) | 1 | 0.05949525 | FDR correction BH |
| Nup214~Q9W1X4 | VMAT~Control | VMAT | Control | 16.10194788 | 0.137271644 | 18.8156298 | 0.046357205 | 0.152440493 | -2.713681916 | -2.548546728 | 1.44E-12 | 11.84033532 | Unpaired t-test | 2.19E-06 | permutation FDR (250 perm) | 1 | 1.21E-10 | FDR correction BH |
| MED26~Q9V4F9 | VMAT~Control | VMAT | Control | 13.40469908 | 0.357720397 | 16.12151275 | 0.131150161 | 0.15210994 | -2.716813665 | -2.321288632 | 1.94E-08 | 7.711862056 | Unpaired t-test | 1.93E-05 | permutation FDR (250 perm) | 1 | 1.26E-07 | FDR correction BH |
| CstF64~Q9VE52 | VMAT~Control | VMAT | Control | 13.56185135 | 0.415150464 | 16.28438827 | 0.096867118 | 0.151507706 | -2.722536918 | -2.286601164 | 5.60E-08 | 7.251633451 | Unpaired t-test | 2.13E-05 | permutation FDR (250 perm) | 1 | 3.00E-07 | FDR correction BH |
| klar~M9PDJ0 | VMAT~Control | VMAT | Control | 13.07706079 | 0.725692893 | 15.84149156 | 0.699687148 | 0.147171398 | -2.764430772 | -1.905426745 | 0.000110915 | 3.955011275 | Unpaired t-test | 0.000215606 | permutation FDR (250 perm) | 1 | 0.000215068 | FDR correction BH |
| CG3950~M9PDW8 | VMAT~Control | VMAT | Control | 16.79364675 | 0.205537456 | 19.56520051 | 0.093360909 | 0.146446564 | -2.771553751 | -2.51740344 | 9.53E-11 | 10.0209942 | Unpaired t-test | 1.31E-05 | permutation FDR (250 perm) | 1 | 2.02E-09 | FDR correction BH |
| Dmel\CG18259~Q9VWQ6 | VMAT~Control | VMAT | Control | 13.37675257 | 0.279191987 | 16.15535953 | 0.506346668 | 0.145732347 | -2.778606961 | -2.207720145 | 8.19E-07 | 6.086527379 | Unpaired t-test | 2.75E-05 | permutation FDR (250 perm) | 1 | 2.83E-06 | FDR correction BH |
| Fmr1~A0A126GUS7 | VMAT~Control | VMAT | Control | 13.82867852 | 0.139073402 | 16.61070622 | 0.489414016 | 0.145387213 | -2.782027703 | -2.266347696 | 2.45E-07 | 6.610914926 | Unpaired t-test | 2.55E-05 | permutation FDR (250 perm) | 1 | 1.02E-06 | FDR correction BH |
| RanGAP~M9PD65 | VMAT~Control | VMAT | Control | 14.73336185 | 0.293687568 | 17.52263208 | 0.123902361 | 0.144659179 | -2.789270227 | -2.441265036 | 2.66E-09 | 8.575428691 | Unpaired t-test | 1.30E-05 | permutation FDR (250 perm) | 1 | 2.67E-08 | FDR correction BH |
| Dmel\CG5676~Q9VL16 | VMAT~Control | VMAT | Control | 13.30045013 | 0.423666367 | 16.09143619 | 0.296367467 | 0.144487234 | -2.79098606 | -2.266835253 | 2.77E-07 | 6.55830392 | Unpaired t-test | 2.55E-05 | permutation FDR (250 perm) | 1 | 1.13E-06 | FDR correction BH |
| Set~P53997 | VMAT~Control | VMAT | Control | 13.54928265 | 0.189350204 | 16.35221751 | 0.141055633 | 0.143295493 | -2.802934858 | -2.535230394 | 1.33E-10 | 9.876701278 | Unpaired t-test | 6.56E-06 | permutation FDR (250 perm) | 1 | 2.65E-09 | FDR correction BH |
| v~P20351 | VMAT~Control | VMAT | Control | 13.30365487 | 0.629842513 | 16.10736079 | 3.601429224 | 0.143218928 | -2.803705923 | -1.064003556 | 0.117158547 | 0.931226022 | Unpaired t-test | 0.011287879 | permutation FDR (250 perm) | 1 | 0.139218183 | FDR correction BH |
| Amph~A0A0B4KEW6 | VMAT~Control | VMAT | Control | 14.00514193 | 0.936435283 | 16.81027687 | 0.20742558 | 0.143077137 | -2.805134939 | -1.96309157 | 6.56E-05 | 4.183272427 | Unpaired t-test | 0.000152577 | permutation FDR (250 perm) | 1 | 0.000134109 | FDR correction BH |
| lig~A0A126GUM8 | VMAT~Control | VMAT | Control | 18.7834999 | 0.16832992 | 21.59141755 | 0.291695696 | 0.142801432 | -2.807917652 | -2.440366755 | 4.26E-09 | 8.370589692 | Unpaired t-test | 1.30E-05 | permutation FDR (250 perm) | 1 | 3.83E-08 | FDR correction BH |
| Prp5~Q9VXW2 | VMAT~Control | VMAT | Control | 14.27672395 | 0.43700842 | 17.08503213 | 0.33866043 | 0.142762782 | -2.808308176 | -2.251596823 | 4.89E-07 | 6.31062738 | Unpaired t-test | 2.76E-05 | permutation FDR (250 perm) | 1 | 1.82E-06 | FDR correction BH |
| hebe~Q7K568 | VMAT~Control | VMAT | Control | 18.05698594 | 0.18184502 | 20.86612179 | 0.122870167 | 0.142680902 | -2.809135853 | -2.558067813 | 6.31E-11 | 10.19998616 | Unpaired t-test | 2.19E-06 | permutation FDR (250 perm) | 1 | 1.52E-09 | FDR correction BH |
| Zasp52~A1ZA47 | VMAT~Control | VMAT | Control | 14.06398631 | 0.538009447 | 16.897673 | 0.219790799 | 0.140273395 | -2.833686685 | -2.249121002 | 7.16E-07 | 6.145075688 | Unpaired t-test | 2.76E-05 | permutation FDR (250 perm) | 1 | 2.51E-06 | FDR correction BH |
| CG13096-RA~H0RNN8 | VMAT~Control | VMAT | Control | 13.03897498 | 0.326409862 | 15.8755189 | 0.246192265 | 0.139995861 | -2.83654392 | -2.398077246 | 2.53E-08 | 7.597113375 | Unpaired t-test | 1.73E-05 | permutation FDR (250 perm) | 1 | 1.56E-07 | FDR correction BH |
| nop5~Q9VM69 | VMAT~Control | VMAT | Control | 16.02993228 | 0.204817847 | 18.86862621 | 0.256578491 | 0.139787384 | -2.838693931 | -2.475270779 | 2.99E-09 | 8.52474349 | Unpaired t-test | 1.31E-05 | permutation FDR (250 perm) | 1 | 2.93E-08 | FDR correction BH |
| Dmel\CG2926~Q9VNE0 | VMAT~Control | VMAT | Control | 17.63060382 | 0.530300672 | 20.47291353 | 0.623562408 | 0.139437479 | -2.842309707 | -2.080641925 | 1.53E-05 | 4.81571259 | Unpaired t-test | 6.27E-05 | permutation FDR (250 perm) | 1 | 3.65E-05 | FDR correction BH |
| futsch~M9NGG5 | VMAT~Control | VMAT | Control | 20.88982155 | 0.164737574 | 23.7452921 | 0.308403308 | 0.138171258 | -2.85547055 | -2.469349182 | 5.21E-09 | 8.283084748 | Unpaired t-test | 1.31E-05 | permutation FDR (250 perm) | 1 | 4.51E-08 | FDR correction BH |
| Pkcdelta~Q9VYN1 | VMAT~Control | VMAT | Control | 13.19321642 | 0.371141275 | 16.0537608 | 0.493347252 | 0.137686175 | -2.860544384 | -2.24164216 | 1.15E-06 | 5.940218354 | Unpaired t-test | 2.76E-05 | permutation FDR (250 perm) | 1 | 3.81E-06 | FDR correction BH |
| Dmel\CG31373~Q8INL3 | VMAT~Control | VMAT | Control | 13.17812844 | 0.489195559 | 16.04285894 | 0.17189514 | 0.137287244 | -2.864730506 | -2.325479666 | 2.22E-07 | 6.653330738 | Unpaired t-test | 1.93E-05 | permutation FDR (250 perm) | 1 | 9.46E-07 | FDR correction BH |
| TfIIS~M9PFV9 | VMAT~Control | VMAT | Control | 13.58991106 | 0.402714247 | 16.46758101 | 0.225021205 | 0.136061429 | -2.877669952 | -2.385519997 | 7.02E-08 | 7.153961547 | Unpaired t-test | 1.73E-05 | permutation FDR (250 perm) | 1 | 3.51E-07 | FDR correction BH |
| xmas~Q9U3V9-1 | VMAT~Control | VMAT | Control | 13.30074716 | 0.407281256 | 16.18095915 | 0.325809632 | 0.135821898 | -2.880211993 | -2.335463098 | 2.23E-07 | 6.651402254 | Unpaired t-test | 1.93E-05 | permutation FDR (250 perm) | 1 | 9.48E-07 | FDR correction BH |
| yuri~Q8IP47 | VMAT~Control | VMAT | Control | 13.4785288 | 0.371332214 | 16.36271962 | 0.273886728 | 0.135447829 | -2.884190823 | -2.390840876 | 6.88E-08 | 7.162491765 | Unpaired t-test | 1.73E-05 | permutation FDR (250 perm) | 1 | 3.45E-07 | FDR correction BH |
| anon-WO0140519.237~B7YZI7 | VMAT~Control | VMAT | Control | 13.2804291 | 0.488939541 | 16.17389168 | 0.159037449 | 0.134580139 | -2.893462575 | -2.352529371 | 1.87E-07 | 6.728930917 | Unpaired t-test | 1.72E-05 | permutation FDR (250 perm) | 1 | 8.18E-07 | FDR correction BH |
| wupA~M9NH88 | VMAT~Control | VMAT | Control | 19.53196225 | 0.394564085 | 22.44027321 | 0.528602386 | 0.133202128 | -2.908310965 | -2.245813906 | 1.81E-06 | 5.742245609 | Unpaired t-test | 2.76E-05 | permutation FDR (250 perm) | 1 | 5.64E-06 | FDR correction BH |
| Sps1~O18373 | VMAT~Control | VMAT | Control | 15.9424925 | 0.104620915 | 18.8613345 | 0.270304905 | 0.132233352 | -2.918841999 | -2.583908669 | 6.71E-10 | 9.173071741 | Unpaired t-test | 2.20E-06 | permutation FDR (250 perm) | 1 | 9.19E-09 | FDR correction BH |
| Dera~Q7K209 | VMAT~Control | VMAT | Control | 14.20890557 | 0.841996153 | 17.14765637 | 0.660513429 | 0.1304211 | -2.938750807 | -1.987537774 | 0.00010969 | 3.959834225 | Unpaired t-test | 0.000119959 | permutation FDR (250 perm) | 1 | 0.00021322 | FDR correction BH |
| Dmel\CG2841~Q9W4Z1 | VMAT~Control | VMAT | Control | 13.73172736 | 0.593585728 | 16.6723019 | 0.537542305 | 0.130256337 | -2.940574538 | -2.165159693 | 9.37E-06 | 5.028313321 | Unpaired t-test | 3.17E-05 | permutation FDR (250 perm) | 1 | 2.38E-05 | FDR correction BH |
| CG4751~Q9VKJ1 | VMAT~Control | VMAT | Control | 13.60116182 | 0.307196265 | 16.54290835 | 0.165589114 | 0.130150564 | -2.94174653 | -2.544609243 | 3.83E-09 | 8.417018053 | Unpaired t-test | 2.19E-06 | permutation FDR (250 perm) | 1 | 3.51E-08 | FDR correction BH |
| Dmel\CG14480~Q7JWU9 | VMAT~Control | VMAT | Control | 13.39217787 | 0.454904994 | 16.33690291 | 0.293523146 | 0.129882139 | -2.944725041 | -2.37073776 | 2.57E-07 | 6.589416856 | Unpaired t-test | 1.72E-05 | permutation FDR (250 perm) | 1 | 1.06E-06 | FDR correction BH |
| CRMP~Q8IPQ2 | VMAT~Control | VMAT | Control | 16.9147311 | 0.053034849 | 19.86687968 | 0.061834613 | 0.129215534 | -2.952148582 | -2.848378328 | 2.00E-15 | 14.69855855 | Unpaired t-test | 0 | permutation FDR (250 perm) | 1 | 1.18E-12 | FDR correction BH |
| Psi~A1ZAK8 | VMAT~Control | VMAT | Control | 13.48751238 | 0.427123901 | 16.44750082 | 0.406616221 | 0.128515258 | -2.959988437 | -2.342260509 | 5.47E-07 | 6.262309323 | Unpaired t-test | 1.93E-05 | permutation FDR (250 perm) | 1 | 2.01E-06 | FDR correction BH |
| CG6239~M9NE59 | VMAT~Control | VMAT | Control | 13.9911577 | 0.340869391 | 16.9548197 | 0.898631178 | 0.128188434 | -2.963662001 | -2.072750543 | 4.22E-05 | 4.374743042 | Unpaired t-test | 6.68E-05 | permutation FDR (250 perm) | 1 | 9.08E-05 | FDR correction BH |
| Mp20~P14318 | VMAT~Control | VMAT | Control | 16.23269911 | 0.291926188 | 19.20597688 | 0.259861617 | 0.12733688 | -2.973277775 | -2.5309123 | 1.04E-08 | 7.983851355 | Unpaired t-test | 1.31E-05 | permutation FDR (250 perm) | 1 | 7.63E-08 | FDR correction BH |
| Su(var)205~P05205 | VMAT~Control | VMAT | Control | 13.24874981 | 0.538970062 | 16.22658208 | 0.118970819 | 0.12693552 | -2.977832268 | -2.388308989 | 2.78E-07 | 6.556124641 | Unpaired t-test | 1.73E-05 | permutation FDR (250 perm) | 1 | 1.13E-06 | FDR correction BH |
| Dmel\CG6106~Q9VWW1 | VMAT~Control | VMAT | Control | 13.49902122 | 0.453506601 | 16.47784177 | 0.316775174 | 0.126848596 | -2.978820543 | -2.388038266 | 2.83E-07 | 6.548299807 | Unpaired t-test | 1.73E-05 | permutation FDR (250 perm) | 1 | 1.14E-06 | FDR correction BH |
| CrebB~A0A1Z1CK36 | VMAT~Control | VMAT | Control | 13.43537928 | 0.244154262 | 16.42797564 | 0.250341894 | 0.125643126 | -2.992596355 | -2.587887819 | 3.30E-09 | 8.481049305 | Unpaired t-test | 2.20E-06 | permutation FDR (250 perm) | 1 | 3.13E-08 | FDR correction BH |
| anon-EST:fe2A9~Q59DP4 | VMAT~Control | VMAT | Control | 14.75142348 | 0.18851926 | 17.75027143 | 0.622565929 | 0.125099858 | -2.998847941 | -2.323059046 | 1.20E-06 | 5.920045414 | Unpaired t-test | 1.93E-05 | permutation FDR (250 perm) | 1 | 3.95E-06 | FDR correction BH |
| Zasp66~Q95TZ7 | VMAT~Control | VMAT | Control | 16.21574492 | 0.319641889 | 19.21748665 | 0.227821058 | 0.124849182 | -3.001741727 | -2.553497918 | 9.87E-09 | 8.005793331 | Unpaired t-test | 2.19E-06 | permutation FDR (250 perm) | 1 | 7.40E-08 | FDR correction BH |
| Sf3b2~Q9VQK7 | VMAT~Control | VMAT | Control | 13.3828092 | 0.472103732 | 16.39620563 | 0.356052186 | 0.12384466 | -3.01339643 | -2.383177646 | 4.74E-07 | 6.323855856 | Unpaired t-test | 1.73E-05 | permutation FDR (250 perm) | 1 | 1.78E-06 | FDR correction BH |
| stwl~M9PI58 | VMAT~Control | VMAT | Control | 13.32136208 | 0.247965827 | 16.33662839 | 0.490561835 | 0.123684249 | -3.015266308 | -2.420306075 | 2.38E-07 | 6.624115213 | Unpaired t-test | 1.30E-05 | permutation FDR (250 perm) | 1 | 9.94E-07 | FDR correction BH |
| HmgZ~A0A0B4LG36 | VMAT~Control | VMAT | Control | 13.73004026 | 0.774610507 | 16.75528243 | 0.19495831 | 0.122831955 | -3.025242169 | -2.22899932 | 7.13E-06 | 5.147165703 | Unpaired t-test | 2.76E-05 | permutation FDR (250 perm) | 1 | 1.89E-05 | FDR correction BH |
| Sf3b6~Q9VRV7 | VMAT~Control | VMAT | Control | 13.4110699 | 0.361510688 | 16.43716801 | 0.218425793 | 0.1227591 | -3.026098118 | -2.545310798 | 1.85E-08 | 7.731709754 | Unpaired t-test | 2.19E-06 | permutation FDR (250 perm) | 1 | 1.23E-07 | FDR correction BH |
| Men-b~Q9VB69 | VMAT~Control | VMAT | Control | 14.16546121 | 0.121537927 | 17.19568892 | 0.135806931 | 0.122408215 | -3.030227713 | -2.801863019 | 4.72E-12 | 11.32618079 | Unpaired t-test | 0 | permutation FDR (250 perm) | 1 | 2.37E-10 | FDR correction BH |
| Hrb98DE~A4V3J6 | VMAT~Control | VMAT | Control | 13.47802237 | 0.470330045 | 16.51909161 | 0.130486642 | 0.121491792 | -3.041069243 | -2.496193003 | 7.10E-08 | 7.148995036 | Unpaired t-test | 1.31E-05 | permutation FDR (250 perm) | 1 | 3.54E-07 | FDR correction BH |
| Rtf1~H0RNK7 | VMAT~Control | VMAT | Control | 13.20710211 | 0.439546988 | 16.25353984 | 0.281757366 | 0.121040543 | -3.046437728 | -2.469770173 | 1.32E-07 | 6.877794272 | Unpaired t-test | 1.31E-05 | permutation FDR (250 perm) | 1 | 6.14E-07 | FDR correction BH |
| east~Q7KW02 | VMAT~Control | VMAT | Control | 13.11452336 | 0.53210217 | 16.16146933 | 0.524150005 | 0.12099791 | -3.046945967 | -2.284023385 | 3.67E-06 | 5.435804272 | Unpaired t-test | 2.35E-05 | permutation FDR (250 perm) | 1 | 1.05E-05 | FDR correction BH |
| hfp~Q8T6B9 | VMAT~Control | VMAT | Control | 13.18979615 | 0.321617758 | 16.24071267 | 0.181331533 | 0.12066536 | -3.050916524 | -2.618548142 | 4.65E-09 | 8.332744513 | Unpaired t-test | 0 | permutation FDR (250 perm) | 1 | 4.12E-08 | FDR correction BH |
| grh~A0A0B4LFT7 | VMAT~Control | VMAT | Control | 13.3235361 | 0.398896172 | 16.37671849 | 0.398839474 | 0.120475994 | -3.053182393 | -2.438125765 | 2.70E-07 | 6.569338691 | Unpaired t-test | 1.30E-05 | permutation FDR (250 perm) | 1 | 1.10E-06 | FDR correction BH |
| Phax~Q6NL95 | VMAT~Control | VMAT | Control | 13.60328518 | 0.243212908 | 16.65680948 | 0.289421258 | 0.120447445 | -3.0535243 | -2.611934091 | 5.80E-09 | 8.236455952 | Unpaired t-test | 0 | permutation FDR (250 perm) | 1 | 4.89E-08 | FDR correction BH |
| Nab2~Q9V3H9 | VMAT~Control | VMAT | Control | 13.2122823 | 0.458188245 | 16.27306467 | 0.430934031 | 0.119843007 | -3.060782369 | -2.388815289 | 7.29E-07 | 6.137049116 | Unpaired t-test | 1.73E-05 | permutation FDR (250 perm) | 1 | 2.55E-06 | FDR correction BH |
| san~Q9NHD5 | VMAT~Control | VMAT | Control | 13.32100352 | 0.388435565 | 16.38339808 | 0.330168021 | 0.119709159 | -3.06239456 | -2.493830596 | 1.01E-07 | 6.997796484 | Unpaired t-test | 1.31E-05 | permutation FDR (250 perm) | 1 | 4.83E-07 | FDR correction BH |
| Edc3~Q9VVI2 | VMAT~Control | VMAT | Control | 14.42654445 | 0.448019661 | 17.49679035 | 0.208358912 | 0.119059455 | -3.070245897 | -2.514599091 | 7.28E-08 | 7.137884179 | Unpaired t-test | 1.31E-05 | permutation FDR (250 perm) | 1 | 3.62E-07 | FDR correction BH |
| CtBP~O46036 | VMAT~Control | VMAT | Control | 13.64546328 | 0.265864366 | 16.75375084 | 0.246027819 | 0.115961069 | -3.108287556 | -2.674955429 | 3.22E-09 | 8.492422182 | Unpaired t-test | 0 | permutation FDR (250 perm) | 1 | 3.10E-08 | FDR correction BH |
| fs(1)h~M9PJC7 | VMAT~Control | VMAT | Control | 13.46810235 | 0.479262801 | 16.58503767 | 0.112975565 | 0.115268057 | -3.116935321 | -2.554430939 | 6.10E-08 | 7.214839602 | Unpaired t-test | 2.19E-06 | permutation FDR (250 perm) | 1 | 3.16E-07 | FDR correction BH |
| mahe~Q9W3M7 | VMAT~Control | VMAT | Control | 13.86366165 | 0.712649575 | 16.98904435 | 0.655275426 | 0.114595103 | -3.1253827 | -2.181073454 | 2.86E-05 | 4.543303246 | Unpaired t-test | 3.17E-05 | permutation FDR (250 perm) | 1 | 6.39E-05 | FDR correction BH |
| Dlg5~M9PCQ6 | VMAT~Control | VMAT | Control | 12.97928255 | 0.304551215 | 16.11190443 | 0.186121168 | 0.114021526 | -3.132621884 | -2.701421333 | 2.58E-09 | 8.588137833 | Unpaired t-test | 0 | permutation FDR (250 perm) | 1 | 2.63E-08 | FDR correction BH |
| Dmel\CG8086~M9PCX6 | VMAT~Control | VMAT | Control | 13.51639129 | 0.430551181 | 16.65078548 | 0.208013923 | 0.11388154 | -3.134394191 | -2.582207278 | 4.36E-08 | 7.360069212 | Unpaired t-test | 2.19E-06 | permutation FDR (250 perm) | 1 | 2.45E-07 | FDR correction BH |
| larp~A0A0B4K7Y7 | VMAT~Control | VMAT | Control | 14.45545049 | 0.255220561 | 17.59055078 | 0.468395022 | 0.113825816 | -3.13510029 | -2.531265847 | 1.24E-07 | 6.907682914 | Unpaired t-test | 6.56E-06 | permutation FDR (250 perm) | 1 | 5.79E-07 | FDR correction BH |
| eIF4EHP~Q8T3K5 | VMAT~Control | VMAT | Control | 13.24084579 | 0.479639472 | 16.40504222 | 0.281707749 | 0.111553182 | -3.164196424 | -2.533866178 | 1.69E-07 | 6.773108787 | Unpaired t-test | 6.56E-06 | permutation FDR (250 perm) | 1 | 7.55E-07 | FDR correction BH |
| Rnp4F~A8JUY0 | VMAT~Control | VMAT | Control | 13.37579916 | 0.589608293 | 16.54158955 | 0.382593514 | 0.111430001 | -3.165790383 | -2.408672978 | 1.49E-06 | 5.82712568 | Unpaired t-test | 1.73E-05 | permutation FDR (250 perm) | 1 | 4.78E-06 | FDR correction BH |
| Cdep~A8JQT5 | VMAT~Control | VMAT | Control | 15.37569427 | 0.20588954 | 18.54309301 | 0.148822084 | 0.111305845 | -3.167398745 | -2.84425683 | 8.18E-11 | 10.08713055 | Unpaired t-test | 0 | permutation FDR (250 perm) | 1 | 1.83E-09 | FDR correction BH |
| bt~L0MN91 | VMAT~Control | VMAT | Control | 13.08018153 | 0.291333704 | 16.25868136 | 0.305382854 | 0.110452668 | -3.178499825 | -2.673815456 | 1.14E-08 | 7.941235783 | Unpaired t-test | 0 | permutation FDR (250 perm) | 1 | 8.24E-08 | FDR correction BH |
| Uba2~Q7KUA4 | VMAT~Control | VMAT | Control | 13.67601877 | 0.606664324 | 16.8619301 | 0.130564788 | 0.109886698 | -3.185911334 | -2.493823599 | 4.43E-07 | 6.353929059 | Unpaired t-test | 1.31E-05 | permutation FDR (250 perm) | 1 | 1.67E-06 | FDR correction BH |
| Bap170~A1Z6M0 | VMAT~Control | VMAT | Control | 13.66941625 | 0.268683165 | 16.88094941 | 0.153615847 | 0.107952371 | -3.211533162 | -2.821065723 | 5.00E-10 | 9.300945121 | Unpaired t-test | 0 | permutation FDR (250 perm) | 1 | 7.18E-09 | FDR correction BH |
| ZAP3~M9NDT5 | VMAT~Control | VMAT | Control | 13.52277232 | 0.31723773 | 16.73727285 | 0.381863019 | 0.10773056 | -3.214500534 | -2.630486216 | 4.91E-08 | 7.308944512 | Unpaired t-test | 0 | permutation FDR (250 perm) | 1 | 2.68E-07 | FDR correction BH |
| Nup44A~Q7K2X8 | VMAT~Control | VMAT | Control | 12.79790465 | 0.344134704 | 16.0186114 | 0.458505364 | 0.107268118 | -3.220706749 | -2.563479527 | 1.90E-07 | 6.721898958 | Unpaired t-test | 2.19E-06 | permutation FDR (250 perm) | 1 | 8.30E-07 | FDR correction BH |
| mdcds_23070~A1Z8Z9 | VMAT~Control | VMAT | Control | 15.05277042 | 0.704726151 | 18.27944961 | 0.194418616 | 0.10682497 | -3.226679188 | -2.431675563 | 1.79E-06 | 5.74646732 | Unpaired t-test | 1.30E-05 | permutation FDR (250 perm) | 1 | 5.60E-06 | FDR correction BH |
| 4E-T~Q8IH18-2 | VMAT~Control | VMAT | Control | 14.02063774 | 0.50013648 | 17.24794819 | 0.279381339 | 0.106778238 | -3.227310453 | -2.56910701 | 1.85E-07 | 6.733219101 | Unpaired t-test | 2.19E-06 | permutation FDR (250 perm) | 1 | 8.12E-07 | FDR correction BH |
| wupA~E4NKJ3 | VMAT~Control | VMAT | Control | 19.78123284 | 0.213247458 | 23.01092794 | 0.224617946 | 0.106601889 | -3.229695097 | -2.836768911 | 4.77E-10 | 9.321927272 | Unpaired t-test | 0 | permutation FDR (250 perm) | 1 | 6.92E-09 | FDR correction BH |
| EMSY~Q9VQ58 | VMAT~Control | VMAT | Control | 13.16098083 | 0.244760469 | 16.39147685 | 0.434893809 | 0.106542724 | -3.230496022 | -2.641069522 | 4.92E-08 | 7.307922548 | Unpaired t-test | 0 | permutation FDR (250 perm) | 1 | 2.68E-07 | FDR correction BH |
| mdlc~Q9VDP8 | VMAT~Control | VMAT | Control | 13.57778667 | 0.55411494 | 16.80870403 | 0.190259041 | 0.106511613 | -3.230917361 | -2.560139454 | 2.26E-07 | 6.645832193 | Unpaired t-test | 2.19E-06 | permutation FDR (250 perm) | 1 | 9.55E-07 | FDR correction BH |
| Rcd1~A1Z9L2 | VMAT~Control | VMAT | Control | 13.54158542 | 0.291168905 | 16.78224358 | 0.21555605 | 0.105794889 | -3.240658161 | -2.788827273 | 2.14E-09 | 8.668836 | Unpaired t-test | 0 | permutation FDR (250 perm) | 1 | 2.26E-08 | FDR correction BH |
| Dmel\CG12065~Q8MRM6 | VMAT~Control | VMAT | Control | 13.59175146 | 0.649839636 | 16.83245245 | 0.101080341 | 0.105791748 | -3.240700993 | -2.504189332 | 6.49E-07 | 6.187787229 | Unpaired t-test | 1.31E-05 | permutation FDR (250 perm) | 1 | 2.31E-06 | FDR correction BH |
| FoxK~Q0E8F8 | VMAT~Control | VMAT | Control | 13.34221715 | 0.417727623 | 16.5992031 | 0.261027483 | 0.104604299 | -3.256985952 | -2.669032834 | 4.02E-08 | 7.396229646 | Unpaired t-test | 0 | permutation FDR (250 perm) | 1 | 2.28E-07 | FDR correction BH |
| Dmel\CG8223~A0A126BEE6 | VMAT~Control | VMAT | Control | 13.42116018 | 0.24803411 | 16.6799183 | 0.295134433 | 0.104475885 | -3.258758113 | -2.779539055 | 3.73E-09 | 8.42876773 | Unpaired t-test | 0 | permutation FDR (250 perm) | 1 | 3.45E-08 | FDR correction BH |
| by~Q9VHC3 | VMAT~Control | VMAT | Control | 14.13864966 | 0.344017341 | 17.41514947 | 0.20128308 | 0.10319895 | -3.276499802 | -2.780822158 | 4.89E-09 | 8.310806066 | Unpaired t-test | 0 | permutation FDR (250 perm) | 1 | 4.29E-08 | FDR correction BH |
| Myo10A~Q0KHU0 | VMAT~Control | VMAT | Control | 13.37531634 | 0.970185689 | 16.65969097 | 0.684031935 | 0.102637182 | -3.284374625 | -2.145418205 | 0.000103237 | 3.986166144 | Unpaired t-test | 3.58E-05 | permutation FDR (250 perm) | 1 | 0.000202854 | FDR correction BH |
| tyf~Q9W4M7 | VMAT~Control | VMAT | Control | 13.22844423 | 0.776706653 | 16.52079818 | 0.456712454 | 0.102071079 | -3.292353951 | -2.346729852 | 9.78E-06 | 5.009489441 | Unpaired t-test | 1.93E-05 | permutation FDR (250 perm) | 1 | 2.46E-05 | FDR correction BH |
| ldbr~Q9VSD7 | VMAT~Control | VMAT | Control | 14.30365034 | 1.193342073 | 17.60425323 | 0.399896476 | 0.101489129 | -3.30060289 | -2.111917131 | 0.000158551 | 3.799832323 | Unpaired t-test | 5.03E-05 | permutation FDR (250 perm) | 1 | 0.000298344 | FDR correction BH |
| cora~A0A0B4LFX4 | VMAT~Control | VMAT | Control | 13.80843385 | 0.288430631 | 17.11000947 | 0.152297077 | 0.101420724 | -3.301575623 | -2.88128919 | 6.38E-10 | 9.19501119 | Unpaired t-test | 0 | permutation FDR (250 perm) | 1 | 8.94E-09 | FDR correction BH |
| Larp7~Q9W362 | VMAT~Control | VMAT | Control | 13.99043275 | 0.837849187 | 17.29676437 | 0.397763367 | 0.10108693 | -3.306331613 | -2.336995569 | 1.22E-05 | 4.915382661 | Unpaired t-test | 1.93E-05 | permutation FDR (250 perm) | 1 | 2.98E-05 | FDR correction BH |
| Dmel\CG1674~Q8IMB8 | VMAT~Control | VMAT | Control | 13.62008238 | 0.399136458 | 16.92675365 | 0.678871923 | 0.101063134 | -3.306671274 | -2.445425267 | 2.82E-06 | 5.549108586 | Unpaired t-test | 1.30E-05 | permutation FDR (250 perm) | 1 | 8.33E-06 | FDR correction BH |
| vig~Q9V426 | VMAT~Control | VMAT | Control | 15.94197958 | 1.149052409 | 19.26006318 | 1.006031264 | 0.100266836 | -3.318083593 | -1.971533399 | 0.000663267 | 3.178311552 | Unpaired t-test | 0.000134228 | permutation FDR (250 perm) | 1 | 0.001089554 | FDR correction BH |
| Not3~Q7K126 | VMAT~Control | VMAT | Control | 13.60835812 | 0.332476032 | 16.92725061 | 0.233715764 | 0.100210634 | -3.31889249 | -2.808458211 | 5.21E-09 | 8.282938388 | Unpaired t-test | 0 | permutation FDR (250 perm) | 1 | 4.51E-08 | FDR correction BH |
| Zn72D~Q86BI3 | VMAT~Control | VMAT | Control | 12.6268676 | 0.847996458 | 15.96297449 | 0.864710737 | 0.099022015 | -3.336106889 | -2.164010784 | 0.000106998 | 3.970624144 | Unpaired t-test | 3.17E-05 | permutation FDR (250 perm) | 1 | 0.000209197 | FDR correction BH |
| Nup62~Q7JXF5 | VMAT~Control | VMAT | Control | 14.02568683 | 0.477390033 | 17.3862432 | 0.293284708 | 0.097358019 | -3.360556372 | -2.687227839 | 1.02E-07 | 6.991513839 | Unpaired t-test | 0 | permutation FDR (250 perm) | 1 | 4.87E-07 | FDR correction BH |
| Top3beta~X2JEB4 | VMAT~Control | VMAT | Control | 13.29284261 | 0.54913999 | 16.66779308 | 0.821974147 | 0.096391486 | -3.374950465 | -2.340326822 | 1.77E-05 | 4.752029852 | Unpaired t-test | 1.93E-05 | permutation FDR (250 perm) | 1 | 4.15E-05 | FDR correction BH |
| Dmel\CG1737~X2JEU3 | VMAT~Control | VMAT | Control | 13.22736981 | 0.32749629 | 16.62121912 | 0.379389361 | 0.095137023 | -3.393849314 | -2.772439227 | 3.19E-08 | 7.495662703 | Unpaired t-test | 0 | permutation FDR (250 perm) | 1 | 1.89E-07 | FDR correction BH |
| Trm1~Q9VK89 | VMAT~Control | VMAT | Control | 13.25786566 | 0.114090465 | 16.65594018 | 0.49408616 | 0.094858804 | -3.398074518 | -2.769921568 | 3.53E-08 | 7.452007705 | Unpaired t-test | 0 | permutation FDR (250 perm) | 1 | 2.05E-07 | FDR correction BH |
| CG12562~Q9VP02 | VMAT~Control | VMAT | Control | 13.05257026 | 0.52850701 | 16.45153824 | 0.423758669 | 0.094800075 | -3.398967986 | -2.608672697 | 5.48E-07 | 6.260904042 | Unpaired t-test | 0 | permutation FDR (250 perm) | 1 | 2.01E-06 | FDR correction BH |
| wupA~P36188-4 | VMAT~Control | VMAT | Control | 17.2677592 | 0.4459932 | 20.66676875 | 0.523289757 | 0.094797344 | -3.399009546 | -2.599650203 | 6.30E-07 | 6.200800961 | Unpaired t-test | 0 | permutation FDR (250 perm) | 1 | 2.26E-06 | FDR correction BH |
| ps~A0A0B4KGY6-7 | VMAT~Control | VMAT | Control | 15.85995311 | 0.222443751 | 19.27365787 | 0.104062153 | 0.093836645 | -3.413704753 | -3.075888314 | 2.79E-11 | 10.55455535 | Unpaired t-test | 0 | permutation FDR (250 perm) | 1 | 8.01E-10 | FDR correction BH |
| EndoGI~Q9V3V9 | VMAT~Control | VMAT | Control | 15.82506521 | 0.239278407 | 19.23945731 | 0.109990497 | 0.093791949 | -3.414392104 | -3.054638969 | 5.56E-11 | 10.25513079 | Unpaired t-test | 0 | permutation FDR (250 perm) | 1 | 1.39E-09 | FDR correction BH |
| EG:152A3.3~O46052 | VMAT~Control | VMAT | Control | 13.05925238 | 0.219464736 | 16.47907923 | 0.207991603 | 0.093439292 | -3.419826854 | -3.012473434 | 2.14E-10 | 9.669342289 | Unpaired t-test | 0 | permutation FDR (250 perm) | 1 | 3.79E-09 | FDR correction BH |
| Gug~Q8IQA6 | VMAT~Control | VMAT | Control | 13.34302062 | 0.243872511 | 16.77294947 | 0.357307214 | 0.092787299 | -3.429928849 | -2.873926555 | 6.95E-09 | 8.15809473 | Unpaired t-test | 0 | permutation FDR (250 perm) | 1 | 5.67E-08 | FDR correction BH |
| PIP82~Q9W3E2 | VMAT~Control | VMAT | Control | 13.5244038 | 0.406093555 | 16.96060918 | 0.116000655 | 0.092384499 | -3.436205382 | -2.890300876 | 5.40E-09 | 8.26721307 | Unpaired t-test | 0 | permutation FDR (250 perm) | 1 | 4.63E-08 | FDR correction BH |
| sals~Q58CJ5 | VMAT~Control | VMAT | Control | 13.29022155 | 0.291058103 | 16.74562467 | 0.348547658 | 0.091163295 | -3.455403125 | -2.872139199 | 1.04E-08 | 7.98489569 | Unpaired t-test | 0 | permutation FDR (250 perm) | 1 | 7.63E-08 | FDR correction BH |
| D1~A4V2K7 | VMAT~Control | VMAT | Control | 14.87665454 | 0.528680759 | 18.33851943 | 0.112376478 | 0.090755891 | -3.461864891 | -2.787969549 | 5.45E-08 | 7.263732827 | Unpaired t-test | 0 | permutation FDR (250 perm) | 1 | 2.93E-07 | FDR correction BH |
| sls~M9PDZ6 | VMAT~Control | VMAT | Control | 14.33353894 | 0.39243929 | 17.80327702 | 0.714245402 | 0.09026196 | -3.469738081 | -2.542938845 | 2.49E-06 | 5.603828193 | Unpaired t-test | 2.19E-06 | permutation FDR (250 perm) | 1 | 7.44E-06 | FDR correction BH |
| Spf45~Q7PL82 | VMAT~Control | VMAT | Control | 13.1058367 | 0.369315219 | 16.58706769 | 0.110465524 | 0.089545765 | -3.48123099 | -2.969338461 | 1.95E-09 | 8.709268316 | Unpaired t-test | 0 | permutation FDR (250 perm) | 1 | 2.09E-08 | FDR correction BH |
| Patj~A4V1B2 | VMAT~Control | VMAT | Control | 12.97783933 | 0.053622678 | 16.46097023 | 0.077662482 | 0.089427918 | -3.483130898 | -3.342074353 | 1.67E-15 | 14.7777466 | Unpaired t-test | 0 | permutation FDR (250 perm) | 1 | 1.18E-12 | FDR correction BH |
| mip120~E1JH59 | VMAT~Control | VMAT | Control | 13.91124194 | 0.690270602 | 17.394886 | 0.601993257 | 0.089396114 | -3.483644064 | -2.471366972 | 6.86E-06 | 5.163568974 | Unpaired t-test | 1.31E-05 | permutation FDR (250 perm) | 1 | 1.82E-05 | FDR correction BH |
| spoon~Q9W4C4 | VMAT~Control | VMAT | Control | 13.23630782 | 0.328183513 | 16.72135649 | 0.292855335 | 0.089309121 | -3.485048665 | -2.912198227 | 6.99E-09 | 8.15522176 | Unpaired t-test | 0 | permutation FDR (250 perm) | 1 | 5.68E-08 | FDR correction BH |
| Cdc5~Q9W0R0 | VMAT~Control | VMAT | Control | 13.5272348 | 0.321868225 | 17.03418122 | 0.137138632 | 0.087963791 | -3.506946415 | -3.032471401 | 7.03E-10 | 9.153262203 | Unpaired t-test | 0 | permutation FDR (250 perm) | 1 | 9.37E-09 | FDR correction BH |
| Marc~A1Z803 | VMAT~Control | VMAT | Control | 13.26531864 | 0.179813072 | 16.77778367 | 1.002381738 | 0.087627954 | -3.512465023 | -2.413344997 | 1.62E-05 | 4.790377463 | Unpaired t-test | 1.51E-05 | permutation FDR (250 perm) | 1 | 3.85E-05 | FDR correction BH |
| eEF1delta~Q9VL18 | VMAT~Control | VMAT | Control | 14.13782984 | 0.408798408 | 17.66444308 | 0.351265838 | 0.086772804 | -3.526613242 | -2.841657312 | 4.44E-08 | 7.352475741 | Unpaired t-test | 0 | permutation FDR (250 perm) | 1 | 2.47E-07 | FDR correction BH |
| Sdb~A1Z9M5 | VMAT~Control | VMAT | Control | 13.00500718 | 0.613224738 | 16.54924277 | 0.886019764 | 0.085719332 | -3.544235582 | -2.391704283 | 2.44E-05 | 4.612663933 | Unpaired t-test | 1.73E-05 | permutation FDR (250 perm) | 1 | 5.55E-05 | FDR correction BH |
| Papss~Q961A8 | VMAT~Control | VMAT | Control | 12.55149868 | 0.65138998 | 16.10083366 | 0.66343804 | 0.08541688 | -3.549334981 | -2.506942016 | 6.64E-06 | 5.177685659 | Unpaired t-test | 1.31E-05 | permutation FDR (250 perm) | 1 | 1.77E-05 | FDR correction BH |
| tral~M9PFB2 | VMAT~Control | VMAT | Control | 14.83976786 | 0.361665928 | 18.39030391 | 0.135356582 | 0.085345799 | -3.550536049 | -3.027663479 | 1.64E-09 | 8.785506962 | Unpaired t-test | 0 | permutation FDR (250 perm) | 1 | 1.82E-08 | FDR correction BH |
| BEAF-32~Q94513 | VMAT~Control | VMAT | Control | 13.35373987 | 0.261905957 | 16.92567773 | 0.141325129 | 0.084089073 | -3.571937853 | -3.152380892 | 1.19E-10 | 9.923863273 | Unpaired t-test | 0 | permutation FDR (250 perm) | 1 | 2.42E-09 | FDR correction BH |
| Dmel\CG14712~Q9VGL0 | VMAT~Control | VMAT | Control | 13.75154294 | 0.496045043 | 17.35602008 | 0.433012198 | 0.082213713 | -3.604477138 | -2.784521581 | 2.42E-07 | 6.615579134 | Unpaired t-test | 0 | permutation FDR (250 perm) | 1 | 1.01E-06 | FDR correction BH |
| Nlp~A0A0B4KI24 | VMAT~Control | VMAT | Control | 14.34087629 | 0.122194154 | 17.95064502 | 0.163132427 | 0.081912718 | -3.609768729 | -3.308217492 | 2.52E-12 | 11.59867509 | Unpaired t-test | 0 | permutation FDR (250 perm) | 1 | 1.59E-10 | FDR correction BH |
| Fim~Q8IQZ7 | VMAT~Control | VMAT | Control | 13.48644456 | 0.265740212 | 17.10141694 | 0.416244312 | 0.081617799 | -3.614972383 | -2.961025179 | 1.51E-08 | 7.821900995 | Unpaired t-test | 0 | permutation FDR (250 perm) | 1 | 1.03E-07 | FDR correction BH |
| sordd1~Q9VXP1 | VMAT~Control | VMAT | Control | 14.09762869 | 1.465519299 | 17.71421238 | 0.3303727 | 0.081526693 | -3.616583694 | -2.163226288 | 0.000308754 | 3.510387403 | Unpaired t-test | 3.17E-05 | permutation FDR (250 perm) | 1 | 0.000547769 | FDR correction BH |
| mor~A0A0B4JDA0 | VMAT~Control | VMAT | Control | 13.38714711 | 0.178682061 | 17.03720882 | 0.098250802 | 0.079656632 | -3.650061708 | -3.345020424 | 2.27E-12 | 11.64468954 | Unpaired t-test | 0 | permutation FDR (250 perm) | 1 | 1.59E-10 | FDR correction BH |
| Dmel\CG2681~Q9W4W4 | VMAT~Control | VMAT | Control | 14.79542851 | 0.474887795 | 18.44580005 | 0.256701633 | 0.079639527 | -3.650371545 | -2.940484557 | 3.24E-08 | 7.48980131 | Unpaired t-test | 0 | permutation FDR (250 perm) | 1 | 1.91E-07 | FDR correction BH |
| TpnC4~Q7K860 | VMAT~Control | VMAT | Control | 13.70833754 | 0.465987043 | 17.37791233 | 0.340915282 | 0.078586494 | -3.669574795 | -2.916499627 | 5.87E-08 | 7.231515331 | Unpaired t-test | 0 | permutation FDR (250 perm) | 1 | 3.09E-07 | FDR correction BH |
| Bap111~Q9W384 | VMAT~Control | VMAT | Control | 13.21152669 | 0.185427236 | 16.89287338 | 0.169195577 | 0.077947864 | -3.681346692 | -3.309792708 | 1.64E-11 | 10.78518032 | Unpaired t-test | 0 | permutation FDR (250 perm) | 1 | 5.51E-10 | FDR correction BH |
| Map205~Q8IMF5 | VMAT~Control | VMAT | Control | 14.42423679 | 0.548340708 | 18.11430264 | 0.13142082 | 0.077478195 | -3.690065848 | -2.946936405 | 4.43E-08 | 7.353180698 | Unpaired t-test | 0 | permutation FDR (250 perm) | 1 | 2.47E-07 | FDR correction BH |
| Nuak~Q9VH03 | VMAT~Control | VMAT | Control | 12.75821333 | 0.718304629 | 16.44971972 | 0.650785498 | 0.077400871 | -3.69150639 | -2.57522316 | 6.78E-06 | 5.168656211 | Unpaired t-test | 2.19E-06 | permutation FDR (250 perm) | 1 | 1.81E-05 | FDR correction BH |
| eIF6~P56538 | VMAT~Control | VMAT | Control | 13.30770726 | 0.486132421 | 17.01570346 | 0.677329428 | 0.076521226 | -3.707996202 | -2.70094028 | 1.67E-06 | 5.776743532 | Unpaired t-test | 0 | permutation FDR (250 perm) | 1 | 5.27E-06 | FDR correction BH |
| Slik~Q8MLP1 | VMAT~Control | VMAT | Control | 14.43204926 | 1.326707709 | 18.15460577 | 0.397563768 | 0.075752824 | -3.722556508 | -2.298742155 | 0.000130351 | 3.884886329 | Unpaired t-test | 1.92E-05 | permutation FDR (250 perm) | 1 | 0.000249232 | FDR correction BH |
| Usp10~Q9W0L7 | VMAT~Control | VMAT | Control | 13.4230703 | 0.62443471 | 17.15239934 | 0.283625802 | 0.075398047 | -3.72932904 | -2.853978585 | 2.58E-07 | 6.588224921 | Unpaired t-test | 0 | permutation FDR (250 perm) | 1 | 1.06E-06 | FDR correction BH |
| Jupiter~Q9I7K0-5 | VMAT~Control | VMAT | Control | 13.50016498 | 0.396694914 | 17.23045069 | 0.236702805 | 0.075348066 | -3.730285704 | -3.091595984 | 5.82E-09 | 8.235393217 | Unpaired t-test | 0 | permutation FDR (250 perm) | 1 | 4.89E-08 | FDR correction BH |
| IRSp53~Q8IQE6 | VMAT~Control | VMAT | Control | 13.22286577 | 0.874382827 | 17.02094298 | 0.446669268 | 0.071889396 | -3.798077214 | -2.639196978 | 5.91E-06 | 5.228774858 | Unpaired t-test | 0 | permutation FDR (250 perm) | 1 | 1.61E-05 | FDR correction BH |
| kis~Q9VPL6 | VMAT~Control | VMAT | Control | 13.72976158 | 0.72456748 | 17.56075201 | 0.447658664 | 0.070267899 | -3.830990432 | -2.774285346 | 1.51E-06 | 5.821727132 | Unpaired t-test | 0 | permutation FDR (250 perm) | 1 | 4.82E-06 | FDR correction BH |
| Ars2~Q9V9K7 | VMAT~Control | VMAT | Control | 13.42660943 | 0.427844007 | 17.27563432 | 0.273884335 | 0.06939498 | -3.849024887 | -3.136468947 | 1.08E-08 | 7.966806065 | Unpaired t-test | 0 | permutation FDR (250 perm) | 1 | 7.89E-08 | FDR correction BH |

| | | | | | | | | | | | | | | | | | | |
|---|---|---|---|---|---|---|---|---|---|---|---|---|---|---|---|---|---|---|
| TpnC73F~X2JCI6 | VMAT~Control | VMAT | Control | 13.61831351 | 0.192231102 | 17.49015403 | 0.47103375 | 0.068306159 | -3.871840515 | -3.154198812 | 1.03E-08 | 7.985475053 | Unpaired t-test | 0 | permutation FDR (250 perm) | 1 | 7.63E-08 | FDR correction BH |
| REG~Q9V3P3 | VMAT~Control | VMAT | Control | 13.36034587 | 0.223585551 | 17.23614928 | 0.321185304 | 0.068118788 | -3.875803418 | -3.298515201 | 7.91E-10 | 9.101958724 | Unpaired t-test | 0 | permutation FDR (250 perm) | 1 | 1.02E-08 | FDR correction BH |
| Dmel\CG1246~Q9VZZ0 | VMAT~Control | VMAT | Control | 13.46804969 | 0.289805653 | 17.34432051 | 0.486398408 | 0.068096723 | -3.876270818 | -3.093079288 | 2.87E-08 | 7.541465802 | Unpaired t-test | 0 | permutation FDR (250 perm) | 1 | 1.73E-07 | FDR correction BH |
| Nup50~Q7K0D8 | VMAT~Control | VMAT | Control | 16.19694674 | 0.165868931 | 20.0841951 | 0.11669211 | 0.067580538 | -3.887248361 | -3.564004542 | 1.15E-12 | 11.94020937 | Unpaired t-test | 0 | permutation FDR (250 perm) | 1 | 1.04E-10 | FDR correction BH |
| Dmel\CG5023~Q9I7J0 | VMAT~Control | VMAT | Control | 15.55995364 | 0.204288823 | 19.45125352 | 0.18210727 | 0.067391018 | -3.891299878 | -3.46697555 | 2.23E-11 | 10.65197041 | Unpaired t-test | 0 | permutation FDR (250 perm) | 1 | 6.98E-10 | FDR correction BH |
| Ciz1~Q9VT61 | VMAT~Control | VMAT | Control | 13.74663201 | 0.310535878 | 17.64830947 | 0.141401405 | 0.066908 | -3.901677459 | -3.385123017 | 1.92E-10 | 9.716215965 | Unpaired t-test | 0 | permutation FDR (250 perm) | 1 | 3.56E-09 | FDR correction BH |
| Dmel\CG7054~Q9VD02 | VMAT~Control | VMAT | Control | 13.70128309 | 0.717350099 | 17.60459971 | 0.09137608 | 0.066832024 | -3.903316618 | -2.949458913 | 2.77E-07 | 6.558032635 | Unpaired t-test | 0 | permutation FDR (250 perm) | 1 | 1.13E-06 | FDR correction BH |
| CG13657~Q9VBT4 | VMAT~Control | VMAT | Control | 13.57721596 | 0.247354858 | 17.48284468 | 0.132225652 | 0.066725003 | -3.905628723 | -3.470332166 | 2.74E-11 | 10.56209429 | Unpaired t-test | 0 | permutation FDR (250 perm) | 1 | 7.97E-10 | FDR correction BH |
| fln~M9PD14 | VMAT~Control | VMAT | Control | 16.01506356 | 0.429743182 | 19.94335509 | 0.113549612 | 0.065685033 | -3.928291524 | -3.27690054 | 2.41E-09 | 8.617487708 | Unpaired t-test | 0 | permutation FDR (250 perm) | 1 | 2.49E-08 | FDR correction BH |
| how~E1JIT0 | VMAT~Control | VMAT | Control | 13.48767064 | 0.259673041 | 17.48782794 | 0.382551293 | 0.062493186 | -4.000157307 | -3.314755109 | 2.97E-09 | 8.52722608 | Unpaired t-test | 0 | permutation FDR (250 perm) | 1 | 2.93E-08 | FDR correction BH |
| caz~Q27294 | VMAT~Control | VMAT | Control | 13.16624192 | 0.399709012 | 17.16641401 | 0.112483393 | 0.062492545 | -4.000172085 | -3.373683804 | 1.04E-09 | 8.984046982 | Unpaired t-test | 0 | permutation FDR (250 perm) | 1 | 1.25E-08 | FDR correction BH |
| CtBP~A4V2S3 | VMAT~Control | VMAT | Control | 16.19907768 | 0.0875769 | 20.23016156 | 0.095464617 | 0.061167797 | -4.031083882 | -3.810326778 | 9.17E-15 | 14.03770067 | Unpaired t-test | 0 | permutation FDR (250 perm) | 1 | 3.08E-12 | FDR correction BH |
| Patronin~A0A0B4KEY6 | VMAT~Control | VMAT | Control | 13.84430199 | 1.025371221 | 17.88451829 | 0.310569054 | 0.06078182 | -4.040216305 | -2.73147712 | 7.40E-06 | 5.13052619 | Unpaired t-test | 0 | permutation FDR (250 perm) | 1 | 1.94E-05 | FDR correction BH |
| Mical~A0A126GUS4 | VMAT~Control | VMAT | Control | 15.03693734 | 0.678981237 | 19.0850069 | 0.441109664 | 0.060451856 | -4.048069561 | -2.97192597 | 5.67E-07 | 6.246346551 | Unpaired t-test | 0 | permutation FDR (250 perm) | 1 | 2.07E-06 | FDR correction BH |
| cg~A8DYD2 | VMAT~Control | VMAT | Control | 13.36443275 | 0.280330419 | 17.45153763 | 0.258427812 | 0.058838127 | -4.087104876 | -3.491726885 | 3.63E-10 | 9.439882532 | Unpaired t-test | 0 | permutation FDR (250 perm) | 1 | 5.71E-09 | FDR correction BH |
| cno~A0A0B4KG43 | VMAT~Control | VMAT | Control | 11.72029247 | 0.381395443 | 15.80822958 | 0.941511294 | 0.058804195 | -4.087937115 | -2.810946619 | 4.14E-06 | 5.382544689 | Unpaired t-test | 0 | permutation FDR (250 perm) | 1 | 1.17E-05 | FDR correction BH |
| FBpp0072605~M9PBH8 | VMAT~Control | VMAT | Control | 13.75014573 | 0.41283752 | 17.84035519 | 0.398334151 | 0.058711647 | -4.090209465 | -3.25509524 | 1.94E-08 | 7.711363444 | Unpaired t-test | 0 | permutation FDR (250 perm) | 1 | 1.26E-07 | FDR correction BH |
| NAT1~A1Z968 | VMAT~Control | VMAT | Control | 13.67218697 | 0.127922047 | 17.77055335 | 0.499222576 | 0.058380631 | -4.098366381 | -3.330726449 | 6.75E-09 | 8.170805916 | Unpaired t-test | 0 | permutation FDR (250 perm) | 1 | 5.54E-08 | FDR correction BH |
| scrib~E1JIZ0 | VMAT~Control | VMAT | Control | 13.62076602 | 0.215697762 | 17.72579396 | 0.139919866 | 0.058111683 | -4.105027942 | -3.681702315 | 7.06E-12 | 11.15135804 | Unpaired t-test | 0 | permutation FDR (250 perm) | 1 | 3.13E-10 | FDR correction BH |
| nonA~X2JFR1 | VMAT~Control | VMAT | Control | 13.69242185 | 0.328979998 | 17.81072683 | 0.314021793 | 0.057579338 | -4.118304982 | -3.422252863 | 1.90E-09 | 8.72076197 | Unpaired t-test | 0 | permutation FDR (250 perm) | 1 | 2.05E-08 | FDR correction BH |
| woc~A0A0B4KI43 | VMAT~Control | VMAT | Control | 13.26920466 | 0.174443903 | 17.39984794 | 0.107914874 | 0.057089005 | -4.13064328 | -3.783559504 | 7.02E-13 | 12.15353246 | Unpaired t-test | 0 | permutation FDR (250 perm) | 1 | 7.51E-11 | FDR correction BH |
| ens~M9PBN6 | VMAT~Control | VMAT | Control | 13.22352954 | 0.476047796 | 17.3606913 | 0.237131193 | 0.056831643 | -4.13716176 | -3.342227558 | 8.36E-09 | 8.077710197 | Unpaired t-test | 0 | permutation FDR (250 perm) | 1 | 6.54E-08 | FDR correction BH |
| CtBP~A0A0B4KFX5 | VMAT~Control | VMAT | Control | 13.466637 | 0.255354122 | 17.60588685 | 0.254593283 | 0.056749447 | -4.139249855 | -3.564448009 | 1.85E-10 | 9.73284449 | Unpaired t-test | 0 | permutation FDR (250 perm) | 1 | 3.48E-09 | FDR correction BH |
| Svil~X2JAM4 | VMAT~Control | VMAT | Control | 15.02102357 | 0.18449748 | 19.17483977 | 0.24773337 | 0.056179352 | -4.153816202 | -3.649658123 | 3.87E-11 | 10.41212137 | Unpaired t-test | 0 | permutation FDR (250 perm) | 1 | 1.04E-09 | FDR correction BH |
| brm~M9PFM5 | VMAT~Control | VMAT | Control | 13.43773395 | 0.448532588 | 17.63934236 | 0.228587665 | 0.054348785 | -4.201608406 | -3.429500298 | 4.22E-09 | 8.374938603 | Unpaired t-test | 0 | permutation FDR (250 perm) | 1 | 3.80E-08 | FDR correction BH |
| zormin~M9PE03 | VMAT~Control | VMAT | Control | 12.38496366 | 1.447520811 | 16.58730074 | 0.761928666 | 0.054321342 | -4.202337079 | -2.42691742 | 0.000186591 | 3.729109037 | Unpaired t-test | 1.30E-05 | permutation FDR (250 perm) | 1 | 0.000345583 | FDR correction BH |
| bt~D1YSG0 | VMAT~Control | VMAT | Control | 15.24747152 | 0.393536128 | 19.46222478 | 0.20696293 | 0.053855845 | -4.214753257 | -3.515666912 | 1.22E-09 | 8.91521636 | Unpaired t-test | 0 | permutation FDR (250 perm) | 1 | 1.40E-08 | FDR correction BH |
| nocte~Q9W2U7 | VMAT~Control | VMAT | Control | 14.45335111 | 0.732931718 | 18.70330998 | 0.304569262 | 0.052557524 | -4.249958866 | -3.136614255 | 2.98E-07 | 6.525333553 | Unpaired t-test | 0 | permutation FDR (250 perm) | 1 | 1.19E-06 | FDR correction BH |
| Chro~Q86BS3 | VMAT~Control | VMAT | Control | 13.35962134 | 0.416240528 | 17.62269165 | 0.532205926 | 0.052082037 | -4.263070313 | -3.273848567 | 6.29E-08 | 7.201369531 | Unpaired t-test | 0 | permutation FDR (250 perm) | 1 | 3.23E-07 | FDR correction BH |
| htk~Q9VWP5 | VMAT~Control | VMAT | Control | 13.53555246 | 0.4390389 | 17.81280318 | 0.134921503 | 0.051572625 | -4.277250719 | -3.548388893 | 1.45E-09 | 8.839796861 | Unpaired t-test | 0 | permutation FDR (250 perm) | 1 | 1.64E-08 | FDR correction BH |
| Dmel\CG1674~Q8IMB9 | VMAT~Control | VMAT | Control | 13.50883893 | 0.427892515 | 17.78640216 | 0.107133071 | 0.051561455 | -4.277563224 | -3.572775805 | 9.72E-10 | 9.012234259 | Unpaired t-test | 0 | permutation FDR (250 perm) | 1 | 1.19E-08 | FDR correction BH |
| SmE~Q9VLV5 | VMAT~Control | VMAT | Control | 13.2827273 | 0.704173086 | 17.57003956 | 0.09296789 | 0.051214202 | -4.287312269 | -3.253760282 | 9.60E-08 | 7.017690339 | Unpaired t-test | 0 | permutation FDR (250 perm) | 1 | 4.63E-07 | FDR correction BH |
| RpS10b~M9NEQ9 | VMAT~Control | VMAT | Control | 13.36345921 | 0.365252569 | 17.72956992 | 0.180440945 | 0.048491957 | -4.366110711 | -3.693235317 | 3.64E-10 | 9.438935659 | Unpaired t-test | 0 | permutation FDR (250 perm) | 1 | 5.71E-09 | FDR correction BH |
| Hcf~Q9V4C8 | VMAT~Control | VMAT | Control | 15.22152322 | 0.729667203 | 19.58939526 | 0.452795342 | 0.048432791 | -4.367872039 | -3.155883178 | 4.83E-07 | 6.316109328 | Unpaired t-test | 0 | permutation FDR (250 perm) | 1 | 1.80E-06 | FDR correction BH |
| jv~M9NE38-2 | VMAT~Control | VMAT | Control | 13.28924764 | 0.263904145 | 17.67129273 | 0.245517647 | 0.047959316 | -4.382045098 | -3.773727055 | 1.05E-10 | 9.978823202 | Unpaired t-test | 0 | permutation FDR (250 perm) | 1 | 2.17E-09 | FDR correction BH |
| Sin3A~A0A0B4LF82 | VMAT~Control | VMAT | Control | 13.38923678 | 0.260341735 | 17.78724023 | 0.281722924 | 0.047431738 | -4.398003445 | -3.754006642 | 1.87E-10 | 9.727628253 | Unpaired t-test | 0 | permutation FDR (250 perm) | 1 | 3.50E-09 | FDR correction BH |
| CG6043~A8DZ00 | VMAT~Control | VMAT | Control | 13.18268067 | 0.383990411 | 17.58128997 | 0.360770449 | 0.047411824 | -4.398609296 | -3.559815343 | 4.21E-09 | 8.376100708 | Unpaired t-test | 0 | permutation FDR (250 perm) | 1 | 3.80E-08 | FDR correction BH |
| Prp39~Q7KRW8 | VMAT~Control | VMAT | Control | 13.21262119 | 0.619128091 | 17.62629293 | 0.203632267 | 0.046919396 | -4.413671736 | -3.417545739 | 3.19E-08 | 7.495891571 | Unpaired t-test | 0 | permutation FDR (250 perm) | 1 | 1.89E-07 | FDR correction BH |
| DmRH27~Q9VTC1 | VMAT~Control | VMAT | Control | 13.34017081 | 0.506180252 | 17.75446255 | 0.097629597 | 0.046899237 | -4.414291744 | -3.587271919 | 3.28E-09 | 8.483568255 | Unpaired t-test | 0 | permutation FDR (250 perm) | 1 | 3.13E-08 | FDR correction BH |
| scrib~A0A0B4K6I1 | VMAT~Control | VMAT | Control | 13.911623 | 0.402733469 | 18.37819271 | 0.101495315 | 0.045230203 | -4.466569709 | -3.766906848 | 3.52E-10 | 9.453731584 | Unpaired t-test | 0 | permutation FDR (250 perm) | 1 | 5.63E-09 | FDR correction BH |
| Wnk~M9PIH3 | VMAT~Control | VMAT | Control | 14.91189879 | 2.217228544 | 19.38220045 | 0.32728437 | 0.045113353 | -4.47030166 | -2.232562109 | 0.001216209 | 2.914991904 | Unpaired t-test | 2.76E-05 | permutation FDR (250 perm) | 1 | 0.001918871 | FDR correction BH |
| Bacc~Q9VQF7 | VMAT~Control | VMAT | Control | 13.48891835 | 0.149194559 | 17.97224249 | 0.243595263 | 0.04470797 | -4.483324136 | -3.975466516 | 8.37E-12 | 11.07745878 | Unpaired t-test | 0 | permutation FDR (250 perm) | 1 | 3.40E-10 | FDR correction BH |
| Dmel\CG8149~Q9VHC8 | VMAT~Control | VMAT | Control | 13.26251982 | 0.252907272 | 17.76928332 | 0.321259424 | 0.043987472 | -4.506763498 | -3.810090364 | 2.76E-10 | 9.559111662 | Unpaired t-test | 0 | permutation FDR (250 perm) | 1 | 4.64E-09 | FDR correction BH |
| jbug~Q9W205 | VMAT~Control | VMAT | Control | 13.16061633 | 0.401522941 | 17.69228051 | 0.181977275 | 0.04323477 | -4.531664178 | -3.785383841 | 5.48E-10 | 9.260881322 | Unpaired t-test | 0 | permutation FDR (250 perm) | 1 | 7.78E-09 | FDR correction BH |
| His4r~A0A0B4KFZ9 | VMAT~Control | VMAT | Control | 14.32978051 | 0.245201351 | 18.88742024 | 0.402387906 | 0.042463299 | -4.557639736 | -3.764367344 | 9.97E-10 | 9.001107327 | Unpaired t-test | 0 | permutation FDR (250 perm) | 1 | 1.22E-08 | FDR correction BH |
| Dmel\CG9667~A0A0B4LGT9 | VMAT~Control | VMAT | Control | 12.81133835 | 0.515904491 | 17.39922521 | 0.855000724 | 0.041582293 | -4.587886859 | -3.171532901 | 1.24E-06 | 5.90629268 | Unpaired t-test | 0 | permutation FDR (250 perm) | 1 | 4.06E-06 | FDR correction BH |
| Dmel\CG44195~M9PEG1 | VMAT~Control | VMAT | Control | 12.93451387 | 0.382546811 | 17.58682804 | 0.177650244 | 0.039766181 | -4.65231417 | -3.914021983 | 2.74E-10 | 9.561998594 | Unpaired t-test | 0 | permutation FDR (250 perm) | 1 | 4.64E-09 | FDR correction BH |
| qkr58E-1~Q9W255 | VMAT~Control | VMAT | Control | 13.22470665 | 0.325772158 | 17.92186926 | 0.161030831 | 0.038549004 | -4.697162618 | -4.040511867 | 5.73E-11 | 10.24196314 | Unpaired t-test | 0 | permutation FDR (250 perm) | 1 | 1.41E-09 | FDR correction BH |
| Top2~P15348 | VMAT~Control | VMAT | Control | 13.16428977 | 0.609091096 | 17.86545773 | 0.249061947 | 0.038442129 | -4.701167958 | -3.632245724 | 1.91E-08 | 7.719776576 | Unpaired t-test | 0 | permutation FDR (250 perm) | 1 | 1.25E-07 | FDR correction BH |
| HIPP1~M9PID3 | VMAT~Control | VMAT | Control | 13.37012778 | 0.150162095 | 18.18643296 | 0.356720749 | 0.035493406 | -4.816305178 | -4.105661845 | 8.34E-11 | 10.07890004 | Unpaired t-test | 0 | permutation FDR (250 perm) | 1 | 1.85E-09 | FDR correction BH |
| sm~A0A0B4K7I2 | VMAT~Control | VMAT | Control | 12.47098815 | 0.051218082 | 17.33633542 | 0.233265269 | 0.034307141 | -4.865347273 | -4.395850749 | 6.26E-13 | 12.20371285 | Unpaired t-test | 0 | permutation FDR (250 perm) | 1 | 7.36E-11 | FDR correction BH |
| Pbgd~Q9W0G4 | VMAT~Control | VMAT | Control | 12.66075447 | 0.590552274 | 17.53567999 | 0.307084287 | 0.034080126 | -4.874925526 | -3.756660657 | 1.50E-08 | 7.824036885 | Unpaired t-test | 0 | permutation FDR (250 perm) | 1 | 1.03E-07 | FDR correction BH |
| Decay~Q9VET9 | VMAT~Control | VMAT | Control | 12.27997997 | 1.208888981 | 17.21181667 | 0.740033396 | 0.03276191 | -4.931836705 | -3.018469141 | 1.50E-05 | 4.823485296 | Unpaired t-test | 0 | permutation FDR (250 perm) | 1 | 3.60E-05 | FDR correction BH |
| Larp4B~Q9I7T7-2 | VMAT~Control | VMAT | Control | 12.70175995 | 0.429525695 | 17.67132107 | 0.523689161 | 0.031916337 | -4.969561118 | -3.814230063 | 1.47E-08 | 7.831707897 | Unpaired t-test | 0 | permutation FDR (250 perm) | 1 | 1.02E-07 | FDR correction BH |
| Smr~Q9VYK0 | VMAT~Control | VMAT | Control | 15.69126064 | 3.324123516 | 20.6677845 | 0.24326062 | 0.031762673 | -4.976523862 | -1.998147724 | 0.007507254 | 2.124518893 | Unpaired t-test | 0.000113813 | permutation FDR (250 perm) | 1 | 0.010626624 | FDR correction BH |
| dHspB8~Q9VWG1 | VMAT~Control | VMAT | Control | 15.33450723 | 2.016287137 | 20.52880333 | 0.440251861 | 0.027312476 | -5.194296099 | -2.701204337 | 0.00021905 | 3.659456484 | Unpaired t-test | 0 | permutation FDR (250 perm) | 1 | 0.000400656 | FDR correction BH |
| PP2Cg~E2QC56 | VMAT~Control | VMAT | Control | 13.14878866 | 0.454688611 | 18.47979866 | 0.110908696 | 0.024843117 | -5.331009994 | -4.408326211 | 2.00E-10 | 9.699654952 | Unpaired t-test | 0 | permutation FDR (250 perm) | 1 | 3.64E-09 | FDR correction BH |
| anon-WO0118547.398~Q9V9U3 | VMAT~Control | VMAT | Control | 13.3822832 | 0.360514354 | 18.72191009 | 0.16455526 | 0.024695177 | -5.33962689 | -4.535762269 | 3.80E-11 | 10.42049225 | Unpaired t-test | 0 | permutation FDR (250 perm) | 1 | 1.03E-09 | FDR correction BH |
| Zasp67~M9PEZ7 | VMAT~Control | VMAT | Control | 12.78822636 | 0.764942497 | 18.14717652 | 0.602962513 | 0.024366618 | -5.358950158 | -3.732920728 | 2.31E-07 | 6.636299132 | Unpaired t-test | 0 | permutation FDR (250 perm) | 1 | 9.71E-07 | FDR correction BH |
| Acbp3~Q9VS22 | VMAT~Control | VMAT | Control | 13.43652639 | 0.450366689 | 18.84956509 | 0.298268218 | 0.023469994 | -5.413038694 | -4.359813672 | 7.05E-10 | 9.152038459 | Unpaired t-test | 0 | permutation FDR (250 perm) | 1 | 9.37E-09 | FDR correction BH |
| Msp300~M9MRJ4 | VMAT~Control | VMAT | Control | 13.21299698 | 0.603447195 | 18.62849865 | 0.167172849 | 0.02342996 | -5.41550168 | -4.230748221 | 2.98E-09 | 8.525711757 | Unpaired t-test | 0 | permutation FDR (250 perm) | 1 | 2.93E-08 | FDR correction BH |
| Ge-1~Q9VKK1 | VMAT~Control | VMAT | Control | 13.01232074 | 0.289166046 | 18.53785107 | 0.134888766 | 0.02170949 | -5.525530331 | -4.835517174 | 3.15E-12 | 11.50172649 | Unpaired t-test | 0 | permutation FDR (250 perm) | 1 | 1.77E-10 | FDR correction BH |
| Sf3a1~Q9VEP9 | VMAT~Control | VMAT | Control | 13.09548495 | 0.571323397 | 18.63141585 | 0.795404932 | 0.021553547 | -5.535930906 | -3.849829253 | 1.79E-07 | 6.747269646 | Unpaired t-test | 0 | permutation FDR (250 perm) | 1 | 7.92E-07 | FDR correction BH |
| Lmpt~Q7KUQ6 | VMAT~Control | VMAT | Control | 13.69904829 | 0.462162174 | 19.29864611 | 0.133248081 | 0.020623059 | -5.599597824 | -4.608327552 | 1.61E-10 | 9.793141382 | Unpaired t-test | 0 | permutation FDR (250 perm) | 1 | 3.14E-09 | FDR correction BH |
| uncNacbeta~Q9VPR2 | VMAT~Control | VMAT | Control | 13.41853497 | 1.155502888 | 19.31559372 | 0.422024485 | 0.016780636 | -5.897058751 | -3.80420084 | 8.38E-07 | 6.076686039 | Unpaired t-test | 0 | permutation FDR (250 perm) | 1 | 2.89E-06 | FDR correction BH |
| MTA1-like~Q9VNF7 | VMAT~Control | VMAT | Control | 13.22637838 | 0.225812155 | 19.1463219 | 0.061836289 | 0.016516554 | -5.919943516 | -5.35884943 | 7.27E-14 | 13.1386508 | Unpaired t-test | 0 | permutation FDR (250 perm) | 1 | 1.56E-11 | FDR correction BH |
| Cp190~Q24478 | VMAT~Control | VMAT | Control | 12.53571044 | 1.139226628 | 18.54074336 | 0.602549713 | 0.015570586 | -6.005032917 | -3.809451192 | 1.09E-06 | 5.962676353 | Unpaired t-test | 0 | permutation FDR (250 perm) | 1 | 3.64E-06 | FDR correction BH |
| None~2xHA-TurboID_bait | VMAT~Control | VMAT | Control | 13.21628704 | 0.463238648 | 19.39166523 | 0.106744988 | 0.01383649 | -6.175378191 | -5.092693698 | 5.45E-11 | 10.26344541 | Unpaired t-test | 0 | permutation FDR (250 perm) | 1 | 1.38E-09 | FDR correction BH |
| Dsp1~F2FBA6 | VMAT~Control | VMAT | Control | 13.06717079 | 0.224450424 | 19.24679758 | 0.290323271 | 0.013795803 | -6.179626786 | -5.308441796 | 4.16E-12 | 11.38107868 | Unpaired t-test | 0 | permutation FDR (250 perm) | 1 | 2.22E-10 | FDR correction BH |
| CG8154~M9PF57 | VMAT~Control | VMAT | Control | 12.33051093 | 0.493202886 | 18.53446006 | 0.334100056 | 0.013565169 | -6.203949137 | -4.898846527 | 4.83E-10 | 9.316448582 | Unpaired t-test | 0 | permutation FDR (250 perm) | 1 | 6.97E-09 | FDR correction BH |
| Dmel\CG18304~M9PCS5 | VMAT~Control | VMAT | Control | 11.09820592 | 0.705119071 | 17.56882867 | 0.666737891 | 0.011275828 | -6.470622751 | -4.512323798 | 3.71E-08 | 7.43118323 | Unpaired t-test | 0 | permutation FDR (250 perm) | 1 | 2.13E-07 | FDR correction BH |
| Rcc1~P25171 | VMAT~Control | VMAT | Control | 10.31378459 | 0.332306329 | 16.83631194 | 0.397678026 | 0.010877363 | -6.522527351 | -5.295268837 | 7.45E-11 | 10.12755481 | Unpaired t-test | 0 | permutation FDR (250 perm) | 1 | 1.69E-09 | FDR correction BH |
| hnRNP U~A1ZBB4 | VMAT~Control | VMAT | Control | 14.17583231 | 1.894570174 | 21.01270239 | 0.214824303 | 0.008747763 | -6.836870073 | -3.690205897 | 1.15E-05 | 4.937518267 | Unpaired t-test | 0 | permutation FDR (250 perm) | 1 | 2.85E-05 | FDR correction BH |
| RpA-70~Q24492 | VMAT~Control | VMAT | Control | 10.1190796 | 0.748338932 | 17.42976568 | 0.257323756 | 0.006298893 | -7.310686086 | -5.399723124 | 1.56E-09 | 8.805786856 | Unpaired t-test | 0 | permutation FDR (250 perm) | 1 | 1.75E-08 | FDR correction BH |
| sls~Q9I7U4-3 | VMAT~Control | VMAT | Control | 12.91330684 | 0.850416406 | 20.35590344 | 0.215428144 | 0.005748508 | -7.442596606 | -5.345422594 | 3.60E-09 | 8.444186663 | Unpaired t-test | 0 | permutation FDR (250 perm) | 1 | 3.37E-08 | FDR correction BH |
| Dmel\CG1674~H9XVM8 | VMAT~Control | VMAT | Control | 15.98616737 | 0.108039849 | 23.42939337 | 0.088841835 | 0.005746001 | -7.443226004 | -7.005028247 | 4.31E-17 | 16.36531433 | Unpaired t-test | 0 | permutation FDR (250 perm) | 1 | 1.02E-13 | FDR correction BH |
| Nup153~A8JV18 | VMAT~Control | VMAT | Control | 10.8349301 | 0.302147501 | 18.34108373 | 0.198407567 | 0.005500759 | -7.506153634 | -6.461613565 | 5.17E-13 | 12.28644366 | Unpaired t-test | 0 | permutation FDR (250 perm) | 1 | 6.76E-11 | FDR correction BH |
| Brr2~Q9VUV9 | VMAT~Control | VMAT | Control | 9.314068307 | 0.508499933 | 16.83126323 | 0.820737216 | 0.005458821 | -7.517194921 | -5.250234275 | 8.29E-09 | 8.081429663 | Unpaired t-test | 0 | permutation FDR (250 perm) | 1 | 6.51E-08 | FDR correction BH |
| Dmel\CG44245~Q9W2J5 | VMAT~Control | VMAT | Control | 10.79789211 | 1.126613671 | 18.82273472 | 0.109922731 | 0.003839562 | -8.02484261 | -5.327768963 | 2.05E-08 | 7.687484465 | Unpaired t-test | 0 | permutation FDR (250 perm) | 1 | 1.32E-07 | FDR correction BH |
| Sf3b1~Q9VPR5 | VMAT~Control | VMAT | Control | 11.8233422 | 0.699166227 | 19.86389452 | 0.096932461 | 0.003797979 | -8.040552326 | -6.111387085 | 2.00E-10 | 9.699910477 | Unpaired t-test | 0 | permutation FDR (250 perm) | 1 | 3.64E-09 | FDR correction BH |
| Mtor~A0A0B4KFE9 | VMAT~Control | VMAT | Control | 9.281873381 | 0.45605647 | 18.43391075 | 0.199128731 | 0.001757771 | -9.152037365 | -7.486030542 | 1.73E-12 | 11.76156131 | Unpaired t-test | 0 | permutation FDR (250 perm) | 1 | 1.31E-10 | FDR correction BH |
| Pep~M9NG39 | VMAT~Control | VMAT | Control | 9.621201399 | 0.217948019 | 18.95249798 | 0.183598471 | 0.001552386 | -9.331296585 | -8.276506342 | 5.51E-15 | 14.25883837 | Unpaired t-test | 0 | permutation FDR (250 perm) | 1 | 2.35E-12 | FDR correction BH |
| Dmel\CG1674~Q9V4C1 | VMAT~Control | VMAT | Control | 12.37928805 | 0.205808954 | 22.92553879 | 0.377962962 | 0.000668748 | -10.54625074 | -8.844077846 | 9.93E-14 | 13.00298769 | Unpaired t-test | 0 | permutation FDR (250 perm) | 1 | 1.95E-11 | FDR correction BH |
